# Architecture of the linker-scaffold in the nuclear pore

**DOI:** 10.1101/2021.10.26.465796

**Authors:** Stefan Petrovic, Dipanjan Samanta, Thibaud Perriches, Christopher J. Bley, Karsten Thierbach, Bonnie Brown, Si Nie, George W. Mobbs, Taylor A. Stevens, Xiaoyu Liu, André Hoelz

**Author notes:** Correspondence (A.H.). these authors contributed equally to this work.

## Abstract

The nuclear pore complex (NPC) is the sole bidirectional gateway for nucleocytoplasmic transport. Despite recent progress in elucidating the arrangement of the structured scaffold building blocks in the NPC symmetric core, their cohesion by multivalent unstructured linker proteins remained elusive. Combining biochemical reconstitution, high resolution structure determination, docking into cryo-electron tomographic reconstructions, and physiological validation, we elucidated the architecture of the entire linker-scaffold, yielding a near-atomic composite structure of the symmetric core accounting for ∼77 MDa of the human NPC. Whereas linkers generally play a rigidifying role, the linker-scaffold of the NPC provides the plasticity and robustness necessary for the reversible constriction and dilation of its central transport channel. Our results complete the structural characterization of the NPC symmetric core, providing a rich foundation for future functional studies.

**One sentence summary:** An interdisciplinary analysis established the near-atomic molecular architecture and evolutionary conservation of the linker-scaffold of the human nuclear pore complex.

## Introduction

The enclosure of genetic material in the nucleus requires the selective transport of folded proteins and ribonucleic acids across the nuclear envelope, for which the nuclear pore complex (NPC) is the sole gateway (*1–4*). Beyond its function as a selective, bidirectional channel for macromolecules, the role of the NPC extends to genome organization, transcription regulation, mRNA maturation, and ribosome assembly (*1, 2*). The NPC and its components are implicated in the etiology of many human diseases, including viral infections (*5, 6*). The building blocks of the NPC are a set of ∼34 different proteins collectively termed nucleoporins (nups). In the NPC, nups assemble into defined subcomplexes that are generally present in multiples of eight, adding up to a mass of ∼110 MDa in the human NPC (*1–4*). The NPC architecture consists of a symmetric core with asymmetric decorations on its nuclear and cytoplasmic faces (Fig. 1A). The symmetric core displays eight-fold rotational symmetry along the nucleocytoplasmic axis and two-fold symmetry along an axis coplanar with the nuclear envelope. It consists of two outer rings that sit on top of the nuclear envelope and an inner ring that lines the lumen generated by the fusion of the two lipid bilayers of the nuclear envelope. From the inner ring, unstructured phenylalanine-glycine (FG) repeats are projected into the central transport channel to establish a diffusion barrier that bars particles larger than ∼40 kDa from freely diffusing across the nuclear envelope. Transport is mediated by karyopherins, whose high affinity for FG repeats and ultrafast exchange kinetics allow karyopherin-bound cargo to traverse the diffusion barrier (*7–10*).

**Fig. 1.**
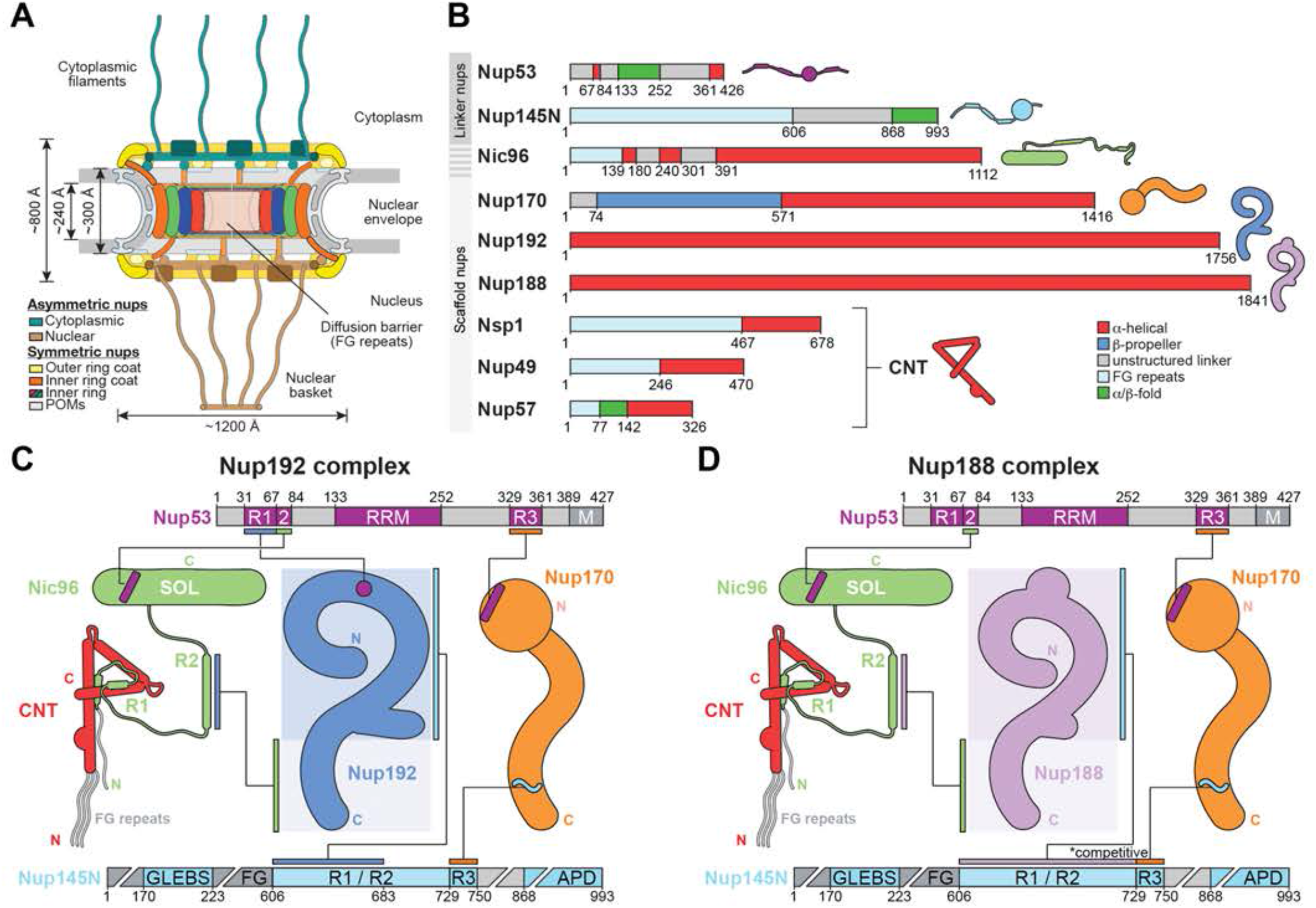
Outline of the symmetric core inner ring architecture. (**A**) Schematic conceptualization of the nuclear pore complex (NPC), shown as a cross-section. Outer rings coat the nuclear envelope on both cytoplasmic and nuclear faces, which are further decorated by asymmetric nups. Pore membrane proteins (POMs) are transmembrane proteins associated with the NPC. The inner ring is composed of concentric layers of nups. Phenylalanine-glycine (FG)-repeats are projected into the central transport channel to form the diffusion barrier. (**B**) Domain structures of the *Chaetomium thermophilum* inner ring linker and scaffold nucleoporins (nups) represented as horizontal boxes, with observed or predicted domain folds color-coded according to the legend and domain boundary residue numbers indicated. Nic96 consists of a linker (residues 1-390) and a scaffold (residues 391-1112) region. Schematic stylizations of the nup structures are used throughout this work as a visual aid. The channel nup hetero-trimer (CNT) components Nsp1, Nup49, and Nup57 form a stable complex and are drawn as a unit. (**C**, **D**) Schematic map of previously established linker-scaffold interactions in alternative, mutually exclusive inner ring complexes organized around the Nup192 and Nup188 scaffold hubs (*30*). Colored bars connected by black lines indicate interactions between nup regions.

The structural characterization of the NPC has progressed through efforts to reconstitute and crystallize ever larger portions of it, from small nup domain fragments to complexes as large as the ∼400 kDa hetero-heptameric Y-shaped coat nup complex (CNC) (*11–27*). In parallel, progress has been driven by efforts to push the resolution of cryo-electron tomographic (cryo-ET) reconstructions of intact NPCs(*5, 6*). The docking of the CNC crystal structure into an ∼32 Å cryo-ET map of the intact human NPC was the first demonstration that biochemical reconstitution and crystal structures could be used to interpret cryo-ET maps and unraveled the head-to-tail tandem arrangement of CNCs in the outer rings. The reconstitution and piecemeal structural analysis of two hetero-nonameric ∼425 kDa inner ring complexes (IRCs) provided the basis for docking 17 symmetric core nups into an ∼23 Å cryo-ET map of the intact human NPC (*25, 28*), yielding a near-atomic composite structure of the entire ∼56 MDa symmetric core of the human NPC (*29, 30*). A similar approach elucidated the near-atomic architectures of constricted and dilated states of the *S. cerevisiae* NPC using nup and nup complex crystal structures to interpret ∼25 Å cryo-ET maps (*31, 32*). Apart from an additional distal CNC ring and associated nups present in the outer rings of the human NPC, the human and *S. cerevisiae* NPC present equivalent nup arrangements (*29–33*).

The inner ring of the human NPC is composed of six scaffold nups NUP155, NUP188, NUP205, NUP54, NUP58, and NUP62, two primarily unstructured linker nups NUP53 and NUP98, and NUP93, which is a hybrid of both (*29, 30*). The doughnut-shaped inner ring adopts a concentric cylinder architecture, in which membrane-anchored NUP155 forms the outermost coat, followed by layers of NUP93, NUP205/NUP188, and the NUP54•NUP58•NUP62 channel nucleoporin hetero-trimer (CNT) in the center, providing the FG repeats to form the diffusion barrier in the central transport channel. Unlike the extensive interactions of large, folded domains found in the CNC (*13-17, 19, 22, 24, 26, 34*), the structured domains of the inner ring nups do not interact directly. Instead, the inner ring is held together by the linker nups NUP53, NUP98, and an unstructured region of NUP93, which are proposed to connect the scaffolds of the four layers (*25, 28, 30, 35, 36*). The resulting linker-scaffold architecture allows for a substantial ∼200 Å dilation of the inner ring’s central transport channel, accompanied by the generation of lateral gaps between the eight spokes, as observed in recent cryo-ET analyses of purified and *in situ* human and yeast NPCs (*31, 37–40*). The linker-scaffold is expected to play a fundamental role in establishing an architectural framework to accommodate the structural changes associated with the reversible constriction and dilation of the inner ring.

Whereas our previous work achieved the identification of the majority of the scaffold nup locations in the NPC, a comprehensive understanding and the molecular details of the linker-scaffold interaction network that mediates the cohesion of the symmetric core has remained elusive. Here, we report the characterization of all linker-scaffold interactions through residue-level biochemical mapping of scaffold-binding regions in linker nups and the determination of crystal and single particle cryo-EM structures of linker-scaffold complexes. Our analysis revealed a common linker-scaffold binding mode, whereby linkers are anchored by central structured motifs whose binding is reinforced by disperse interactions of flanking regions. We quantitatively docked the complete set of linker-scaffold structures into an ∼12Å cryo-ET reconstruction of the human NPC (provided by Martin Beck) (*41*), and an ∼25Å *in situ* cryo-ET reconstruction of the *S. cerevisiae* NPC (*31*). In the inner ring, our new linker-scaffold structures allowed for the unambiguous assignment of NUP188 and NUP205 to 16 peripheral and 16 equatorial positions, respectively. From the nuclear envelope to the central transport channel, linkers bridge the layers of the inner ring to coalesce scaffold nups into eight relatively rigid spokes that are flexibly inter-connected, allowing for the formation of lateral gates. The linker-scaffold confers the plasticity necessary for the reversible dilation and constriction of the inner ring’s central transport channel in response to alterations in the nuclear envelope membrane tension. The topology of linker-scaffold interactions between inner ring nups is conserved from fungi to humans. We carried out systematic functional analyses of the linker-scaffold network, including the development of a minimal linker S. *cerevisiae* strain, establishing its robustness and essential nature. Our quantitative docking analysis of the human NPC revealed eight NUP205•NUP93 complexes that cross-link adjacent spokes in both nuclear and cytoplasmic outer rings. Facing the central transport channel, an additional eight NUP205•NUP93 copies are exclusively anchored at the base of the cytoplasmic outer ring. NUP93 emerges as a versatile linker-scaffold hybrid that recruits and positions the FG repeat-harboring CNT to the inner ring and reinforces the tandem head-to-tail CNC arrangement in the outer rings, explaining its fundamental role in maintaining the integrity of the entire NPC. Our analysis completes the structural characterization of the ∼77 MDa symmetric core and lays out a roadmap for future studies on the NPC assembly and function.

## RESULTS

### Biochemical and structural analysis of *C. thermophilum* scaffold-linker interactions

Nup192 and Nup188 are the scaffold keystones of two alternative eight-protein inner ring complexes, which both include the linkers Nup145N, Nup53, and the scaffold-linker hybrid Nic96 (Fig. 1, B to D) (*26, 28, 30, 35, 36*). Previously, a composite structure of the full-length ∼200 kDa Nup192 was determined by superposing overlapping structures of its N- and C-terminal parts, revealing an extended *α*-helical solenoid with a question mark-shaped architecture, composed of five Huntingtin, elongation factor 3 (EF3), protein phosphatase 2A (PP2A), and the yeast kinase TOR1 (HEAT), fifteen Armadillo (ARM) repeats, and a prominent central Tower (*30, 35, 36, 42*). A question mark shape was also observed for Nup188 by negative-stain electron microscopy (EM) (*35*) and high-resolution structures of its ∼130 kDa N-terminal domain (NTD), which contains a central SH3-like domain insertion, and ∼45 kDa C-terminal Tail region again revealed extended *α*-helical solenoids composed of ARM and HEAT repeats (*25, 43*). However, no structural information could so far be obtained for the ∼32 kDa Nup188 central region equivalent to the Nup192 Tower. Furthermore, although binding of Nup192 and Nup188 has biochemically been mapped to extensive unstructured regions in the linkers Nic96, Nup53 and Nup145N, the molecular details of these interactions have yet to be elucidated.

### Nup192 interaction with Nic96

Having previously determined the crystal structure of a *C. thermophilum* Nup192 fragment only missing the N-terminal Head subdomain (Nup192^ΔHead^, residues 153-1756) (*30*), we were able to obtain co-crystals with our previously mapped Nic96^187-301^ fragment that diffracted to 3.6 Å resolution, allowing for structure determination by molecular replacement with the Nup192^ΔHead^ structure as search model (Fig. 2). The Nic96 sequence register was unambiguously assigned by calculating anomalous difference Fourier maps to locate seleno-L-methionine (SeMet)-labeled endogenous M263 and substituted A289M residues (fig. S1). In the Nup192^ΔHead^•Nic96^187-301^ complex structure, Nup192 adopts the previously described question mark shaped *α*-helical solenoid (*30*), on which binding of Nic96 (residues 240-301; R2) extends from the midpoint to the base, burying ∼3,700 Å^2^ of combined surface area. Nic96^R2^ forms two amphipathic *α*-helices connected via a sharply kinking connector. The longer N-terminal *α*-helix is cradled by an extensive concave Nup192 surface formed by ten ARM and HEAT repeats and the central Tower, while the shorter C-terminal one packs against a hydrophobic patch formed from the C-terminal Nup192 *α*-helices *α*75-77 (ARM20). Nic96 residues 187-239 were not resolved in the complex structure, but were found to be dispensable for Nup192 binding by Isothermal titration calorimetry (ITC), as both Nic96^187-301^ and Nic96^R2^ retained a dissociation constant (*KD*) of ∼75 nM (Fig. 2E, fig. S2, A and D). To obtain the structure of full-length Nup192 with Nic96^R2^ despite its resistance to crystallization, we determined a single particle cryo-EM reconstruction of the complex at 3.8 Å resolution from a refined set of 176,609 particles (Fig. 2B, fig. S3). The cryo-EM reconstruction confirmed the molecular details of Nic96^R2^ binding, but showed a slightly wider gap between the Nup192 Head subdomain and Tower (fig. S4). To validate the molecular details of the Nup192-Nic96^R2^ interface, we performed structure-guided mutagenesis and assessed binding by both size-exclusion chromatography coupled to multi-angle light scattering (SEC-MALS) and ITC. Consistent with the extensive hydrophobic nature of the interface, binding was not strongly affected by individual substitutions, but abolished by triple mutations in Nic96 (FFF mutant; F275E, F278E, and F298E) (Fig. 2, C, E and F, figs. S5, S2B, and S6) or Nup192 (LAF mutant; L1584E, A1648E, and F1735E) (Fig. 2, D, E and F figs. S2C, S6, and S7).

**Fig. 2.**
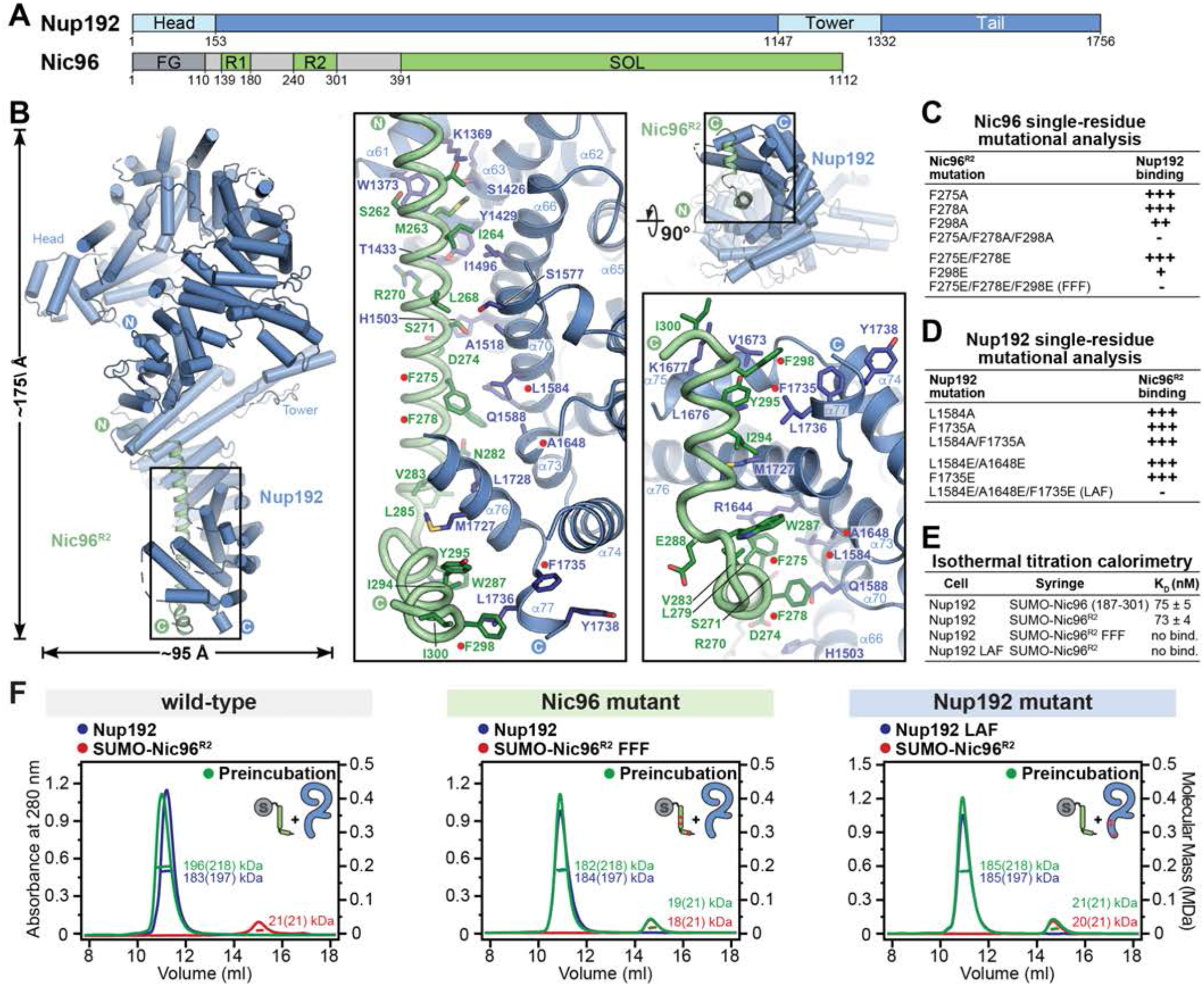
Structural and biochemical analyses of the Nup192-Nic96 interaction. (**A**) Domain structures of Nup192 and Nic96. (**B**) Two views of a cartoon representation of the Nup192•Nic96^R2^ single particle cryo-EM structure. The insets indicate regions magnified to illustrate the molecular details of the Nup192-Nic96 interaction. Red circles indicate residues found to be involved in the Nup192-Nic96 interaction. (**C**, **D**) Summary of the effect of structure-guided mutations in (C) SUMO-Nic96^R2^ and (D) Nup192 on Nup192•SUMO-Nic96^R2^ complex formation, assayed by size exclusion chromatography (SEC); (+++) no effect, (++) weak effect, (+) moderate effect, (-) abolished binding. (**E**) Dissociation constants (*KD*s) determined by isothermal titration calorimetry (ITC). All experiments were performed in triplicate, with the mean and its standard error reported. Experiments for which no heat exchange to indicate binding could be detected are indicated as “no bind.”. (**F**) Size-exclusion chromatography coupled to multi-angle light scattering (SEC-MALS) analysis of Nup192•SUMO-Nic96^R2^ and interaction-abolishing mutants. SEC-MALS profiles are shown for individual nups and for their preincubation. Measured molecular masses are indicated, with respective theoretical masses provided in parentheses.

### Nup188 interaction with Nic96

Having identified the Nic96^R2^ fragment as sufficient for Nup192 binding, we tested whether the same Nic96 region was also sufficient for Nup188 binding (Fig. 3). Indeed, ITC measurements revealed that Nup188 binds both Nic96^R2^ and Nic96^187-301^ with similar *KDs* of ∼90 nM (Fig. 3E and fig. S8, A and D). We were able to determine a crystal structure of Nup188•Nic96^R2^ at 4.4 Å resolution as well as a crystal structure of Nup188^NTD^ (residues 1-1134) at 2.8 Å resolution, which was used to aid with the phasing, model building, and sequence assignment. The Nup188 and Nic96^R2^ sequence registers were unambiguously assigned by identifying SeMet-labeled residues in anomalous difference Fourier maps (Fig. 3B, fig. S9). Similar to the Nup192•Nic96^R2^ complex, Nup188 adopts an overall question mark-shaped architecture, composed of an N-terminal Head subdomain, 9 HEAT, 13 ARM repeats, and a central, comparatively more compact Tower (Fig. 3B), with Nic96^R2^ binding an ∼3,700 Å^2^ concave surface between the midpoint and the base of the Nup188 molecule. However, although Nup188-bound Nic96^R2^ again forms two amphipathic *α*-helices, their secondary structure is switched compared to its Nup192-bound form (Fig. 3B): Whereas Nup192-bound Nic96^R2^ has a longer N-terminal helix and a shorter C-terminal helix which wraps around the Nup192^Tail^, Nup188-bound Nic96^R2^ has a shorter N-terminal helix that binds to the central Tower, and a longer C-terminal helix cradled in the concave surface at the base of Nup188 (Fig. 3B). Validation of the Nup188•Nic96^R2^ interface through structure-guided mutagenesis and binding assessment by SEC-MALS and ITC again failed to show an effect for single substitutions (Fig. 3, Cand D, figs. S8, S10, and S11). Remarkably, the Nic96^R2^ FFF mutation that abolishes binding to Nup192 had the same effect on Nup188 binding, despite the structural polymorphism between Nup192- and Nup188-bound Nic96^R2^ (Fig. 3, C to F, figs. S8, S10 and S12). Analogous to the Nup192 LAF mutant, we were also able to identify a triple Nup188 mutation (FLV mutant; F1478E, L1689E, and V1760E) that disrupted binding to Nic96^R2^ (Fig. 3, D to F, figs. S8, S11, and S12).

**Fig. 3.**
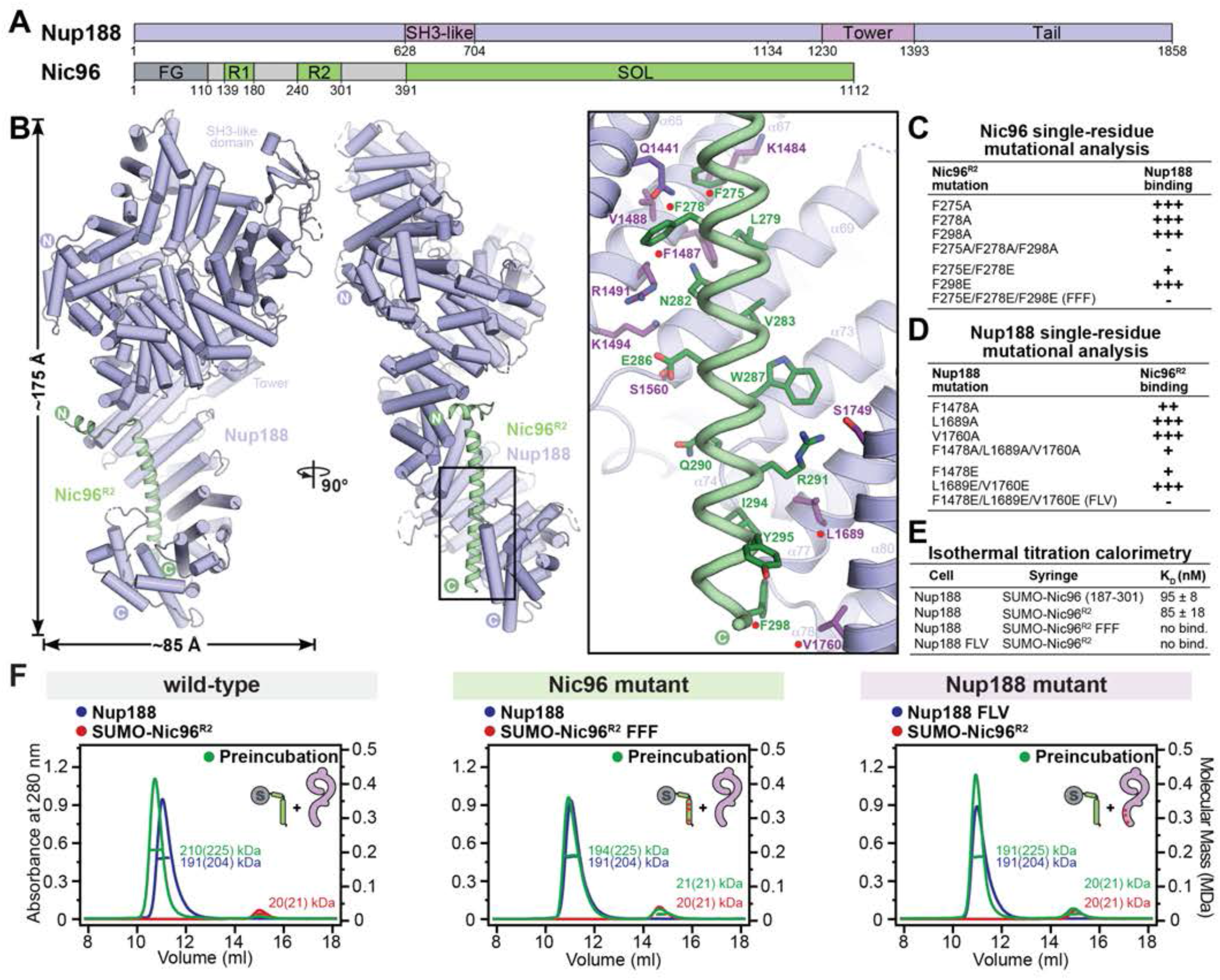
Structural and biochemical analyses of the Nup188-Nic96 interaction. (**A**) Domain structures of Nup188 and Nic96. (**B**) Two views of a cartoon representation of the Nup188•Nic96^R2^ crystal structure. The insets indicate regions magnified to illustrate the molecular details of the Nup188-Nic96 interaction. Red circles indicate residues found to be involved in Nup188-Nic96 binding. (**C**, **D**) Summary of the effect of structure-driven mutations in (C) SUMO-Nic96^R2^ and (D) Nup188 on Nup188•SUMO-Nic96^R2^ complex formation, assayed by SEC; (+++) no effect, (++) weak effect, (+) moderate effect, (-) abolished binding. (**E**) *KD*s determined by ITC. All experiments were performed in triplicate, with the mean and its standard error reported. Experiments for which no heat exchange to indicate binding could be detected are indicated as “no bind.”. (**F**) SEC-MALS analysis of Nup188•SUMO-Nic96^R2^ and interaction-abolishing mutants. SEC-MALS profiles are shown for individual nups and for their preincubation. Measured molecular masses are indicated, with respective theoretical masses provided in parentheses.

### Nup192 interaction with Nup145N and Nup53

To identify the Nup145N regions necessary and sufficient for Nup192 binding, we used a combination of five-alanine scanning mutagenesis and truncation analysis (Fig. 4A). Substituting blocks of five consecutive residues at a time to alanines, we found a hotspot between residues 626-655 that displayed diminished affinity for binding to Nup192 when mutated (Fig. 4A and fig. S13). N- and C-terminal Nup145N truncation resulted in a minimal Nup145N^R1^ peptide (residues 616-683) that recapitulated the Nup192-Nup145N interaction (Fig. 4B and fig. S14). Although shorter Nup145N fragments still bound to Nup192 as well, they had progressively reduced affinity and ITC measurements confirmed that the majority of the interaction is encapsulated in the Nup145N^R1^ region, with measured *KD*s of ∼825 nM and ∼1,500 nM for Nup145N and Nup145N^R1^, respectively (Fig. 4G and fig. S15).

**Fig. 4.**
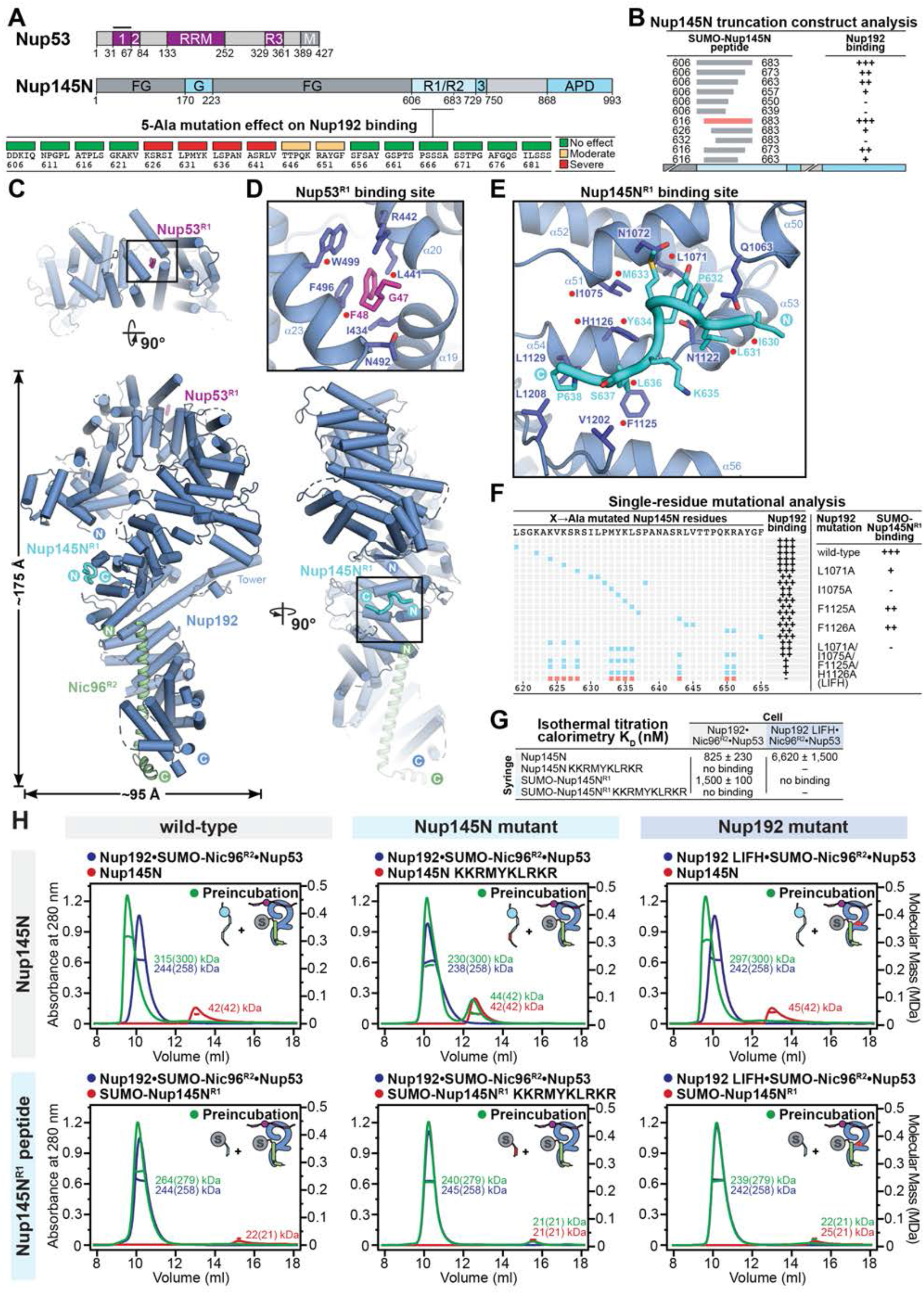
Structural and biochemical analyses of the Nup192-Nup145N interaction. (**A**) Domain structures of Nup53 and Nup145N, with the Nup145N primary sequence analyzed by 5-Ala scanning mutagenesis indicated. The effect of each 5-Ala substitution on Nup145N binding to Nup192, assessed by SEC, is indicated by the color-coded box above the respective mutated five-residue block. (**B**) The minimal Nup145N region sufficient for binding to Nup192 was identified by progressive peptide truncation from either terminus. Peptide boundaries are shown as gray bars above a stylized domain structure of Nup145N. The Nup145N^R1^ peptide is indicated by a red bar. Binding of SUMO-Nup145N peptides to Nup192, assessed by SEC, is summarized according to the measured effect: (+++) no effect, (++) weak effect, (+) moderate effect, and (-) abolished binding. (**C**) Three views of the Nup192•Nic96^R2^•Nup53^R1^•Nup145N^R1^ single particle cryo-EM structure in cartoon representation. The insets indicate regions magnified to illustrate the molecular details of (**D**) the Nup192-Nup53 and (**E**) the Nup192-Nup145N interactions. Red circles indicate residues found to be involved in the Nup192-Nup53 (*36*) and the Nup192-Nup145N interactions. (**F**) On the left, cyan squares indicate alanine substitutions of Nup145N residues and their effect on Nup145N binding to Nup192, assayed by SEC, is summarized as in (B). The Nup145N KKRMYKLRKR mutation that abolishes binding to Nup192 is indicated in red. Structure-driven alanine substitutions in the Nup145N binding site of Nup192 and their effect on the binding of Nup192 to SUMO-Nup145N^R1^ is tabulated on the right. (**G**) *KD*s determined by ITC. All experiments were performed in triplicate, with the mean and its standard error reported. Experiments for which no heat exchange to indicate binding could be detected are indicated as “no binding”. (**H**) SEC- MALS analysis of Nup192•SUMO-Nic96^R2^•Nup53•Nup145N and Nup192•SUMO- Nic96^R2^•Nup53•SUMO-Nup145N^R1^ complex formation and disruption by mutants. SEC-MALS profiles are shown for individual nups and for their preincubation. Measured molecular masses are indicated, with respective theoretical masses provided in parentheses.

Adding the newly mapped Nup145N^R1^ and our previously mapped minimal Nup53^R1^ fragment (residues 31-67) (*36*), we reconstituted an ∼220 kDa Nup192•Nic96^R2^•Nup145N^R1^•Nup53^R1^ complex and were able to obtain a single particle cryo-EM reconstruction at 3.2 Å resolution from a selected set of 484,910 particles (Fig. 4C and fig. S16). The Nup145N^R1^ binding site is proximal to the Nic96^R2^ binding site, at the midpoint of the question mark-shaped Nup192, where a hydrophobic pocket of *α−*helices *α*51, *α*53, and *α*54 anchors a Nup145N^R1^ MYKL motif (residues 633-636) that runs perpendicular to the long axis of the question mark, with its N-terminus oriented towards the N-terminus of Nic96^R2^ (Fig. 4E). For Nup53^R1^, the cryo-EM map only resolved the central phenylalanine-glycine (FG) dipeptide, which bound a hydrophobic Nup192 surface pocket at the top of the molecule formed by *α*-helices *α*19, *α*20, and *α*23. Key contacts involve L441 and W499 of Nup192 and F48A of Nup53, consistent with our previous observation that substitution of either of these residues abolishes the interactions (*36*) (Fig. 4D). Although we previously established that basic regions around the central Nup53 FG motif are also required for binding (*36*), the flanking residues were not resolved in the cryo-EM density for Nup53 or Nup145N. Overall, comparison of the Nup192 structures in complex with different linkers demonstrates that linker binding does not induce conformational rearrangements in the scaffold Nup192 (fig. S17).

Validation of the Nup192-Nup145N^R1^ interface through structure-guided mutagenesis and subsequent binding assessment confirmed the importance of the central hydrophobic Nup145N MYKL motif (Fig. 4F and fig. S18), but complete ablation of binding was only observed when the three flanking basic residues on either side were also mutated to alanine (10-residue KKR-MYKL-RKR mutant; K624A, K626A, R628A, M633A, Y634A, K635A, L636A, R643A, K650A, and R651A) (Fig. 4, F to H, and figs. S15, S18C, and S19). Reciprocally, targeting of the Nup145N^R1^ MYKL binding site in Nup192 identified a quadruple mutant (LIFH; L1071A, I1075A, F1125A, and H1126A) that specifically abolished Nup192 binding to Nup145N^R1^, but not Nic96^R2^ and Nup53 (Fig. 4, F and H, and figs. S19 to S21).

### Nup188 interaction with Nup145N

To identify the minimal Nup145N fragment and key residues required for Nup188 binding, we refined our previously identified binding region (residues 606-750) (*30*), utilizing the same alanine scanning and fragment truncation approach used to map its binding to Nup192. This resulted in an extended NUP145N^R2^ region between residues 640 and 732, within which mutation of 706-715 diminished Nup188^NTD^ binding (Fig. 5, A and B, and figs. S22 and S23). Consistent with our previous findings (*25*), we also confirmed that Nup188 does not bind Nup53, even in the presence of Nic96^R2^ and Nup145N (fig. S24).

**Fig. 5.**
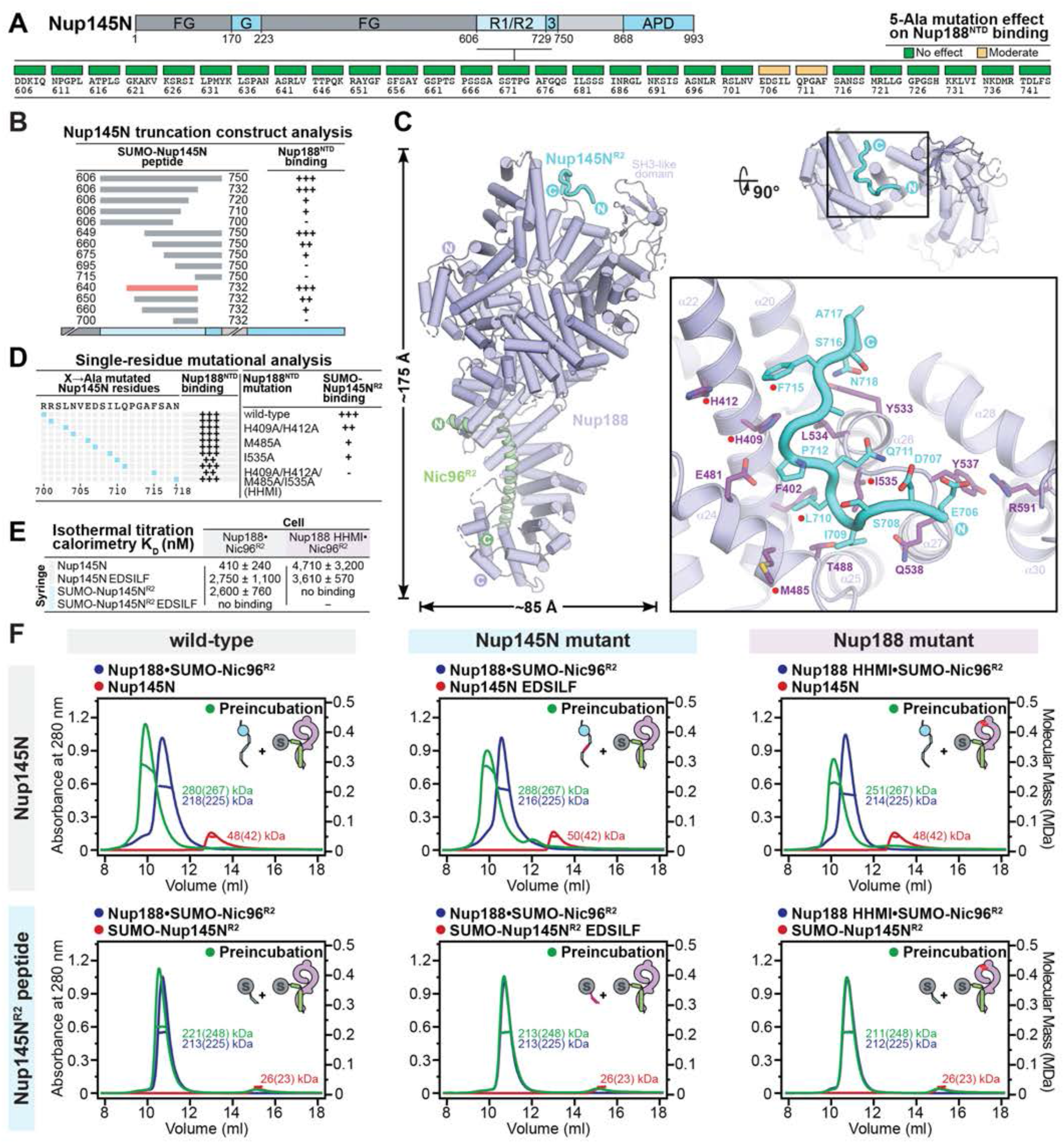
Structural and biochemical analyses of the Nup188-Nup145N interaction. (**A**) Domain structure of Nup145N, with the Nup145N primary sequence analyzed by 5-Ala scanning mutagenesis indicated. The effect of each 5-Ala substitution on Nup145N binding to Nup188^NTD^, assessed by SEC, is indicated by the color-coded box above the respective mutated five-residue block. (**B**) The minimal Nup145N region sufficient for binding to Nup188 was identified by progressive peptide truncation from either terminus. Peptide boundaries are shown as gray bars above a stylized domain structure of Nup145N. The Nup145N^R2^ peptide is indicated by a red bar. Binding of SUMO-Nup145N peptides to Nup188^NTD^, assessed by SEC, is summarized according to the measured effect: (+++) no effect, (++) weak effect, (+) moderate effect, and (-) abolished binding. (**C**) Two views of the Nup188•Nic96^R2^•Nup145N^R2^ single particle cryo-EM structure in cartoon representation. The inset indicates a region magnified to illustrate the molecular details of the Nup188-Nup145N interaction. Red circles indicate residues found to be involved in the Nup188-Nup145N interaction. (**D**) On the left, cyan squares indicate alanine substitutions of Nup145N residues and their effect on Nup145N binding to Nup188^NTD^, assayed by SEC, is summarized as in (B). Structure-driven alanine substitutions in the Nup145N binding site of Nup188 and their effect on the binding of Nup188^NTD^ to SUMO-Nup145N^R2^ is tabulated on the right. (**E**) *KD*s determined by ITC. All experiments were performed in triplicate, with the mean and its standard error reported. Experiments for which no heat exchange to indicate binding could be detected are indicated as “no binding”. (**F**) SEC-MALS analysis of Nup188•SUMO-Nic96^R2^•Nup145N and Nup188•SUMO-Nic96^R2^•SUMO-Nup145N^R2^ complex formation and disruption by mutants. SEC-MALS profiles are shown for individual nups and for their preincubation. Measured molecular masses are indicated, with respective theoretical masses provided in parentheses.

To structurally characterize the Nup188-Nup145N interaction, we reconstituted a ∼220 kDa Nup188•Nic96^R2^•Nup145N^R2^ complex and determined its structure by single particle cryo-EM. An initial set of 709,123 particles produced a reconstruction of Nup188•Nic96^R2^ at 2.4 Å resolution that presented weak excess density at the top of the question mark-shaped Nup188 molecule (Fig. 5C). Further 3D local classification identified a subset of 298,317 particles that yielded a reconstruction of Nup188•Nic96^R2^•Nup145N^R2^ at 2.8 Å resolution (Fig. 5C and fig. S25). Nup145N^R2^ buries residues I709, L710 and F715 in a hydrophobic cradle formed by *α*-helices *α*22, *α*24, and the loop connecting *α*-helices *α*27-*α*28, braced by the SH3-like domain. As with Nup192, only a portion of the Nup145N^R2^ peptide was resolved in the cryo-EM density (residues 706-718) indicating that Nup188 also forms multivalent interactions around a core Nup145N^R2^ motif, enhanced by weaker binding events distributed across flanking sequences on either side (Fig. 5C). As with Nup192, no significant conformational changes were observed between the different Nup188 structures in response to linker binding (fig. S26).

Structure-guided mutagenesis of Nup188 hydrophobic residues lining the Nup145N^R2^-binding cradle identified a Nup188 HHMI mutant (H409A, H412A, M485A, and I535A) that abolished binding to Nup145N^R2^, but not to Nup145N (Fig. 5Da and fig. S27). On the Nup145N side, mutation of two structurally resolved residues, L710A and F715A, moderately disrupted Nup188^NTD^ binding (Fig. 5D and fig. S28). Further systematic mutagenesis led to a Nup145N EDSILF mutant (E706A, D707A, S708A, I709A, L710A, and F715A), which respectively abolished and reduced Nup188 binding to Nup145N^R2^ and Nup145N (Fig. 5, E and F, and figs. S29 and S30). Overall, the greater tolerance of the Nup188- Nup145N interaction to binding site mutations compared to the Nup192-Nup145N interaction demonstrates an even greater reliance on promiscuous binding events in flanking regions dispersed well beyond the structurally resolved core.

### Comparison of the Nup192- and Nup188-linker complexes

The determination of full-length structures of both Nup192 and Nup188 scaffolds bound to their respective linkers permits a direct comparison of these two distantly homologous *α*-helical solenoids (∼28% sequence similarity). Although both structures share the same overall question mark-shaped architecture, composed of HEAT and ARM repeats, the Nup188 *α*-helical solenoid displays a tighter super-helical twist, resulting in an ∼10 Å narrower molecule with a compacted upper ring (Fig. 6, A and B). A Tower protrudes from the midpoint of the *α*-helical solenoid towards the Head subdomain in both structures, but extends further in Nup192 than the more compressed Nup188 version. Nic96^R2^ binds both scaffolds at the base of the question mark, but remarkably adopts different secondary structures to do so, switching between which requires breaking and re-forming *α*-helices (Fig. 6C). By contrast, Nup145N binds to different parts of the Nup192 and Nup188 question mark shape in the center or at the top, respectively. In contrast to Nup192, Nup188 does not bind Nup53, but, interestingly, the Nup145N^R2^ binding site at the top of Nup188 is nearly congruent with that of the Nup53^R1^ binding site on Nup192 (Fig. 6A).

**Fig. 6.**
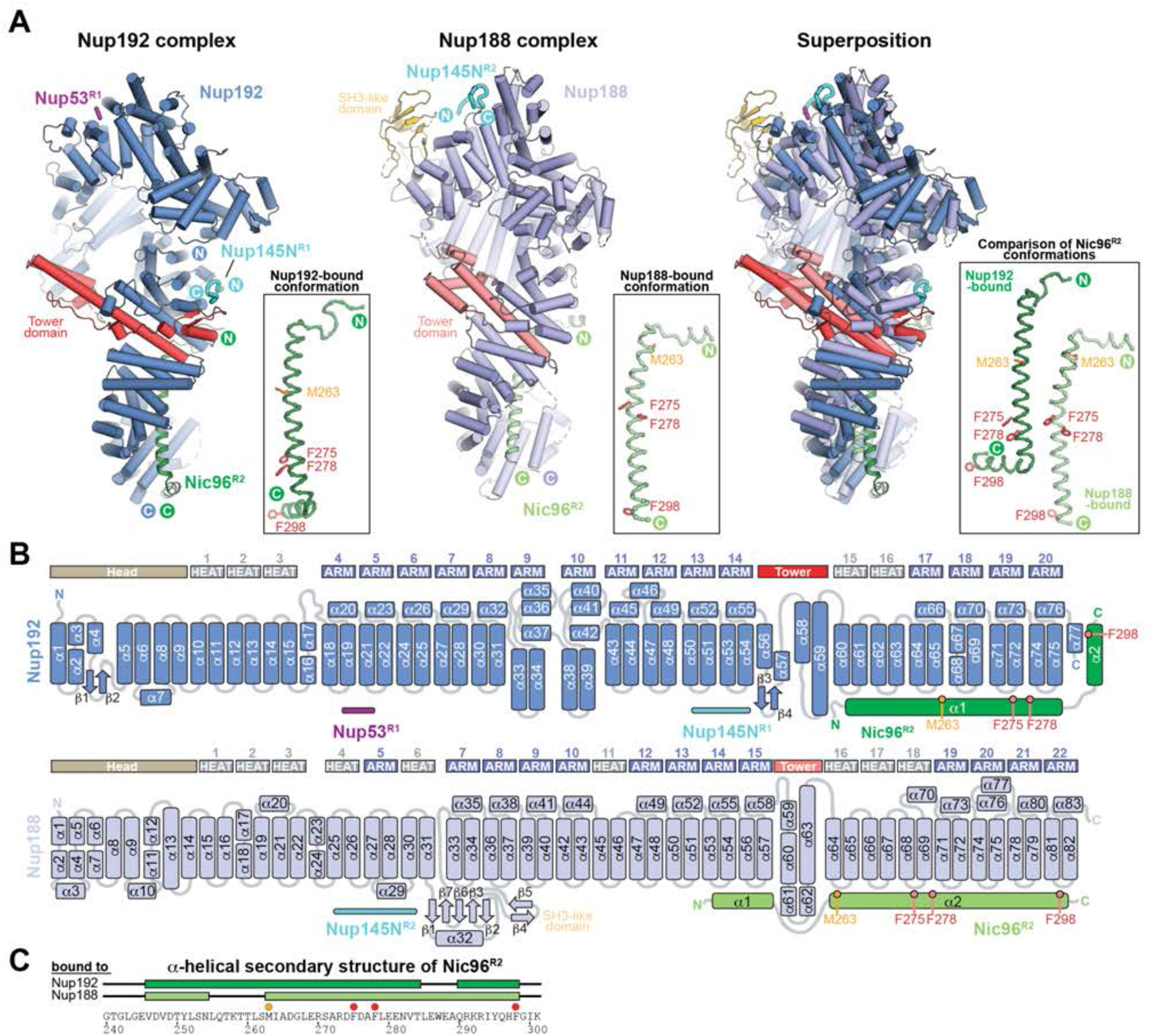
Structural comparison of the Nup192 and Nup188 scaffold-linker complexes. (**A**) Cartoon representations of the Nup192•Nic96^R2^•Nup53^R1^•Nup145N^R1^ and Nup188•Nic96^R2^•Nup145N^R2^ single particle cryo-EM structures, and their superposition. The Nup192 and Nup188 Tower domains (red and light red) and the Nup188 SH3-like domain (gold) are highlighted. In boxes, isolated and magnified views of conformations of Nic96^R2^ bound to Nup192 and Nup188 are shown to illustrate the Nic96^R2^ structural polymorphism. The two Nic96^R2^ conformations were superposed based on a common *α*-helical region between residues 263-284. Residues found to be involved in Nic96^R2^ binding to Nup192 and Nup188 (red) and residue M263 (orange), uniquely identifiable in anomalous difference Fourier maps derived from SeMet-derivatized Nup192^ΔHEAD^•Nic96^R2^ and Nup188•Nic96^R2^ crystals, are shown in ball-and-stick representation. (**B**) Schematic of the Nup192 and Nup188 fold architectures. Rounded rectangles and arrows represent *α*-helices and *β*-strands, respectively. (**C**) Schematic of the secondary structure that Nic96^R2^ assumes in binding Nup192 and Nup188. Green boxes and black lines above the primary sequence indicate *α*-helices and loops, respectively. Orange and red circles indicate residues highlighted in (A).

Our structures identify two distinct types of linker-scaffold interactions. Nic96^R2^ binds with high affinity, utilizing the same well-defined ∼60-residue motif in binding to both Nup192 and Nup188. On the contrary, Nup145N binds to Nup192 and Nup188 through extensive, overlapping, binding regions and a distinctive common binding mode, whereby both Nup145N interactions depend on a central ∼10-residue anchor motif that is structurally well-characterized, yet tight binding depends on extensive ∼20-60-residue N- and C-terminal flanking regions with high basic character. A similar binding mode is also observed for the Nup53-Nup192 interaction. The observation that flanking regions present in all three of these linker-scaffold interactions evade structural characterization suggests highly dynamic and promiscuous interactions with the scaffold surface. Notably, the previously characterized ∼14-residue Nup170-binding motif of Nup145N (residues 729-750) does not depend on binding-enhancing flanking regions, suggesting that the uncovered Nup145N/Nup53 mode of binding to the Nup192 and Nup188 scaffolds is a desirable evolutionary outcome and architectural principle of the NPC inner ring.

Together, these data complete the structural characterization of the linker-scaffold interactions that coordinate the larger structured nups of the inner ring into two distinct complexes, containing either Nup192 or Nup188. Nup145N interacts at distinct sites on Nup192 and Nup188 and can only form mutually exclusive interactions with either Nup192 and Nup170 or Nup188, which can be explained by the significant overlap in the mapped binding regions for Nup192 (R1, 616-683), Nup188 (R2, 640-732) and Nup170 (R3, 729-750). Whereas Nic96^R2^ binds to large surfaces of both Nup188 and Nup192 with ∼80 nM high affinity, binding of Nup145N and Nup53 utilizes relatively short central core motifs that form well-defined interactions with scaffold surface patches and are enhanced by extensively distributed binding through the N- and C-terminal flanking regions. This binding mode is reminiscent of Velcro, in which the accumulation of weak binding events builds towards a robust interaction that provides flexibility because there are manifold productive binding configurations in terms of spatial distribution or occupancy of binding residues. As an architectural principle, a Velcro-like binding between linkers and scaffolds is advantageous in accommodating movements of scaffold domains without fully breaking linker-scaffold interactions, as the NPC expands and contracts.

### Architecture of the *S. cerevisiae* linker-scaffold

The inner ring of the NPC adopts an overall doughnut-shaped architecture with eight-fold rotational symmetry across a nucleocytoplasmic axis and two-fold symmetry in the plane of the nuclear envelope (*29, 30*). Each of the 16 inner ring protomers were proposed to consist of a Nup192 inner ring complex (IRC) and a Nup188 IRC, located at the equatorial and peripheral positions, respectively (*29, 30*). Whereas quantitative docking of the folded scaffold nups Nup170, Nic96, Nup192, Nup188 and CNT into cryo-ET maps of intact NPCs revealed their positioning to form four concentric cylinders, the linker network that connects them has remained elusive. Combined with our previously determined structures of Nup170•Nup53^R3^, Nup170•Nup145N^R3^, Nic96•Nup53^R2^, and Nic96^R1^•CNT (*29, 30*), the Nup192•Nic96^R2^•Nup145N^R1^•Nup53^R1^ and Nup188•Nic96^R2^•Nup145N^R2^ structures now allow us to identify the locations of the entire linker-scaffold network showing all binding sites in the NPC’s inner ring (Fig. 1, C and D).

With full-length experimental structures available, we statistically scored the fit of a million randomly placed and locally refined resolution-matched densities simulated from Nup192•Nic96^R2^•Nup145N^R1^•Nup53^R1^ and Nup188•Nic96^R2^•Nup145N^R2^ structures against a ∼25 Å *in situ* cryo-ET map of the *S. cerevisiae* NPC that captures a dilated inner ring state (fig. S31) (*31*). Nup192 and Nup188 were quantitatively docked with high confidence into inner ring equatorial and peripheral question mark-shaped cryo-ET densities, respectively, confirming previous proposals (*31, 32*). Structural differences, including the reduced width due to tighter superhelical winding of the Nup188 *α*-helical solenoid and of the presence of an SH3-like domain in Nup188, closely matched discriminating features of the cryo-ET density (fig. S32). Linker-scaffold Nup170•Nup145N^R3^•Nup53^R3^, Nic96^SOL^•Nup53^R2^, and CNT•Nic96^R1^ structures were placed by superposition with the previously determined composite model of the *S. cerevisiae* NPC inner ring (Fig. 7 and fig. S33) (*29, 30*).

**Fig. 7.**
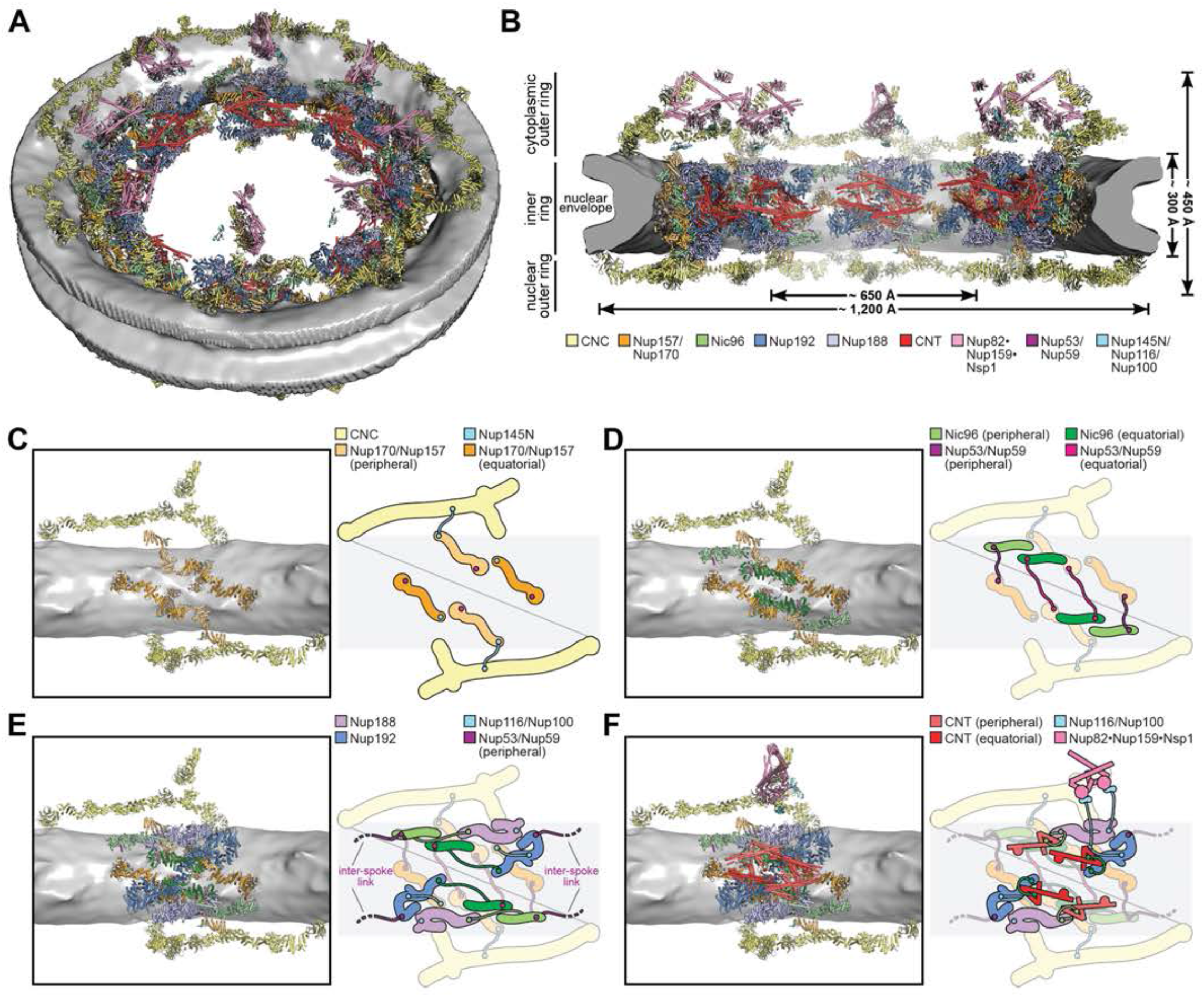
Architecture of the *S. cerevisiae* NPC linker-scaffold. Composite structure of the *S. cerevisiae* NPC generated by docking linker-scaffold structures into an ∼25 Å sub-tomogram averaged cryo-electron tomography (cryo-ET) map of the *in situ* imaged *S. cerevisiae* NPC (EMD-10198) (*31*), viewed from (**A**) above the cytoplasmic face, and (**B**) a cross-section of the central transport channel. The nuclear envelope is rendered as a gray isosurface and the docked structures are represented as cartoons. (**C**-**F**) Cartoon representation of a single spoke’s layered architecture and schematic illustrating linker paths determined by assuming maximum distance and linker copy parsimony. Linker binding sites on scaffold nup surfaces are schematically indicated with colored circles. (C) Nup145N can simultaneously bind the coat nup complex (CNC) component Nup145C in the outer rings and Nup157 in the inner ring. (D) Nup53/Nup59 can link equatorial Nic96^SOL^ with peripheral Nup157 and peripheral Nic96^SOL^ with equatorial Nup170, across the spoke midplane. (E) Nup192 and Nup188 bind to equatorial and peripheral Nic96^R2^, respectively. Nup116/Nup100 can link Nup192 and the equatorial Nup170. Nup53/Nup59 can link Nup192 and the peripheral Nic96 of adjacent spokes. (F) The CNT is recruited by Nic96^R1^. Nup116/Nup100 can simultaneously bind to the cytoplasmic asymmetric Nup82•Nup159•Nsp1 complex and to Nup188 or Nup170 in the inner ring.

The docking shows that Nup192 and Nup188 are linker-scaffold hubs crucially positioned between the membrane-coating Nup157/Nup170 and the central transport channel-interfacing CNT layers of the inner ring. The placement of Nup170•Nup145N^R3^•Nup53^R3^ in the Nup157/Nup170 layer depicts it as a pegboard onto which linkers are anchored (Fig. 7C). Because the integrity of stochiometric Nup192 and Nup188 inner ring complexes in solution depends on linker-scaffold interactions, we wondered if the same interactions could be identified in the composite structure of the inner ring. Indeed, Nic96^R2^ bound to Nup192 and Nup188, connected to respective equatorial and peripheral copies of Nic96^SOL^ and CNT•Nic96^R1^ (Fig. 7, E and F). Nic96^SOL^-bound Nup53^R2^ connected to Nup170-bound Nup53^R3^, Nup192-bound Nup53^R1^ connected to Nic96^SOL^-bound Nup53^R2^, and Nup192-bound Nup145N^R1^ connected to Nup170-bound Nup145N^R3^ (Fig. 7, D and E). Because linkers between the different Nup145N scaffold binding regions in the same molecule are not sufficiently long to span the physical distances between binding sites in the composite structure, Nup188-bound Nup145N^R2^ and peripheral Nup170-bound Nup145N^R3^ could not be connected to other inner ring Nup145N linker-scaffold binding sites, consistent with the mutually exclusive binding events observed in solution (*30*). Importantly, the biochemical incompatibility corresponding to physically impossible connections suggests that the assembly of inner ring complexes is *a priori* encoded by their biochemical properties to prevent the formation of spurious linker-scaffold complexes.

We considered whether connections between linker regions bound to scaffold surfaces were constrained to nearest neighbors by the physical length of the linker spanning two binding sites. We found that to be true for the peripheral and equatorial connections mediated by the N-terminal Nic96 linker that connects to Nup192/Nup188 and the CNT (fig. S34) and the Nup145N connection between Nup192 and the equatorial Nup170 (fig. S34F). As explained above, Nup188 and the peripheral Nup170 each are required to bind their own Nup145N copy. Although a stretched Nup53 linker can reach at least three different Nup170 positions from each peripheral and equatorial Nic96^SOL^, the nearest neighbor is only ∼50 Å away in both cases, in contrast to all the other copies that are at least twice as distant. Therefore, Nup53 is expected to connect the peripheral Nup170 with the equatorial Nic96^SOL^, and *vice versa*, the equatorial Nup170 with the peripheral Nic96^SOL^, both across the inner ring midplane (fig. S34, B and C). The Nup53-mediated connection between Nup192 and Nic96^SOL^ is the only linker-scaffold interaction that occurs between adjacent spokes (fig. S34G), as we previously proposed (*30*). This linker-scaffold arrangement suggested the Nup53^RRM^ homodimer is placed between spokes, despite the lack of corresponding cryo-ET density.

Together, these data elucidate the architecture of the *S. cerevisiae* inner ring linker-scaffold. Apart from Nup53-mediated spoke bridging, all linker-scaffold connections in the *S. cerevisiae* inner ring occur within the same spoke, thereby allowing inter-spoke gaps to form. Thus, the linker-scaffold architecture provides a molecular explanation for the inner ring’s ability to exist in constricted and dilated states (*31*). Future work needs to address the forces that govern the inner ring’s reversible dilation and constriction, with membrane tension being the most likely candidate.

### *S. cerevisiae* linker-scaffold is robust and essential

Due to ancestral gene duplication events in *S. cerevisiae*, there are several linker and scaffold nup paralogs, including Nup145N paralogs Nup116 and Nup100, Nup53 paralog Nup59, and Nup170 paralog Nup157 (*2*). Evolutionary retention of distinct inner ring paralogs suggests they are either speciated to occupy unique positions and roles within the assembled NPC, or are broadly redundant and capable of accommodating several binding functions at once. A multispecies sequence alignment uncovered that the *S. cerevisiae* Nup100 and Nup116 paralogs contained sequences homologous to the Nup192, Nup188, and Nup170 binding regions characterized in *C. thermophilum* Nup145N, but only Nup116 possesses the Gle2 binding site (GLEBS) motif that recruits the mRNA export factor Gle2 (fig. S35) (*44*). In addition, all three paralogs contain C-terminal domains (CTDs) that were previously shown to form interactions with either Nup145C or Nup82, which are components of the CNC and cytoplasmic filaments, respectively (*30, 45, 46*).

To interrogate the function of individual scaffold binding regions in the linker Nup116, we first established a *S. cerevisiae* minimal linker strain in which mutations would not be complemented by the Nup145N and Nup100 paralogs (Fig. 8). Because Nup145N results from the cleavage of a Nup145 precursor protein whose other product Nup145C incorporates into the CNC and is required for survival (*47*), all experiments involving Nup145 deletion were carried out in the presence of *NUP145C* ectopically expressed from a centromeric plasmid. Building on previous studies of synthetic lethality (*48*), we systematically established that all double knockout combinations of *NUP100*, *NUP116* and *NUP145* can be rescued by a chimeric *NUP116-NUP145C* (fig. S36). Consequently, we found that *NUP116* is able to complement the *nup100Δnup116Δnup145Δ* strain (in the presence of ectopic *NUP145C*), albeit with background nuclear mRNA retention above wildtype levels (Fig. 8E, figs. S36, C and D, and S37). Unlike previously described for Nup100 (*44*), Nup145N chimeras containing either a GLEBS motif or the entire GLEBS-containing FG-repeat domain of Nup116 could not rescue the *nup100Δnup116Δnup145Δ/NUP145C* strain, suggesting an essential function for the R1/R2 regions that are uniquely absent in Nup145N but not the other paralogs (fig. S36, E and F).

**Fig. 8.**
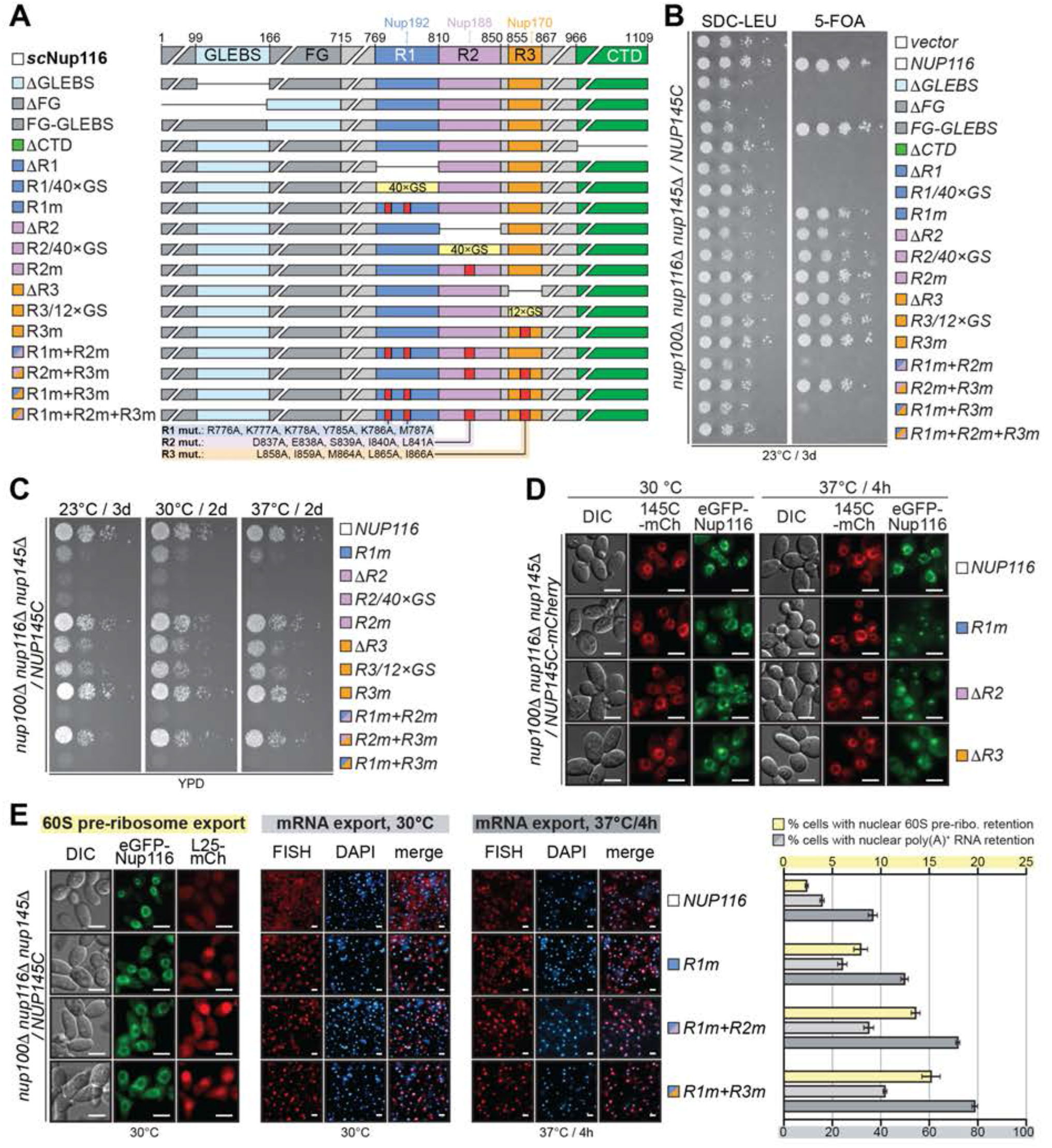
Functional *in vivo* dissection of Nup116 in a minimal *S. cerevisiae* linker strain. (**A**) Domain structure of Nup116 variants transformed into *nup100*Δ*nup116*Δ*nup145*Δ *S. cerevisiae* strains, with color-coded squares indicating the functional region targeted by each mutation; pale cyan, Gle2-binding sequence (GLEBS); dark gray, FG repeats; green, C-terminal domain (CTD); blue, Nup192-binding region (R1); purple, Nup188-binding region (R2); orange, Nup170-binding region (R3). (**B**) Viability analysis of a ten-fold dilution series of a *nup100*Δ*nup116*Δ*nup145*Δ*/NUP145C* strain expressing Nup116 variants and subjected to 5-FOA selection for loss of the rescuing wildtype plasmid. (**C**) Growth analysis at different temperatures of a ten-fold dilution series of a *nup100*Δ*nup116*Δ*nup145*Δ*/NUP145C* strain expressing viable Nup116 variants. (**D**) Subcellular localization at permissive and growth-challenging temperatures of a representative subset of eGFP-Nup116 variants in a *nup100*Δ*nup116*Δ*nup145*Δ*/NUP145C-mCherry* strain. (**E**) Representative images of the subcellular localization 60S pre-ribosomal export reporter Rpl25-mCherry (*left*) and poly(A)^+^ RNA at 30 °C and 37 °C (*middle*) in the presence of a representative subset of Nup116 variants. Quantitation of the proportion of cells (n>500) with poly(A)^+^ RNA and 60S pre-ribosome nuclear retention is shown (*right*). Experiments were performed in triplicate, and the mean with the associated standard error reported. All scalebars are 5 μm.

Based on this result, we systematically mutated all functional elements in the Nup116 sequence, and found that the GLEBS, FG and CTD are essential (Fig. 8, A and B, and fig. S37, A to C). We targeted the Nup192, Nup188 and Nup170 scaffold binding regions (R1, R2 and R3, respectively) with three types of mutations: deleting entire binding regions (*ΔR1*, *ΔR2*, and *ΔR3*), replacing entire binding regions with a glycine-serine (GS)-linker of equivalent length (*R1/40×GS*, *R2/40×GS*, and *R3/12×GS*), or substituting sequence-conserved residues shown to disrupt binding *of the C. thermophilum* Nup145N to the respective scaffolds (*R1m*, *R2m*, and *R3m*) (Fig. 8A and fig. S35). All Nup116 variants were expressed at comparable levels (fig. S37, C and E). After establishing viability, non-lethal mutants were further assayed for growth rates at different temperatures, localization of eGFP-Nup116, and nuclear retention of mRNA and 60S pre-ribosomes. Mutations in each of the Nup192, Nup188 and Nup170 binding regions resulted in altered phenotypes, albeit to different extents (Fig. 8 and fig. S37). Deletions and GS-linker replacements, being aggressive types of mutations, were lethal if targeting R1, and impacted growth and mRNA/60S pre-ribosome export if introduced in R2 and R3. The less aggressive combination of substitutions, *R1m*, allowed us to perturb R1 with significant yet non-lethal phenotypic effects, which were further exacerbated through combination with R2m (*R1m+R2m*) or R3m (*R1m+R3m*), culminating with the lethal *R1m+R2m+R3m* triple mutation (Fig. 8 and fig. S37). Interestingly, all Nup116 mutations resulted in temperature-dependent loss of eGFP-Nup116 from the nuclear envelope rim and concomitant emergence of eGFP-Nup116 foci, as previously reported (Fig. 8C and fig. S37F) (*45, 49, 50*). These results demonstrate the essential linker function of Nup116 in the *S. cerevisiae* NPC and illustrate the physiological relevance of the biochemically and structurally characterized Nup192, Nup188 and Nup170 binding regions.

Perturbing the linker-scaffold interactions on the scaffold side, absence of Nup192 results in lethality, matching the severe consequences of deleting the Nup192 binding region in Nup116 (Fig. 9B). Additionally, we had previously found that truncation of just the Nup192 Tail region has a deleterious effect, attributable to the perturbation of the Nic96^R2^ binding site (*36*). Our new Nup192-Nic96^R2^ structures provide a molecular explanation for this result, showing Nic96 binding at the Nup192 Tail and the base of the Tower (Fig. 6, A and B). Based on sequence conservation and the insight from our structural and biochemical characterization, we transposed the *C. thermophilum* Nup192 *LAF* and *LIFH* substitutions that ablated binding to Nic96^R2^ and Nup145N^R1^, respectively, *in vitro* into *S. cerevisiae* Nup192, along with designing aggressive *ΔTail-Tower* and *Tail-Tower* truncations deleting the respective linker binding subdomains (Fig. 9A and fig. S38). Individually, the *LAF* and *LIFH* mutations had no effect on growth, although wildtype Nup192 could outcompete both for localization to the nuclear envelope rim. However, the combination of *LAF* and *LIFH* substitutions (*LAF+LIFH*) and both Nup192 truncations failed to rescue the lethal *nup192Δ* phenotype (Fig. 9, A to D, and fig. S39).

**Fig. 9.**
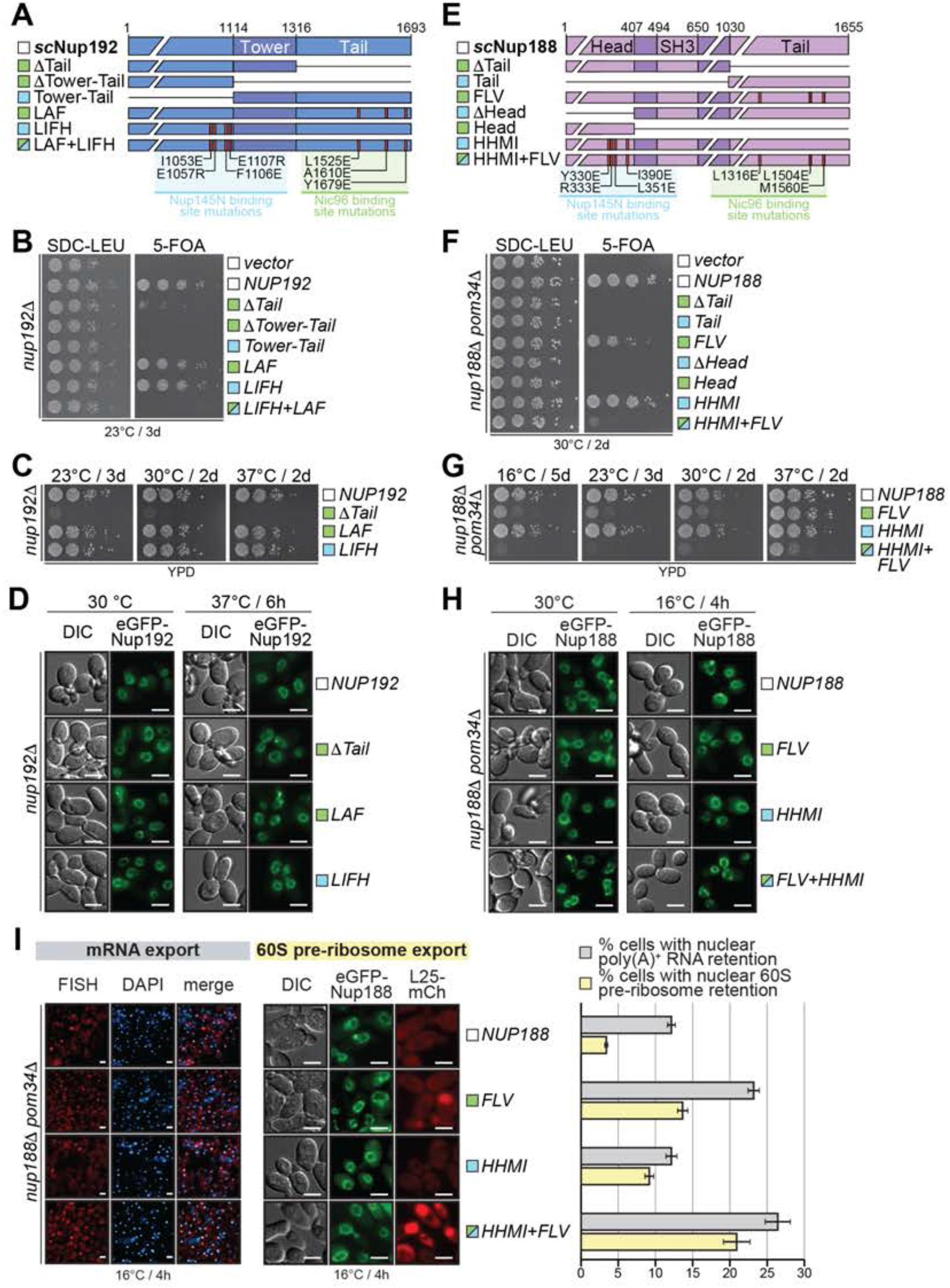
Functional *in vivo* validation of the Nup192 and Nup188 linker binding sites. (**A**, **E**) Domain structure of (A) Nup192 variants introduced into the *nup192*Δ *S. cerevisiae* strain, and (E) Nup188 variants introduced into the *nup188*Δ*pom34*Δ *S. cerevisiae* strain, with colored squares indicating whether the mutation affects the Nup145N binding site (cyan), the Nic96 binding site (pale green), or both (cyan and pale green). (**B**, **F**) Viability analysis of a ten-fold dilution series of (B) a *nup192*Δ strain transformed with Nup192 variants, and (F) a *nup188*Δ*pom34*Δ strain transformed with Nup188 variants, subjected to 5-fluoroorotic acid (5-FOA) selection for loss of the rescuing wildtype plasmids. (**C**, **G**) Growth analysis at different temperatures of a ten-fold dilution series of (C) a *nup192*Δ strain expressing viable Nup192 variants, and (G) a *nup188*Δ*pom34*Δ strain expressing viable Nup188 variants. (**D**, **H**) Subcellular localization at permissive and growth-challenging temperatures of (D) eGFP-Nup192 variants, and (H) eGFP-Nup188 variants. (**I**) Representative images of the subcellular localization of poly(A)^+^ RNA (*left*) and 60S pre-ribosomal export reporter Rpl25-mCherry (*middle*) in the presence of different Nup188 variants in a *nup188*Δ*pom34*Δ strain. Quantitation of the proportion of cells (n>500) with poly(A)^+^ RNA and 60S pre-ribosome nuclear retention is shown (*right*). Experiments were performed in triplicate, and the mean with the associated standard error reported. All scalebars are 5 μm.

Because *NUP188* is not an essential gene, we analyzed the effect of transposed Nup188 *FLV* and *HHMI* substitutions and truncations of its Nic96^R2^ and Nup145N^R2^ binding domains on the viability and growth rate of previously identified synthetic lethal *nup188Δpom34Δ* and *nup188Δpom152Δ* strains (Fig. 9E and fig. S40) (*51, 52*). Though all truncations were lethal, the *ΔTail* and *ΔHead* mutants still showed nuclear envelope rim staining, suggesting that either Nic96^R2^ and Nup145N^R2^ homolog binding site is sufficient for NPC localization (Fig. 9F and fig. S41). The *FLV* mutation was lethal to the *nup188Δpom152Δ* strain and led to slow growth with mRNA/60S pre-ribosome export defects in the *nup188Δpom34Δ* strain. The *HHMI* mutation alone resulted in only a moderate 60S pre-ribosome export defect, but the combined *HHMI+FLV* mutations further amplified the *FLV* mutant phenotype (Fig. 9, G to I, and fig. 41, E and F). Interestingly, mutant phenotypes were intensified at lower temperatures, consistent with previous reports of cold-sensitive *NUP188* mutations (*53*).

Finally, we transposed the *FFF* substitutions of evolutionarily conserved hydrophobic residues that abolished Nic96^R2^ binding to Nup192 and Nup188 *in vitro* into *S. cerevisiae* Nic96 and also constructed a *ΔR2* mutant lacking Nic96^R2^ (Fig. 10 and fig. S42). Surprisingly, neither mutation resulted in a significant phenotype when introduced into a *nic96Δ* strain (Fig. 10, A to D and fig. S43). The composite structure of the NPC linker-scaffold suggests that Nic96^R2^ binding to Nup192 and Nup188 restricts the diffusive path of the N-terminal Nic96 linker, thereby correctly positioning the Nic96^R1^ assembly sensor that recruits the CNT complex (Fig. 10G). Because the inner ring midplane is lined with a narrow band of 32 CNT copies that project FG repeats into the central transport channel, we reasoned that their mispositioning would affect the spatial distribution and local concentration of FG repeats, with consequences on nucleocytoplasmic transport. Therefore, we replaced the Nic96^R2^ region with GS-linkers matching the number of residues (*R2/66×GS*) or approximating its *α*-helical length (*R2/32×GS*) (Fig. 10A). Despite not affecting expression levels, nuclear envelope rim localization, and CNT recruitment by the Nic96^R1^ region, the *R2/66×GS* and *R2/32×GS* mutations respectively resulted in lethal and severely deleterious effects on growth and mRNA/60S pre-ribosome export (Fig. 10 and fig. S43). Taken together, these data demonstrate the physiological relevance of Nic96^R2^ binding to Nup192 and Nup188 and its role in restricting the diffusive path of the Nic96 N-terminal linker region.

**Fig. 10.**
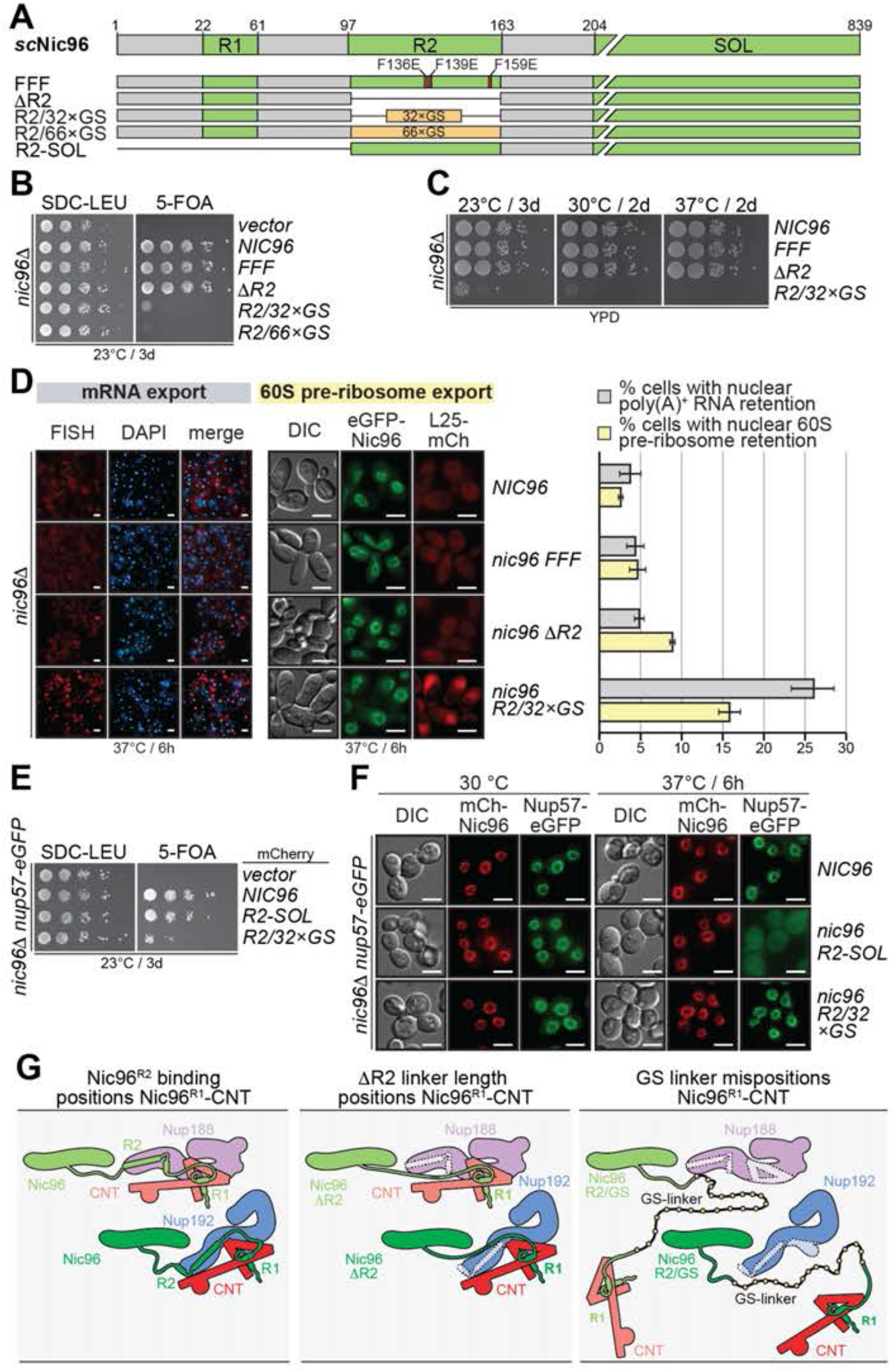
The Nup192/Nup188-binding Nic96^R2^ linker region spatially restricts CNT localization. (**A**) Domain structure of Nic96 variants introduced into the *nic96*Δ *S. cerevisiae* strain. (**B**) Viability analysis of a ten-fold dilution series of a *nic96*Δ strain transformed with Nic96 variants and subjected to 5-FOA selection for loss of the rescuing wildtype plasmid. (**C**) Growth analysis at different temperatures of a ten- fold dilution series of a *nic96*Δ strain expressing viable Nic96 variants. (**D**) Representative images of the subcellular localization of poly(A)^+^ RNA (*left*) and 60S pre-ribosomal export reporter Rpl25-mCherry (*middle*) in the presence of different Nic96 variants. Quantitation of the proportion of cells (n>500) with poly(A)^+^ RNA and 60S pre-ribosome nuclear retention is shown (*right*). Experiments were performed in triplicate, and the mean with the associated standard error reported. (**E**) Viability analysis of a ten-fold dilution series of a *nic96*Δ*nup57-eGFP* strain transformed with mCherry-Nic96 variants and subjected to 5-FOA selection for loss of the rescuing wildtype plasmid. (**F**) Subcellular localization at permissive (30 °C) and growth-challenging (37 °C) temperatures of the CNT subunit Nup57-eGFP in the presence of different mCherry-Nic96 variants. (**G**) Nic96^R2^ positions the CNT in the central transport channel. The N-terminal Nic96 linker region presents scaffold-binding regions R1 and R2. Nic96^R1^ recruits a CNT to the NPC’s inner ring. Binding of Nic96^R2^ to Nup192 and Nup188 positions the Nic96^R1^-anchored CNT to the midplane of the inner ring (*left*). The deletion of Nic96^R2^ in the *ΔR2* variant ablates the spatial restraint, but is tolerated by maintaining CNT location close to the inner ring midplane, confounding the effect from the loss of Nup188/Nup192 binding (*middle*). In the Nic96 *R2/32×GS* and *R2/66×GS* variants, the introduction of glycine-serine (GS)-linkers in lieu of Nic96^R2^ results in the mispositioning of Nic96^R1^-bound CNTs and the FG-repeat mass they contribute to the central transport channel, thus hampering nucleocytoplasmic transport (*right*). All scalebars are 5 μm.

Overall, our data disambiguate the positioning of Nup192 and Nup188 in the inner ring and elucidate the architecture of the linker-scaffold in the *S. cerevisiae* NPC. Through comprehensive experiments that dissect the various linker regions and scaffold binding sites in *S. cerevisiae*, we established the physiological relevance of our biochemical and structural data. Nup192 and Nup188 are keystone scaffold hubs of the inner ring that integrate connections between the Nup157/Nup170 membrane-coating layer and the central transport channel-interfacing CNT layer through respective interactions with Nup145N paralogs and the N-terminal Nic96 linker region.

The fact that *NUP188* is not an essential gene while *NUP192* is, raises the question about what differences in the linker-scaffold interactions prevent Nup188 from taking Nup192’s role in the inner ring. Indeed, the loss of Nup188 can be compensated by recruitment of additional Nup192 to the NPC, but not *vice versa* (*54*). Our analysis concludes that unlike Nup188, Nup192 can bind to both Nup53 and Nup145N concomitantly bound with Nup170. The viability of the *nup53Δnup59Δ* strain (*55*) implies that connection between Nup192 and the equatorial Nup170, attaching the Nup192 complex to the Nup157/Nup170 coat of the inner ring, is an essential architecture feature that distinguishes Nup192 from Nup188. The wildtype phenotype of the *nup53Δnup59Δ* strain precludes analysis of the Nup53/Nup59 interactions in *S. cerevisiae*. However, this fact coupled with our knockout of all but one of the Nup145N paralogs highlight the robustness of the *S. cerevisiae* inner ring architecture, which is able to tolerate a considerable loss of linker-scaffold interactions. Robustness is also found in nup-nup interactions, whereby perturbing a linker-scaffold interaction requires multiple residue substitutions in both structured motifs and flanking linker regions. Moreover, FG-repeat sequences of the Nup145N paralogs contribute to the overall binding to scaffold proteins (*43, 50, 56-58*). Given that *S. cerevisiae* contains three Nup145N paralogs and two Nup53 paralogs, future work needs to elucidate whether their localization in the NPC is deterministic and possibly fine-tuning the function of the NPC. A prominent example may be the asymmetric localization of Nup116, which binds most tightly to the cytoplasmic filament component Nup82 and was previously shown to be specifically anchored at the cytoplasmic face of the NPC (*31, 45, 48, 59*).

### Evolutionary conservation of the human linker-scaffold

Despite low sequence conservation, composite structures of the human and *S. cerevisiae* NPC reveal an identical positioning of the scaffold nups, suggesting that the linker-scaffold architecture is evolutionarily conserved (*29–32*). Specifically, the human linker-scaffold interactions, the topology of scaffold-binding regions in the linkers, and the location of linker-binding sites in the scaffolds are expected to match the ones established with the *C. thermophilum* nups (*25, 28, 30, 35, 36*). To test this prediction, we carried out a systematic nup-nup interaction analysis utilizing a full complement of the human inner ring nup orthologs: NUP53 (ctNup53), NUP98 (ctNup145N), NUP93 (ctNic96), NUP155 (ctNup170), NUP188 (ctNup188), NUP205 (ctNup192), NUP62 (ctNsp1), NUP58 (ctNup49) and NUP54 (ctNup57) (Fig. 11, A and B). We developed bacterial expression and purification protocols for all recombinant human nups, apart from NUP205 and NUP188, which were expressed in *S. cerevisiae*. For linker nups, we generated truncation and sequence variants, aided by multispecies sequence alignments that either identified conserved linker motifs in all eukaryotes or only in fungal or metazoan species (figs. S35, S42, and S44 to S46).

**Fig. 11.**
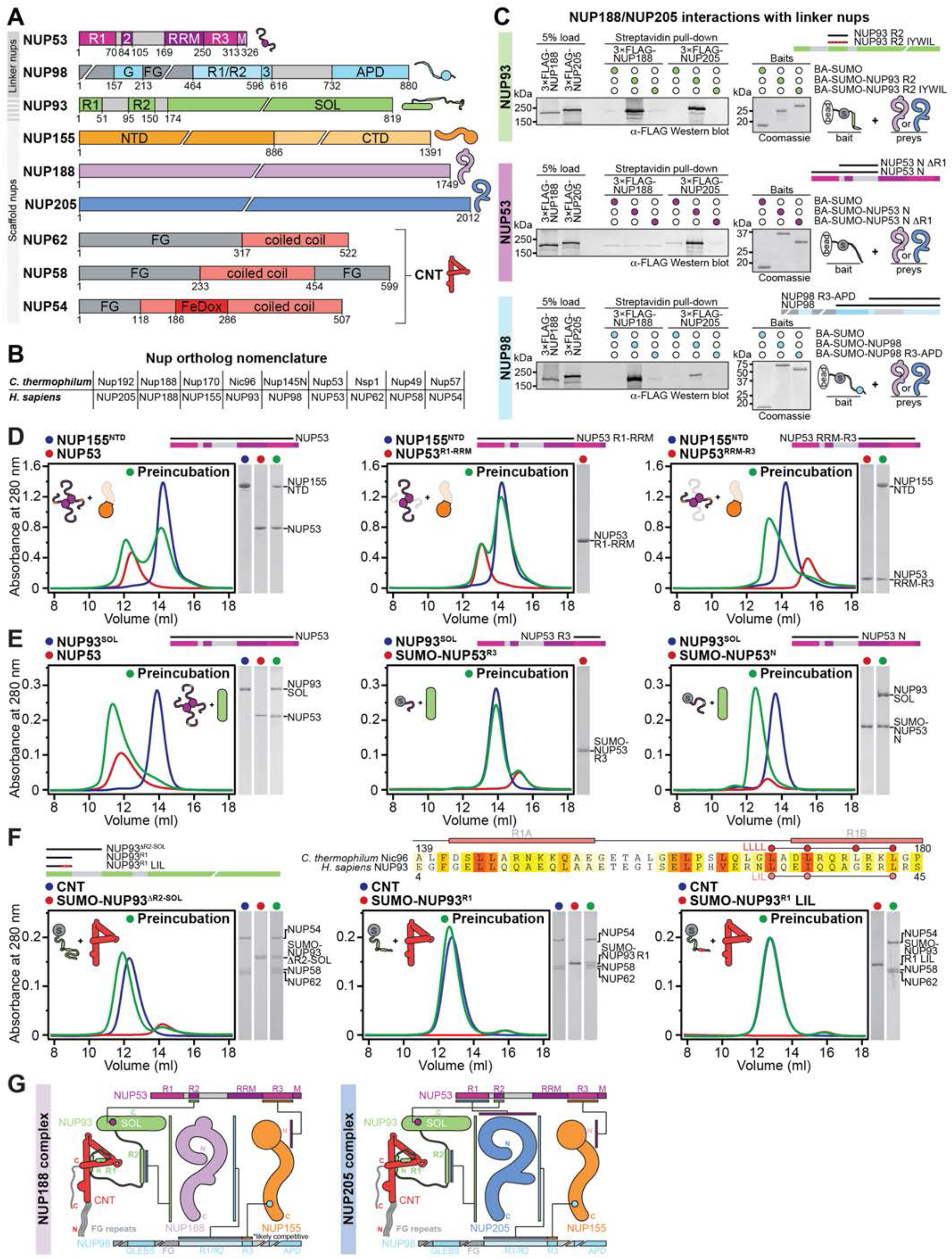
Evolutionary conservation of the human linker-scaffold network. (**A**) Domain structures of the human inner ring linker and scaffold nups. NUP93 consists of linker (residues 1-173) and scaffold (residues 174-819) regions. The human CNT components NUP62, NUP58, and NUP54 are drawn as a unit. (**B**) Nomenclature of *C. thermophilum* and human nup orthologs. (**C**) Pulldown analysis of NUP188 and NUP205 interactions with NUP93^R2^, NUP53, and NUP98. Biotinylated His6-Avi-SUMO (BA-SUMO) fused with linker NUP93^R2^, NUP53 and NUP98 fragments were loaded onto streptavidin-conjugated magnetic beads and used as baits to pull down 3×FLAG-NUP188 or 3×FLAG-NUP205 from lysate. Linker nup truncations and substitutions are indicated with black lines and red circles above the stylized domain structures, respectively. Load and pulldown fractions were resolved by SDS-PAGE and visualized by western blot. Baits were resolved by SDS-PAGE and visualized by Coomassie staining. (**D**-**F**) Analysis of the human linker-scaffold binding topology. For each experiment schematics of linker nup truncations and substitutions are indicated as in (C). SEC profiles are shown for individual nups or nup complexes and their preincubation, along with Coomassie-stained SDS-PAGE gel slices of peak fractions. In (F), a sequence alignment of orthologous *C. thermophilum* Nic96^R1^ and human NUP93^R1^ regions shows the evolutionary conservation of the LLLL mutation that abolishes binding of *C. thermophilum* Nic96^R1^ to the CNT (*25*). Residues are colored according to a multispecies sequence alignment, from white (less than 55 % similarity), to yellow (55 % similarity), to red (100 % identity) using the BLOSUM62 substitution matrix. Lines and rectangles above the sequence indicate loop and *α*-helical segments of Nic96^R1^, respectively. (**G**) Schematic summary of the established linker-scaffold interactions in the complexes organized about the NUP188 and N205 scaffold hubs. Colored bars connected by black lines indicate interactions between nup regions.

In *C. thermophilum*, Nup53^R3^ and Nup145N^R3^ bind to the Nup170 N-terminal *β*-propeller and C-terminal *α*-helical solenoid domains, respectively and Nup53^R2^ binds the Nic96^SOL^ scaffold (*30, 35*). We have previously elucidated the molecular details of the human NUP155^CTD^-NUP98^R3^ interaction (*30*). SEC interaction assays showed that, like its fungal ortholog, NUP53 binds to NUP155^NTD^ via a C-terminal region located between the RRM-like domain and the predicted C-terminal amphipathic *α*-helix (Fig. 11D, and fig. S47). Although NUP53 does not contain a motif resembling the *C. thermophilum* Nup53^R2^ amphipathic *α*-helix that interacts with Nic96^SOL^, we find that it binds to NUP93^SOL^ via a region N-terminal to the RRM-like domain (Fig. 11E and fig. S48). Next, we were also able to reconstitute an intact, stoichiometric, heterotrimeric human CNT (*hs*CNT), consisting of the C-terminal coiled-coil domains of NUP62 (residues 317-552), NUP58 (residues 233-454), and NUP54 (residues 118-507). *hs*CNT formed an interaction with a sequence conserved NUP93^R1^ region (residues 2-51), enhanced by a C-terminal flanking region (residues 52-95), and abolished by a LIL substitution (L33A, I36A, and L43A) of residues homologous to the previously reported *C. thermophilum* Nic96^R1^ LLLL mutation that abolishes CNT binding (Fig. 11F and fig. S49) (*25*).

We probed the interactions between the NUP205 and NUP188 keystone scaffolds and the NUP93^R2^, NUP53, and NUP98 linkers by pulling down overexpressed NUP205 and NUP188 from lysate with linker baits (Fig. 11C). The NUP93^R2^ fragment, homologous to Nic96^R2^, bound to both NUP205 and NUP188. Both interactions were abolished by a NUP93^R2^ IYWIL mutation, combining substitutions Y128E and L148E (homologous to F278 and F298 of the Nic96^R2^ FFF mutant) with other conserved residues I114E, W137E, and I144E on the predicted hydrophobic face NUP93^R2^ amphipathic *α*-helix (Fig. 11C and fig. S42). As in *C. thermophilum,* NUP53 did not bind to NUP188, and its interaction with NUP205 required an N-terminal R1 region (residues 1-70). Lastly, the NUP98 interaction with both NUP205 and NUP188 was mapped to a region topologically homologous to the Nup192/Nup188-binding R1/R2 regions of Nup145N (Fig. 11C).

Together, these data establish that the linker-scaffold is evolutionarily conserved from *C. thermophilum* to humans, including the linker-binding sites in the scaffolds and the topology of the scaffold-binding regions in the linkers (Fig. 11G).

### Biochemical and structural analysis of the human NUP93-NUP53 interaction

Our dissection of the human linker-scaffold interaction network identified an interaction between NUP93^SOL^ and a NUP53 region N-terminal of the RRM-like domain (residues 1-169; N) (Fig. 11E, and fig. S48) that was surprisingly devoid of sequence homology to the corresponding *C. thermophilum* Nup53^R2^ amphipathic *α*-helix motif that fits into a hydrophobic groove of the Nic96^SOL^ scaffold (*30*), suggesting a distinct binding mode between the human proteins (figs. S45 and S46). Through fragment truncation and five-alanine scanning mutagenesis, we identified a NUP53^R2^ region (residues 84-150) that formed a stable complex with NUP93^SOL^, and a shorter, more weakly binding 84-105 fragment, within which residues 86-100 were required for binding to NUP93^SOL^ (Fig. 12, A and B, and figs. S50 and S51). To elucidate the molecular details of binding between the divergent NUP53^R2^ motif and NUP93^SOL^, we determined crystal structures of *apo* NUP93^SOL^ and NUP93^SOL^•NUP53^R2^ at 2.0 Å and 3.4 Å resolution, respectively. The NUP53^R2^ sequence assignment was confirmed by identifying a peak in the anomalous difference Fourier map corresponding to a SeMet-labeled I94M substitution (Fig. 12C and fig. S52).

**Fig. 12.**
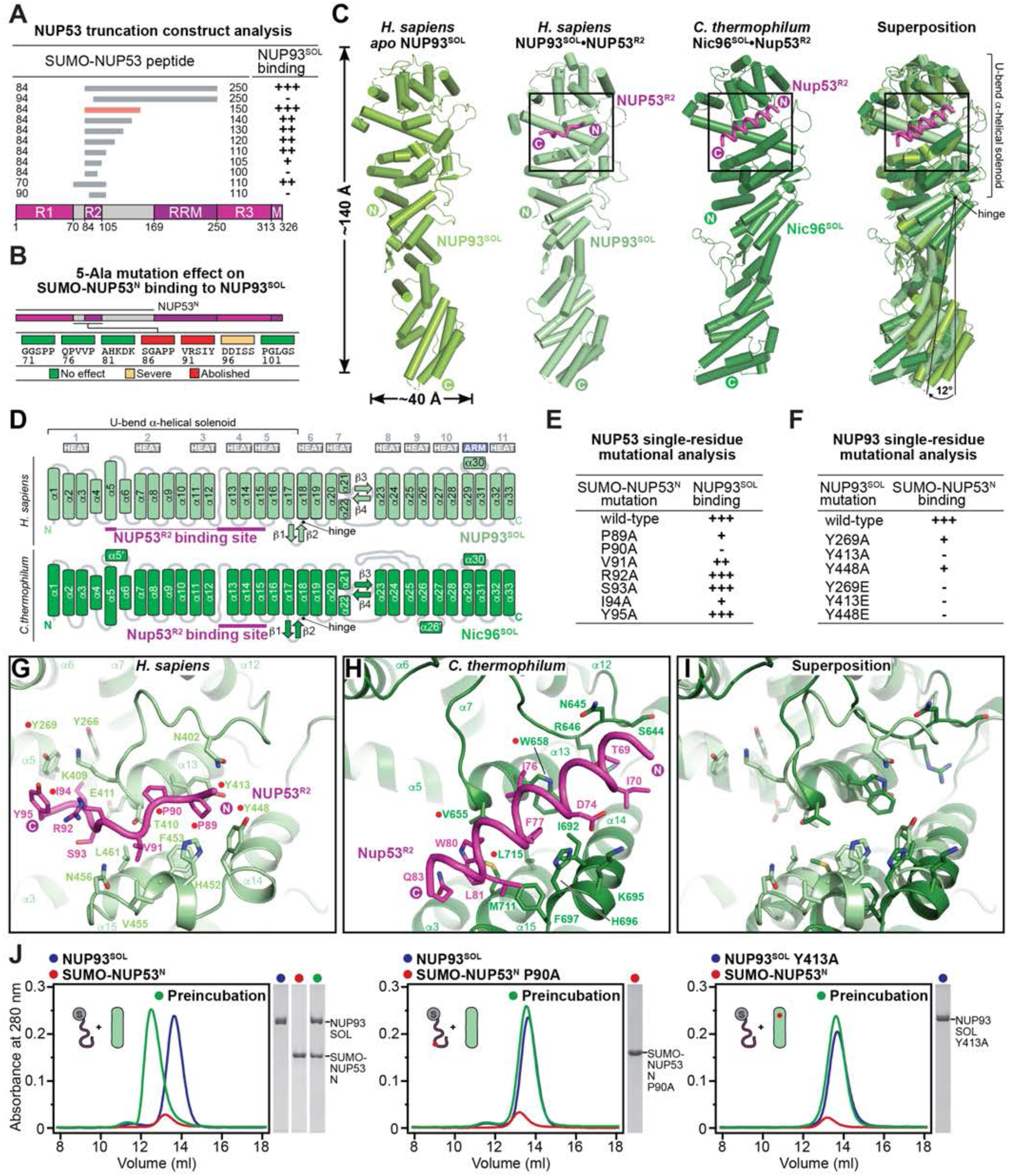
Structural and biochemical analyses of the NUP93-NUP53 interaction. (**A**) The minimal region sufficient for binding to NUP93^SOL^ was identified by progressively truncating NUP53 peptide constructs from either terminus. Peptide boundaries are shown as gray bars above the domain structure of NUP53. The NUP53^R2^ peptide is indicated by a red bar. Binding of SUMO-NUP53 peptides to NUP93^SOL^, assessed by SEC, is summarized according to the measured effect: (+++) no effect, (++) weak effect, (+) moderate effect, and (-) abolished binding. (**B**) The effect of each 5-Ala substitution on SUMO-NUP53^N^ binding to NUP93^SOL^, assessed by SEC, is indicated by the color-coded box above the respective mutated five-residue block of the NUP53 primary sequence. (**C**) Cartoon representation of *apo* NUP93^SOL^, NUP93^SOL^•NUP53^R2^, Nic96^SOL^•Nup53^R2^ (PDB ID 5HB3) (*30*), and their superposition based on the rigid N-terminal U-bend *α*-helical solenoid. A ∼12° displacement of the C-terminal *α*-helical solenoid, pivoted about the hinge loop, is observed between the *apo* NUP93^SOL^ and NUP93^SOL^•NUP53^R2^ structures. (**D**) Schematic of the human NUP93^SOL^ and *C. thermophilum* Nic96^SOL^ fold architectures. Rectangles and arrows represent *α*-helices and *β*-strands, respectively. (**E**, **F**) Summary of the effect of structure-driven mutations in (E) SUMO-NUP53^N^ and (D) NUP93^SOL^ on NUP93^SOL^•SUMO-NUP53^N^ complex formation, assayed by SEC and scored as in (A). (**G**-**I**) Closeup views of regions indicated with insets in (C), comparing the molecular details of the human NUP53^R2^ and *C. thermophilum* Nup53^R2^ binding sites. (**J**) Point mutations abolish SUMO-NUP53^N^ binding to NUP93^SOL^. SEC profiles are shown for individual nup variants and their preincubation, along with Coomassie-stained SDS-PAGE gel slices of peak fractions.

*C. thermophilum* and human Nic96^SOL^ orthologs display an equivalent secondary structure consisting of eleven HEAT repeats that form an *α*-helical solenoid, the first half of which is arranged in a U-bend fold (Fig. 12D) (*30, 57, 60*). Comparing the *apo* NUP93^SOL^ and NUP93^SOL^•NUP53^R2^ structures obtained from different crystal packing arrangements reveals significant structural plasticity of NUP93^SOL^, with a ∼12° rotation between the N-terminal U-bend and C-terminal halves of the *α*-helical solenoid, indicative of a central hinge (Fig. 12C and fig. S52, A to C).

As with other linker-scaffold interactions we characterized, only a core region (residues 88-95) of the biochemically mapped minimal NUP53^R2^ was resolved in the electron density, occupying a hydrophobic binding site, the shape and location of which are conserved between *C. thermophilum* Nic96^SOL^ and human NUP93^SOL^ (Fig. 12, C and G to I). In contrast to *C. thermophilum* Nup53^R2^, which adopts the form of an amphipathic *α*-helix, human NUP53^R2^ binds as a linear eight-residue motif to a hydrophobic surface groove encompassing *α*-helices *α*5 and *α*13-15 in the middle of the U-bend part of NUP93^SOL^, burying ∼1,100 Å^2^ of combined surface area (Fig. 12, C and G to I and fig. S52E). Inducing only a minor conformational change, NUP53^R2^ inserts the P89-P90 di-proline motif and I94 into hydrophobic pockets of NUP93^SOL^, organized around Y413 and Y448, and Y269, respectively (Fig. 12G and fig. S52, A to C). Systematic alanine substitution of the resolved NUP53^R2^ motif residues confirmed the key role of the P89-P90 di-proline and I94, consequently also illustrating the importance of the NUP93^SOL^ Y269, Y413, and Y448 residues that interface with them (Fig. 12, C and E to J, and fig. S53). Of note, the identified motif is invariant across metazoan NUP53 sequences (fig. S46).

Together, our data establish that despite distinct binding motifs and low sequence conservation, the linker-scaffold interactions are evolutionarily conserved, further highlighting their essential role and indicating that shape conservation of the scaffolds are key determinants of the NPC architecture.

### Architecture of the human NPC symmetric core

#### Quantitative docking of nup complexes in cryo-ET maps of the human NPC

We have previously demonstrated that nup ortholog crystal structures can be successfully used to interpret the density of a ∼23 Å cryo-ET map of the intact human NPC, yielding a near-atomic composite structure of the NPC symmetric core that included linker-scaffold crystal structures of the Nup170•Nup53^R3^•Nup145N^R3^, Nic96^SOL^•Nup53^R2^, and CNT•Nic96^R1^ complexes (*29, 30*). The newly available structures of full-length Nup192 and Nup188 as part of Nup192•Nic96^R2^•Nup145N^R1^•Nup53^R1^ and Nup188•Nic96^R2^•Nup145N^R2^ linker-scaffold complexes, as well as the human NUP93^SOL^•NUP53^R2^ complex, allowed us to build on our previous analysis with an improved ∼12 Å cryo-ET map of the intact human NPC (provided by Martin Beck) (*41*). As for the *S. cerevisiae* NPC described above, our quantitative docking approach consisted of statistically scoring the fit of resolution-matched densities simulated from crystal and single particle cryo-EM structures that were randomly placed and locally refined in cryo-ET maps of the human NPC. Structures of the CNC, Nup192•Nic96^R2^•Nup145N^R1^•Nup53^R1^, Nup188•Nic96^R2^•Nup145N^R2^, and NUP358^NTD^ (reported in the accompanying manuscript) (*61*) were readily placed in cryo-ET maps of the entire NPC or of the inner ring portion. Accounted density was then iteratively subtracted from the maps to reduce the subsequent search space for NUP93^SOL^•NUP53^R2^, Nup170•Nup53^R3^•Nup145N^R3^, CNT•Nic96^R1^, and NUP53^RRM^ (fig. S54).

As previously established, we identified 32 copies of the CNC, arranged in two concentric rings both on the nuclear and the cytoplasmic faces of the NPC (fig. S55A) (*26, 62*). The hetero-nonameric Y-shaped CNCs are generally rigid structures with an unstructured linker between the N-terminal *β*-propeller domain and the C-terminal *α*-helical solenoid of Nup133, which accommodates the difference in the circumference of narrower proximal and larger distal CNC rings, as we showed previously (fig. S55, B and C) (*26*).

Due to their overall shape similarity and the lack of full-length structures, our previous analysis had left ambiguity in the assignments of NUP205 and NUP188, which we are now able to resolve (figs. S56, S57, and S58). In the inner ring, 16 NUP188 and 16 NUP205 copies explain the question mark-shaped densities at peripheral and equatorial positions, respectively, in agreement with our data and previous assignments of Nup192 and Nup188 homologs in the *S. cerevisiae* NPC (*31, 32*). In the cytoplasmic outer ring, 16 copies of NUP205 are placed into eight distal and eight proximal question mark-shaped densities, but only 8 copies of NUP205 were identified in the nuclear outer ring, at the distal position. In all cases, the crystal conformation of Nup192, presenting a constricted gap between the Head subdomain and the Tower, resulted in a better fit than the cryo-EM conformation (figs. S4, S56, S57, S59, and S60). Outer ring positions assigned to NUP205 also appeared as high-scoring solutions in searches for NUP188, albeit with lower confidence than NUP205. The quantitative assignment of cryo-ET density to either NUP205 or NUP188 was supported by structural differences such as the Nup188 SH3- like domain, or the *α*-helical solenoid super-helical twist that determines the width of the question mark-shaped cryo-ET density (figs. S61 and S62). For reference, we also quantitatively docked the novel structures into the previously used ∼23 Å cryo-ET map (*33*), obtaining similar albeit less confident solutions, particularly in the novel assignment of the proximal outer ring NUP205 (figs. S59, S60, and S63). As with docking into the *S. cerevisiae* cryo-ET map, the novel full-length Nup188 structure was essential for differentiating NUP188 from NUP205 in the inner ring of the human cryo-ET maps.

Searches for NUP93^SOL^•NUP53^R2^ in ∼12 Å and ∼23 Å cryo-ET human NPC maps from which hereto assigned density had been subtracted, identified a total of 56 copies of NUP93: 32 copies previously assigned in the inner ring at equatorial and peripheral positions, 16 novel copies discovered at equivalent distal positions in the nuclear and cytoplasmic outer rings, and 8 novel copies manually placed into weak but matching rod-like densities at proximal positions in the cytoplasmic outer ring (figs. S64 and S65). This stoichiometry of placed NUP93^SOL^ molecules also matches the stoichiometry of NUP93^R2^ binding sites identified through placement of NUP205 and NUP188.

NUP155 locations have previously been determined by searching for Nup170•Nup53^R3^•Nup145N^R3^ crystal composite structures of different Nup170 conformations in an ∼23 Å cryo-ET map of the intact human NPC (*30, 33*). Quantitative docking into the improved ∼12 Å cryo-ET map with hereto assigned density subtracted allowed for assignment of the bridge and equatorial NUP155 positions (figs. S66 and S67A), but did not yield high confidence solutions for the peripheral NUP155 copies due to an imperfect match of the Nup170 conformation and poor local quality of the cryo-ET density, necessitating their assignment by visual inspection (fig. S67B).

For the numeral “4”-shaped CNT•Nic96^R1^, inspection of the top scoring hits from searches in an ∼12 Å cryo-ET density-subtracted map of the inner ring identified their equatorial and peripheral positions (fig. S68). As previously observed (*29, 30*), globular density proximal to the *α*/*β* insertion domain of NUP54 (*25, 63*), at the base of the first coiled-coil segment, indicated the location and flexible orientation of Ferredoxin-like NUP54 domains (fig. S69).

Finally, we searched for density corresponding to the NUP53^RRM^ homodimer (*30*) and NUP98^APD^ (*11*) in an ∼12 Å cryo-ET map of the intact NPC with hereto assigned density subtracted. Two globular discontinuous densities found between spokes of the inner ring and related by a C2-symmetry operator about the midplane were identified as the top two non-redundant solutions, though the orientation or identity of the docked NUP53^RRM^ homodimer and NUP98^APD^ domains could not be confidently assigned due to lack of distinct shape features and small size. The placement of 32 homodimerizing NUP53^RRM^ copies that, unlike the 16 copies of monomeric NUP98^APD^, would be spatially restrained by opposing tethers from adjacent spokes, is consistent with density that is discontinuous from the rest of the cryo-ET map withstanding sub-tomogram averaging (fig. S70).

We further confirmed the architecture of the cytoplasmic outer ring of the metazoan NPC by placing our composite structure into an ∼5.5 Å region of an anisotropic composite single particle cryo-EM map of the cytoplasmic face of the *X. laevis* NPC (*64*). Though the masking of the *X. laevis* single particle cryo-EM map excluded the region in which the proximal NUP93^SOL^ would be expected, we noted a close correspondence between the NUP93^SOL^ crystal structure and a secondary structure discernable in the map that had erroneously been referred to as the ‘bridge domain of the NUP358 complex’. Indeed, our docking analysis unambiguously confirmed the assignment of the rod-like *α*-helical solenoid bisecting the stalks of Y-shaped proximal and distal CNCs to the cytoplasmic outer ring distal copy of NUP93^SOL^. We also uncovered long tubular density perpendicular to the C-terminal ARM and HEAT repeats of both proximal and distal outer ring NUP205 copies, likely corresponding to NUP205-bound NUP93^R2^ (fig. S71).

After this docking procedure, the only regions of unexplained density left in the inner ring of the ∼12 Å cryo-ET human NPC map are two small globular densities symmetrically distributed on both nuclear and cytoplasmic sides of the inner ring, one atop NUP188 and proximal NUP93^SOL^, and the other adjacent to the nuclear envelope and the bridge and proximal NUP155. Asymmetric clusters of unexplained density that remained on the cytoplasmic and nuclear faces of the NPC are addressed in the accompanying manuscript (fig. S72) (*61*).

In conclusion, the novel structures and improved cryo-ET map of the intact human NPC have disambiguated the placement of NUP188 and NUP205 complexes in the inner ring, led to the discovery of a proximal NUP205 complex in the cytoplasmic outer ring, identified NUP93^SOL^ in the outer rings, placed NUP53^RRM^ homodimers in between inner ring spokes, and revealed a comprehensive map of linker-scaffold binding sites on scaffold nup surfaces of the symmetric core (fig. S73). The composite structure includes ∼400,000 ordered residues that explain ∼77 MDa of the human NPC mass.

#### Architecture of cytoplasmic and nuclear outer rings

In both the cytoplasmic and nuclear outer rings of the human NPC, 16 copies of the Y-shaped CNC are arranged in two concentric proximal and distal rings. At equivalent locations on both cytoplasmic and nuclear sides, eight copies of NUP205 are intercalated between the proximal NUP75 arms and distal NUP107 stalks of Y-shaped CNCs from adjacent spokes. Eight NUP93^SOL^ copies are inserted between the distal NUP107 and proximal NUP96 *α*-helical solenoids, bisecting the stalks of tandem-arranged Y-shaped CNCs of a single spoke (Fig. 13 and fig. S74). Stretched out, the ∼25 residue unstructured linker connecting the R2 and SOL regions of NUP93 could bridge the ∼95 Å gap between the distal NUP205-bound NUP93^R2^ and the distal NUP93^SOL^ of an adjacent spoke, thus crosslinking the outer ring spokes (Fig. 13, figs. S74 and S75). Compared to the Nup192 ortholog, NUP205 presents an additional ∼240 residues that elongate the C-terminal Tail region, suggesting that ∼95 Å is an upper estimate for the distance between distal NUP93^R2^ and NUP93^SOL^ from adjacent spokes.

**Fig. 13.**
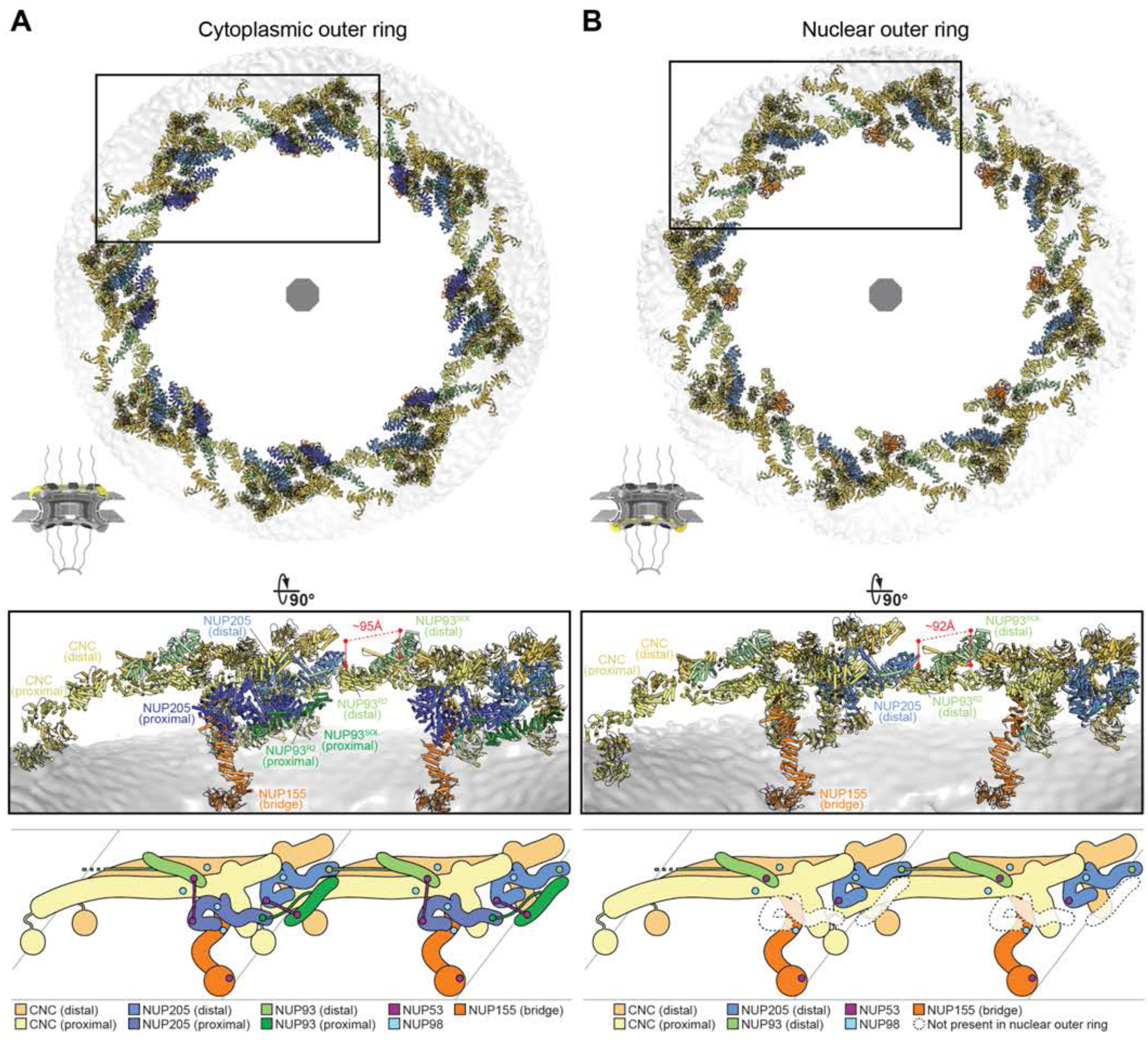
Architecture of the human NPC symmetric core outer rings. A composite structure of the human NPC was generated by quantitatively docking crystal and single particle cryo-EM structures into an ∼12 Å cryo-ET map of the intact human NPC (EMD-XXXX) (*41*). The nuclear envelope and docked structures are rendered in isosurface and cartoon representation, respectively. Views from above the (**A**) cytoplasmic, and (**B**) nuclear faces of the NPC. The insets indicate regions encompassing two spokes, shown in 90°-rotated and magnified views (*middle*). Distances between the distal NUP205-bound NUP93^R2^ and distal NUP93^SOL^ from adjacent spokes are indicated in red. Schematics illustrate the architecture of cytoplasmic and nuclear outer rings spokes (*bottom*). Linker binding sites on scaffold nup surfaces are indicated by colored circles. Proximal NUP205 and NUP93 were not found in the nuclear outer ring but positions equivalent to the cytoplasmic docking sites are indicated with dashed transparent shapes.

Unique to the cytoplasmic face, an additional eight copies of the proximal NUP205 are lodged between the NUP75 arm of the proximal CNC and the bridge NUP155 that connects the outer and inner ring (Fig. 13A and fig. S74). The proximal NUP205-bound NUP93^R2^ can only be linked with a proximal NUP93^SOL^ of the same spoke (Fig. 13A and figs. S74 and S75A). Furthermore, the arrangement of NUP53 binding sites on NUP93^SOL^ and NUP205 copies in the cytoplasmic outer rings is compatible with the NUP53-mediated linkage of the distal NUP205 with the proximal NUP93^SOL^ and, conversely, the proximal NUP205 with the distal NUP93^SOL^ (Fig. 13 and figs. S74 and S75, D and E). NUP53 could also mediate long-range links between the bridge NUP155 and the R1 and R2 binding sites on outer ring NUP205 and NUP93^SOL^, respectively (fig. S75, F, H and I).

On both nuclear and cytoplasmic sides, the NUP98^APD^-binding NUP96 sites present in the 16 CNC copies recruit 16 copies of NUP98 that can simultaneously satisfy the outer ring NUP205 binding sites thanks to the ∼115 residue linker between the APD and the R1-R2-R3 regions (Fig. 13, fig S74). The cytoplasmic proximal NUP205 and bridge NUP155 scaffolds could in principle be linked by NUP98, although NUP98^R3^ is likely outcompeted from its bridge NUP155 binding site by the asymmetric cytoplasmic filament nups GLE1•NUP42, as explained in the accompanying manuscript (Fig. 13A and fig. S75C) (*61*). To maximize nup copy parsimony while satisfying all available scaffold binding sites, the outer rings would recruit 16 copies of NUP53 and NUP98 on each side of the NPC.

#### Architecture of the inner ring linker-scaffold

The quantitative docking confirmed evolutionary conservation of the inner ring scaffold-linker architecture between *H. sapiens* and *S. cerevisiae* NPCs (Fig. 14). A NUP155•NUP53 linker-scaffold coats the nuclear envelope, anchored by membrane curvature-sensing Arf1GAP1 lipid packing sensor (ALPS) motifs and the C-terminal NUP53 amphipathic helix (*30, 65–67*), with a peripheral and equatorial copy on each side of a spoke midplane. The homodimerizing NUP53^RRM^ domains link spoke halves across the midplane (Fig. 14C and figs. S76 and S77, A and B). A second cross-midplane link between NUP53 and NUP93 connects NUP188•NUP93•CNT and NUP205•NUP93•CNT modules to the coat at equatorial and peripheral positions, respectively (Fig. 14, D to F, and fig. S77, C to J). Restrained by their length, NUP93 N-terminal linkers unambiguously connect NUP188 with the peripheral, and NUP205 with the equatorial inner ring NUP93^SOL^ and CNT copies (Fig. 14, D to F, and fig. S77, G to J).

**Fig. 14.**
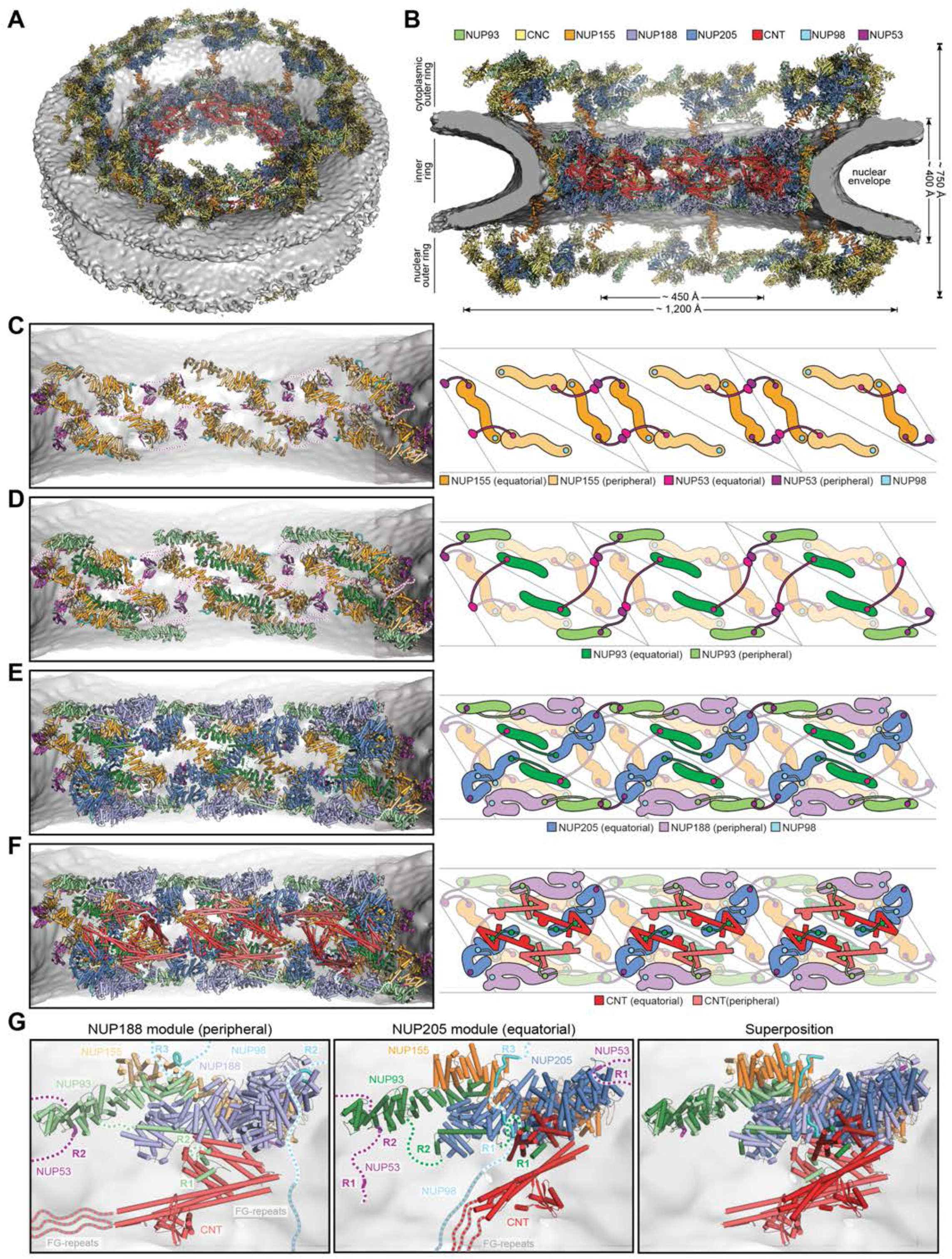
Architecture of the linker-scaffold of the human NPC inner ring. Composite structure generated by quantitatively docking crystal and single particle cryo-EM structures into an ∼12 Å cryo-ET map of the intact human NPC (EMD-XXXX) (*41*), viewed from (**A**) above the cytoplasmic face, and (**B**) a cross-section of the central transport channel. The nuclear envelope is rendered as a gray isosurface and the docked structures are represented as cartoons. (**C**-**F**) Starting from the nuclear envelope, successive layers of the inner ring in cartoon representation are added to reveal the architecture of three inner ring spokes of the human NPC. Corresponding schematics illustrate linker paths between binding sites on scaffold surfaces determined by considering maximum distance and linker copy parsimony. Linker binding sites on scaffold nup surfaces are indicated with colored circles. (C) NUP53^RRM^ domains dimerize between spokes to link cytoplasmic peripheral with nuclear equatorial, and conversely cytoplasmic equatorial with nuclear peripheral copies of NUP155. (D) The dimerized NUP53^RRM^ domains also link cytoplasmic peripheral with nuclear equatorial, and conversely cytoplasmic equatorial with nuclear peripheral copies of NUP93. (E) NUP205 and NUP188 bind to the equatorial and peripheral NUP93^R2^, respectively. NUP98 can simultaneously bind NUP205 and the equatorial NUP155. NUP53 can link NUP205 and the peripheral NUP93^SOL^ from adjacent spokes. (F) CNT is recruited by NUP93^R1^ and positioned by the binding of NUP93^R2^ to NUP188 and NUP205. (**G**) Closeup views of inner ring modules assembled on the NUP188 and NUP205 scaffold hubs, and their superposition. Dashed lines indicate unstructured protein regions, FG-repeats and linker nup segments connecting resolved linker regions bound to scaffold surfaces.

The equatorial position of NUP205 is further solidified by a NUP98-mediated linkage with the equatorial NUP155 and by a NUP53-mediated linkage with a peripheral NUP93 from an adjacent spoke (Fig. 14E and fig. S77, E and F). As in the *S. cerevisiae* NPC and the corresponding nup orthologs, the NUP205, NUP188, and peripheral NUP155 binding sites for the NUP98 R1, R2, and R3 regions, respectively, are too far apart to be linked by the same NUP98, suggesting that three copies are required to satisfy all binding sites on each side of a spoke midplane instead. On the cytoplasmic side of the inner ring, two of these three NUP98 copies could bind the cytoplasmic filament NUP88^NTD^ via their NUP98^APD^ domains (placed in the accompanying manuscript) (*61*), thus linking the inner ring with the asymmetric portion of the cytoplasmic face of the NPC.

The peripheral NUP170-NUP188-NUP93-CNT and equatorial NUP170-NUP205-NUP93-CNT scaffold modules that together form a protomer for a D8-symmetric inner ring can themselves be superposed, if NUP188 and NUP205 are considered as equivalent organizing hubs (Fig. 14G). Nevertheless, the peripheral NUP155 is not linked to the peripheral NUP188•NUP93•CNT complex from the same midplane of a spoke. Instead, within each spoke, linker-mediated complexes form between a peripheral NUP188•NUP98•NUP93•CNT•NUP53 and an equatorial NUP155 from across the midplane (fig. S77K). On the contrary, the equatorial NUP205•NUP98•NUP93•CNT•NUP53 complex is linked with a peripheral NUP155 from the same and an equatorial NUP155 from the opposite side of a spoke midplane, through NUP98- and NUP53-mediated links, respectively (fig. S77K). The inner ring scaffold architecture is remarkably interwoven by linker interactions.

To maximize nup copy parsimony while satisfying all available scaffold binding sites, the inner ring would recruit 32 and 48 copies of NUP53 and NUP98, respectively, symmetrically distributed in accordance with the eight- and two-fold rotational symmetries of the inner ring. Including copies recruited to outer rings, the human NPC would total 56 and 80 copies of NUP53 and NUP98, respectively. Though the rest of the symmetric core composite structure agrees with previous estimates of nup stoichiometry, the implied NUP98 and NUP53 copy number exceeds the empirical measurements (*68*). Whereas the discrepancy may be explained by available NUP98 and NUP53 binding sites not being fully occupied, the NUP98 and NUP53 linker nups are known to exchange comparatively rapidly at the NPC (*69, 70*), and might get depleted as nuclear envelopes are separated from cellular NUP98 and NUP53 reservoirs as part of the preparation for mass spectrometric analysis.

Docking the composite structure of the NPC into a ∼37 Å *in situ* cryo-ET map of the dilated human NPC revealed that the inner ring spokes move as relatively rigid bodies, accommodating the dilation by increasing the NUP53 linker-bridged gaps between spokes (fig. S78) (*40*). Spatial restraints in the dilated NPC confirm the intra-spoke topology of linkages established by N-terminal NUP93 and NUP98 linkers. Remarkably, the dilation of the NPC does not induce a significant increase in the gap between the adjacent spokes of the outer rings, consistent with the crosslinking purported by the linker between distal NUP93 R2 and SOL regions of adjacent spokes (fig. S78).

## Conclusions

The linker-scaffold is a fundamental architectural principle of the NPC structure. Despite the continuing improvement in resolution attained by cryo-ET reconstructions of intact NPCs over the last decade, the fine molecular detail of the linker-scaffold has remained out of reach. We employed comprehensive residue-level biochemical reconstitution and mapping, crystallographic and single particle cryo-EM structure determination, *in vivo* validation, and quantitative docking into improved and diversified cryo-ET NPC reconstructions to delineate the near-atomic structure and evolutionary conservation of the linker-scaffold interactions that underpin the integrity of the NPC.

The near-atomic resolution structures reported in this study complete the set of structures capturing all linker-scaffold interactions of the symmetric core. Docked into cryo-ET maps of the human and *S. cerevisiae* NPC, they reveal the topology and restrain distances between linker binding sites on scaffold surfaces, outlining how the multivalent linkers Nup145N/NUP98, Nup53/NUP53 and the N-terminal region of Nic96/NUP93 connect different parts of the NPC. Linkers mediate the formation of inner ring complexes that coalesce into relatively rigid spokes spanning from the nuclear envelope to the central transport channel. They also cross-stitch the inner ring scaffolds with connections across spoke midplanes and flexible links between spokes. In the outer rings, linker-scaffold interactions connect spokes and project ties towards the inner ring.

Our biochemical analysis of linker scaffold interactions involving Nup145N and Nup53 revealed another architectural principle, invisible to structural methods: linker-scaffold interactions are driven by structurally defined motifs that present canonical two-component binding dynamics, but are potentiated by disperse, structurally elusive interactions between flanking residues and promiscuous binding sites on the scaffold surfaces. These types of interactions, sometimes referred to as “fuzzy,” are found in systems involving intrinsically disordered proteins across biology, a prominent example being the ultrafast exchange of nucleocytoplasmic transport receptors on FG repeats (*9*).

The physical and chemical properties of linkers are advantageous for the assembly of giant complexes like the NPC. The unfolded property of linkers enables long-range interactions and confers flexibility that can accommodate large movements or shock-absorb deformations in the nuclear envelope.

The disperse nature of linker-scaffold interactions is conducive to the reuse of linker interactions in different chemical and steric environments of the NPC. The ensemble of binding modes provides robustness in the face of conformational changes of the NPC that might otherwise be incompatible with a unique mode of binding. Considering the avidity that results from multiple scaffold valences per linker, and the allovalency mediated by flanking residues (*71, 72*), it is unsurprising that our *in vivo* perturbation of interactions required extensive mutations to exacerbate deleterious phenotypes. For this reason and because of the lack of constraints imposed by a protein fold, new linker sequences are readily evolvable. Remarkably, the linker-scaffold network topology and modular binding site distribution on linkers is conserved from fungi to humans, despite striking divergence in linker sequences, most extremely exemplified by the complete divergence of the Nup53/NUP53 motif that binds to a conserved site on Nic96/NUP93.

The binding of linkers is amenable to exchange and regulation. The linearity of linkers imposes few obstacles to the deposition of post-translational modifications by the same machinery along the entire sequence to rapidly ablate the multiple binding valences. Indeed, patterns of Cdk1 and Nek-driven phosphorylation that lead to the choreographed depletion of both NUP98 and NUP53 from the NPC during mitotic nuclear envelope breakdown include the R1, R2 and R3 regions of both linkers (*73, 74*). Structural defects in the NPC resulting in aberrant nucleocytoplasmic transport may affect gene expression, mRNA maturation and export, leading to downstream tumorigenic processes. Therefore, the depletion of NUP98 from the NPC as a result of gene fusion mutations associated with various hematopoietic malignancies should be considered in the study of the disease-causing mechanisms triggered by these mutations (*75*).

The unexpected discovery of the presence and unique role of NUP93 in crosslinking the outer ring spokes of the human NPC, along with its organizing role in the inner ring as both scaffold and linker, exemplifies the reuse of linker-scaffold functional units at completely different locations of the NPC. Its ubiquity rationalizes the observation that rapid degron-induced depletion of NUP93 leads to the concomitant loss of both inner and outer rings from the nuclear pore, legitimizing NUP93 as a ‘lynchpin’ of the NPC (*76*). These findings further inform the mechanistic bases for pathologies like steroid-resistant nephrotic syndrome (SRNS), which is associated with mutations in NUP93 and NUP205, the most poignant of which is NUP205 F1995S, located in the NUP93^R2^ binding site of NUP205 and shown to abolish the NUP205-NUP93 interaction (*77*). Nonsense mutations that omit the NUP93^R2^ binding site in NUP188 are also associated with neurologic, ocular, and cardiac abnormalities (*78*).

Lastly, the docking of our NPC composite model into an ∼37 Å *in situ* cryo-ET map of the dilated human NPC demonstrates the magnitude of the movements that must be withstood by the linkers that connect the inner ring spokes (*40*). Moreover, the dilated inner ring reveals unencumbered lateral channels between its spokes and the nuclear envelope, providing an opening for the transport of large cytosolic domains of inner nuclear membrane integral membrane proteins bound to nucleocytoplasmic transport receptors (*79–82*). Finally, the capacity of the NPC to exist in dilated and constricted states dependent on tension imparted by the surrounding nuclear envelope portrays the NPC as the cell’s largest mechanosensitive channel, with implications in cellular energy state sensing and transport of transcriptional regulators in response to mechanical stress on the cell. Future studies will have to establish the causal links between the transmission of tension from the nuclear envelope to the NPC, the dilation of its inner ring, and mechanisms by which dilation and constriction may modulate nucleocytoplasmic transport.

Our results shed light on the elusive linker-scaffold molecular interactions that maintain the integrity of the NPC, providing a comprehensive characterization of a final major aspect of the NPC symmetric core. Building on this roadmap, future studies can address NPC assembly and disassembly mechanisms, the emergence of NPC polarity and attachment of the asymmetric cytoplasmic filaments and nuclear basket to the symmetric core, mechanisms of NPC-associated diseases, and mechanisms of NPC dilation and constriction along with their implications in nucleocytoplasmic transport.

## Supporting information

Hoelz_NPC_LinkerScaffold

## Acknowledgements

We thank Alina Patke for critical reading and editing of the manuscript and insightful discussions; Martin Beck for sharing an ∼12 Å cryo-ET reconstruction of the intact human HeLa cell NPC prior to publication; Ed Hurt, Susan Wente, Beatriz Fountura, David Baltimore and the Kazusa DNA Research Institute for providing material, and Felice Liang, Alex Lyons, Aaron Tang, Jimmy Thai, Giovani Pinton Tomaleri, and Lucas Schaus for experimental support. We are grateful to Dominika Borek, Shyam Saladi, and members of the Hoelz lab for discussion and suggestions and Valerie Altounian for the preparation of NPC schematics. We acknowledge Jens Kaiser, the scientific staff of the SSRL beamline 12-2, and the National Institute of General Medical Sciences and National Cancer Institute Structural Biology Facility (GM/CA) at the Advanced Photon Source (APS) for their support with x-ray diffraction measurements. We acknowledge the Gordon and Betty Moore Foundation, the Beckman Institute, and the Sanofi-Aventis Bioengineering Research Program for their support of the Molecular Observatory at the California Institute of Technology (Caltech). The operations at the SSRL are supported by the U.S. Department of Energy (DOE) and the National Institutes of Health (NIH). We acknowledge Songye Chen and Andrey Malyutin of the Beckman Institute Resource Center for Transmission Electron Microscopy at the Caltech for their support with cryo-electron microscopy (cryo-EM) imaging. We also acknowledge Janette Myers and the scientific staff of the Pacific Northwest CryoEM Center (PNCC) at the Oregon Health and Science University (OHSU) and the Environmental Molecular Sciences Laboratory (EMSL) for their support with cryo-EM imaging. A portion of this research was supported by NIH grant U24GM129547 and performed at the PNCC at OHSU and accessed through EMSL (grid.436923.9), a DOE Office of Science User Facility sponsored by the Office of Biological and Environmental Research. We acknowledge the Center for Molecular Medicine at Caltech for access to an isothermal titration calorimetry instrument. The establishment of the Center for Molecular Medicine was made possible by a generous grant from the Gordon and Betty Moore Foundation. We acknowledge the Cold Spring Harbor Laboratory (CSHL) cryo-EM course, the CSHL X-ray Methods in Structural Biology course, and the Michigan Life Science Institute cryo-EM workshop and their instructors Michael Cianfrocco, William Furey, Gary Gilliland, Justin Kollman, Gabe Lander, Melanie Ohi, James Pflugrath, Alexander McPherson, and Matthijn Vos, along with and all the course staff and lecturers for valuable expert training.

S.P. was supported by a pre-doctoral fellowship from the Boehringer Ingelheim Fonds and by an Amgen Graduate Fellowship through the Caltech-Amgen Research Collaboration. A.H. was supported by a Camille-Dreyfus Teacher Scholar Award, NIH grants R01-GM117360 and R01-GM111461, and is a Faculty Scholar with the Howard Hughes Medical Institute and an Investigator with the Heritage Medical Research Institute. The coordinates and structure factors of crystal structures have been deposited in the PDB with accession numbers 7MVT (Nup192 residues 185-1756; Nic96 residues 187-301), 7MVW (Nup188^NTD^), 7MVX (Nup188•Nic96^R2^), 7MW0 (NUP93^SOL^), and 7MW0 (NUP93^SOL^•NUP53^R2^). The coordinates of single particle cryo-EM structures have been deposited in the PDB with accession numbers 7MVU (Nup192•Nic96R2), 7MVV (Nup192•Nic96^R2^•Nup145N^R1^•Nup53^R1^), 7MVY (Nup188•Nic96^R2^), and 7MVZ (Nup188•Nic96^R2^•Nup145N^R2^). The maps of single particle cryo-EM structures have been deposited in the EMDB with accession numbers EMD-24056 (Nup192•Nic96^R2^), EMD-24057 (Nup192•Nic96^R2^•Nup145N^R1^•Nup53^R1^), EMD-24058 (Nup188•Nic96^R2^), EMD-24059 (Nup188•Nic96^R2^•Nup145N^R2^). PyMol and Chimera sessions containing the composite structures of the dilated *S. cerevisiae*, constricted human and dilated human NPC symmetric core can be obtained from our webpage (http://ahweb.caltech.edu) and are deposited in the PDB with the respective accession numbers XXXX, XXXX, XXXX. All quantitative docking data and code were deposited on CaltechDATA (XXXX, XXXX). The authors declare no financial conflicts of interest. A.H. is a co-founder of AMPlify Biosciences, Inc., a biotech company working in the hearing field.

## Supplementary Materials

Materials and Methods

Figs. S1-S78

Tables S1-S14 References (83–130)

## REFERENCES AND NOTES

1. A. Hoelz, E. W. Debler, G. Blobel, The structure of the nuclear pore complex. Annu. Rev. Biochem. 80, 613–643 (2011).

2. D. H. Lin, A. Hoelz, The Structure of the Nuclear Pore Complex (An Update). Annu. Rev. Biochem. 88, 725–783 (2019).

3. B. Hampoelz, A. Andres-Pons, P. Kastritis, M. Beck, Structure and Assembly of the Nuclear Pore Complex. Annu. Rev. Biophys. 48, 515–536 (2019).

4. K. E. Knockenhauer, T. U. Schwartz, The Nuclear Pore Complex as a Flexible and Dynamic Gate. Cell 164, 1162–1171 (2016).

5. A. Kohler, E. Hurt, Gene regulation by nucleoporins and links to cancer. Mol. Cell 38, 6–15 (2010).

6. M. L. Yarbrough, M. A. Mata, R. Sakthivel, B. M. Fontoura, Viral subversion of nucleocytoplasmic trafficking. Traffic 15, 127–140 (2014).

7. Y. M. Chook, K. E. Suel, Nuclear import by karyopherin-betas: recognition and inhibition. Biochim. Biophys. Acta 1813, 1593–1606 (2011).

8. A. Cook, F. Bono, M. Jinek, E. Conti, Structural biology of nucleocytoplasmic transport. Annu. Rev. Biochem. 76, 647–671 (2007).

9. S. Milles et al., Plasticity of an ultrafast interaction between nucleoporins and nuclear transport receptors. Cell 163, 734–745 (2015).

10. R. A. Pumroy, G. Cingolani, Diversification of importin-alpha isoforms in cellular trafficking and disease states. Biochem. J. 466, 13–28 (2015).

11. A. E. Hodel et al., The three-dimensional structure of the autoproteolytic, nuclear pore-targeting domain of the human nucleoporin Nup98. Mol. Cell 10, 347–358 (2002).

12. I. C. Berke, T. Boehmer, G. Blobel, T. U. Schwartz, Structural and functional analysis of Nup133 domains reveals modular building blocks of the nuclear pore complex. J. Cell Biol. 167, 591–597 (2004).

13. K. C. Hsia, P. Stavropoulos, G. Blobel, A. Hoelz, Architecture of a coat for the nuclear pore membrane. Cell 131, 1313–1326 (2007).

14. T. Boehmer, S. Jeudy, I. C. Berke, T. U. Schwartz, Structural and functional studies of Nup107/Nup133 interaction and its implications for the architecture of the nuclear pore complex. Mol. Cell 30, 721–731 (2008).

15. S. G. Brohawn, N. C. Leksa, E. D. Spear, K. R. Rajashankar, T. U. Schwartz, Structural evidence for common ancestry of the nuclear pore complex and vesicle coats. Science 322, 1369–1373 (2008).

16. E. W. Debler et al., A fence-like coat for the nuclear pore membrane. Mol. Cell 32, 815–826 (2008).

17. S. G. Brohawn, T. U. Schwartz, Molecular architecture of the Nup84-Nup145C-Sec13 edge element in the nuclear pore complex lattice. Nat. Struct. Mol. Biol. 16, 1173–1177 (2009).

18. N. C. Leksa, S. G. Brohawn, T. U. Schwartz, The structure of the scaffold nucleoporin Nup120 reveals a new and unexpected domain architecture. Structure 17, 1082–1091 (2009).

19. V. Nagy et al., Structure of a trimeric nucleoporin complex reveals alternate oligomerization states. Proc. Natl. Acad. Sci. U.S.A. 106, 17693–17698 (2009).

20. H. S. Seo et al., Structural and functional analysis of Nup120 suggests ring formation of the Nup84 complex. Proc. Natl. Acad. Sci. U.S.A. 106, 14281–14286 (2009).

21. J. R. R. Whittle, T. U. Schwartz, Architectural nucleoporins Nup157/170 and Nup133 are structurally related and descend from a second ancestral element. J. Biol. Chem. 284, 28442–28452 (2009).

22. S. Bilokapic, T. U. Schwartz, Molecular basis for Nup37 and ELY5/ELYS recruitment to the nuclear pore complex. Proc. Natl. Acad. Sci. U.S.A. 109, 15241–15246 (2012).

23. S. Bilokapic, T. U. Schwartz, Structural and functional studies of the 252 kDa nucleoporin ELYS reveal distinct roles for its three tethered domains. Structure 21, 572–580 (2013).

24. K. Kelley, K. E. Knockenhauer, G. Kabachinski, T. U. Schwartz, Atomic structure of the Y complex of the nuclear pore. Nat. Struct. Mol. Biol. 22, 425–431 (2015).

25. T. Stuwe et al., Architecture of the fungal nuclear pore inner ring complex. Science 350, 56–64 (2015).

26. T. Stuwe, et al., Nuclear pores. Architecture of the nuclear pore complex coat. Science 347, 1148–1152 (2015).

27. C. Xu et al., Crystal structure of human nuclear pore complex component NUP43. FEBS Lett. 589, 3247–3253 (2015).

28. J. Fischer, R. Teimer, S. Amlacher, R. Kunze, E. Hurt, Linker Nups connect the nuclear pore complex inner ring with the outer ring and transport channel. Nat. Struct. Mol. Biol. 22, 774–781 (2015).

29. J. Kosinski et al., Molecular architecture of the inner ring scaffold of the human nuclear pore complex. Science 352, 363–365 (2016).

30. D. H. Lin et al., Architecture of the symmetric core of the nuclear pore. Science 352, aaf1015 (2016).

31. M. Allegretti et al., In-cell architecture of the nuclear pore and snapshots of its turnover. Nature 586, 796–800 (2020).

32. S. J. Kim et al., Integrative structure and functional anatomy of a nuclear pore complex. Nature 555, 475–482 (2018).

33. A. von Appen et al., In situ structural analysis of the human nuclear pore complex. Nature 526, 140–143 (2015).

34. S. A. Nordeen, D. L. Turman, T. U. Schwartz, Yeast Nup84-Nup133 complex structure details flexibility and reveals conservation of the membrane anchoring ALPS motif. Nat. Commun. 11, 6060 (2020).

35. S. Amlacher et al., Insight into structure and assembly of the nuclear pore complex by utilizing the genome of a eukaryotic thermophile. Cell 146, 277–289 (2011).

36. T. Stuwe, D. H. Lin, L. N. Collins, E. Hurt, A. Hoelz, Evidence for an evolutionary relationship between the large adaptor nucleoporin Nup192 and karyopherins. Proc. Natl. Acad. Sci. U.S.A. 111, 2530–2535 (2014).

37. S. Mosalaganti et al., In situ architecture of the algal nuclear pore complex. Nat. Commun. 9, 2361 (2018).

38. C. E. Zimmerli et al., Nuclear pores constrict upon energy depletion. bioRxiv, 2020.2007.2030.228585 (2020).

39. A. P. Schuller et al., The cellular environment shapes the nuclear pore complex architecture. Nature, (2021).

40. V. Zila et al., Cone-shaped HIV-1 capsids are transported through intact nuclear pores. Cell 184, 1032–1046 e1018 (2021).

41. M. Beck, Cryo ET of the human nuclear pore complex. Accompanying Manuscript, (2021).

42. P. Sampathkumar et al., Structure, dynamics, evolution, and function of a major scaffold component in the nuclear pore complex. Structure 21, 560–571 (2013).

43. K. R. Andersen et al., Scaffold nucleoporins Nup188 and Nup192 share structural and functional properties with nuclear transport receptors. Elife 2, e00745 (2013).

44. S. M. Bailer et al., Nup116p and nup100p are interchangeable through a conserved motif which constitutes a docking site for the mRNA transport factor gle2p. EMBO J. 17, 1107–1119 (1998).

45. K. Yoshida, H. S. Seo, E. W. Debler, G. Blobel, A. Hoelz, Structural and functional analysis of an essential nucleoporin heterotrimer on the cytoplasmic face of the nuclear pore complex. Proc. Natl. Acad. Sci. U.S.A. 108, 16571–16576 (2011).

46. A. K. Ho et al., Assembly and preferential localization of Nup116p on the cytoplasmic face of the nuclear pore complex by interaction with Nup82p. Mol. Cell. Biol. 20, 5736–5748 (2000).

47. M. T. Teixeira et al., Two functionally distinct domains generated by in vivo cleavage of Nup145p: a novel biogenesis pathway for nucleoporins. EMBO J. 16, 5086–5097 (1997).

48. S. R. Wente, G. Blobel, NUP145 encodes a novel yeast glycine-leucine-phenylalanine-glycine (GLFG) nucleoporin required for nuclear envelope structure. J. Cell Biol. 125, 955–969 (1994).

49. K. J. Ryan, S. R. Wente, Isolation and characterization of new Saccharomyces cerevisiae mutants perturbed in nuclear pore complex assembly. BMC Genet. 3, 17 (2002).

50. S. S. Patel, B. J. Belmont, J. M. Sante, M. F. Rexach, Natively unfolded nucleoporins gate protein diffusion across the nuclear pore complex. Cell 129, 83–96 (2007).

51. M. Miao, K. J. Ryan, S. R. Wente, The integral membrane protein Pom34p functionally links nucleoporin subcomplexes. Genetics 172, 1441–1457 (2006).

52. U. Nehrbass, M. P. Rout, S. Maguire, G. Blobel, R. W. Wozniak, The yeast nucleoporin Nup188p interacts genetically and physically with the core structures of the nuclear pore complex. J. Cell Biol. 133, 1153–1162 (1996).

53. A. de Bruyn Kops, C. Guthrie, Identification of the Novel Nup188-brr7 Allele in a Screen for Cold-Sensitive mRNA Export Mutants in Saccharomyces cerevisiae. G3 (Bethesda) 8, 2991–3003 (2018).

54. S. Rajoo, P. Vallotton, E. Onischenko, K. Weis, Stoichiometry and compositional plasticity of the yeast nuclear pore complex revealed by quantitative fluorescence microscopy. Proc. Natl. Acad. Sci. U.S.A. 115, E3969–E3977 (2018).

55. M. Marelli, J. D. Aitchison, R. W. Wozniak, Specific binding of the karyopherin Kap121p to a subunit of the nuclear pore complex containing Nup53p, Nup59p, and Nup170p. J. Cell Biol. 143, 1813–1830 (1998).

56. N. P. Allen, L. Huang, A. Burlingame, M. Rexach, Proteomic analysis of nucleoporin interacting proteins. J. Biol. Chem. 276, 29268–29274 (2001).

57. N. Schrader et al., Structural basis of the nic96 subcomplex organization in the nuclear pore channel. Mol. Cell 29, 46–55 (2008).

58. E. Onischenko et al., Natively Unfolded FG Repeats Stabilize the Structure of the Nuclear Pore Complex. Cell 171, 904–917 e919 (2017).

59. M. P. Rout et al., The yeast nuclear pore complex: composition, architecture, and transport mechanism. J. Cell Biol. 148, 635–651 (2000).

60. S. Jeudy, T. U. Schwartz, Crystal structure of nucleoporin Nic96 reveals a novel, intricate helical domain architecture. J. Biol. Chem. 282, 34904–34912 (2007).

61. C. J. Bley et al., Architechture of the cytoplasmic face of the nuclear pore. Accompanying Manuscript, (2021).

62. K. H. Bui et al., Integrated structural analysis of the human nuclear pore complex scaffold. Cell 155, 1233–1243 (2013).

63. H. Chug, S. Trakhanov, B. B. Hulsmann, T. Pleiner, D. Gorlich, Crystal structure of the metazoan Nup62*Nup58*Nup54 nucleoporin complex. Science 350, 106–110 (2015).

64. G. Huang et al., Structure of the cytoplasmic ring of the Xenopus laevis nuclear pore complex by cryo-electron microscopy single particle analysis. Cell Res. 30, 520–531 (2020).

65. G. Drin et al., A general amphipathic alpha-helical motif for sensing membrane curvature. Nat. Struct. Mol. Biol. 14, 138–146 (2007).

66. N. Eisenhardt, J. Redolfi, W. Antonin, Interaction of Nup53 with Ndc1 and Nup155 is required for nuclear pore complex assembly. J. Cell Sci. 127, 908–921 (2014).

67. E. Onischenko, L. H. Stanton, A. S. Madrid, T. Kieselbach, K. Weis, Role of the Ndc1 interaction network in yeast nuclear pore complex assembly and maintenance. J. Cell Biol. 185, 475–491 (2009).

68. A. Ori et al., Cell type-specific nuclear pores: a case in point for context-dependent stoichiometry of molecular machines. Mol. Syst. Biol. 9, 648 (2013).

69. E. R. Griffis, N. Altan, J. Lippincott-Schwartz, M. A. Powers, Nup98 is a mobile nucleoporin with transcription-dependent dynamics. Mol. Biol. Cell 13, 1282–1297 (2002).

70. G. Rabut, V. Doye, J. Ellenberg, Mapping the dynamic organization of the nuclear pore complex inside single living cells. Nat. Cell Biol. 6, 1114–1121 (2004).

71. A. Levchenko, Allovalency: a case of molecular entanglement. Curr. Biol. 13, R876–878 (2003).

72. J. G. Olsen, K. Teilum, B. B. Kragelund, Behaviour of intrinsically disordered proteins in protein-protein complexes with an emphasis on fuzziness. Cell. Mol. Life Sci. 74, 3175–3183 (2017).

73. E. Laurell et al., Phosphorylation of Nup98 by multiple kinases is crucial for NPC disassembly during mitotic entry. Cell 144, 539–550 (2011).

74. M. I. Linder et al., Mitotic Disassembly of Nuclear Pore Complexes Involves CDK1- and PLK1- Mediated Phosphorylation of Key Interconnecting Nucleoporins. Dev. Cell 43, 141–156 e147 (2017).

75. S. M. Gough, C. I. Slape, P. D. Aplan, NUP98 gene fusions and hematopoietic malignancies: common themes and new biologic insights. Blood 118, 6247–6257 (2011).

76. S. G. Regmi et al., The Nuclear Pore Complex consists of two independent scaffolds. bioRxiv, 2020.2011.2013.381947 (2020).

77. D. A. Braun et al., Mutations in nuclear pore genes NUP93, NUP205 and XPO5 cause steroid-resistant nephrotic syndrome. Nat. Genet. 48, 457–465 (2016).

78. A. M. Muir et al., Bi-allelic Loss-of-Function Variants in NUP188 Cause a Recognizable Syndrome Characterized by Neurologic, Ocular, and Cardiac Abnormalities. Am. J. Hum. Genet. 106, 623–631 (2020).

79. M. C. King, C. P. Lusk, G. Blobel, Karyopherin-mediated import of integral inner nuclear membrane proteins. Nature 442, 1003–1007 (2006).

80. A. C. Meinema et al., Long unfolded linkers facilitate membrane protein import through the nuclear pore complex. Science 333, 90–93 (2011).

81. R. K. Lokareddy et al., Distinctive Properties of the Nuclear Localization Signals of Inner Nuclear Membrane Proteins Heh1 and Heh2. Structure 23, 1305–1316 (2015).

82. P. Popken, A. Ghavami, P. R. Onck, B. Poolman, L. M. Veenhoff, Size-dependent leak of soluble and membrane proteins through the yeast nuclear pore complex. Mol. Biol. Cell 26, 1386–1394 (2015).

83. B. M. Fontoura, G. Blobel, M. J. Matunis, A conserved biogenesis pathway for nucleoporins: proteolytic processing of a 186-kilodalton precursor generates Nup98 and the novel nucleoporin, Nup96. J. Cell Biol. 144, 1097–1112 (1999).

84. F. Kendirgi, D. J. Rexer, A. R. Alcazar-Roman, H. M. Onishko, S. R. Wente, Interaction between the shuttling mRNA export factor Gle1 and the nucleoporin hCG1: a conserved mechanism in the export of Hsp70 mRNA. Mol. Biol. Cell 16, 4304–4315 (2005).

85. A. Hoelz, A. C. Nairn, J. Kuriyan, Crystal structure of a tetradecameric assembly of the association domain of Ca2+/calmodulin-dependent kinase II. Mol. Cell 11, 1241–1251 (2003).

86. E. Mossessova, C. D. Lima, Ulp1-SUMO crystal structure and genetic analysis reveal conserved interactions and a regulatory element essential for cell growth in yeast. Mol. Cell 5, 865–876 (2000).

87. C. Romier et al., Co-expression of protein complexes in prokaryotic and eukaryotic hosts: experimental procedures, database tracking and case studies. Acta Crystallogr. D Biol. Crystallogr. 62, 1232–1242 (2006).

88. S. Doublie, Preparation of selenomethionyl proteins for phase determination. Methods Enzymol. 276, 523–530 (1997).

89. P. J. Wyatt, Multiangle light scattering: The basic tool for macromolecular characterization. Instrum. Sci. Technol. 25, 1–18 (1997).

90. W. Kabsch, Xds. Acta Crystallogr. D Biol. Crystallogr. 66, 125–132 (2010).

91. P. Emsley, B. Lohkamp, W. G. Scott, K. Cowtan, Features and development of Coot. Acta Crystallogr. D Biol. Crystallogr. 66, 486–501 (2010).

92. D. Liebschner et al., Macromolecular structure determination using X-rays, neutrons and electrons: recent developments in Phenix. Acta Crystallogr. D Struct. Biol. 75, 861–877 (2019).

93. C. J. Williams et al., MolProbity: More and better reference data for improved all-atom structure validation. Protein Sci. 27, 293–315 (2018).

94. C. Vonrhein, E. Blanc, P. Roversi, G. Bricogne, Automated structure solution with autoSHARP. Methods Mol. Biol. 364, 215–230 (2007).

95. G. Winter et al., DIALS: implementation and evaluation of a new integration package. Acta Crystallogr. D Struct. Biol. 74, 85–97 (2018).

96. M. D. Winn et al., Overview of the CCP4 suite and current developments. Acta Crystallogr. D Biol. Crystallogr. 67, 235–242 (2011).

97. J. Painter, E. A. Merritt, Optimal description of a protein structure in terms of multiple groups undergoing TLS motion. Acta Crystallogr. D Biol. Crystallogr. 62, 439–450 (2006).

98. A. J. McCoy, et al., Phaser crystallographic software. J. Appl. Crystallogr. 40, 658–674 (2007).

99. S. P. Tiwari et al., WEBnm@ v2.0: Web server and services for comparing protein flexibility. BMC Bioinformatics 15, 427 (2014).

100. T. C. Terwilliger et al., Iterative model building, structure refinement and density modification with the PHENIX AutoBuild wizard. Acta Crystallogr. D Biol. Crystallogr. 64, 61–69 (2008).

101. J. W. Wang et al., SAD phasing by combination of direct methods with the SOLVE/RESOLVE procedure. Acta Crystallogr. D Biol. Crystallogr. 60, 1244–1253 (2004).

102. T. Terwilliger, SOLVE and RESOLVE: automated structure solution, density modification and model building. J. Synchrotron Radiat. 11, 49–52 (2004).

103. D. N. Mastronarde, Automated electron microscope tomography using robust prediction of specimen movements. J. Struct. Biol. 152, 36–51 (2005).

104. R. F. Thompson, M. G. Iadanza, E. L. Hesketh, S. Rawson, N. A. Ranson, Collection, pre-processing and on-the-fly analysis of data for high-resolution, single-particle cryo-electron microscopy. Nat. Protoc. 14, 100–118 (2019).

105. S. Q. Zheng et al., MotionCor2: anisotropic correction of beam-induced motion for improved cryo-electron microscopy. Nat. Methods 14, 331–332 (2017).

106. A. Punjani, J. L. Rubinstein, D. J. Fleet, M. A. Brubaker, cryoSPARC: algorithms for rapid unsupervised cryo-EM structure determination. Nat. Methods 14, 290–296 (2017).

107. J. Zivanov et al., New tools for automated high-resolution cryo-EM structure determination in RELION-3. Elife 7, (2018).

108. Y. Z. Tan et al., Addressing preferred specimen orientation in single-particle cryo-EM through tilting. Nat. Methods 14, 793–796 (2017).

109. E. F. Pettersen et al., UCSF Chimera--a visualization system for exploratory research and analysis. J. Comput. Chem. 25, 1605–1612 (2004).

110. D. Asarnow,,, E. Palovcak, Y. Cheng, asarnow/pyem: UCSF pyem v0.5., (2019).

111. P. V. Afonine et al., Real-space refinement in PHENIX for cryo-EM and crystallography. Acta Crystallogr. D Struct. Biol. 74, 531–544 (2018).

112. B. A. Barad et al., EMRinger: side chain-directed model and map validation for 3D cryo-electron microscopy. Nat. Methods 12, 943–946 (2015).

113. T. C. Terwilliger, O. V. Sobolev, P. V. Afonine, P. D. Adams, Automated map sharpening by maximization of detail and connectivity. Acta Crystallogr. D Struct. Biol. 74, 545–559 (2018).

114. A. Punjani, H. Zhang, D. J. Fleet, Non-uniform refinement: adaptive regularization improves single-particle cryo-EM reconstruction. Nat. Methods 17, 1214–1221 (2020).

115. A. J. Jakobi, M. Wilmanns, C. Sachse, Model-based local density sharpening of cryo-EM maps. Elife 6, (2017).

116. J. Pei, B. H. Kim, N. V. Grishin, PROMALS3D: a tool for multiple protein sequence and structure alignments. Nucleic Acids Res. 36, 2295–2300 (2008).

117. K. Katoh, D. M. Standley, MAFFT multiple sequence alignment software version 7: improvements in performance and usability. Mol. Biol. Evol. 30, 772–780 (2013).

118. G. J. Barton, ALSCRIPT: a tool to format multiple sequence alignments. Protein Eng. 6, 37–40 (1993).

119. R. S. Sikorski, P. Hieter, A system of shuttle vectors and yeast host strains designed for efficient manipulation of DNA in Saccharomyces cerevisiae. Genetics 122, 19–27 (1989).

120. C. Janke et al., A versatile toolbox for PCR-based tagging of yeast genes: new fluorescent proteins, more markers and promoter substitution cassettes. Yeast 21, 947–962 (2004).

121. M. S. Longtine et al., Additional modules for versatile and economical PCR-based gene deletion and modification in Saccharomyces cerevisiae. Yeast 14, 953–961 (1998).

122. J. S. Wadia, R. V. Stan, S. F. Dowdy, Transducible TAT-HA fusogenic peptide enhances escape of TAT-fusion proteins after lipid raft macropinocytosis. Nat. Med. 10, 310–315 (2004).

123. M. P. Yaffe, G. Schatz, Two nuclear mutations that block mitochondrial protein import in yeast. Proc. Natl. Acad. Sci. U.S.A. 81, 4819–4823 (1984).

124. C. Gwizdek et al., Ubiquitin-associated domain of Mex67 synchronizes recruitment of the mRNA export machinery with transcription. Proc. Natl. Acad. Sci. U.S.A. 103, 16376–16381 (2006).

125. 125. M. L. Waskom, seaborn:statistical data visualization. J. Open Source Sotfw. 6, (2021).

126. C. P. Lusk, T. Makhnevych, M. Marelli, J. D. Aitchison, R. W. Wozniak, Karyopherins in nuclear pore biogenesis: a role for Kap121p in the assembly of Nup53p into nuclear pore complexes. J. Cell Biol. 159, 267–278 (2002).

127. D. H. Lin et al., Structural and functional analysis of mRNA export regulation by the nuclear pore complex. Nat. Commun. 9, 2319 (2018).

128. J. Schindelin et al., Fiji: an open-source platform for biological-image analysis. Nat. Methods 9, 676–682 (2012).

129. F. Jeanmougin, J. D. Thompson, M. Gouy, D. G. Higgins, T. J. Gibson, Multiple sequence alignment with Clustal X. Trends Biochem. Sci. 23, 403–405 (1998).

130. T. W. Christianson, R. S. Sikorski, M. Dante, J. H. Shero, P. Hieter, Multifunctional yeast high-copy-number shuttle vectors. Gene 110, 119–122 (1992).

