## Supplementary material for "Architecture of the linker-scaffold in the nuclear pore": Hoelz_NPC_LinkerScaffold

**This PDF file includes:**

Materials and Methods
Figs. S1 to S78
Tables S1 to S14

### Materials and Methods

#### *Materials and reagents*

The sources of all materials and reagents are summarized in Table S1.

#### *Bacterial expression constructs*

The generation of cDNAs encoding *Chaetomium thermophilum* Nup192, Nup188, Nic96, Nup145N, and Nup53 was previously reported (*25, 30*). NUP98 cDNA was a gift of Beatriz Fontoura (UT Southwestern Medical Center, USA) (*83*). NUP155 cDNA was a gift of Susan Wente (Vanderbilt University, USA) (*84*). NUP53, NUP93, NUP62, NUP58, and NUP54 cDNAs were obtained commercially (Open Biosystems).

DNA fragments were amplified by PCR using *C. thermophilum* and *Homo sapiens* cDNA and cloned into either a modified pET28a vector encoding an N-terminal $His_6$-tag followed by a PreScission protease cleavage site (pET28a-PreS) (*85*), a modified pET28a vector (Novagen) encoding an N-terminal $His_6$-tag followed by a small ubiquitin-like modifier (SUMO) with or without an uncleavable C-terminal $His_6$-tag (pET28a-SUMO) (*86*), a modified pET28a vector (Novagen) encoding an N-terminal $His_6$-tag followed by an Avi-tag and SUMO (pET28a-Avi-SUMO), a modified pET-Duet1 vector (Novagen) encoding N-terminal SUMO-tags in both expression sites (pET-Duet1-SUMO2), or a modified pET-MCN vector (*87*) encoding an N-terminal $His_6$-tag followed by SUMO (pET-MCN-SUMO) (*25*). pET28a-PreS-Avi and pET28a-SUMO-Avi expression constructs were generated by subsequently cloning an Avi-tag-encoding DNA cassette into the NdeI and BamHI sites, respectively. If necessary, the SUMO protease ULP1 cleavage site between SUMO and the rest of the open reading frame was ablated by introducing a glycine to arginine or tyrosine substitution. Substitution mutants were generated by QuikChange mutagenesis using PfuUltra II Fusion HotStart DNA Polymerase (Agilent Technologies). Truncation mutants were generated by PCR amplification of desired fragments with primers encoding restriction sites followed by enzymatic restriction and ligation into linearized plasmid vectors. All mutagenesis was confirmed by Sanger sequencing. Details of bacterial expression constructs are summarized in Table S2.

#### *Protein expression and purification.*

Proteins were expressed in *Escherichia coli* BL21-CodonPlus(DE3)-RIL cells (Agilent Technologies) in Luria-Bertani media (Fisher bioreagent), induced at an $OD_{600}$ (optical density at 600 nm) of 0.8 with 0.5 mM isopropyl-β-D-thiogalactopyranoside (IPTG). A detailed summary of expression times and temperatures is found in Table S2. Seleno-L-methionine (SeMet)-labeled proteins were expressed with a methionine pathway inhibition protocol, as previously described (*88*). Cells were harvested by centrifugation and resuspended in a buffer containing 20 mM tris(hydroxymethyl)aminomethane (TRIS, pH 8.0), 500 mM sodium chloride (NaCl), and 5 mM 2-mercaptoethanol, supplemented with complete EDTA-free protease inhibitor cocktail (Roche), 2 μM bovine lung aprotinin (Sigma-Aldrich), and 1 mM phenylmethylsulfonyl fluoride (PMSF, Gold Biotechnology). Cell suspensions were supplemented with deoxyribonuclease I (Roche) and lysed with a cell disruptor (Avestin). Lysates were cleared by centrifugation at 30,000×*g* for 1 hour. The supernatants were filtered through a 0.45 μm filter (Millipore) and purified using standard chromatography methods, as outlined in Table S3. Proteins were concentrated to ~5-30 mg/ml for biochemical interaction experiments, complex reconstitution, and crystallization.

#### *Analytical size exclusion chromatography (SEC)*

For binding interaction analyses, all purified proteins and complexes were mixed and preincubated on ice for 30 minutes before injecting 200 μl of the sample on a Superdex 200 10/300 GL column (GE Healthcare) equilibrated in a buffer containing 20 mM TRIS (pH 8.0), 100 mM NaCl, and 5 mM dithiothreitol (DTT). Protein-containing fractions were analyzed by sodium dodecyl sulphate-polyacrylamide gel electrophoresis (SDS-PAGE) and visualized with Coomassie Brilliant Blue staining.

*Mutational analysis of the Nup192-Nic96 and Nup188-Nic96 interactions.* Purified Nup192 and Nup188 variants were mixed with purified SUMO-$Nic96^{R2}$ variants at final concentrations of 15 μM and 30 μM, respectively, prior to injection on a Superdex 200 10/300 GL column.

*5-Ala and scanning and single-residue Nup145N mutant analysis of the Nup192-Nup145N interaction.* Purified Nup192 was mixed with a purified Nup145N variants at final concentrations of 10 μM and 45 μM, respectively, prior to injection on a Superdex 200 10/300 GL column.

*Nup145N truncation mutant analysis of the Nup192-Nup145N interaction.* Purified Nup192 was mixed with a purified SUMO-Nup145N truncation variants at final concentrations of 10 μM and 45 μM, respectively, prior to injection on a Superdex 200 10/300 GL column.

*Single-residue Nup192 mutant analysis of the Nup192-Nup145N interaction.* Purified Nup192•SUMO-Nic96$^{R2}$ containing Nup192 variants was mixed with a 1.2-fold molar excess of purified Nup53 and injected on a Superdex 200 10/300 GL column. Protein containing fractions corresponding to the Nup192•SUMO-Nic96$^{R2}$•Nup53 containing Nup192 variants were collected, concentrated, and mixed with purified SUMO-Nup145N truncation variants at final concentrations of 10 μM and 45 μM, respectively, prior to injection on a Superdex 200 10/300 GL column.

*5-Ala scanning and single-residue Nup145N mutant analysis of the Nup188-Nup145N interaction.* Purified Nup188$^{NTD}$ was mixed with a purified Nup145N variants at final concentrations of 10 μM and 45 μM, respectively, prior to injection on a Superdex 200 10/300 GL column.

*Nup145N truncation mutant analysis of the Nup188-Nup145N interaction.* Purified Nup188$^{NTD}$ was mixed with a purified SUMO-Nup145N truncation variants at final concentrations of 10 μM and 45 μM, respectively, prior to injection on a Superdex 200 10/300 GL column.

*Single-residue Nup188 mutant analysis of the Nup188-Nup145N interaction.* Purified Nup188$^{NTD}$ variants were mixed with a purified SUMO-Nup145N$^{R2}$ at final concentrations of 10 μM and 45 μM, respectively, prior to injection on a Superdex 200 10/300 GL column.

*NUP53 truncation mutant analysis of the NUP155$^{NTD}$-NUP53 interaction.* Purified NUP155$^{NTD}$ was mixed with a purified NUP53 variants at final concentrations of 45 μM and 135 μM, respectively, prior to injection on a Superdex 200 10/300 GL column.

*NUP93 truncation and mutant analysis of CNT-NUP93 interaction.* Purified CNT was mixed with a purified SUMO-NUP93 variants at final concentrations of 20 μM and 60 μM, respectively, prior to injection on a Superdex 200 10/300 GL column.

*NUP53 truncation analysis of the NUP93$^{SOL}$-NUP53 interaction.* Purified NUP93$^{SOL}$ was mixed with purified NUP53 and SUMO-NUP53 truncation variants at final concentrations of 15 μM and 45 μM, respectively, prior to injection on a Superdex 200 10/300 GL column.

*5-Ala and single-residue mutant analysis of the NUP93$^{SOL}$-NUP53 interaction.* Purified NUP93$^{SOL}$ variants were mixed with purified SUMO-NUP53$^{N}$ variants at final concentrations of 15 μM and 45 μM, respectively, prior to injection on a Superdex 200 10/300 GL column.

***Multi-angle light scattering coupled to analytical size-exclusion chromatography (SEC-MALS)***

The mass of purified proteins and complexes was measured by inline multiangle light scattering (MALS) after separation on a Superdex 200 Increase 10/300 GL column (GE Healthcare) equilibrated in a buffer containing 20 mM TRIS (pH 8.0), 100 mM NaCl, and 5 mM DTT. The chromatographic system was connected in series with an 18-angle light-scattering detector (DAWN HELEOS II, Wyatt Technology), a dynamic light-scattering detector (DynaPro Nanostar, Wyatt Technology), and a refractive index detector (Optilab t-rEX, Wyatt Technology). Data were sampled every 1 second at a flow rate of 0.4 ml/minute, at 21 °C, and analyzed using ASTRA 6 (Wyatt Technology) to determine the molar mass and mass distribution (polydispersity) of the sample (*89*). For binding interaction analyses, proteins were mixed and preincubated on ice for 30 minutes prior to the injection of 200 μl of sample on the size exclusion column. Protein-containing fractions were analyzed by SDS-PAGE and visualized with Coomassie stain. Experimental and theoretical masses of all proteins and complexes are listed in Table S4.

*SEC-MALS analysis of the Nup192-Nic96 and the Nup188-Nic96 interaction.* Purified Nup192 and Nup188 variants were mixed with purified SUMO-Nic96$^{R2}$ variants at final concentrations of 15 μM and 22.5 μM, respectively, prior to SEC-MALS analysis.

*SEC-MALS analysis of the Nup192-Nup53 interaction and lack of the Nup188-Nup53 interaction.* Purified Nup192•SUMO-Nic96$^{R2}$ and Nup188•SUMO-Nic96$^{R2}$ were mixed with purified Nup53 at final concentrations of 15 μM and 22.5 μM, respectively, prior to SEC-MALS analysis.

*SEC-MALS analysis of the Nup192-Nup145N interaction.* Purified Nup192•SUMO-Nic96$^{R2}$ containing Nup192 variants were mixed with a 1.2-fold molar excess of purified Nup53 and injected on a Superdex 200 Increase 10/300 GL. Protein containing fractions corresponding to the Nup192•SUMO-Nic96$^{R2}$•Nup53 containing Nup192 variants were collected, concentrated, and mixed with purified Nup145 or SUMO-Nup145N$^{R1}$ variants at final concentrations of 15 μM and 22.5 μM, respectively, prior to SEC-MALS analysis.

*SEC-MALS analysis of the Nup188-Nup145N interaction.* Purified Nup188•SUMO-Nic96$^{R2}$ containing Nup188 variants were mixed purified Nup145 or SUMO-Nup145N$^{R2}$ variants at final concentrations of 15 μM and 22.5 μM, respectively, prior to SEC-MALS analysis.

#### *Isothermal titration calorimetry (ITC)*
Isothermal titration calorimetry (ITC) measurements were performed on an Affinity ITC instrument (TA Instruments). The buffer conditions were equalized between titrant and titrand by overnight dialysis against of the same buffer. Buffer conditions, cell temperatures, and initial concentrations of titrants and titrands for each ITC experiment are summarized in Table S5. Each titration was performed by injecting 2.5 μL with an injection frequency that allowed for baseline equilibration between injections. The ITC data was analyzed and plotted with the NanoAnalyze software (TA Instruments). We employed nonlinear least squares fit of a single-site binding model to estimate the dissociation constant ($K_D$), binding site multiplicity parameter $n$, and the enthalpy of binding ($\Delta°H$). All titrations were performed in triplicate, and mean and standard error of the mean values were estimated for $K_D$, $n$, and $\Delta°H$ quantities. Free energy ($\Delta°G$) and entropy ($\Delta°S$) of binding were calculated according to the thermodynamic identities $\Delta°G = \Delta°H - T\Delta°S$ *and* $\Delta°G = RT\, ln(K_D)$, and the error was propagated accordingly. The mean estimates and associated errors for thermodynamic parameters are reported in Table S5.

#### *Structure determination by X-ray crystallography*
Crystallization of proteins were performed at 21 °C in hanging drops containing 1 μl of protein and 1 μl of reservoir solution. Details about the crystallization and cryoprotection conditions are provided in Table S6. Crystals were cryoprotected by supplementing the drop with well solution containing cryoprotectant and vitrified by rapid transfer into liquid nitrogen. X-ray diffraction data were collected at 100 K at beamline BL12-2 at the Stanford Synchrotron Radiation Source (SSRL) and processed using XDS (*90*), unless specified otherwise. Iterative rounds of model building and refinement were performed using Coot (*91*) and PHENIX (*92*). All models presented excellent stereochemistry and geometry, as determined by MolProbity (*93*). Details regarding diffraction data processing and model refinement are provided in Tables S7-S10.

*Structure determination of Nup188$^{NTD}$.* Native and SeMet-derivatized Nup188$^{NTD}$ crystals grew in 0.1 M potassium thiocyanate and 10% (w/v) PEG-3,350 and diffracted to a resolution of 2.8 Å and 3.5 Å, respectively. Crystals were cryoprotected by gradually supplementing the drop with ethylene glycol in 1% (v/v) steps, to 25% (v/v). Initial phases were calculated in SHARP (*94*) using multiple anomalous dispersion (MAD) X-ray diffraction data obtained from the SeMet-derivatized Nup188$^{NTD}$ crystal. The initial model was built in the experimental electron map and refined against the higher resolution native data.

*Structure determination of Nup188•Nic96$^{R2}$.* Crystals of Nup188•Nic96$^{R2}$ were grown in 50 mM HEPES (pH7.5) and 6.5% (w/v) PEG-20,000. Initial crystals did not diffract beyond ~7 Å resolution and presented diffraction pathologies that precluded us from assigning the unit cell parameters and space group. We developed a dehydration protocol that involved a gradual increase of the PEG-20,000 concentration in the well by 1% (w/v) every 24 hours over the span of 7 days, by briefly unsealing the EasyXtal (Qiagen) tray well and pipetting the appropriate amount of 35% (w/v) PEG-20,000 stock solution. The dehydration allowed us to identify a SeMet-derivatized crystal that diffracted to 4.4 Å. Crystals were cryoprotected by gradually supplementing the drop with ethylene glycol in 1% steps, to 25% (v/v). X-ray diffraction data were integrated and scaled using DIALS (*95*). Further scaling and reduction were performed in CCP4 suite programs AIMLESS, POINTLESS, and ctruncate (*96*). Initial phases were calculated with PHASER using the PHENIX MR-SAD routine to combine single-wavelength anomalous dispersion (SAD) phases with molecular replacement (MR) phases obtained by placing the Nup188$^{NTD}$ and Nup188$^{Tail}$ (PDB ID 5CWU) (*25, 92, 97*). The sequence register of the region not modeled by the Nup188$^{NTD}$ and Nup188$^{Tail}$ fragments (residues 1135 to 1146), as well as the sequence register of the Nup188-bound Nic96$^{R2}$ polypeptide chain, were confirmed by calculating anomalous difference Fourier maps to locate the SeMet positions. Model B-factor refinement was performed with residue-group restraints and Translation-Libration-Screw-rotation (TLS) groups that were identified with the TLSMD server (*97*).

*Structure determination of Nup192$^{\Delta Head}$•Nic96$^{R2}$.* To favor crystallization of the Nup192$^{\Delta Head}$•Nic96$^{R2}$ complex, we employed a Nup192 mutant, encompassing residues 167-184 replaced with a GSGS linker previously established to improve crystal quality (*30*). Crystals were grown in 50 mM HEPES (pH 7.5), 8% (w/v) PEG-3350, 0.1 M ammonium sulfate, and 1% (v/v) 2-propanol. Two rounds of micro-seeding with a cat whisker were necessary to grow crystals of appropriate size, subsequently cryoprotected by gradually supplementing the drop with ethylene glycol in 1% steps, to 25% (v/v). The structure was phased by molecular replacement with PHASER, using a Nup192$^{\Delta Head}$ model (PDB ID 5HB3) (*30*). Because the map lacked electron density that could be assigned to Nic96 residues 187-239, we generated a shorter Nic96$^{R2}$ construct, comprising residues 240-301. Using an equivalent crystallization approach, we obtained crystals of SeMet-derivatized Nup192$^{\Delta Head}$•Nic96$^{R2}$ and Nup192$^{\Delta Head}$•Nic96$^{R2}$ A289M complexes. Phases for the Nup192$^{\Delta Head}$•Nic96$^{R2}$ dataset were obtained with PHASER by using the MR-SAD routine in PHENIX with Nup192$^{\Delta Head}$ (PDB ID 5HB4) as search model (*30, 98*). Sequence registers of the two $\alpha$-helices comprised by the Nup192-bound Nic96$^{R2}$ polypeptide were confirmed by calculating anomalous difference Fourier maps to locate the positions of the SeMet-labeled endogenous M263, as well as the A289M substitution.

*Structure determination of apo NUP93$^{SOL}$ and NUP93$^{SOL}$•NUP53$^{R2}$.* NUP93$^{SOL}$ crystals were grown by equilibrating a mixture of 1 $\mu$l protein solution and 1 $\mu$l of 75 mM TRIS (pH 7.5) and 11% (w/v) PEG-20,000 against a well solution consisting of 75 mM TRIS (pH 7.5) and 19% (w/v) PEG-20,000. Crystals were cryoprotected with mother liquor supplemented with 20% (v/v) ethylene glycol. Phasing was conducted by molecular replacement in PHASER with a search model generated from the *C. thermophilum* Nic96$^{SOL}$ structure (PDB ID 5HB3) (*30, 98*). Successful model placement and packing was achieved by accounting for conformational differences through the iterative placement of five normal mode fragments of the search model identified with the WebNMA server (*99*). Phases were subsequently improved by rebuilding the model with Autobuild (*100*). To aid with modeling, we calculated an experimental SAD map with PHASER from the anomalous contributions in the X-ray diffraction data contributed from sulfur atoms whose coordinates had been determined by initial modeling (*101*). The maps were improved by density modification with RESOLVE (*102*). Analogously to the crystallization of NUP93$^{SOL}$, co-crystals of NUP93$^{SOL}$•NUP53$^{R2}$ and NUP93$^{SOL}$•NUP53$^{R2}$ I94M were grown by equilibrating a mixture of 1 $\mu$l protein solution and 1 $\mu$l of 75 mM TRIS (pH 7.5) and 11% (w/v) PEG-20,000 against a well solution consisting of 75 mM TRIS (pH 7.5) and 17% (w/v) PEG-20,000. Crystals were cryoprotected with mother liquor supplemented with 20% (v/v) ethylene glycol and NUP53$^{R2}$ peptide. We maintained a ten-fold molar excess of the NUP53$^{R2}$ peptide to NUP93$^{SOL}$ throughout the crystallization and cryoprotection steps. The NUP93$^{SOL}$•NUP53$^{R2}$ structure was phased by molecular replacement with *apo* NUP93$^{SOL}$ as search model. The sequence register and directionality of the NUP53$^{R2}$ polypeptide were confirmed by calculating anomalous difference Fourier maps from diffraction data collected on crystals of NUP93$^{SOL}$•NUP53$^{R2}$ I94M reconstituted with SeMet-derivatized NUP53$^{R2}$ I94M.

***Structure determination by single particle cryo-electron microscopy (cryo-EM)***

*Cryo-EM sample preparation and data collection.* For all cryo-EM grid preparation, 3 $\mu$l of sample at 0.5 mg/ml were applied to glow-discharged 300 mesh R2/2 Quanitfoil grids (Micro Tools, Germany). Grids were blotted for 6-8 seconds with blot force -5 at 4 °C and 100% humidity before plunging into to liquid nitrogen temperature ethane using a FEI Vitrobot (Thermo Fisher Scientific, USA) automatic plunger. The grids were imaged using a 300 keV Titan Krios electron microscope (Thermo Fisher Scientific, USA) equipped with an energy filter and Gatan K2 Summit or Gatan K3 (Thermo Fisher Scientific, USA) direct electron detectors. High magnification movies were automatically recorded with SerialEM or EPU (*103, 104*). Data collection, processing, and model refinement parameters and statistics are provided in Tables S7 and S8.

*Cryo-EM data processing.* Micrographs were dose-weighted, binned and motion corrected in MotionCor2 (*105*). Contrast transfer function (CTF) parameters were estimated using cryoSPARC PatchCTF (*106*). We selected a subset of micrographs that presented intact thin ice, homogeneous particle distribution, and excellent CTF fits from which particles were manually picked to generate initial particle templates, followed by reference-based automated picking in cryoSPARC (*106*). *Ab initio* reconstruction models were generated in cryoSPARC (*106*). Particle datasets were cleaned up with 2D and 3D classification in cryoSPARC (*106*) and RELION (*107*). Specific steps in the data processing are outlined in detail in the respective workflow diagrams (figs. S3, S16, and S25). Pixel and box size parameters specific to each structure are outlined in Tables S11 and S12. Refinement of particle orientations was carried out in cryoSPARC (*106*) and RELION (*107*). Fourier shell correlation (FSC) curves were calculated with 3DFSC (*108*). Angular distribution plots were generated with RELION (*107*) and rendered with UCSF Chimera (*109*). Local resolution was calculated with cryoSPARC Local Resolution (*106*). In the general workflow, particle and micrograph files were converted and manipulated with PyEM (*110*) .

*Model building and refinement.* All initial models were rigid body fit into cryo-EM maps with UCSF Chimera (*109*). Further model building and adjustment was performed iteratively through manual modification in Coot (*91*) and real-space refinement in PHENIX (*111*). Model quality was assessed with MolProbity (*93*) and EMRinger (*112*).

*Structure determination of Nup192•Nic96$^{R2}$.* We vitrified a purified stoichiometric Nup192•Nic96$^{R2}$ complex in a buffer solution consisting of 100 mM NaCl, 20 mM TRIS (pH 8.0), 5 mM DTT on cryo-EM grids. Movie collection was automated with EPU and consisted in applying stage shift to record three acquisitions per hole (*104*). The processing workflow applied to these data is outlined in fig. S3. The final 3.8 Å reconstruction was calculated from particle orientations refined with the cryoSPARC homogeneous refinement module (*106*). The cryo-EM map was largely explained by the Nup192•Nic96$^{R2}$ composite crystal structure. A conformational difference between the crystal and single particle cryo-EM Nup192 structures could be rigid body refined to fit the Nup192•Nic96$^{R2}$ composite crystal structure into the cryo-EM density with Coot (*91*) and further real-space refined in PHENIX (fig. S4) (*111*).

*Structure determination of Nup192•Nic96$^{R2}$•Nup145N$^{R1}$•Nup53$^{R1}$.* The sample was prepared by incubating a 50-fold molar excess of purified Nup145N$^{R1}$ and Nup53$^{R1}$ peptides with purified Nup192•Nic96$^{R2}$ in a buffer solution consisting of 150 mM NaCl, 20 mM TRIS (pH 8.0), 5 mM DTT, before applying the mixture to cryo-EM grids. The movie collection strategy relied on beam tilt to collect four acquisitions per hole over nine holes before moving the stage. Beam tilt compensation was applied in SerialEM to maintain coma-free alignment during collection (*103*). The presence of systematic beam tilt was further assessed for micrographs split according to collection tilt groups in RELION (*107*). The processing workflow applied to the data is outlined in fig. S16. Following 2D classification, an *ab initio* reconstruction and ~3.7 Å refined map of the Nup192•Nic96$^{R2}$•Nup145N$^{R1}$•Nup53$^{R1}$ complex were calculated in cryoSPARC (*106*). The particle dataset was cleaned up further through heterogeneous 3D refinement in cryoSPARC with a template set consisting of the ~3.7 Å map low pass filtered to 20 Å, with all but one template's phases randomized in RELION (*107*) in order to capture spurious particle picks and low-quality particles (in two iterations, with phases randomized beyond 40 Å and 25 Å, respectively). A final 3.2 Å 3D reconstruction was obtained by non-uniform refinement in cryoSPARC, with map interpretability improved in regions corresponding to the surface-bound peptides Nic96$^{R2}$, Nup145N$^{R1}$, and Nup53$^{R1}$ through local sharpening with PHENIX Autosharpen (*113, 114*). The single particle cryo-EM Nup192•Nic96$^{R2}$ structure was rigid-body fit into the density for further modelling in Coot (*91*) and real-space refinement in PHENIX (*111*). Despite the Nup145N$^{R1}$ sequence consisting of 67 residues, the resolved cryo-EM density was consistent with an eight-residue chain. The Nup145N$^{R1}$ peptide was modeled by considering all possible eight residue sequence windows to identify one most congruent with the side chain features present in the cryo-EM density and the binding site chemical properties. The Nup145N$^{R1}$ chain structure was then validated by mutational analysis of the Nup145N sequence (Fig. 4). Cryo-EM density that could be modeled as a dipeptide containing a bulky residue was observed in a hydrophobic groove on the Nup192 surface previously mapped as the Nup53 binding site. Of the 36 residues in the Nup53$^{R1}$ sequence, the F48-G49 motif was most consistent with density and previous mutational analysis that identified F48 as the fulcrum of the Nup192-Nup53 interaction (*36*).

*Structure determination of Nup188•Nic96$^{R2}$ and Nup188•Nic96$^{R2}$•Nup145N$^{R2}$.* The sample was prepared by incubating a 20-fold molar excess of purified Nup145N$^{R2}$ peptide with purified Nup188•Nic96$^{R2}$ in a buffer solution consisting of 100 mM NaCl, 20 mM TRIS (pH 8.0), 5 mM DTT, before applying the mixture to cryo-EM grids. The movie collection strategy relied on beam tilt to collect four acquisitions per hole over nine holes before moving the stage. Beam tilt compensation was applied in SerialEM to maintain coma-free alignment during collection (*103*). The presence of systematic beam tilt was further assessed for micrographs split according to collection tilt groups in RELION (*107*). The processing workflow applied to the data is outlined in fig. S25. Briefly, following 2D classification and *ab initio* 3D reconstruction in cryoSPARC (*106*), the particle set was further narrowed by iterative 3D refinement, classification, CTF refinement and Bayesian polishing in RELION to obtain a 2.4 Å reconstruction (*107*). The Nup188•Nic96$^{R2}$ crystal structure was found to fit well in the 2.4 Å Nup188•Nic96$^{R2}$ map and further real-space refined in PHENIX (*111*). Upon inspection of the initial map at low contour levels, a weak yet distinctly tubular density was observed in the saddle-like groove proximal to the SH3-like domain of Nup188, later confirmed to correspond to the Nup145N$^{R2}$ density. Iterative 3D classification of particles subtracted around the extra density into two classes was performed in RELION to isolate a homogenous subset of particles containing the Nup145N$^{R2}$ density. The T-regularization parameter in RELION was optimized to maximize the difference between the classes (*107*). After reverting the selected subset of subtracted particles to non-subtracted particles, a final 2.8 Å reconstruction of the Nup188•Nic96$^{R2}$•Nup145N$^{R2}$ complex was obtained by non-uniform refinement in cryoSPARC (*106*). Despite 88 residues being present in the Nup145N$^{R2}$ sequence, the resolved cryo-EM density was consistent with a 13-residue chain. The Nup145N$^{R2}$ peptide was modeled by considering all possible 13-residue sequence windows and identifying the sequence most congruent with side chain features present in the cryo-EM density and binding site chemical properties. The Nup145N$^{R2}$ chain structure was then validated by mutational analysis of the Nup145N sequence (Fig. 5). The cryo-EM map interpretability was iteratively improved by real-space refinement in PHENIX (*111*) and by applying refined atomic B factor-based sharpening in LocScale (*115*).

#### *Pulldown assay for NUP188 and NUP205 binding to linkers NUP93, NUP98, and NUP53*

*NUP205 and NUP188 expression in S. cerevisiae. H. sapiens* NUP205 cDNA was obtained commercially (Kazusa Genome Technology). *H. sapiens* NUP188 cDNA lacking the coding region for residues 1738-1749 was obtained commercially (DNAsu Plasmid Repository) and modified by primer extension PCR to include the missing codons. The human NUP205 and NUP188 open reading frames were amplified by PCR and cloned into the multiple cloning site of a pRS426 2μ yeast plasmids by enzymatic restriction with BamHI and NotI and ligation, downstream of a $P_{GAL1-10}$ promoter inserted between SalI and EcoRI restriction sites. A DNA cassette encoding 3×FLAG-His$_6$ was subsequently cloned into the BamHI site. Plasmids were transformed into the BY4741 *S. cerevisiae* strain by the lithium acetate method and selected on synthetic dextrose complete without uracil (SDC-URA) solid media. Large scale 1 L cultures in synthetic raffinose complete without uracil (SRC-URA) liquid media were inoculated 1:100, grown at 30 ºC to an $OD_{600}$ of 1.0, and supplemented with galactose to a final concentration of 2% (w/v) to induce expression. After induction, yeast cultures were incubated at 30 ºC for an additional 6 hours before harvesting by centrifugation. Cells were resuspended at an $OD_{600}$ of 400 in lysis buffer containing 20 mM TRIS (pH 7.5), 150 mM NaCl, 5 mM DTT, 5% (v/v) glycerol, 5 mM ethylenediaminetetraacetic acid (EDTA, pH 8.0), and supplemented with 2 μM bovine lung aprotinin (Sigma-Al), 1 mM PMSF and complete EDTA-free protease inhibitor cocktail (Roche). The suspension was flash frozen in liquid nitrogen before cryo-milling (Retsch). The pulverized frozen lysate was stored at -80 ºC. To generate clarified lysate for the pulldown assay, thawed aliquots of the pulverized lysate were centrifuged twice for 10 minutes at 30,130×*g* and 4 ºC, transferring the supernatant into a clean Eppendorf tube between centrifugations.

*In vitro biotinylation of Avi-tagged bait proteins.* Purified His$_6$-Avi-SUMO-tagged NUP93$^{R2}$, NUP98 and NUP53 variants were biotinylated at a protein concentration of 40 μM in a buffer solution composed of 50 mM bicine (pH 8.3), 100 mM biotin, 10 mM adenosine triphosphate (ATP), 10 mM magnesium acetate, and 20 μg/ml biotin ligase (BirA) at 30 °C for 4 hours. Subsequently, biotinylated protein was purified from the excess biotin with a 5 ml HiTrap Desalting (GE Healthcare) column equilibrated with a buffer containing 20 mM TRIS (pH 8.0), 100 mM NaCl, and 5 mM DTT. The efficiency of biotinylation was evaluated by incubating the biotinylated protein with Streptavidin MagneSphere Paramagnetic Particles (Promega) and assessing the ratio of bound to unbound protein by SDS-PAGE and Coomassie brilliant blue staining of the two fractions.

*In vitro pulldown and immunoblotting.* Streptavidin MagneSphere Paramagnetic Particles (Promega) resin was equilibrated with a buffer containing 20 mM TRIS (pH 8.0), 100 mM NaCl and 5 mM DTT. The resin was then incubated with 200 ng of biotinylated bait protein per 1 μl of resin for 30 minutes at room temperature, followed by removal of the supernatant and three resin-volume washes with a washing buffer composed of 20 mM TRIS (pH 7.5), 150 mM NaCl, 1 mM DTT, and 0.05% (v/v) Tween-20. For each pulldown, 50 μl of resin were aliquoted and resuspended in 25 μl of washing buffer before incubation with 300 μl of clarified *S. cerevisiae* lysate for 30 minutes on ice. After the removal of the supernatant, the resin was washed with 50 μl of washing buffer, transferred to a clean Eppendorf tube, and resuspended in 25 μl of washing buffer. Finally, 8.33 μl of 4×SDS-PAGE loading buffer were added to the resin and the mixture was incubated for 2 minutes at 98 ºC. Load, bait and pulldown samples were resolved with SDS-PAGE. The bait-only samples were visualized with Coomassie staining, whereas the pulldown and load samples were transferred to a PVDF membrane, which was blocked for 1 hour at room temperature in 5% (w/v) fat-free milk powder resuspended in phosphate buffered saline supplemented with Tween 20 (PBS-T) buffer. All antibody blotting was performed in SuperBlock-PBS blocking buffer (Thermo Fisher Scientific) and subsequent washes were performed at room temperature in PBS-T buffer. To detect 3×FLAG-tagged NUP205 and NUP188, the PVDF membrane was incubated for 1 hour at room temperature with an mouse monoclonal anti-FLAG primary antibody (Sigma-Aldrich; 1:6,000 dilution), washed three times, and subsequently incubated for 1 hour at room temperature with a secondary goat anti-mouse antibody fused to an IR800 fluorescent protein (LI-COR; 1:6,000 dilution), which was detected with a Li-Cor Odyssey imager using the 800 nm channel at 169 μm resolution and a scan intensity of 2.5.

#### *Multispecies sequence alignment*

To infer evolutionary conservation between *C. thermophilum*, *S. cerevisiae*, and *H. sapiens*, we sampled primary sequences from species broadly representative of the phylogenetic diversity between fungi and metazoans. Primary sequences of Nup188, Nup192 and Nic96 scaffold nup homologs were aligned with the PROMALS3D server (*116*). The highest resolution available structures from different organisms were used as inputs to facilitate the alignment. Linker nup alignments were generated with MAFFT (*117*). Whereas it was possible to align Nup145N sequences from fungi to metazoans, some binding region motifs in Nup53 could not be aligned, in some cases even between the fungal species *S. cerevisiae* and *C. thermophilum*. To study sequence conservation of Nup53 homologs on a narrower evolutionary scale, we generated individual multispecies alignments of Nup53 homolog sequences from species related to *C. thermophilum*, *S. cerevisiae*, and *H. sapiens*. Sequence alignments were colored with ALSCRIPT by sequence similarity according to the BLOSUM62 matrix (*118*).

#### *S. cerevisiae reverse genetics and functional assays*

*S. cerevisiae culture and expression plasmids.* Preparation of all media (yeast peptone dextrose, YPD; synthetic dextrose complete, SDC; synthetic galactose complete, SGC) and lithium acetate yeast transformation were carried out according to standard protocols. For plasmid marker selection, selective media was formulated by omitting the appropriate amino acids or nucleobase (leucine, -LEU; histidine, -HIS; methionine, -MET; uracil -URA). To construct *S. cerevisiae* nup expression plasmids, fragments were amplified by PCR from S. *cerevisiae* genomic DNA and inserted by enzymatic restriction and ligation into the multiple cloning site of plasmids from the pRS series containing a $P_{Nop1}$ promoter (*119*). Mutants were generated by QuikChange mutagenesis using PfuUltra II Fusion HotStart DNA Polymerase (Agilent Technologies), or by PCR amplification of desired fragments with primers encoding restriction sites, followed by enzymatic restriction and ligation into linearized plasmid vectors. According to need, nup genes and gene variants were subcloned into vectors containing different N-terminal tags (eGFP, mCherry, 3×FLAG, and 3×HA) or C-terminal tags (mCherry and 3×HA) with different selection markers (*MET15*, pRS411; *HIS3*, pRS413; *LEU2*, pRS415; *URA3*, pRS416) by restriction enzyme digestion and ligation. The pRS415-$P_{Nop1}$-3×FLAG and pRS415-$P_{Nop1}$-3×HA plasmids were generated by inserting 3×FLAG or 3×HA DNA cassettes into HindIII and BamHI or HindIII and NotI sites by enzymatic restriction and ligation, respectively. All mutagenesis and subcloning was confirmed by Sanger sequencing. Details of *S. cerevisiae* constructs employed in this study are summarized in Table 13.

*S. cerevisiae genetic manipulation.* Starting from a parent BY4741 strain, haploid knockout strains were generated by homologous recombination of genomic loci with PCR-amplified *KanMX6* or the *natNT2* cassettes introduced into cells by lithium acetate transformation, as previously described (*120, 121*). Subsequently, *KanMX6* and *natNT2* genomic integration was selected for on YPD solid media containing G418 (Gold Biotechnology) or nourseothricin (Gold Biotechnology), respectively, and confirmed by PCR amplification of genomic DNA and Sanger sequencing. Prior to knocking out a lethal gene, parental strains were complemented with a pRS416 plasmid carrying the respective wildtype *S. cerevisiae* gene controlled by a $P_{Nop1}$ promoter. The generation of *nup192Δ, nic96Δ,* and *nic96Δnup57-eGFP* strains was described previously (*25*). Details about haploid yeast strains employed in this study are summarized in Table 14.

*Generation of the nup100Δnup116Δnup145Δ strain.* We generated a *loxP-natNT2-loxP* cassette by flanking the *natNT2* cassette with upstream and downstream *loxP* sequences (5'-ATAACTTCGTATAATGTATACTATACGAAGTTAT-3') via QuikChange mutagenesis of the previously described pFA6a-natNT2 plasmid (*120*). We used the *loxP-natNT2-loxP* cassette to knock out the *NUP116* gene in the *nup100Δ* strain transformed with the pRS416-$P_{Nop1}$-*scnup116-nup145C* chimera plasmid. The *scnup116-nup145C* chimera gene was generated by fusing DNA fragments coding for the Nup116 residues 1-967 and Nup145C residues 1-712 by overlap PCR extension. The *loxP-natNT2-loxP* was excised from the *nup100Δnup116Δ* genome by expressing the Cre recombinase from the pRS413-$P_{GAL1\text{-}10}$-*Cre* plasmid on SGC-HIS solid media. The Cre cDNA was obtained as a gift from David Baltimore (California Institute of Technology, USA) (*122*), amplified by PCR, and cloned into a pRS413 plasmid by enzymatic restriction with XbaI and NotI and ligation, following the insertion of a $P_{GAL1\text{-}10}$ promoter cassette by enzymatic restriction with SalI and EcoRI and ligation. The pRS413-$P_{GAL1\text{-}10}$-*Cre* plasmid was then removed from the strain by outgrowth on solid YPD media followed by two rounds of selection against growth on SDC-HIS solid media. Next, we knocked the *NUP145* gene out with a *natNT2* cassette. Although the pRS416-$P_{Nop1}$-*scnup116-nup145C* chimera plasmid rescued the lethal triple gene knockout in the *nup100Δnup116Δnup145Δ* strain, the strain displayed a slow growth phenotype. We established that complementation of the *nup100Δnup116Δnup145Δ* strain with pRS415-$P_{Nop1}$-*scNUP116* and pRS413-$P_{Nop1}$-*nup145C* plasmids resulted in wildtype growth rates. Hence, before further shuffling with *NUP116* variants, we substituted the pRS416-$P_{Nop1}$-*scnup116-nup145C* chimera rescue plasmid with the pRS416-$P_{Nop1}$-*scNUP116* plasmid and either a pRS413-$P_{Nop1}$-*nup145C*, pRS413-$P_{Nop1}$-*nup145C-mCherry,* or pRS413-$P_{Nop1}$-*nup145C*-3×HA plasmid.

*Mutant viability and growth assays.* To test the effect of nup variants, pRS415, pRS415-3×HA, pRS415-3×FLAG, pRS415-eGFP or pRS415-mCherry constructs containing nup gene variants controlled by a $P_{Nop1}$ promoter were transformed into a knockout strain and selected twice on SDC-LEU solid media. To 'shuffle' the complementing pRS416 plasmids, transformants were grown until mid-log phase at 30°C, diluted to an $OD_{600}$ of 0.2 and 7.5 μl, and 15 μl of a ten-fold dilution series were spotted onto SDC-LEU and SDC with 5-fluoroorotic acid (SDC+5-FOA) solid media, respectively, and incubated at favorable growth temperatures (23°C for all strains except for *nup188Δpom34Δ* and *nup188Δpom152Δ*, which were incubated at 30 °C). The viable transformants were subjected to an ulterior round of selection on SDC+5-FOA. Growth of viable transformants was analyzed by diluting mid-log phase cells to an $OD_{600}$ of 0.2, spotting 5 μl of a ten-fold dilution series on YPD solid media, and incubating the spotted cultures at different temperatures.

*Western blot analysis of protein expression.* To ascertain the expression of plasmid constructs, we repeated all growth and viability assays with transformants carrying plasmids with 3×FLAG and 3×HA-tagged variants. Protein extraction from mid-log phase yeast cells was performed via NaOH and SDS treatment, as previously described (*123*). The total protein extracts were resolved by SDS-PAGE and transferred to a PVDF membrane, which was blocked for 1 hour at room temperature in 5% (w/v) fat-free milk powder resuspended in PBS-T buffer. All antibody blotting was performed in SuperBlock-

PBS blocking buffer and subsequent washes were performed at room temperature in PBS-T buffer. Western blotting of Nup145C-3×HA and 3×HA-Nup192 variants was performed with a 1-hour incubation at room temperature with a mouse monoclonal anti-HA antibody (Biolegend; 1:5,000 dilution). Western blotting of 3×FLAG-Nic96, 3×FLAG-Nup116 and 3×FLAG-Nup188 variants was performed with a 1-hour incubation at room temperature with an mouse monoclonal anti-FLAG antibody (Sigma-Aldrich; 1:6,000 dilution). Equal loading was established with an overnight incubation at 4 °C with rabbit anti-hexokinase antibody (US Biological; 1:6,000 dilution). Primary antibody binding was detected by 1 hour incubation at room temperature with a goat anti-mouse antibody fused to an IR800 fluorescent protein (LI-COR, 1:6,000 dilution) or a goat anti-rabbit antibody fused to an IR800 fluorescent probe (LI-COR, 1:6,000 dilution), respectively, and imaged with an Li-Cor Odyssey imager using the 800 nm channels in a single scan at 169 μm resolution and a scan intensity of 5.

*Live cell fluorescence for subcellular localization studies.* Liquid cultures of shuffled and non-shuffled strains were grown at 30°C in YPD and SDC-LEU media, respectively. To assess the effect of shifting temperature, liquid cultures were grown at 37 °C for 6 hours (*nic96Δ* and *nup192Δ* strains), 37 °C for 4 hours (*nup100Δnup116Δnup145Δ* strain), or 16 °C for 4 hours (*nup188Δ* and *nup188Δpom34Δ* strains). For fluorescence and differential interference contrast (DIC) microscopy imaging, cells were pelleted by centrifugation for 2 minutes at 1000 × g, resuspended in 100 mM potassium phosphate (pH 6.4), and rapidly imaged at room temperature using a Carl Zeiss Observer Z.1 equipped with a Hamamatsu camera C10600 Orca-R2.

*60S pre-ribosome export assay.* Knockout strains were co-transformed with pRS415-$P_{Nop1}$ plasmids carrying variants of the knocked-out gene, and with a pRS411-$P_{Nop1}$-*sc*Rpl25 plasmid expressing an mCherry-tagged L25 ribosomal protein. Following shuffling of the rescue plasmids on SDC-MET+5-FOA solid media, liquid cultures of yeast transformants were grown at 30°C in SDC-MET media to an $OD_{600}$ of 0.4 and subsequently shifted to 37°C for 6 hours (*nic96Δ* strain), 16°C for 4 hours (*nup188Δpom34Δ* strain) or not shifted but allowed to reach $OD_{600}$ of 1.0 (*nup100Δnup116Δnup145Δ* strain). For fluorescence and differential interference contrast (DIC) microscopy, cells were pelleted by centrifugation for 2 minutes at 1000 × g, resuspended in 100 mM potassium phosphate (pH 6.4), and rapidly imaged at room temperature using a Carl Zeiss Observer Z.1 equipped with a Hamamatsu camera C10600 Orca-R2.

*Fluorescent in situ hybridization (FISH) mRNA export assay.* The preparation of *S. cerevisiae* cells for FISH was adapted from previously established protocols (*20, 124*). Liquid cultures of shuffled yeast transformants were grown at 30 °C in YPD media to an $OD_{600}$ of 0.4 and subsequently shifted to 37°C for 6 hours (*nic96Δ* strain), 30 °C and 37 °C for 4 hours (*nup100Δnup116Δnup145Δ* strain), or 16 °C for 4 hours (*nup188Δpom34Δ* strain). Cell washes were carried out by centrifuging cells for 2 minutes at 1000 × g followed by gentle resuspension. For each experimental group, cells were resuspended at $OD_{600}$ of 1.0 in 1 ml of 4% (v/v) formaldehyde, 100 mM potassium phosphate (pH 6.4), and fixed for 1.5 hours at 23 °C. Following two washes in 100 mM potassium phosphate (pH 6.4) and two washes in 100 mM potassium phosphate (pH 6.4), 1.2 M sorbitol, cell walls were digested in 100 mM potassium phosphate (pH 6.4), 1.2 M sorbitol, 2.5 mg/ml T20 zymolyase (US Biological) for 40 minutes at 30 °C. Following two gentle washes with 100 mM potassium phosphate (pH 6.4), 1.2 M sorbitol, cells were resuspended in 200 μl of the same buffer and cell walls further sheared by pipetting the cells 20 times. The resulting spheroplasts were placed on freshly poly-lysinated coverslips for 10 minutes and the unadhered spheroplasts pipetted off before the coverslips were submerged in -80 °C methanol for 6 minutes followed by a 5 second submersion in -80 °C acetone and rapid drying under an air stream for 30 minutes. The following washes and incubations were carried out by pipetting 500 μl of solution on and off the spheroplasts fixed onto coverslips. The coverslips were incubated with 100 mM triethanolamine (pH 8.0) for 2 minutes at 23 °C before they were incubated with 100 mM triethanolamine (pH 8.0), 0.25% (v/v) acetic anhydride for 10 minutes at 23 °C. After two washes with 2× saline sodium citrate buffer (SSC), coverslips were inverted and incubated for 30 minutes at 37 °C onto 40 μl of hybridization buffer, consisting of 50% (v/v) formamide (Thermo Fisher), 10% (w/v) dextran sulfate (Sigma-Aldrich), 4×SSC, 1×Denhardt's solution (Sigma-Aldrich), 125 μg/ml *E. coli* tRNA (Roche), 500 μg/ml salmon sperm DNA (Invitrogen). Next, coverslips were incubated on hybridization buffer including 1.67 μg/ml Alexa Fluor

647-labeled 50-mer oligo(dT) probe (Integrated DNA Technologies) for 16 hours at 37 °C in a humid chamber equilibrated with 2×SSC. Following hybridization, coverslips were sequentially incubated in the dark for 1 hour in 2×SSC at 23 °C, for 30 minutes in 1×SSC at 23 °C, for 30 minutes in 0.5×SSC at 37 °C, and for 30 minutes in 0.5×SSC at 23 °C before mounting in DAPI-containing mounting solution (ProLong Gold Antifade, Invitrogen), and imaging with a Carl Zeiss Observer Z.1 microscope equipped with a Hamamatsu camera C10600 Orca-R2.

*Quantitation of 60S pre-ribosome and mRNA export assays.* In the 60S pre-ribosome export assay, the nuclear rim was identified by the localization of GFP-tagged nup variants to the nuclear envelope and the nuclear retention of S60 pre-ribosomal particles was assessed based on the relative intensity between the nucleus and the cytoplasm of the signal from the mCherry-tagged L25 ribosomal protein. In the mRNA export assay, the DAPI staining of chromatin was used to identify nuclei and nuclear retention of poly(A)$^+$ RNA as assessed based on the relative intensity between the nucleus and the cytoplasm of the signal from the Alexa647-labeled 50-mer oligo(dT) probe used to FISH-label poly(A)$^+$ RNA. For both ribosomal and mRNA export assays, five individuals independently assessed >500 cells for each imaged variant without knowing the identity of the images. The ratios of cells with nuclear retention phenotype to the total number of assessed cells reported by the five individuals were averaged to produce a mean value. Three independent biological replicas (independent transformants of a variant plasmid) were assayed and quantitated for each experiment. The mean of the three replicas' ratio of cells with nuclear retention of mRNA or 60S pre-ribosomes along with the associated standard error were reported.

#### *Docking of near-atomic resolution structures into cryo-electron tomographic (cryo-ET) maps*

*Quantitative docking by global searching and statistical scoring.* For each quantitative docking experiment, we used the Chimera fitmap tool to generate 1 million random placements of volumes simulated from crystal and single particle cryo-EM structures low pass filtered to the resolution of the cryo-ET map. Placements were limited to an asymmetric subunit of the C8 symmetry averaged cryo-ET map from which nuclear envelope regions had been eliminated by segmentation, and a volume overlap cutoff of 20% was employed to limit the random placements to map density. To evaluate the docking within a statistical framework, we applied an analysis similar to previous statistical approaches to docking high resolution structures into lower resolution cryo-ET maps (*62*). After local rigid body optimization, correlations about the mean (Pearson correlations) were calculated for every placed volume and the map, and then Fisher z-transformed and normalized to approximate a normal sampling distribution. Because correct placements are rare events, we used the Fisher z-score distribution as a null distribution from which to calculate one-tailed p-values for the top-scoring solutions. We evaluated several of the top scoring solutions visually for shape complementarity between the fit volume and the cryo-ET map, and for clashes with other placed structures.

*Docking of near-atomic resolution structures into the in situ S. cerevisiae cryo-ET map.* We performed global searches in an ~25 Å sub-tomogram averaged cryo-ET map of the *in situ* imaged *S. cerevisiae* NPC (EMD-10198) with the single particle cryo-EM Nup188•Nic96$^{R2}$•Nup145N$^{R2}$ and Nup192•Nic96$^{R2}$•Nup145N$^{R1}$•Nup53$^{R1}$ structures, and the composite crystal Nup192•Nic96$^{R2}$•Nup145N$^{R1}$•Nup53$^{R1}$ structure that included the crystal structure conformation of Nup192 (*31*). Top scoring solutions placed 16 Nup188•Nic96$^{R2}$•Nup145N$^{R2}$ copies in the peripheral question mark shaped densities on both nuclear and cytoplasmic sides of the inner ring. Likewise, top scoring solutions placed 16 Nup192•Nic96$^{R2}$•Nup145N$^{R1}$•Nup53$^{R1}$ copies in the equatorial question mark shaped densities on both nuclear and cytoplasmic sides of the inner ring, with marginally higher confidence scores for the placement of the crystal structure conformation of Nup192. Next, we placed 32 copies of Nup170•Nup145N$^{R3}$•Nup53$^{R3}$, Nic96$^{SOL}$•Nup53$^{R2}$, and CNT•Nic96$^{R1}$ into the inner ring, respectively, by superposition with the previously determined integrative model of the *S. cerevisiae* NPC and local rigid body fitting into the cryo-ET map density (*31*). The C-terminal $\alpha$-helical solenoid domain of Nup170 was shown to be conformationally flexible (*30*). To improve the fit into the cryo-ET density, the Nup170 NTD and CTD portions of the peripheral Nup170•Nup145N$^{R3}$•Nup53$^{R3}$ were independently rigid body refined against the cryo-ET map. Finally, the CNC and Nup82•Nup159•Nsp1 complex models were retained from the previously available integrative model of the *S. cerevisiae* NPC (*31*).

*Incremental approach to quantitative docking into the human NPC cryo-ET map.* The incremental docking approach employed in this study was similar to the one previously applied to the docking of symmetric core high resolution structures into a ~23 Å cryo-ET map of the intact human NPC and is graphically outlined in fig. S54 (*30*). Structures of CNC, Nup192•Nic96$^{R2}$•Nup145N$^{R1}$•Nup53$^{R1}$, Nup188•Nic96$^{R2}$•Nup145N$^{R2}$, and the NUP358$^{NTD}$ (reported in the accompanying manuscript) were quantitively docked into the full ~12 Å cryo-ET map of the intact human NPC (*41, 61*). Subsequently, the density explained by the docked structures was subtracted from the cryo-ET density of the full NPC map, as well as a map segmented to cover only the inner ring. Iteratively, quantitative docking was performed with the NUP93$^{SOL}$•NUP53$^{R2}$, Nup170•Nup145N$^{R3}$•Nup53$^{R3}$ (multiple conformations of Nup170), CNT•Nic96$^{R1}$, and NUP53$^{RRM}$, with the assigned cryo-ET density subtracted from the maps before each subsequent round of global searching.

*Docking of CNC into the human NPC cryo-ET map.* Previously reported composite structures of the Y-shaped CNC hetero-nonamer, composed of the yeast CNC-hexamer (PDB ID 4XMM) (*26*), the NUP84•NUP133 hetero-dimer (PDB ID 3I4R) (*21*), NUP43 (PDB ID 4I79) (*27*), NUP37 (PDB ID 4FHM) (*22*), and NUP133$^{NTE}$ (PDB ID 1XKS) (*12*)crystal structures, were quantitatively docked into an ~12 Å cryo-ET map of the intact human NPC. Top solutions identified two concentric rings of eight CNCs sat atop each side of the nuclear envelope, for a total of 32 CNC copies per NPC (*30*). The composite CNC structures agreed well with an ~12 Å cryo-ET map density, apart from the NUP133$^{NTE}$ domains, which had to be locally rigid body fit at each of the four unique locations. Details about the docking of CNC and local fitting of NUP133$^{NTE}$ are reported in fig. S54.

*Docking of NUP358$^{NTD}$ into the human NPC cryo-ET map.* As described in more detail in the accompanying manuscript, five copies of the NUP358$^{NTD}$ open conformation were identified as top scoring solutions per human NPC spoke of the ~12 Å cryo-ET map of the intact human NPC, for a total of 40 copies per NPC (*61*).

*Docking of Nup192•Nic96$^{R2}$•Nup145N$^{R1}$•Nup53$^{R1}$ into the human NPC cryo-ET map.* We performed global searches with the single particle cryo-EM Nup192•Nic96$^{R2}$•Nup145N$^{R1}$•Nup53$^{R1}$ structure and a composite Nup192•Nic96$^{R2}$•Nup145N$^{R1}$•Nup53$^{R1}$ structure that included the crystal structure conformation of Nup192, in an ~12 Å cryo-ET map of the intact human NPC. In both docking experiments, the three highest ranking solutions identified 16 copies of NUP205 at proximal and distal positions in the cytoplasmic outer ring and eight copies of NUP205 at a distal position in the nuclear outer ring equivalent to the cytoplasmic distal position. The top two results from global searches in the inner ring, as well as the 4th and 5th top results in the full map, were placed in equatorial question mark-shaped densities in the inner ring. Thus, a total of 40 copies of NUP205 were assigned in the human NPC. As a benchmark for quantitative docking in the new ~12 Å cryo-ET map, we replicated the quantitative docking experiments with the novel Nup192•Nic96$^{R2}$•Nup145N$^{R1}$•Nup53$^{R1}$ structures in a previously available ~23 Å cryo-ET map of the intact human NPC (*33*). Overall, the top solutions were of lower confidence in the lower resolution ~23 Å cryo-ET map. The top four scoring solutions from global searches in the full map with Nup192•Nic96$^{R2}$•Nup145N$^{R1}$•Nup53$^{R1}$ structures of both conformations identified distal copies in the outer rings and equatorial copies in the inner ring. The novel placement of NUP205 at the proximal cytoplasmic outer ring position came up as the 8th and 9th highest scoring solution for searches in the ~23 Å cryo-ET full map with the single particle cryo-EM and crystal structure conformations of Nup192•Nic96$^{R2}$•Nup145N$^{R1}$•Nup53$^{R1}$, respectively. Top solutions for searches limited to the inner ring of the ~23 Å cryo-ET map identified the same equatorial positions as in the ~12 Å cryo-ET map, albeit with lower confidence. In all cases, the crystal structure conformation of Nup192 resulted in higher confidence solutions than the conformation found in single particle cryo-EM structures. Details about the docking are reported in figs. S56, S57, S59, S60.

*Docking of Nup188•Nic96$^{R2}$•Nup145N$^{R2}$ into the human NPC cryo-ET map.* We performed global searches in the ~12 Å cryo-ET map of the intact human NPC with the single particle cryo-EM Nup188•Nic96$^{R2}$•Nup145N$^{R2}$ structure. The top three scoring solutions corresponded to question mark shaped densities in the outer rings assigned to the NUP205 complex, albeit with lower scores than the corresponding solutions with either conformation of the Nup192•Nic96$^{R2}$•Nup145N$^{R1}$•Nup53$^{R1}$ structure. Thus, the outer ring solutions were discarded. The top two results from global searches in the inner ring, as well as the 4th and 5th top results in the full map, were placed in peripheral question mark-shaped densities in the inner ring. Thus, a total of 16 copies of the NUP188 complex were placed in the human NPC. As a benchmark for quantitative docking in the new ~12 Å cryo-ET map, we replicated the quantitative docking experiments with the novel Nup188•Nic96$^{R2}$•Nup145N$^{R2}$ structure in a previously available ~23 Å cryo-ET map of the intact human NPC (*33*). In contrast with the solutions from docking in the ~12 Å cryo-ET map, the 1st and 3rd highest scoring solutions corresponded to inner ring peripheral placements, whereas the 2nd, 4th and 11th highest scores corresponded to the nuclear distal, cytoplasmic distal, and cytoplasmic proximal question mark-shaped densities of the outer rings. Top solutions for searches limited to the inner ring of the ~23 Å cryo-ET map identified the same peripheral positions as in the ~12 Å cryo-ET map, albeit with lower confidence. Details about the docking are reported in figs. S58 and S63.

*Docking of NUP93$^{SOL}$•NUP53$^{R2}$ into the human NPC cryo-ET map.* We performed global searches with the NUP93$^{SOL}$•NUP53$^{R2}$ crystal structure in the ~12 Å cryo-ET map of the intact human NPC from which densities assigned to the CNC, the NUP205 complex, the NUP188 complex, and NUP358$^{NTD}$ had been subtracted. The top four solutions identified the nuclear outer ring distal, inner ring cytoplasmic peripheral, inner ring nuclear peripheral, and cytoplasmic outer ring distal copies, respectively. Global searches in the subtracted inner ring map identified cytoplasmic peripheral, nuclear peripheral and cytoplasmic equatorial placements as top solutions. A placement at the nuclear equatorial position of the inner ring could be inferred by a C2 symmetry relationship with the top-scoring cytoplasmic equatorial copy (3$^{rd}$ highest scoring) and matched with the 36th highest scoring solution. Lastly, the presence of a proximal NUP205 complex in the cytoplasmic ring suggested the existence of a proximal outer ring copy of NUP93$^{SOL}$•NUP53$^{R2}$, which was readily manually placed into matching but weak density nearby. Thus, a total of 56 copies of the NUP93$^{SOL}$•NUP53$^{R2}$ were placed in the human NPC. As a benchmark for quantitative docking in the new ~12 Å cryo-ET map, we replicated the quantitative docking experiments with the novel NUP93$^{SOL}$•NUP53$^{R2}$ structure in a previously available ~23 Å cryo-ET map of the intact human NPC from which the densities assigned to the CNC, NUP205 complex, NUP188 complex, and NUP358$^{NTD}$ had been subtracted (*33*). The distal placements in both cytoplasmic and nuclear outer rings were identified as the two top scoring solutions, but none of the other placements segregated from the bulk of randomly placed structures. Global searches in the inner ring of the subtracted ~23 Å cryo-ET map identified the previously assigned peripheral and equatorial placements on both cytoplasmic and nuclear sides of the inner ring as the top four scoring solutions. Details about the docking are reported in figs. S64 and S65.

*Docking of Nup170•Nup53$^{R3}$•Nup145N$^{R3}$ into the human NPC cryo-ET map.* Subsequently to subtracting assigned NUP93$^{SOL}$•NUP53$^{R2}$ densities from the hereto subtracted ~12 Å cryo-ET map we performed global searches with three composite crystal Nup170•Nup53$^{R3}$•Nup145N$^{R3}$ structures that differed in the conformation of the Nup170 $\alpha$-helical solenoid, as determined previously (*30*). The top two solutions identified from global searches with conformation II of the Nup170•Nup53$^{R3}$•Nup145N$^{R3}$ composite crystal structure corresponded to the cytoplasmic and nuclear bridge NUP155 complex placements, respectively. Furthermore, the top two solutions identified from global searches with conformation I of the Nup170•Nup53$^{R3}$•Nup145N$^{R3}$ composite crystal structure in the inner ring of the subtracted ~12 Å cryo-ET map identified the nuclear and cytoplasmic equatorial inner ring placements of the NUP155 complex. Finally, as in the previous docking of the ~23 Å cryo-ET map, it was not possible to identify inner ring peripheral placements of the NUP155 complex with global searches due to the imperfect match of the Nup170 conformation as well as the poor local quality of the cryo-ET density (*30*). We placed conformation III of the Nup170•Nup53$^{R3}$•Nup145N$^{R3}$ composite crystal structure at cytoplasmic and nuclear peripheral positions and rigid body fit them locally into the cryo-ET density. Thus, a total of 48 copies of the NUP155 complex was assigned to the human NPC. Details about the docking are reported in figs. S66 and S67.

*Docking of CNT•Nic96$^{R1}$ into the human NPC cryo-ET map.* Subsequently to subtracting assigned NUP155 complex densities from the ~12 Å cryo-ET map we performed global searches in the inner ring with the CNT•Nic96$^{R1}$ crystal structure (PDB ID 5CWS) (*25*). The four inner ring placements were identified by the 2nd, 3rd, 6th, and 9th highest scoring solutions, assigning a total of 32 copies of CNT•NUP93$^{R1}$ per human NPC. Details about the docking are reported in fig. S68.

*Docking of the NUP53$^{RRM}$ homodimer and NUP98$^{APD}$ into the human NPC cryo-ET map.*

Subsequently to subtracting assigned CNT•NUP93$^{R1}$ complex densities from the hereto subtracted ~12 Å cryo-ET map we performed global searches with the NUP53$^{RRM}$ homodimer (PDB ID 4LIR) and NUP98$^{APD}$ crystal structures (PDB ID 1KO6) (*11*). Top solutions from global searches with either structure identified globular densities between spokes of the inner ring that are related by a C2 symmetry operator about the midplane of the inner ring. For the NUP53$^{RRM}$ homodimer, the top ten scoring solutions identified the same cytoplasmic inner ring placement in slightly different orientations due to the lack of discriminating shape features in the small protein complex. The 11th highest scoring solution identified the symmetry-related placement on the nuclear side of the inner ring. Likewise, the top ten scoring solutions for the docking of NUP98$^{APD}$ identified the same cytoplasmic inner ring placement in slightly different orientations, whereas the 8th top scoring solution identified the symmetry-related placement on the nuclear side of the inner ring. The globular densities found by the searches appeared to be discontinuous with the rest of the cryo-ET map, suggesting that they are held in place by unresolved tethers, such as flexible linker sequences. Unlike NUP98$^{APD}$, the NUP53$^{RRM}$ homodimer could be held in place by opposing tethers from adjacent spokes and withstand cancellation of its signal by sub-tomogram averaging. Thus, we assigned 32 copies of NUP53$^{RRM}$ to the inner ring of the human NPC. Details about the docking are reported in fig. S70.

*Docking into the dilated in situ cryo-ET map of the human NPC.* A composite structure of a spoke of the cytoplasmic outer ring built into the constricted ~12 Å cryo-ET sub-tomogram averaged human NPC map (EMD-XXXX) (*41*) was manually placed in the dilated *in situ* ~37 Å cryo-ET sub-tomogram averaged human NPC map (EMD-11967) (*40*). The structure fit into the dilated human NPC ~37 Å cryo-ET map was further locally refined as two rigid bodies composed of all distal and all proximal subunits. A C8 symmetry operation was applied to place the docked spoke to the rest of the dilated human NPC ~37 Å cryo-ET map. Details about the docking are reported in fig. S78.

*Docking into an anisotropic ~8 Å X. laevis cytoplasmic face protomer map.* The anisotropic ~8 Å composite single particle cryo-EM map of the *X. laevis* cytoplasmic outer ring protomer (EMD-0909) (*64*) was superposed to the ~12 Å cryo-ET map of the intact human NPC (EMD-XXXX) (*41*). The placement of individual structures was further refined by local rigid-body fitting into *X. laevis* map. The map did not include regions corresponding to the proximal copy of NUP93$^{SOL}$•NUP53$^{R2}$, presumably due to omission by the masking applied during map processing. Details about the docking are reported in fig. S71.

***Figures and movies***

Gel filtration profiles and MALS graphs were generated in IGOR (WaveMetrics) and assembled in Adobe Illustrator. Structure figures were created using PyMol (www.pymol.org) and UCSF Chimera (*109*). Quantitative map fitting histograms were generated with the Python seaborn library (*125*).

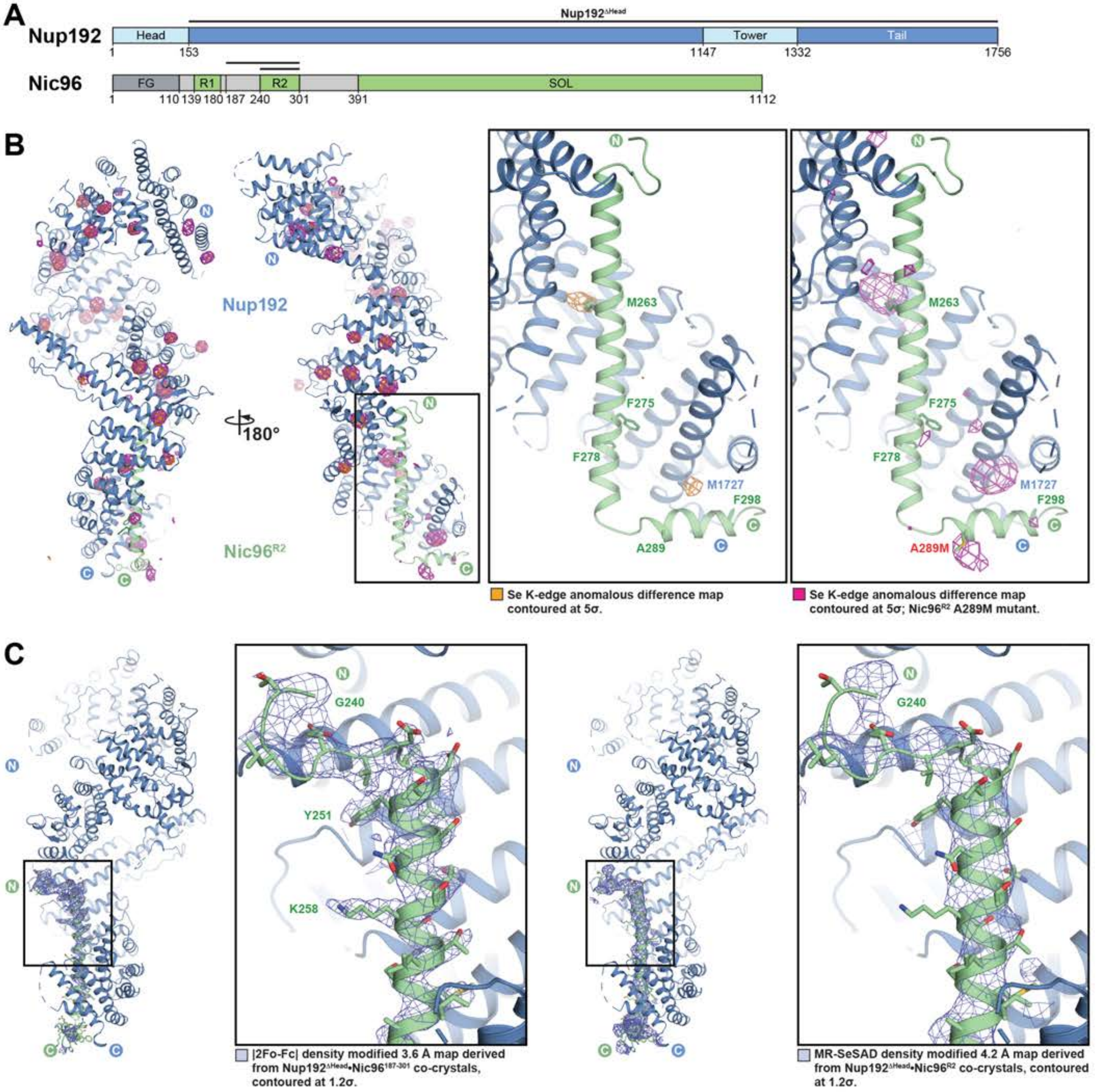

A
Nup192
Nup192ΔHead
Head
Tower
Tail
1
153
1147
1332
1756
Nic96
FG
R1
R2
SOL
1
110
139
180
187
240
301
391
1112
B
Nup192
Nic96R2
180°
M263
F275
F278
M1727
F298
A289
A289M
Se K-edge anomalous difference map contoured at 5σ.
Se K-edge anomalous difference map contoured at 5σ; Nic96R2 A289M mutant.
C
G240
Y251
K258
|2Fo-Fc| density modified 3.6 Å map derived from Nup192ΔHead•Nic96187-301 co-crystals, contoured at 1.2σ.
MR-SeSAD density modified 4.2 Å map derived from Nup192ΔHead•Nic96R2 co-crystals, contoured at 1.2σ.

Supp. Fig. S1

**Fig.S1. Crystallographic analysis of Nup192$^{\Delta HEAD}$-bound Nic96 fragments.** (**A**) Domain structures of Nup192 and Nic96. Black lines indicate the co-crystallized fragments. (**B**) Isomesh representation of anomalous difference Fourier maps calculated from X-ray diffraction data collected at the selenium anomalous peak wavelengths for Nup192$^{\Delta Head}$•Nic96$^{R2}$ (orange) and Nup192$^{\Delta Head}$•Nic96$^{R2}$ A289M (magenta) SeMet-derivatized co-crystals and superposed with the cartoon representation of the Nup192$^{\Delta Head}$•Nic96$^{187\text{-}301}$ structure, validating the sequence register of the Nic96$^{R2}$ peptide. (**C**) Comparison of electron density maps derived from Nup192$^{\Delta Head}$•Nic96$^{187\text{-}301}$ and Nup192$^{\Delta Head}$•Nic96$^{R2}$ co-crystals demonstrating that Nic96 residues 187-239 are not resolved in the electron density.

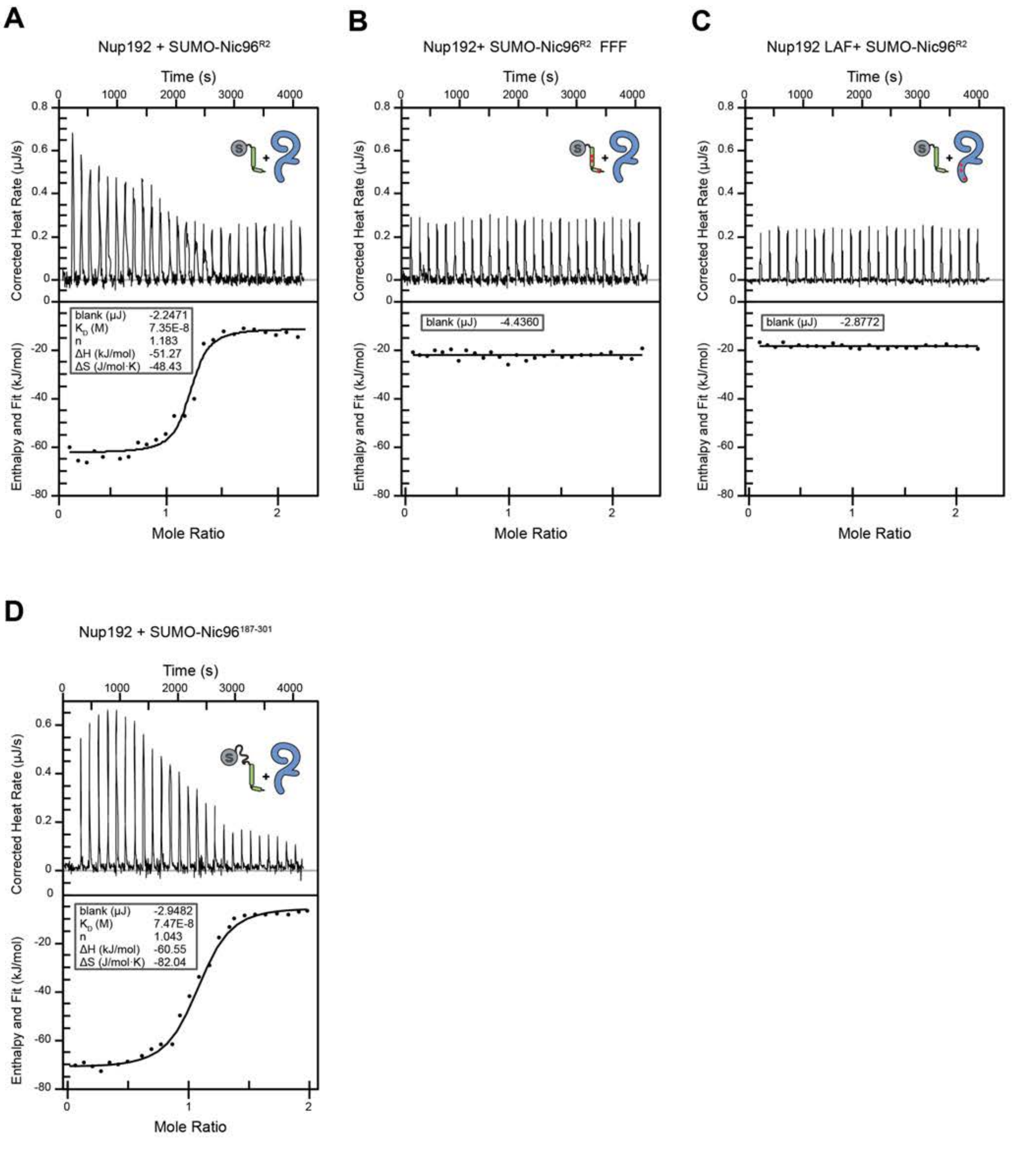

Supp. Fig. S2

Hoelz et al., 2021

**Fig.S2. ITC analysis of the Nup192-Nic96$^{R2}$ interaction.** Representative baseline-corrected ITC experiments of (**A**) wildtype SUMO-Nic96$^{R2}$ titrated against wildtype Nup192, (**B**) SUMO-Nic96$^{R2}$ FFF mutant titrated against wildtype Nup192, (**C**) wildtype SUMO-Nic96$^{R2}$ titrated against Nup192 LAF mutant, and (**D**) wildtype SUMO-Nic96$^{187\text{-}301}$ titrated against wildtype Nup192. The least-squares fit thermodynamic parameters for a single binding site model are shown in the inset boxes. Overall ITC experimental conditions and results of experiments conducted in triplicate are reported in Table S5.

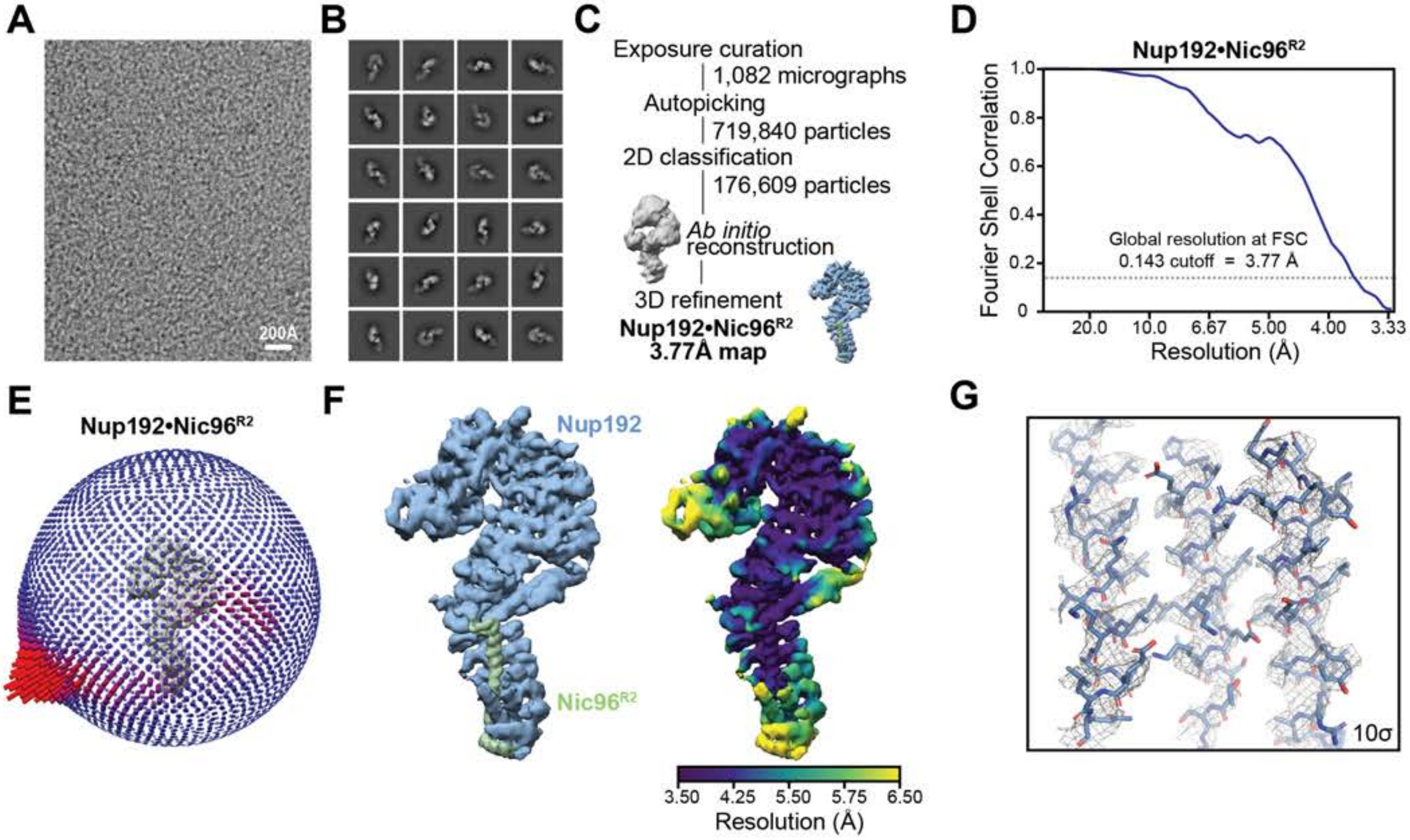
A
200Å
B
C
Exposure curation
1,082 micrographs
Autopicking
719,840 particles
2D classification
176,609 particles
Ab initio reconstruction
3D refinement
Nup192•Nic96R2
3.77Å map
D
Nup192•Nic96R2
Fourier Shell Correlation
Global resolution at FSC 0.143 cutoff = 3.77 Å
Resolution (Å)
E
Nup192•Nic96R2
F
Nup192
Nic96R2
3.50 4.25 5.50 5.75 6.50
Resolution (Å)
G
10σ

 

**Fig.S3. Single particle cryo-EM analysis of the Nup192•Nic96$^{R2}$ complex.** (**A**) Representative motion corrected cryo-electron micrograph. (**B**) 2D class averages representative of different particle orientations. (**C**) Summary of single particle data processing workflow with indicated number of micrographs or particles involved at each step. (**D**) Gold-standard Fourier Shell Correlation curve estimated with 3DFSC and global resolution of 3.8 Å at the 0.143 cutoff. (**E**) Three-dimensional angular distribution plot of particles. (**F**) Isosurface representation of the Nup192•Nic96$^{R2}$ cryo-EM density (EMD-24056) colored according to protein chain identity (*left*) or local resolution estimation (*right*). (**G**) Isomesh representation of representative sharpened cryo-electron microscopy (cryo-EM) density contoured at a 10 $\sigma$ cutoff level and ball-and-stick representation of corresponding atomic coordinates.

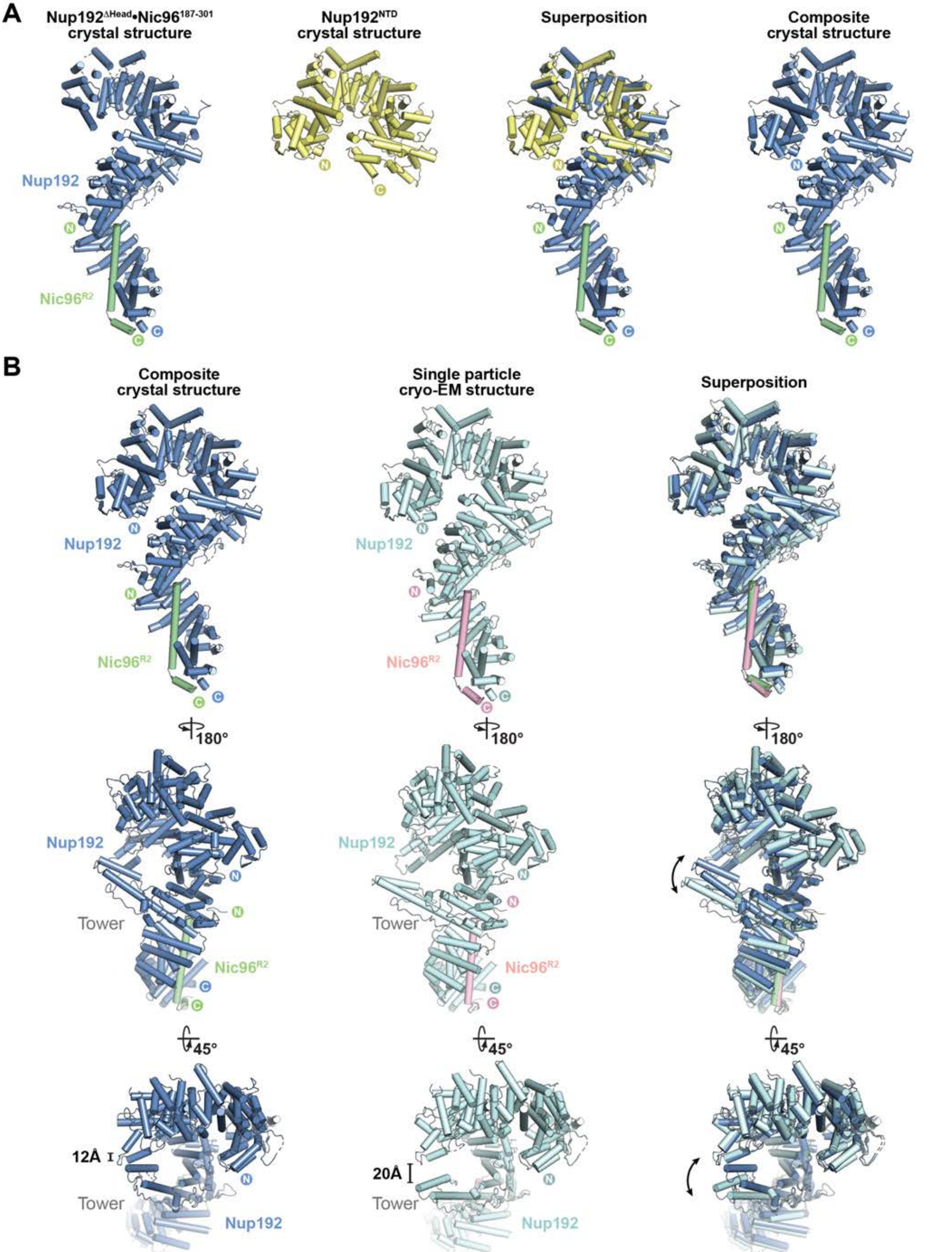

Supp. Fig. S4

Hoelz et al., 2021

**Fig.S4. Comparison of crystal composite and single particle cryo-EM structures of Nup192•Nic96$^{R2}$.** (**A**) Cartoon representation of the crystal structures of Nup192$^{\Delta Head}$•Nic96$^{187\text{-}301}$ (blue and pale green) and the previously solved Nup192$^{NTD}$ structure (yellow, PDB ID 4KNH) (*36*), and their superposition are shown. The Nup192•Nic96$^{R2}$ composite crystal structure was generated by aligning and merging residues in common between the two structures. (**B**) Cartoon representation of the Nup192•Nic96$^{R2}$ composite crystal structure (blue and pale green) and the 3.8 Å Nup192•Nic96$^{R2}$ single particle cryo-EM structure (light cyan and pink), shown from three points of view to illustrate the conformational difference between the two structures. Curved arrows indicate the displacement of the Tower domain induced by the conformational change.

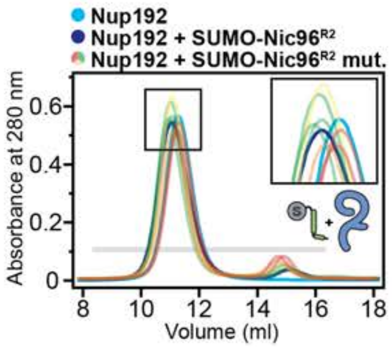

**Nic96 single-residue mutational analysis**

| Nic96^R2 mutation | Nup192 binding |
|---|---|
| wild type | +++ |
| F275A | +++ |
| F278A | +++ |
| F298A | ++ |
| F275A/F278A/F298A | - |
| F275E/F278E | +++ |
| F298E | + |
| F275E/F278E/F298E (FFF) | - |

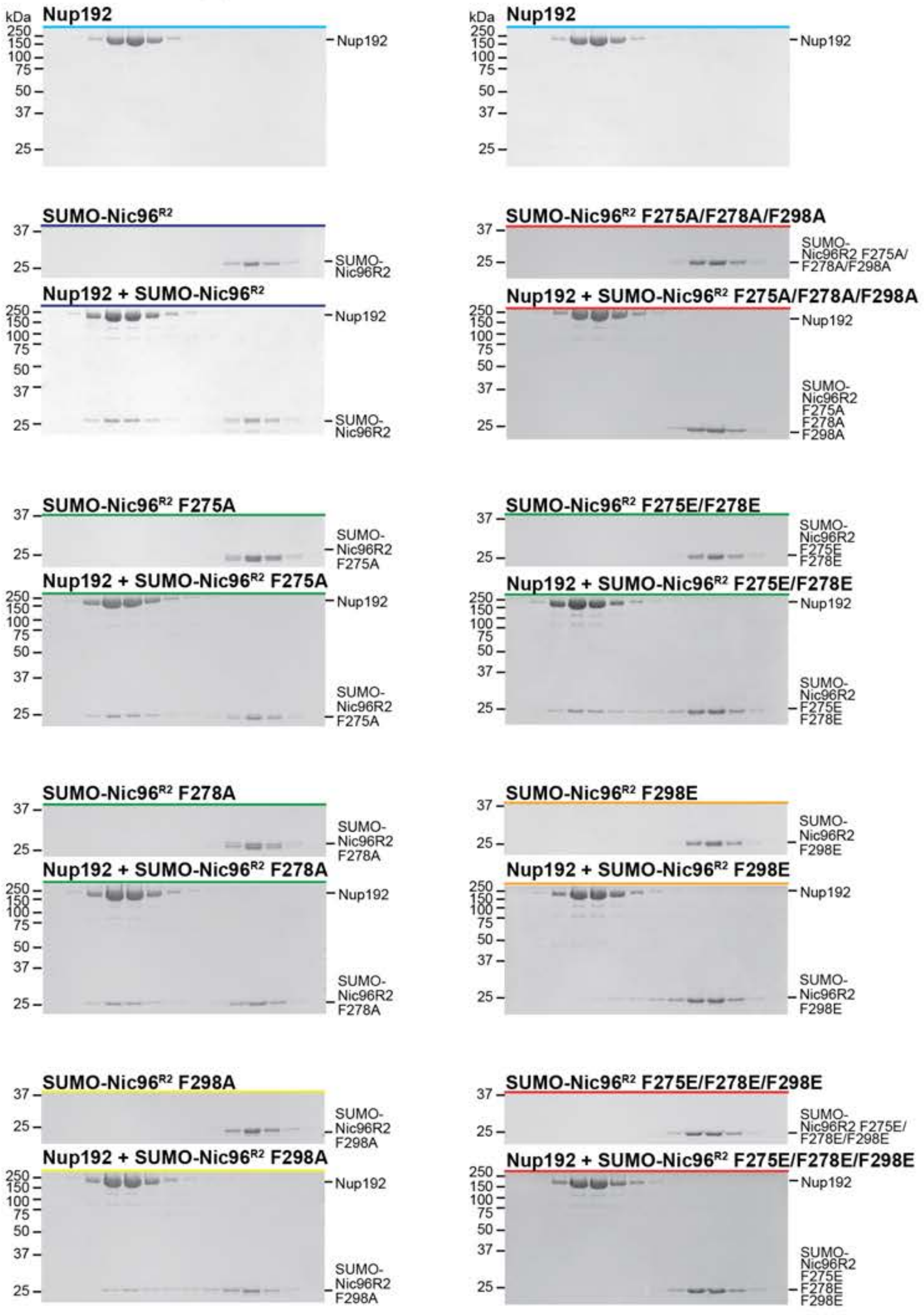

 

**Fig.S5. Perturbation of the Nup192-Nic96$^{R2}$ interaction by Nic96 point mutants.** SEC and SDS-PAGE analysis corresponding to Fig. 2C. SEC profiles of wildtype Nup192 alone (cyan) and Nup192 preincubated with wildtype SUMO-Nic96$^{R2}$ (dark blue) are shown. SEC profiles of SUMO-Nic96$^{R2}$ mutants preincubated with Nup192 are colored according to the measured effect: no effect (green, +++), weak effect (yellow, ++), moderate effect (orange, +), and abolished binding (red, -). The inset box shows a closeup view of elution profile shifts at the peak fractions. The gray bar indicates fractions that were resolved on SDS-PAGE gels and visualized by Coomassie staining. All SEC profiles were obtained using a Superdex 200 10/300 GL column. The results are summarized in the table (*top right*).

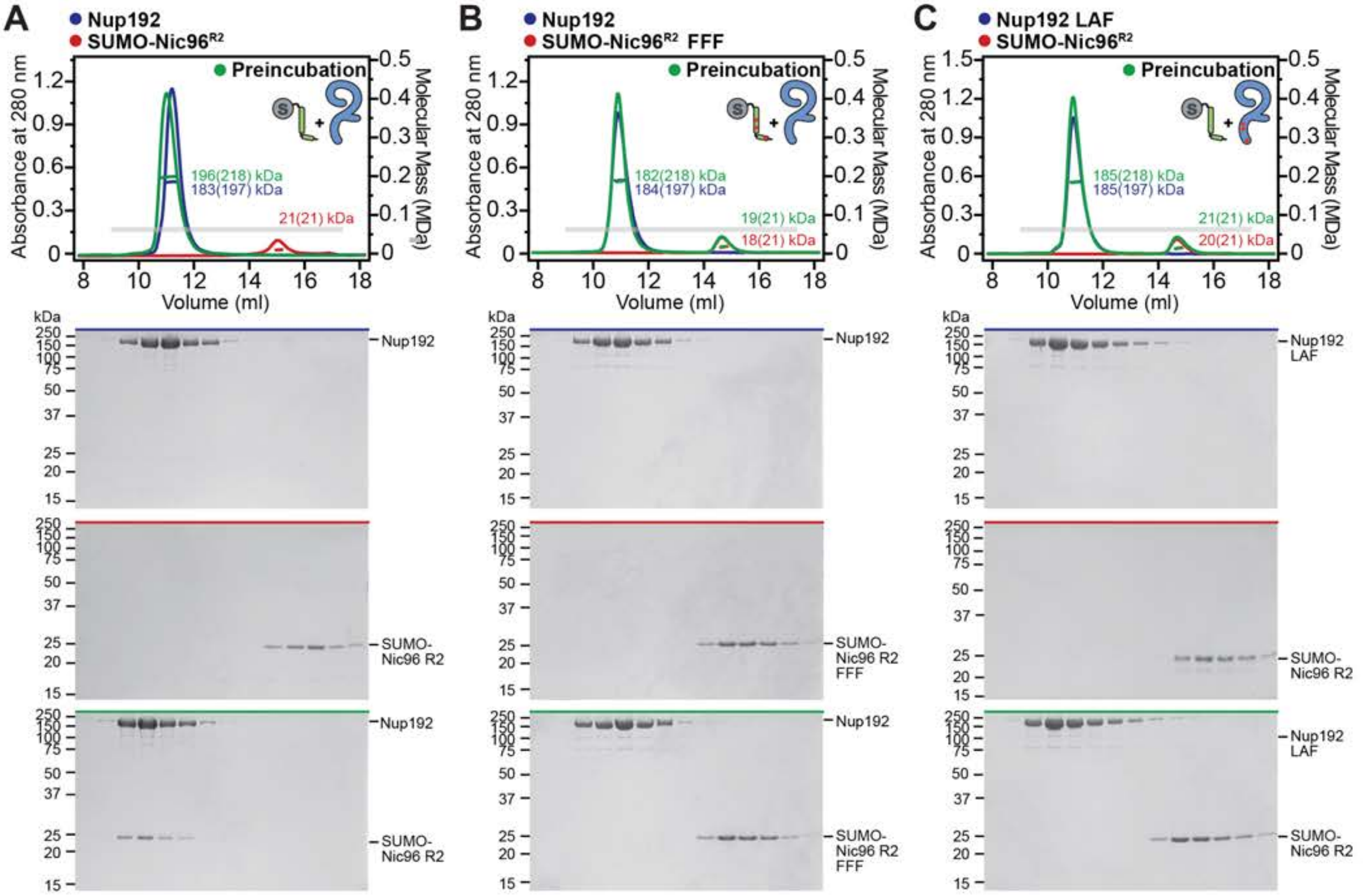

A
Nup192
SUMO-Nic96R2
Preincubation
196(218) kDa
183(197) kDa
21(21) kDa
Absorbance at 280 nm
Molecular Mass (MDa)
Volume (ml)
Nup192
SUMO-Nic96 R2
B
Nup192
SUMO-Nic96R2 FFF
Preincubation
182(218) kDa
184(197) kDa
19(21) kDa
18(21) kDa
SUMO-Nic96 R2 FFF
C
Nup192 LAF
SUMO-Nic96R2
185(218) kDa
185(197) kDa
21(21) kDa
20(21) kDa
Nup192 LAF

 

**Fig.S6. SEC-MALS analysis of the Nup192-Nic96$^{R2}$ interaction.** SEC-MALS and SDS-PAGE analysis corresponding to Fig. 2F. SEC-MALS profiles of nups are shown individually (red and blue) and after their preincubation (green) for: (**A**) wildtype SUMO-Nic96$^{R2}$ binding to wildtype Nup192, (**B**) the SUMO-Nic96$^{R2}$ FFF mutant binding to wildtype Nup192, (**C**) wildtype SUMO-Nic96$^{R2}$ binding to the Nup192 LAF mutant. Measured molecular masses are indicated for the peak fractions and corresponding theoretical molecular masses are reported in parenthesis. Gray bars indicate fractions that were resolved on SDS-PAGE gels and visualized by Coomassie staining. All SEC profiles were obtained using a Superdex 200 Increase 10/300 GL column. All SEC-MALS results are summarized in Table S4.

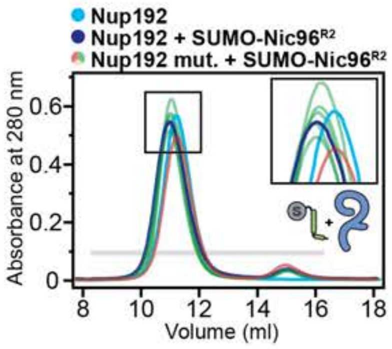

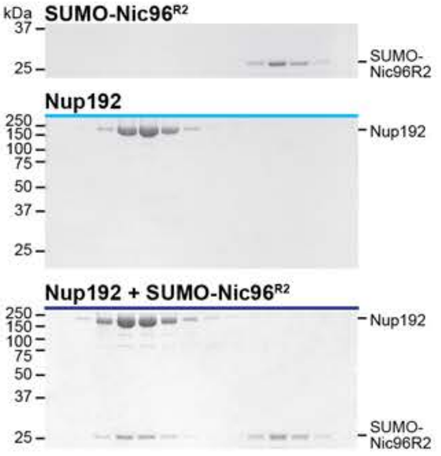

**Nup192 single-residue mutational analysis**

| Nup192 mutation | Nic96R2 binding |
|---|---|
| wild-type | +++ ● |
| L1584A | +++ ● |
| F1735A | +++ ● |
| L1584A/F1735A | +++ ● |
| L1584E/A1648E | +++ ● |
| F1735E | +++ ● |
| L1584E/A1648E/F1735E (LAF) | - ● |

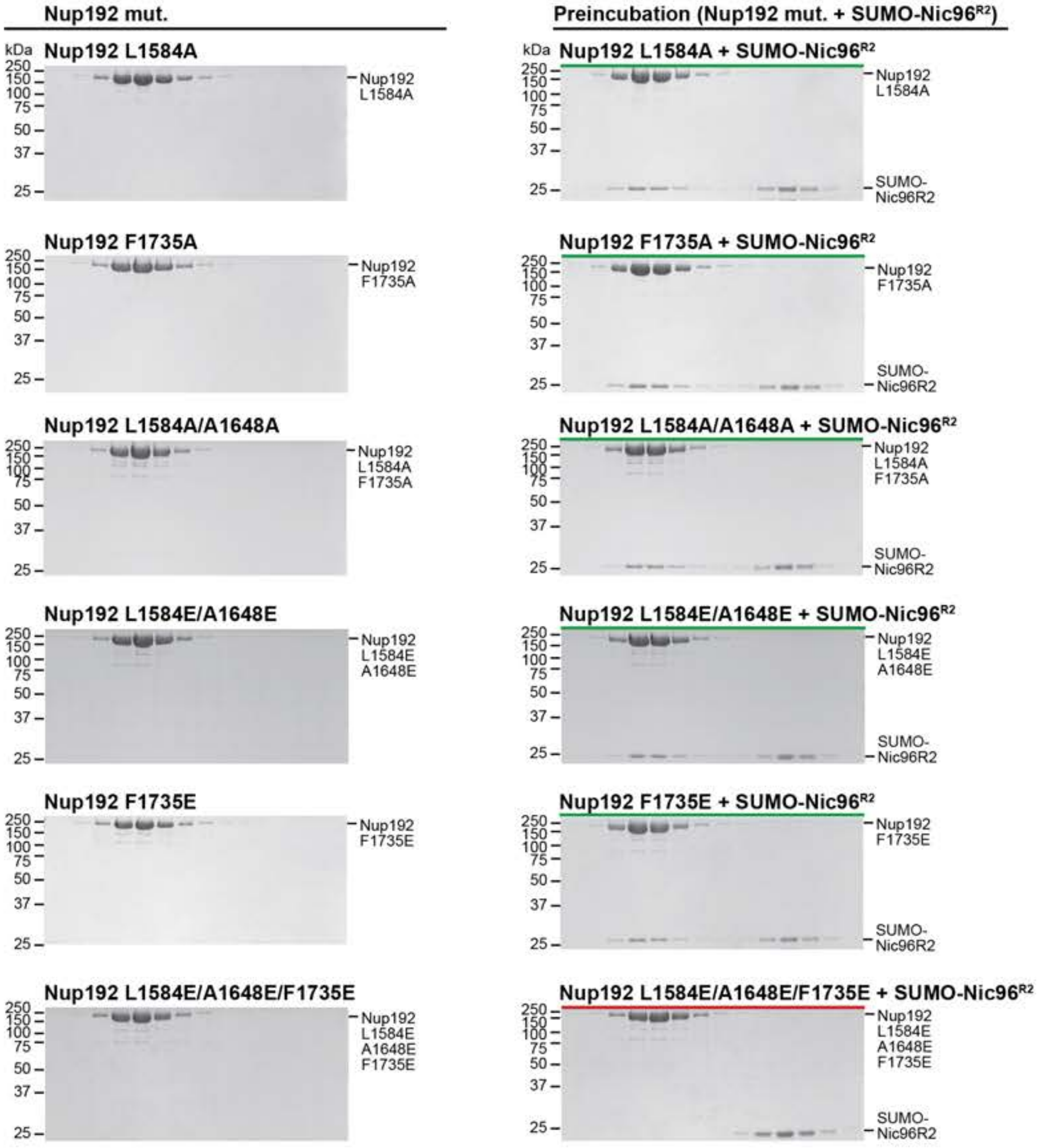

**Fig.S7. Perturbation of the Nup192-Nic96$^{R2}$ interaction by Nup192 point mutants.** SEC and SDS-PAGE analysis corresponding to Fig. 2D. SEC profiles of wildtype Nup192 alone (cyan) and Nup192 preincubated with wildtype SUMO-Nic96$^{R2}$ (dark blue) are shown. SEC profiles of Nup192 mutants preincubated with SUMO-Nic96$^{R2}$ are colored according to the measured effect: no effect (green, +++) and abolished binding (red, -). The inset box shows a closeup view of elution profile shifts at the peak fractions. The gray bar indicates fractions that were resolved on SDS-PAGE gels and visualized by Coomassie staining. All SEC profiles were obtained using a Superdex 200 10/300 GL column. The results are summarized in the table (*top left*).

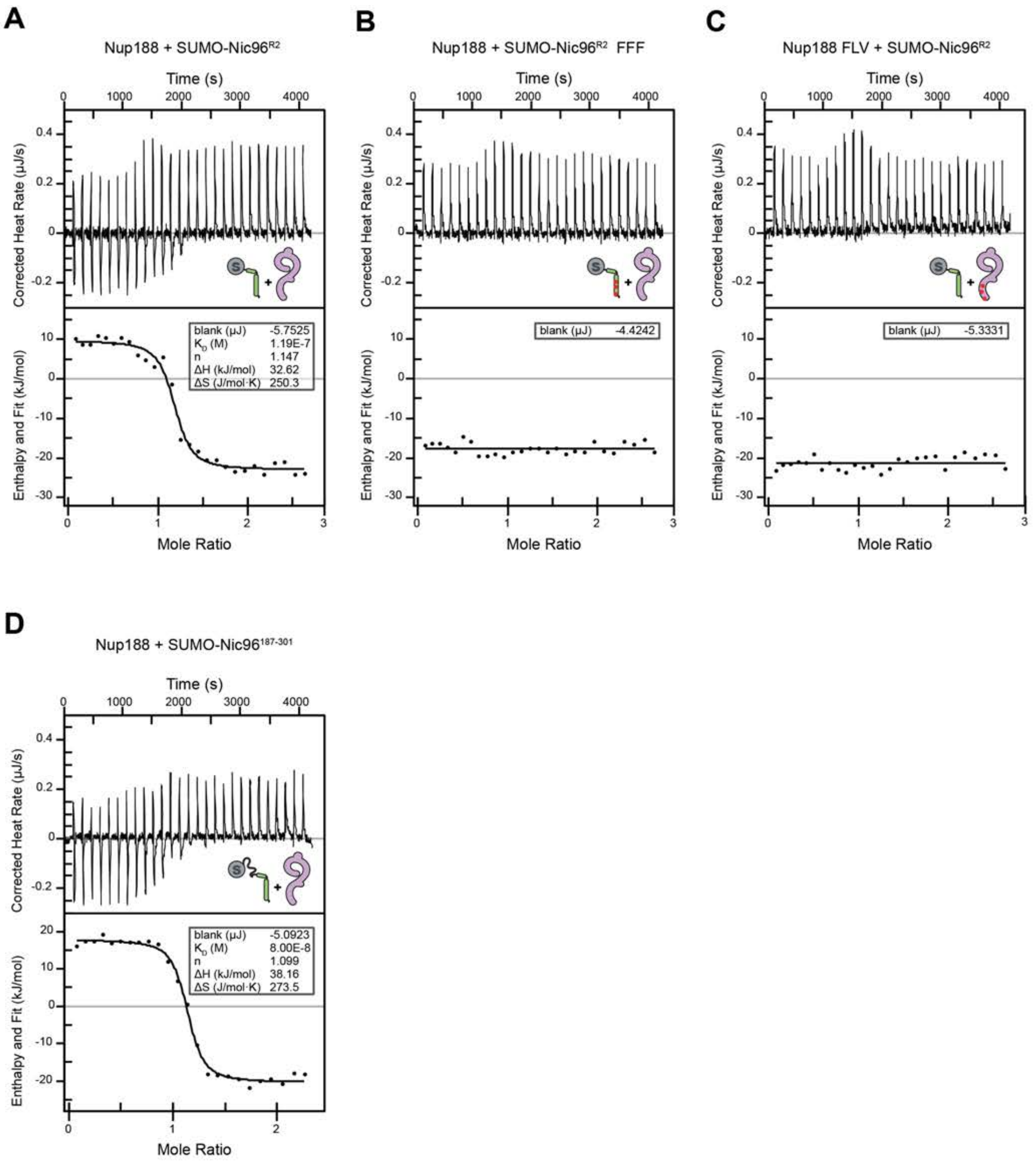

A
Nup188 + SUMO-Nic96R2
Time (s)
Corrected Heat Rate (µJ/s)
Enthalpy and Fit (kJ/mol)
Mole Ratio
blank (µJ) -5.7525
KD (M) 1.19E-7
n 1.147
ΔH (kJ/mol) 32.62
ΔS (J/mol·K) 250.3
B
Nup188 + SUMO-Nic96R2 FFF
blank (µJ) -4.4242
C
Nup188 FLV + SUMO-Nic96R2
blank (µJ) -5.3331
D
Nup188 + SUMO-Nic96187-301
blank (µJ) -5.0923
KD (M) 8.00E-8
n 1.099
ΔH (kJ/mol) 38.16
ΔS (J/mol·K) 273.5

Supp. Fig. S8

**Fig.S8. ITC analysis of the Nup188-Nic96$^{R2}$ interaction.** Representative baseline-corrected ITC experiments for (**A**) wildtype SUMO-Nic96$^{R2}$ titrated against wildtype Nup188, (**B**) SUMO-Nic96$^{R2}$ FFF mutant titrated against wildtype Nup188, (**C**) wildtype SUMO-Nic96$^{R2}$ titrated against Nup188 FLV mutant, and (**D**) wildtype SUMO-Nic96$^{187-301}$ titrated against wildtype Nup188. The least-squares fit thermodynamic parameters for a single binding site model are shown in the inset boxes. Overall ITC experimental conditions and results of experiments conducted in triplicate are reported in Table S5.

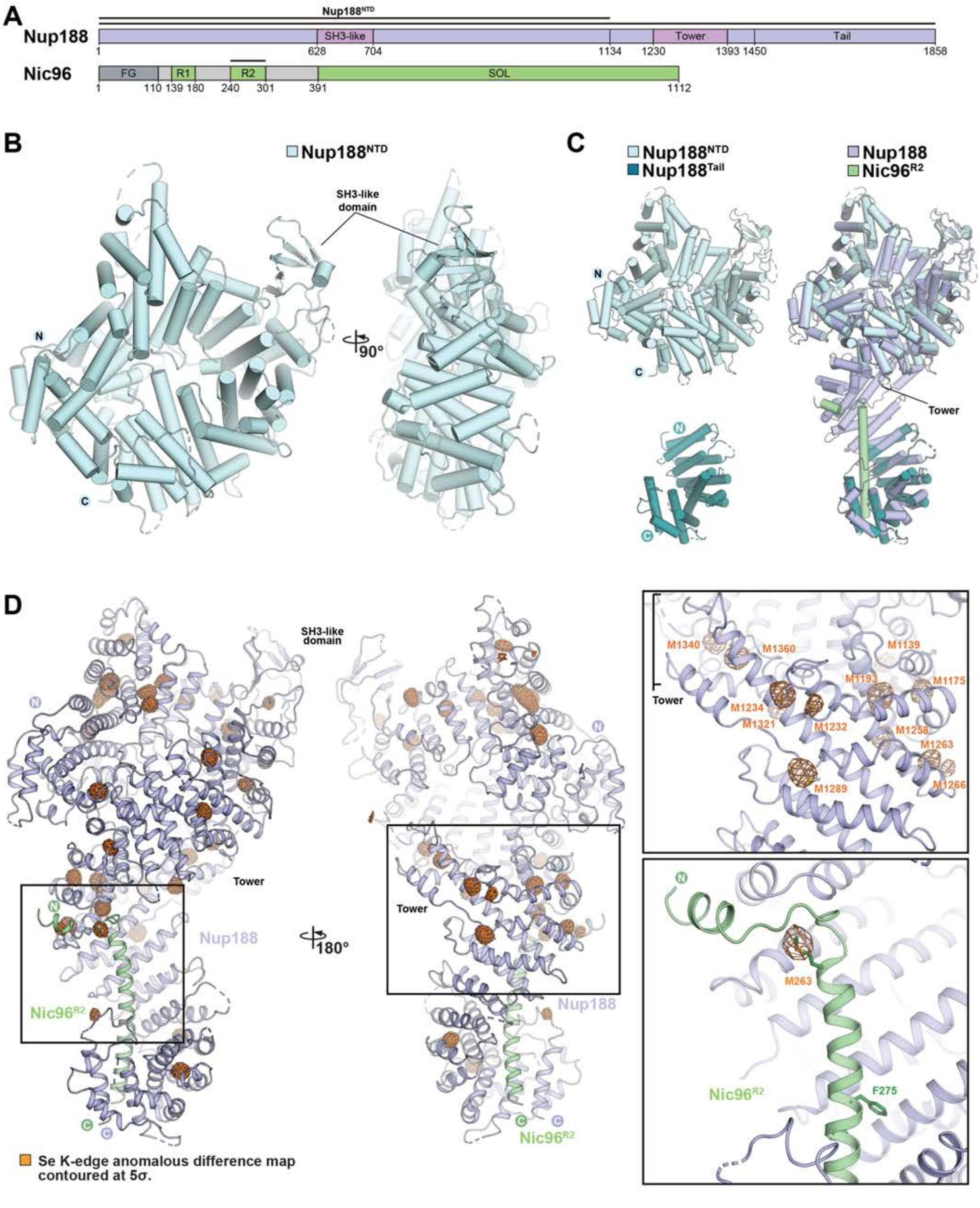

Supp. Fig. S9

Hoelz et al., 2021

**Fig.S9. Crystallographic analysis of the Nup188•Nic96$^{R2}$ complex.** (**A**) Domain structures of Nup188 and Nic96. Black lines indicate the fragments used for crystallization. (**B**) Cartoon representation of the 2.8 Å crystal structure of Nup188$^{NTD}$. (**C**) Cartoon representations of the 2.8 Å Nup188$^{NTD}$ (light cyan) and previously solved 3.4 Å Nup188$^{TAIL}$ (teal; PDB ID 5CWU) (*25*) crystal structures (*left*) used as starting models for the 4.4 Å Nup188•Nic96$^{R2}$ crystal structure (light purple and pale green), and their superposition (*right*). (**D**) Isomesh representation of anomalous difference Fourier map (orange) calculated from X-ray diffraction data collected at the selenium anomalous peak wavelengths for a Nup188•Nic96$^{R2}$ seleno-L-methionine (SeMet)-derivatized co-crystal and superposed with the cartoon representation of the Nup188•Nic96$^{R2}$ structure, validating the sequence register of Nic96$^{R2}$ and the previously unmodeled Nup188 region.

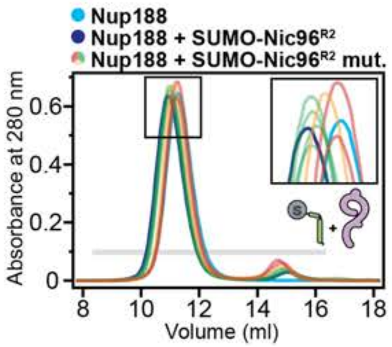

**Nic96 single-residue mutational analysis**

| Nic96R2 mutation | Nup188 binding |
|---|---|
| wild type | +++ (blue) |
| F275A | +++ (green) |
| F278A | +++ (green) |
| F298A | +++ (green) |
| F275A/F278A/F298A | - (red) |
| F275E/F278E | + (orange) |
| F298E | +++ (green) |
| F275E/F278E/F298E (FFF) | - (red) |

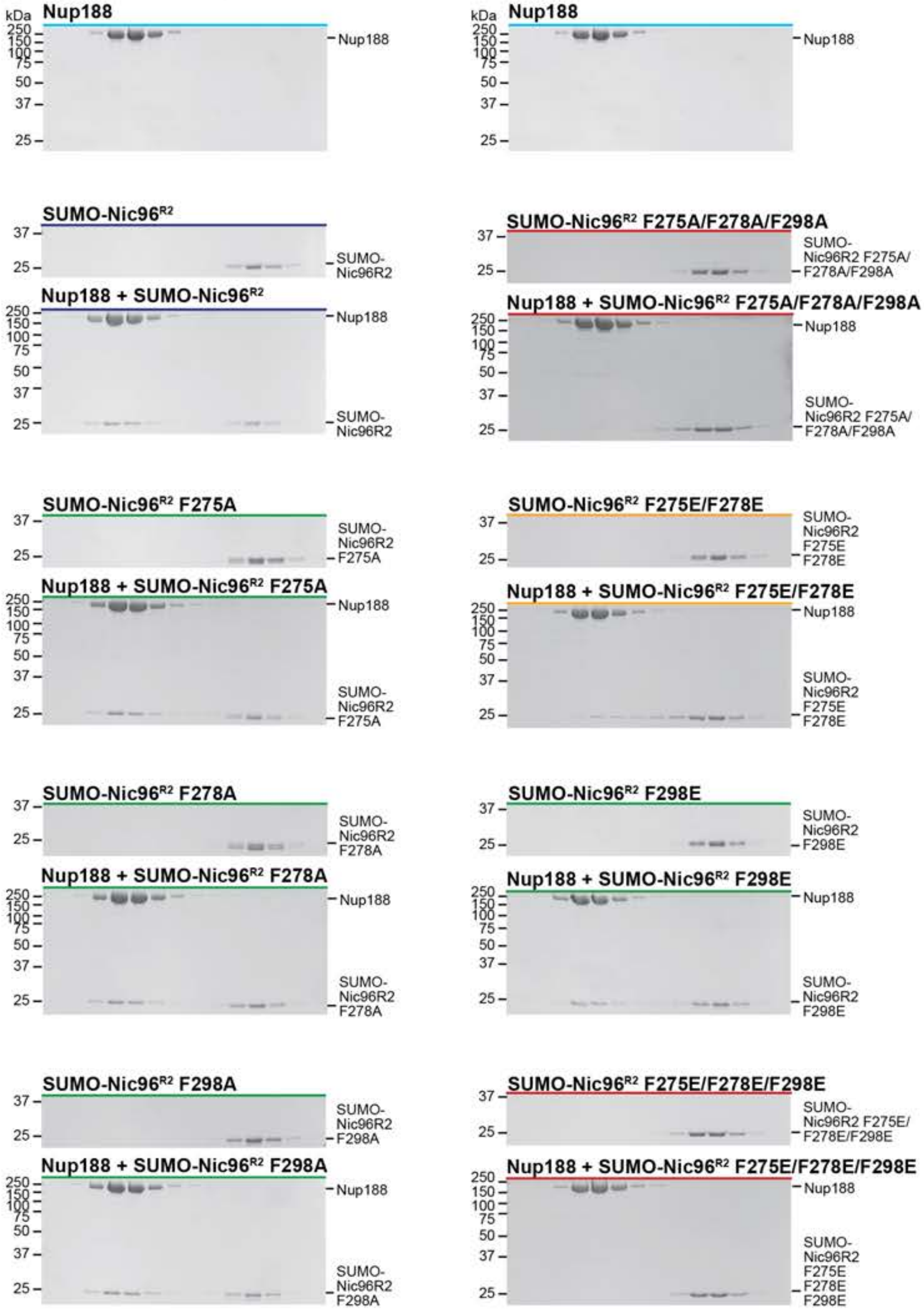

**Fig.S10. Perturbation of the Nup188-Nic96$^{R2}$ interaction by Nic96 point mutants.** SEC and SDS-PAGE analysis corresponding to Fig. 3C. SEC profiles of wildtype Nup188 alone (cyan) and Nup188 preincubated with wildtype SUMO-Nic96$^{R2}$ (dark blue) are shown. SEC profiles of SUMO-Nic96$^{R2}$ mutants preincubated with Nup188 are colored according to the measured effect: no effect (green, +++), moderate effect (orange, +), and abolished binding (red, -). The inset box shows a closeup view of elution profile shifts at the peak fractions. The gray bar indicates fractions that were resolved on SDS-PAGE gels and visualized by Coomassie staining. All SEC profiles were obtained using a Superdex 200 10/300 GL column. The results are summarized in the table (*top right*).

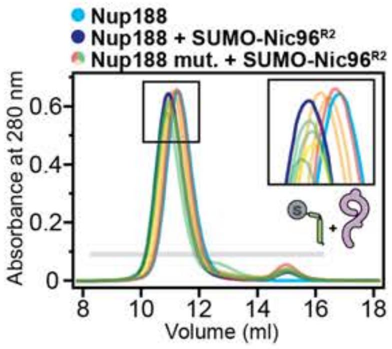

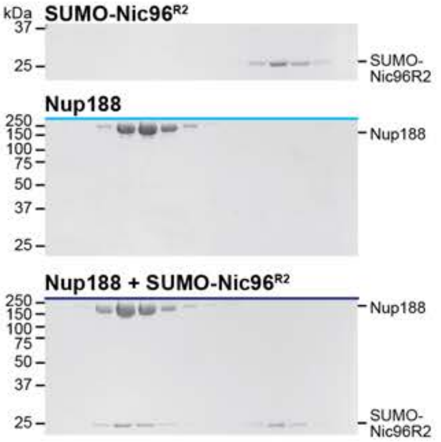

**Nup188 single-residue mutational analysis**

| Nup188 mutation | Nic96R2 binding |
|---|---|
| wild-type | +++ |
| F1478A | ++ |
| L1689A | +++ |
| V1760A | +++ |
| F1478A/L1689A/V1760A | + |
| F1478E | + |
| L1689E/V1760E | +++ |
| F1478E/L1689E/V1760E (FLV) | - |

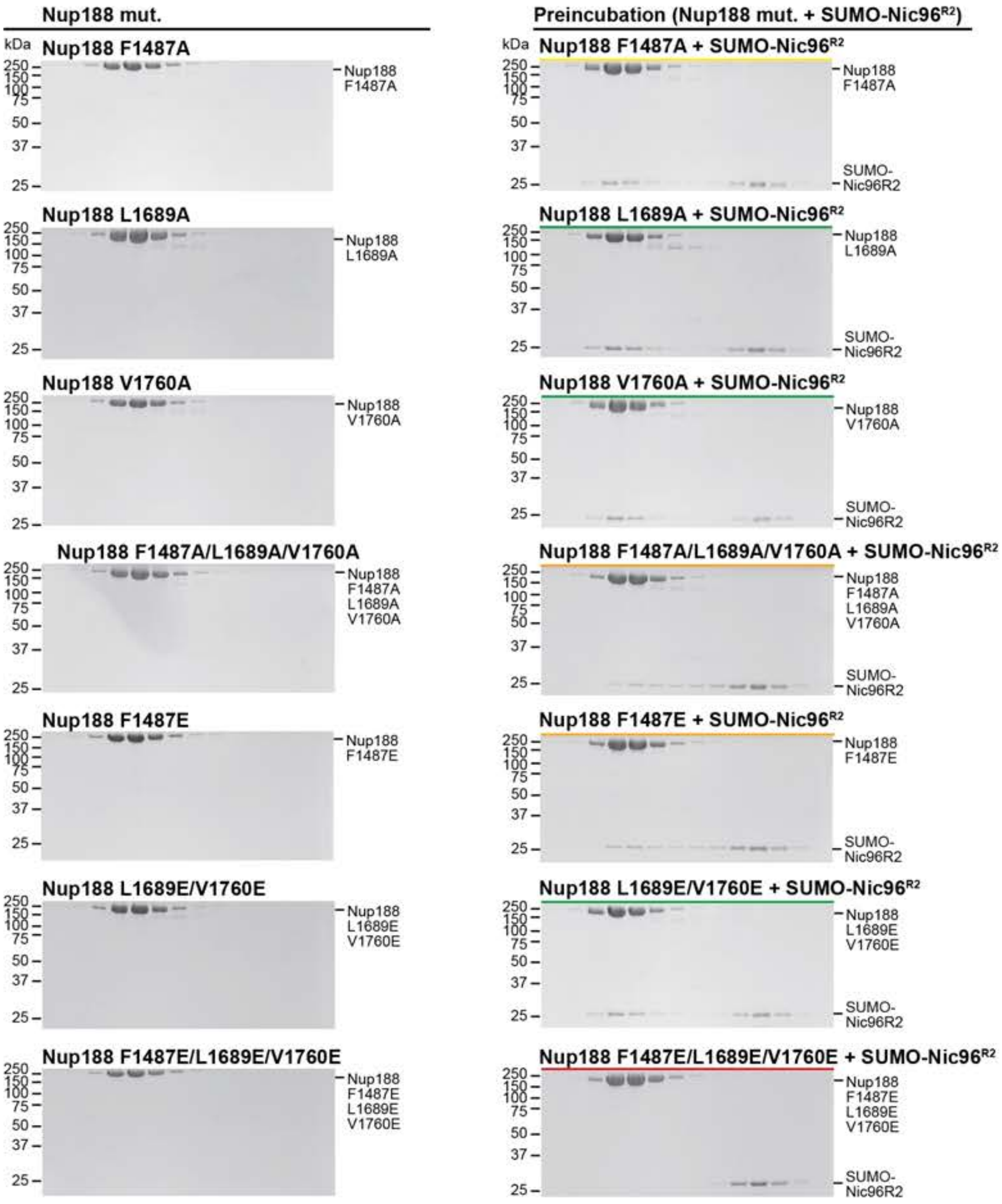

**Fig.S11. Perturbation of the Nup188-Nic96$^{R2}$ interaction by Nup188 point mutants.** SEC and SDS-PAGE analysis corresponding to Fig. 3D. SEC profiles of wildtype Nup188 alone (cyan) and Nup188 preincubated with wildtype SUMO-Nic96$^{R2}$ (dark blue) are shown. SEC profiles of Nup188 mutants preincubated with SUMO-Nic96$^{R2}$ are colored according to the measured effect: no effect (green, +++), weak effect (yellow, ++), moderate effect (orange, +), and abolished binding (red, -). The inset box shows a closeup view of elution profile shifts at the peak fractions. The gray bar indicates fractions that were resolved on SDS-PAGE gels and visualized by Coomassie staining. All SEC profiles were obtained using a Superdex 200 10/300 GL column. The results are summarized in the table (*top left*).

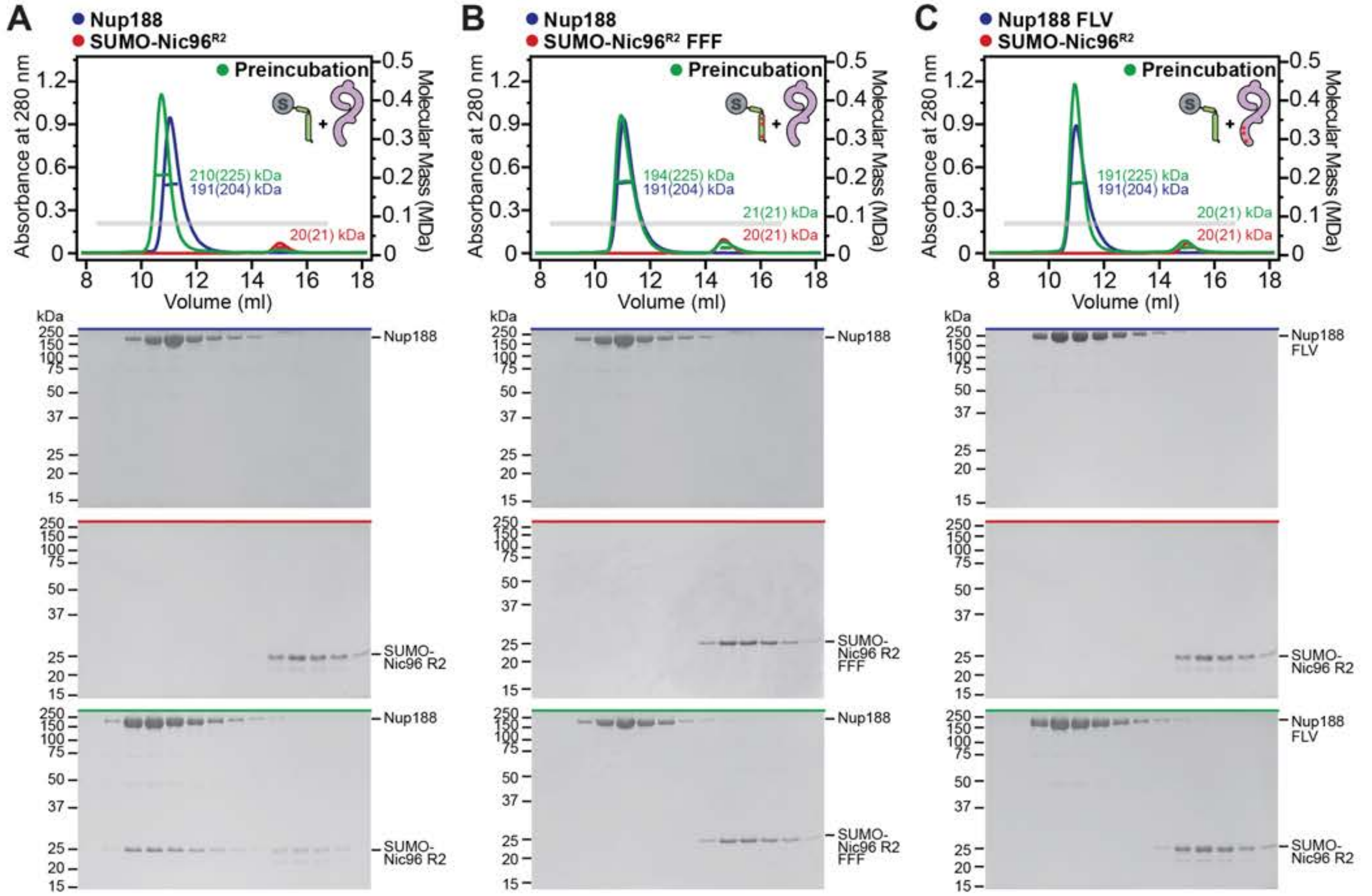

A
Nup188
SUMO-Nic96R2
Preincubation
Absorbance at 280 nm
Molecular Mass (MDa)
Volume (ml)
210(225) kDa
191(204) kDa
20(21) kDa
B
Nup188
SUMO-Nic96R2 FFF
Preincubation
194(225) kDa
191(204) kDa
21(21) kDa
20(21) kDa
C
Nup188 FLV
SUMO-Nic96R2
Preincubation
191(225) kDa
191(204) kDa
20(21) kDa
20(21) kDa
kDa
Nup188
Nup188 FLV
SUMO-Nic96 R2
SUMO-Nic96 R2 FFF

Supp. Fig. S12

**Fig.S12. SEC-MALS analysis of the Nup188-Nic96$^{R2}$ interaction.** SEC-MALS and SDS-PAGE analysis corresponding to Fig. 3F. SEC-MALS profiles of nups are shown individually (red and blue) and after their preincubation (green) for: (**A**) wildtype SUMO-Nic96$^{R2}$ binding to wildtype Nup188, (**B**) the SUMO-Nic96$^{R2}$ FFF mutant binding to wildtype Nup188, (**C**) wildtype SUMO-Nic96$^{R2}$ binding to the Nup188 FLV mutant. Measured molecular masses are indicated for the peak fractions and corresponding theoretical molecular masses are reported in parenthesis. Gray bars indicate fractions that were resolved on SDS-PAGE gels and visualized by Coomassie staining. All SEC profiles were obtained using a Superdex 200 Increase 10/300 GL column. All SEC-MALS results are summarized in Table S4.

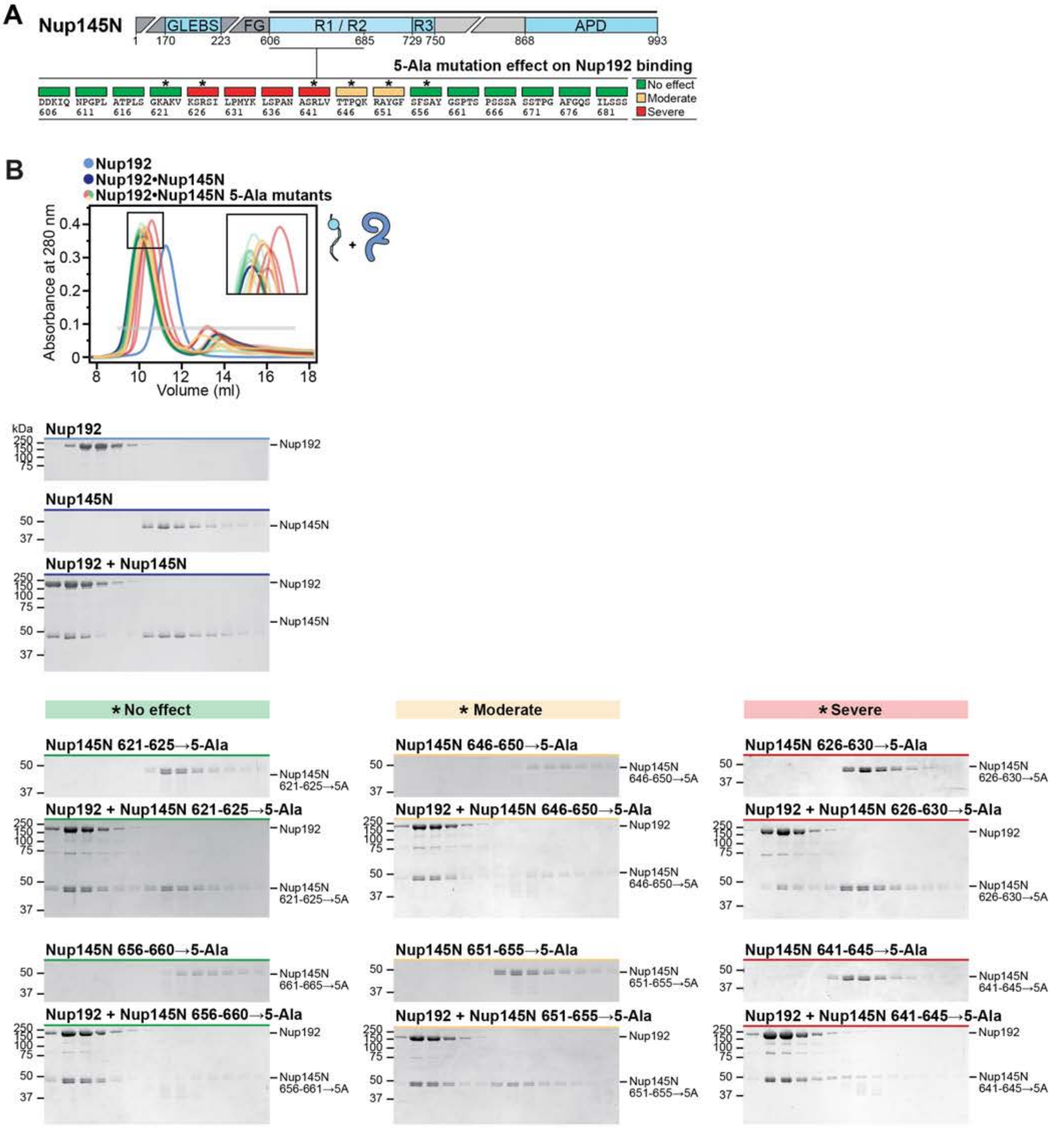

A
Nup145N
GLEBS
FG
R1 / R2
R3
APD
1 170 223 606 685 729 750 868 993
5-Ala mutation effect on Nup192 binding
DDKIQ 606
NPGPL 611
ATPLS 616
GKAKV 621
KSRSI 626
LPMYK 631
LSPAN 636
ASRLV 641
TTPQK 646
RAYGF 651
SFSAY 656
GSPTS 661
PSSSA 666
SSTPG 671
AFGQS 676
ILSSS 681
No effect
Moderate
Severe
B
Nup192
Nup192•Nup145N
Nup192•Nup145N 5-Ala mutants
Absorbance at 280 nm
Volume (ml)
kDa
Nup192
Nup145N
Nup192 + Nup145N
* No effect
* Moderate
* Severe
Nup145N 621-625→5-Ala
Nup192 + Nup145N 621-625→5-Ala
Nup145N 646-650→5-Ala
Nup192 + Nup145N 646-650→5-Ala
Nup145N 626-630→5-Ala
Nup192 + Nup145N 626-630→5-Ala
Nup145N 656-660→5-Ala
Nup192 + Nup145N 656-660→5-Ala
Nup145N 651-655→5-Ala
Nup192 + Nup145N 651-655→5-Ala
Nup145N 641-645→5-Ala
Nup192 + Nup145N 641-645→5-Ala
Nup145N 621-625→5A
Nup145N 646-650→5A
Nup145N 626-630→5A
Nup145N 661-665→5A
Nup145N 656-661→5A
Nup145N 651-655→5A
Nup145N 641-645→5A

Supp Fig. S13

**Fig.S13. 5-Ala scanning mutagenesis of Nup145N sequence for residues necessary for Nup192 binding.** SEC and SDS-PAGE analysis corresponding to Fig. 4A. (**A**) Domain structure of Nup145N. Mutations were introduced into the Nup145N construct indicated by the black line above the domain structure. The positions of the 5-Ala mutated residues are indicated above the primary Nup145N sequence with boxes colored according to the measured effect on Nup192 binding: no effect (green), moderate effect (yellow), and severe effect (red). Asterisks above boxes indicate SEC experiments for which representative SDS-PAGE gels are shown in (B). (**B**) SEC profiles of wildtype Nup192 alone (cyan) and Nup192 preincubated with wildtype Nup145N (dark blue) are shown. SEC profiles of Nup145N 5-Ala mutants preincubated with Nup192 are colored according to the measured effect, as in (A). The inset box shows a closeup view of elution profile shifts at the peak fractions. The gray bar indicates fractions that were resolved on SDS-PAGE gels and visualized by Coomassie staining. All SEC experiments were analyzed by SDS-PAGE, but only experiments representative of the three observed effect levels are shown for brevity. All SEC profiles were obtained using a Superdex 200 10/300 GL column.

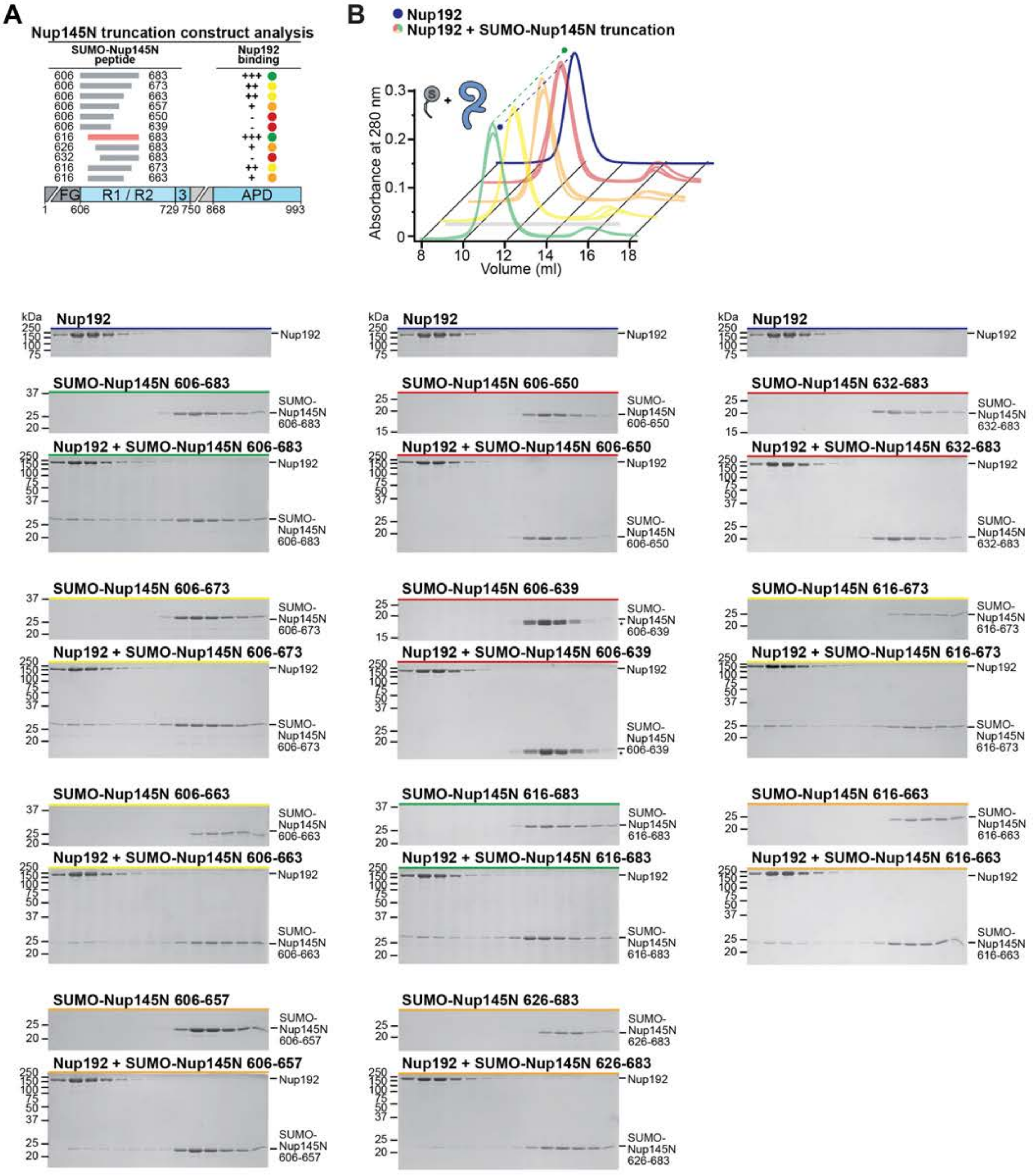
A
Nup145N truncation construct analysis
SUMO-Nup145N peptide
Nup192 binding
606 683 +++
606 673 ++
606 663 ++
606 657 +
606 650 -
606 639 -
616 683 +++
626 683 +
632 683 -
616 673 ++
616 663 +
FG
R1 / R2
3
APD
1 606 729 750 868 993
B
Nup192
Nup192 + SUMO-Nup145N truncation
Absorbance at 280 nm
Volume (ml)
Nup192
SUMO-Nup145N 606-683
Nup192 + SUMO-Nup145N 606-683
SUMO-Nup145N 606-673
Nup192 + SUMO-Nup145N 606-673
SUMO-Nup145N 606-663
Nup192 + SUMO-Nup145N 606-663
SUMO-Nup145N 606-657
Nup192 + SUMO-Nup145N 606-657
SUMO-Nup145N 606-650
Nup192 + SUMO-Nup145N 606-650
SUMO-Nup145N 606-639
Nup192 + SUMO-Nup145N 606-639
SUMO-Nup145N 616-683
Nup192 + SUMO-Nup145N 616-683
SUMO-Nup145N 626-683
Nup192 + SUMO-Nup145N 626-683
SUMO-Nup145N 632-683
Nup192 + SUMO-Nup145N 632-683
SUMO-Nup145N 616-673
Nup192 + SUMO-Nup145N 616-673
SUMO-Nup145N 616-663
Nup192 + SUMO-Nup145N 616-663
kDa

**Fig.S14. Mapping of the minimal Nup145N region sufficient for Nup192 binding.** SEC and SDS-PAGE analysis corresponding to Fig. 4B. (**A**) Domain structure of Nup145N with gray bars indicating truncation construct boundaries analyzed. The Nup145N$^{R1}$ peptide is indicated by a red bar. The binding of SUMO-Nup145N peptides to Nup192, as assessed by SEC and SDS-PAGE, is summarized and colored according to the measured effect: no effect (green, +++), weak effect (yellow, ++), moderate effect (orange, +), and abolished binding (red, -). (**B**) SEC profile of wildtype Nup192 (dark blue) is shown as reference. SEC profiles of Nup192 preincubated with SUMO-Nup145N peptides are colored according to the measured effect, as in (A). Dashed lines indicate the peak elution volumes across the offset dimension. The gray bar indicates fractions that were resolved on SDS-PAGE gels and visualized by Coomassie staining. All SEC profiles were obtained using a Superdex 200 10/300 GL column. Asterisks indicate degradation products.

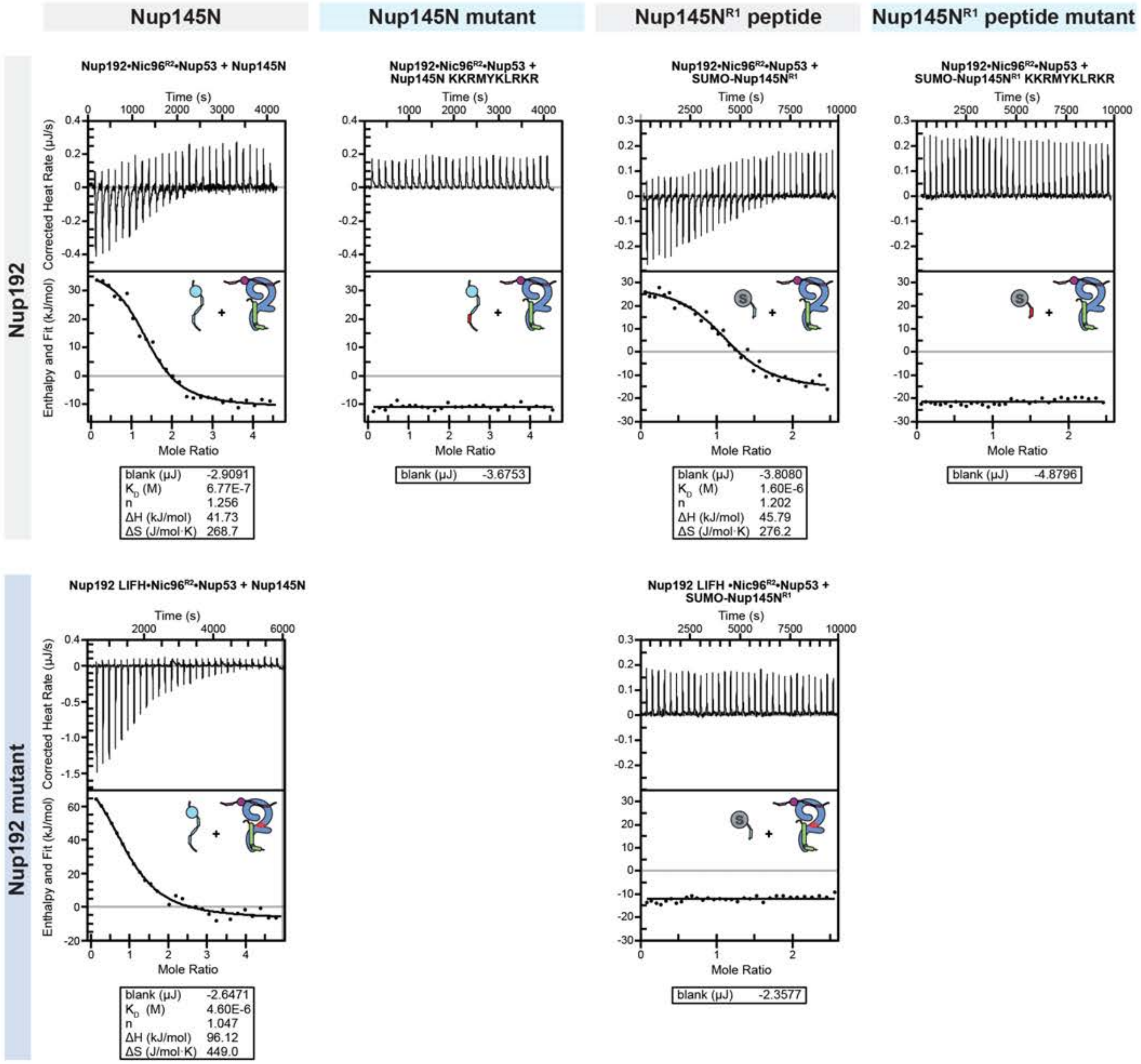
Nup145N
Nup145N mutant
Nup145N$^{R1}$ peptide
Nup145N$^{R1}$ peptide mutant
Nup192
Nup192 mutant
Nup192•Nic96$^{R2}$•Nup53 + Nup145N
Nup192•Nic96$^{R2}$•Nup53 + Nup145N KKRMYKLRKR
Nup192•Nic96$^{R2}$•Nup53 + SUMO-Nup145N$^{R1}$
Nup192•Nic96$^{R2}$•Nup53 + SUMO-Nup145N$^{R1}$ KKRMYKLRKR
Nup192 LIFH•Nic96$^{R2}$•Nup53 + Nup145N
Nup192 LIFH •Nic96$^{R2}$•Nup53 + SUMO-Nup145N$^{R1}$
Time (s)
Corrected Heat Rate (μJ/s)
Enthalpy and Fit (kJ/mol)
Mole Ratio
blank (μJ) -2.9091
$K_D$ (M) 6.77E-7
n 1.256
ΔH (kJ/mol) 41.73
ΔS (J/mol·K) 268.7
blank (μJ) -3.6753
blank (μJ) -3.8080
$K_D$ (M) 1.60E-6
n 1.202
ΔH (kJ/mol) 45.79
ΔS (J/mol·K) 276.2
blank (μJ) -4.8796
blank (μJ) -2.6471
$K_D$ (M) 4.60E-6
n 1.047
ΔH (kJ/mol) 96.12
ΔS (J/mol·K) 449.0
blank (μJ) -2.3577

Supp. Fig. S15

**Fig.S15. Isothermal titration calorimetry analysis of Nup192-Nup145N binding.** Representative baseline-corrected ITC experiments for Nup145N, Nup145N KKRMYKLRKR, SUMO-Nup145N$^{R1}$, and SUMO-Nup145N$^{R1}$ KKRMYKLRKR titrated against a preformed complex of Nic96$^{R2}$, Nup53, and either wildtype Nup192 or mutant Nup192 LIFH. The least-squares fit thermodynamic parameters for a single binding site model are shown in box beneath ITC plot. Overall ITC experimental conditions and results of experiments conducted in triplicate are reported in Table S5.

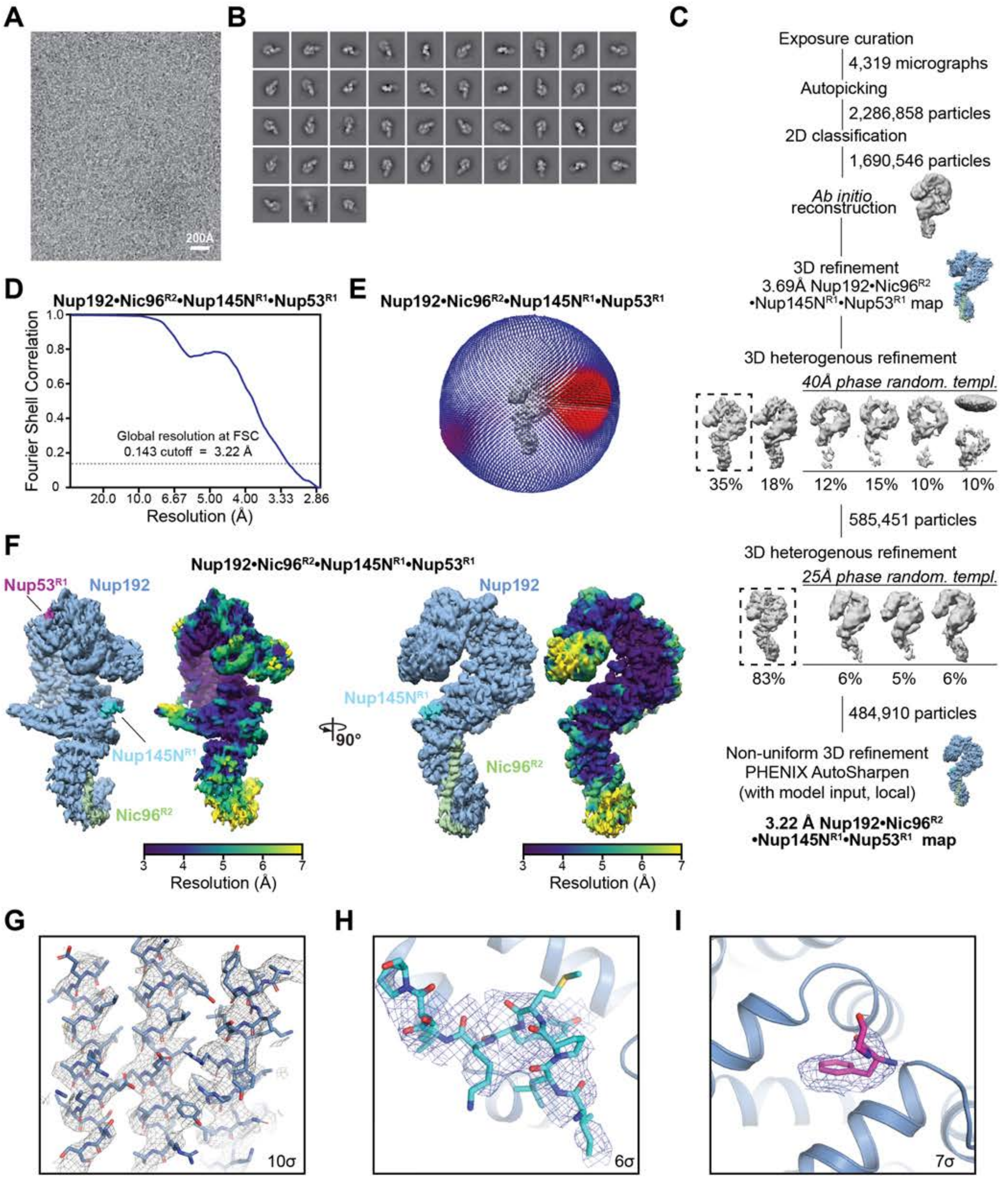
A
200Å
B
C
Exposure curation
4,319 micrographs
Autopicking
2,286,858 particles
2D classification
1,690,546 particles
Ab initio reconstruction
3D refinement
3.69Å Nup192•Nic96R2 •Nup145NR1•Nup53R1 map
3D heterogenous refinement
40Å phase random. templ.
35% 18% 12% 15% 10% 10%
585,451 particles
3D heterogenous refinement
25Å phase random. templ.
83% 6% 5% 6%
484,910 particles
Non-uniform 3D refinement
PHENIX AutoSharpen
(with model input, local)
3.22 Å Nup192•Nic96R2 •Nup145NR1•Nup53R1 map
D
Nup192•Nic96R2•Nup145NR1•Nup53R1
Fourier Shell Correlation
Global resolution at FSC 0.143 cutoff = 3.22 Å
Resolution (Å)
E
Nup192•Nic96R2•Nup145NR1•Nup53R1
F
Nup192•Nic96R2•Nup145NR1•Nup53R1
Nup53R1
Nup192
Nup145NR1
Nic96R2
90°
Resolution (Å)
G
10σ
H
6σ
I
7σ

**Fig.S16. Single particle cryo-EM analysis of the Nup192•Nic96$^{R2}$•Nup145N$^{R1}$•Nup53$^{R1}$ complex.** (**A**) Representative motion corrected cryo-electron micrograph. (**B**) 2D class averages representative of different particle orientations. (**C**) Summary of single particle data processing workflow with indicated number of micrographs or particles involved at each step. Dashed boxes indicate classes that were selected for subsequent steps in the workflow. (**D**) Gold-standard Fourier Shell Correlation curve estimated with 3DFSC and global resolution of 3.2 Å at the 0.143 cutoff. (**E**) Three-dimensional angular distribution plot of particles. (**F**) Isosurface representation of the Nup192•Nic96$^{R2}$•Nup145N$^{R1}$•Nup53$^{R1}$ cryo-EM density (EMD-24057) colored according to protein chain identity (*left*) or local resolution estimation (*right*). (**G-I**) Isomesh representation of representative sharpened cryo-EM density contoured (G) at a 10$\sigma$ cutoff level and stick representation of the corresponding Nup192 polypeptide chain of the Nup192•Nic96$^{R2}$•Nup145N$^{R1}$•Nup53$^{R1}$ structure, (H) contoured at 6$\sigma$ cutoff level around the Nup145N$^{R1}$ peptide shown in stick representation (cyan), and (I) contoured at 7$\sigma$ cutoff level around the Nup53$^{R1}$ peptide shown in stick representation (magenta).

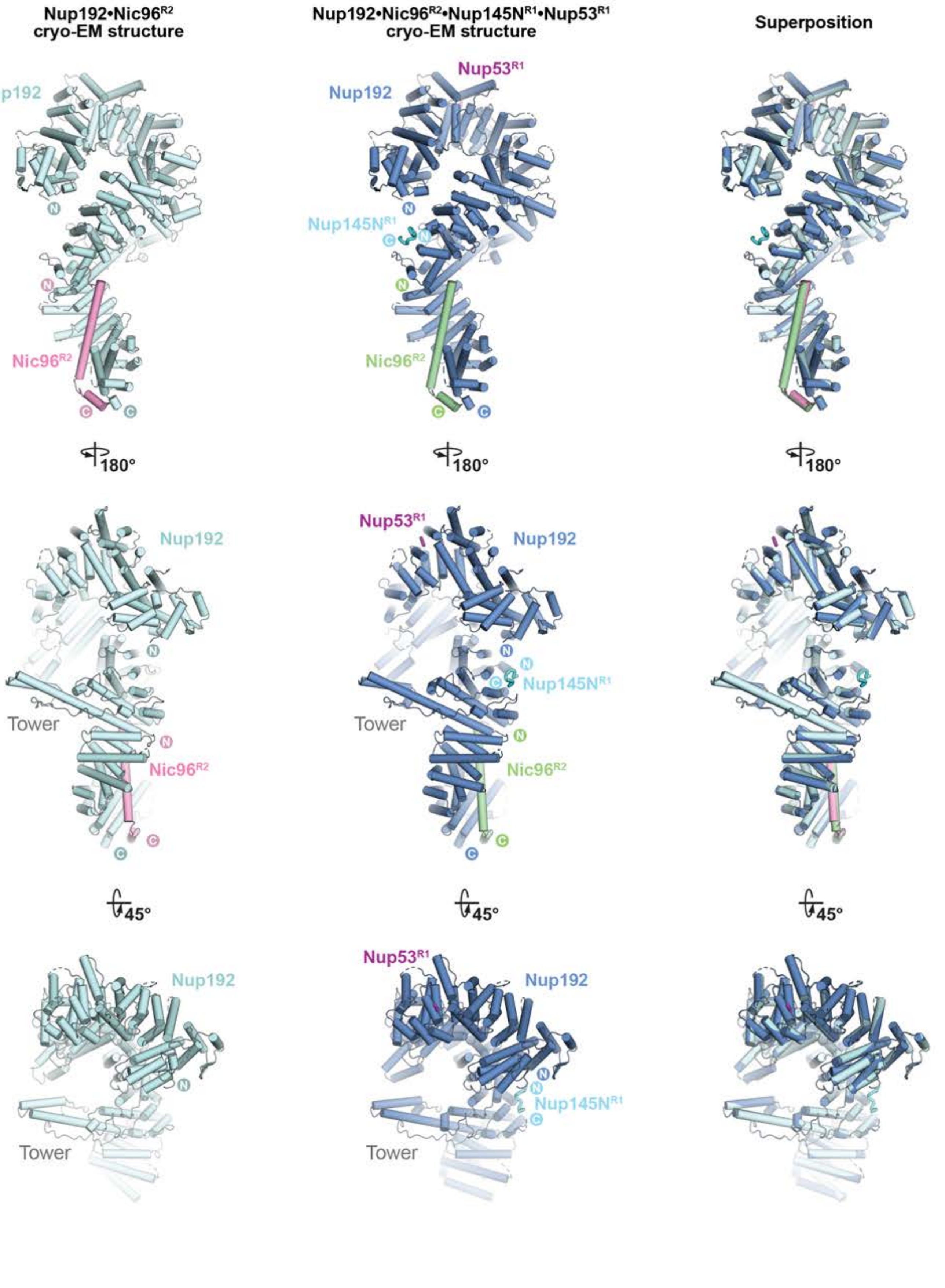

Supp. Fig. S17

Hoelz et al., 2021

**Fig.S17. Nup145N$^{R1}$ and Nup53$^{R1}$ binding to Nup192•Nic96$^{R2}$ does not induce conformational changes**. Cartoon representation of the 3.8 Å Nup192•Nic96$^{R2}$ (pale cyan and pink) and the 3.2 Å Nup192•Nic96$^{R2}$•Nup145N$^{R1}$•Nup53$^{R1}$ (blue, pale green, cyan, and magenta) single particle cryo-EM structures, and their superposition, viewed from three sides.

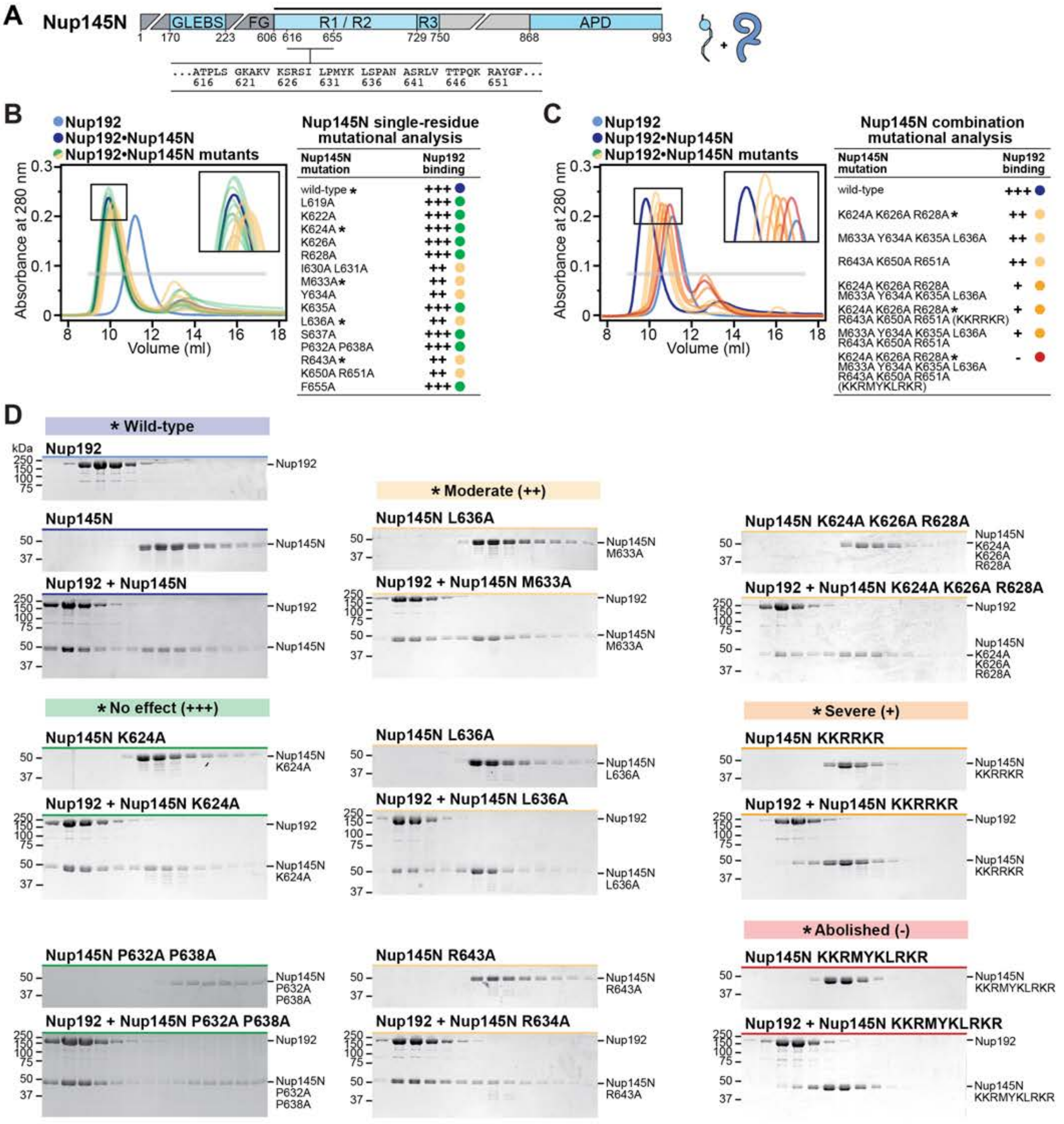
A
Nup145N
GLEBS
FG
R1 / R2
R3
APD
1 170 223 606 616 655 729 750 868 993
...ATPLS GKAKV KSRSI LPMYK LSPAN ASRLV TTPQK RAYGF...
616 621 626 631 636 641 646 651
B
Nup192
Nup192•Nup145N
Nup192•Nup145N mutants
Absorbance at 280 nm
Volume (ml)
Nup145N single-residue mutational analysis
Nup145N mutation | Nup192 binding
wild-type * +++
L619A +++
K622A +++
K624A * +++
K626A +++
R628A +++
I630A L631A ++
M633A* ++
Y634A ++
K635A +++
L636A * ++
S637A +++
P632A P638A +++
R643A* ++
K650A R651A ++
F655A +++
C
Nup192
Nup192•Nup145N
Nup192•Nup145N mutants
Nup145N combination mutational analysis
Nup145N mutation | Nup192 binding
wild-type +++
K624A K626A R628A* ++
M633A Y634A K635A L636A ++
R643A K650A R651A ++
K624A K626A R628A M633A Y634A K635A L636A +
K624A K626A R628A* R643A K650A R651A (KKRRKR) +
M633A Y634A K635A L636A R643A K650A R651A +
K624A K626A R628A* M633A Y634A K635A L636A R643A K650A R651A (KKRMYKLRKR) -
D
* Wild-type
kDa
Nup192
Nup145N
Nup192 + Nup145N
* No effect (+++)
Nup145N K624A
Nup192 + Nup145N K624A
Nup145N P632A P638A
Nup192 + Nup145N P632A P638A
* Moderate (++)
Nup145N L636A
Nup192 + Nup145N M633A
Nup145N L636A
Nup192 + Nup145N L636A
Nup145N R643A
Nup192 + Nup145N R634A
Nup145N K624A K626A R628A
Nup192 + Nup145N K624A K626A R628A
* Severe (+)
Nup145N KKRRKR
Nup192 + Nup145N KKRRKR
* Abolished (-)
Nup145N KKRMYKLRKR
Nup192 + Nup145N KKRMYKLRKR

**Fig.S18. Perturbation of the Nup192-Nup145N interaction by Nup145N point mutants.** SEC and SDS-PAGE analysis corresponding to Fig. 4F. (**A**) Domain structure of Nup145N. Mutations were introduced into the Nup145N construct indicated by the black line above the domain structure. Primary sequence containing the mutated residues is shown beneath the domain structure. (**B-C**) SEC profiles of wildtype Nup192 alone (blue) and Nup192 preincubated with wildtype Nup145N (dark blue) are shown. SEC profiles of Nup192 preincubated with Nup145N mutants are colored according to the measured effect and the results summarized in the adjacent table: no effect (green, +++), weak effect (yellow, ++), moderate effect (orange, +), and abolished binding (red, -). The inset boxes show a closeup view of elution profile shifts at the peak fractions. Asterisks indicate SEC experiments for which representative SDS-PAGE gels are shown in (D). Gray bars indicate fractions that were resolved on SDS-PAGE gels and visualized by Coomassie staining. All SEC profiles were obtained using a Superdex 200 10/300 GL column. (**D**) A selection of SDS-PAGE gels representative of the effect levels observed in (B-C) is shown for brevity.

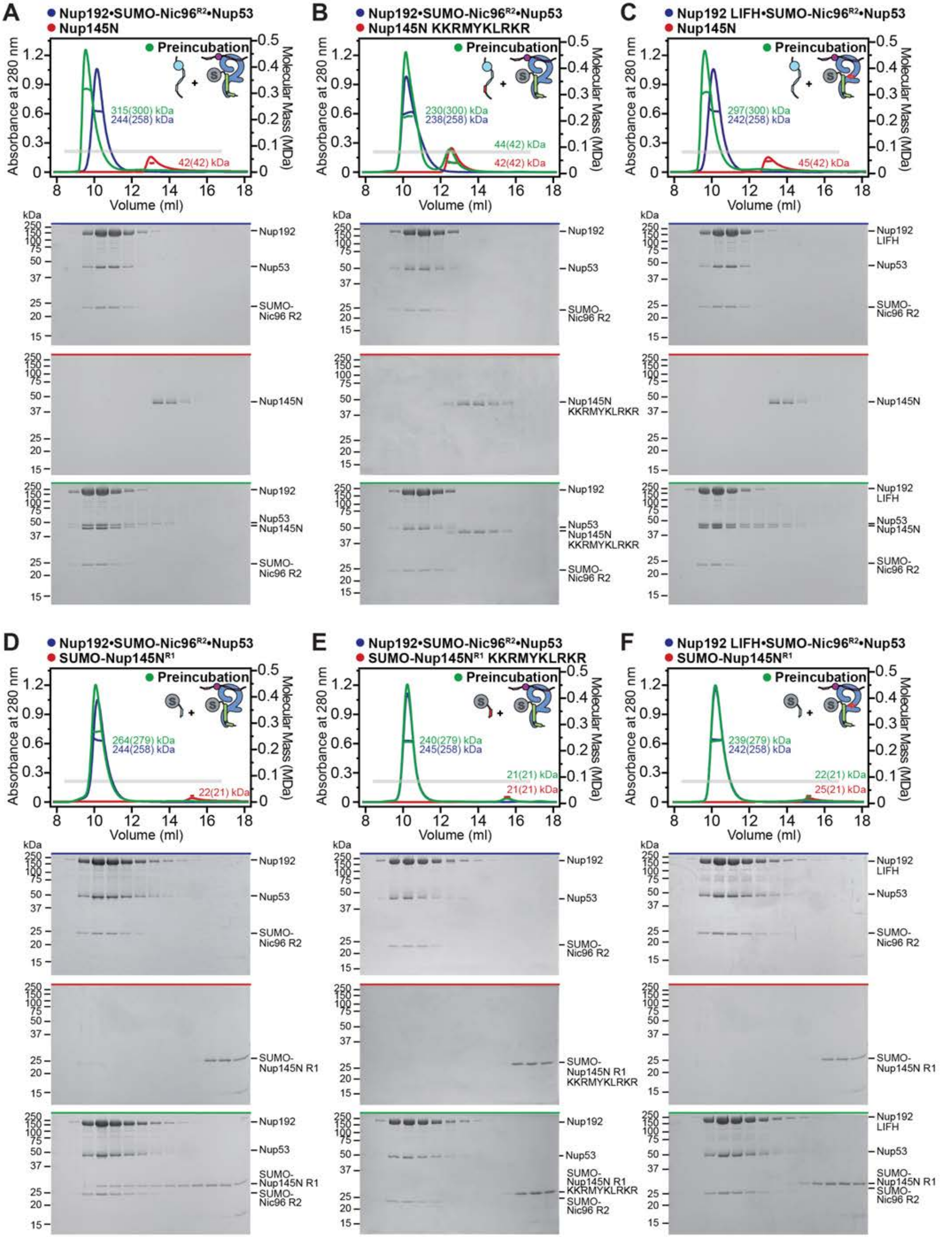
A
Nup192•SUMO-Nic96R2•Nup53
Nup145N
Preincubation
315(300) kDa
244(258) kDa
42(42) kDa
B
Nup192•SUMO-Nic96R2•Nup53
Nup145N KKRMYKLRKR
Preincubation
230(300) kDa
238(258) kDa
44(42) kDa
42(42) kDa
C
Nup192 LIFH•SUMO-Nic96R2•Nup53
Nup145N
Preincubation
297(300) kDa
242(258) kDa
45(42) kDa
D
Nup192•SUMO-Nic96R2•Nup53
SUMO-Nup145NR1
Preincubation
264(279) kDa
244(258) kDa
22(21) kDa
E
Nup192•SUMO-Nic96R2•Nup53
SUMO-Nup145NR1 KKRMYKLRKR
Preincubation
240(279) kDa
245(258) kDa
21(21) kDa
21(21) kDa
F
Nup192 LIFH•SUMO-Nic96R2•Nup53
SUMO-Nup145NR1
Preincubation
239(279) kDa
242(258) kDa
22(21) kDa
25(21) kDa
Absorbance at 280 nm
Molecular Mass (MDa)
Volume (ml)
kDa
Nup192
Nup192 LIFH
Nup53
SUMO-Nic96 R2
Nup145N
Nup145N KKRMYKLRKR
Nup53 Nup145N
SUMO-Nup145N R1
SUMO-Nup145N R1 KKRMYKLRKR

Supp. Fig. S19

**Fig.S19. SEC-MALS analysis of the Nup192-Nup145N interaction.** SEC-MALS and SDS-PAGE analysis corresponding to Fig. 4H. (**A-F**) SEC-MALS profiles of nups are shown individually (red and blue) and after their preincubation (green) for either wilt-type Nup192•SUMO-Nic96$^{R2}$•Nup53 or mutant Nup192 LIFH•SUMO-Nic96$^{R2}$•Nup53 binding to either wildtype or KKRMYKLRKR mutant of Nup145N and SUMO-Nup145N$^{R1}$. Measured molecular masses are indicated for the peak fractions and corresponding theoretical molecular masses are reported in parenthesis. Gray bars indicate fractions that were resolved on SDS-PAGE gels and visualized by Coomassie staining. All SEC profiles were obtained using a Superdex 200 Increase 10/300 GL column. All SEC-MALS results are summarized in Table S4.

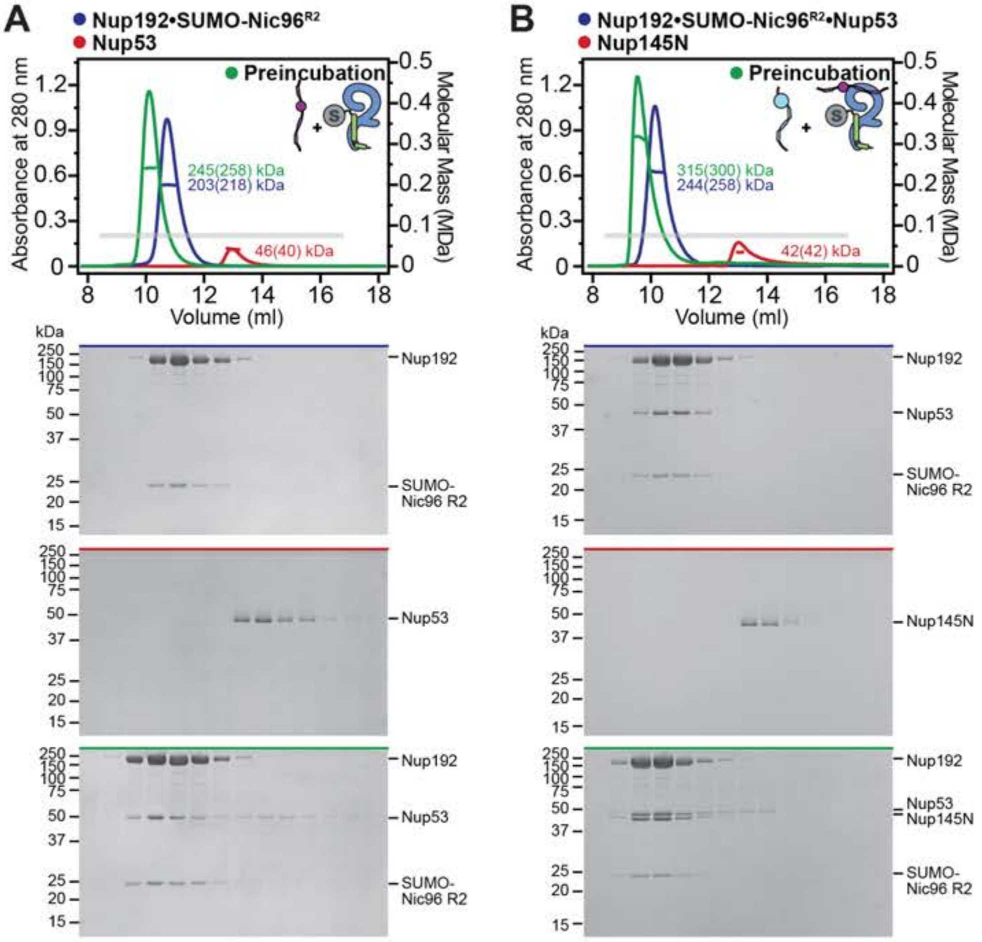
A
Nup192•SUMO-Nic96R2
Nup53
Preincubation
Absorbance at 280 nm
Molecular Mass (MDa)
245(258) kDa
203(218) kDa
46(40) kDa
Volume (ml)
kDa
Nup192
SUMO-Nic96 R2
Nup53
B
Nup192•SUMO-Nic96R2•Nup53
Nup145N
Preincubation
315(300) kDa
244(258) kDa
42(42) kDa
Nup192
Nup53
SUMO-Nic96 R2
Nup145N
Nup53
Nup145N

**Fig.S20. Reconstitution of the Nup192•SUMO-Nic96$^{R2}$•Nup53 complex.** SEC-MALS profiles of nups are shown individually (red and blue) and after their preincubation (green) for (**A**) the complexation of Nup192 and SUMO-Nic96$^{R2}$, and (**B**) subsequent complexation of Nup53 with Nup192•SUMO-Nic96$^{R2}$. Measured molecular masses are indicated for the peak fractions and corresponding theoretical molecular masses are reported in parenthesis. Gray bars indicate fractions that were resolved on SDS-PAGE gels and visualized by Coomassie staining. All SEC profiles were obtained using a Superdex 200 Increase 10/300 GL column. All SEC-MALS results are summarized in Table S4.

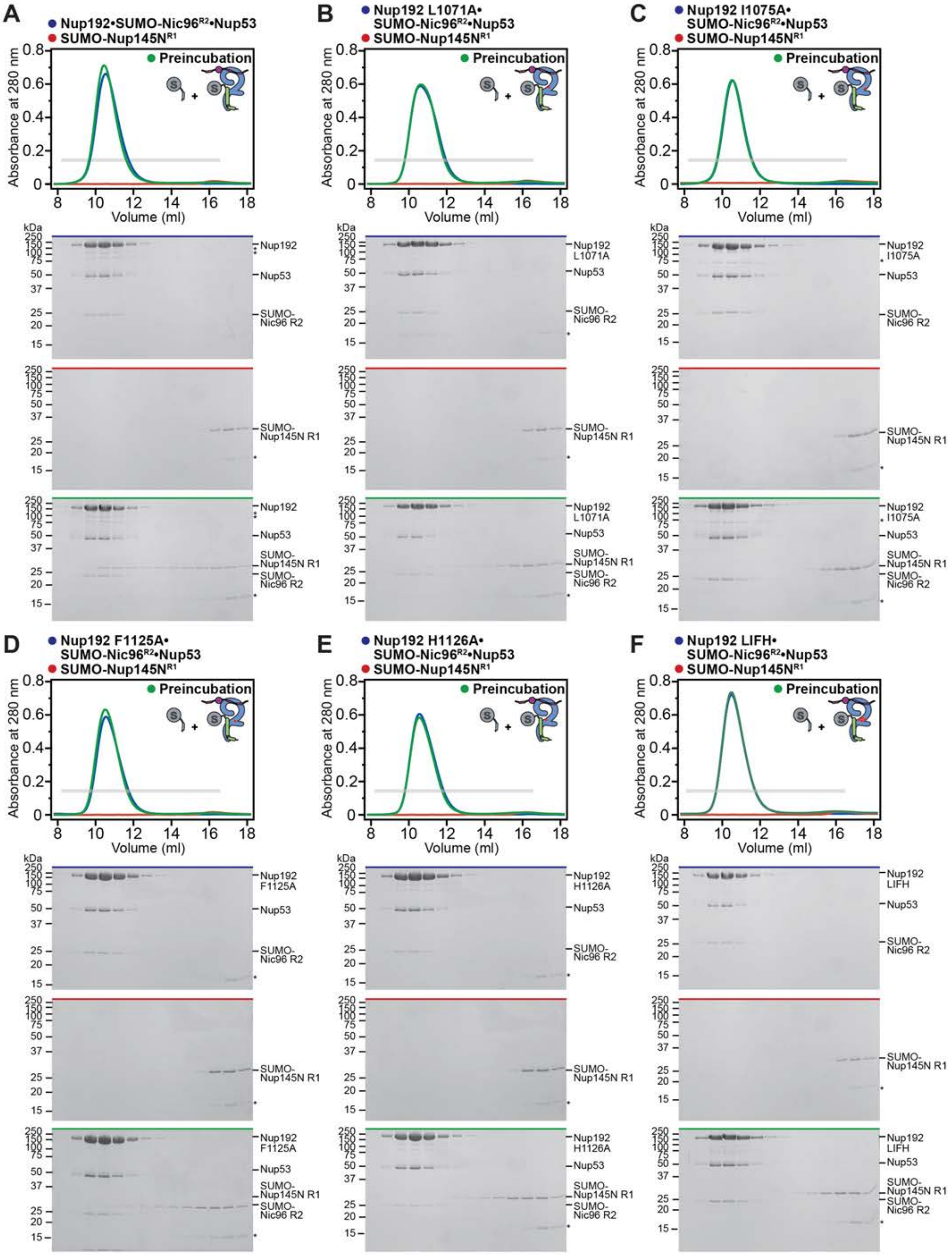

Supp. Fig. S21

Hoelz et al., 2021

**Fig.S21. Perturbation of the Nup192-Nup145N interaction by Nup192 point mutants.** (**A-F**) SEC and SDS-PAGE analysis corresponding to Fig. 4F. SEC profiles are shown for nups or nup complexes individually (red and blue) and after their preincubation (green) for SUMO-Nup145N$^{R1}$ binding to pre-assembled complexes of SUMO-Nic96$^{R2}$, Nup53, and wildtype or mutant Nup192. Gray bars indicate fractions that were resolved on SDS-PAGE gels and visualized by Coomassie staining. All SEC profiles were obtained using a Superdex 200 10/300 GL column. Asterisks indicate SUMO contaminant.

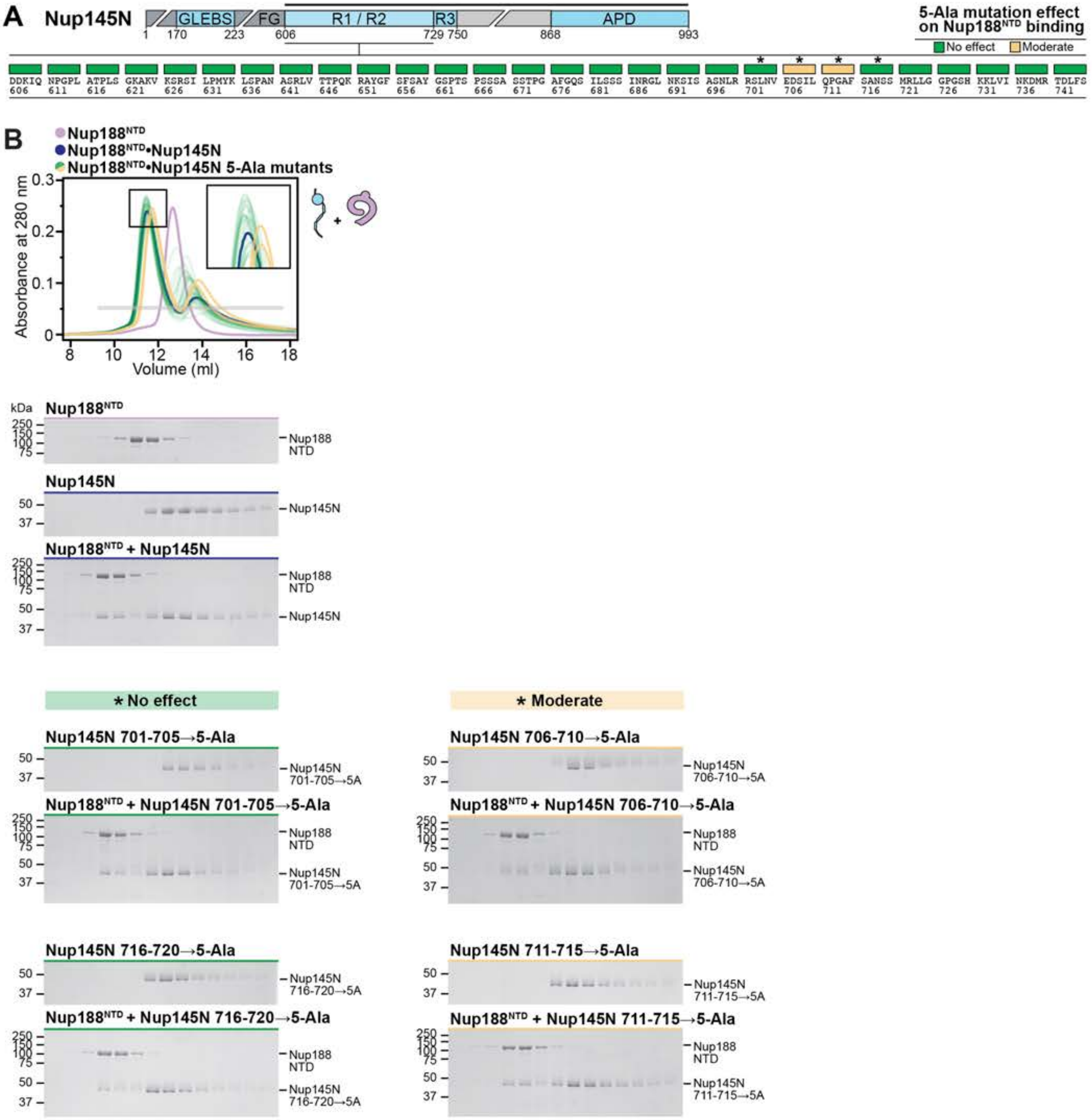
A
Nup145N
GLEBS
FG
R1 / R2
R3
APD
1
170
223
606
729
750
868
993
5-Ala mutation effect on Nup188NTD binding
No effect
Moderate
DDKIQ 606
NPGPL 611
ATPLS 616
GKAKV 621
KSRSI 626
LPMYK 631
LSPAN 636
ASRLV 641
TTPQK 646
RAYGF 651
SFSAY 656
GSPTS 661
PSSSA 666
SSTPG 671
AFGQS 676
ILSSS 681
INRGL 686
NKSIS 691
ASNLR 696
RSLNV 701
EDSIL 706
QPGAF 711
SANSS 716
MRLLG 721
GPGSH 726
KKLVI 731
NKDMR 736
TDLFS 741
B
Nup188NTD
Nup188NTD•Nup145N
Nup188NTD•Nup145N 5-Ala mutants
Absorbance at 280 nm
Volume (ml)
kDa
Nup188NTD
Nup145N
Nup188NTD + Nup145N
Nup188 NTD
* No effect
* Moderate
Nup145N 701-705→5-Ala
Nup188NTD + Nup145N 701-705→5-Ala
Nup145N 701-705→5A
Nup145N 706-710→5-Ala
Nup188NTD + Nup145N 706-710→5-Ala
Nup145N 706-710→5A
Nup145N 716-720→5-Ala
Nup188NTD + Nup145N 716-720→5-Ala
Nup145N 716-720→5A
Nup145N 711-715→5-Ala
Nup188NTD + Nup145N 711-715→5-Ala
Nup145N 711-715→5A

**Fig.S22. 5-Ala scanning mutagenesis of Nup145N sequence for residues necessary for Nup188$^{NTD}$ binding.** SEC and SDS-PAGE analysis corresponding to Fig. 5A. (**A**) Domain structure of Nup145N. Mutations were introduced into the Nup145N construct indicated by the black line above the domain structure. The positions of the 5-Ala mutated residues are indicated above the primary Nup145N sequence with boxes colored according to the measured effect on Nup188$^{NTD}$ binding: no effect (green), moderate effect (yellow). Asterisks above boxes indicate SEC experiments for which representative SDS-PAGE gels are shown in (B). (**B**) SEC profiles of wildtype Nup188$^{NTD}$ alone (light purple) and Nup188$^{NTD}$ preincubated with wildtype Nup145N (dark blue) are shown. SEC profiles of Nup145N 5-Ala mutants preincubated with Nup188$^{NTD}$ are colored according to the measured effect, as in (A). The inset box shows a closeup view of elution profile shifts at the peak fractions. The gray bar indicates fractions that were resolved on SDS-PAGE gels and visualized by Coomassie staining. All SEC experiments were analyzed by SDS-PAGE, but only experiments representative of the three observed effect levels are shown for brevity. All SEC profiles were obtained using a Superdex 200 10/300 GL column.

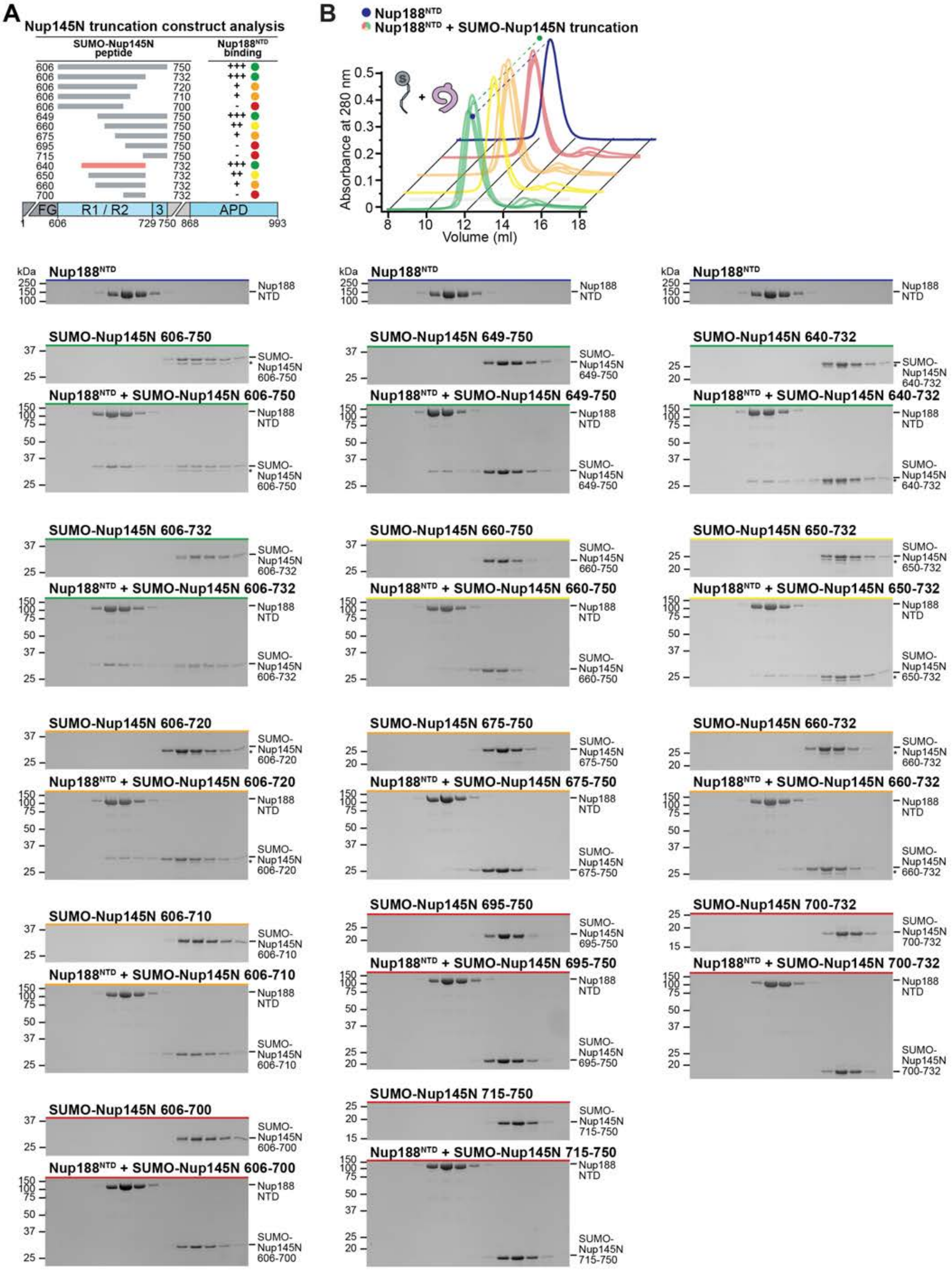

A
Nup145N truncation construct analysis
SUMO-Nup145N peptide
Nup188NTD binding
606 750 +++
606 732 +++
606 720 +
606 710 +
606 700 -
649 750 +++
660 750 ++
675 750 +
695 750 -
715 750 -
640 732 +++
650 732 ++
660 732 +
700 732 -
FG
R1 / R2
3
APD
1
606
729 750 868
993
B
Nup188NTD
Nup188NTD + SUMO-Nup145N truncation
Absorbance at 280 nm
Volume (ml)
kDa
Nup188NTD
Nup188 NTD
SUMO-Nup145N 606-750
Nup188NTD + SUMO-Nup145N 606-750
SUMO-Nup145N 649-750
Nup188NTD + SUMO-Nup145N 649-750
SUMO-Nup145N 640-732
Nup188NTD + SUMO-Nup145N 640-732
SUMO-Nup145N 606-732
Nup188NTD + SUMO-Nup145N 606-732
SUMO-Nup145N 660-750
Nup188NTD + SUMO-Nup145N 660-750
SUMO-Nup145N 650-732
Nup188NTD + SUMO-Nup145N 650-732
SUMO-Nup145N 606-720
Nup188NTD + SUMO-Nup145N 606-720
SUMO-Nup145N 675-750
Nup188NTD + SUMO-Nup145N 675-750
SUMO-Nup145N 660-732
Nup188NTD + SUMO-Nup145N 660-732
SUMO-Nup145N 606-710
Nup188NTD + SUMO-Nup145N 606-710
SUMO-Nup145N 695-750
Nup188NTD + SUMO-Nup145N 695-750
SUMO-Nup145N 700-732
Nup188NTD + SUMO-Nup145N 700-732
SUMO-Nup145N 606-700
Nup188NTD + SUMO-Nup145N 606-700
SUMO-Nup145N 715-750
Nup188NTD + SUMO-Nup145N 715-750

Supp. Fig. S23

**Fig.S23. Mapping of the minimal Nup145N region sufficient for Nup188$^{NTD}$ binding.** SEC and SDS-PAGE analysis corresponding to Fig. 5B. (**A**) Domain structure of Nup145N with gray bars indicating truncation construct boundaries analyzed. The Nup145N$^{R2}$ peptide is indicated by a red bar. The binding of SUMO-Nup145N peptides to Nup188$^{NTD}$, as assessed by SEC and SDS-PAGE, is summarized and colored according to the measured effect: no effect (green, +++), weak effect (yellow, ++), moderate effect (orange, +), and abolished binding (red, -). (**B**) SEC profile of wildtype Nup188$^{NTD}$ (dark blue) is shown as reference. SEC profiles of Nup188$^{NTD}$ preincubated with SUMO-Nup145N peptides are offset for clarity and colored according to the measured effect, as in (A). Dashed lines indicate the peak elution volumes across the offset dimension. The gray bar indicates fractions that were resolved on SDS-PAGE gels and visualized by Coomassie staining. All SEC profiles were obtained using a Superdex 200 10/300 GL column. Asterisks indicate degradation products.

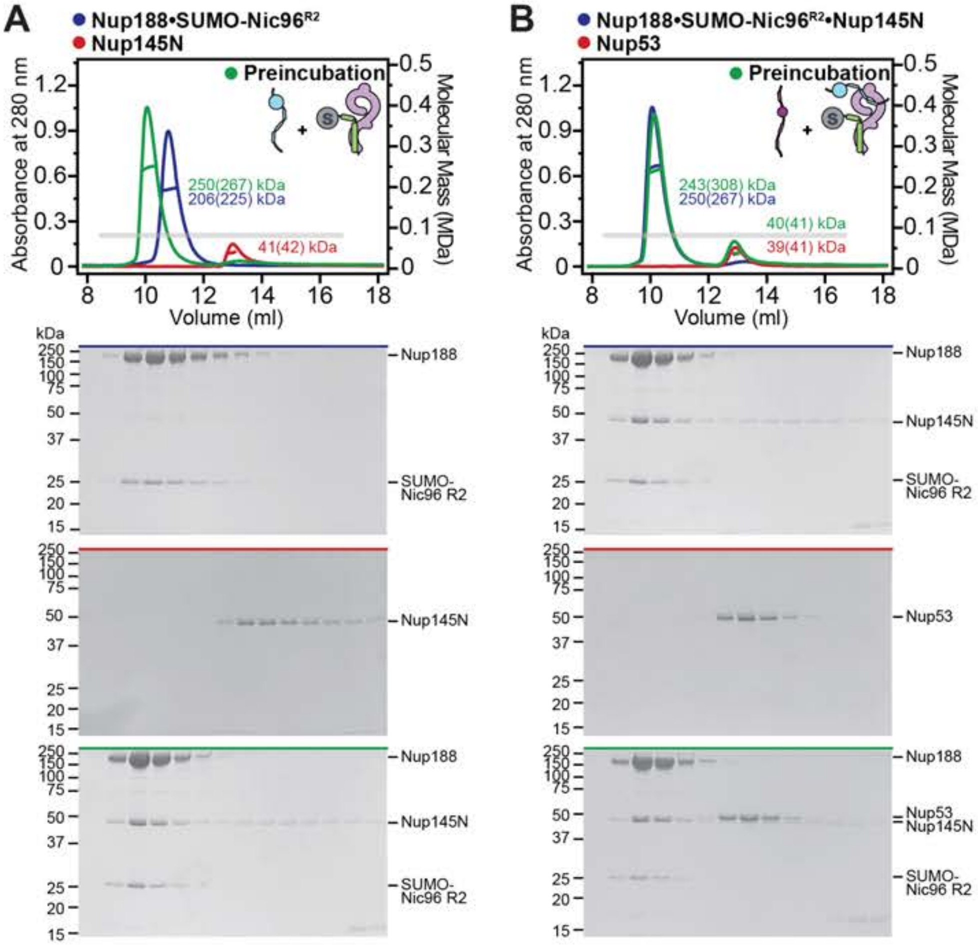
A
Nup188•SUMO-Nic96R2
Nup145N
Preincubation
250(267) kDa
206(225) kDa
41(42) kDa
Absorbance at 280 nm
Molecular Mass (MDa)
Volume (ml)
kDa
Nup188
SUMO-Nic96 R2
Nup145N
B
Nup188•SUMO-Nic96R2•Nup145N
Nup53
243(308) kDa
250(267) kDa
40(41) kDa
39(41) kDa
Nup53
Nup145N

**Fig.S24. Nup53 is not part of the Nup188 symmetric core complex.** SEC-MALS profiles of nups are shown individually (red and blue) and after their preincubation (green) for (**A**) the complexation of Nup188•SUMO-Nic96$^{R2}$ and Nup145N, and (**B**) subsequent binding experiment between Nup188•SUMO-Nic96$^{R2}$•Nup145N and Nup53. Measured molecular masses are indicated for the peak fractions and corresponding theoretical molecular masses are reported in parenthesis. Gray bars indicate fractions that were resolved on SDS-PAGE gels and visualized by Coomassie staining. All SEC profiles were obtained using a Superdex 200 Increase 10/300 GL column. All SEC-MALS results are summarized in Table S4.

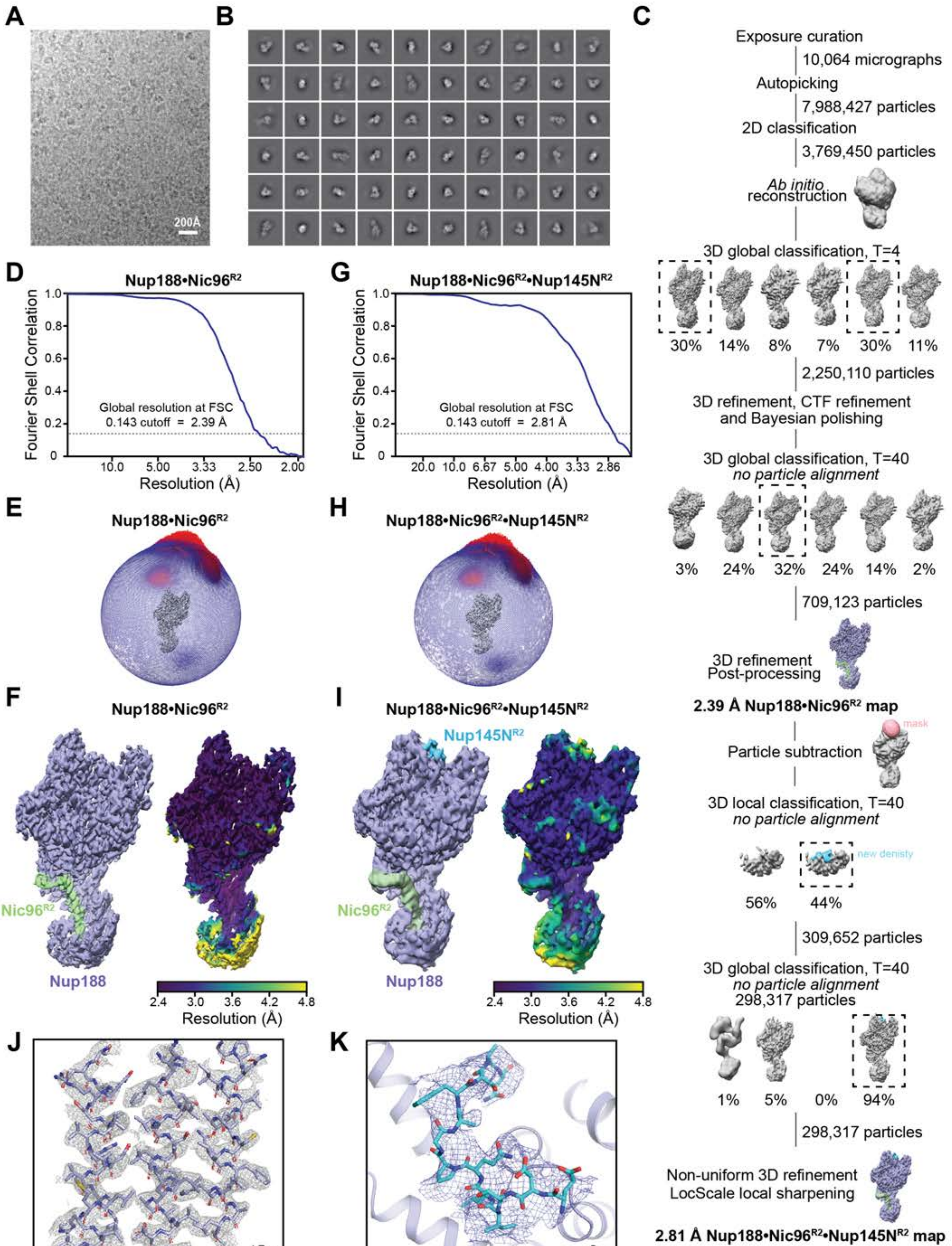
A
200Å
B
C
Exposure curation
10,064 micrographs
Autopicking
7,988,427 particles
2D classification
3,769,450 particles
Ab initio reconstruction
3D global classification, T=4
30% 14% 8% 7% 30% 11%
2,250,110 particles
3D refinement, CTF refinement and Bayesian polishing
3D global classification, T=40
no particle alignment
3% 24% 32% 24% 14% 2%
709,123 particles
3D refinement
Post-processing
2.39 Å Nup188•Nic96R2 map
mask
Particle subtraction
3D local classification, T=40
no particle alignment
new denisty
56% 44%
309,652 particles
3D global classification, T=40
no particle alignment
298,317 particles
1% 5% 0% 94%
298,317 particles
Non-uniform 3D refinement
LocScale local sharpening
2.81 Å Nup188•Nic96R2•Nup145NR2 map
D
Nup188•Nic96R2
Fourier Shell Correlation
Global resolution at FSC 0.143 cutoff = 2.39 Å
Resolution (Å)
G
Nup188•Nic96R2•Nup145NR2
Global resolution at FSC 0.143 cutoff = 2.81 Å
E
Nup188•Nic96R2
H
Nup188•Nic96R2•Nup145NR2
F
Nup188•Nic96R2
Nic96R2
Nup188
I
Nup188•Nic96R2•Nup145NR2
Nup145NR2
Nic96R2
Nup188
Resolution (Å)
J
15σ
K
8σ

**Fig.S25. Single particle cryo-EM analysis of the Nup188•Nic96$^{R2}$•Nup145N$^{R2}$ complex.** (**A**) Representative motion corrected cryo-electron micrograph. (**B**) 2D class averages representative of different particle orientations. (**C**) Summary of single particle data processing workflow with indicated number of micrographs or particles involved at each step. Dashed boxes indicate classes that were selected for subsequent steps in the workflow. (**D-I**) Resolution and particle orientation estimates (D, G) Gold-standard Fourier Shell Correlation curve estimated with 3DFSC and global resolution at the 0.143 cutoff, (E, H) three-dimensional angular distribution plot of particles, and (F, I) isosurface representation of the cryo-EM density colored according to protein chain identity (*left*) or local resolution estimation (*right*) for: (D-F) the Nup188•Nic96$^{R2}$ set of particles, or (G-I) the Nup188•Nic96$^{R2}$•Nup145N$^{R2}$ subset of particles. (**J**) Isomesh representation of representative sharpened cryo-EM density of the 2.4 Å Nup188•Nic96$^{R2}$ map (EMD-24058) contoured at a 15$\sigma$ cutoff level and ball-and-stick representation of the corresponding Nup188 polypeptide chain of the atomic Nup188•Nic96$^{R2}$ model (light purple). (**K**) Isomesh representation of representative sharpened cryo-EM density of the 2.8 Å Nup188•Nic96$^{R2}$•Nup145N$^{R2}$ map (EMD-24059) contoured at an 8 $\sigma$ cutoff level around the Nup145N$^{R2}$ peptide shown in stick representation (cyan).

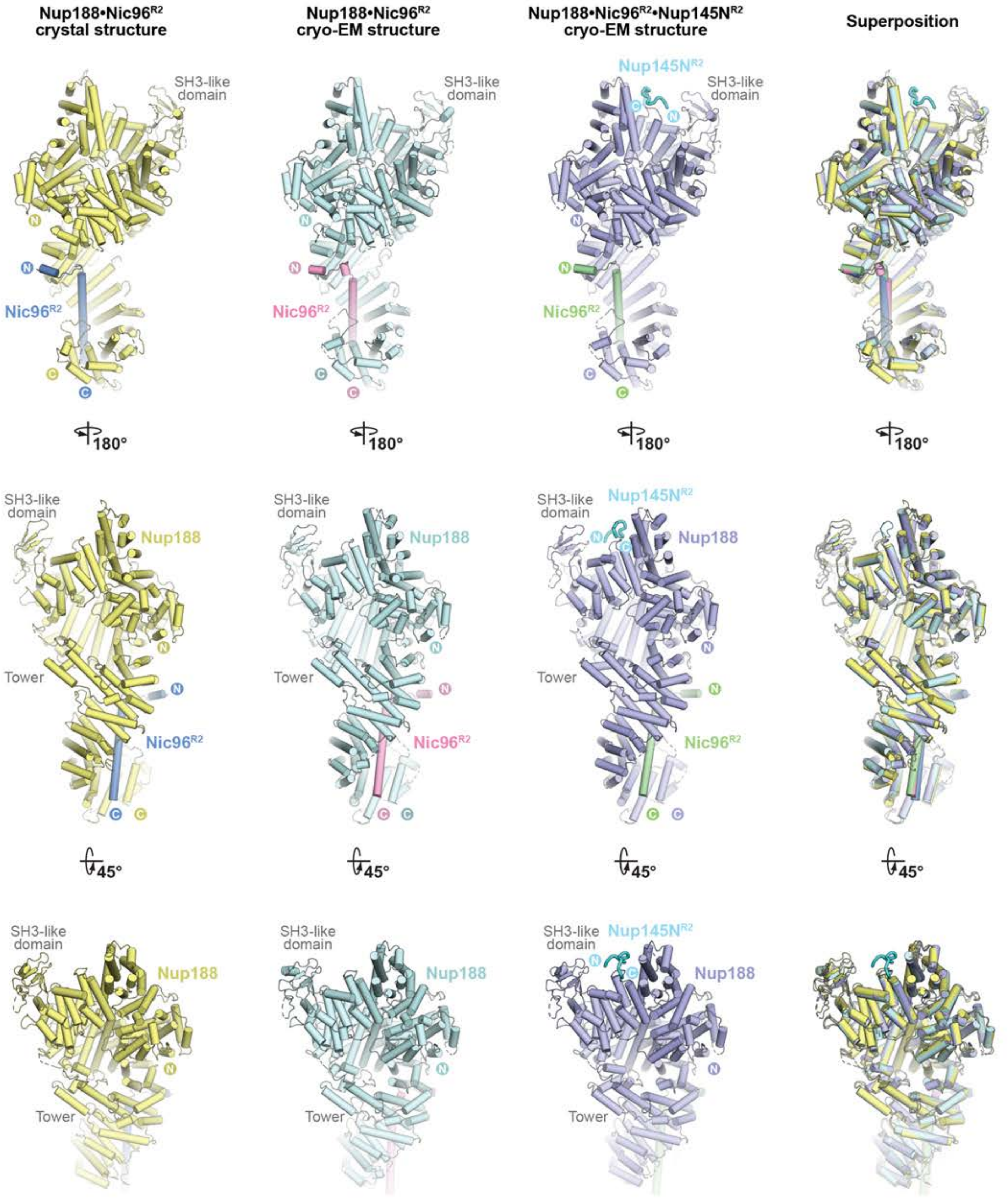

Nup188•Nic96R2
crystal structure
Nup188•Nic96R2
cryo-EM structure
Nup188•Nic96R2•Nup145NR2
cryo-EM structure
Superposition
SH3-like domain
Nup145NR2
Nic96R2
Nup188
Tower
180°
45°
N
C

Supp. Fig. S26

**Fig.S26. Nup188•Nic96$^{R2}$ crystal structure and single particle cryo-EM structures of Nup188•Nic96$^{R2}$ and Nup188•Nic96$^{R2}$•Nup145N$^{R2}$ do not present major conformational differences**. Cartoon representation of the 4.4 Å Nup188•Nic96$^{R2}$ crystal structure (yellow and blue), the 2.4 Å Nup188•Nic96$^{R2}$ (pale cyan and pink) and the 2.8 Å Nup188•Nic96$^{R2}$•Nup145N$^{R2}$ (light purple, pale green, cyan) single particle cryo-EM structures, and their superposition, viewed from three sides.

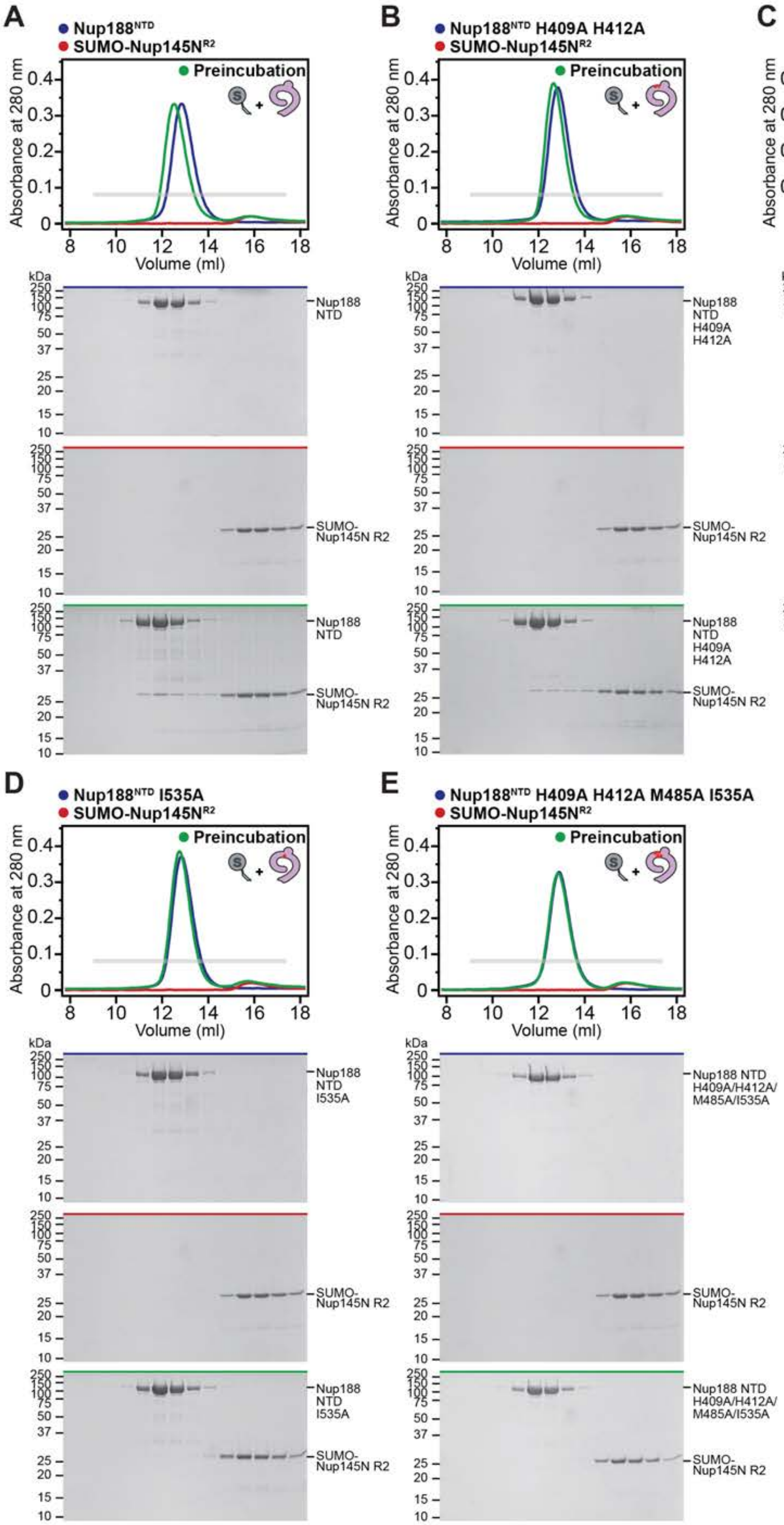
A
Nup188^NTD
SUMO-Nup145N^R2
Preincubation
Absorbance at 280 nm
Volume (ml)
kDa
Nup188 NTD
SUMO-Nup145N R2
B
Nup188^NTD H409A H412A
SUMO-Nup145N^R2
Preincubation
Nup188 NTD H409A H412A
SUMO-Nup145N R2
C
Nup188^NTD M485A
SUMO-Nup145N^R2
Preincubation
Nup188 NTD M485A
SUMO-Nup145N R2
D
Nup188^NTD I535A
SUMO-Nup145N^R2
Preincubation
Nup188 NTD I535A
SUMO-Nup145N R2
E
Nup188^NTD H409A H412A M485A I535A
SUMO-Nup145N^R2
Preincubation
Nup188 NTD H409A/H412A/M485A/I535A
SUMO-Nup145N R2

**Fig.S27. Perturbation of the Nup188$^{NTD}$-Nup145N interaction by Nup188$^{NTD}$ point mutants.** (**A-E**) SEC and SDS-PAGE analysis corresponding to Fig. 5D. SEC profiles are shown for nups individually (red and blue) and after their preincubation (green) for SUMO-Nup145N$^{R2}$ binding to wildtype or mutant Nup188$^{NTD}$. Gray bars indicate fractions that were resolved on SDS-PAGE gels and visualized by Coomassie staining. All SEC profiles were obtained using a Superdex 200 10/300 GL column.

**A**

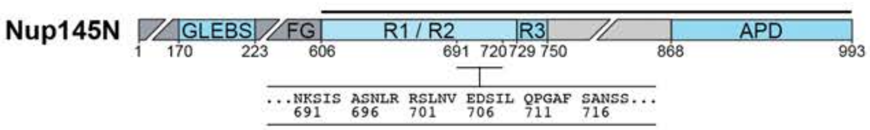

**B**

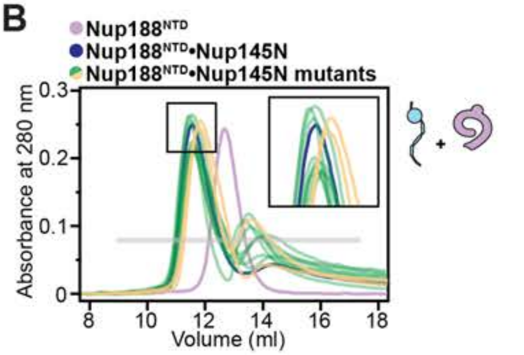

**Nup145N single-residue mutational analysis**

| Nup145N mutation | Nup188NTD binding |
|---|---|
| wild-type * | +++ |
| R700A | +++ |
| R701A | +++ |
| L703A | +++ |
| N704A | +++ |
| E706A * | +++ |
| D707A * | +++ |
| I709A * | +++ |
| L710A * | ++ |
| Q711A * | +++ |
| F715A * | ++ |
| N718A | +++ |

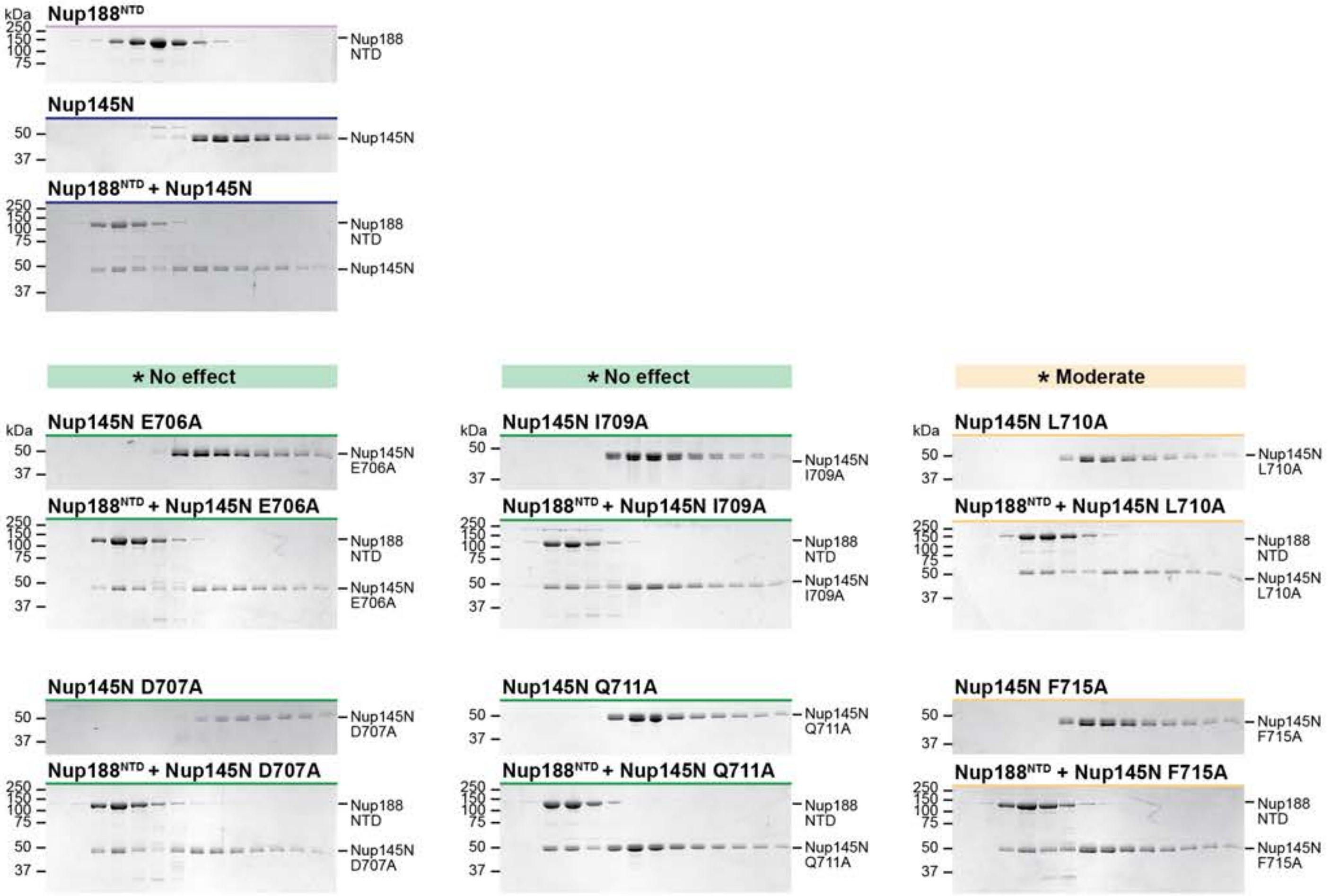

**Fig.S28. Perturbation of the Nup188$^{NTD}$-Nup145N interaction by Nup145N point mutants.** SEC and SDS-PAGE analysis corresponding to Fig. 5D. (**A**) Domain structure of Nup145N. Mutations were introduced into the Nup145N construct indicated by the black line above the domain structure. Primary sequence containing the mutated residues is shown beneath the domain structure. (**B**) SEC profiles of wildtype Nup188$^{NTD}$ alone (light purple) and Nup188$^{NTD}$ preincubated with wildtype Nup145N (dark blue) are shown. SEC profiles of Nup188$^{NTD}$ preincubated with Nup145N mutants are colored according to the measured effect and the results summarized in the adjacent table: no effect (green, +++), weak effect (yellow, ++). The inset boxes show a closeup view of elution profile shifts at the peak fractions. Asterisks indicate SEC experiments for which SDS-PAGE gels representative of the effects observed are shown. Gray bars indicate fractions that were resolved on SDS-PAGE gels and visualized by Coomassie staining. All SEC profiles were obtained using a Superdex 200 10/300 GL column.

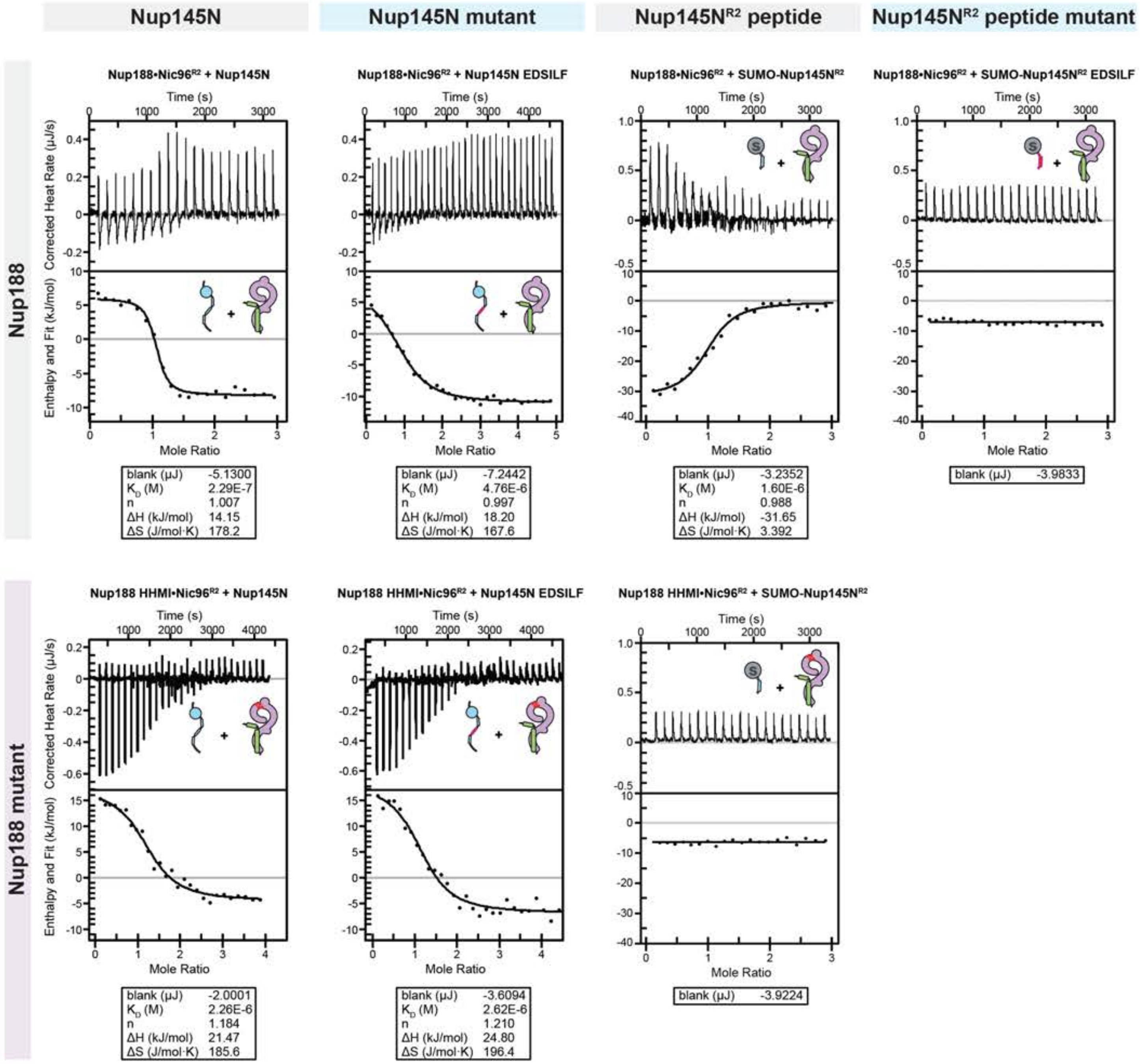

Nup145N
Nup145N mutant
Nup145N$^{R2}$ peptide
Nup145N$^{R2}$ peptide mutant
Nup188
Nup188•Nic96$^{R2}$ + Nup145N
Nup188•Nic96$^{R2}$ + Nup145N EDSILF
Nup188•Nic96$^{R2}$ + SUMO-Nup145N$^{R2}$
Nup188•Nic96$^{R2}$ + SUMO-Nup145N$^{R2}$ EDSILF
Time (s)
Corrected Heat Rate (µJ/s)
Enthalpy and Fit (kJ/mol)
Mole Ratio
blank (µJ) -5.1300
K$_D$ (M) 2.29E-7
n 1.007
ΔH (kJ/mol) 14.15
ΔS (J/mol·K) 178.2
blank (µJ) -7.2442
K$_D$ (M) 4.76E-6
n 0.997
ΔH (kJ/mol) 18.20
ΔS (J/mol·K) 167.6
blank (µJ) -3.2352
K$_D$ (M) 1.60E-6
n 0.988
ΔH (kJ/mol) -31.65
ΔS (J/mol·K) 3.392
blank (µJ) -3.9833
Nup188 mutant
Nup188 HHMI•Nic96$^{R2}$ + Nup145N
Nup188 HHMI•Nic96$^{R2}$ + Nup145N EDSILF
Nup188 HHMI•Nic96$^{R2}$ + SUMO-Nup145N$^{R2}$
blank (µJ) -2.0001
K$_D$ (M) 2.26E-6
n 1.184
ΔH (kJ/mol) 21.47
ΔS (J/mol·K) 185.6
blank (µJ) -3.6094
K$_D$ (M) 2.62E-6
n 1.210
ΔH (kJ/mol) 24.80
ΔS (J/mol·K) 196.4
blank (µJ) -3.9224

Supp. Fig. S29

**Fig.S29. Isothermal titration calorimetry analysis of Nup188-Nup145N binding.** Representative baseline-corrected ITC experiments for Nup145N, Nup145N EDSILF, SUMO-Nup145N$^{R2}$, and SUMO-Nup145N$^{R2}$ EDSILF titrated against a preformed complex of Nic96$^{R2}$and either wildtype Nup188 or mutant Nup188 HHMI. The least-squares fit thermodynamic parameters for a single binding site model are shown in box beneath ITC plot. Overall ITC experimental conditions and results of experiments conducted in triplicate are reported in Table S5.

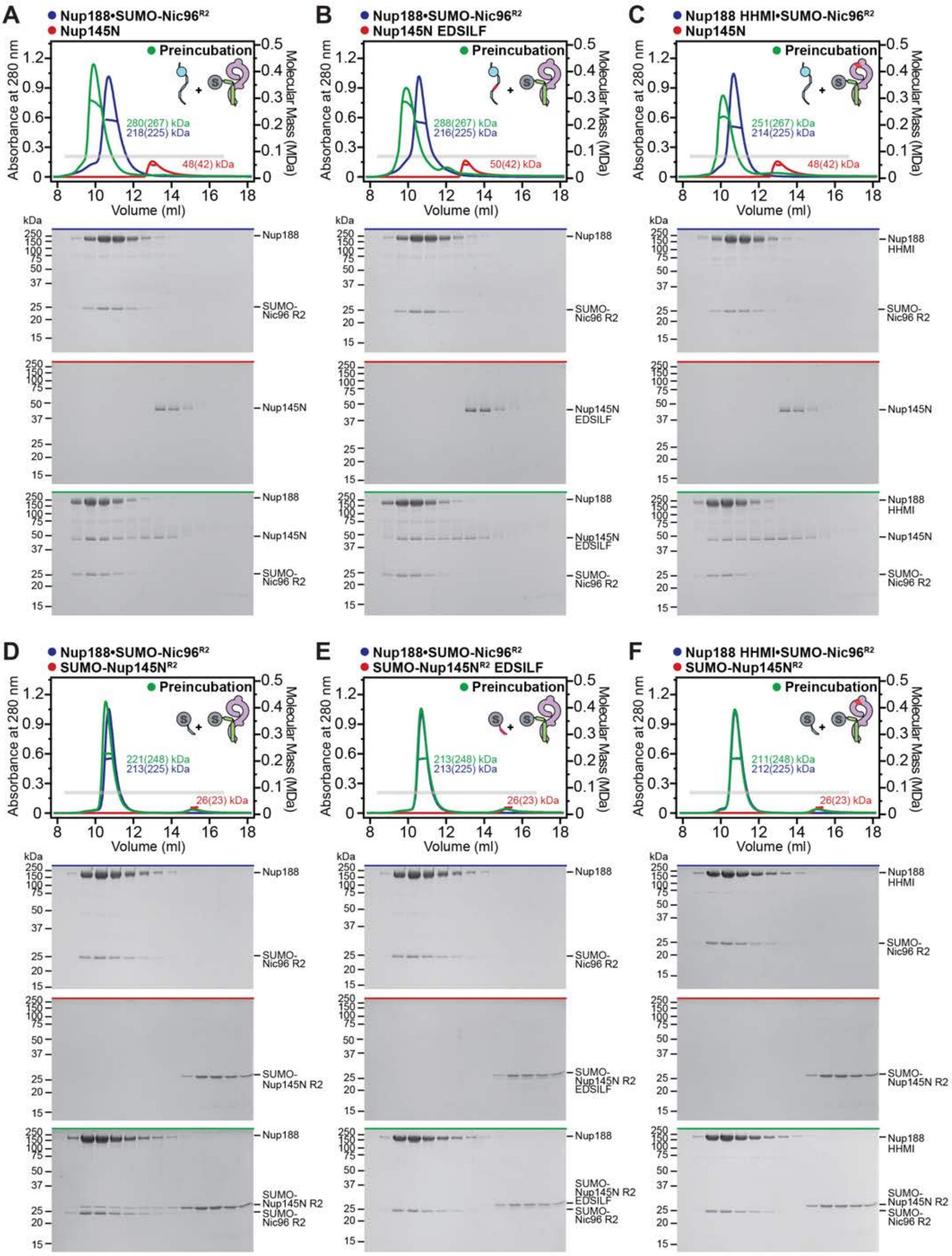

A
Nup188•SUMO-Nic96R2
Nup145N
Preincubation
280(267) kDa
218(225) kDa
48(42) kDa
B
Nup188•SUMO-Nic96R2
Nup145N EDSILF
288(267) kDa
216(225) kDa
50(42) kDa
C
Nup188 HHMI•SUMO-Nic96R2
Nup145N
251(267) kDa
214(225) kDa
48(42) kDa
D
Nup188•SUMO-Nic96R2
SUMO-Nup145NR2
221(248) kDa
213(225) kDa
26(23) kDa
E
Nup188•SUMO-Nic96R2
SUMO-Nup145NR2 EDSILF
213(248) kDa
213(225) kDa
26(23) kDa
F
Nup188 HHMI•SUMO-Nic96R2
SUMO-Nup145NR2
211(248) kDa
212(225) kDa
26(23) kDa
Absorbance at 280 nm
Molecular Mass (MDa)
Volume (ml)
kDa
Nup188
Nup188 HHMI
SUMO-Nic96 R2
Nup145N
Nup145N EDSILF
SUMO-Nup145N R2
SUMO-Nup145N R2 EDSILF

**Fig.S30. SEC-MALS analysis of the Nup188-Nup145N interaction.** SEC-MALS and SDS-PAGE analysis corresponding to Fig. 5F. (**A-F**) SEC-MALS profiles of nups or nup complexes are shown individually (red and blue) and after their preincubation (green) for either wildtype Nup188•SUMO-Nic96$^{R2}$ or mutant Nup188 LIFH•SUMO-Nic96$^{R2}$ binding to either wildtype or EDSILF mutant of Nup145N and SUMO-Nup145N$^{R2}$. Measured molecular masses are indicated for the peak fractions and corresponding theoretical molecular masses are reported in parenthesis. Gray bars indicate fractions that were resolved on SDS-PAGE gels and visualized by Coomassie staining. All SEC profiles were obtained using a Superdex 200 Increase 10/300 GL column. All SEC-MALS results are summarized in Table S4.

**A**

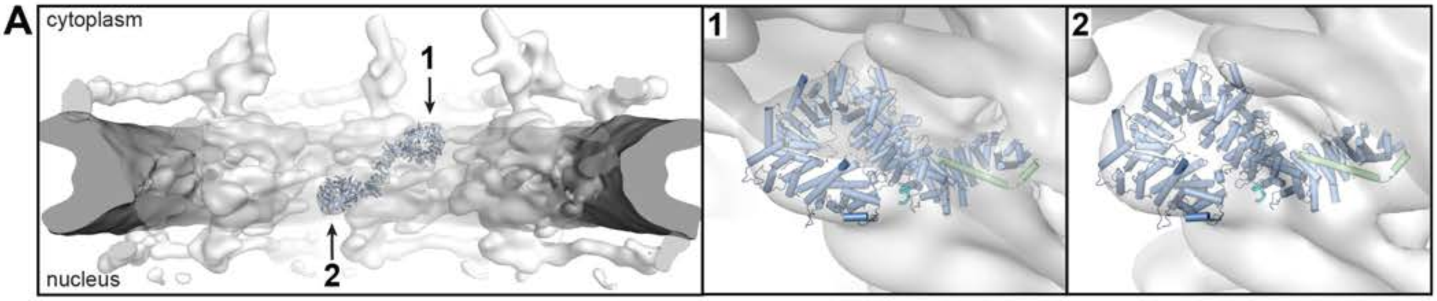

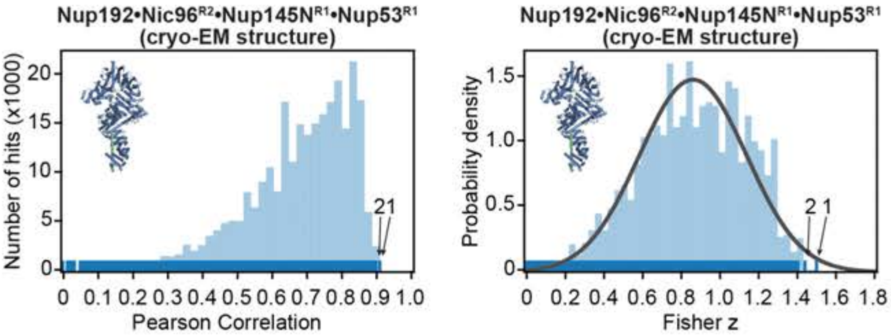

**Cryo-ET map fitting statistics**

| solution rank | Pearson correlation | Fisher z | p-value |
|---|---|---|---|
| 1 | 0.904 | 1.49 | 1.1 e-2 |
| 2 | 0.892 | 1.43 | 1.8 e-2 |

**B**

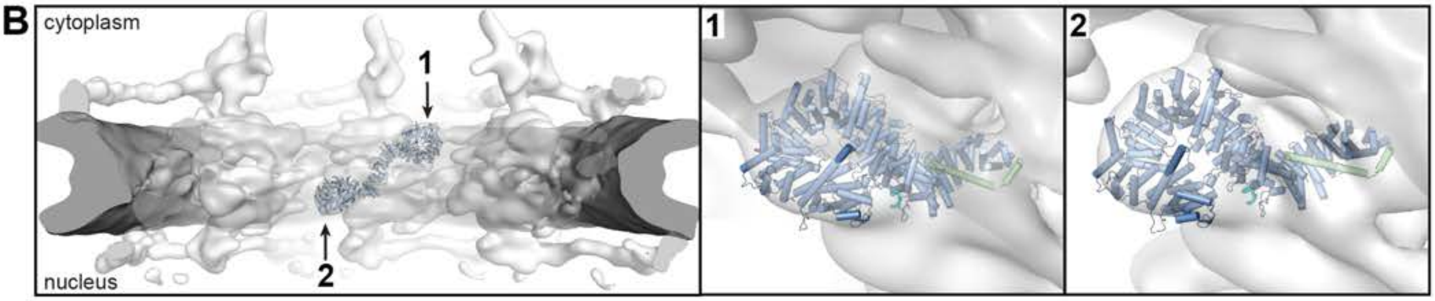

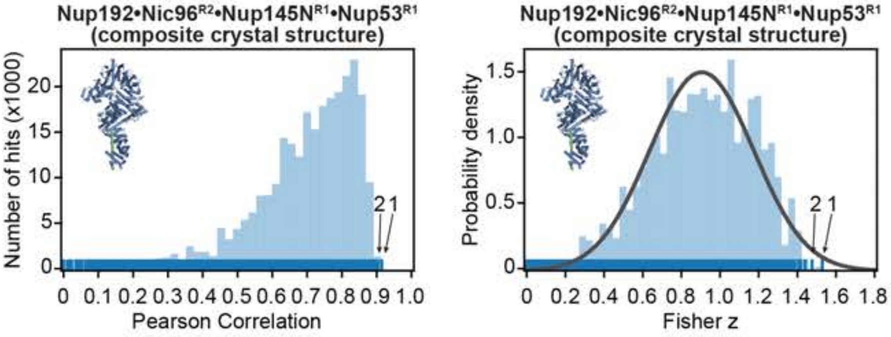

**Cryo-ET map fitting statistics**

| solution rank | Pearson correlation | Fisher z | p-value |
|---|---|---|---|
| 1 | 0.909 | 1.52 | 1.0 e-2 |
| 2 | 0.899 | 1.47 | 1.7 e-2 |

**C**

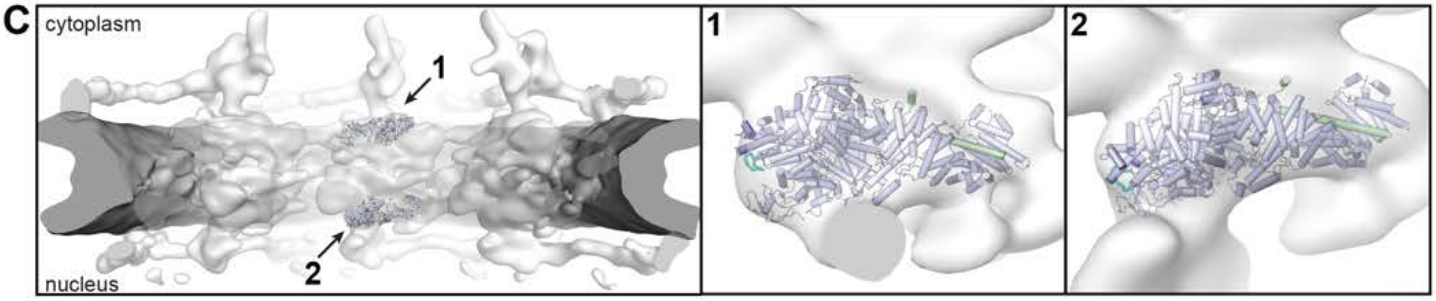

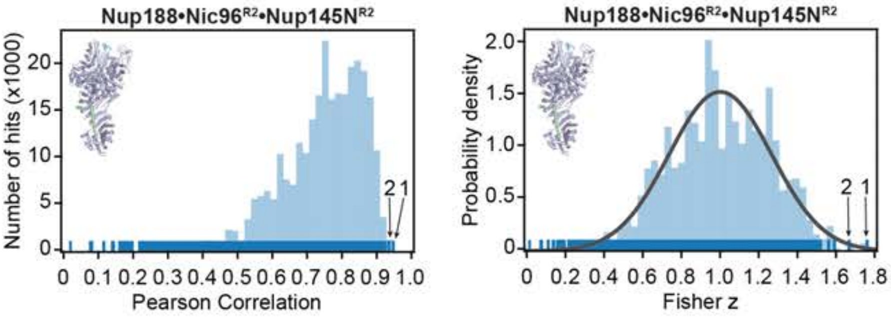

**Cryo-ET map fitting statistics**

| solution rank | Pearson correlation | Fisher z | p-value |
|---|---|---|---|
| 1 | 0.941 | 1.75 | 2.1 e-3 |
| 2 | 0.930 | 1.56 | 5.9 e-3 |

**Supp. Fig. S31**

**Fig.S31. Docking of the Nup192 and Nup188 complex structures into an ~25 Å *in situ* cryo-ET map of the *Saccharomyces cerevisiae* NPC**. Resolution-matched simulated cryo-EM densities of side-chain resolution structures were quantitatively docked into the ~25 Å sub-tomogram averaged cryo-electron tomography (cryo-ET) map of the *in situ* imaged *S. cerevisiae* NPC (EMD-10198) (*31*). The placement of unique solutions in a single spoke is shown (*top left*). Arrows and numbers indicate the accepted solutions and their corresponding rank. Closeup views of the accepted solutions are shown on the right, with the rank indicated on the top left corner of the box. Isosurface representation of the nuclear envelope and the NPC are colored in dark gray and white, respectively. The docked structures are displayed in cartoon representation. Rug plots (blue) and histograms (light blue) of Pearson correlation scores and derived Fisher z scores fit with a normalized Gaussian curve (black) from a global search with 1 million random initial placements are shown (*bottom left* and *middle*). Arrows and numbers indicate accepted solutions and their rank, respectively. A tabular summary of the accepted solutions fitting statistics, along with one-tailed p-values calculated from the Fisher z score distribution, is shown (*bottom right*). The quantitative docking analysis of (**A**) the Nup192•Nic96$^{R2}$•Nup145N$^{R1}$•Nup53$^{R1}$ single particle cryo-EM structure, (**B**) the alternative-conformation Nup192•Nic96$^{R2}$ composite crystal structure with Nup145N$^{R1}$ and Nup53$^{R1}$ peptides included by superposition with the single particle cryo-EM structure, and (**C**) the Nup188•Nic96$^{R2}$•Nup145N$^{R1}$ single particle cryo-EM structure, collocates the Nup192 and Nup188 complexes to the equatorial and peripheral positions of the inner ring, respectively.

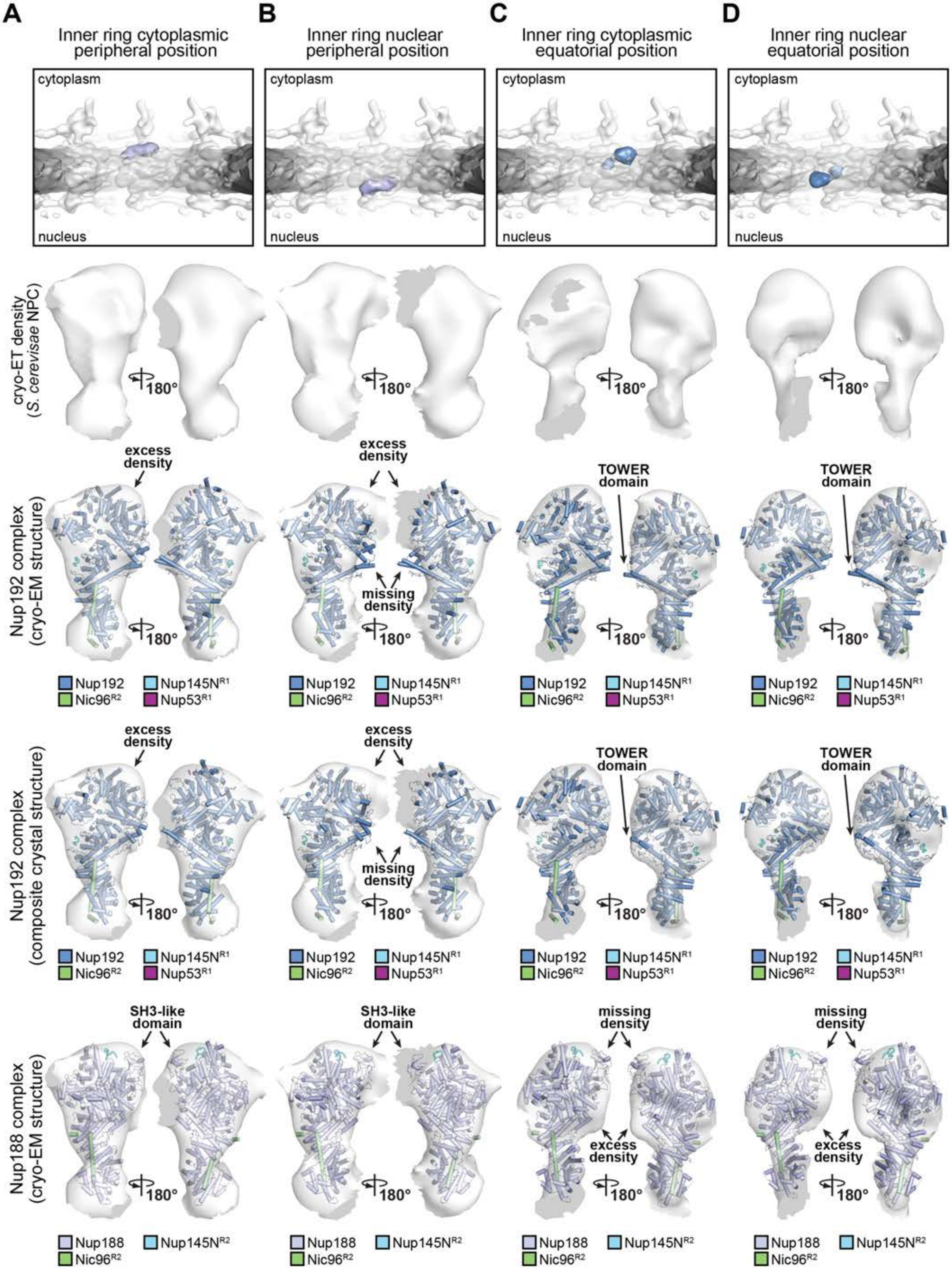
A
Inner ring cytoplasmic peripheral position
B
Inner ring nuclear peripheral position
C
Inner ring cytoplasmic equatorial position
D
Inner ring nuclear equatorial position
cytoplasm
nucleus
cryo-ET density (S. cerevisae NPC)
180°
Nup192 complex (cryo-EM structure)
excess density
missing density
TOWER domain
Nup192
Nup145N$^{R1}$
Nic96$^{R2}$
Nup53$^{R1}$
Nup192 complex (composite crystal structure)
Nup188 complex (cryo-EM structure)
SH3-like domain
Nup188
Nup145N$^{R2}$

Supp. Fig. S32

**Fig.S32. Comparison of the Nup192 and Nup188 complex placement *in situ* cryo-ET map of the *S. cerevisiae* NPC.** Question mark-shaped cryo-ET densities carved out from (**A**) cytoplasmic peripheral, (**B**) nuclear peripheral, (**C**) cytoplasmic equatorial, and (**D**) nuclear equatorial positions of the inner ring of the 25 Å sub-tomogram averaged cryo-ET map of the *in situ* imaged *S. cerevisiae* NPC (EMD-10198) (*31*), where either Nup192•Nic96$^{R2}$•Nup145N$^{R1}$•Nup53$^{R1}$ or Nup188•Nic96$^{R2}$•Nup145N$^{R1}$ were quantitatively docked, as indicated in the boxes (*top*). To compare the goodness of fit by visual inspection, two isosurface represented views of the carved density with cartoon representations of the quantitatively docked composite crystal Nup192, single particle cryo-EM Nup192, or single particle cryo-EM Nup188 complex structures are shown. At the peripheral positions, a tentative placement of the two conformations of the Nup192 complex (single particle cryo-EM and composite crystal structures, respectively) results in unexplained excess density that can be accounted for by the SH3-like domain of Nup188 if the Nup188 complex (single particle cryo-EM structure) is placed in the same position. Unlike the shorter Nup188 Tower, the longer Nup192 Tower exceeds the cryo-ET density when tentatively placed at the peripheral positions. On the contrary, the longer Tower is accommodated by the cryo-ET density at the equatorial position, but the SH3-like domain of the tentatively placed Nup188 exceeds the cryo-ET density at the equatorial position. Overall, the Nup188 $\alpha$-helical solenoid presents a narrower superhelical twist compared to Nup192, which leaves unexplained density if tentatively fit at the equatorial positions. Conversely, the larger superhelical twist of the Nup192 $\alpha$-helical solenoid exceeds the cryo-ET density if tentatively fit at the peripheral positions.

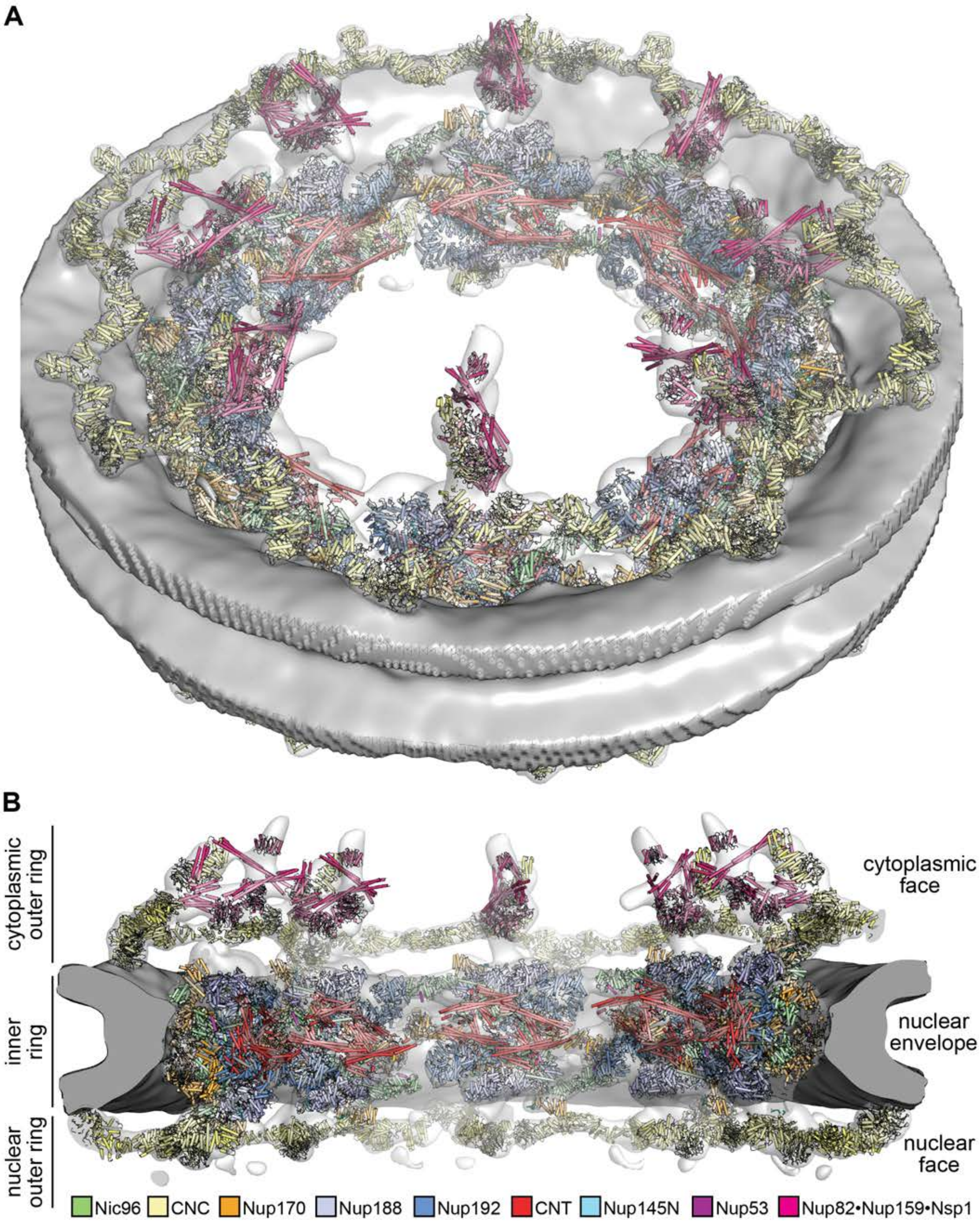
A
B
cytoplasmic outer ring
inner ring
nuclear outer ring
cytoplasmic face
nuclear envelope
nuclear face
Nic96
CNC
Nup170
Nup188
Nup192
CNT
Nup145N
Nup53
Nup82•Nup159•Nsp1

**Fig.S33. Composite structure of the *S. cerevisiae* NPC.** Linker-scaffold structures were docked into an ~25 Å sub-tomogram averaged cryo-ET map of the *in situ* imaged *S. cerevisiae* NPC (EMD-10198) to obtain a composite structure of the *S. cerevisiae* NPC complete with surface-bound linker nup segments. The crystal composite Nup192•Nic96$^{R2}$•Nup145N$^{R1}$•Nup53$^{R1}$ structure and the single particle cryo-EM Nup188•Nic96$^{R2}$•Nup145N$^{R2}$ structure were quantitatively docked (fig. S30). Crystal structures of linker-scaffold complexes Nup170•Nup145N$^{R3}$•Nup53$^{R3}$ (full-length crystal composite structures) (*30*) and CNT•Nic96$^{R1}$ (PDB ID 5CWS) (*25*) were first aligned to the composite structure from (*31*) and then locally rigid-body fit into the cryo-ET map. The structures of the coat nup complex (CNC) and the Nup82•Nup159•Nsp1 complex were carried over from the composite structure from (*31*). (**A**) View from above the cytoplasmic face and (**B**) a cross-sectional view from the central transport channel. Isosurface representations of the nuclear envelope and the NPC are colored in dark gray and white, respectively. The docked structures are displayed in cartoon representation.

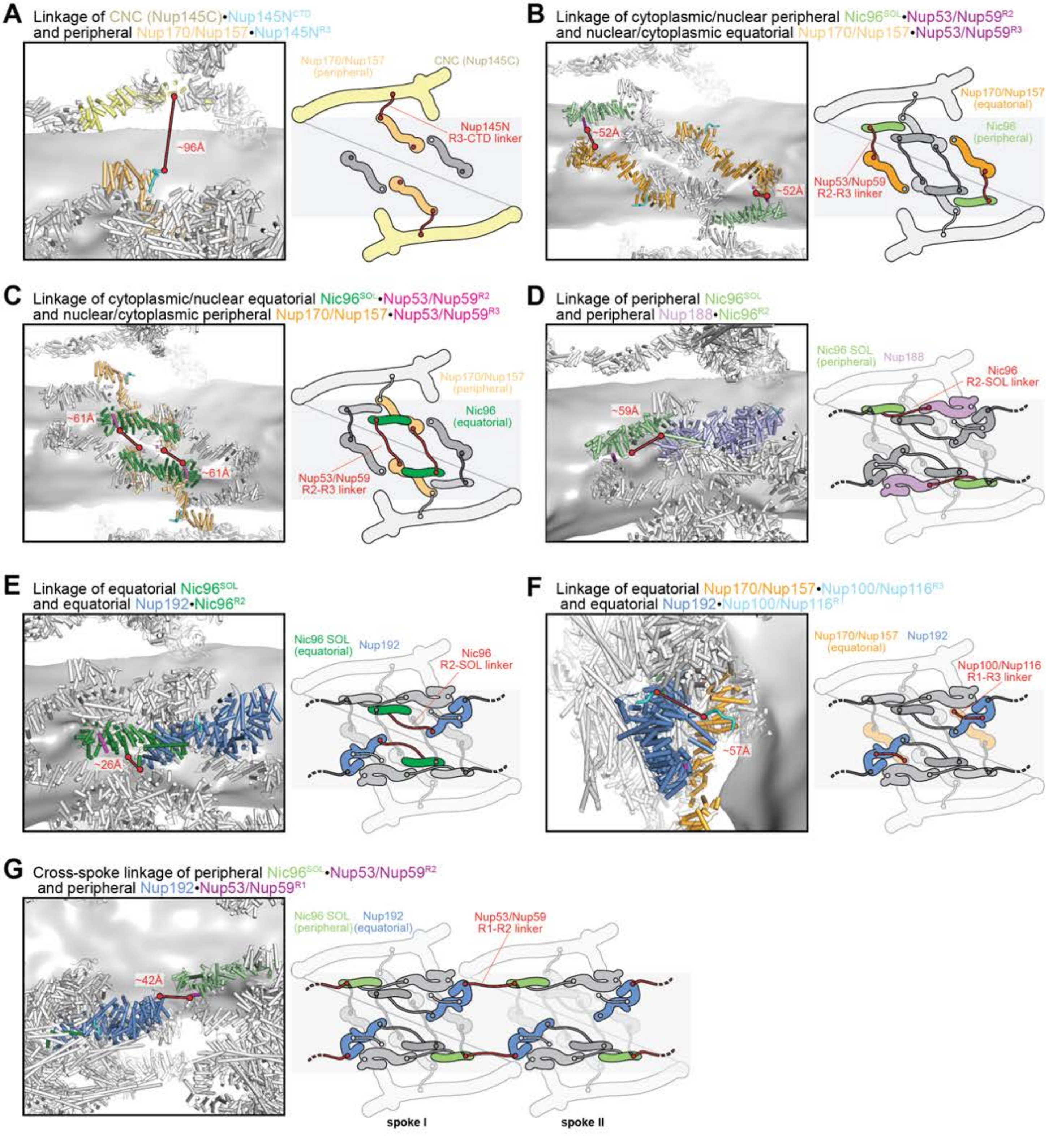

Supp. Fig. S34a

**H** Linkage of peripheral Nup188•Nic96$^{R2}$ and peripheral CNT•Nic96$^{R1}$

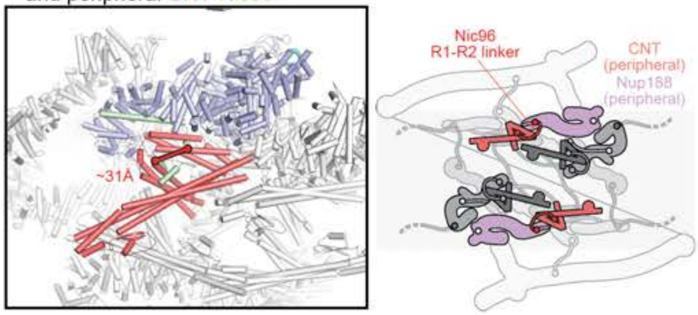

**I** Linkage of equatorial Nup192•Nic96$^{R2}$ and equatorial CNT•Nic96$^{R1}$

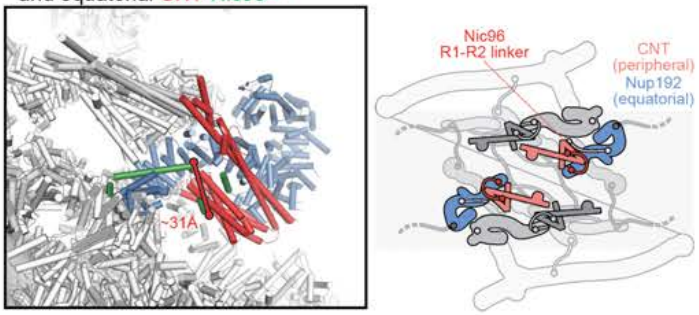

**J** Linkage of peripheral Nup188•Nup100/Nup116$^{R2}$ and distal Nup100/Nup116$^{CTD}$•Nup82$^{NTD}$

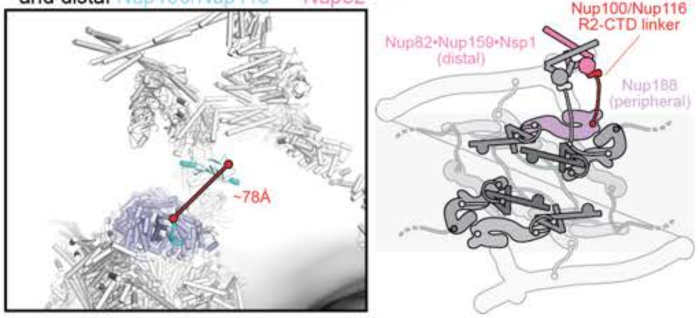

**K** Linkage of equatorial Nup170/Nup157•Nup100/Nup116$^{R3}$ and proximal Nup100/Nup116$^{CTD}$•Nup82$^{NTD}$

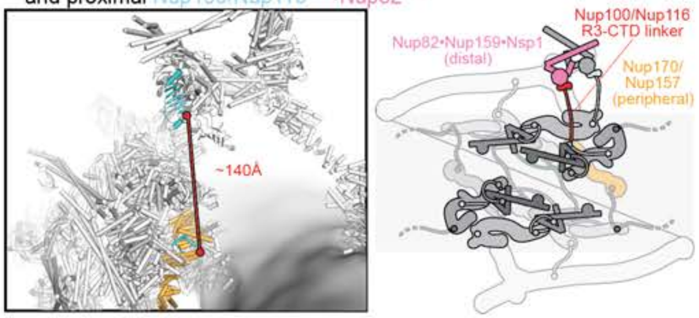

**Fig.S34. Shortest-distance analysis of linker network in the *S. cerevisiae* NPC.** Spatial distances in the composite structure of the *S. cerevisiae* NPC were measured between two copies of linker nup segments that are adjacent in their primary sequence. (**A-K**) In the box on the left, linker nup segments and the scaffold surface that they are bound to are shown in cartoon representation and highlighted in color. The remainder of nups are colored in white. The nuclear envelope is rendered as dark gray isosurface. The straight-line linkage between the two closest copies of linker nup segments adjacent in their primary sequence is indicated by a solid red line. The measured distance is reported in red. On the right, a schematic representation of the scaffold nup binding sites (highlighted in color) and the connecting linker nup segment (red), illustrates the topology of the linkage within the context of an entire NPC spoke (remainder of nups shown in gray).

**Nup145N**

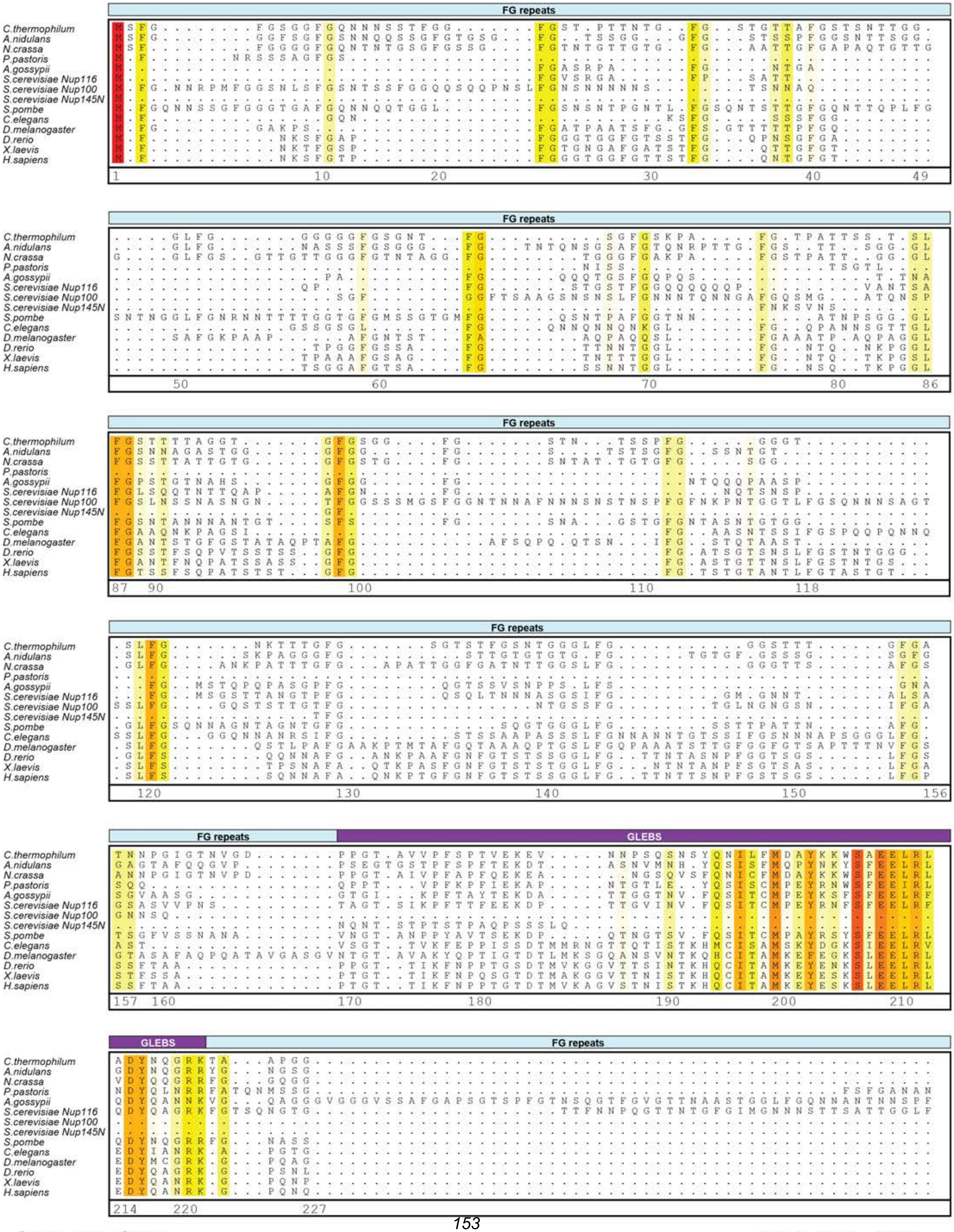

Supp. Fig. S35a

Hoelz et al., 2021

### Nup145N

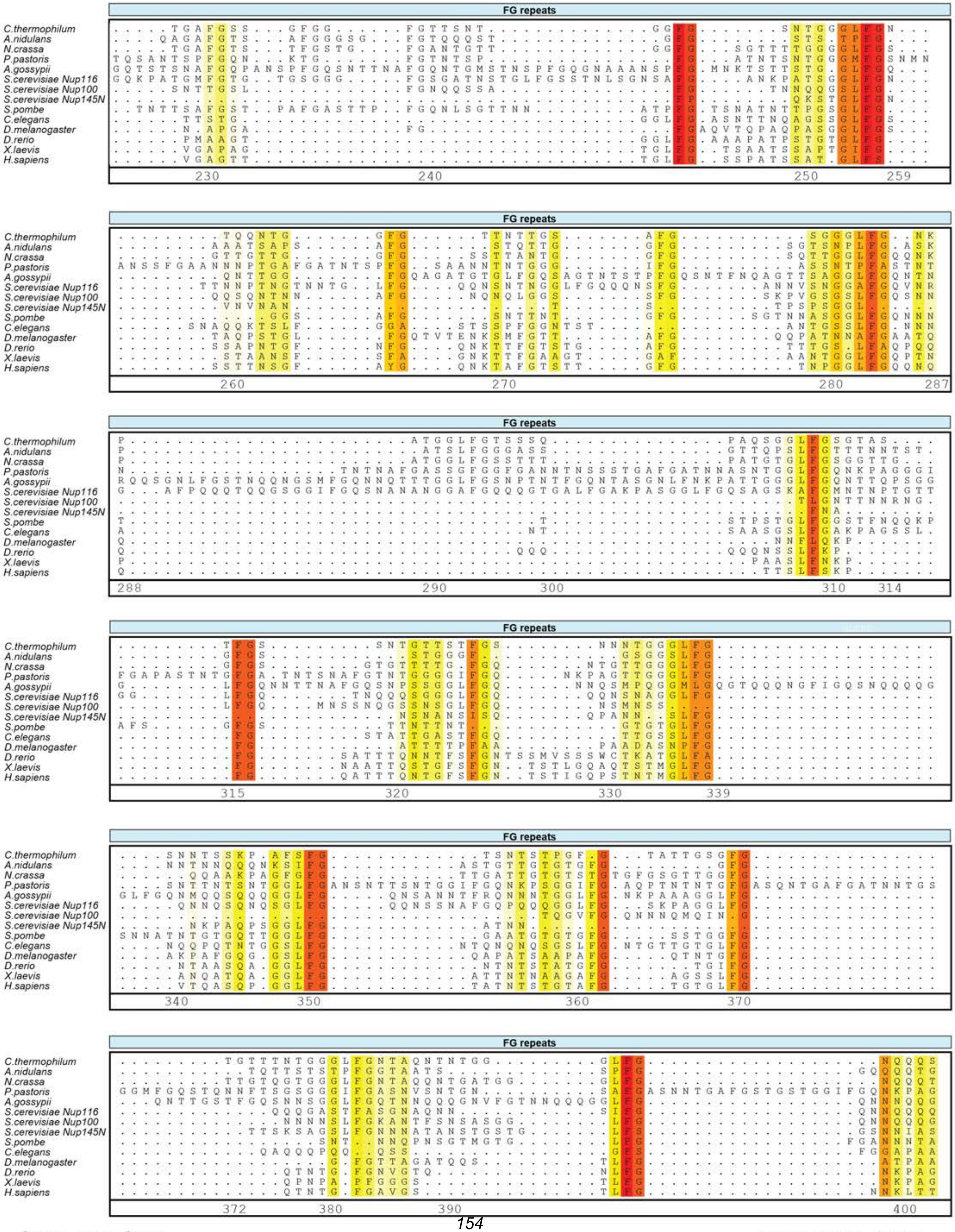
FG repeats
C.thermophilum
A.nidulans
N.crassa
P.pastoris
A.gossypii
S.cerevisiae Nup116
S.cerevisiae Nup100
S.cerevisiae Nup145N
S.pombe
C.elegans
D.melanogaster
D.rerio
X.laevis
H.sapiens

**Nup145N**

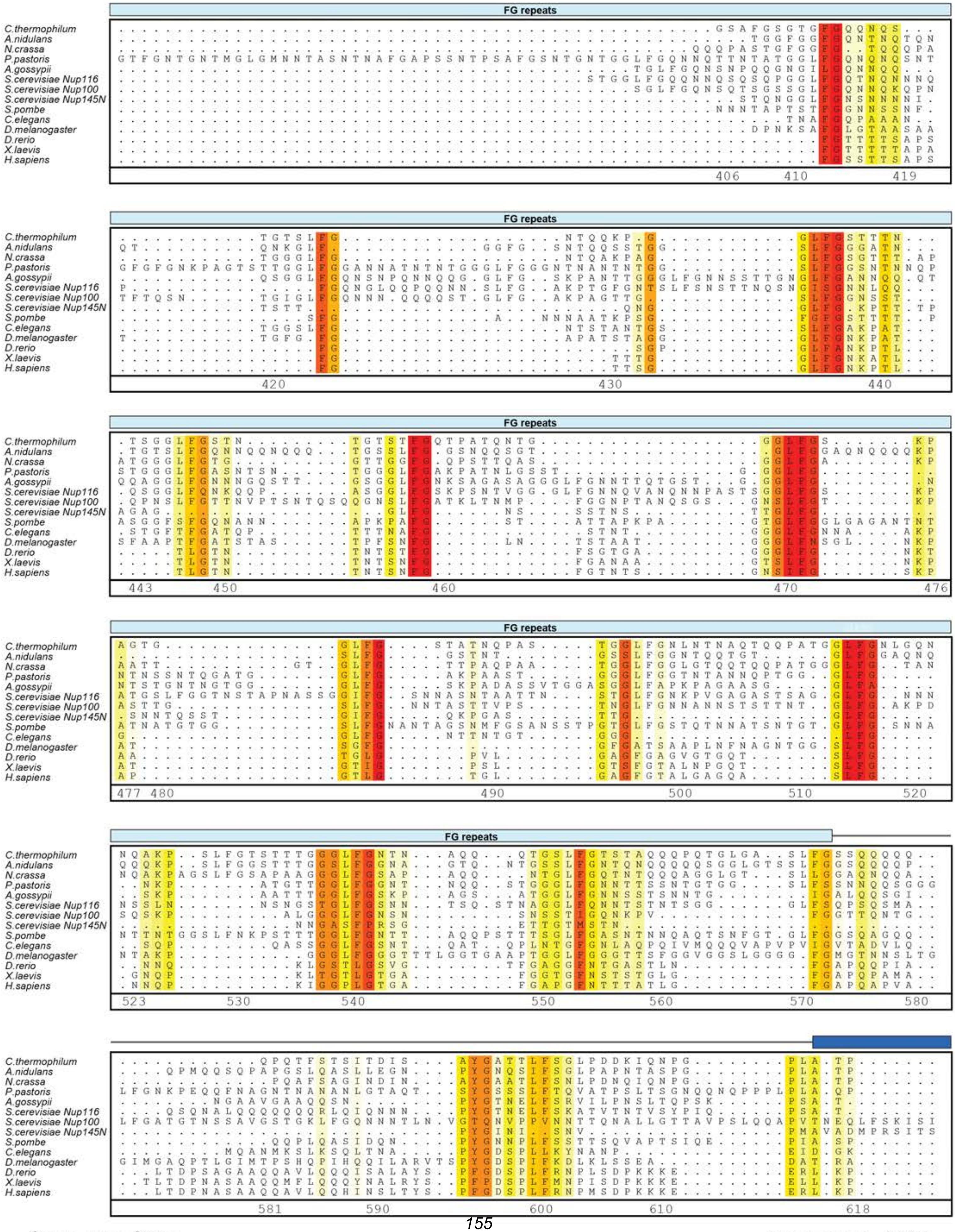

**Supp. Fig. S35c**

**Hoelz et al., 2021**

**Nup145N**

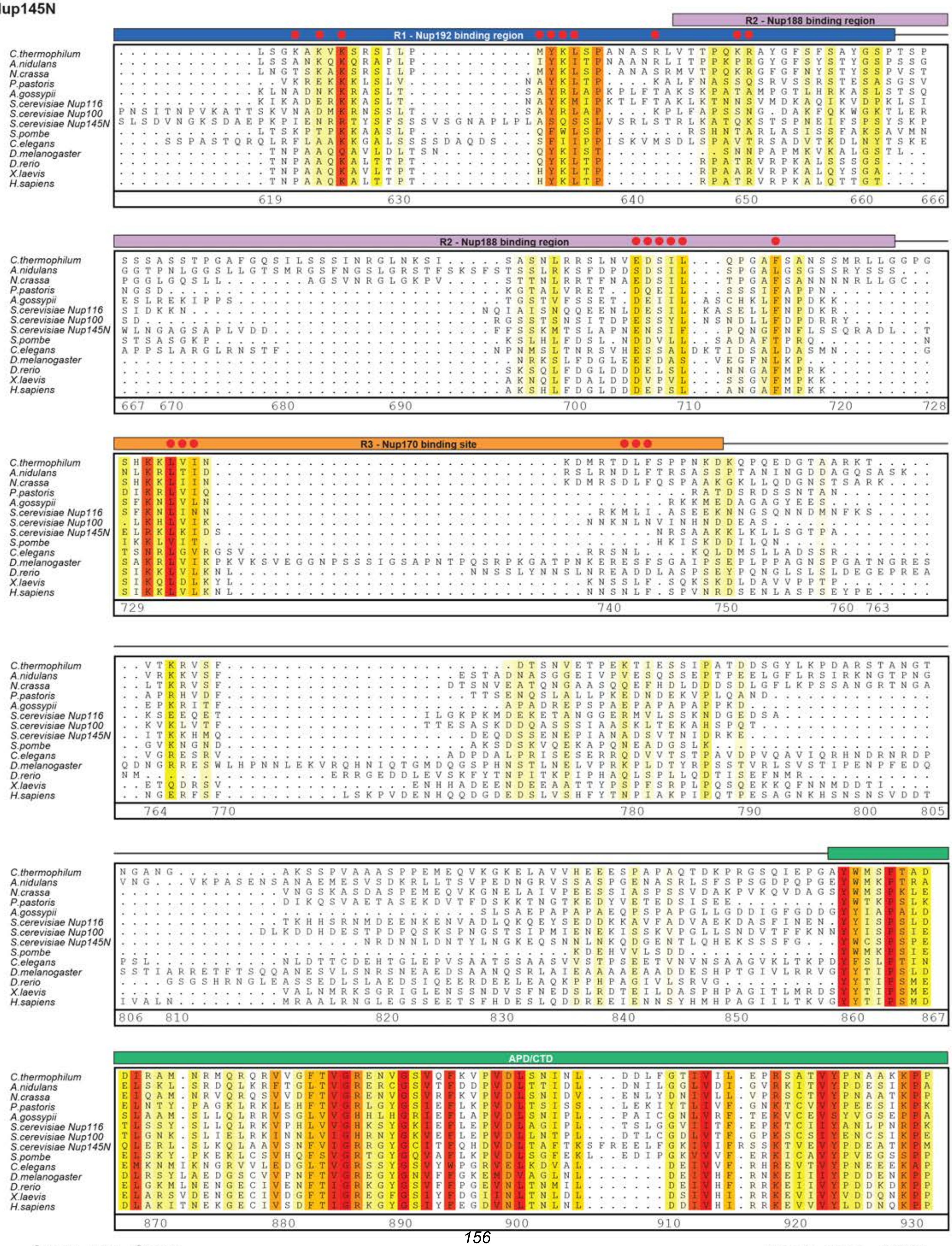

**Supp. Fig. S35d**

Hoelz et al., 2021

**Nup145N**

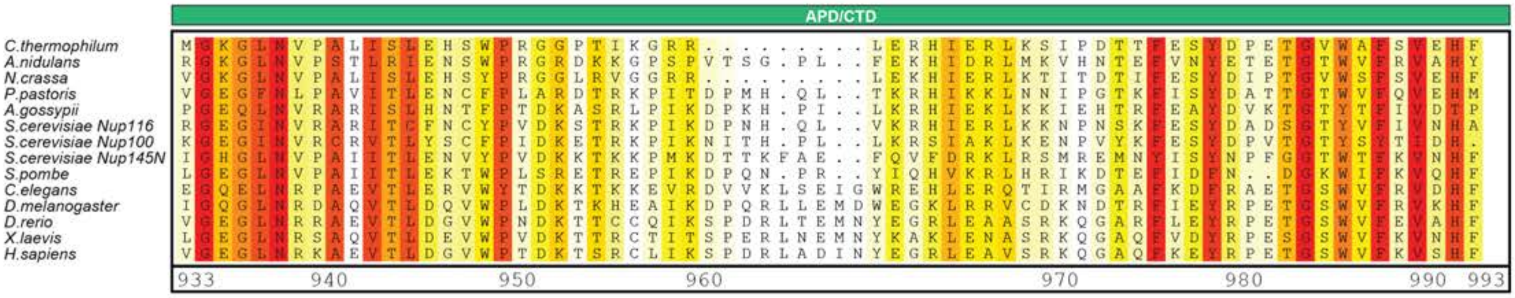
APD/CTD
C.thermophilum
A.nidulans
N.crassa
P.pastoris
A.gossypii
S.cerevisiae Nup116
S.cerevisiae Nup100
S.cerevisiae Nup145N
S.pombe
C.elegans
D.melanogaster
D.rerio
X.laevis
H.sapiens
933
940
950
960
970
980
990
993

**Fig.S35. Multispecies sequence alignment of Nup145N.** Sequences from twelve diverse species, including three *S. cerevisiae* paralogs Nup100, Nup116, and Nup145N, were aligned and colored by sequence similarity according to the BLOSUM62 matrix from white (less than 55 % similarity), to yellow (55 % similarity), to red (100 % identity). Numbering below alignment is relative to the *C. thermophilum* sequence. Regions of the sequence are annotated above the alignment: FG/GLFG repeats (pale cyan), Gle2-binding sequence – GLEBS (purple), R1 – Nup192 binding region (blue), R2 – Nup188 binding region (light purple), R3 – Nup170 binding region (orange), autoproteolytic domain – APD (green). Residues found by mutational analysis to affect Nup192 binding (Fig. 4, figs. S15, S18, and S19), Nup188 binding (Figs. 5, Fig. S28, S29, and S30), or Nup170 binding (*30*) are indicated by red circles above the alignment.

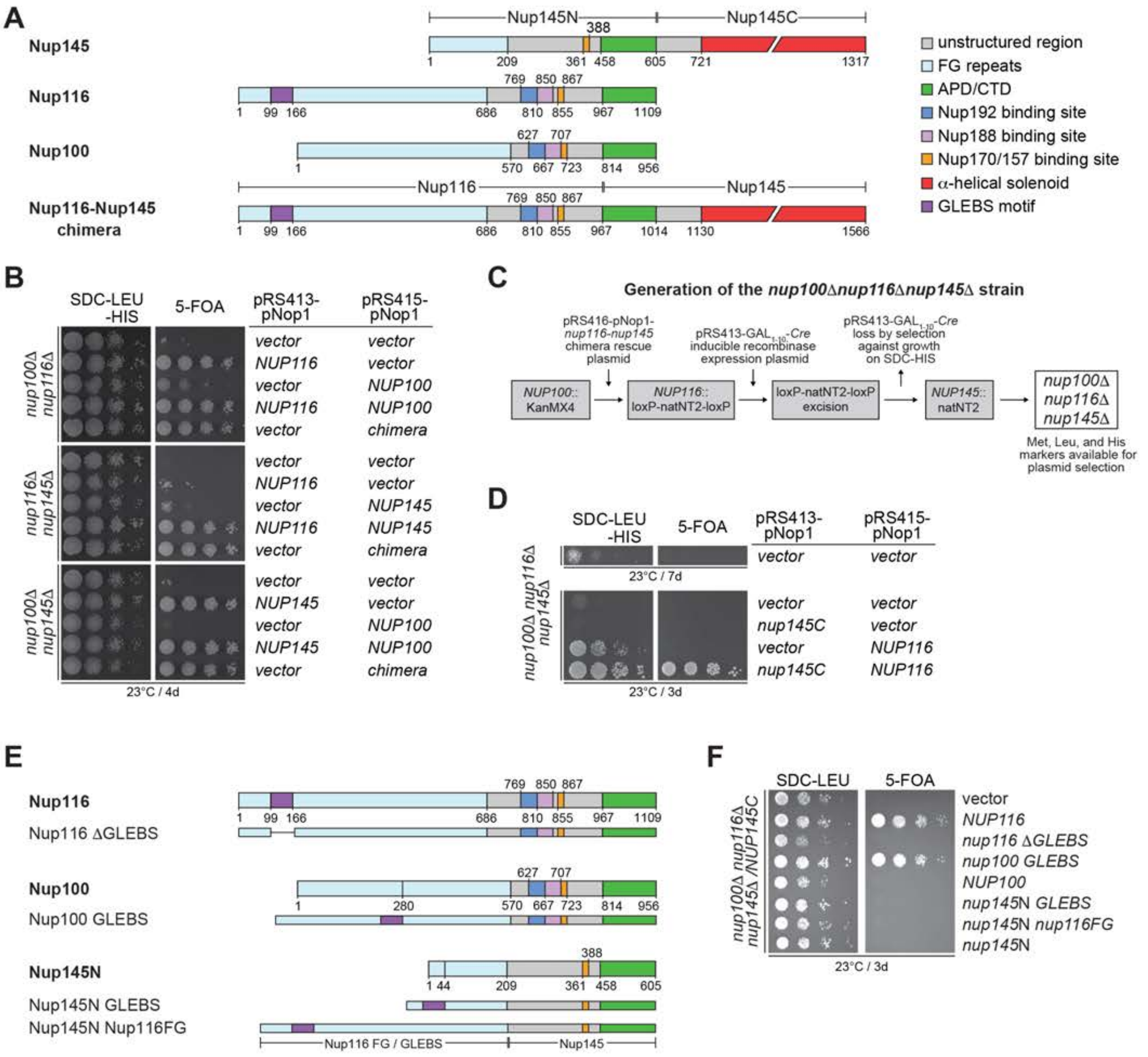
A
Nup145N
Nup145C
Nup145
Nup116
Nup100
Nup116
Nup145
Nup116-Nup145 chimera
unstructured region
FG repeats
APD/CTD
Nup192 binding site
Nup188 binding site
Nup170/157 binding site
α-helical solenoid
GLEBS motif
B
SDC-LEU -HIS
5-FOA
pRS413-pNop1
pRS415-pNop1
nup100Δ nup116Δ
nup116Δ nup145Δ
nup100Δ nup145Δ
vector
NUP116
NUP100
NUP145
chimera
23°C / 4d
C
Generation of the nup100Δnup116Δnup145Δ strain
pRS416-pNop1-nup116-nup145 chimera rescue plasmid
pRS413-GAL1-10-Cre inducible recombinase expression plasmid
pRS413-GAL1-10-Cre loss by selection against growth on SDC-HIS
NUP100::KanMX4
NUP116::loxP-natNT2-loxP
loxP-natNT2-loxP excision
NUP145::natNT2
nup100Δ nup116Δ nup145Δ
Met, Leu, and His markers available for plasmid selection
D
SDC-LEU -HIS
5-FOA
pRS413-pNop1
pRS415-pNop1
nup100Δ nup116Δ nup145Δ
vector
nup145C
NUP116
23°C / 7d
23°C / 3d
E
Nup116
Nup116 ΔGLEBS
Nup100
Nup100 GLEBS
Nup145N
Nup145N GLEBS
Nup145N Nup116FG
Nup116 FG / GLEBS
Nup145
F
SDC-LEU
5-FOA
nup100Δ nup116Δ nup145Δ /NUP145C
vector
NUP116
nup116 ΔGLEBS
nup100 GLEBS
NUP100
nup145N GLEBS
nup145N nup116FG
nup145N
23°C / 3d

**Fig.S36. Generation of the *nup100Δnup116Δnup145Δ* *S. cerevisiae* knockout strain.** (**A**) Domain structure of the *S. cerevisiae* gene products: Nup145, the Nup145N paralogs Nup100 and Nup116, and the Nup116-Nup145C chimera construct. (**B**) Complementation of double knockout strains by co-transformation with a combination of plasmids expressing Nup100, Nup116, Nup145N-Nup145C, or the Nup116-Nup145C chimera. The Nup116-Nup145C chimera is a single construct that complements all double knockout combinations. (**C**) Gene knockout strategy employed in the generation of the *nup100Δnup116Δnup145Δ* strain. The Nup116-Nup145C chimera complements sequential knockouts of *NUP100*, *NUP116*, and *NUP145* by homologous recombination with selection marker cassettes. The inducible expression of Cre results in the efficient excision of the loxP-natNT2-loxP cassette, allowing for a second round of selection using the natNT2 cassette. (**D**) The *nup100Δnup116Δnup145Δ* strain is rescued by the Nup116-Nup145C chimera construct but presents a slow growth phenotype. Growth can be restored to wildtype rates by co-transforming the strain with plasmids expressing Nup145C and Nup116. (**E**) Domain structure of the *S. cerevisiae* Nup145N paralogs and artificial constructs. Nup116 is the only paralog that possesses a GLEBS motif. To test if the GLEBS motif is the sole determinant of viable *nup100Δnup116Δnup145Δ* complementation by plasmids expressing Nup145N paralogs, constructs with the Nup116 GLEBS motif sequence inserted into the Nup100 and Nup145N FG repeat region, as well as a Nup145N construct in which the FG repeat region is replaced with the Nup116 FG-GLEBS repeat region, were generated. (**F**) Viability analysis of the *nup100Δnup116Δnup145Δ* strain carrying a plasmid expressing Nup145C, after transformation with indicated Nup145N paralog constructs and selection of ten-fold serial dilution series spotted onto either SDC-LEU and SDC+5-FOA plates (shuffling).

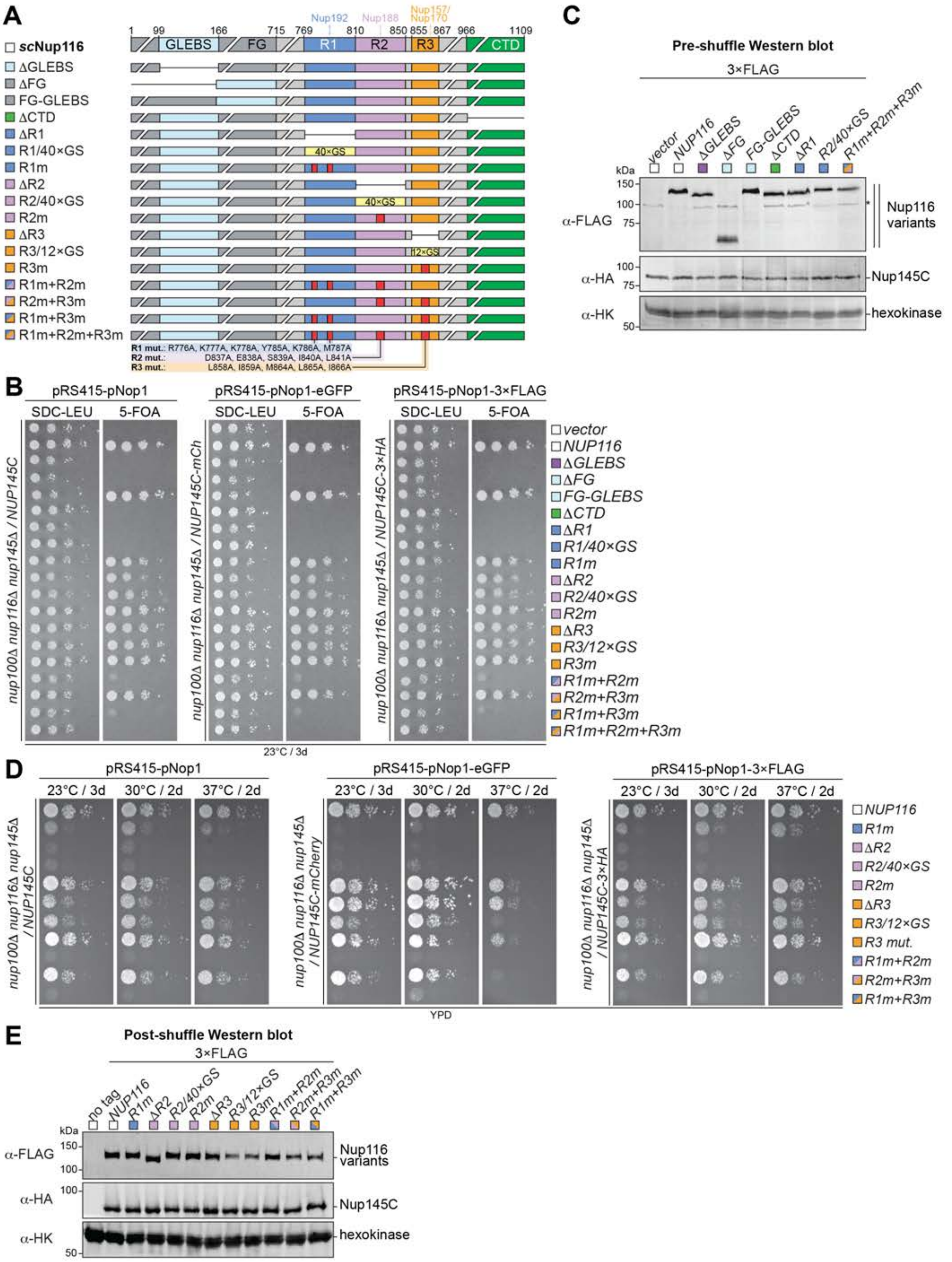

Supp. Fig. S37a

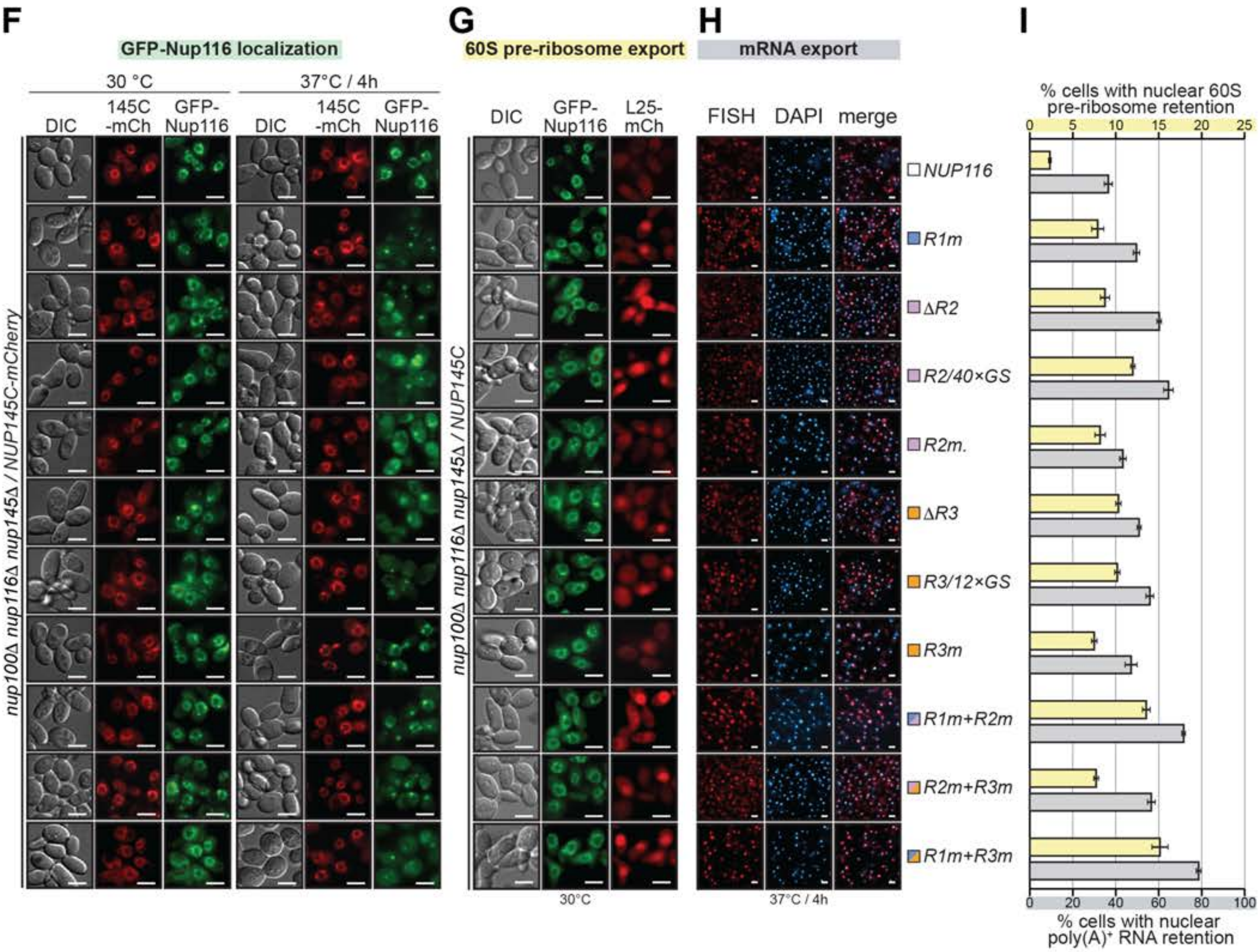
F
GFP-Nup116 localization
30 °C
37°C / 4h
DIC
145C -mCh
GFP-Nup116
nup100Δ nup116Δ nup145Δ / NUP145C-mCherry
G
60S pre-ribosome export
DIC
GFP-Nup116
L25-mCh
nup100Δ nup116Δ nup145Δ / NUP145C
30°C
H
mRNA export
FISH
DAPI
merge
37°C / 4h
NUP116
R1m
ΔR2
R2/40×GS
R2m.
ΔR3
R3/12×GS
R3m
R1m+R2m
R2m+R3m
R1m+R3m
I
% cells with nuclear 60S pre-ribosome retention
0
5
10
15
20
25
0
20
40
60
80
100
% cells with nuclear poly(A)+ RNA retention

**Fig.S37. *In vivo* analysis of Nup116 functional regions and validation of interactions with scaffold proteins.** Supporting data for analysis presented in Fig. 10. (**A**) Domain structure of Nup116 variants introduced into the *nup100Δnup116Δnup145Δ S. cerevisiae* strain, with colored squares indicating the functional region targeted by each mutation: GLEBS (pale cyan); FG repeats (dark gray); R1m, Nup192 binding region (blue); R2m, Nup188 binding region (light purple); R3m, Nup157/Nup170 binding region (orange); R1m+R2m (blue and light purple); R2m+R3m (light purple and orange); R1m+R3m (blue and orange); R1m+R2m+R3m (blue, light purple, and orange); CTD (green). (**B**) Viability analysis of the *nup100Δnup116Δnup145Δ* strains after transformation with indicated Nup116 variants and selection of ten-fold serial dilution series spotted onto either SDC-LEU and SDC+5-FOA plates (shuffling). Untagged or N-terminally tagged with 3×FLAG or eGFP were introduced in strains carrying plasmids expressing Nup145C, Nup145C-3×HA, and Nup145C-mCherry, respectively. (**C**) Western blot analysis of expression levels of 3×FLAG-tagged non-rescuing Nup116 variants in unshuffled *nup100Δnup116Δnup145Δ* strain transformants selected in SDC-LEU media. 3×FLAG-Nup116 variants, Nup145C-3×HA, and the endogenous hexokinase loading control were detected with anti-FLAG, anti-HA, and anti-hexokinase antibodies, respectively. (**D**) Growth analysis at different temperatures of ten-fold serial dilution series of the shuffled *nup100Δnup116Δnup145Δ* strains carrying indicated Nup116 variants, replicated for untagged and N-terminally 3×FLAG or eGFP-tagged constructs. (**E**) Western blot analysis of expression levels of 3×FLAG-tagged Nup116 variants in shuffled *nup100Δnup116Δnup145Δ* strain transformants selected on SDC+5-FOA media and subsequently grown in YPD media at 30 °C. 3×FLAG-Nup116 variants, Nup145C-3×HA, and the endogenous hexokinase loading control were detected with anti-FLAG, anti-HA, and anti-hexokinase antibodies, respectively. (**F**) Subcellular localization at different temperatures eGFP-Nup116 variants in a shuffled *nup100Δnup116Δnup145Δ* strain carrying a plasmid expressing Nup145C-mCherry. (**G**) Subcellular localization of the 60S pre-ribosomal export reporter Rpl25-mCherry and eGFP-Nup116 variants in a shuffled *nup100Δnup116Δnup145Δ* strain. (**H**) Poly(A)$^+$ RNA localization detected by FISH in a shuffled *nup100Δnup116Δnup145Δ* strain carrying plasmids expressing Nup116 variants and grown at 37 °C for 4 hours. All scalebars are 5 μm. (**I**) Quantitation of the 60S pre-ribosomal and Poly(A)$^+$ RNA retention in the nucleus. All experiments were performed in triplicate and mean and its standard error are reported.

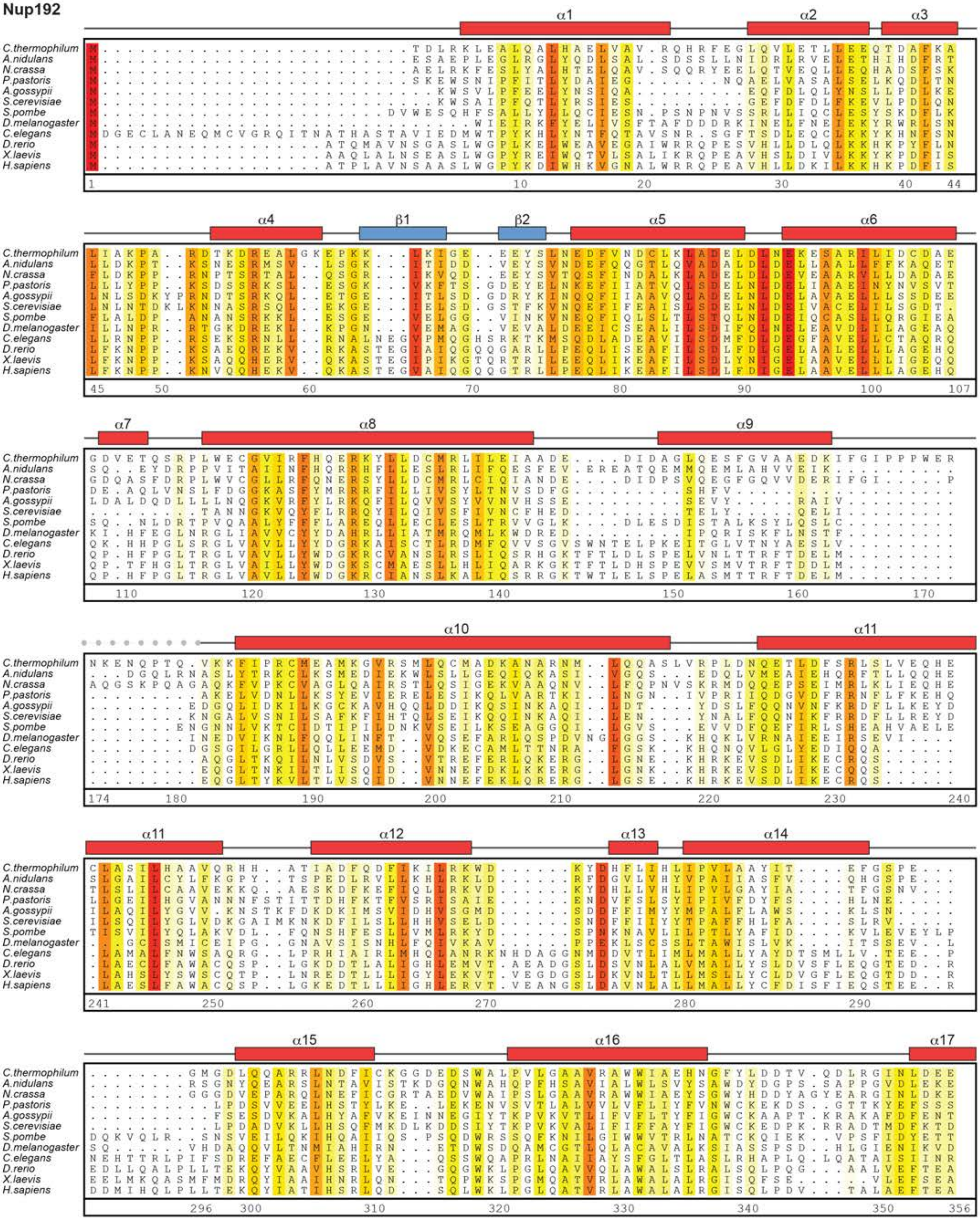
Nup192
α1
α2
α3
α4
β1
β2
α5
α6
α7
α8
α9
α10
α11
α12
α13
α14
α15
α16
α17
C.thermophilum
A.nidulans
N.crassa
P.pastoris
A.gossypii
S.cerevisiae
S.pombe
D.melanogaster
C.elegans
D.rerio
X.laevis
H.sapiens

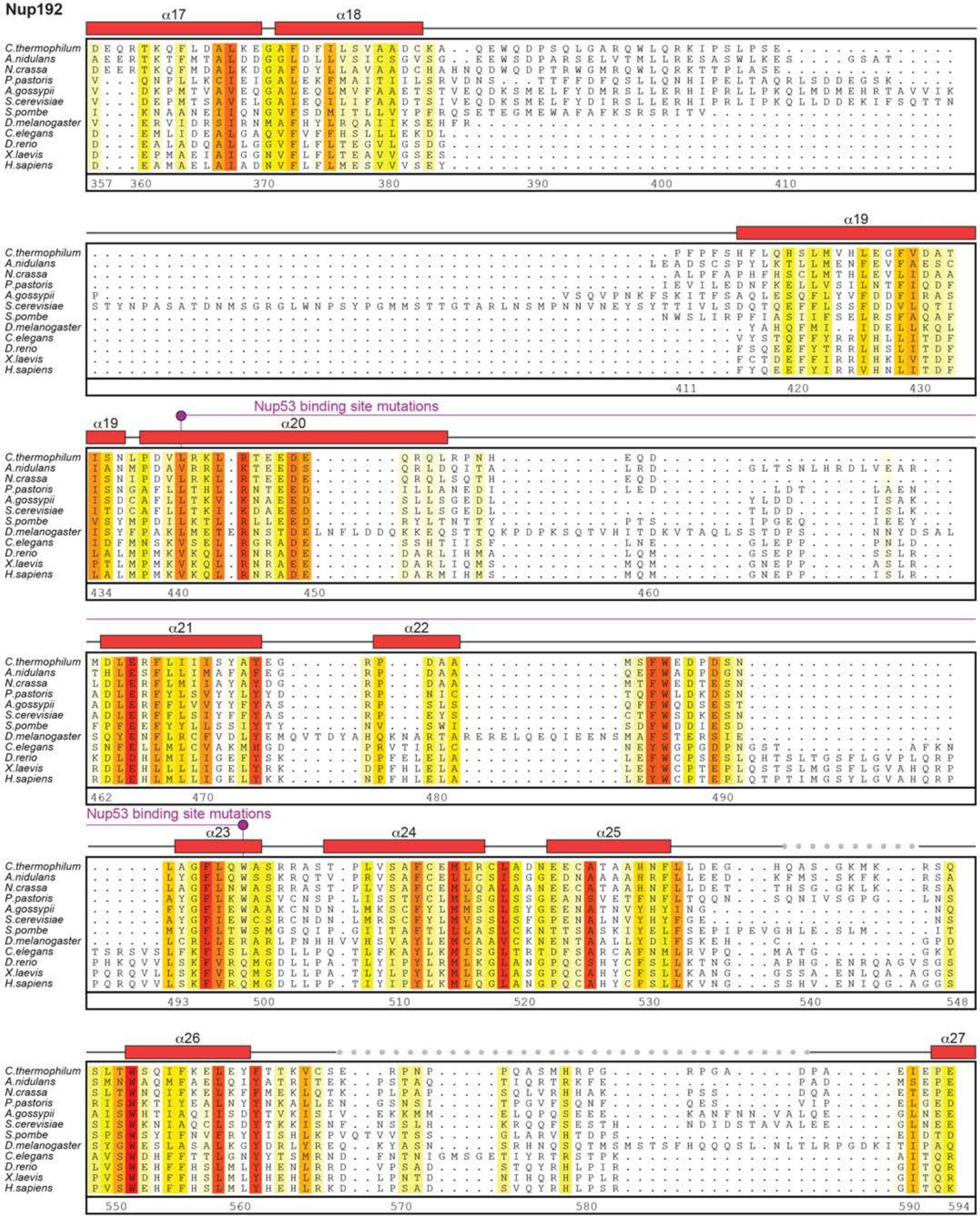

Supp Fig. S38b

Hoelz et al., 2021

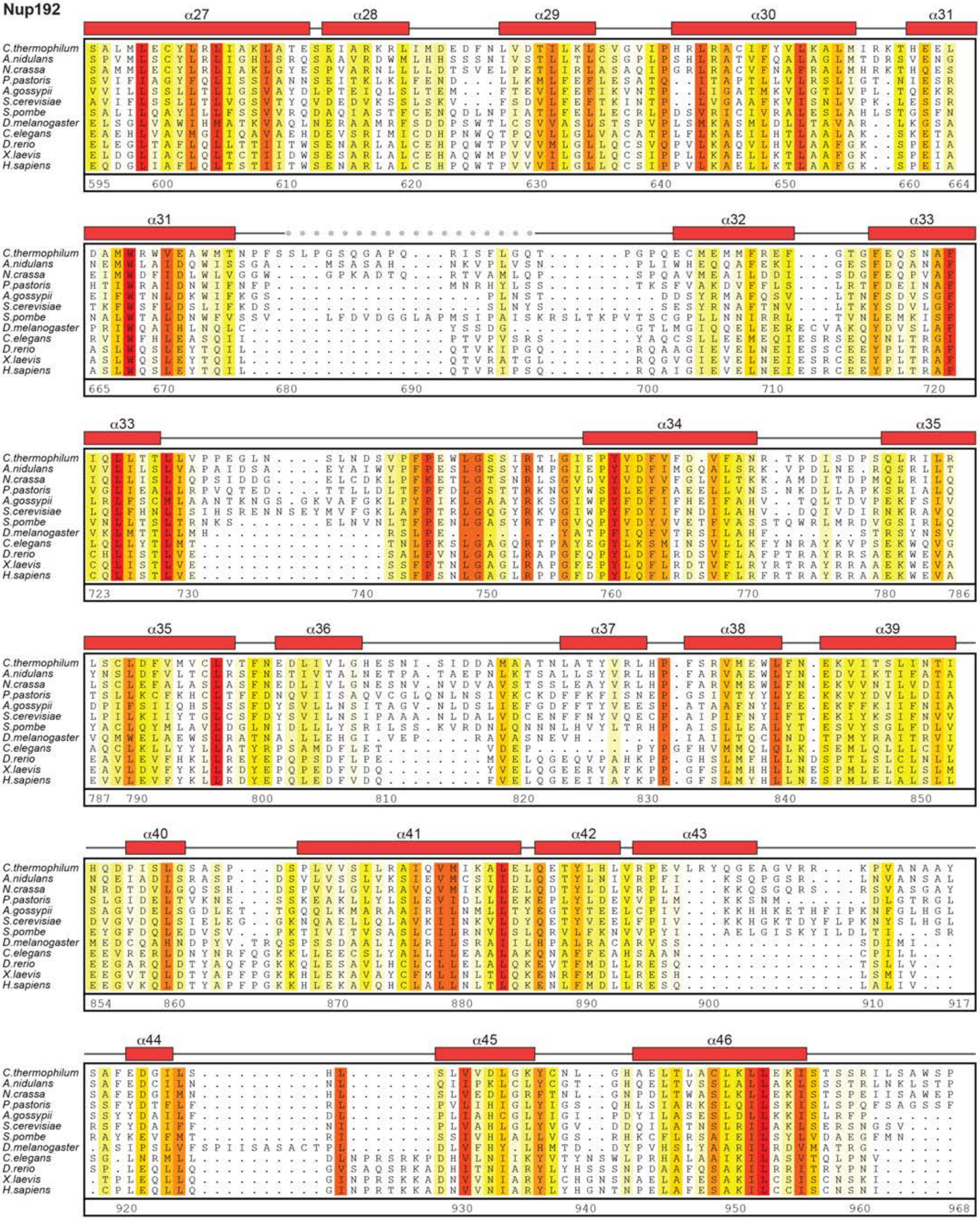

Supp Fig. S38c

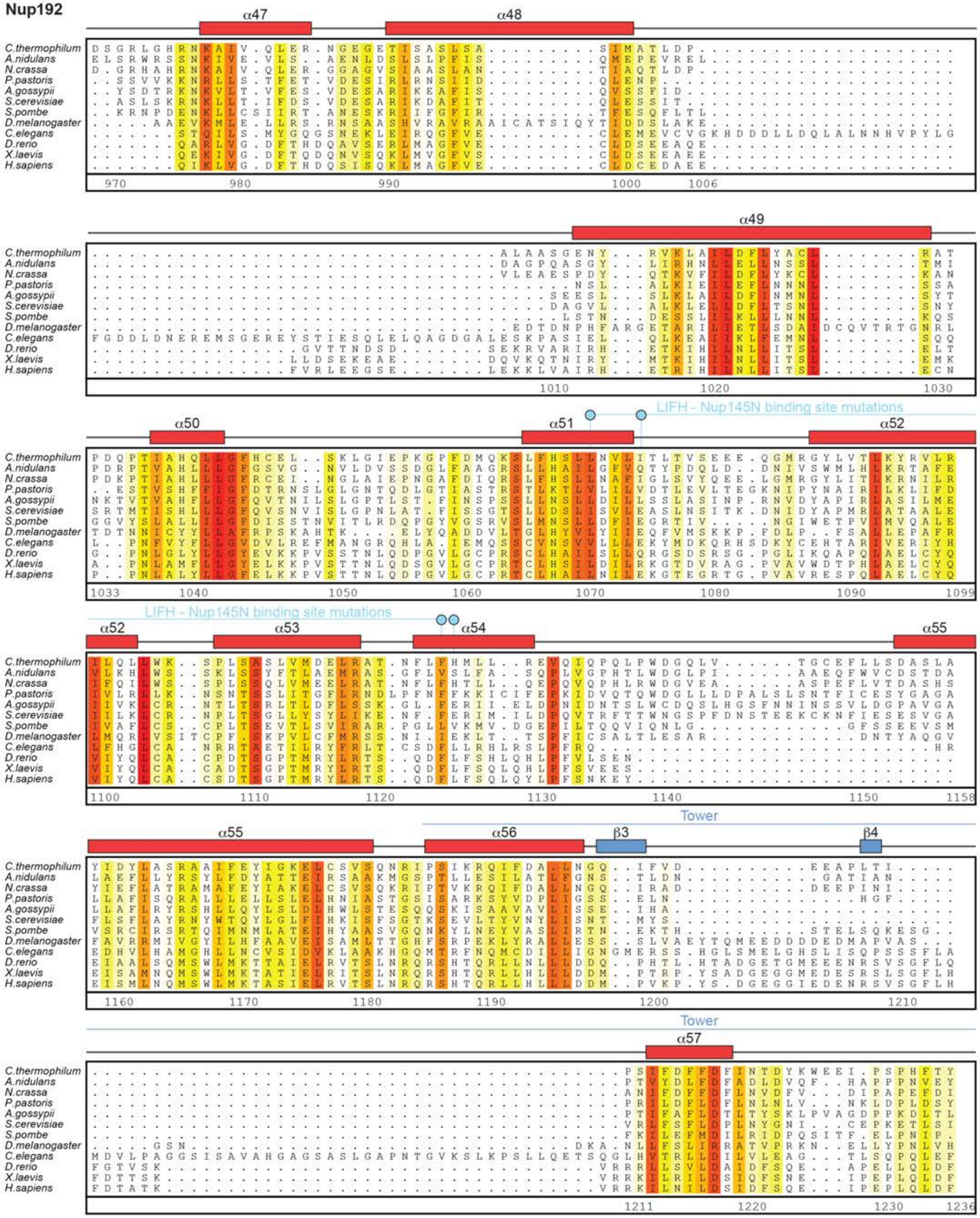
Nup192
α47
α48
α49
α50
α51
α52
α53
α54
α55
α56
α57
β3
β4
LIFH - Nup145N binding site mutations
Tower
C.thermophilum
A.nidulans
N.crassa
P.pastoris
A.gossypii
S.cerevisiae
S.pombe
D.melanogaster
C.elegans
D.rerio
X.laevis
H.sapiens

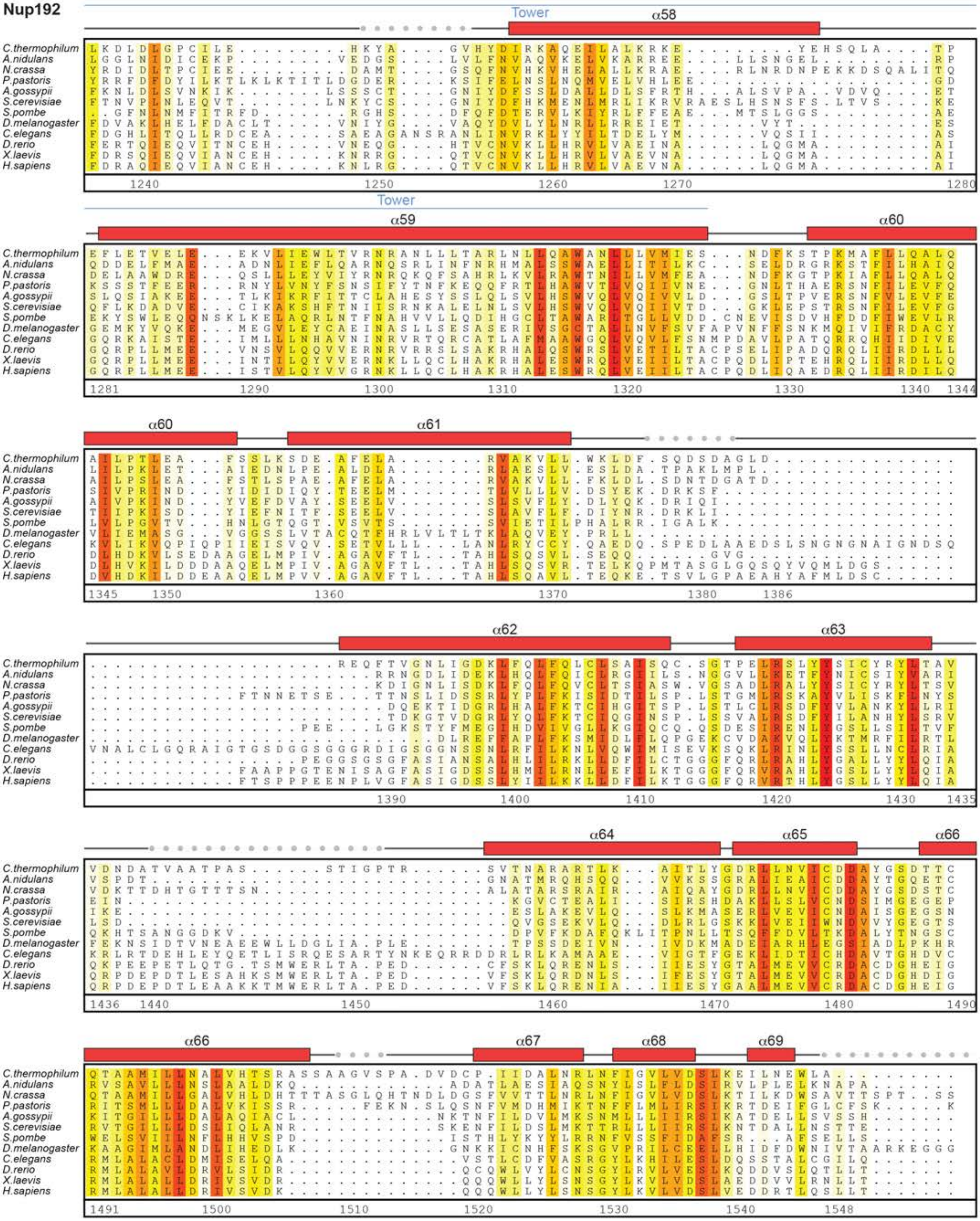

Supp Fig. S38e

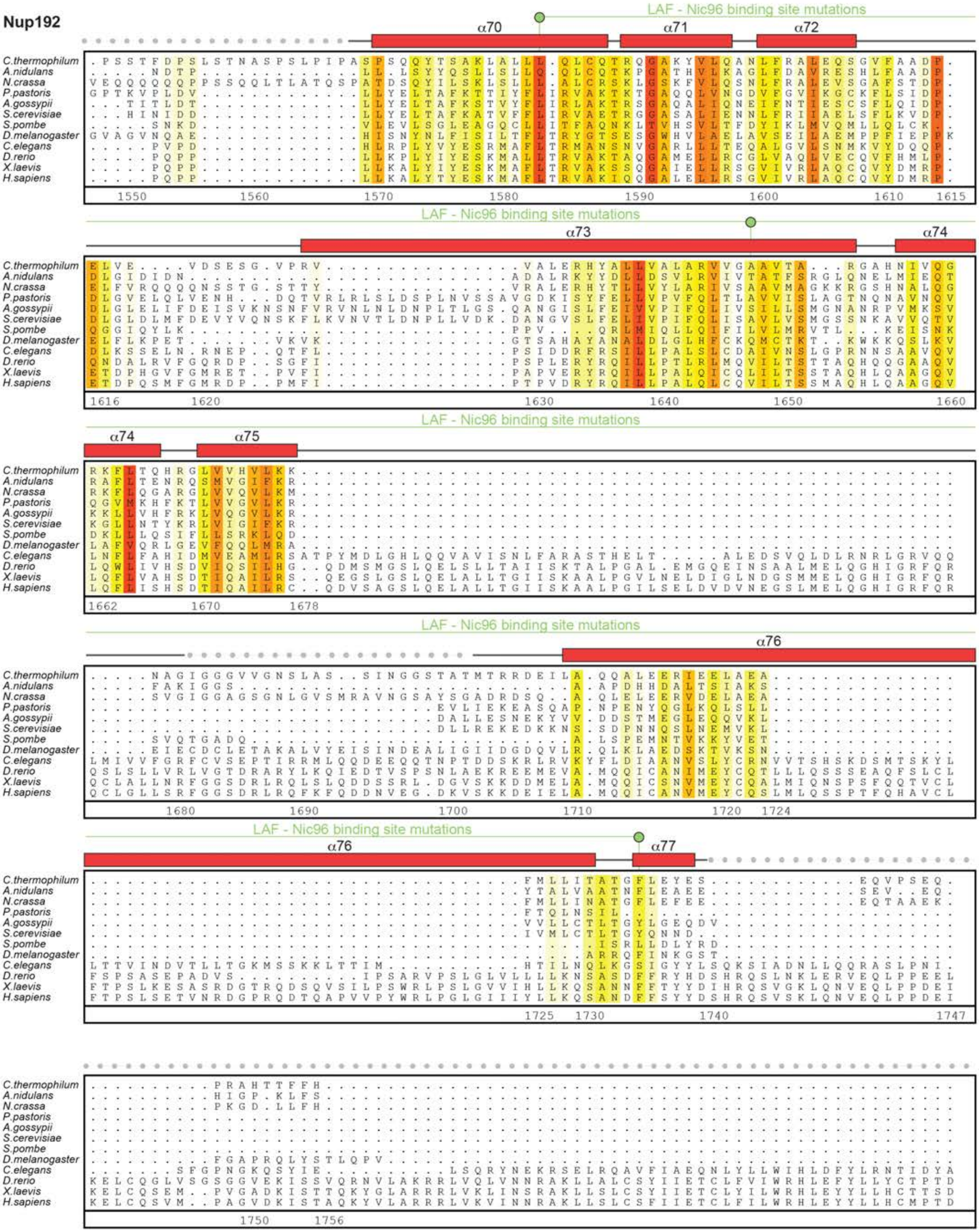

Supp Fig. S38f

### Nup192

```
C.thermophilum  ......................................................................
A.nidulans      ......................................................................
N.crassa        ......................................................................
P.pastoris      ......................................................................
A.gossypii      ......................................................................
S.cerevisiae    ......................................................................
S.pombe         ......................................................................
D.melanogaster  ......................................................................
C.elegans       NEN.............RNAINESNMDDDNDMSVLNASQEEIVQLKQLLISTFNETFCTQLINASEDYSI.
D.rerio         PKDSLLPG.......YRDPAGS......RGLSVSRVSQQDLEQLQSDMSSSFSEPLQRKLLEVEGLYSKS
X.laevis        SQDPVFSNMTFGNRRFQDTFNTDPNMDPRNLRQNKVSQQDVDTLLREGANSFGESLQKRLLDIESLYCKV
H.sapiens       SQDSLFASRTLFKSRRLQDSFASETNLDFRSGLAIVSQHDLDQLQADAINAFGESLQKKLLDIEGLYSKV
```

```
C.thermophilum  ..................................
A.nidulans      ..................................
N.crassa        ..................................
P.pastoris      ..................................
A.gossypii      ..................................
S.cerevisiae    ..................................
S.pombe         ..................................
D.melanogaster  ..................................
C.elegans       .KCKGFNGSLLRRIKALVQFAPITANDVNSSFDS
D.rerio         RSRHTFIQALTRRIRGLVHHAKS...........
X.laevis        RSRHSFIQALVRRIRGLLRVSRV...........
H.sapiens       RSRYSFIQALVRRIRGLLRISRN...........
```

**Fig.S38. Multispecies sequence alignment of Nup192.** Sequences from twelve diverse species were aligned and colored by sequence similarity according to the BLOSUM62 matrix from white (less than 55 % similarity), to yellow (55 % similarity), to red (100 % identity). Numbering below alignment is relative to the *C. thermophilum* sequence. Secondary structure observed in the Nup192 structures is shown above the alignment: $\alpha$-helices (red bars), $\beta$-sheets (blue bars), and unstructured regions (black lines). Disordered regions are indicated by gray dots. Residues found by mutational analysis to affect binding to Nup53 (magenta) (*36*), Nup145N (cyan; Fig. 4, figs. S15, S19, and S21), and Nic96 (pale green; Fig. 2, figs. S2, S6, and S7) are indicated by circles above the alignment.

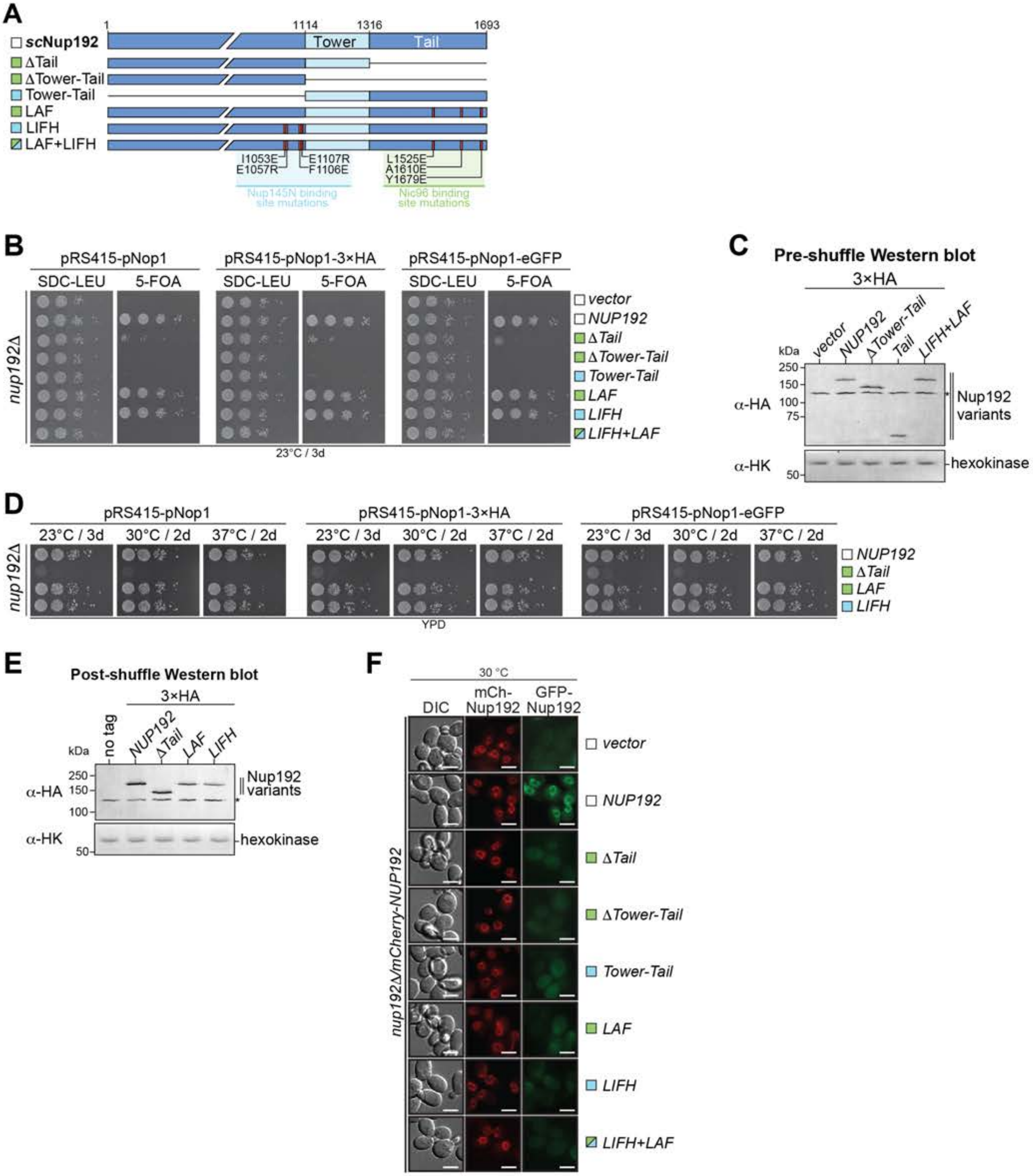
A
1
1114
1316
1693
scNup192
Tower
Tail
ΔTail
ΔTower-Tail
Tower-Tail
LAF
LIFH
LAF+LIFH
I1053E
E1057R
E1107R
F1106E
L1525E
A1610E
Y1679E
Nup145N binding site mutations
Nic96 binding site mutations
B
pRS415-pNop1
pRS415-pNop1-3×HA
pRS415-pNop1-eGFP
SDC-LEU
5-FOA
nup192Δ
vector
NUP192
ΔTail
ΔTower-Tail
Tower-Tail
LAF
LIFH
LIFH+LAF
23°C / 3d
C
Pre-shuffle Western blot
3×HA
kDa
vector
NUP192
ΔTower-Tail
Tail
LIFH+LAF
250
150
100
75
50
α-HA
α-HK
Nup192 variants
hexokinase
D
pRS415-pNop1
pRS415-pNop1-3×HA
pRS415-pNop1-eGFP
23°C / 3d
30°C / 2d
37°C / 2d
nup192Δ
NUP192
ΔTail
LAF
LIFH
YPD
E
Post-shuffle Western blot
3×HA
kDa
no tag
NUP192
ΔTail
LAF
LIFH
250
150
100
50
α-HA
α-HK
Nup192 variants
hexokinase
F
30 °C
DIC
mCh-Nup192
GFP-Nup192
nup192Δ/mCherry-NUP192
vector
NUP192
ΔTail
ΔTower-Tail
Tower-Tail
LAF
LIFH
LIFH+LAF

**Fig.S39. *In vivo* validation of Nup192 interactions with Nic96 and Nup145N.** Supporting data for analysis presented in Fig. 8. (**A**) Domain structure of Nup192 variants introduced into the *nup192*Δ strain, with colored squares indicating whether the mutation affects the Nup145N binding site (cyan), the Nic96 binding site (pale green), or both (cyan and pale green). (**B**) Viability analysis of the *nup192*Δ strain after transformation with indicated Nup192 variants and selection of ten-fold serial dilution series spotted onto either SDC-LEU and SDC+5-FOA plates (shuffling). Variants were either untagged or N-terminally tagged with 3×HA or eGFP. (**C**) Western blot analysis of expression levels of 3×HA-tagged non-rescuing Nup192 variants in unshuffled *nup192*Δ strain transformants selected in SDC-LEU media. 3×HA-Nup192 variants and the endogenous hexokinase loading control were detected with anti-HA and anti-hexokinase antibodies, respectively. Asterisk indicates nonspecific band detected by the anti-HA antibody. (**D**) Growth analysis at different temperatures of ten-fold serial dilution series of the shuffled *nup192*Δ strain carrying indicated Nup192 variants, replicated for untagged and N-terminally 3×HA- or eGFP-tagged constructs. (**E**) Western blot analysis of expression levels of 3×HA-tagged Nup192 variants in shuffled *nup192*Δ strain transformants selected on SDC+5-FOA media and subsequently grown in YPD media at 30 °C. Asterisk indicates nonspecific band detected by the anti-HA antibody. 3×HA-Nup192 variants and the endogenous hexokinase loading control were detected with anti-HA and anti-hexokinase antibodies, respectively. (**F**) Subcellular localization of eGFP-Nup192 variants (green) and wildtype mCherry-Nup192 (red) in an unshuffled *nup192*Δ strain. Scalebars are 5 μm.

**Nup188**

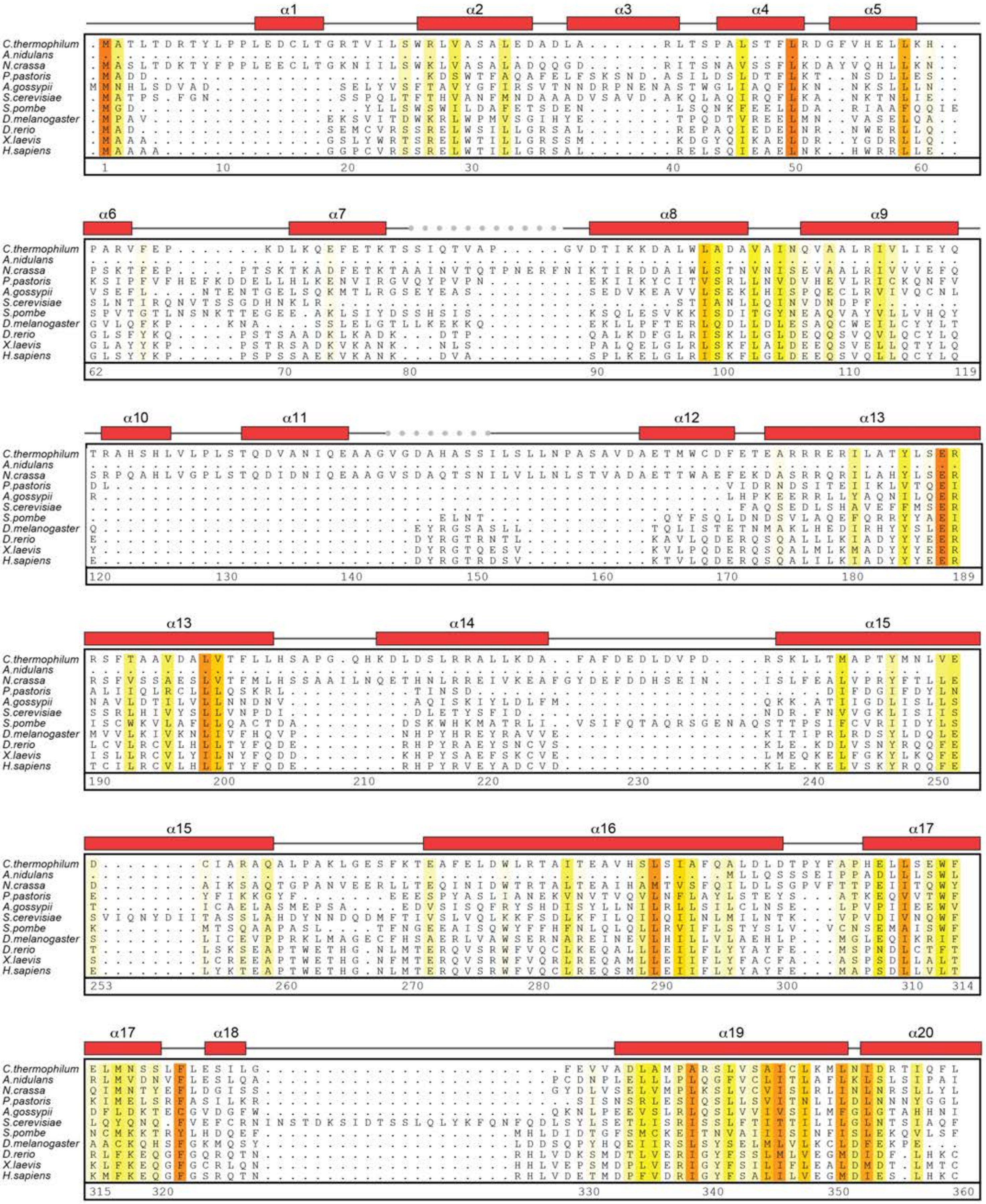

Supp Fig. S40a

Hoelz et al., 2021

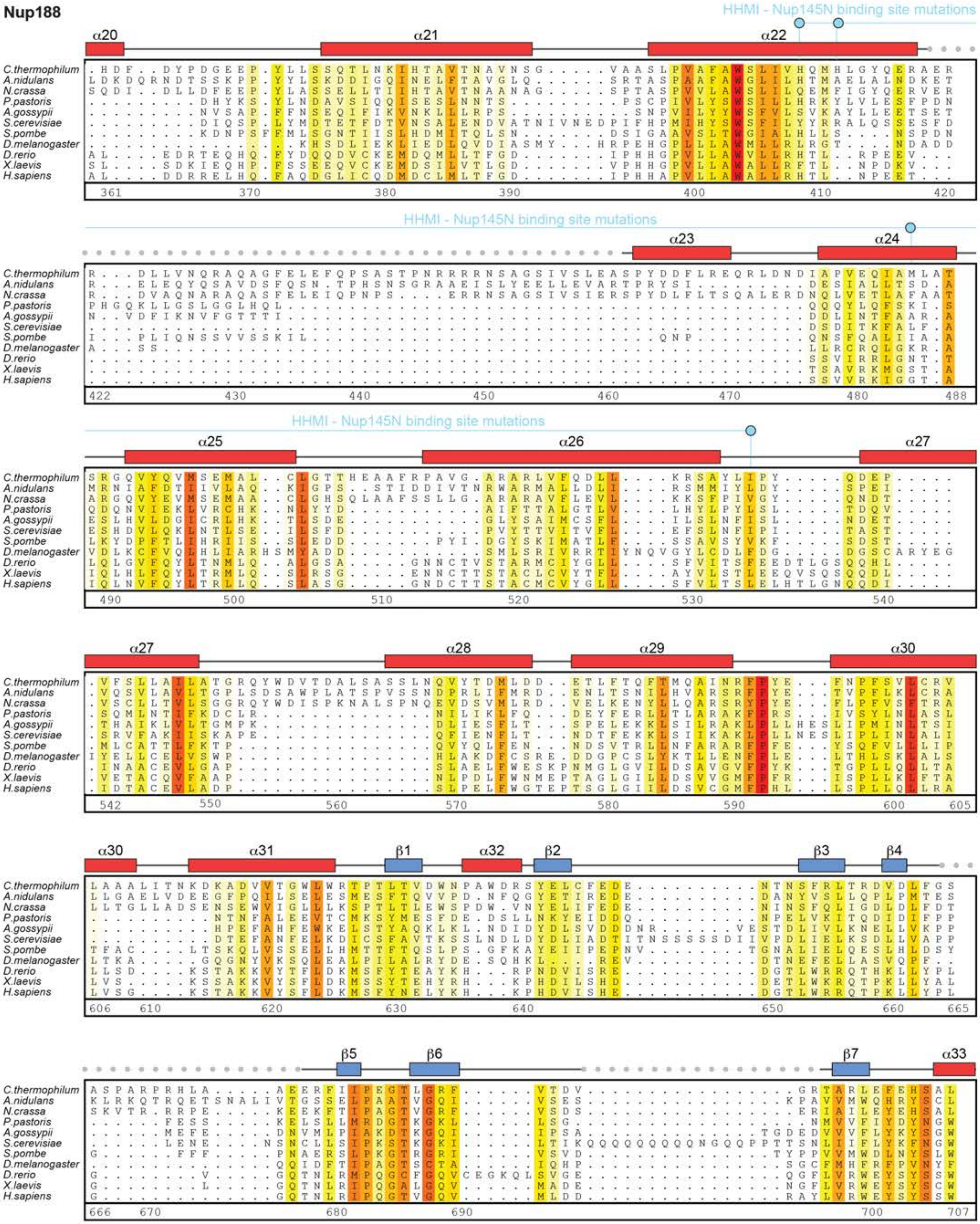
Nup188
HHMI - Nup145N binding site mutations
α20 α21 α22 α23 α24 α25 α26 α27 α28 α29 α30 α31 α32 α33
β1 β2 β3 β4 β5 β6 β7
C.thermophilum
A.nidulans
N.crassa
P.pastoris
A.gossypii
S.cerevisiae
S.pombe
D.melanogaster
D.rerio
X.laevis
H.sapiens

### Nup188

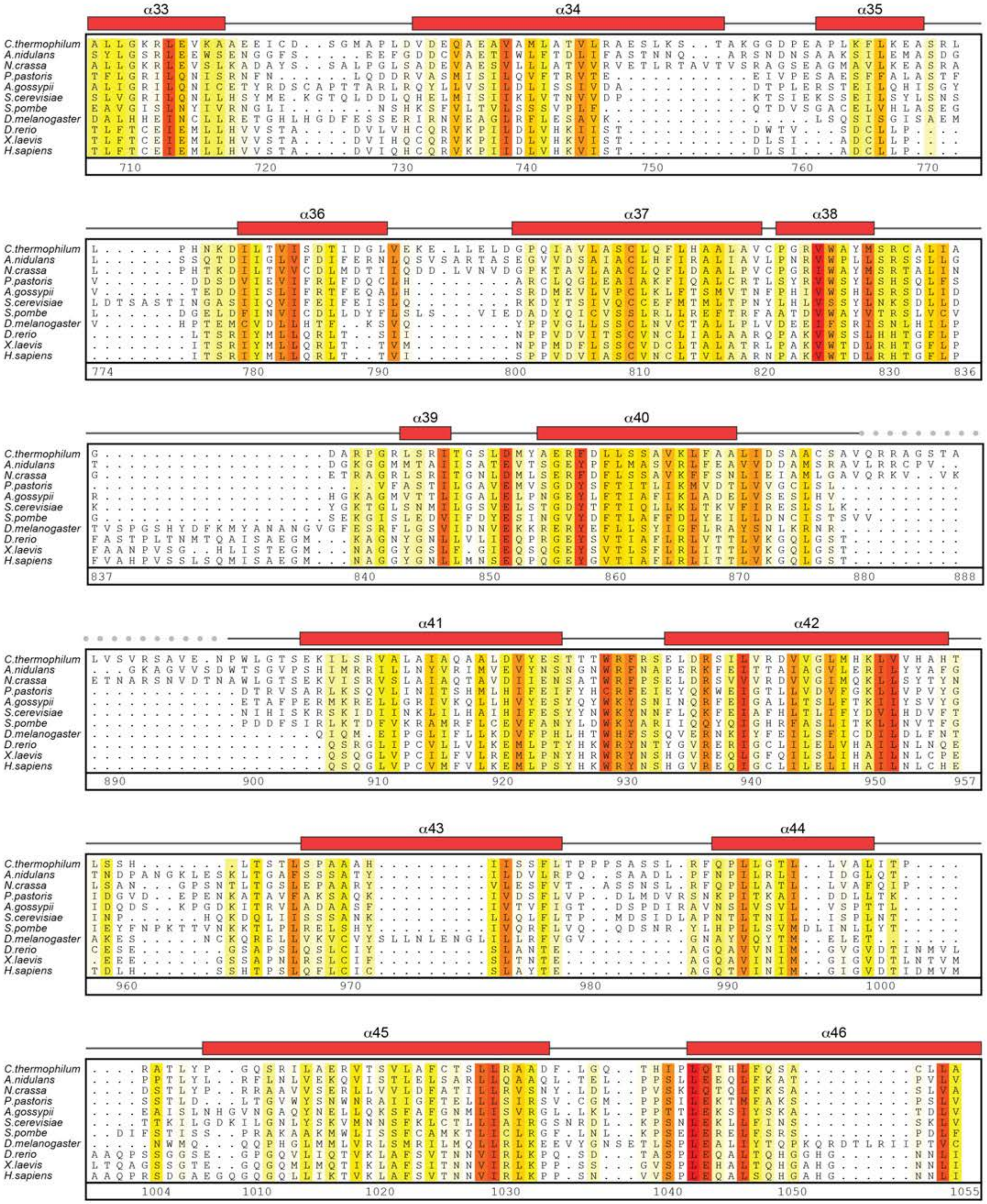
α33
α34
α35
α36
α37
α38
α39
α40
α41
α42
α43
α44
α45
α46
C.thermophilum
A.nidulans
N.crassa
P.pastoris
A.gossypii
S.cerevisiae
S.pombe
D.melanogaster
D.rerio
X.laevis
H.sapiens

### Nup188

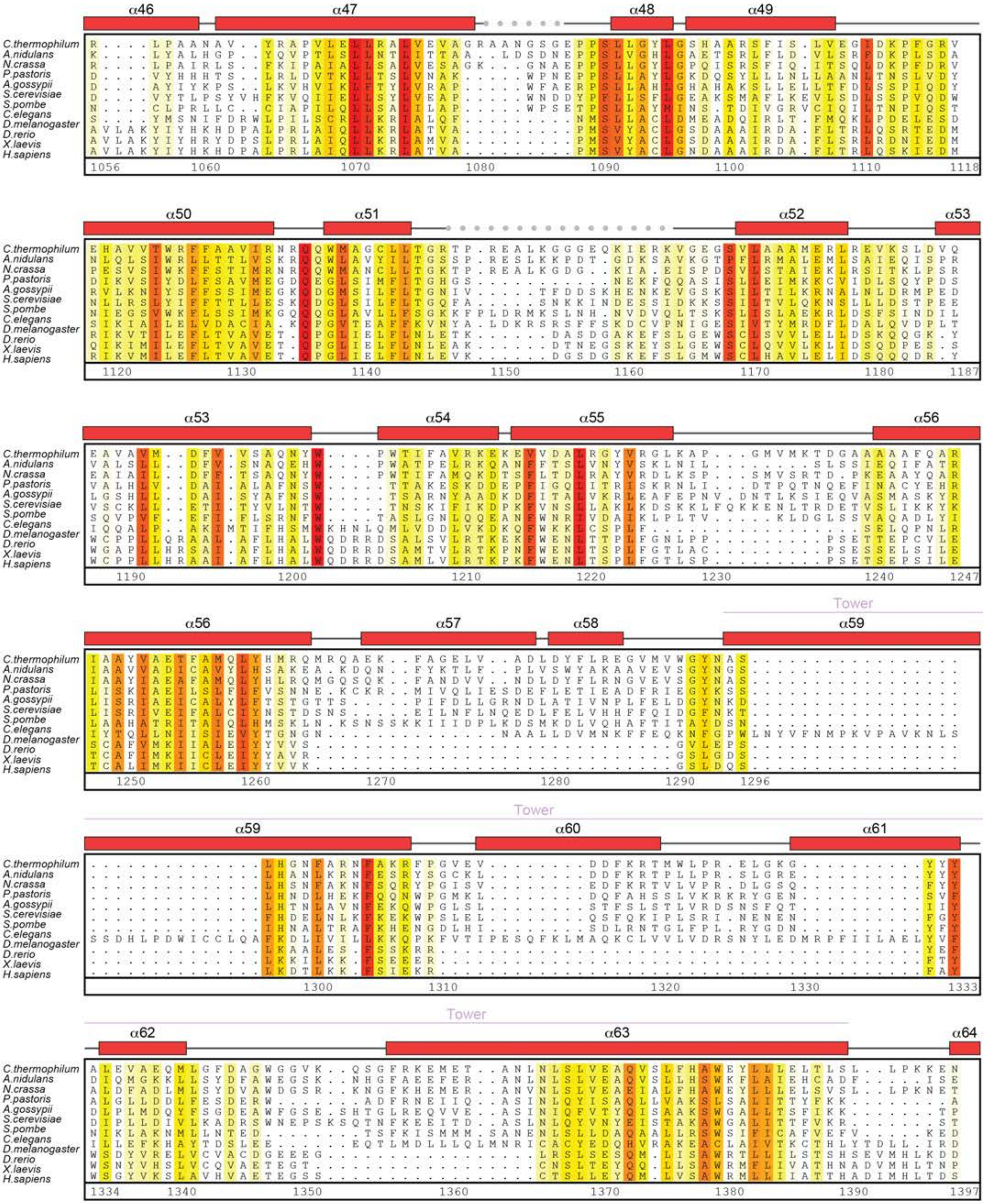

Supp Fig. S40d

Hoelz et al., 2021

**Nup188**

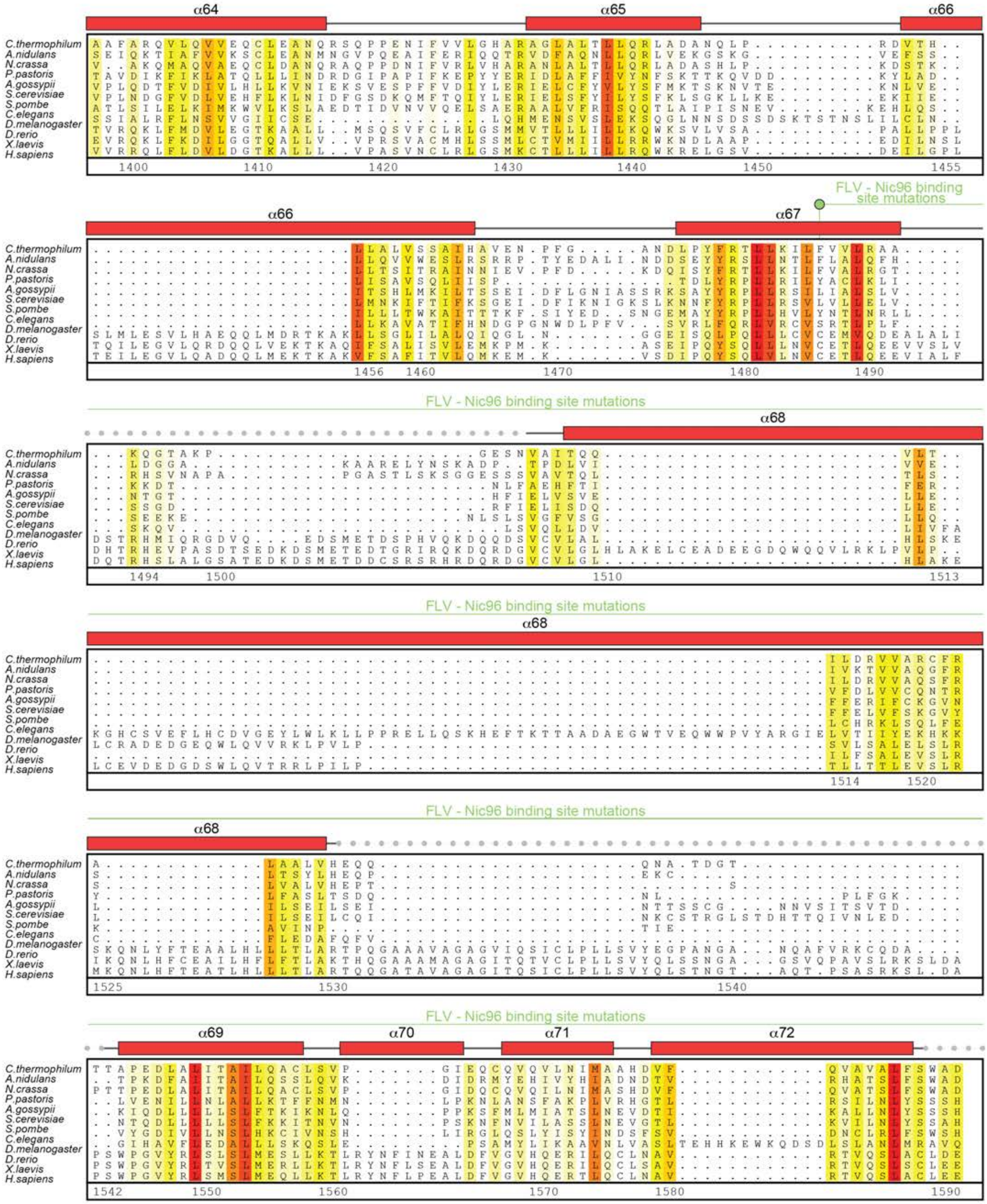

**Supp Fig. S40e**

Hoelz et al., 2021

**Nup188**

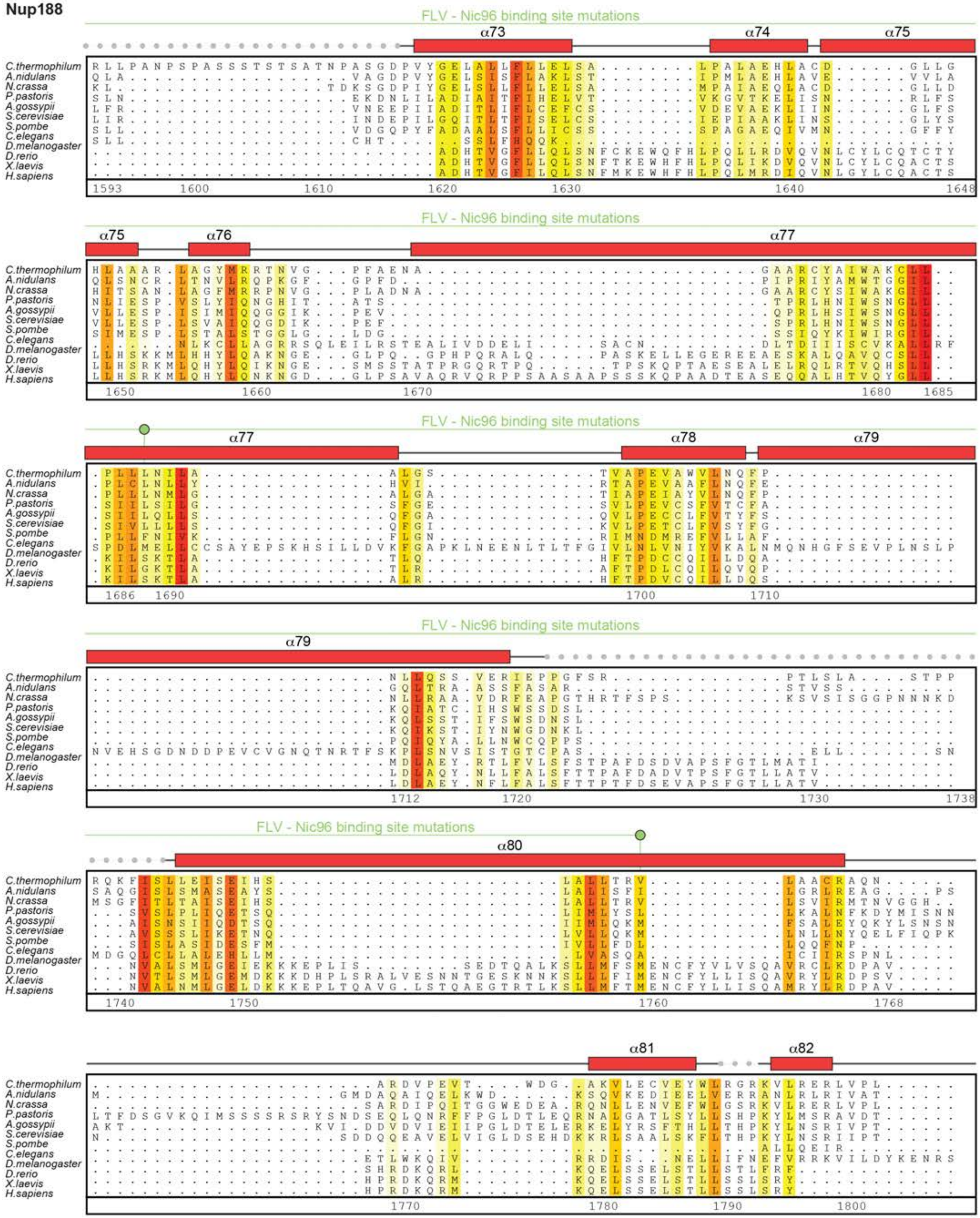

FLV - Nic96 binding site mutations
α73
α74
α75
α76
α77
α78
α79
α80
α81
α82
C.thermophilum
A.nidulans
N.crassa
P.pastoris
A.gossypii
S.cerevisiae
S.pombe
C.elegans
D.melanogaster
D.rerio
X.laevis
H.sapiens

**Nup188**

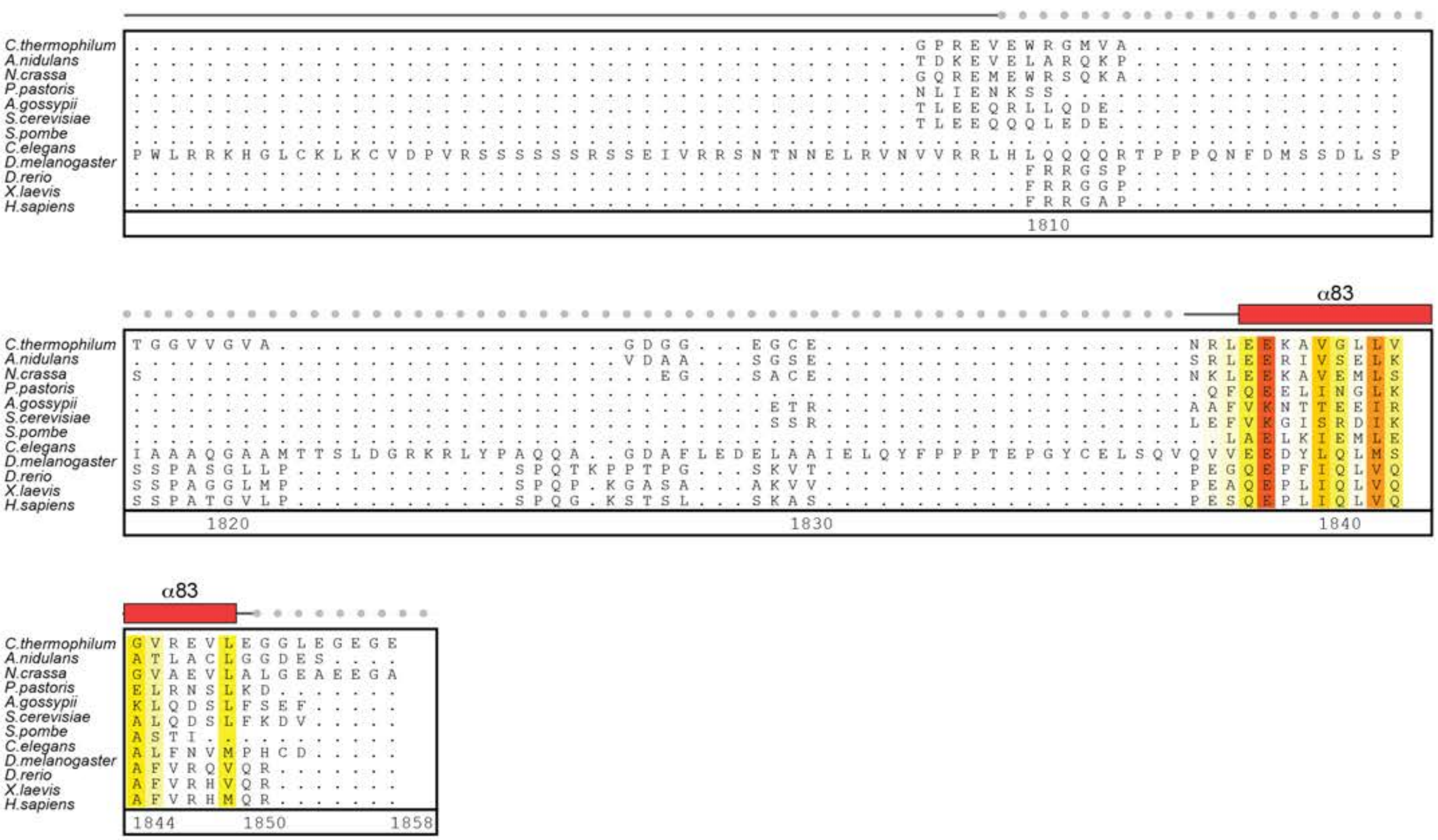

Supp Fig. S40g

Hoelz et al., 2021

**Fig.S40. Multispecies sequence alignment of Nup188.** Sequences from eleven diverse species were aligned and colored by sequence similarity according to the BLOSUM62 matrix from white (less than 55 % similarity), to yellow (55 % similarity), to red (100 % identity). Numbering below alignment is relative to the *C. thermophilum* sequence. Secondary structure observed in the Nup188 structures is shown above the alignment: α-helices (red bars), β-sheets (blue bars), and unstructured regions (black lines). Disordered regions are indicated by gray dots. Residues found by mutational analysis to affect binding to Nup145N (cyan; Fig. 5, figs. S27, S29, and S30), and Nic96 (pale green; Fig. 3, figs. S8, S11, and S12) are indicated by circles above the alignment.

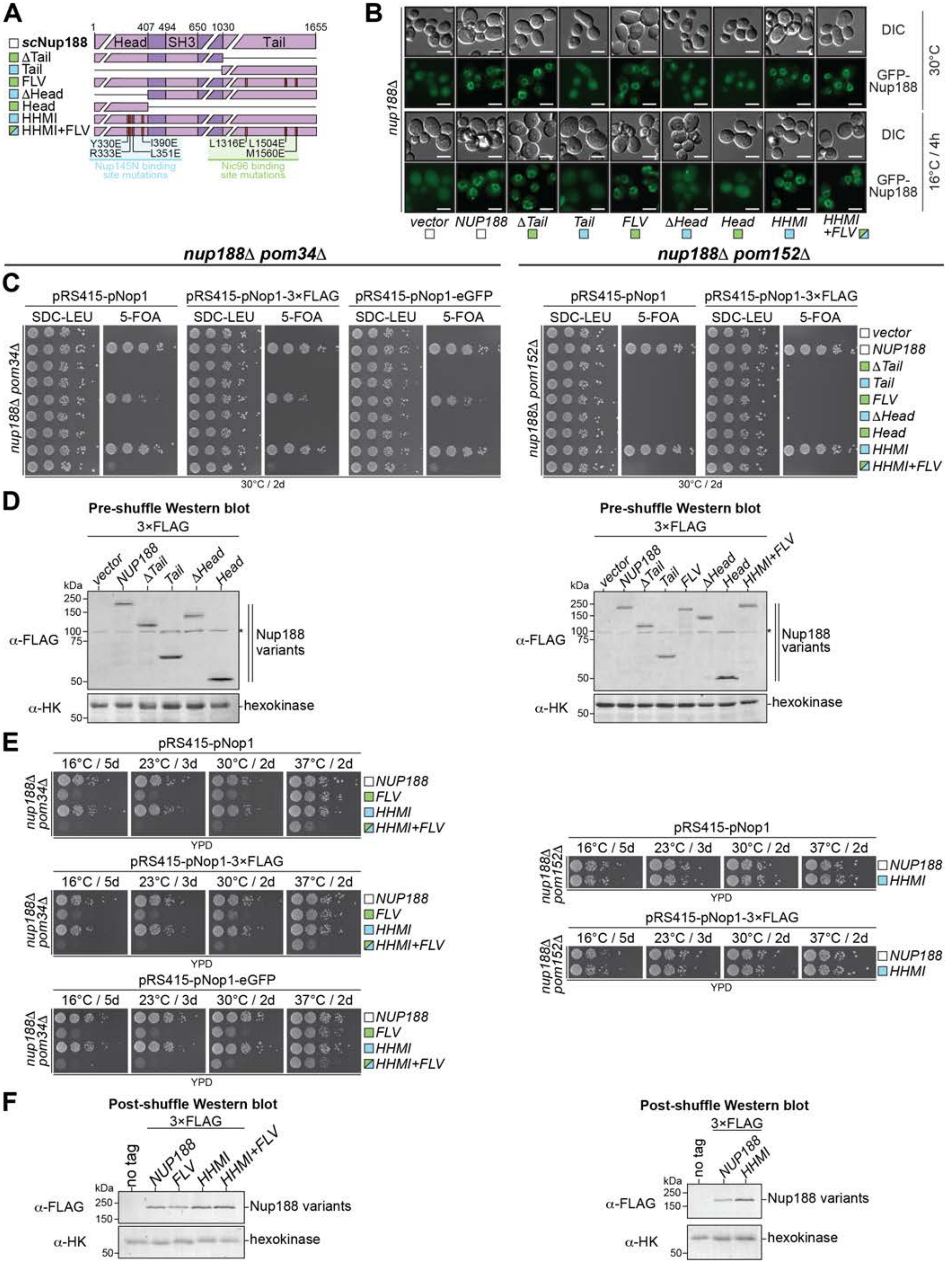

Supp Fig. S41

Hoelz et al., 2021

**Fig.S41. *In vivo* validation of Nup188 interactions with Nic96 and Nup145N.** Supporting data for analysis presented in Fig. 9. (**A**) Domain structure of Nup188 variants introduced into the *nup188∆*, *nup188∆pom34∆*, and *nup188∆pom152∆* strains, with colored squares indicating whether the mutation affects the Nup145N binding site (cyan), the Nic96 binding site (pale green), or both (cyan and pale green). (**B**) Subcellular localization of GFP-Nup188 variants transformed into a *nup188∆* strain. Scalebars are 5 $\mu$m. (**C**) Viability analysis of the *nup188∆pom34∆* and *nup188∆pom152∆* strains after transformation with indicated Nup188 variants and selection of ten-fold serial dilution series spotted onto either SDC-LEU and SDC+5-FOA plates (shuffling). Variants were either untagged or N-terminally tagged with 3×FLAG or eGFP. (**D**) Western blot analysis of expression levels of 3×FLAG-tagged non-rescuing Nup188 variants in unshuffled *nup188∆pom34∆* and *nup188∆pom152∆* strain transformants selected in SDC-LEU media. 3×FLAG-Nup188 variants and the endogenous hexokinase loading control were detected with anti-FLAG and anti-hexokinase antibodies, respectively. Asterisk indicates nonspecific band detected by the anti-FLAG antibody. (**D**) Growth analysis at different temperatures of ten-fold serial dilution series of the shuffled *nup188∆pom34∆* and *nup188∆pom152∆* strains carrying indicated Nup192 variants, replicated for untagged and N-terminally 3×FLAG or eGFP-tagged constructs. (**E**) Western blot analysis of expression levels of 3×FLAG-tagged Nup188 variants in shuffled *nup188∆pom34∆* and *nup188∆pom152∆* transformants selected on SDC+5-FOA media and subsequently grown in YPD media at 30 °C. Asterisk indicates nonspecific band detected by the anti-HA antibody. 3×HA-Nup192 variants and the endogenous hexokinase loading control were detected with anti-FLAG and anti-hexokinase antibodies, respectively.

### Nic96

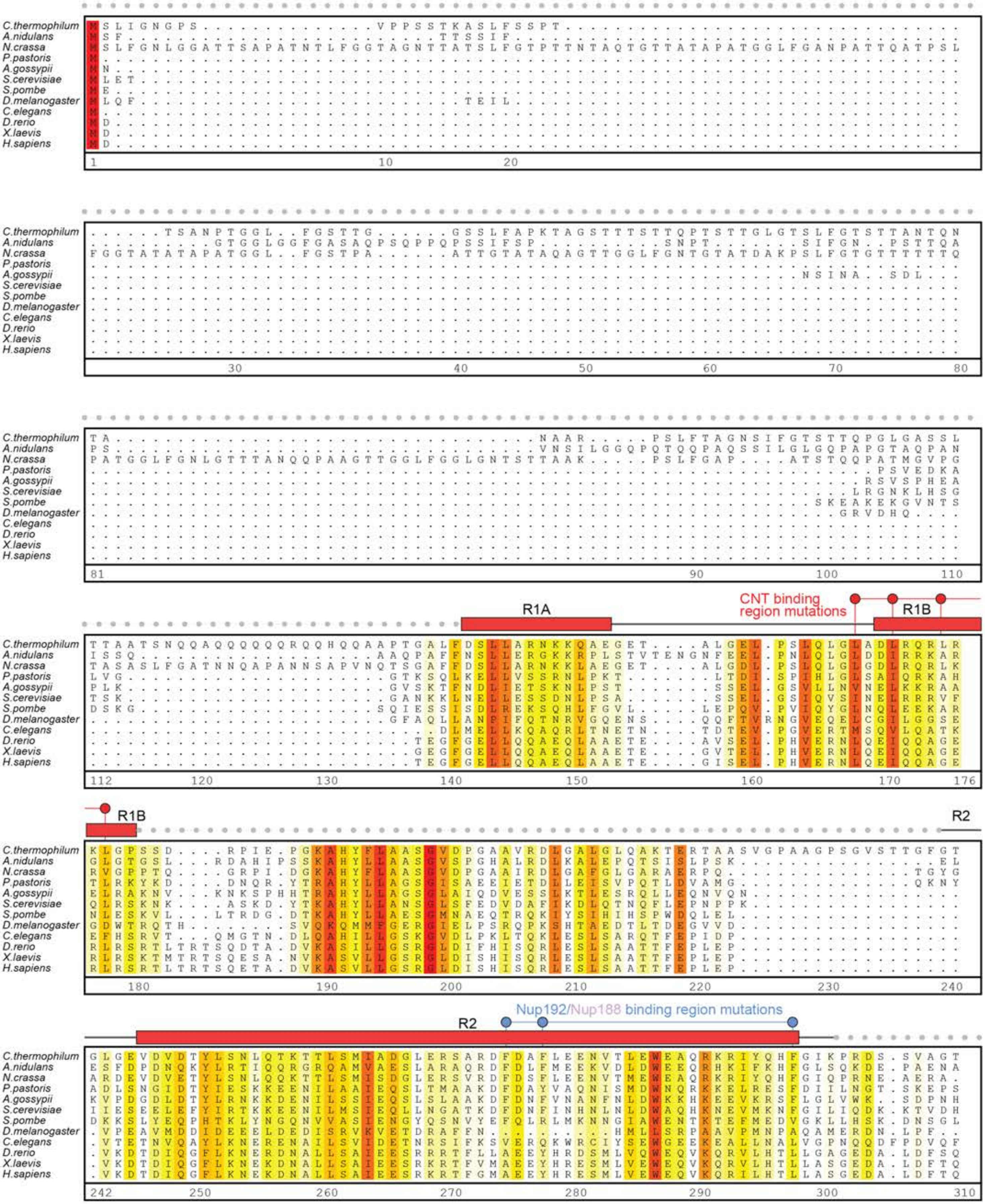

C.thermophilum
A.nidulans
N.crassa
P.pastoris
A.gossypii
S.cerevisiae
S.pombe
D.melanogaster
C.elegans
D.rerio
X.laevis
H.sapiens
R1A
R1B
R2
CNT binding region mutations
Nup192/Nup188 binding region mutations

### Nic96

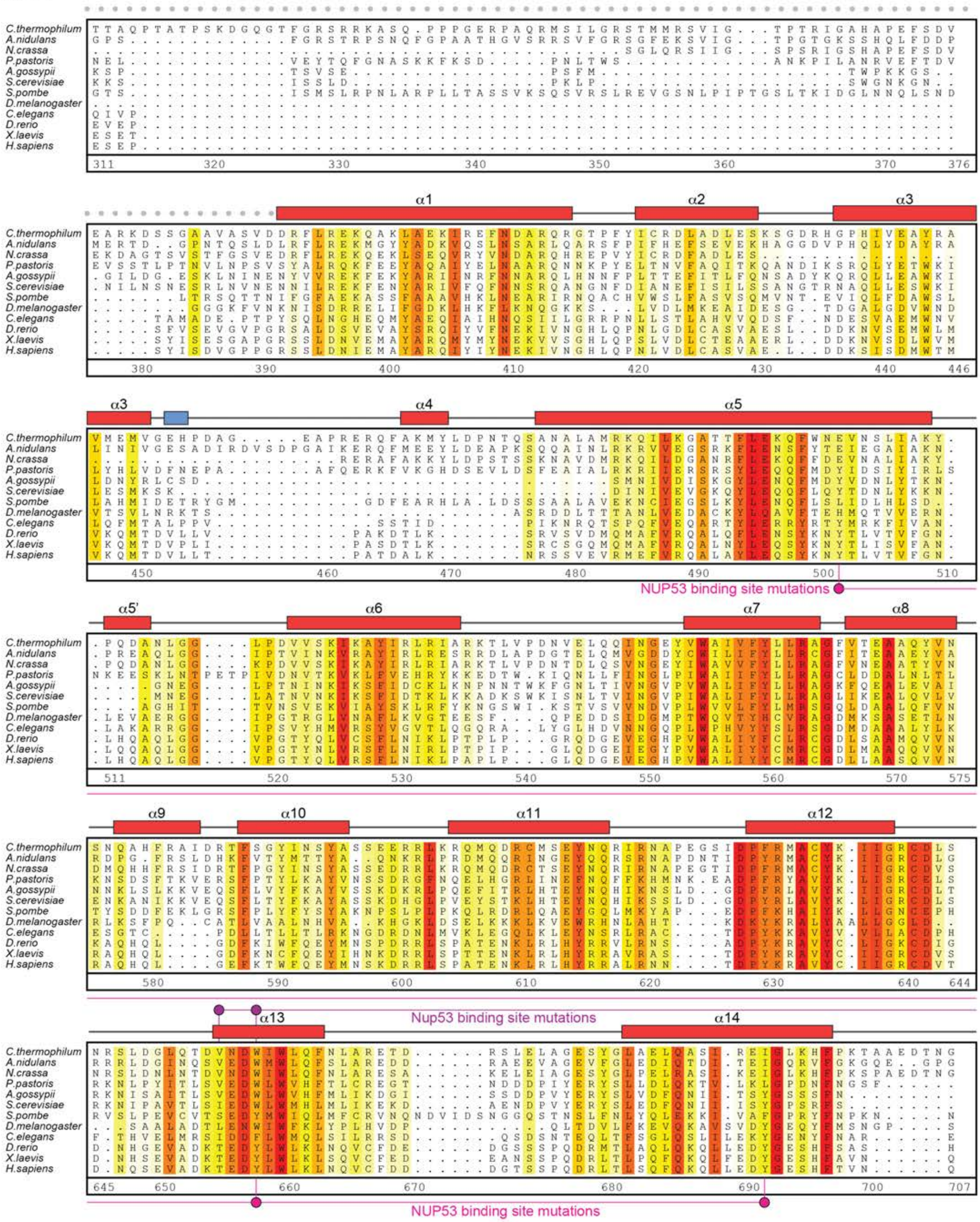

Supp. Fig. S42b

Hoelz et al., 2021

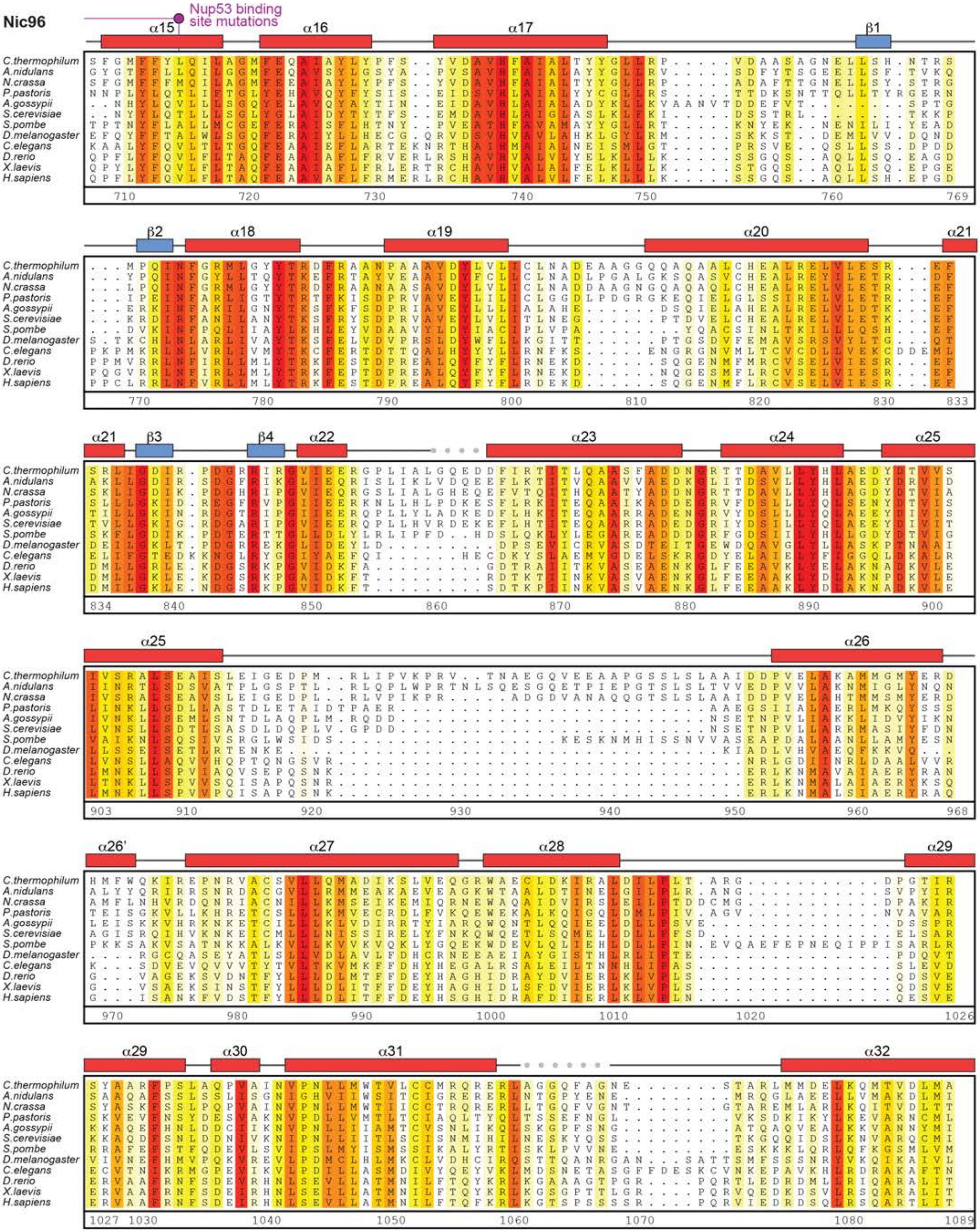

Supp. Fig. S42c

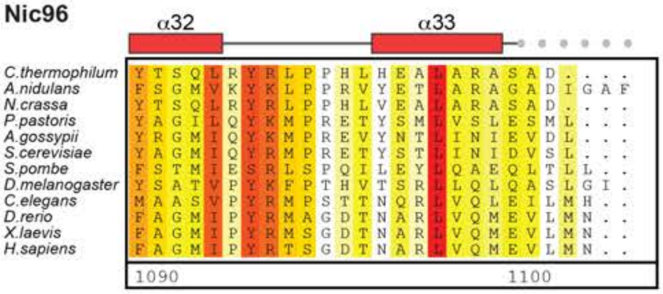

Nic96
α32
α33
C.thermophilum YTSQLRYRLPPHLHEALARASAD....
A.nidulans FSGMVKYKLPPRVYETLARAGADIGAF
N.crassa YTSQLRYRLPPHLVEALARASAD....
P.pastoris YAGILQYKMPRETYSMLVSLESML...
A.gossypii YRGMIQYKMPREVYNTLINIEVDL...
S.cerevisiae YAGMIQYRMPRETYSTLINIDVSL...
S.pombe FSTMIESRLSPQILEYLQAEQLTLL..
D.melanogaster YSATVPYKFPTHVTSRLLQLQASLGI.
C.elegans MAASVPYRMPSTTNQRLVQLEILMH..
D.rerio FAGMIPYRMAGDTNARLVQMEVLMN..
X.laevis FAGMIPYRMSGDTNARLVQMEVLMN..
H.sapiens FAGMIPYRTSGDTNARLVQMEVLMN..
1090
1100

**Fig.S42. Multispecies sequence alignment of Nic96.** Sequences from twelve diverse species were aligned and colored by sequence similarity according to the BLOSUM62 matrix from white (less than 55 % similarity), to yellow (55 % similarity), to red (100 % identity). Numbering below alignment is relative to the *C. thermophilum* sequence. Secondary structure observed in the Nic96 structures is shown above the alignment: α-helices (red bars), β-sheets (blue bars), and unstructured regions (black lines). Disordered regions are indicated by gray dots. Residues found by mutational analysis to affect binding to the CNT (red) (*25*), Nup192 (blue; Fig. 2C, figs. S2, S5, S6), Nup188 (blue; Fig. 3C, figs. S8, S10, S12), or Nup53 (magenta) (*30*) in the *C. thermophilum* system, are indicated by circles above the alignment. Residues found by mutational analysis to affect binding between NUP93 and NUP53 (bright pink; Fig. 12, fig. S53) in the *H. sapiens* system, are indicated by circles below the alignment.

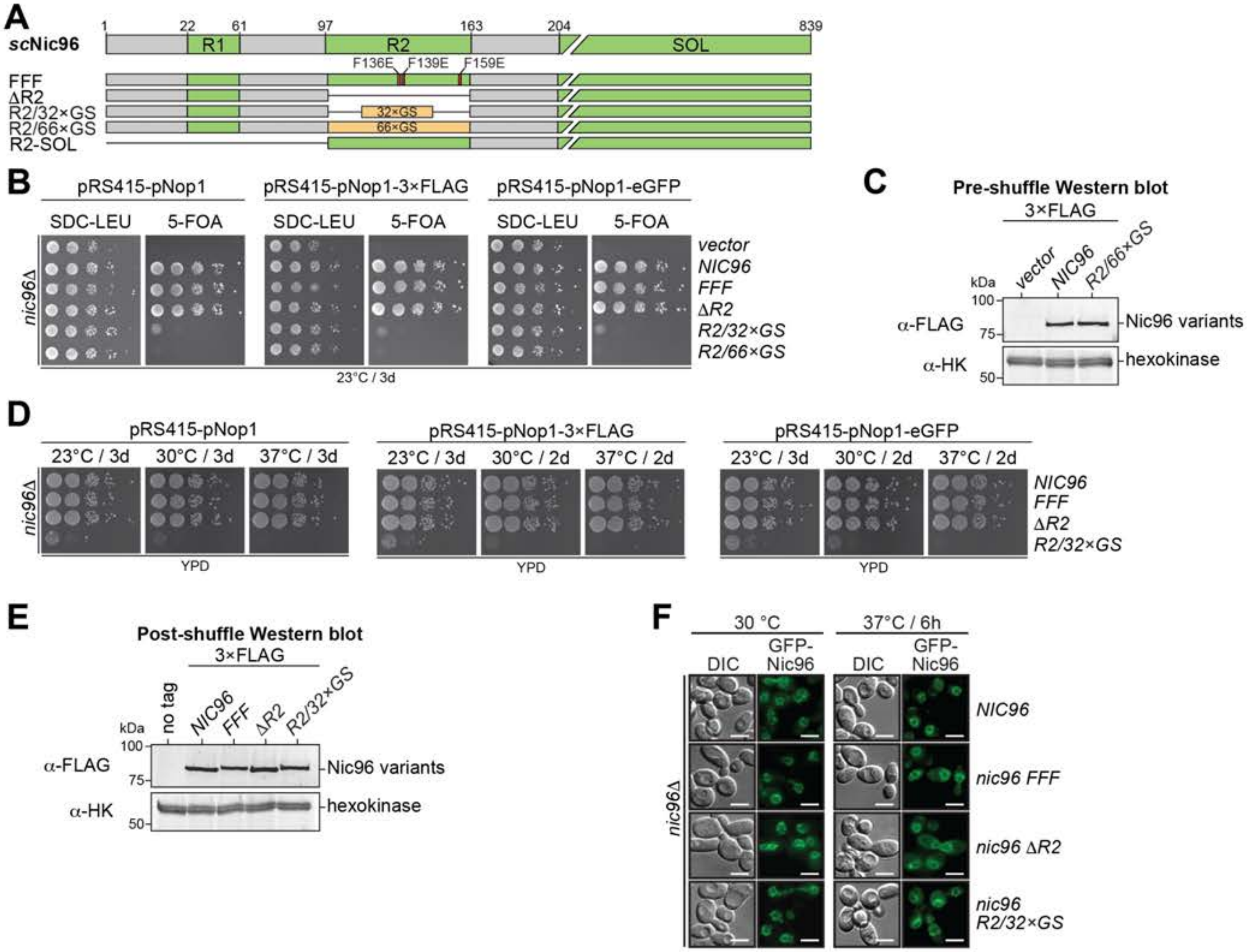

A
scNic96
1
22
61
97
163
204
839
R1
R2
SOL
F136E
F139E
F159E
FFF
ΔR2
R2/32×GS
R2/66×GS
R2-SOL
32×GS
66×GS
B
pRS415-pNop1
pRS415-pNop1-3×FLAG
pRS415-pNop1-eGFP
SDC-LEU
5-FOA
nic96Δ
vector
NIC96
FFF
ΔR2
R2/32×GS
R2/66×GS
23°C / 3d
C
Pre-shuffle Western blot
3×FLAG
vector
NIC96
R2/66×GS
kDa
100
75
50
α-FLAG
α-HK
Nic96 variants
hexokinase
D
pRS415-pNop1
pRS415-pNop1-3×FLAG
pRS415-pNop1-eGFP
23°C / 3d
30°C / 3d
37°C / 3d
30°C / 2d
37°C / 2d
nic96Δ
NIC96
FFF
ΔR2
R2/32×GS
YPD
E
Post-shuffle Western blot
3×FLAG
no tag
NIC96
FFF
ΔR2
R2/32×GS
kDa
100
75
50
α-FLAG
α-HK
Nic96 variants
hexokinase
F
30 °C
37°C / 6h
DIC
GFP-Nic96
nic96Δ
NIC96
nic96 FFF
nic96 ΔR2
nic96 R2/32×GS

**Fig.S43. *In vivo* validation of the Nic96 R2 region.** Supporting data for analysis presented in Fig. 10. (**A**) Domain structure of Nic96 variants introduced into the *nic96Δ* strain. (**B**) Viability analysis of the *nic96Δ* strain after transformation with indicated Nic96 variants and selection of ten-fold serial dilution series spotted onto either SDC-LEU and SDC+5-FOA plates (shuffling). Variants were either untagged or N-terminally tagged with 3×FLAG or eGFP. (**C**) Western blot analysis of expression levels of 3×FLAG-tagged non-rescuing Nic96 variants in unshuffled *nic96Δ* strain transformants selected in SDC-LEU media. 3×FLAG-Nic96 variants and the endogenous hexokinase loading control were detected with anti-FLAG and anti-hexokinase antibodies, respectively. (**D**) Growth analysis at different temperatures of ten-fold serial dilution series of the shuffled *nic96Δ* strain carrying indicated Nic96 variants, replicated for untagged and N-terminally 3×FLAG or eGFP-tagged constructs. (**E**) Western blot analysis of expression levels of 3×FLAG-tagged Nic96 variants in shuffled *nic96Δ* strain transformants selected on SDC+5-FOA media and subsequently grown in YPD media at 30 °C. 3×FLAG-Nic96 variants and the endogenous hexokinase loading control were detected with anti-FLAG and anti-hexokinase antibodies, respectively. (**F**) Subcellular localization of shuffled eGFP-Nic96 variants in the *nic96Δ* strain at different temperatures. Scalebars are 5 μm.

### Nup53 from species related to *S.cerevisae*

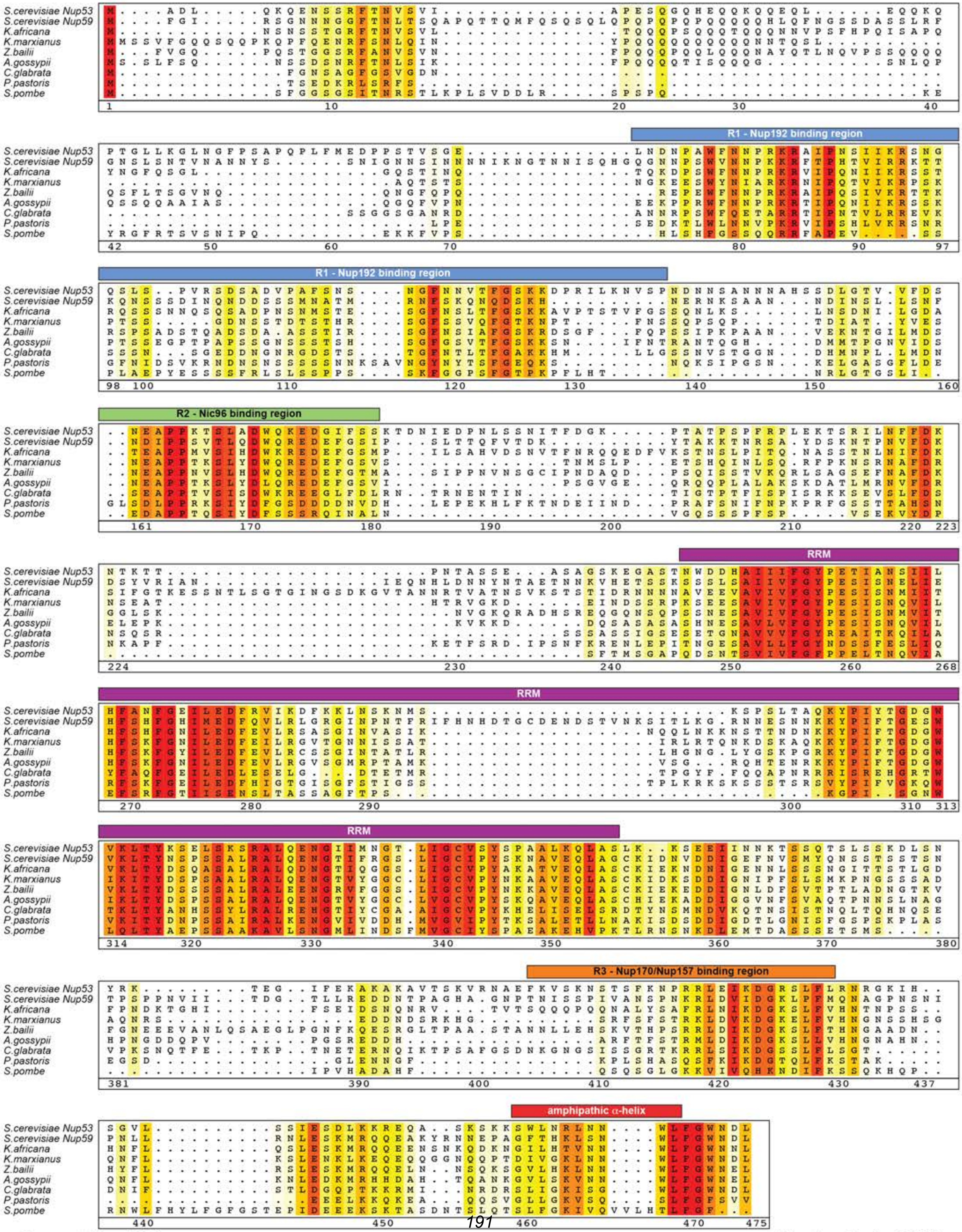

**Supp. Fig. S44** Hoelz et al., 2021

**Fig.S44. Multispecies sequence alignment of Nup53 from species related to *S. cerevisiae*.** Sequences from eight diverse species, including two *S. cerevisiae* paralogs Nup53 and Nup59, were aligned and colored by sequence similarity according to the BLOSUM62 matrix from white (less than 55 % similarity), to yellow (55 % similarity), to red (100 % identity). Numbering below alignment is relative to the *S. cerevisiae* sequence. Regions of the sequence are annotated above the alignment: R1 – Nup192 binding region (blue), R2 – Nic96 binding region (green), RNA recognition motif – RRM (magenta), R3 – Nup157/Nup170 binding region (orange) (*126*), amphipathic $\alpha$-helix (red).

### Nup53 from species related to *C.thermophilum*

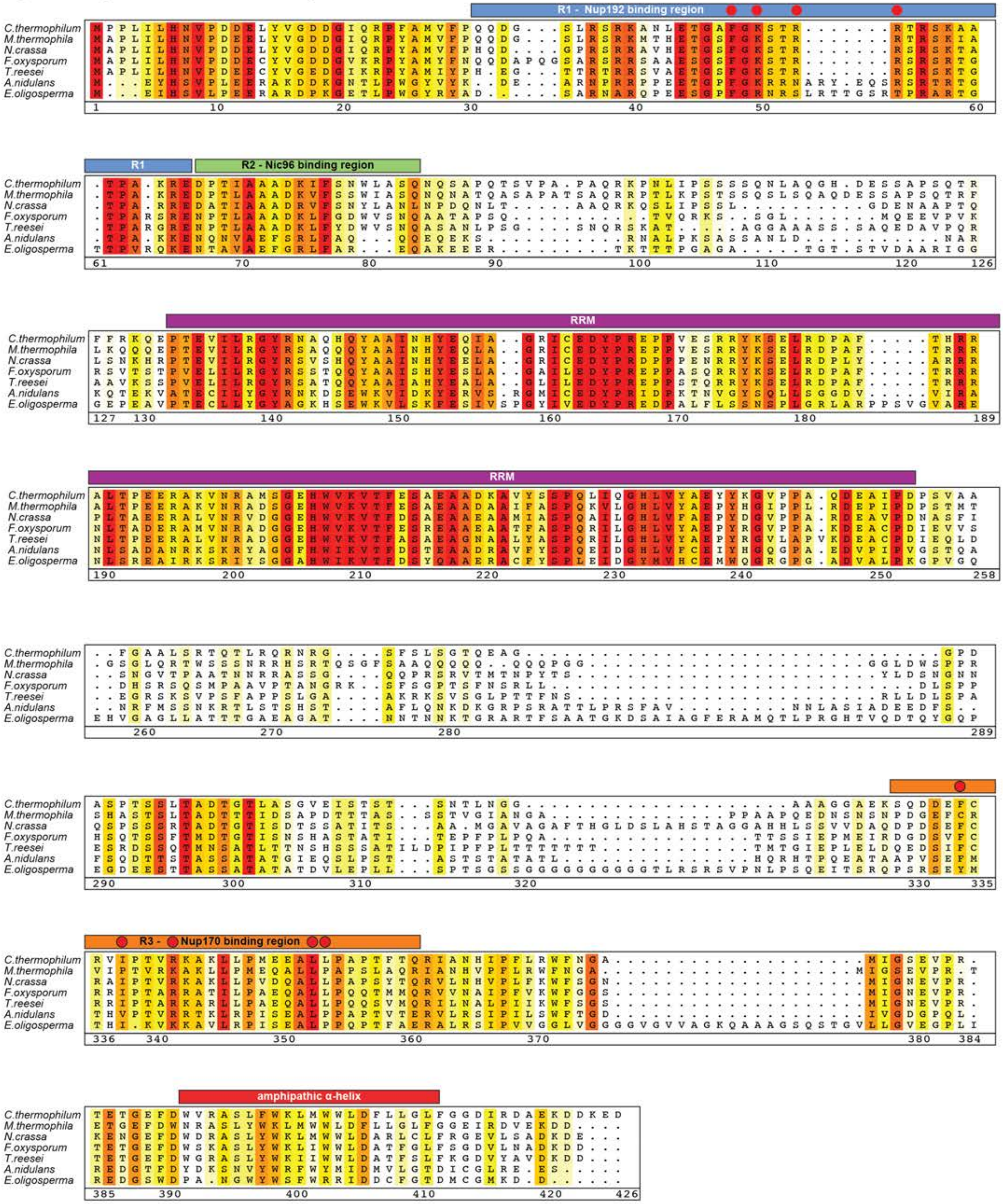

**Fig.S45. Multispecies sequence alignment of Nup53 from species related to *C. thermophilum*.** Sequences from seven diverse species were aligned and colored by sequence similarity according to the BLOSUM62 matrix from white (less than 55 % similarity), to yellow (55 % similarity), to red (100 % identity). Numbering below alignment is relative to the *C. thermophilum* sequence. Regions of the sequence are annotated above the alignment: R1 – Nup192 binding region (blue), R2 – Nic96 binding region (green), RNA recognition motif – RRM (magenta), R3 – Nup157/Nup170 binding region (orange), amphipathic $\alpha$-helix (red). Residues found by mutational analysis to affect Nup192 binding (*36*) or Nup170 biding (*25*) are indicated by red circles above the alignment.

### NUP35 from metazoan species

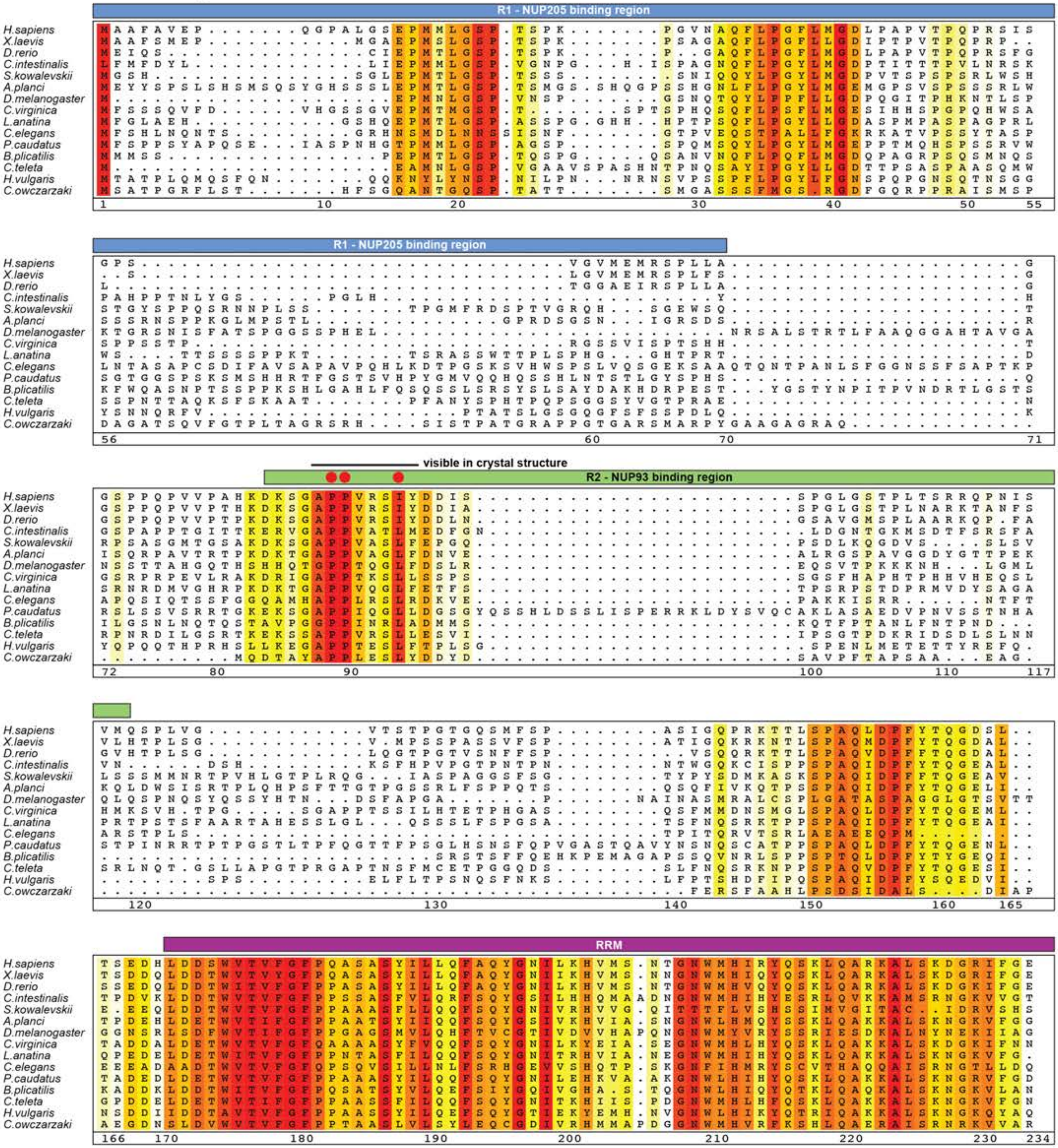

**Supp. Fig. S46a**

**Hoelz et al., 2021**

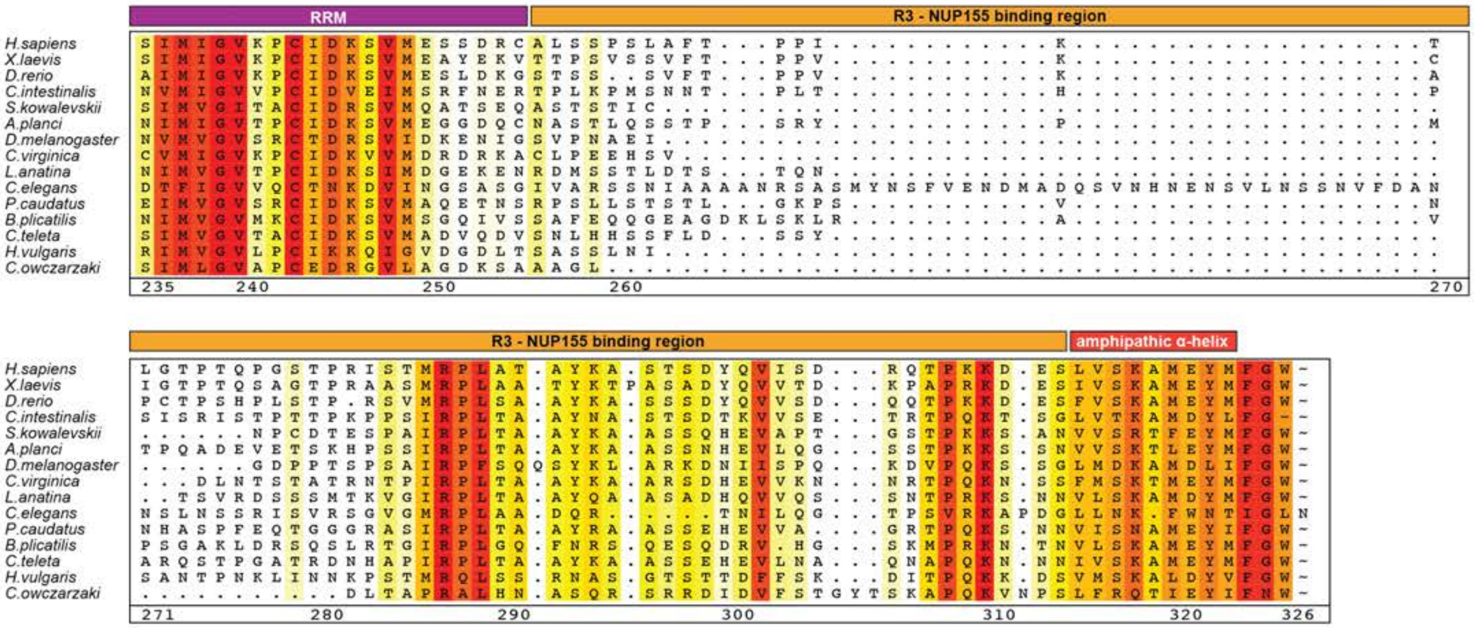

Supp. Fig. S46b

**Fig.S46. Multispecies sequence alignment of Nup53 from species related to *H. sapiens*.** Sequences from fifteen diverse species were aligned and colored by sequence similarity according to the BLOSUM62 matrix from white (less than 55 % similarity), to yellow (55 % similarity), to red (100 % identity). Numbering below alignment is relative to the *H. sapiens* sequence. Regions of the sequence are annotated above the alignment: R1 – NUP205 binding region (blue), R2 – NUP93 binding region (green), RNA recognition motif – RRM (magenta), R3 – NUP155 binding region, amphipathic $\alpha$-helix (red). Residues found by mutational analysis to affect binding to NUP93$^{SOL}$ (Fig. 12, fig. S53) are indicated by red circles above the alignment.

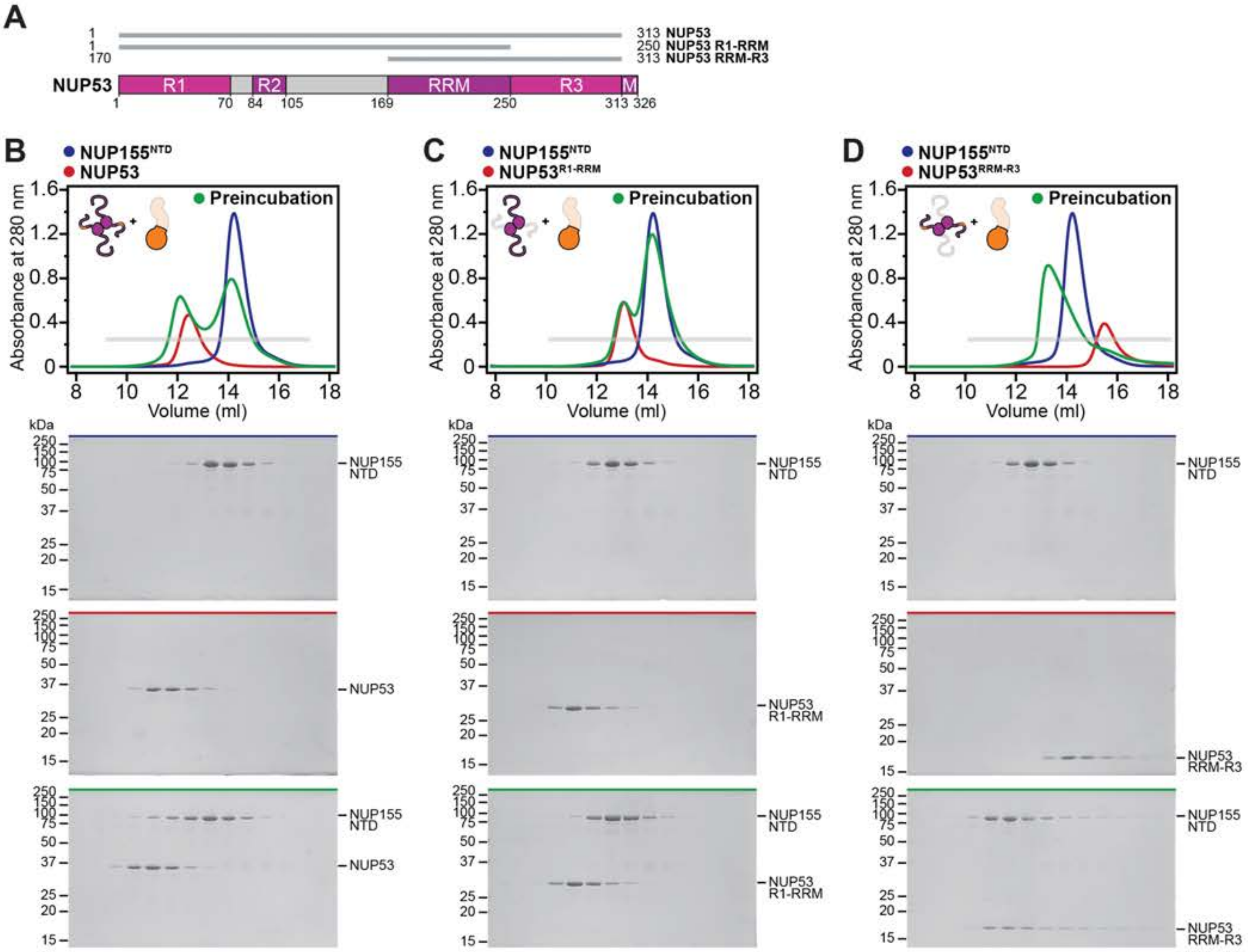

Supp. Fig. S47

Hoelz et al., 2021

**Fig.S47. NUP155$^{NTD}$ binds to a topologically conserved region of NUP53.** (**A**) Domain structure of NUP53 with gray lines indicating truncation construct boundaries analyzed. (**B**-**D**) SEC profiles are shown for nups individually (red and blue) and after their preincubation (green) for NUP155$^{NTD}$ binding to various NUP53 constructs. Gray bars indicate fractions that were resolved on SDS-PAGE gels and visualized by Coomassie staining. All SEC profiles were obtained using a Superdex 200 10/300 GL column.

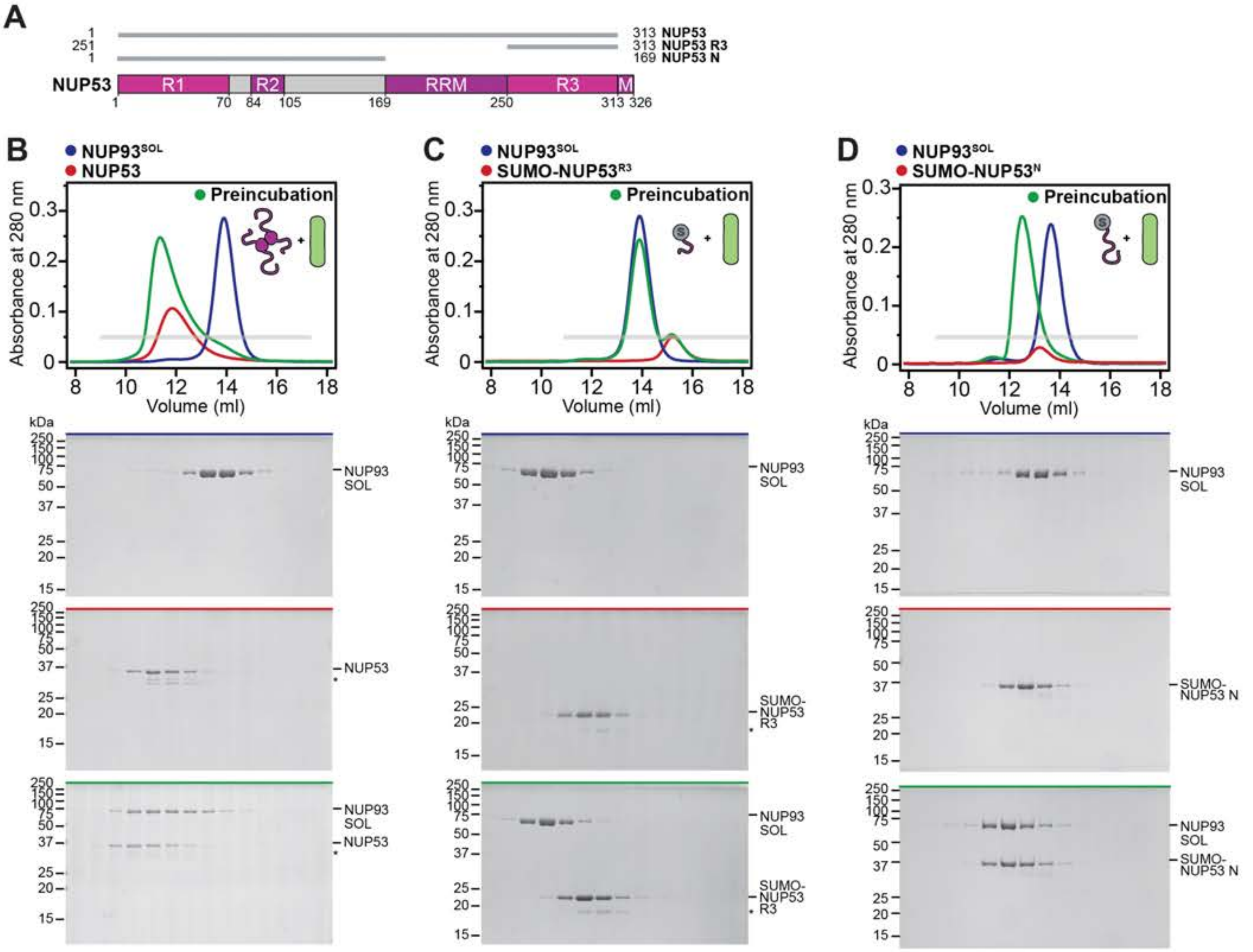
A
1 313 NUP53
251 313 NUP53 R3
1 169 NUP53 N
NUP53
R1
R2
RRM
R3
M
1 70 84 105 169 250 313 326
B
NUP93SOL
NUP53
Preincubation
Absorbance at 280 nm
Volume (ml)
kDa
NUP93 SOL
NUP53
C
NUP93SOL
SUMO-NUP53R3
Preincubation
SUMO-NUP53 R3
D
NUP93SOL
SUMO-NUP53N
Preincubation
SUMO-NUP53 N

**Fig.S48. NUP93$^{SOL}$ binds to a topologically conserved region of NUP53.** (**A**) Domain structure of NUP53 with gray lines indicating truncation construct boundaries analyzed. (**B**-**D**) SEC profiles are shown for nups individually (red and blue) and after their preincubation (green) for NUP93$^{SOL}$ binding to various NUP53 constructs. Gray bars indicate fractions that were resolved on SDS-PAGE gels and visualized by Coomassie staining. All SEC profiles were obtained using a Superdex 200 10/300 GL column. Asterisks indicate degradation products or contamination.

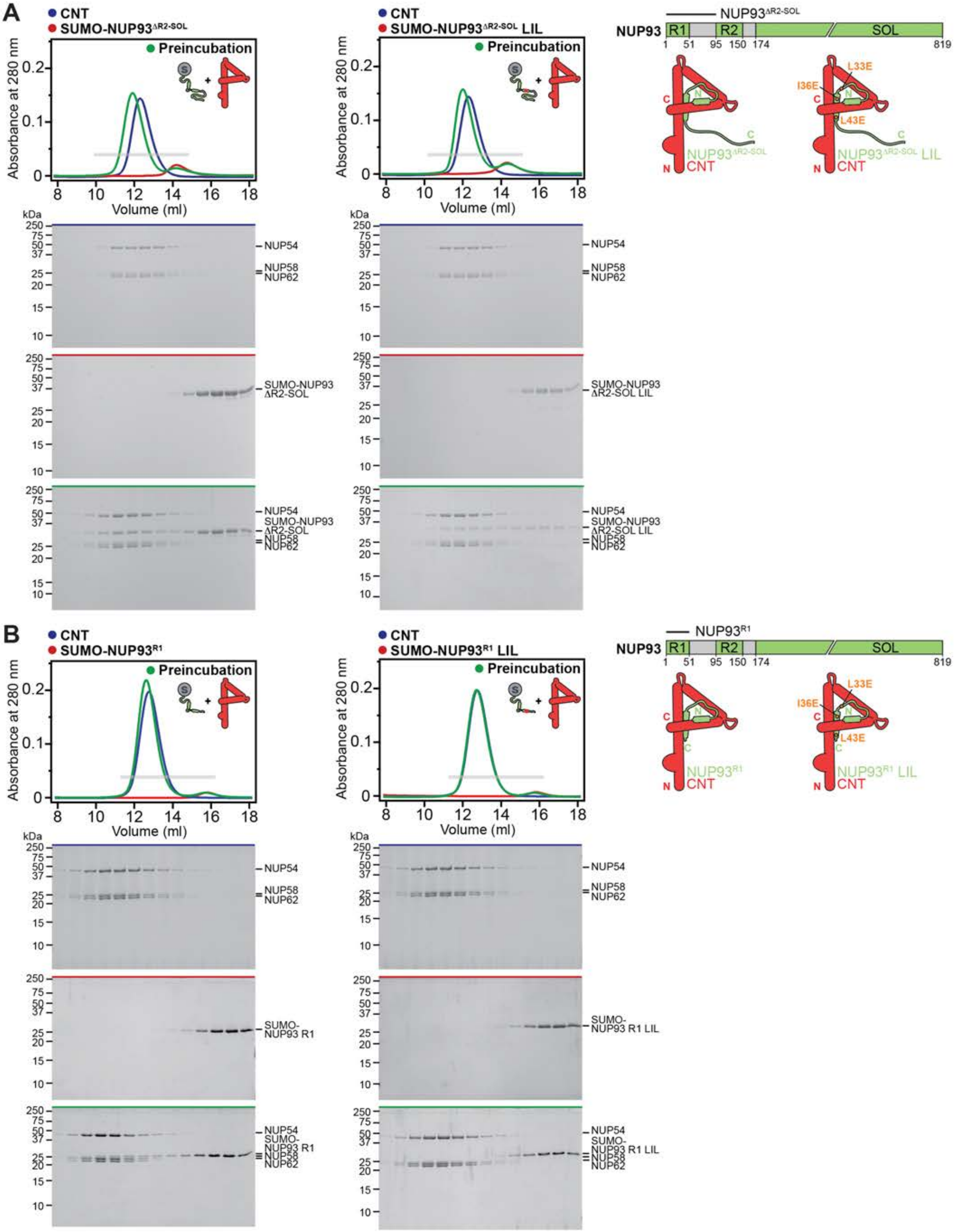

A
CNT
SUMO-NUP93ΔR2-SOL
Preincubation
Absorbance at 280 nm
Volume (ml)
kDa
NUP54
NUP58
NUP62
SUMO-NUP93 ΔR2-SOL
SUMO-NUP93ΔR2-SOL LIL
SUMO-NUP93 ΔR2-SOL LIL
NUP93ΔR2-SOL
NUP93
R1
R2
SOL
1 51 95 150 174 819
L33E
I36E
L43E
NUP93ΔR2-SOL
CNT
NUP93ΔR2-SOL LIL
B
SUMO-NUP93R1
SUMO-NUP93R1 LIL
SUMO-NUP93 R1
SUMO-NUP93 R1 LIL
NUP93R1
NUP93R1 LIL

**Fig.S49. NUP93$^{R1}$ binding to the CNT is disrupted by mutating evolutionarily conserved residues.** (**A-B**) Domain structure of NUP93 with black lines indicating the analyzed construct boundaries, and a schematic of the suggested NUP93$^{R1}$-CNT binding mode and location of evolutionarily conserved LIL residues (*right*). The *H. sapiens* CNT is composed of NUP54, NUP58 and NUP62. SEC profiles are shown for nups or nup complexes individually (red and blue) and after their preincubation (green) for the CNT binding to (A) NUP93$^{\Delta R2\text{-}SOL}$ and the mutant NUP93$^{\Delta R2\text{-}SOL}$ LIL, and (B) NUP93$^{R1}$ and the mutant NUP93$^{R1}$ LIL. Gray bars indicate fractions that were resolved on SDS-PAGE gels and visualized by Coomassie staining. All SEC profiles were obtained using a Superdex 200 10/300 GL column.

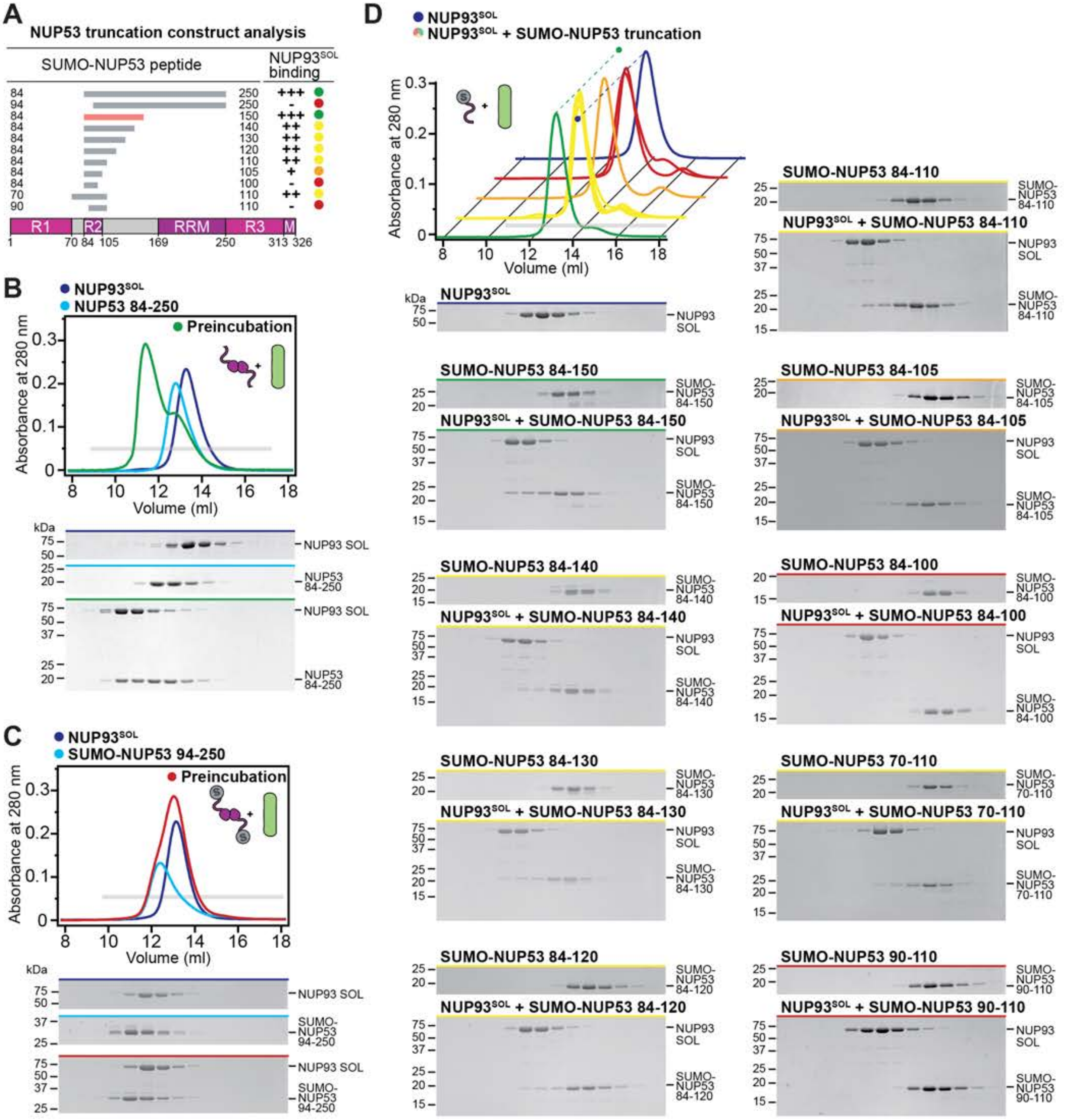
A
NUP53 truncation construct analysis
SUMO-NUP53 peptide
NUP93^SOL binding
84 250 +++
94 250 -
84 150 +++
84 140 ++
84 130 ++
84 120 ++
84 110 ++
84 105 +
84 100 -
70 110 ++
90 110 -
R1 R2 RRM R3 M
1 70 84 105 169 250 313 326
B
NUP93^SOL
NUP53 84-250
Preincubation
Absorbance at 280 nm
Volume (ml)
kDa
NUP93 SOL
NUP53 84-250
C
NUP93^SOL
SUMO-NUP53 94-250
Preincubation
Absorbance at 280 nm
Volume (ml)
kDa
NUP93 SOL
SUMO-NUP53 94-250
D
NUP93^SOL
NUP93^SOL + SUMO-NUP53 truncation
Absorbance at 280 nm
Volume (ml)
kDa
NUP93^SOL
SUMO-NUP53 84-150
NUP93^SOL + SUMO-NUP53 84-150
SUMO-NUP53 84-140
NUP93^SOL + SUMO-NUP53 84-140
SUMO-NUP53 84-130
NUP93^SOL + SUMO-NUP53 84-130
SUMO-NUP53 84-120
NUP93^SOL + SUMO-NUP53 84-120
SUMO-NUP53 84-110
NUP93^SOL + SUMO-NUP53 84-110
SUMO-NUP53 84-105
NUP93^SOL + SUMO-NUP53 84-105
SUMO-NUP53 84-100
NUP93^SOL + SUMO-NUP53 84-100
SUMO-NUP53 70-110
NUP93^SOL + SUMO-NUP53 70-110
SUMO-NUP53 90-110
NUP93^SOL + SUMO-NUP53 90-110
NUP93 SOL

**Fig.S50. Mapping of the minimal NUP53 region sufficient for NUP93$^{SOL}$ binding.** SEC and SDS-PAGE analysis corresponding to Fig. 12A. (**A**) Domain structure of NUP53 with gray bars indicating truncation construct boundaries analyzed. The NUP53$^{R2}$ peptide is indicated by a red bar. The binding of NUP53 truncation constructs to NUP93$^{SOL}$, as assessed by SEC and SDS-PAGE, is summarized and colored according to the measured effect: no effect (green, +++), weak effect (yellow, ++), moderate effect (orange, +), and abolished binding (red, -). (**B**, **C**) SEC profiles are shown for nups individually (dark blue and cyan) and after their preincubation (colored according to the measured effect, as in (A). (**D**) SEC profile of wildtype NUP93$^{SOL}$ (dark blue) is shown as reference. SEC profiles of NUP93$^{SOL}$ preincubated with SUMO-NUP53 truncation constructs are colored according to the measured effect, as in (A). Dashed lines indicate the peak elution volumes across the offset dimension. The gray bar indicates fractions that were resolved on SDS-PAGE gels and visualized by Coomassie staining. All SEC profiles were obtained using a Superdex 200 10/300 GL column.

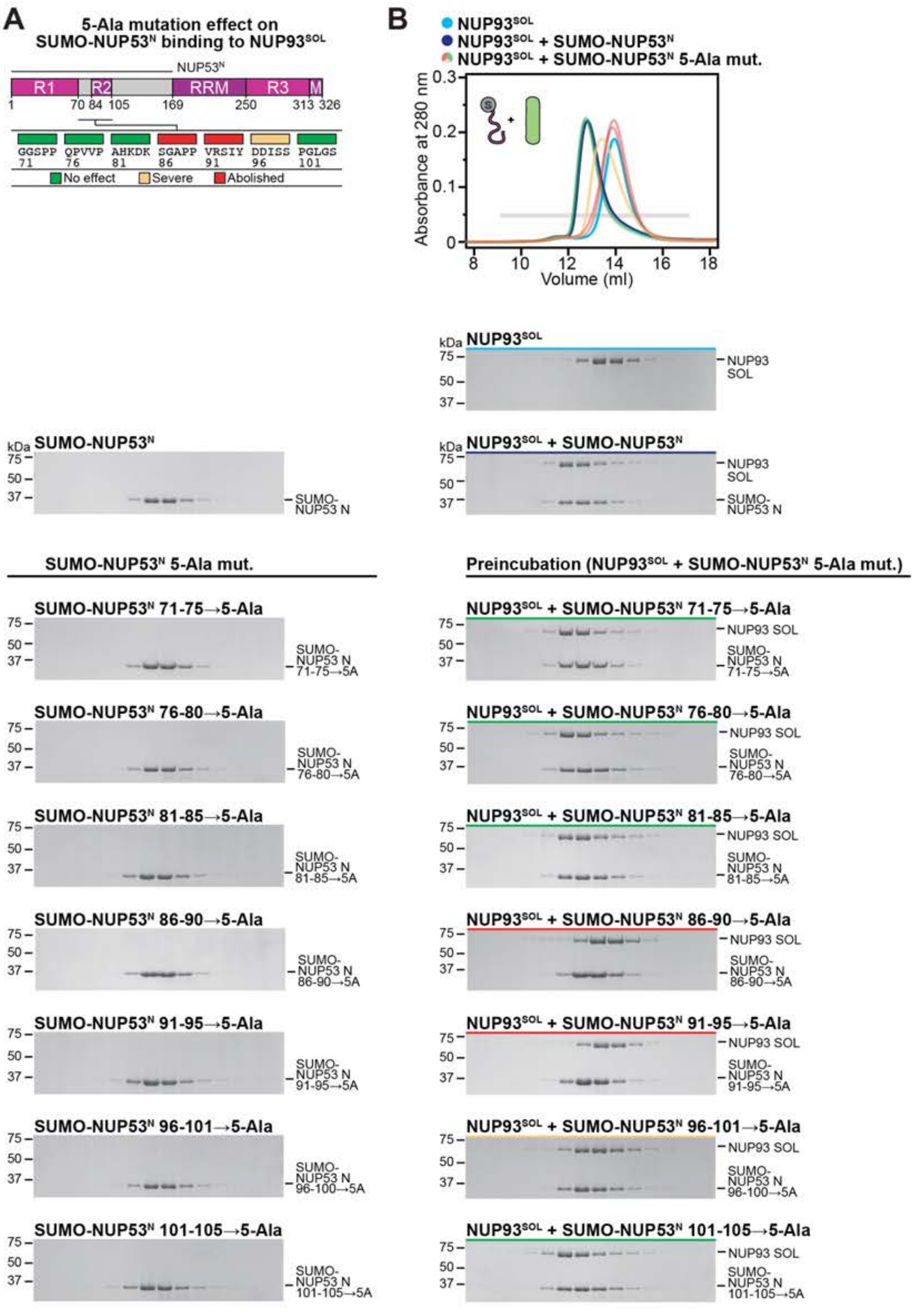
A
5-Ala mutation effect on
SUMO-NUP53N binding to NUP93SOL
NUP53N
R1 R2 RRM R3 M
1 70 84 105 169 250 313 326
GGSPP 71
QPVVP 76
AHKDK 81
SGAPP 86
VRSIY 91
DDISS 96
PGLGS 101
No effect
Severe
Abolished
B
NUP93SOL
NUP93SOL + SUMO-NUP53N
NUP93SOL + SUMO-NUP53N 5-Ala mut.
Absorbance at 280 nm
Volume (ml)
kDa NUP93SOL
NUP93 SOL
kDa SUMO-NUP53N
SUMO-NUP53 N
kDa NUP93SOL + SUMO-NUP53N
SUMO-NUP53N 5-Ala mut.
Preincubation (NUP93SOL + SUMO-NUP53N 5-Ala mut.)
SUMO-NUP53N 71-75→5-Ala
NUP93SOL + SUMO-NUP53N 71-75→5-Ala
SUMO-NUP53N 76-80→5-Ala
NUP93SOL + SUMO-NUP53N 76-80→5-Ala
SUMO-NUP53N 81-85→5-Ala
NUP93SOL + SUMO-NUP53N 81-85→5-Ala
SUMO-NUP53N 86-90→5-Ala
NUP93SOL + SUMO-NUP53N 86-90→5-Ala
SUMO-NUP53N 91-95→5-Ala
NUP93SOL + SUMO-NUP53N 91-95→5-Ala
SUMO-NUP53N 96-101→5-Ala
NUP93SOL + SUMO-NUP53N 96-101→5-Ala
SUMO-NUP53 N 96-100→5A
SUMO-NUP53N 101-105→5-Ala
NUP93SOL + SUMO-NUP53N 101-105→5-Ala
SUMO-NUP53 N 101-105→5A

**Fig.S51. 5-Ala scanning mutagenesis of NUP53 sequence for residues necessary for NUP93$^{SOL}$ binding.** SEC and SDS-PAGE analysis corresponding to Fig. 12B. (**A**) Domain structure of NUP53. Mutations were introduced into the NUP53$^{N}$ construct indicated by the black line above the domain structure. The positions of the 5-Ala mutated residues are indicated above the primary NUP53 sequence with boxes colored according to the measured effect on Nup192 binding: no effect (green), moderate effect (yellow), and abolished binding (red). (**B**) SEC profiles of wildtype NUP93$^{SOL}$ alone (cyan) and NUP93$^{SOL}$ preincubated with wildtype SUMO-NUP53$^{N}$ (dark blue) are shown. SEC profiles of SUMO-NUP53$^{N}$ 5-Ala mutants preincubated with NUP93$^{SOL}$ are colored according to the measured effect, as in (A). The gray bar indicates fractions that were resolved on SDS-PAGE gels and visualized by Coomassie staining. All SEC profiles were obtained using a Superdex 200 10/300 GL column.

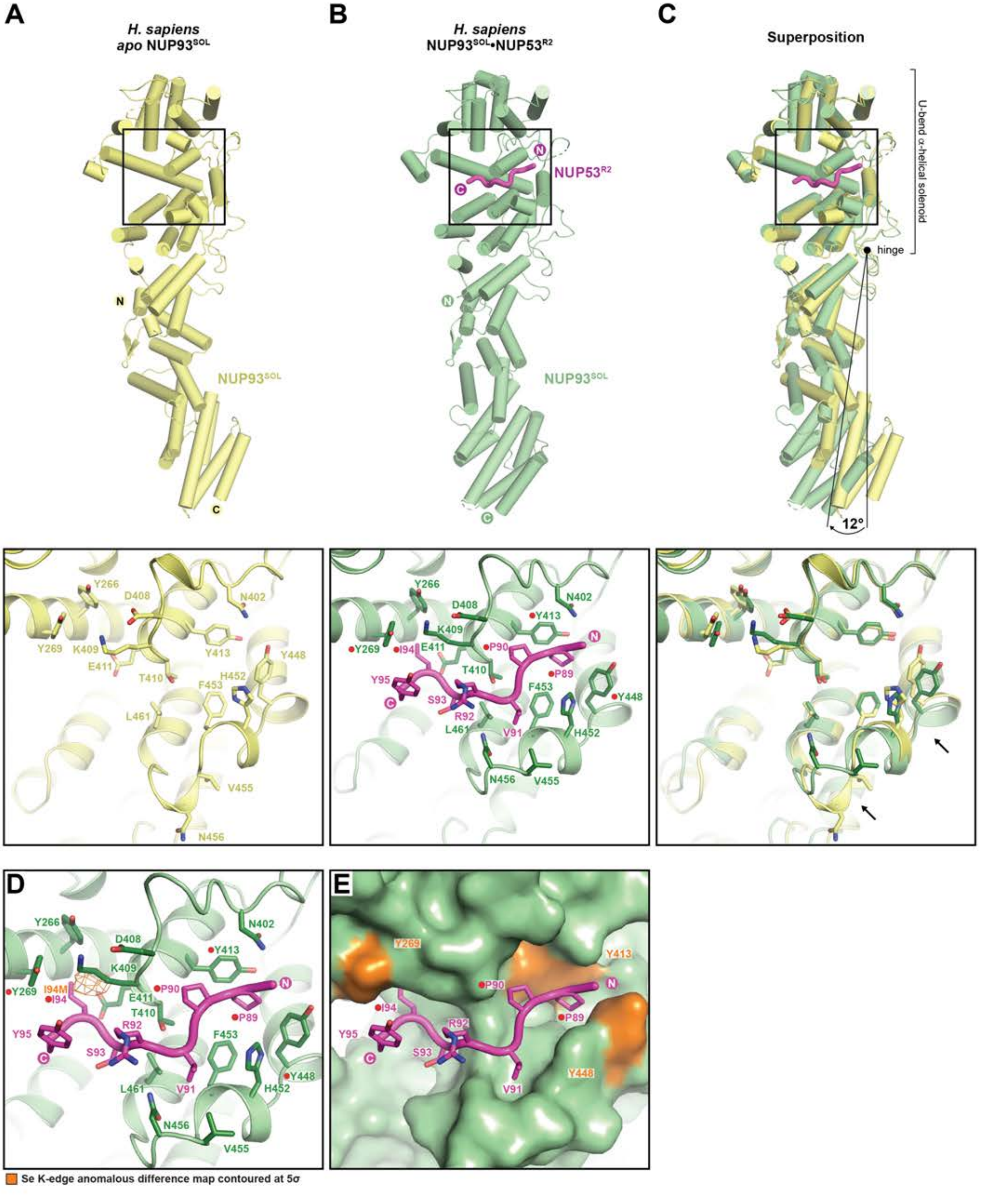
A
H. sapiens
apo NUP93SOL
N
NUP93SOL
C
B
H. sapiens
NUP93SOL•NUP53R2
NUP53R2
C
N
NUP93SOL
C
Superposition
U-bend α-helical solenoid
hinge
12°
Y266
D408
N402
Y269
K409
Y413
E411
T410
H452
Y448
F453
L461
V455
N456
I94
P90
P89
Y95
S93
R92
V91
D
I94M
E
Se K-edge anomalous difference map contoured at 5σ

**Fig.S52. Crystallographic analysis of *apo* NUP93$^{SOL}$ and NUP93$^{SOL}$•NUP53$^{R2}$.** Cartoon representation of *apo* NUP93$^{SOL}$ and NUP93$^{SOL}$•NUP53$^{R2}$ crystal structures. Inset boxes indicate magnified views of the NUP53 binding site of (**A**) the 2.0 Å crystal structure of *apo* NUP93$^{SOL}$ (yellow), (**B**) the 3.4 Å co-crystal structure of the NUP93$^{SOL}$•NUP53$^{R2}$ complex (green and magenta), and (**C**) a structural superposition of the two structures. Red circles indicate residues found by mutational analysis to affect NUP93$^{SOL}$ binding to NUP53 (Fig. 12, fig. S53). Black arrows indicate conformational changes in the loops adjacent to the binding site induced by the binding of the NUP53$^{R2}$ peptide. A global 12° displacement respect to the and N-terminal U-bend $\alpha$-helical solenoid, pivoted about a connecting hinge loop, is observed. (**D**) Anomalous difference Fourier map (orange) calculated from X-ray diffraction data collected at the selenium anomalous peak wavelengths for NUP93$^{SOL}$•NUP53$^{R2}$ I94M SeMet-derivatized co-crystals and superposed with the cartoon and stick representation of the NUP93$^{SOL}$•NUP53$^{R2}$ structure (green and magenta), validating the sequence assignment of the NUP53$^{R2}$ peptide. (**E**) NUP53$^{R2}$ shown in cartoon and ball-and-stick representation. Surface representation of NUP93$^{SOL}$ binding site (pale green) with residues found by mutational analysis to affect NUP53 binding (Fig. 12, fig. S53) highlighted (orange).

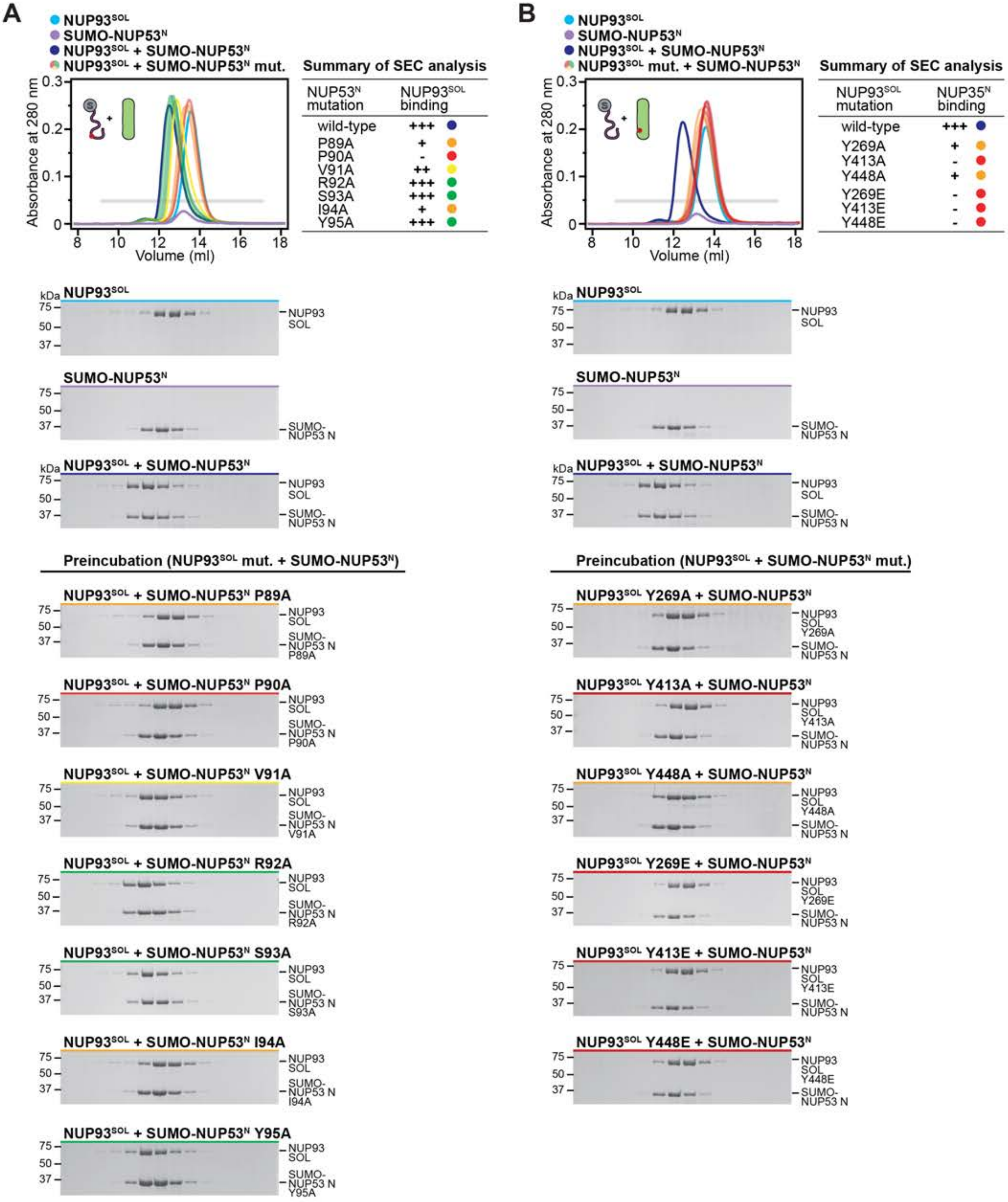
A
NUP93SOL
SUMO-NUP53N
NUP93SOL + SUMO-NUP53N
NUP93SOL + SUMO-NUP53N mut.
Absorbance at 280 nm
Volume (ml)
Summary of SEC analysis
NUP53N mutation | NUP93SOL binding
wild-type +++
P89A +
P90A -
V91A ++
R92A +++
S93A +++
I94A +
Y95A +++
kDa
NUP93 SOL
SUMO-NUP53 N
NUP93SOL + SUMO-NUP53N
Preincubation (NUP93SOL mut. + SUMO-NUP53N)
NUP93SOL + SUMO-NUP53N P89A
NUP93SOL + SUMO-NUP53N P90A
NUP93SOL + SUMO-NUP53N V91A
NUP93SOL + SUMO-NUP53N R92A
NUP93SOL + SUMO-NUP53N S93A
NUP93SOL + SUMO-NUP53N I94A
NUP93SOL + SUMO-NUP53N Y95A
B
NUP93SOL
SUMO-NUP53N
NUP93SOL + SUMO-NUP53N
NUP93SOL mut. + SUMO-NUP53N
Summary of SEC analysis
NUP93SOL mutation | NUP35N binding
wild-type +++
Y269A +
Y413A -
Y448A +
Y269E -
Y413E -
Y448E -
Preincubation (NUP93SOL + SUMO-NUP53N mut.)
NUP93SOL Y269A + SUMO-NUP53N
NUP93SOL Y413A + SUMO-NUP53N
NUP93SOL Y448A + SUMO-NUP53N
NUP93SOL Y269E + SUMO-NUP53N
NUP93SOL Y413E + SUMO-NUP53N
NUP93SOL Y448E + SUMO-NUP53N

Supp Fig. S53

**Fig.S53. Perturbation of the NUP93$^{SOL}$-NUP53 interaction by point mutants.** SEC and SDS-PAGE analysis corresponding to Fig. 12, E and F. SEC profiles of wildtype NUP93$^{SOL}$ alone (cyan) and NUP93$^{SOL}$ preincubated with wildtype SUMO-NUP53$^{N}$ (dark blue) are shown. SEC profiles of (**A**) SUMO-NUP53$^{N}$ mutants preincubated with NUP93$^{SOL}$ (**B**) NUP93$^{SOL}$ mutants preincubated with SUMO-NUP53$^{N}$ are colored according to the measured effect: no effect (green, +++), weak effect (yellow, ++), moderate effect (orange, +), and abolished binding (red, -). The gray bar indicates fractions that were resolved on SDS-PAGE gels and visualized by Coomassie staining. All SEC profiles were obtained using a Superdex 200 10/300 GL column. The results are summarized in the tables (*top right*).

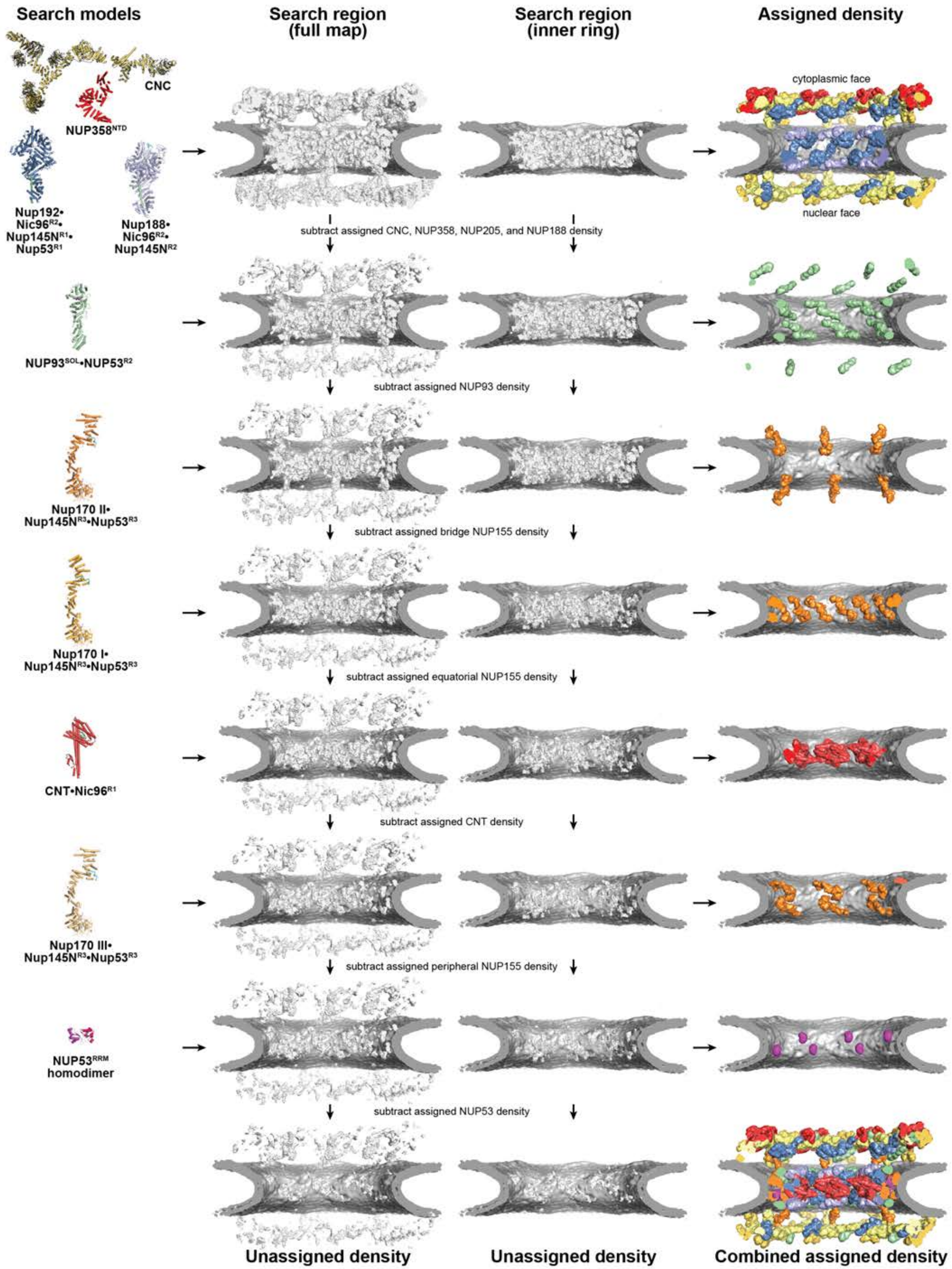

Search models
Search region (full map)
Search region (inner ring)
Assigned density
CNC
NUP358^NTD
Nup192•Nic96^R2•Nup145N^R1•Nup53^R1
Nup188•Nic96^R2•Nup145N^R2
cytoplasmic face
nuclear face
subtract assigned CNC, NUP358, NUP205, and NUP188 density
NUP93^SOL•NUP53^R2
subtract assigned NUP93 density
Nup170 II•Nup145N^R3•Nup53^R3
subtract assigned bridge NUP155 density
Nup170 I•Nup145N^R3•Nup53^R3
subtract assigned equatorial NUP155 density
CNT•Nic96^R1
subtract assigned CNT density
Nup170 III•Nup145N^R3•Nup53^R3
subtract assigned peripheral NUP155 density
NUP53^RRM homodimer
subtract assigned NUP53 density
Unassigned density
Unassigned density
Combined assigned density

Supp Fig. S54

**Fig.S54. Workflow for the sequential quantitative docking of high-resolution structures into the ~12 Å cryo-ET map of the intact human NPC.** Cartoon representations of the high resolution structures used as search models at each step are shown on the left. Global searches were performed in the full map, as well as a map limited to the inner ring. At each step, map density assigned to the search model was removed from the search regions of the subsequent step. Cross-sectional views of the full map and inner ring search region (white) and the nuclear envelope (dark gray) are shown in isosurface representation (*middle*). Isolated contours of the assigned densities are rendered according to the color of the placed search model (*right*). The density left over after all assigned density is removed from the full map and inner ring search regions, along with a colored composite of all the assigned density.

**A**

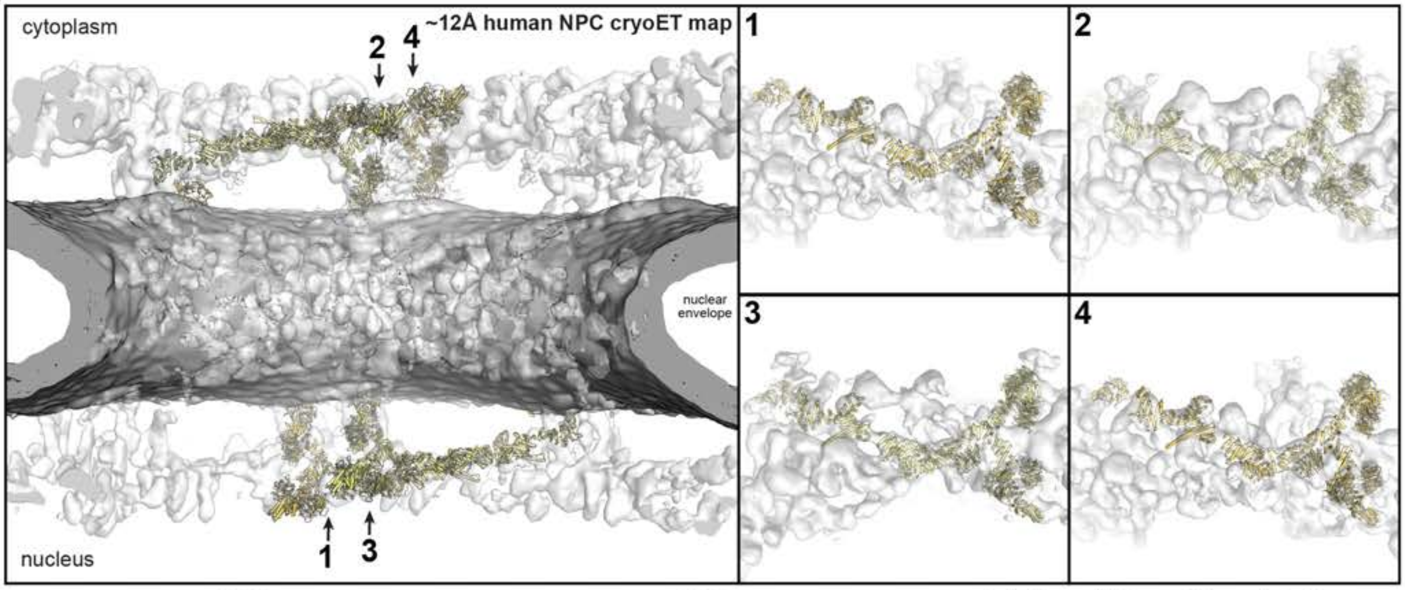

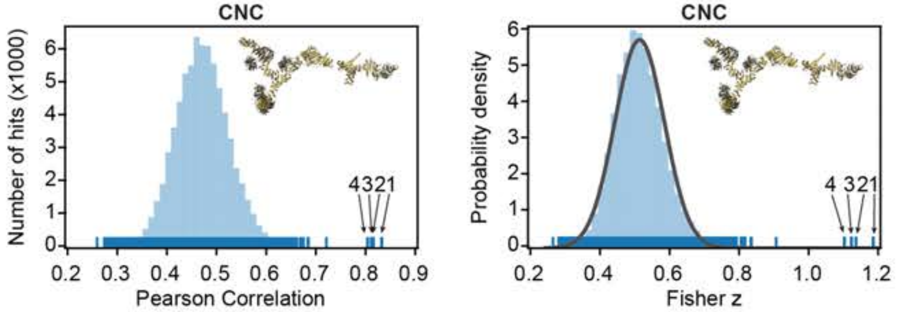

**Intact cryo-ET map fitting statistics**

| solution rank | Pearson correlation | Fisher z | p-value |
|---|---|---|---|
| 1 | 0.827 | 1.18 | 1.7 e-21 |
| 2 | 0.810 | 1.13 | 1.0 e-18 |
| 3 | 0.806 | 1.11 | 5.2 e-18 |
| 4 | 0.798 | 1.09 | 6.3 e-17 |

**B**

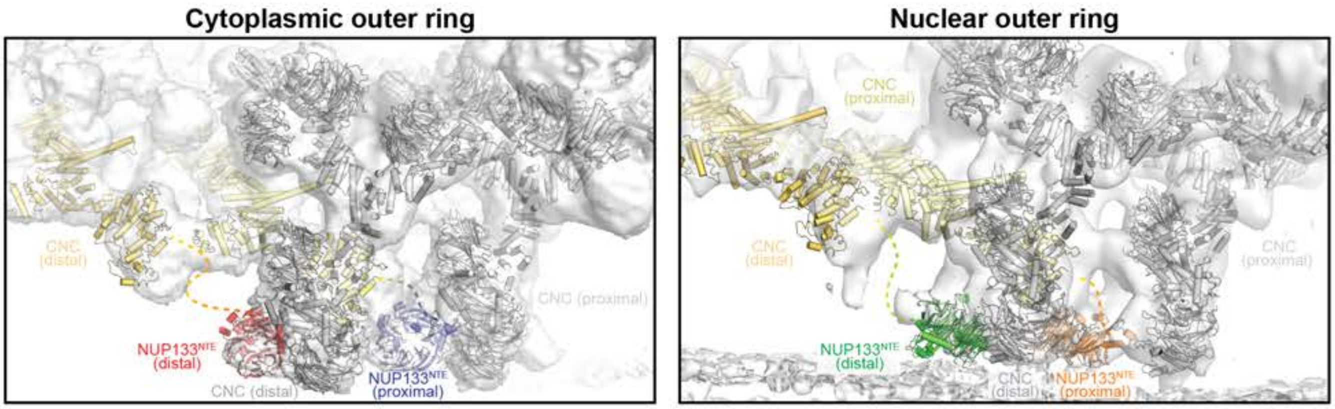

**C**

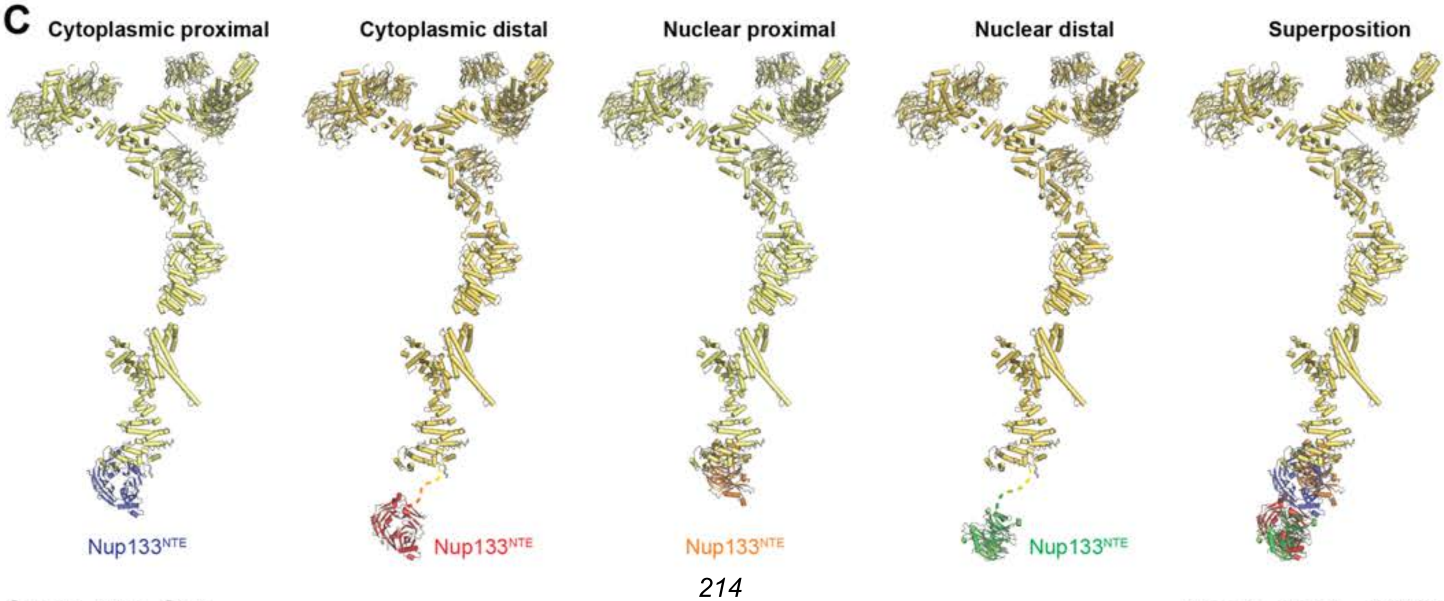

**Fig.S55. Docking of the coat nup complex structures into the ~12 Å cryo-ET map of the intact human NPC**. Resolution-matched simulated cryo-EM densities of side-chain resolution structures of the coat nup complex (CNC) were quantitatively docked into an ~12 Å sub-tomogram averaged cryo-ET map of the intact human NPC (EMD-XXXX) (*41*). The search model was adopted from (*30*) and was composed of the yeast CNC-hexamer (PDB ID 4XMM) (*26*), the NUP84•NUP133 hetero-dimer (PDB ID 3I4R) (*21*), NUP43 (PDB ID 4I79) (*27*), NUP37 (PDB ID 4FHM) (*22*), and $NUP133^{NTE}$ (PDB ID 1XKS) (*12*) crystal structures. (**A**) The placement of unique solutions in a single spoke is shown (*top left*). Arrows and numbers indicate the accepted solutions and their corresponding rank. Closeup views of the accepted solutions are shown on the right, with the rank indicated on the top left corner of the box. Isosurface representations of the nuclear envelope and the NPC are colored in dark gray and white, respectively. The docked structures are displayed as cartoons. Rug plots (blue) and histograms (light blue) of Pearson correlation scores and derived Fisher z scores fit with a normalized Gaussian curve (black) from a global search with 1 million random initial placements are shown (*bottom left* and *middle*, respectively). A tabular summary of the accepted solutions fitting statistics, along with one-tailed p-values calculated from the Fisher z score distribution, is shown (*bottom right*). (**B**) The $NUP133^{NTE}$ rigid-body fit to the cryo-ET map was further locally optimized at each position: cytoplasmic distal (red), cytoplasmic proximal (dark blue), nuclear distal (green), and nuclear proximal (orange). (**C**) Comparison of cartoon representations of the CNC docked at different positions, with $NUP133^{NTE}$ polypeptide chains colored as in (B). To account for the difference in diameter of the proximal and distal CNC rings, the $NUP133^{NTE}$ position and distance from the rest of the structured NUP133 (yellow) varies significantly between the distal and proximal positions.

**A**

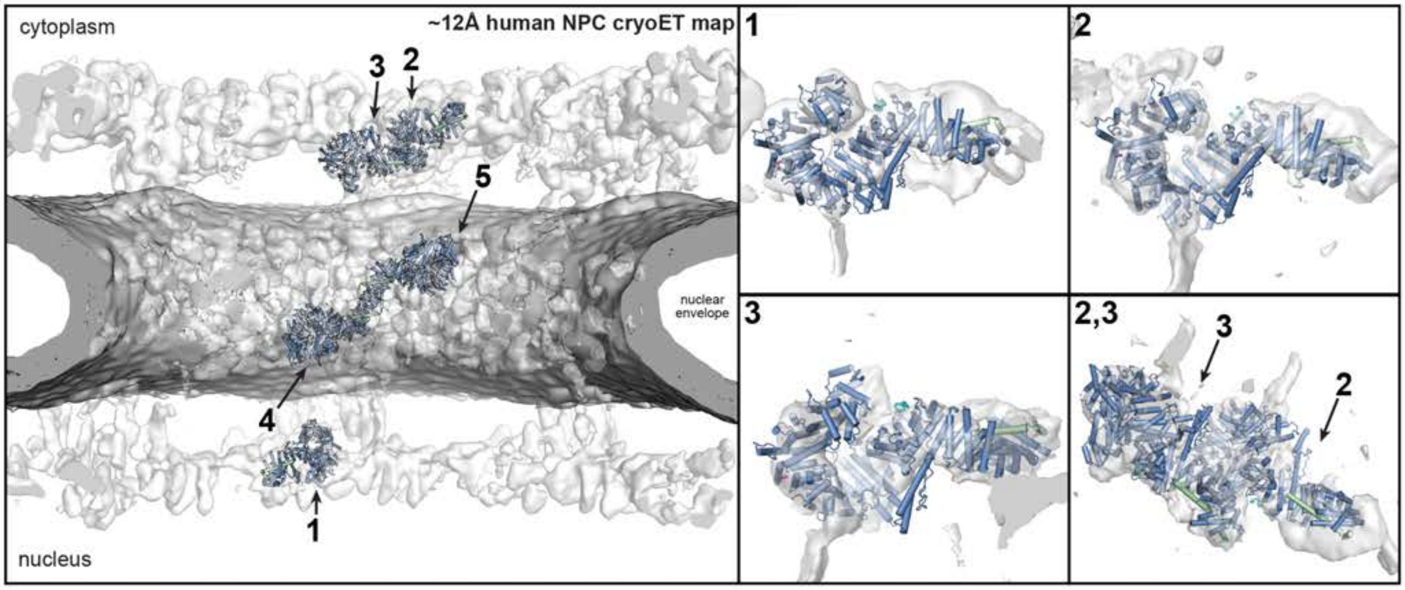

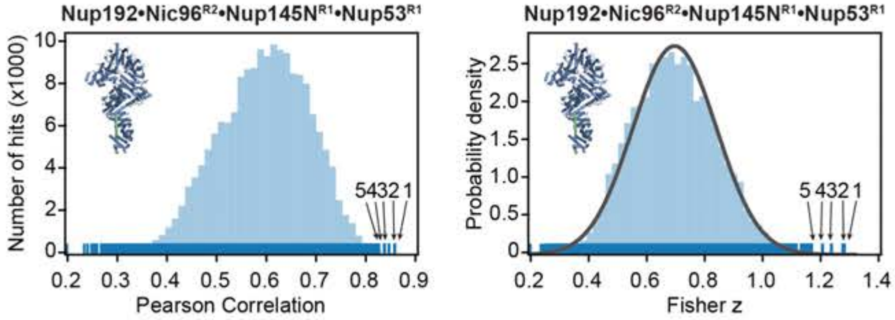

**Intact cryo-ET map fitting statistics**

| solution rank | Pearson correlation | Fisher z | p-value |
|---|---|---|---|
| 1 | 0.855 | 1.27 | 2.6 e-5 |
| 2 | 0.854 | 1.27 | 2.9 e-5 |
| 3 | 0.843 | 1.23 | 8.7 e-5 |
| 4 | 0.833 | 1.20 | 2.1 e-4 |
| 5 | 0.823 | 1.17 | 5.2 e-4 |

**B**

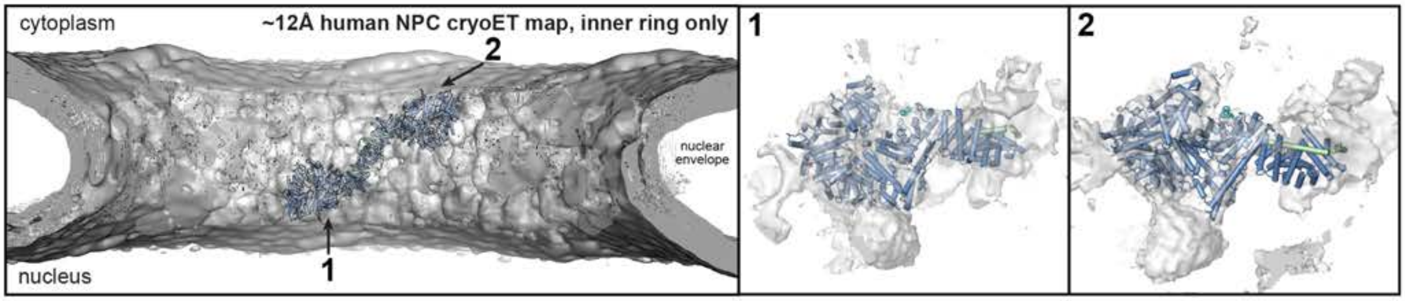

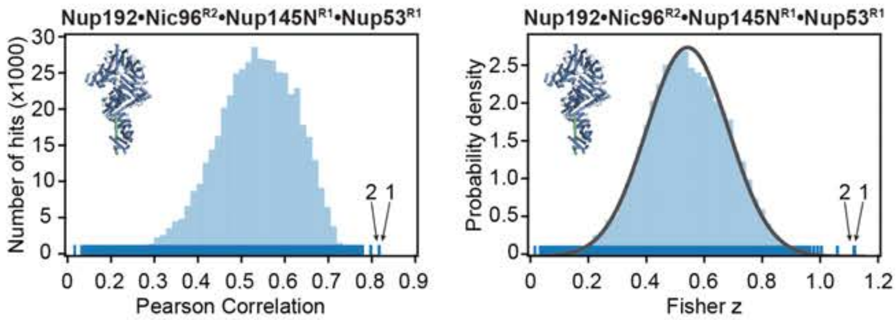

**Inner ring cryo-ET map fitting statistics**

| solution rank | Pearson correlation | Fisher z | p-value |
|---|---|---|---|
| 1 | 0.807 | 1.12 | 2.9 e-3 |
| 2 | 0.806 | 1.11 | 3.1 e-3 |

**Fig.S56. Docking of the single particle cryo-EM Nup192•Nic96$^{R2}$•Nup145N$^{R1}$•Nup53$^{R1}$ structure into the ~12 Å cryo-ET map of the intact human NPC**. Resolution-matched simulated cryo-EM densities of the 3.2 Å Nup192•Nic96$^{R2}$•Nup145N$^{R1}$•Nup53$^{R1}$ single particle cryo-EM structure were quantitatively docked into (**A**) an ~12 Å sub-tomogram averaged cryo-ET map of the intact human NPC (EMD-XXXX) (*41*) and (**B**) a sub-segment of the map comprised solely of the inner ring. The placement of unique solutions in a single spoke is shown (*top left*). Arrows and numbers indicate the accepted solutions and their corresponding rank. Closeup views of the accepted solutions are shown on the right, with the rank indicated on the top left corner of the box. Isosurface representations of the nuclear envelope and the NPC are colored in dark gray and white, respectively. The docked structures are displayed as cartoons. Rug plots (blue) and histograms (light blue) of Pearson correlation scores and derived Fisher z scores fit with a normalized Gaussian curve (black) from a global search with 1 million random initial placements are shown (*bottom left* and *middle*). A tabular summary of the accepted solutions fitting statistics, along with one-tailed p-values calculated from the Fisher z score distribution, is shown (*bottom right*).

**A**

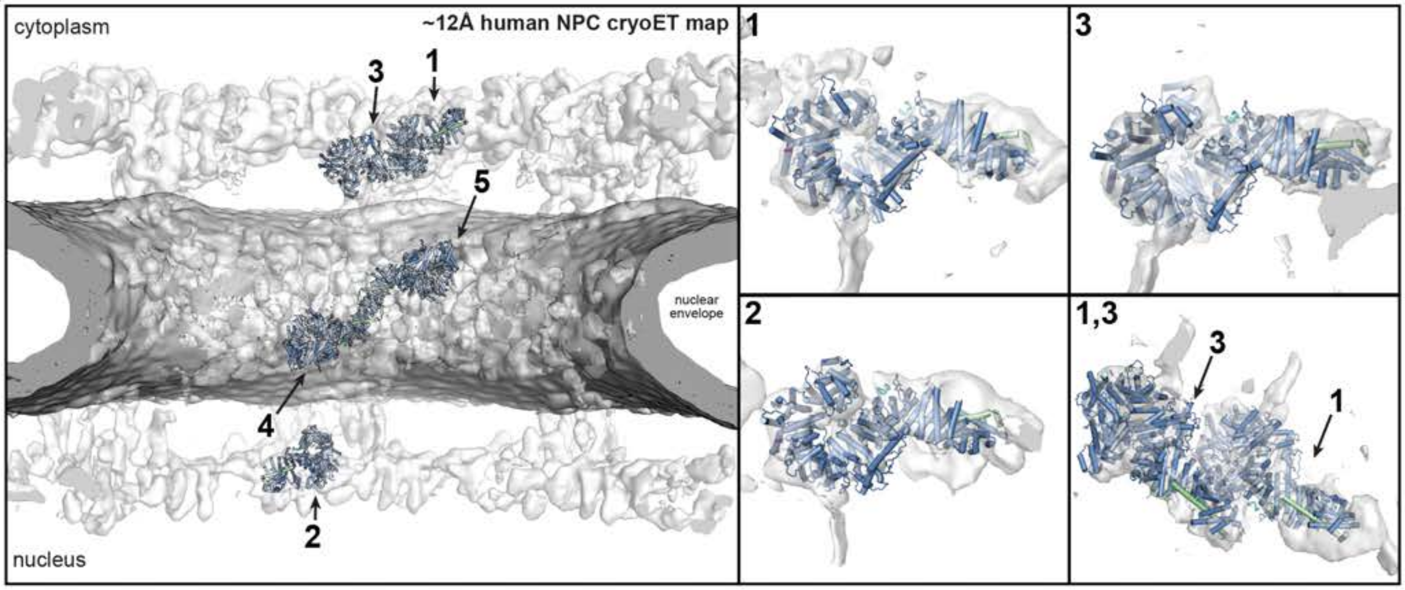

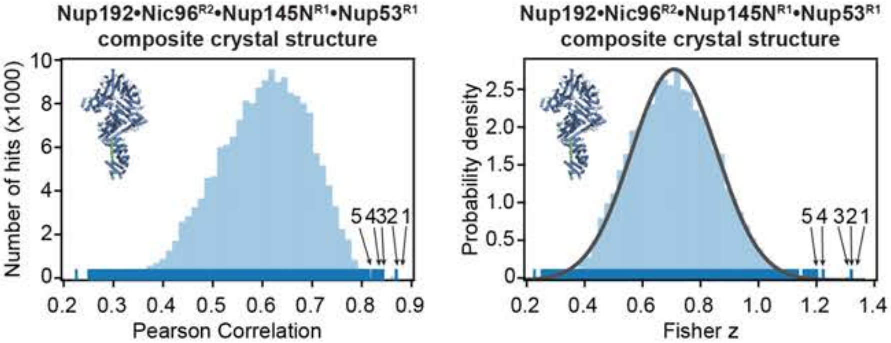

**Intact cryo-ET map fitting statistics**

| solution rank | Pearson correlation | Fisher z | p-value |
|---|---|---|---|
| 1 | 0.866 | 1.31 | 1.8 e-5 |
| 2 | 0.865 | 1.31 | 1.8 e-5 |
| 3 | 0.865 | 1.31 | 1.9 e-5 |
| 4 | 0.839 | 1.22 | 2.5 e-4 |
| 5 | 0.832 | 1.20 | 4.5 e-4 |

**B**

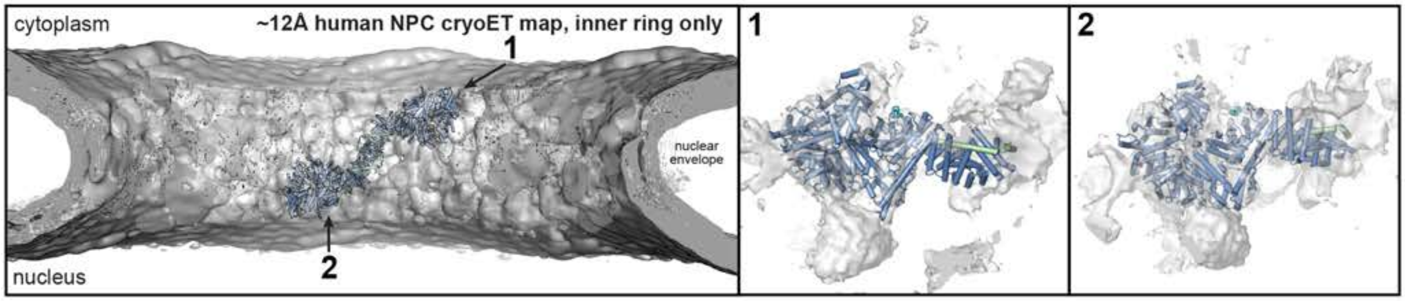

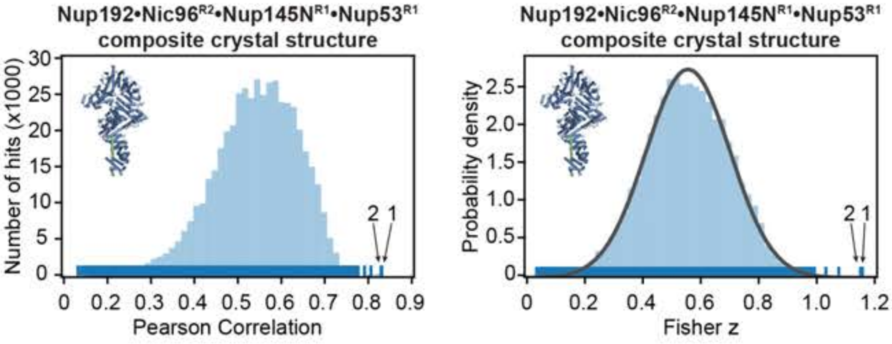

**Inner ring cryo-ET map fitting statistics**

| solution rank | Pearson correlation | Fisher z | p-value |
|---|---|---|---|
| 1 | 0.818 | 1.15 | 2.4 e-3 |
| 2 | 0.816 | 1.14 | 2.6 e-3 |

**Fig.S57. Docking of the composite crystal Nup192•Nic96$^{R2}$•Nup145N$^{R1}$•Nup53$^{R1}$ structure into the ~12 Å cryo-ET map of the intact human NPC**. Resolution-matched simulated cryo-EM densities of the composite crystal Nup192•Nic96$^{R2}$•Nup145N$^{R1}$•Nup53$^{R1}$ structure were quantitatively docked into (**A**) an ~12 Å sub-tomogram averaged cryo-ET map of the intact human NPC (EMD-XXXX) (*41*) and (**B**) a sub-segment of the map comprised solely of the inner ring. The placement of unique solutions in a single spoke is shown (*top left*). Arrows and numbers indicate the accepted solutions and their corresponding rank. Closeup views of the accepted solutions are shown on the right, with the rank indicated on the top left corner of the box. Isosurface representations of the nuclear envelope and the NPC are colored in dark gray and white, respectively. The docked structures are displayed as in cartoon representation. Rug plots (blue) and histograms (light blue) of Pearson correlation scores and derived Fisher z scores fit with a normalized Gaussian curve (black) from a global search with 1 million random initial placements are shown (*bottom left* and *middle*). A tabular summary of the accepted solutions fitting statistics, along with one-tailed p-values calculated from the Fisher z score distribution, is shown (*bottom right*). The crystallized conformation of Nup192 found in the composite crystal structure fits the cryo-ET map density with greater confidence than the conformation of Nup192 found in single particle cryo-EM structure (Fig. S56).

**A**

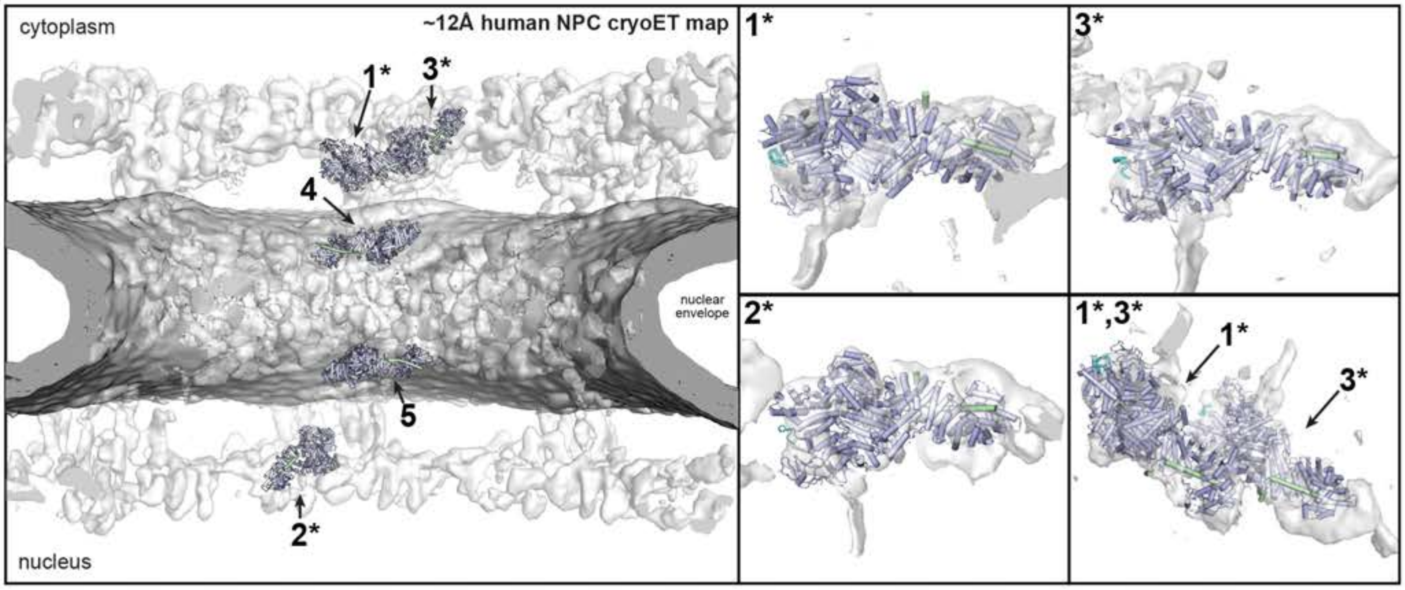

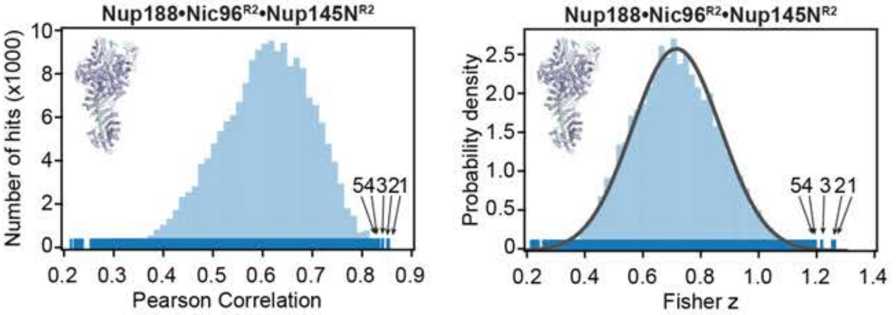

| Intact cryo-ET map fitting statistics | | | |
|---|---|---|---|
| solution rank | Pearson correlation | Fisher z | p-value |
| 1* | 0.850 | 1.26 | 1.8 e-4 |
| 2* | 0.848 | 1.25 | 2.1 e-4 |
| 3* | 0.838 | 1.21 | 5.0 e-4 |
| 4 | 0.830 | 1.19 | 8.7 e-4 |
| 5 | 0.830 | 1.19 | 8.8 e-4 |

**B**

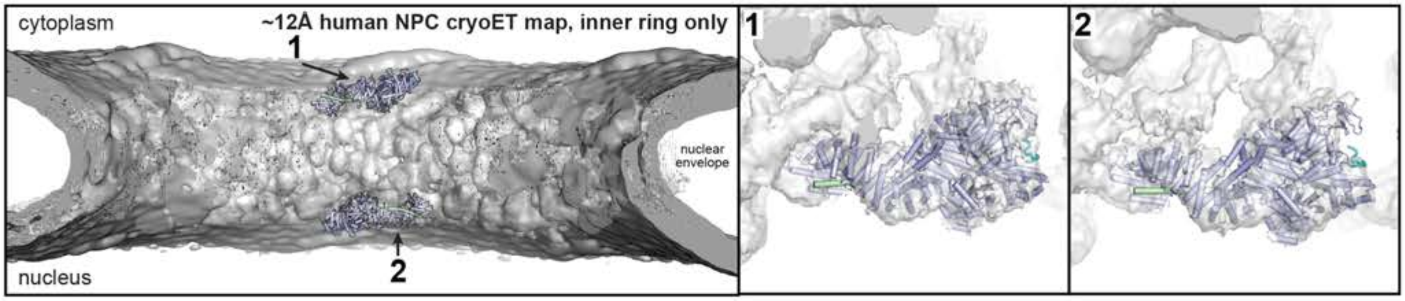

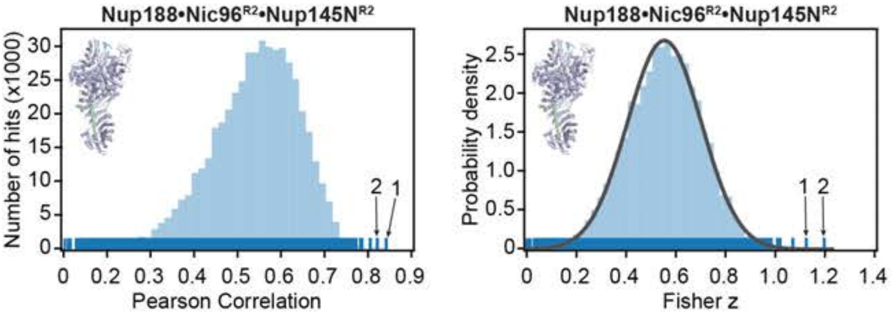

| Inner ring cryo-ET map fitting statistics | | | |
|---|---|---|---|
| solution rank | Pearson correlation | Fisher z | p-value |
| 1 | 0.831 | 1.19 | 8.0 e-6 |
| 2 | 0.807 | 1.12 | 6.3 e-5 |

**Fig.S58. Docking of the single particle cryo-EM Nup188•Nic96$^{R2}$•Nup145N$^{R2}$ structure into the ~12 Å cryo-ET map of the intact human NPC**. Resolution-matched simulated cryo-EM densities of the 2.8 Å Nup188•Nic96$^{R2}$•Nup145N$^{R2}$ single particle cryo-EM structure were quantitatively docked into (**A**) the ~12 Å sub-tomogram averaged cryo-ET map of the intact human NPC (EMD-XXXX) (*41*) and (**B**) a sub-segment of the map comprised solely of the inner ring. The placement of unique solutions in a single spoke is shown (*top left*). Arrows and numbers indicate the accepted and tentative solutions and their corresponding rank. Asterisks indicate that solutions in the outer rings are tentative and shown for illustrative purpose, as quantitative docking in the ~12 Å cryo-ET map places the Nup192 complex with greater confidence into the outer ring positions (figs. S56 and S57). Closeup views of the accepted and tentative solutions are shown on the right, with the rank indicated on the top left corner of the box. Isosurface representations of the nuclear envelope and the NPC are colored in dark gray and white, respectively. The docked structures are displayed in cartoon representation. Rug plots (blue) and histograms (light blue) of Pearson correlation scores and derived Fisher z scores fit with a normalized Gaussian curve (black) from a global search with 1 million random initial placements are shown (*bottom left* and *middle*). A tabular summary of the solutions fitting statistics, along with one-tailed p-values calculated from the Fisher z score distribution, is shown (*bottom right*).

**A**

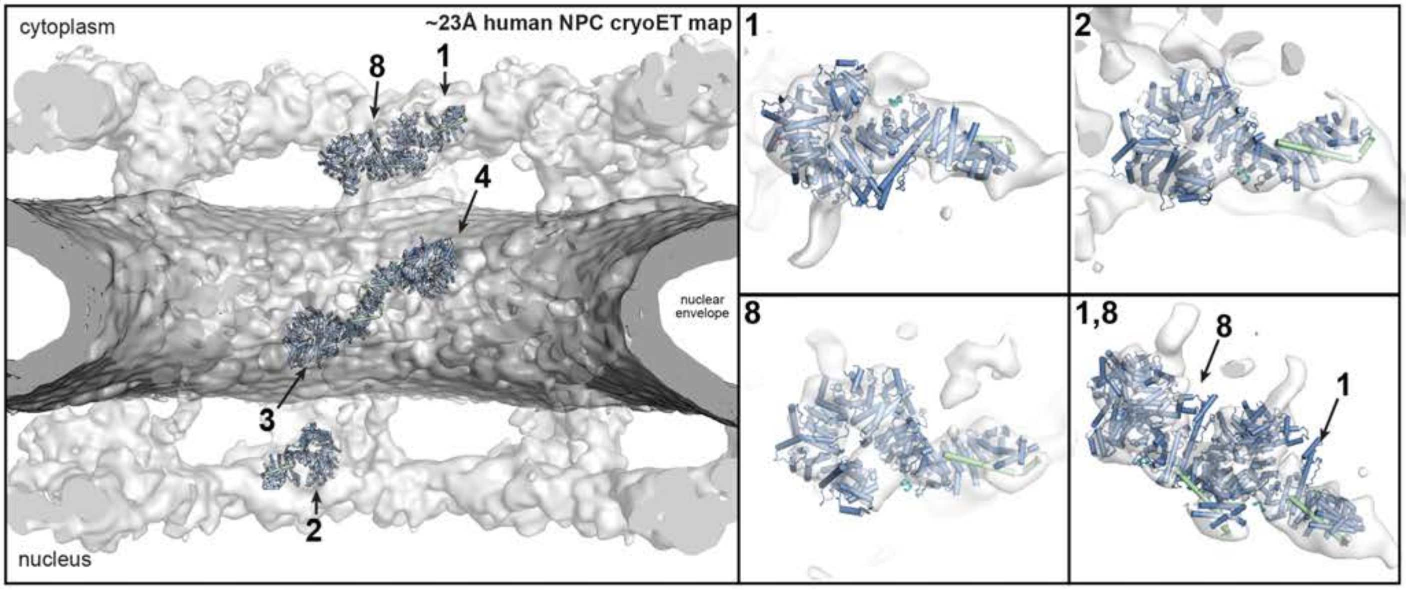

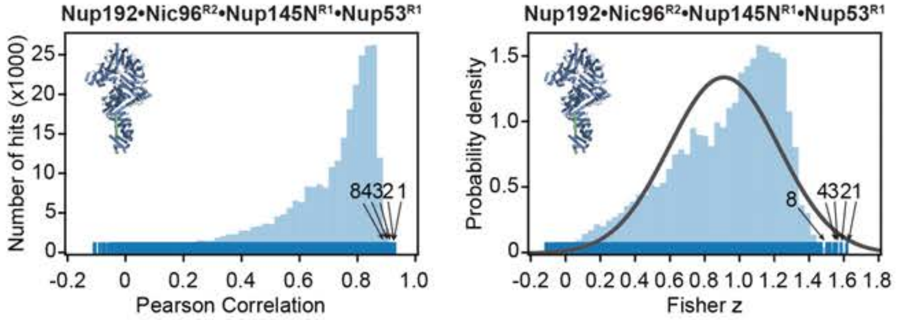

**Intact cryo-ET map fitting statistics**

| solution rank | Pearson correlation | Fisher z | p-value |
|---|---|---|---|
| 1 | 0.922 | 1.61 | 1.3 e-2 |
| 2 | 0.918 | 1.58 | 1.6 e-2 |
| 3 | 0.914 | 1.55 | 2.0 e-2 |
| 4 | 0.911 | 1.53 | 2.2 e-2 |
| 8 | 0.898 | 1.46 | 3.8 e-2 |

**B**

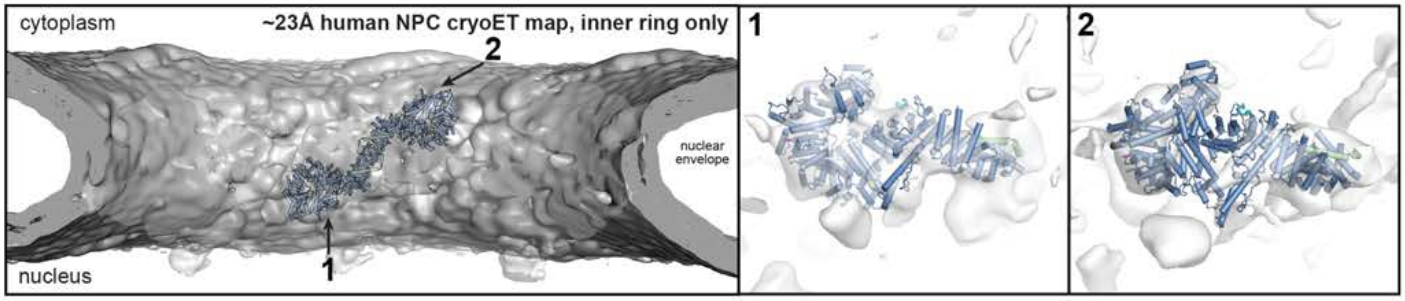

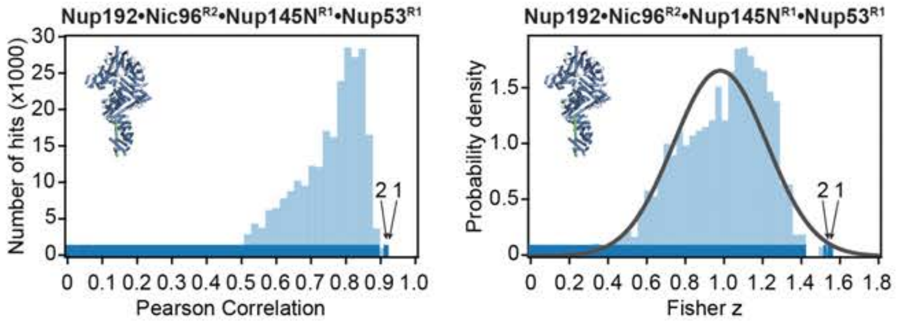

**Inner ring cryo-ET map fitting statistics**

| solution rank | Pearson correlation | Fisher z | p-value |
|---|---|---|---|
| 1 | 0.914 | 1.55 | 8.7 e-3 |
| 2 | 0.911 | 1.53 | 1.0 e-2 |

**Fig.S59. Docking of the single particle cryo-EM Nup192•Nic96$^{R2}$•Nup145N$^{R1}$•Nup53$^{R1}$ structure into an ~23 Å cryo-ET map of the intact human NPC**. Resolution-matched simulated cryo-EM densities of the 3.2 Å Nup192•Nic96$^{R2}$•Nup145N$^{R1}$•Nup53$^{R1}$ single particle cryo-EM structure were quantitatively docked into (**A**) the 23 Å sub-tomogram averaged cryo-ET map of the intact human NPC (EMD-3103) (*33*) and (**B**) a sub-segment of the map comprised solely of the inner ring. The placement of unique solutions in a single spoke is shown (*top left*). Arrows and numbers indicate the accepted solutions and their corresponding rank. Closeup views of the accepted solutions are shown on the right, with the rank indicated on the top left corner of the box. Isosurface representation of the nuclear envelope and the NPC are colored in dark gray and white, respectively. The docked structures are displayed as in cartoon representation. Rug plots (blue) and histograms (light blue) of Pearson correlation scores and derived Fisher z scores fit with a normalized Gaussian curve (black) from a global search with 1 million random initial placements are shown (bottom left and middle, respectively). A tabular summary of the accepted solutions fitting statistics, along with one-tailed p-values calculated from the Fisher z score distribution, is shown (*bottom right*). Top-scoring solutions were identified for four of the five positions identified by docking into the higher resolution ~12 Å cryo-ET map (Fig. S56). The cytoplasmic outer ring proximal position was ranked as the 8th highest scoring solution.

A

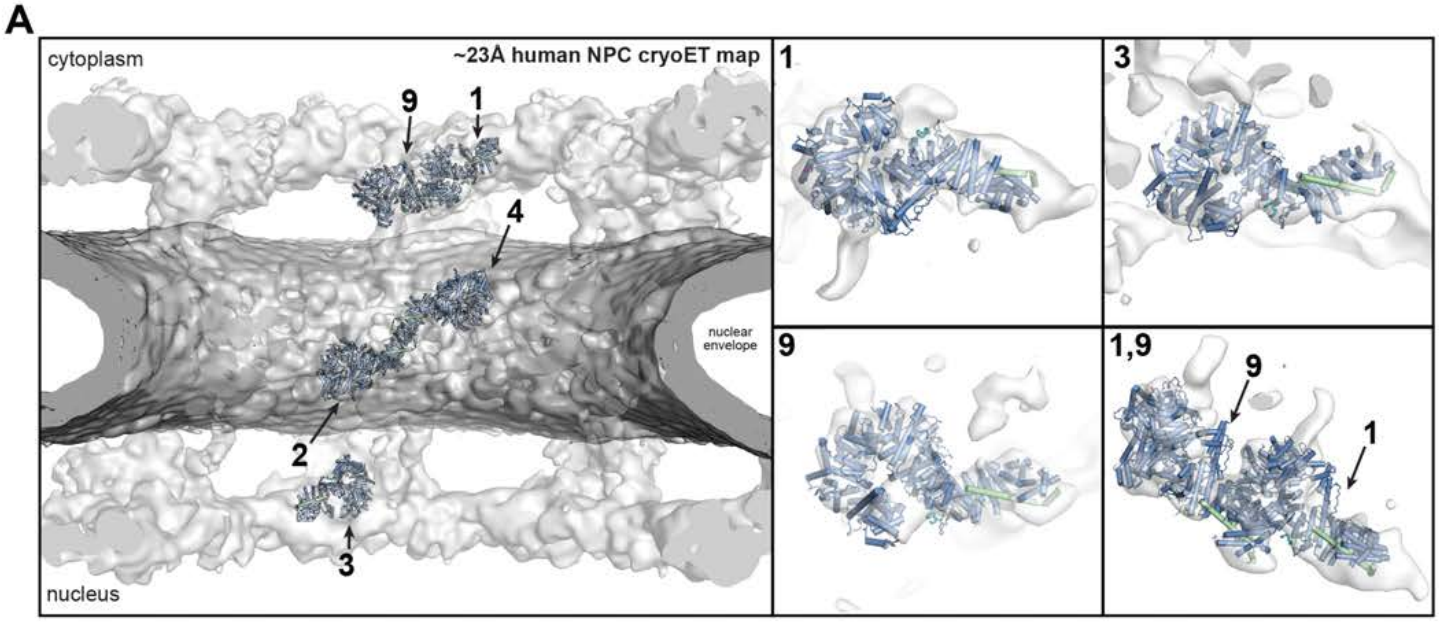

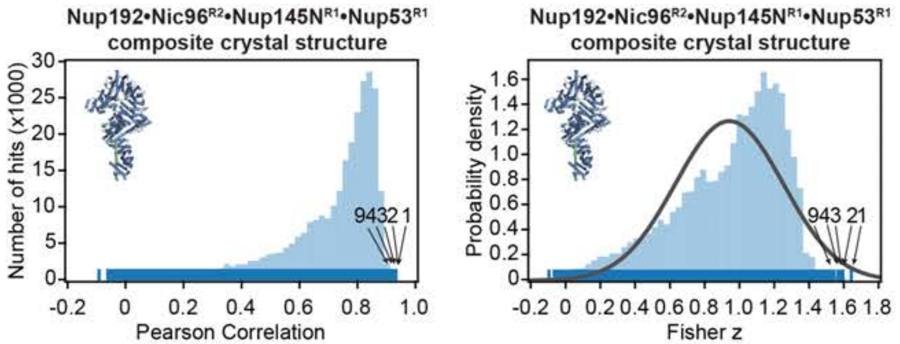

**Intact cryo-ET map fitting statistics**

| solution rank | Pearson correlation | Fisher z | p-value |
|---|---|---|---|
| 1 | 0.927 | 1.63 | 3.4 e-3 |
| 2 | 0.919 | 1.58 | 6.4 e-3 |
| 3 | 0.917 | 1.57 | 7.5 e-3 |
| 4 | 0.915 | 1.56 | 8.8 e-3 |
| 9 | 0.903 | 1.49 | 1.9 e-2 |

B

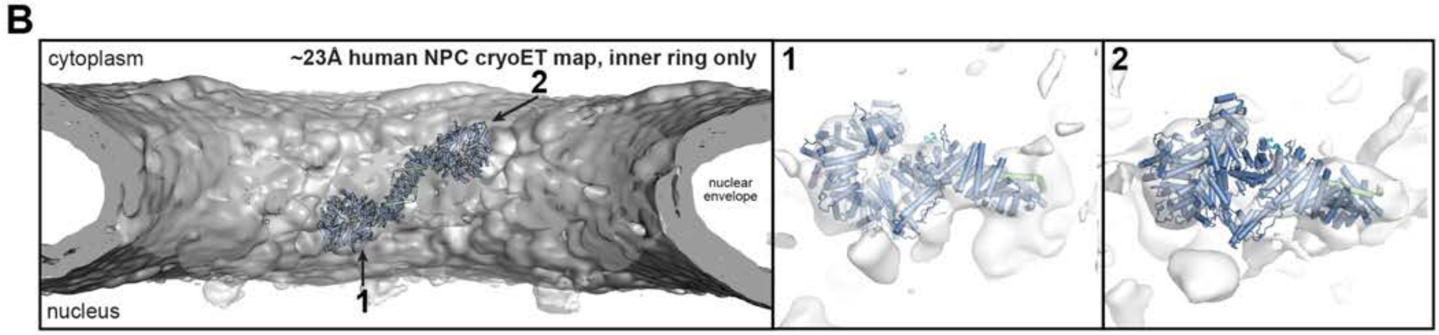

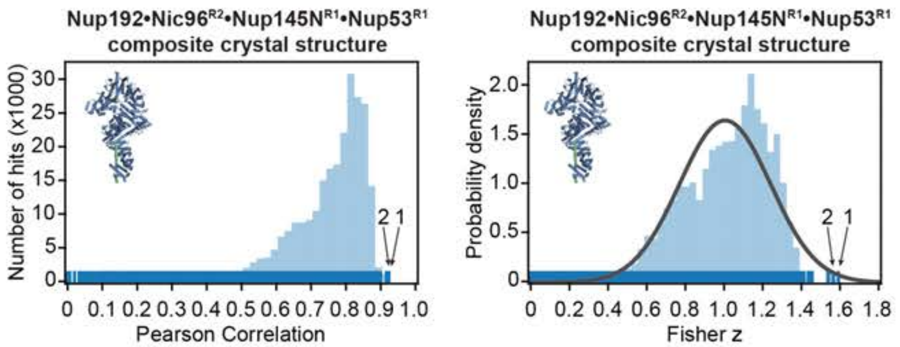

**Inner ring cryo-ET map fitting statistics**

| solution rank | Pearson correlation | Fisher z | p-value |
|---|---|---|---|
| 1 | 0.919 | 1.58 | 6.4 e-3 |
| 2 | 0.915 | 1.56 | 8.8 e-3 |

**Fig.S60. Docking of the composite crystal Nup192•Nic96$^{R2}$•Nup145N$^{R1}$•Nup53$^{R1}$ structure into an ~23 Å cryo-ET map of the intact human NPC**. Resolution-matched simulated cryo-EM densities of the composite crystal Nup192•Nic96$^{R2}$•Nup145N$^{R1}$•Nup53$^{R1}$ structure were quantitatively docked into (**A**) the 23 Å sub-tomogram averaged cryo-ET map of the intact human NPC (EMD-3103) (*33*) and (**B**) a sub-segment of the map comprised solely of the inner ring. The placement of unique solutions in a single spoke is shown (*top left*). Arrows and numbers indicate the accepted solutions and their corresponding rank. Closeup views of the accepted solutions are shown on the right, with the rank indicated on the top left corner of the box. Isosurface representations of the nuclear envelope and the NPC are colored in dark gray and white, respectively. The docked structures are displayed in cartoon representation. Rug plots (blue) and histograms (light blue) of Pearson correlation scores and derived Fisher z scores fit with a normalized Gaussian curve (black) from a global search with 1 million random initial placements are shown (*bottom left* and *middle*). A tabular summary of the accepted solutions fitting statistics, along with one-tailed p-values calculated from the Fisher z score distribution, is shown (*bottom right*). Top-scoring solutions were identified for four of the five positions identified by docking into the higher resolution ~12 Å cryo-ET map (Fig. S57). The cytoplasmic outer ring proximal position was ranked as the 9th highest scoring solution.

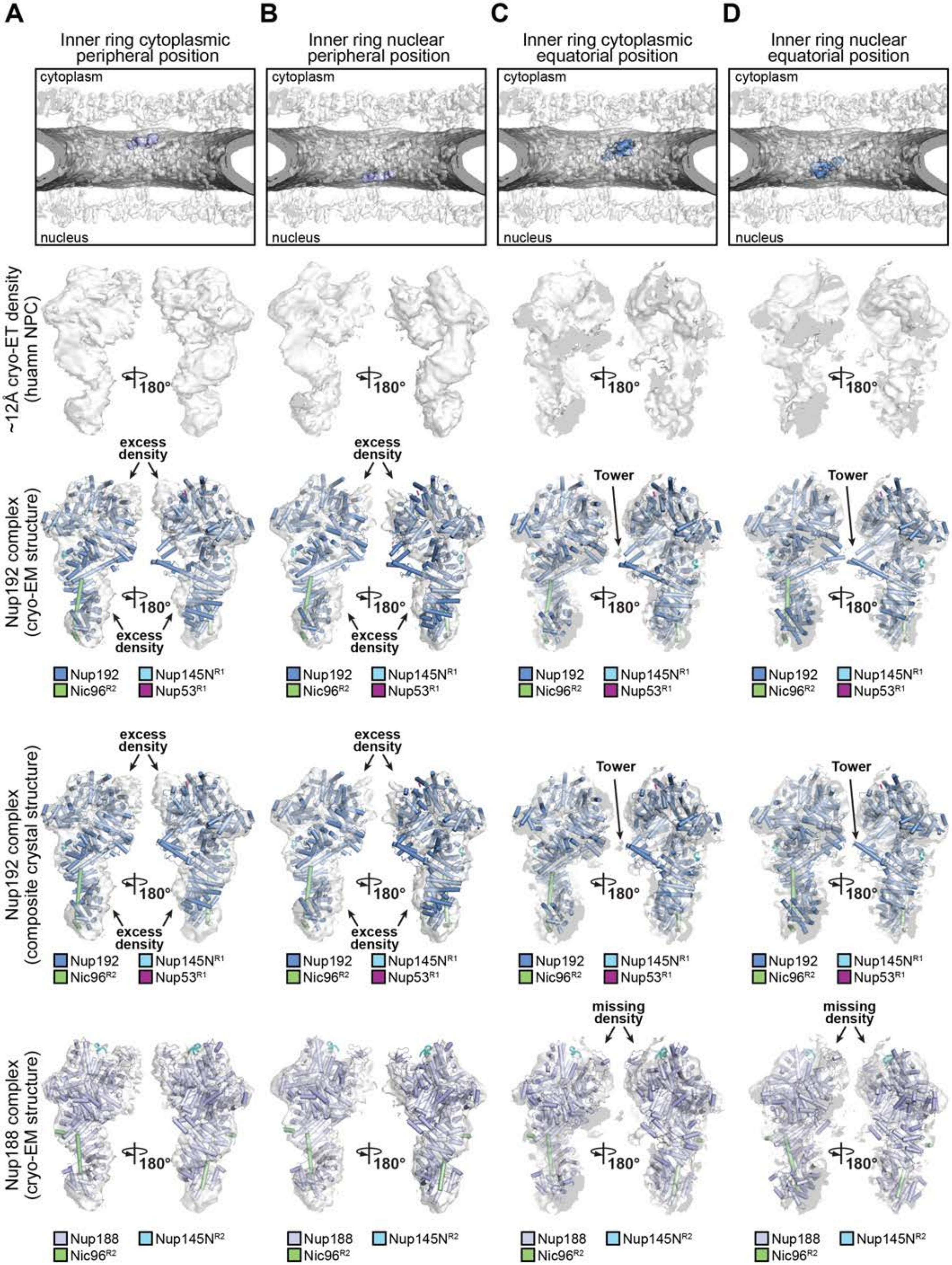

Supp Fig. S61

Hoelz et al., 2021

**Fig.S61. Comparison of the Nup192 and Nup188 complex placement in the inner ring of the ~12 Å cryo-ET map of the intact human NPC.** Question mark-shaped cryo-ET densities carved out from (**A**) cytoplasmic peripheral, (**B**) nuclear peripheral, (**C**) cytoplasmic equatorial, and (**D**) nuclear equatorial positions of the inner ring of an ~12 Å sub-tomogram averaged cryo-ET map of the intact human NPC (EMD-XXXX) (*41*) where either Nup192•Nic96$^{R2}$•Nup145N$^{R1}$•Nup53$^{R1}$ or Nup188•Nic96$^{R2}$•Nup145N$^{R2}$ complexes were quantitatively docked, as indicated in the boxes (*top*). To compare the goodness of fit by visual inspection, two isosurface representation views of the carved density with cartoons of the quantitatively docked composite crystal Nup192•Nic96$^{R2}$•Nup145N$^{R1}$•Nup53$^{R1}$, single particle cryo-EM Nup192•Nic96$^{R2}$•Nup145N$^{R1}$•Nup53$^{R1}$, or single particle cryo-EM Nup188•Nic96$^{R2}$•Nup145N$^{R2}$ structures are shown. At the peripheral positions, a tentative placement of the two conformations of the Nup192•Nic96$^{R2}$•Nup145N$^{R1}$•Nup53$^{R1}$ complex (single particle cryo-EM and composite crystal structures, respectively) results in unexplained excess density that can be accounted for by the SH3-like domain of Nup188 if the Nup188•Nic96$^{R2}$•Nup145N$^{R2}$ complex (single particle cryo-EM structure) is placed in the same position. Unlike the shorter Nup188 Tower, the longer Nup192 Tower exceeds the cryo-ET density if tentatively placed at the peripheral positions. On the contrary, the longer Tower domain is accommodated by the cryo-ET density at the equatorial position, but the SH3-like domain of the tentatively placed Nup188 exceeds the cryo-ET density at the equatorial position. At the equatorial positions, the Nup192 Tower domain conformation found in the composite crystal Nup192 complex structure fits the cryo-ET density better than the Nup192 Tower domain conformation observed in the single particle cryo-EM Nup192 complex structure.

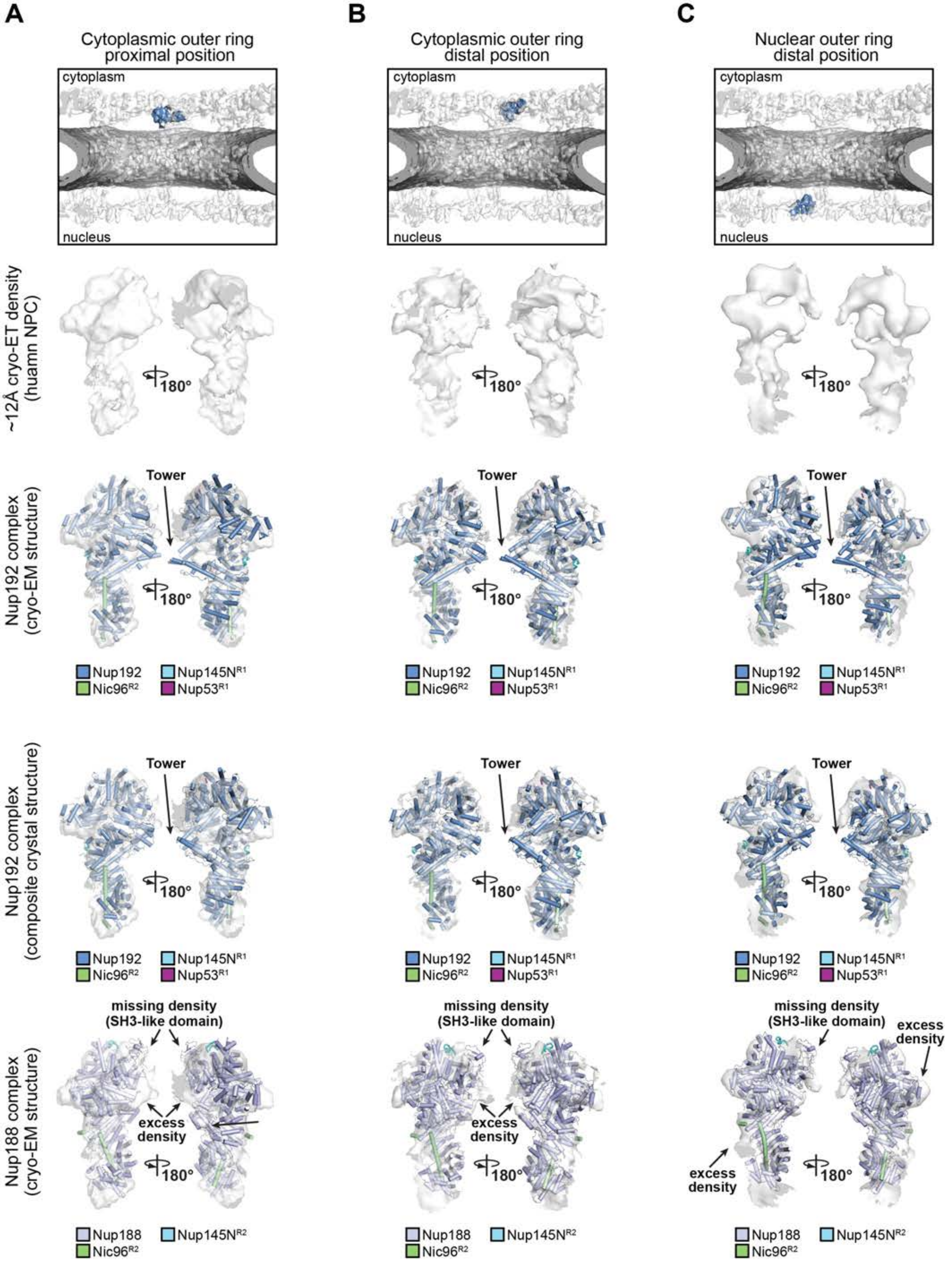

Supp Fig. S62

Hoelz et al., 2021

**Fig.S62. Comparison of the Nup192 and Nup188 complex placement in the outer rings of the ~12 Å cryo-ET map of the intact human NPC.** Question mark-shaped cryo-ET densities carved out from (**A**) cytoplasmic proximal, (**B**) cytoplasmic distal, and (**C**) nuclear distal positions of the outer rings of the ~12 Å sub-tomogram averaged cryo-ET map of the intact human NPC (EMD-XXXX) (*41*) where Nup192•Nic96$^{R2}$•Nup145N$^{R1}$•Nup53$^{R1}$ complexes were quantitatively docked, as indicated in the boxes (*top*). To compare the goodness of fit by visual inspection, two isosurface representation views of the carved density with cartoons of the quantitatively docked composite crystal Nup192•Nic96$^{R2}$•Nup145N$^{R1}$•Nup53$^{R1}$, single particle cryo-EM Nup192•Nic96$^{R2}$•Nup145N$^{R1}$•Nup53$^{R1}$, or single particle cryo-EM Nup188•Nic96$^{R2}$•Nup145N$^{R2}$ structures are shown. The cryo-ET density is exceeded by the Nup188 SH3-like domain if the Nup188•Nic96$^{R2}$•Nup145N$^{R2}$ complex is tentatively placed at all three outer ring positions. On the other hand, the Nup188 $\alpha$-helical solenoid presents a narrower superhelical twist compared to Nup192, which leaves unexplained density in other parts of the cryo-ET density. At all outer ring positions, the Nup192 Tower conformation found in the composite crystal Nup192•Nic96$^{R2}$•Nup145N$^{R1}$•Nup53$^{R1}$ complex structure fits the cryo-ET density better than the Nup192 Tower conformation observed in the single particle cryo-EM Nup192•Nic96$^{R2}$•Nup145N$^{R1}$•Nup53$^{R1}$complex structure.

**A**

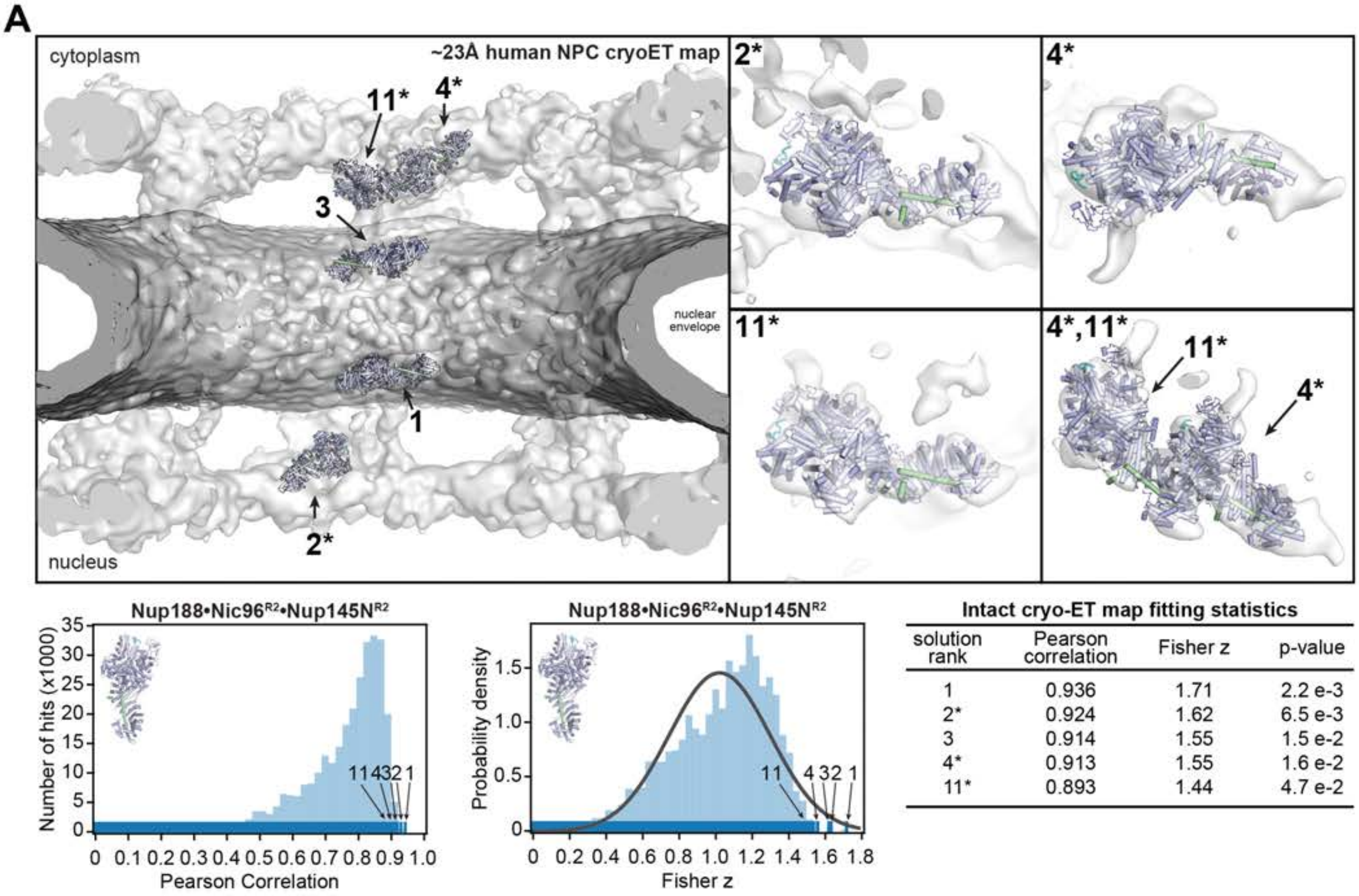

**Intact cryo-ET map fitting statistics**

| solution rank | Pearson correlation | Fisher z | p-value |
|---|---|---|---|
| 1 | 0.936 | 1.71 | 2.2 e-3 |
| 2* | 0.924 | 1.62 | 6.5 e-3 |
| 3 | 0.914 | 1.55 | 1.5 e-2 |
| 4* | 0.913 | 1.55 | 1.6 e-2 |
| 11* | 0.893 | 1.44 | 4.7 e-2 |

**B**

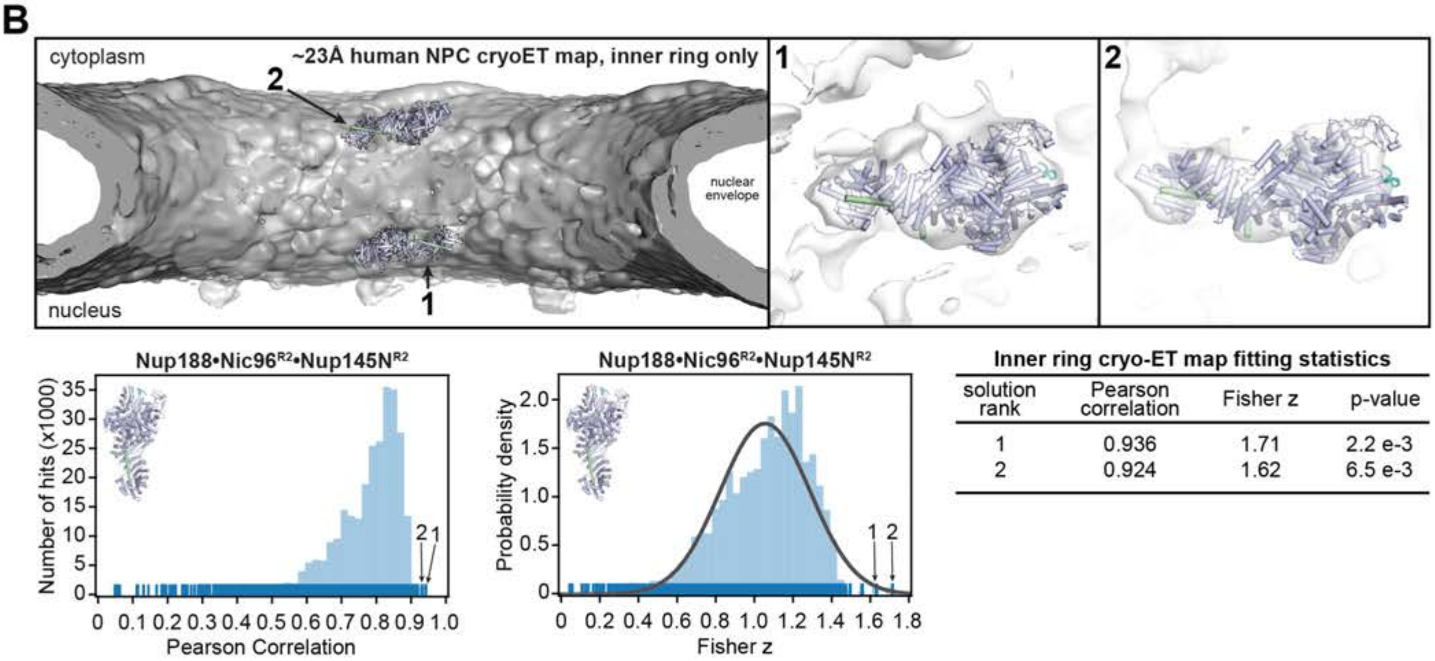

**Inner ring cryo-ET map fitting statistics**

| solution rank | Pearson correlation | Fisher z | p-value |
|---|---|---|---|
| 1 | 0.936 | 1.71 | 2.2 e-3 |
| 2 | 0.924 | 1.62 | 6.5 e-3 |

**Fig.S63. Docking of the single particle cryo-EM Nup188•Nic96$^{R2}$•Nup145N$^{R2}$ structure into an ~23 Å cryo-ET map of the intact human NPC**. Resolution-matched simulated cryo-EM densities of the 2.8 Å Nup188•Nic96$^{R2}$•Nup145N$^{R2}$ single particle cryo-EM structure were quantitatively docked into (**A**) an ~23 Å sub-tomogram averaged cryo-ET map of the intact human NPC (EMD-3103) (*33*) and (**B**) a sub-segment of the map comprised solely of the inner ring. The placement of unique solutions in a single spoke is shown (*top left*). Arrows and numbers indicate the accepted solutions and their corresponding rank. Asterisks indicate that solutions in the outer rings are tentative and shown for illustrative purpose, as quantitative docking in the ~12 Å cryo-ET map places the Nup192 complex with greater confidence into the outer ring positions (figs. S56 and S57). Closeup views of the accepted and tentative solutions are shown on the right, with the rank indicated on the top left corner of the box. Isosurface representations of the nuclear envelope and the NPC are colored in dark gray and white, respectively. The docked structures are displayed in cartoon representation. Rug plots (blue) and histograms (light blue) of Pearson correlation scores and derived Fisher z scores fit with a normalized Gaussian curve (black) from a global search with 1 million random initial placements are shown (*bottom left* and *middle*). A tabular summary of the accepted solutions fitting statistics, along with one-tailed p-values calculated from the Fisher z score distribution, is shown (*bottom right*). Top-scoring solutions were identified for four of the five positions identified by docking into the higher resolution ~12 Å cryo-ET map (fig. S58). The tentative cytoplasmic outer ring proximal position was ranked as the 11th highest scoring solution.

**A**

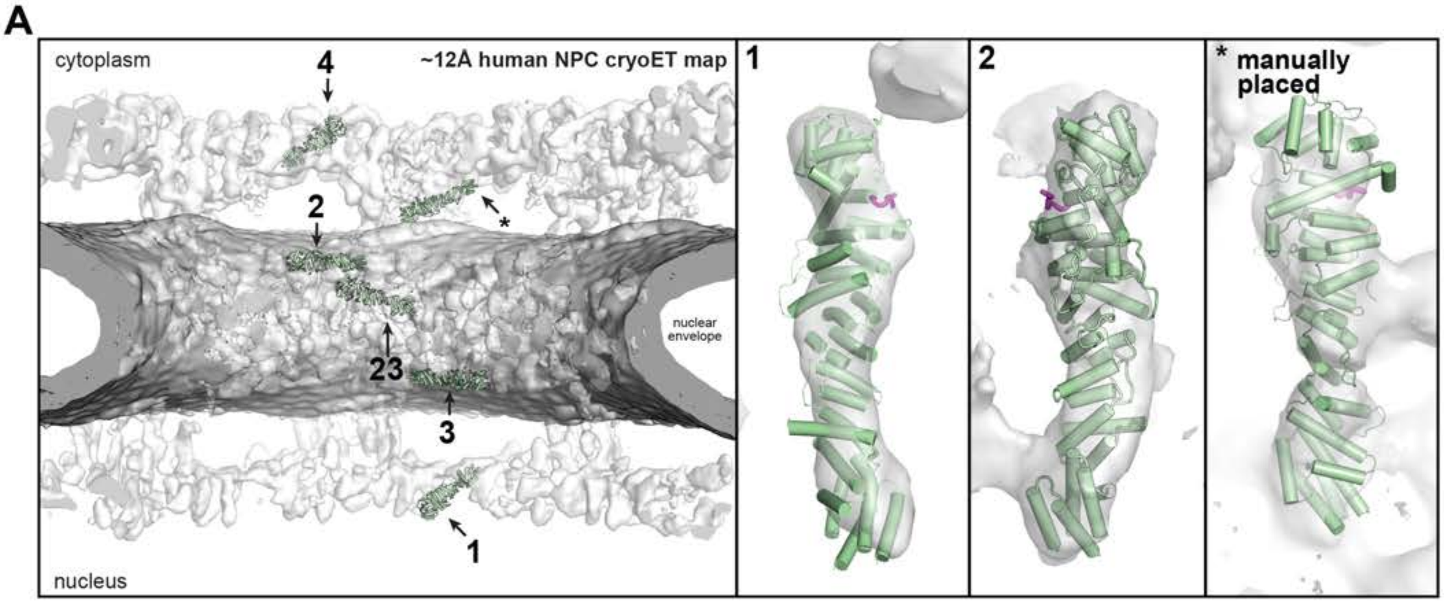

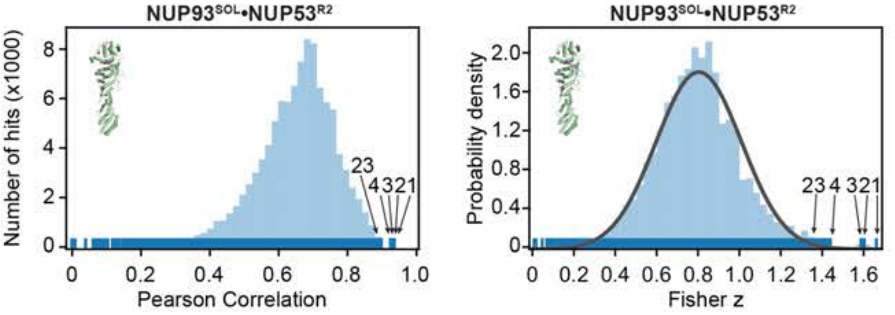

**Subtracted cryo-ET map fitting statistics**

| solution rank | Pearson correlation | Fisher z | p-value |
|---|---|---|---|
| 1 | 0.931 | 1.69 | 1.5 e-5 |
| 2 | 0.923 | 1.61 | 5.2 e-5 |
| 3 | 0.921 | 1.59 | 7.1 e-5 |
| 4 | 0.895 | 1.45 | 1.0 e-3 |
| 23 | 0.878 | 1.37 | 1.3 e-3 |

**B**

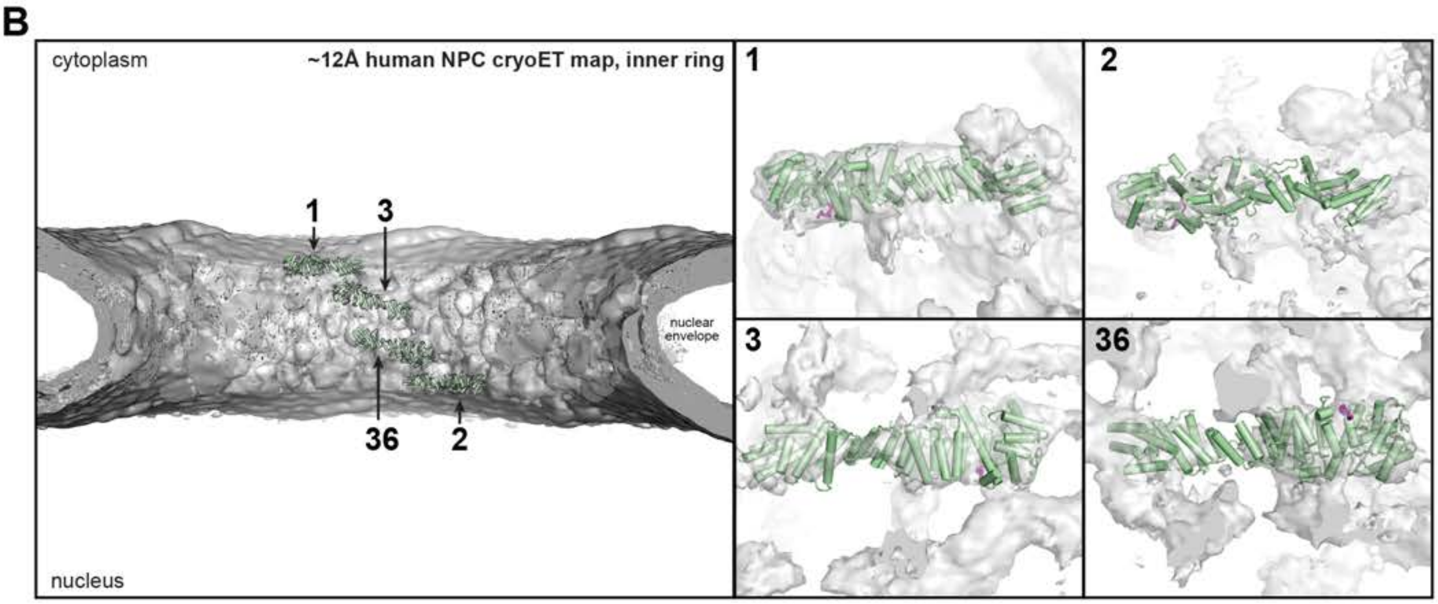

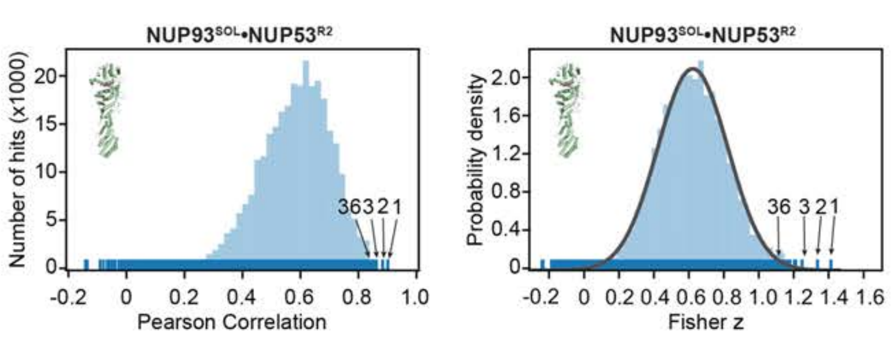

**Subtracted inner ring cryo-ET map fitting statistics**

| solution rank | Pearson correlation | Fisher z | p-value |
|---|---|---|---|
| 1 | 0.889 | 1.42 | 4.3 e-5 |
| 2 | 0.873 | 1.35 | 1.9 e-4 |
| 3 | 0.852 | 1.26 | 9.4 e-4 |
| 36 | 0.820 | 1.16 | 5.3 e-3 |

**Fig.S64. Docking of the NUP93$^{SOL}$•NUP53$^{R2}$ co-crystal structure into the ~12 Å cryo-ET map of the intact human NPC**. Resolution-matched simulated cryo-EM densities of the 3.4 Å NUP93$^{SOL}$•NUP53$^{R2}$ structure were quantitatively docked into (**A**) the ~12 Å sub-tomogram averaged cryo-ET map of the intact human NPC (EMD-XXXX) (*41*) from which cryo-ET density corresponding to hereto docked nups was subtracted (fig. S54) and (**B**) a sub-segment of the map comprised solely of the inner ring from which cryo-ET density corresponding to hereto docked nups was subtracted (fig. S54). The placement of unique solutions in a single spoke is shown (*top left*). Arrows and numbers indicate the accepted solutions and their corresponding rank. The asterisk indicates a copy of NUP93$^{SOL}$•NUP53$^{R2}$ that was placed manually into matching but weak density. The presence of a proximal copy of NUP93$^{SOL}$ is supported by the presence of a proximal copy of NUP205, which binds to NUP93$^{R2}$. Closeup views of the accepted solutions are shown on the right, with the rank indicated on the top left corner of the box. Isosurface representations of the nuclear envelope and the NPC are colored in dark gray and white, respectively. The docked structures are displayed in cartoon representation. Rug plots (blue) and histograms (light blue) of Pearson correlation scores and derived Fisher z scores fit with a normalized Gaussian curve (black) from a global search with 1 million random initial placements are shown (*bottom left* and *middle*). A tabular summary of the solutions fitting statistics, along with one-tailed p-values calculated from the Fisher z score distribution, is shown (*bottom right*). Top scoring solutions in neither full map nor inner ring search regions did not include a nuclear equatorial copy of NUP93$^{SOL}$•NUP53$^{R2}$. However, its position could be inferred by the C2 symmetry relationship with the top-scoring cytoplasmic equatorial copy (ranked 3rd highest) and matched with the 36th highest scoring solution.

**A**

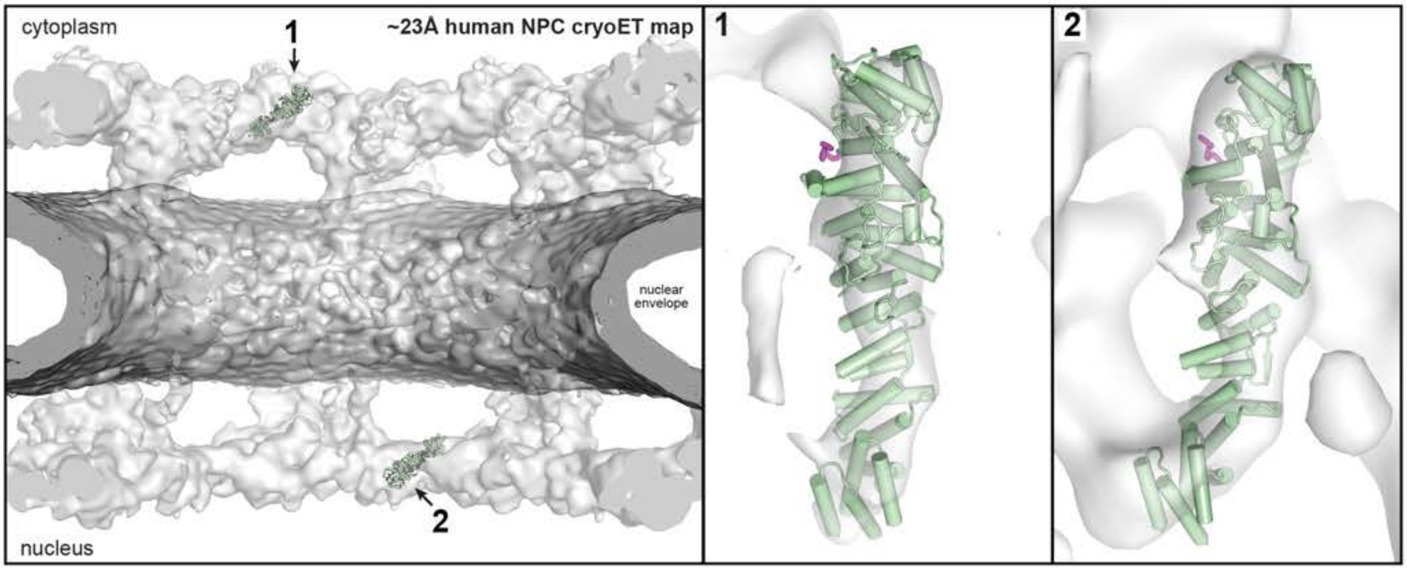

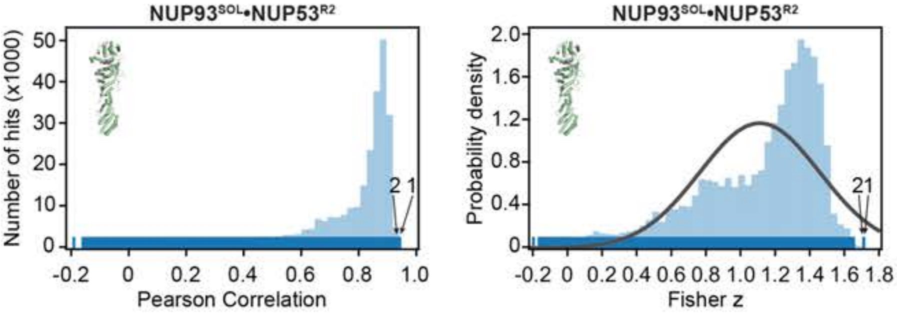

**Subtracted cryo-ET map fitting statistics**

| solution rank | Pearson correlation | Fisher z | p-value |
|---|---|---|---|
| 1 | 0.935 | 1.70 | 4.2 e-2 |
| 2 | 0.935 | 1.70 | 4.2 e-2 |

**B**

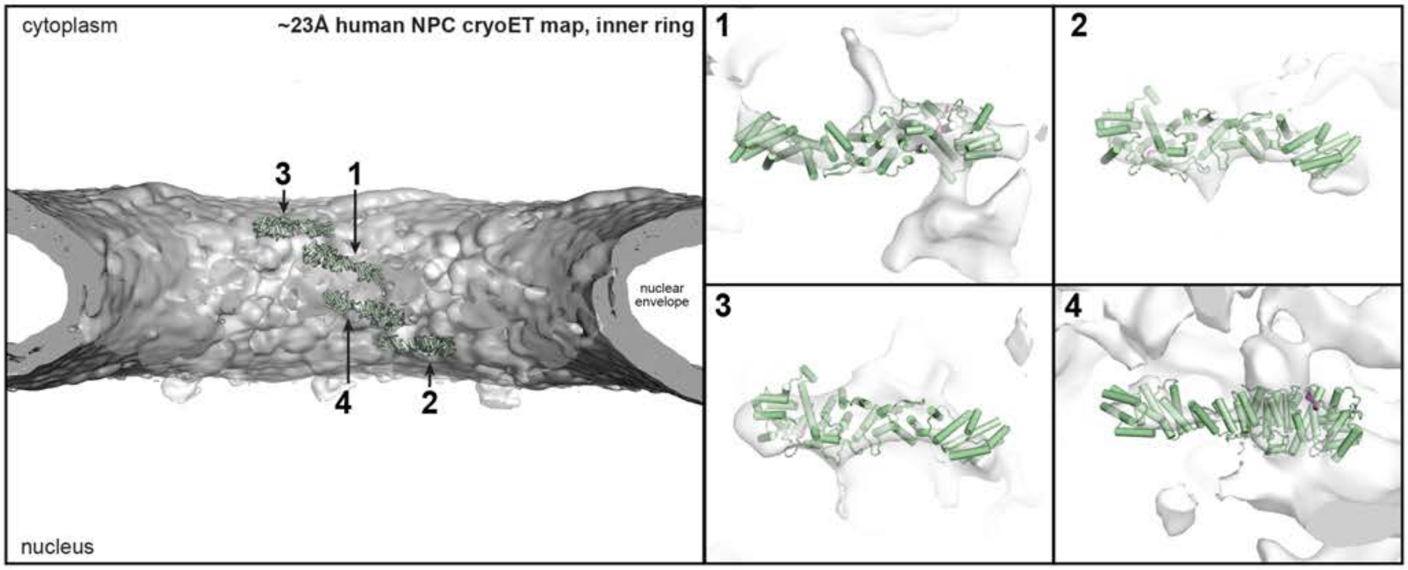

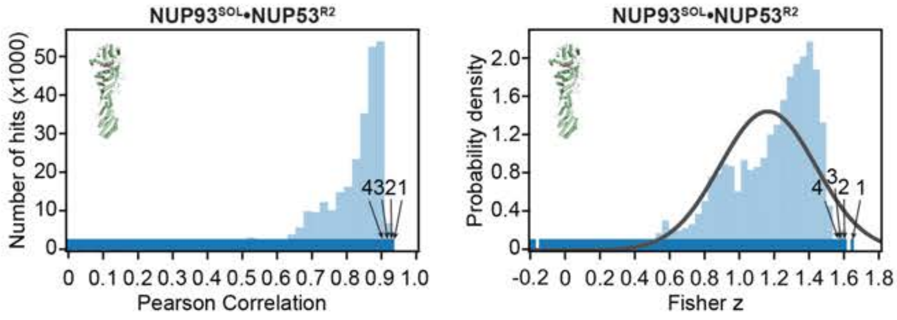

**Subtracted inner ring cryo-ET map fitting statistics**

| solution rank | Pearson correlation | Fisher z | p-value |
|---|---|---|---|
| 1 | 0.927 | 1.64 | 3.9 e-2 |
| 2 | 0.921 | 1.60 | 5.1 e-2 |
| 3 | 0.919 | 1.58 | 6.0 e-2 |
| 4 | 0.918 | 1.58 | 6.1 e-2 |

**Fig.S65. Docking of the NUP93$^{SOL}$•NUP53$^{R1}$ co-crystal structure into an ~23 Å cryo-ET map of the intact human NPC**. Resolution-matched simulated cryo-EM densities of the 3.4 Å NUP93$^{SOL}$•NUP53$^{R2}$ structure were quantitatively docked into (**A**) an ~23 Å sub-tomogram averaged cryo-ET map of the intact human NPC (EMD-3103) (*33*) and (**B**) a sub-segment of the map comprised solely of the inner ring. The placement of unique solutions in a single spoke is shown (*top left*). Arrows and numbers indicate the accepted solutions and their corresponding rank. Closeup views of the accepted solutions are shown on the right, with the rank indicated on the top left corner of the box. Isosurface representations of the nuclear envelope and the NPC are colored in dark gray and white, respectively. The docked structures are displayed in cartoon representation. Rug plots (blue) and histograms (light blue) of Pearson correlation scores and derived Fisher z scores fit with a normalized Gaussian curve (black) from a global search with 1 million random initial placements are shown (bottom left and middle, respectively). A tabular summary of the solutions fitting statistics, along with one-tailed p-values calculated from the Fisher z score distribution, is shown (*bottom right*). Top scoring solutions were identified for a single copy of NUP93$^{SOL}$•NUP53$^{R2}$ at the distal position in each one of the outer rings. No cryo-ET density was observed for a proximal copy of NUP93$^{SOL}$•NUP53$^{R2}$ in the cytoplasmic outer ring. Top scoring solutions for all four copies of NUP93$^{SOL}$•NUP53$^{R2}$ in the inner ring were identified in the inner ring search region.

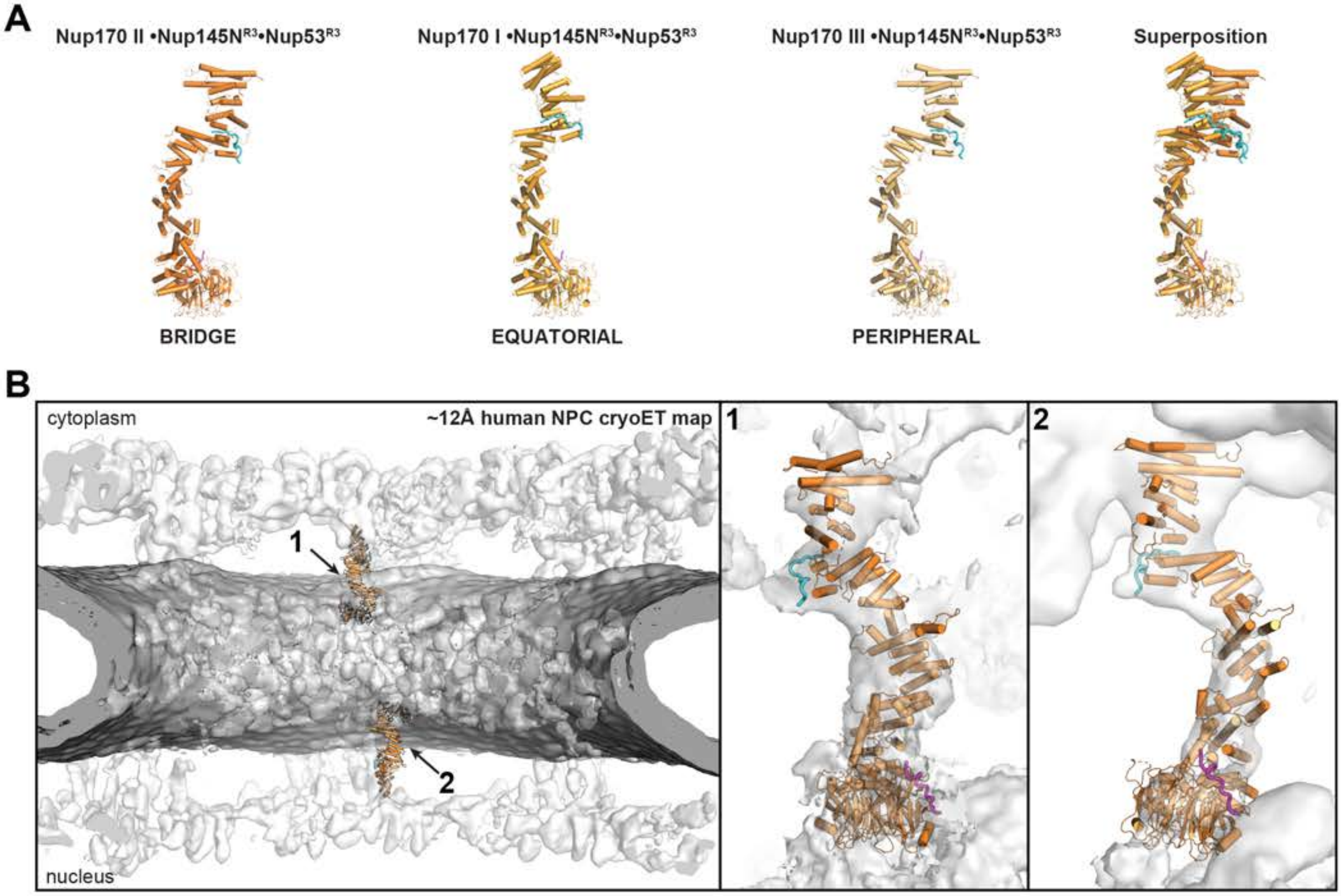

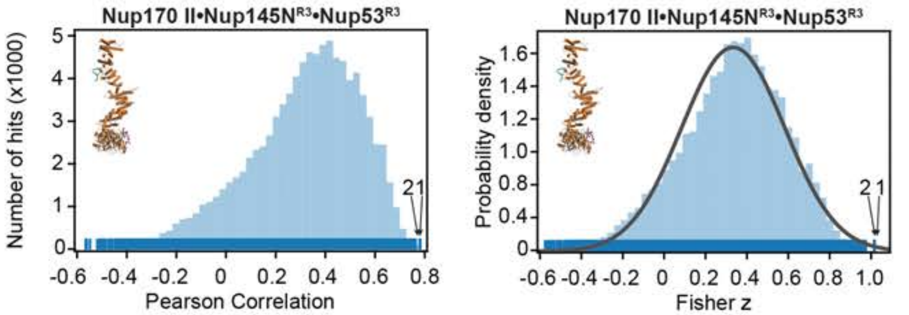

**Subtracted cryo-ET map fitting statistics**

| solution rank | Pearson correlation | Fisher z | p-value |
|---|---|---|---|
| 1 | 0.772 | 1.03 | 3.1 e-3 |
| 2 | 0.754 | 0.98 | 5.2 e-3 |

**Fig.S66. Docking of the bridge composite crystal Nup170•Nup145N $^{R3}$•Nup53$^{R3}$ structure into the ~12 Å cryo-ET map of the intact human NPC**. Resolution-matched simulated cryo-EM densities of the composite crystal Nup170•Nup145N$^{R3}$•Nup53$^{R3}$ conformation II structure (as identified by (*30*) were quantitatively docked into (**A**) the ~12 Å sub-tomogram averaged cryo-ET map of the intact human NPC (EMD-XXXX) (*41*) from which cryo-ET density corresponding to hereto docked nups was subtracted (fig. S54) and (**B**) a sub-segment of the map comprised solely of the inner ring from which cryo-ET density corresponding to hereto docked nups was subtracted (fig. S54). The placement of unique solutions in a single spoke is shown (*top left*). Arrows and numbers indicate the accepted solutions and their corresponding rank. Closeup views of the accepted solutions are shown on the right, with the rank indicated on the top left corner of the box. Isosurface representations of the nuclear envelope and the NPC are colored in dark gray and white, respectively. The docked structures are displayed in cartoon representation. Rug plots (blue) and histograms (light blue) of Pearson correlation scores and derived Fisher z scores fit with a normalized Gaussian curve (black) from a global search with 1 million random initial placements are shown (*bottom left* and *middle*). A tabular summary of the solutions fitting statistics, along with one-tailed p-values calculated from the Fisher z score distribution, is shown (*bottom right*).

**A**

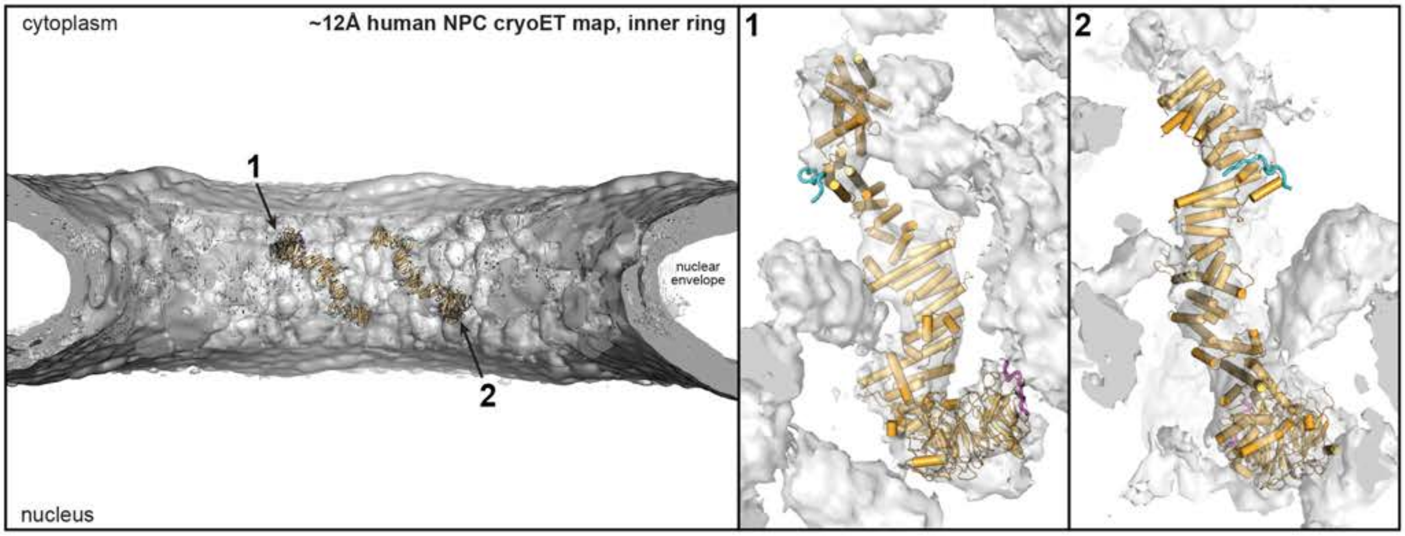

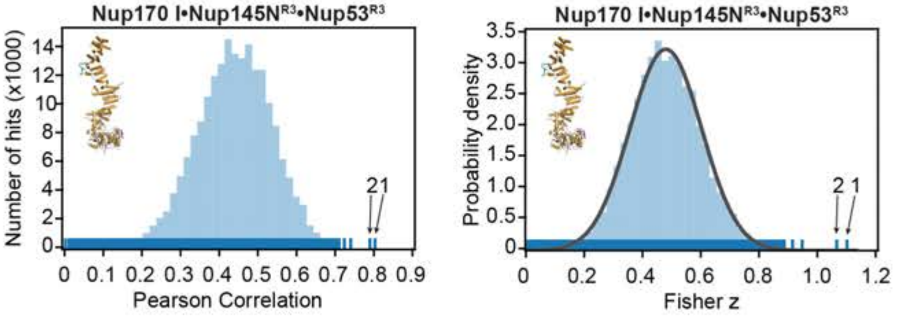

**Subtracted inner ring cryo-ET map fitting statistics**

| solution rank | Pearson correlation | Fisher z | p-value |
|---|---|---|---|
| 1 | 0.799 | 1.09 | 2.2 e-7 |
| 2 | 0.786 | 1.06 | 8.7 e-7 |

**B**

**Manual placement of Nup170 III•Nup145N$^{R3}$•Nup53$^{R3}$**

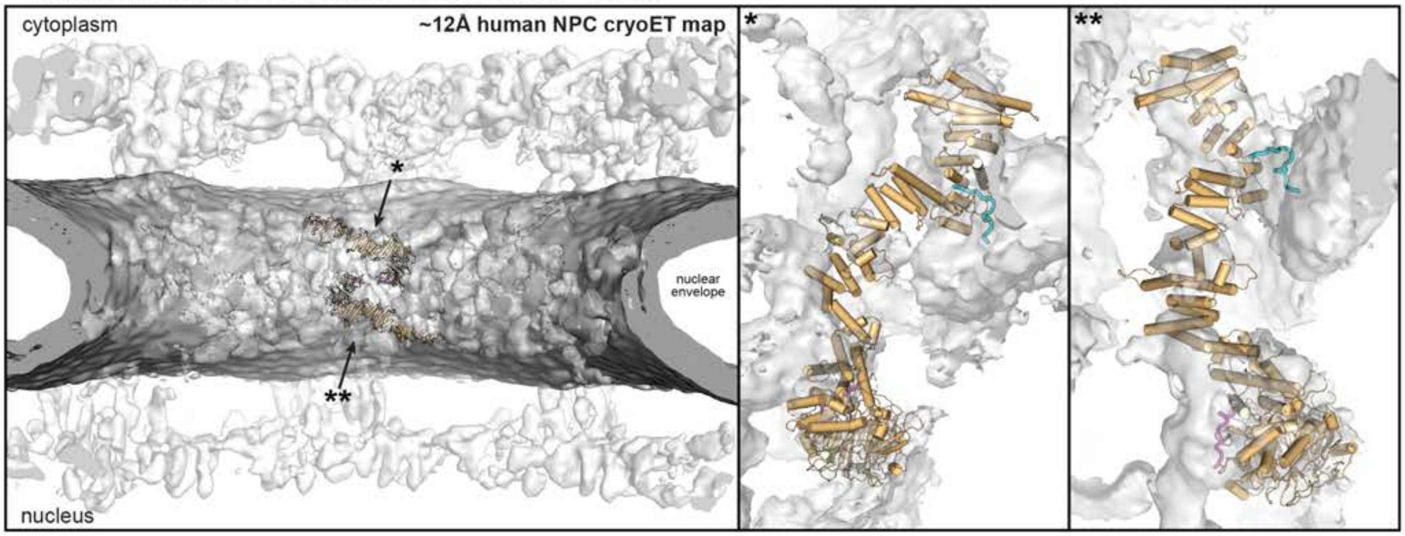

**Fig.S67. Docking of the inner ring composite crystal Nup170•Nup145N $^{R3}$•Nup53$^{R3}$ structures into the ~12 Å cryo-ET map of the intact human NPC**. (**A**) Resolution-matched simulated cryo-EM densities of the composite crystal Nup170•Nup145N$^{R3}$•Nup53$^{R3}$ conformation I structure (as identified by (*30*)) were quantitatively docked into a sub-segment of the ~12 Å sub-tomogram averaged cryo-ET map of the intact human NPC (EMD-XXXX) (*41*) comprised solely of the inner ring from which cryo-ET density corresponding to hereto docked nups was subtracted (fig. S54). The placement of unique solutions in a single spoke is shown (top left). Arrows and numbers indicate the accepted solutions and their corresponding rank. Closeup views of the accepted solutions are shown on the right, with the rank indicated on the top left corner of the box. Isosurface representations of the nuclear envelope and the NPC are colored in dark gray and white, respectively. The docked structures are displayed in cartoon representation. Rug plots (blue) and histograms (light blue) of Pearson correlation scores and derived Fisher z scores fit with a normalized Gaussian curve (black) from a global search with 1 million random initial placements are shown (*bottom left* and *middle*). A tabular summary of the solutions fitting statistics, along with one-tailed p-values calculated from the Fisher z score distribution, is shown (bottom right). The top-scoring solutions identified the two equatorial copies of Nup170•Nup145N$^{R3}$•Nup53$^{R3}$ in the inner ring. (**B**) Attempts of quantitative docking of all available Nup170•Nup145N$^{R3}$•Nup53$^{R3}$ conformations in the peripheral inner ring positions was unsuccessful. The composite crystal Nup170•Nup145N$^{R3}$•Nup53$^{R3}$ conformation III structure (as identified by (*30*) was placed manually at the inner ring cytoplasmic peripheral (*) and nuclear peripheral (**) positions and the fit was locally optimized.

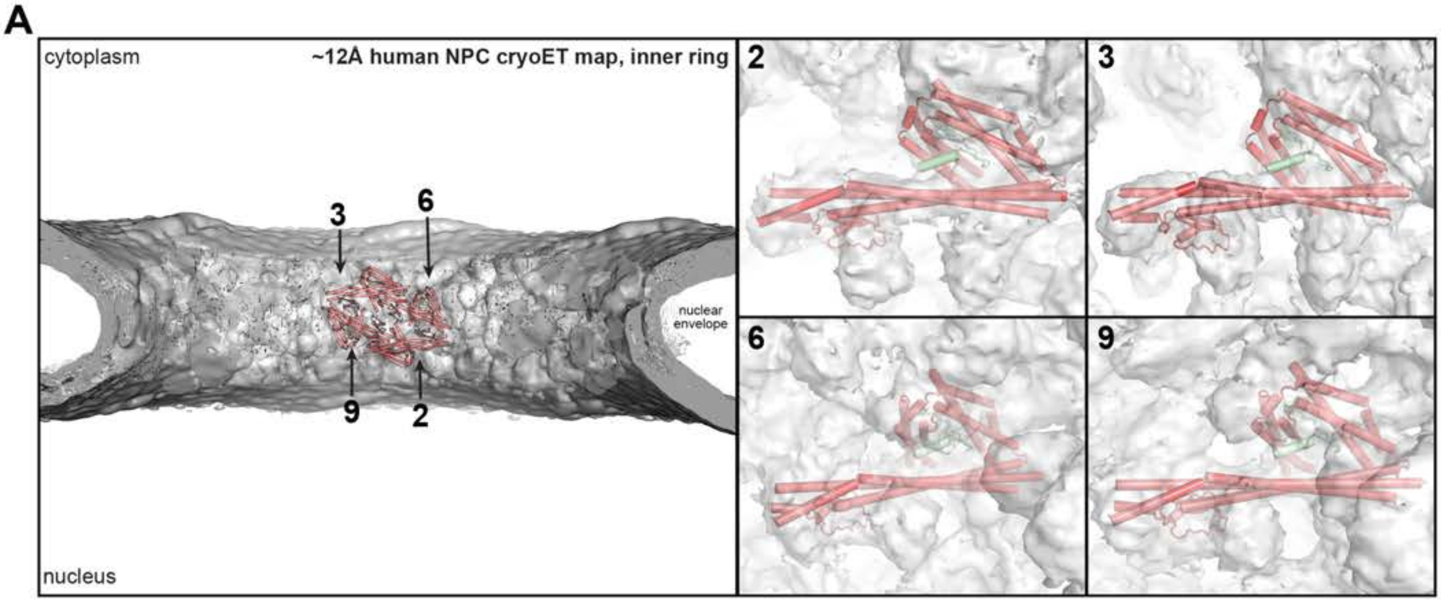

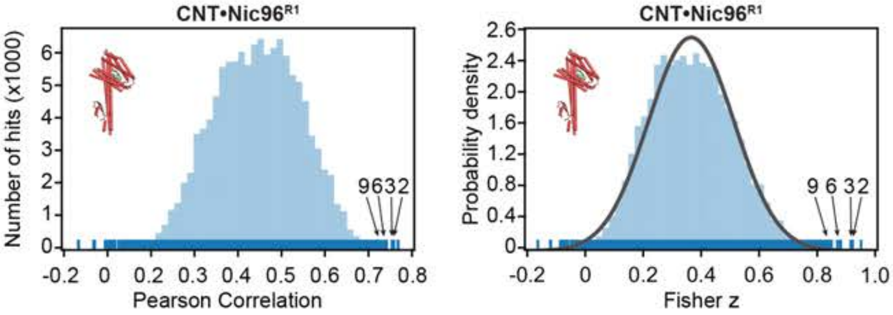

**Subtracted inner ring cryo-ET map fitting statistics**

| solution rank | Pearson correlation | Fisher z | p-value |
|---|---|---|---|
| 2 | 0.724 | 0.916 | 8.3 e-5 |
| 3 | 0.721 | 0.911 | 9.6 e-5 |
| 6 | 0.699 | 0.865 | 3.2 e-4 |
| 9 | 0.684 | 0.836 | 6.4 e-4 |

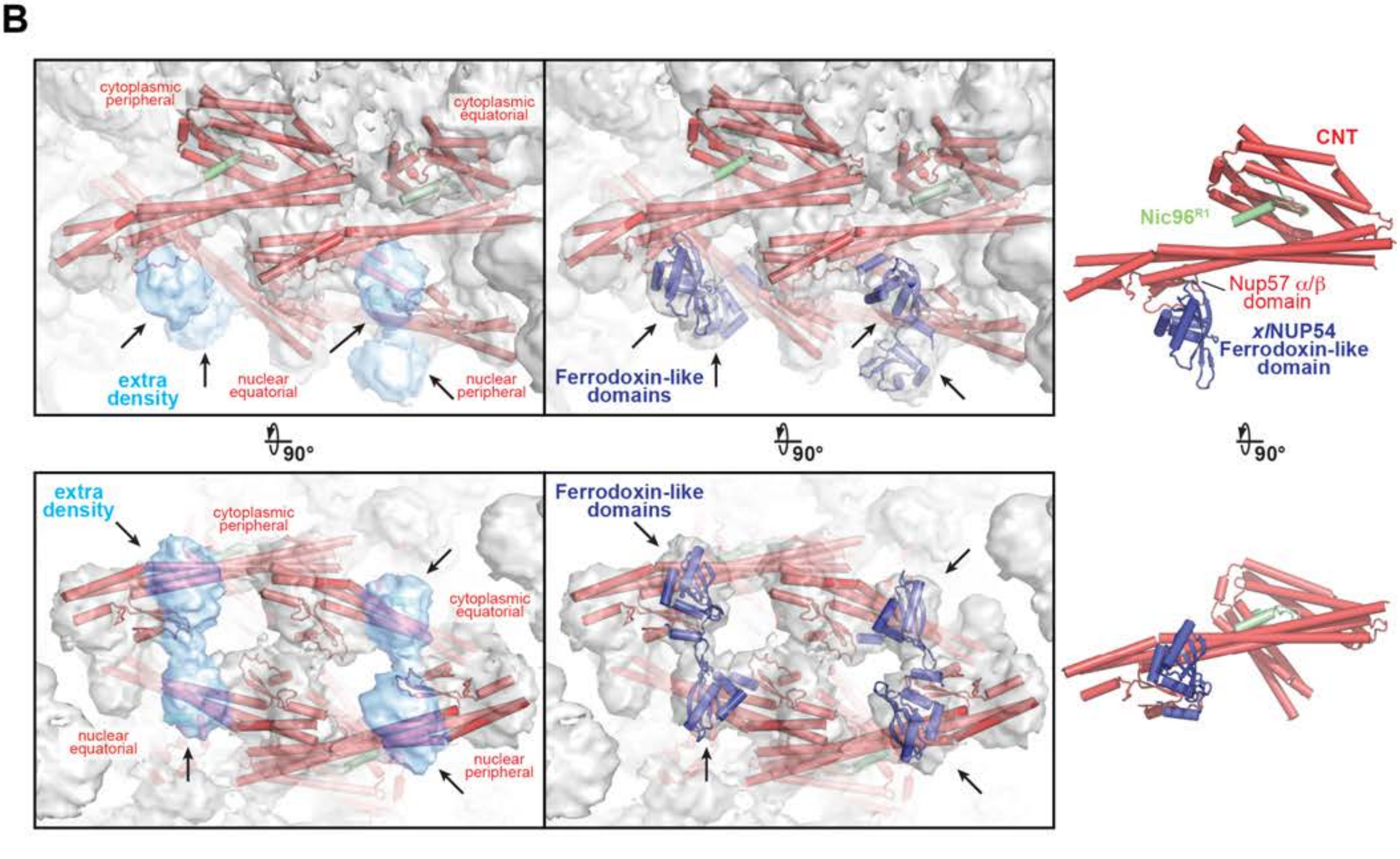

**Fig.S68. Docking of the CNT•Nic96$^{R1}$ co-crystal structure into the ~12 Å cryo-ET map of the intact human NPC**. (**A**) Resolution-matched simulated cryo-EM densities of the CNT•Nic96$^{R1}$ co-crystal structure (PDB ID 5CWS) (*25*) were quantitatively docked into a sub-segment of the ~12 Å sub-tomogram averaged cryo-ET map of the intact human NPC (EMD-XXXX) (*41*) comprised solely of the inner ring from which cryo-ET density corresponding to hereto docked nups was subtracted (fig. S54). The placement of unique solutions in a single spoke is shown (top left). Arrows and numbers indicate the accepted solutions and their corresponding rank. Closeup views of the accepted solutions are shown on the right, with the rank indicated on the top left corner of the box. Isosurface representations of the nuclear envelope and the NPC are colored in dark gray and white, respectively. The docked structures are displayed in cartoon representation. Rug plots (blue) and histograms (light blue) of Pearson correlation scores and derived Fisher z scores fit with a normalized Gaussian curve (black) from a global search with 1 million random initial placements are shown (*bottom left* and *middle*). A tabular summary of the solutions fitting statistics, along with one-tailed p-values calculated from the Fisher z score distribution, is shown (*bottom right*). The four inner ring positions were identified by the 2nd, 3rd, 6th, and 9th highest scoring solutions. (**B**) Two views of cartoon representations of the CNT•Nic96$^{R1}$ co-crystal structures (red and pale green, respectively) docked in the isosurface representation of the ~12 Å cryo-ET map of the human NPC (white). Four globular extra densities (cyan, indicated by arrows) adjacent to the Nup57 $\alpha/\beta$ domain insert (*left*) are shown. NUP54, the metazoan ortholog of Nup57, possesses an additional ferrodoxin-like domain insert adjacent to the $\alpha/\beta$ domain. Ferrodoxin-like domains from *Xenopus laevis xl*NUP54 (PDB ID 5C2U, dark blue) were manually fit into the extra densities and the fit locally optimized (*middle*). A representative pair of CNT•Nic96$^{R1}$ and *xl*NUP54 structures fit in the cryo-ET density is shown to illustrate their relative positions (*right*).

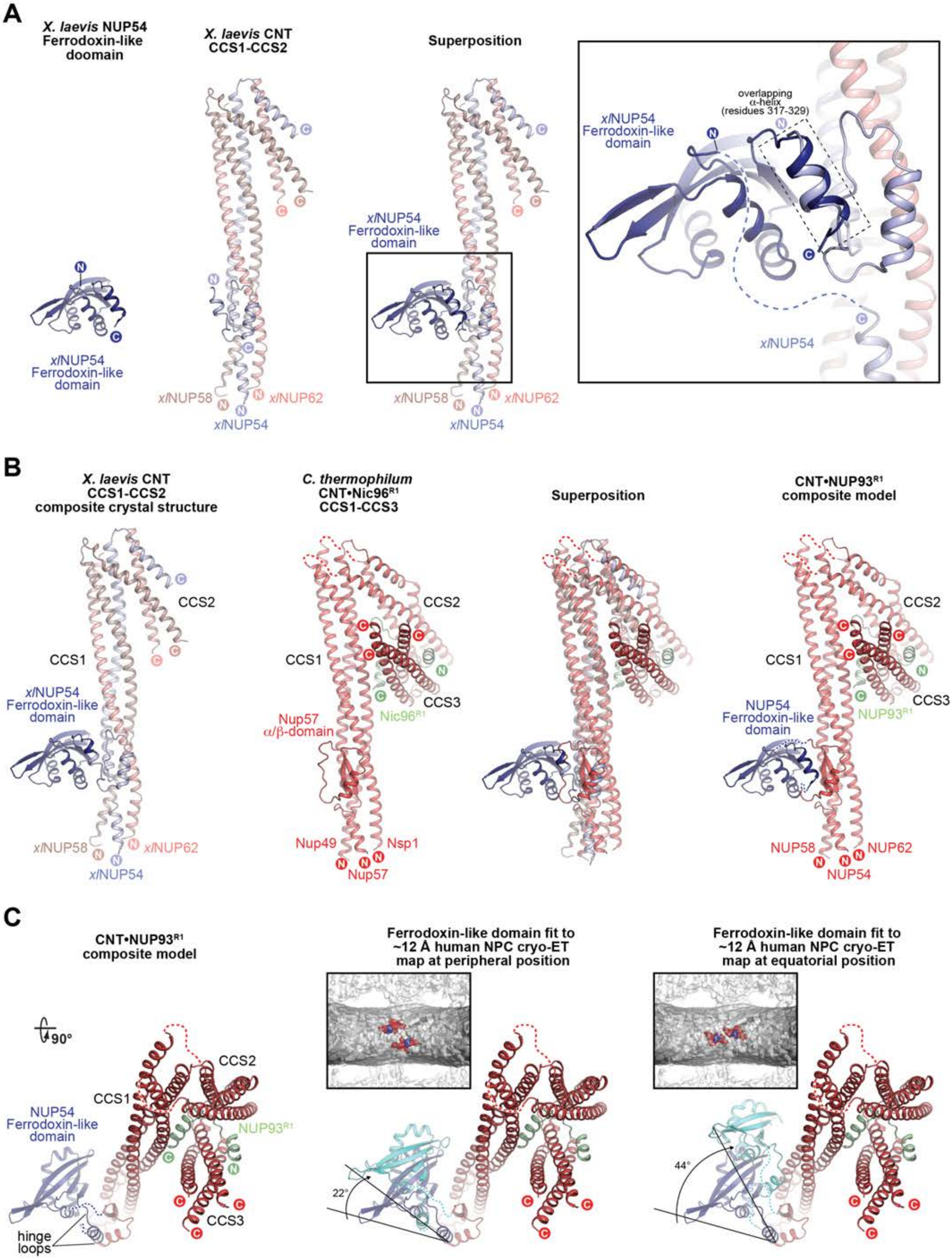

A
X. laevis NUP54 Ferrodoxin-like doomain
X. laevis CNT CCS1-CCS2
Superposition
xlNUP54 Ferrodoxin-like domain
overlapping α-helix (residues 317-329)
xlNUP54
xlNUP58
xlNUP62
B
X. laevis CNT CCS1-CCS2 composite crystal structure
C. thermophilum CNT•Nic96R1 CCS1-CCS3
Superposition
CNT•NUP93R1 composite model
CCS1
CCS2
CCS3
Nup57 α/β-domain
Nic96R1
Nup49
Nup57
Nsp1
NUP54 Ferrodoxin-like domain
NUP93R1
NUP58
NUP54
NUP62
C
CNT•NUP93R1 composite model
Ferrodoxin-like domain fit to ~12 Å human NPC cryo-ET map at peripheral position
Ferrodoxin-like domain fit to ~12 Å human NPC cryo-ET map at equatorial position
90°
hinge loops
22°
44°

**Fig.S69. Composite model of a CNT•NUP93$^{R1}$ inclusive of the NUP54 Ferrodoxin-like domain.** All structures are shown in cartoon representation. (**A**) The *X. laevis* crystal structures *xl*NUP54 (PDB ID 5C2U, dark blue) (*63*) and *xl*CNT (PDB ID 5C2U) (*63*), comprising the first two coiled coil segments CCS1 and CCS2 of *xl*NUP54 (light blue), *xl*NUP58 (brick), and *xl*NUP62 (salmon), were structurally aligned using overlapping residues 317-329 (inset box) to generate a *X. laevis* composite crystal structure. (**B**) The *X. laevis* composite crystal structure was superposed with the *C. thermophilum* CNT•Nic96$^{R1}$ co-crystal structure to generate a composite CNT•NUP93$^{R1}$ model which includes a Ferrodoxin-like domain and all coiled coil segments CCS1-CCS3. (**C**) A top view of the composite CNT•NUP93$^{R1}$ model structurally aligned with the docked CNT•Nic96$^{R1}$ (PDB ID 5CWS) (*25*) co-crystal structure reveals that the ferrodoxin-like domains fit-optimized in the 12 Å cryo-ET map density (cyan) adopt varying degrees of displacement at the peripheral and equatorial positions, which is likely allowed by flexible hinge loops that connect the ferrodoxin-like domain to the rest of the NUP54 chain.

**A**

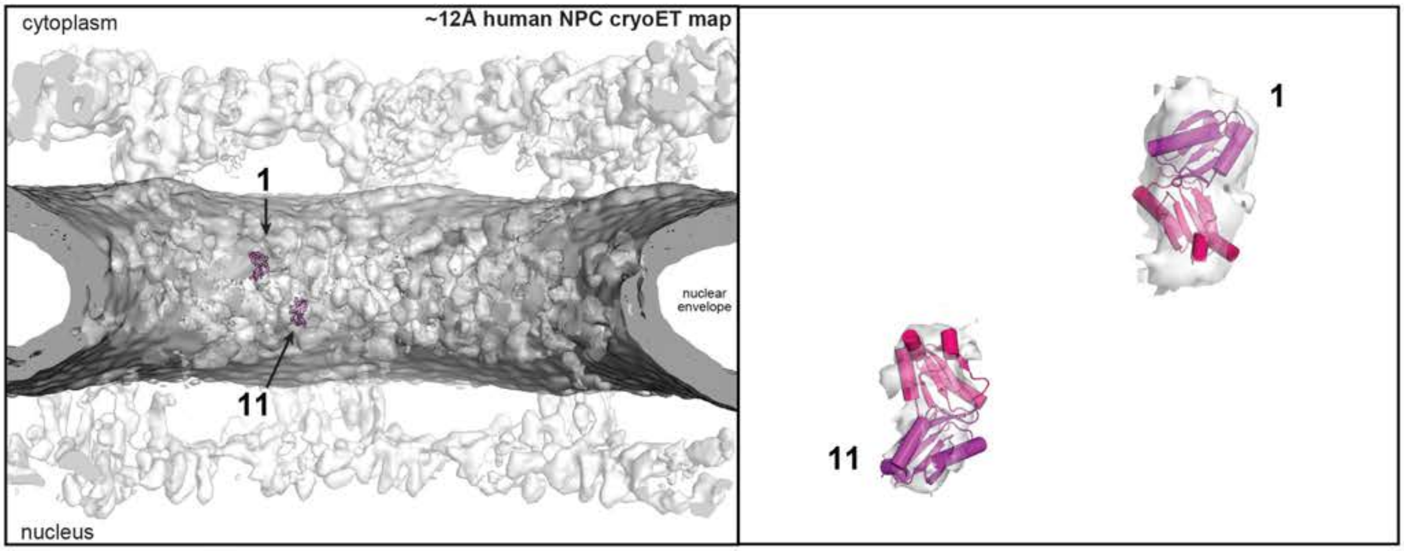

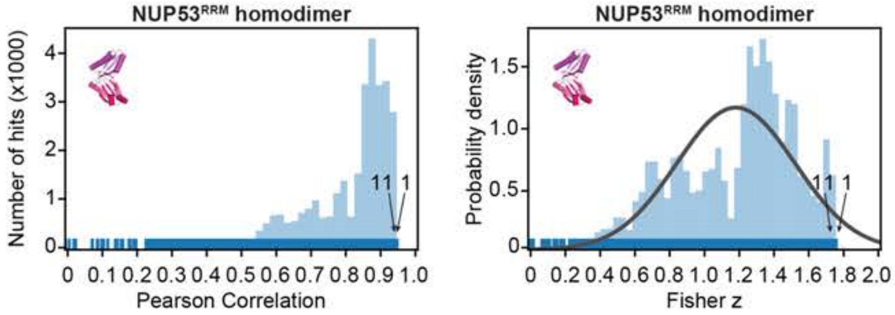

**Subtracted cryo-ET map fitting statistics**

| solution rank | Pearson correlation | Fisher z | p-value |
|---|---|---|---|
| 1 | 0.941 | 1.75 | 4.6 e-2 |
| 11 | 0.936 | 1.70 | 6.0 e-2 |

**B**

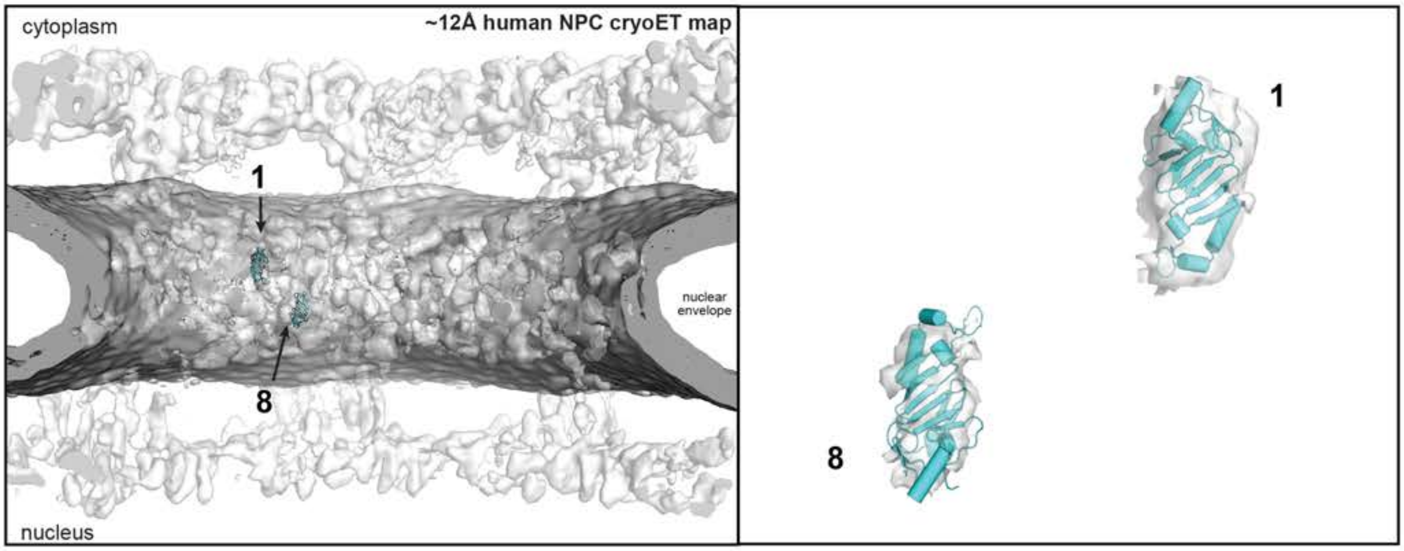

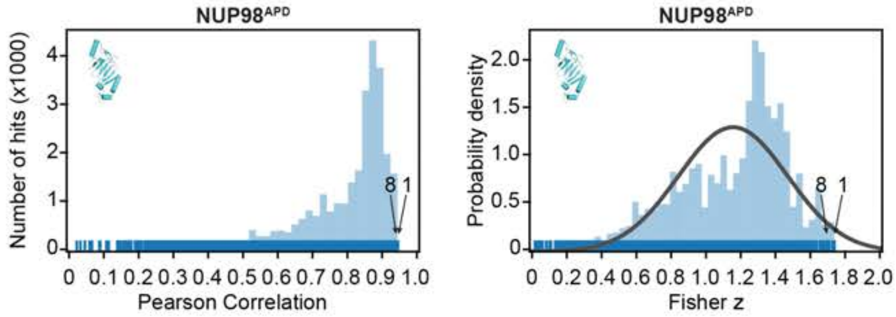

**Subtracted cryo-ET map fitting statistics**

| solution rank | Pearson correlation | Fisher z | p-value |
|---|---|---|---|
| 1 | 0.939 | 1.73 | 3.2 e-2 |
| 8 | 0.936 | 1.71 | 5.8 e-2 |

**Fig.S70. Docking of the NUP53$^{RRM}$ and NUP98$^{APD}$ crystal structures into the ~12 Å cryo-ET map of the intact human NPC**. Resolution-matched simulated cryo-EM densities of the (**A**) NUP53$^{RRM}$ homodimer (PDB ID 4LIR) and (**B**) NUP98$^{APD}$ (PDB ID 1KO6) (*11*) crystal structures were quantitatively docked into the ~12 Å sub-tomogram averaged cryo-ET map of the intact human NPC (EMD-XXXX) (*41*) from which cryo-ET density corresponding to hereto docked nups was subtracted (fig. S54). The placement of unique solutions in a single spoke is shown (*top left*). Arrows and numbers indicate the accepted solutions and their corresponding rank. Closeup views of the accepted solutions are shown on the right, with the rank indicated on the top left corner of the box. Isosurface representations of the nuclear envelope and the NPC are colored in dark gray and white, respectively. The docked structures are displayed in cartoon representation. Rug plots (blue) and histograms (light blue) of Pearson correlation scores and derived Fisher z scores fit with a normalized Gaussian curve (black) from a global search with 1 million random initial placements are shown (*bottom left* and *middle*). A tabular summary of the solutions fitting statistics, along with one-tailed p-values calculated from the Fisher z score distribution, is shown (bottom right). The 2nd-10th highest scoring solutions for the NUP53$^{RRM}$ are placed at the same position as the 1st highest scoring solution, in slightly different orientations. The 2nd-7th highest scoring solutions for the NUP98$^{APD}$ are placed at the same position as the 1st highest scoring solution, in slightly different orientations.

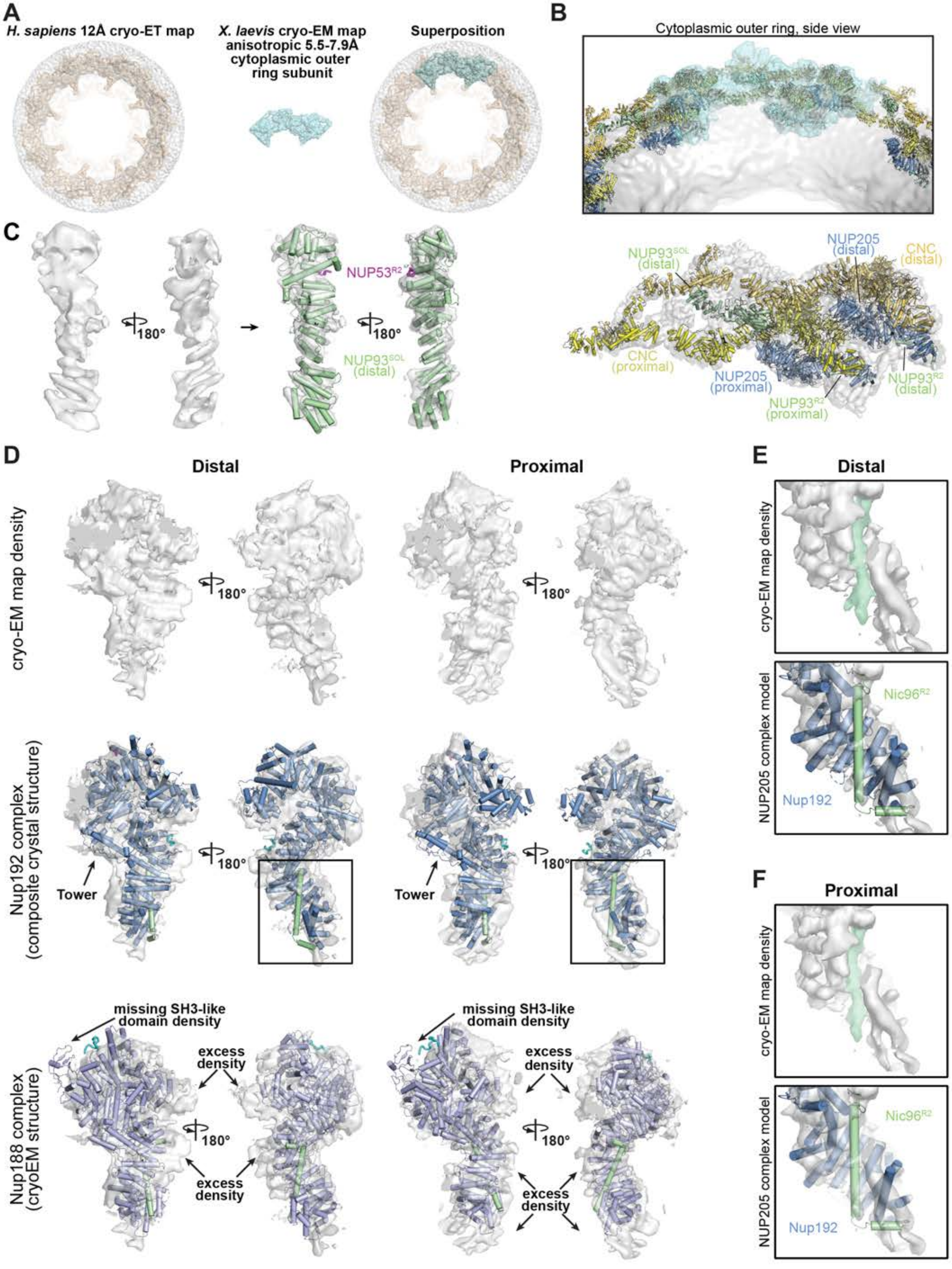

Supp Fig. S71

Hoelz et al., 2021

**Fig.S71. Docking of the *X. leavis* NPC cytoplasmic outer ring**. (**A**) An anisotropic ~8 Å single particle cryo-EM composite map of the *X. laevis* cytoplasmic face of the NPC (light cyan; EMD-0909) (*64*) was superposed to the ~12 Å cryo-ET map of the intact human NPC (wheat; EMD-XXXX) (*41*). (**B**) The superposition of the two maps placed the docked cytoplasmic outer ring nups into the *X. laevis* cytoplasmic face cryo-EM map. The fit of the structures was locally optimized. (**C**) Two views of isosurface representations of carved density and the rigid-body fit NUP93$^{SOL}$•NUP53$^{R2}$ crystal structure shown in cartoon representation (pale green and magenta), revealing correspondence between secondary structure and cryo-EM map features. (**D**) Two views of isosurface representations of proximal and distal question mark-shaped densities carved out from the *X. laevis* cytoplasmic face cryo-EM map and fit with the composite crystal Nup192•Nic96$^{R2}$•Nup145N$^{R1}$•Nup53$^{R1}$ structure. Tentative placement of the Nup188•Nic96$^{R2}$•Nup145N$^{R2}$ single particle cryo-EM structure into the same densities indicates poor fit and confirms the assignment of NUP205 to both proximal and distal question mark-shaped densities of the cytoplasmic outer ring. (**E**, **F**) Closeup on regions indicated by inset boxes in (D). Isosurface of a long tubular cryo-EM density (pale green) running near-perpendicular to surrounding tubular cryo-EM density (white) is shown (*top*). Superposition of cryo-EM density (white) with the Nup192 (blue) and Nic96$^{R2}$ (pale green), shown as cartoons, suggests that the long tubular density engulfed corresponds to the long NUP93$^{R2}$ $\alpha$-helix binding along NUP205 Tail domain (*bottom*).

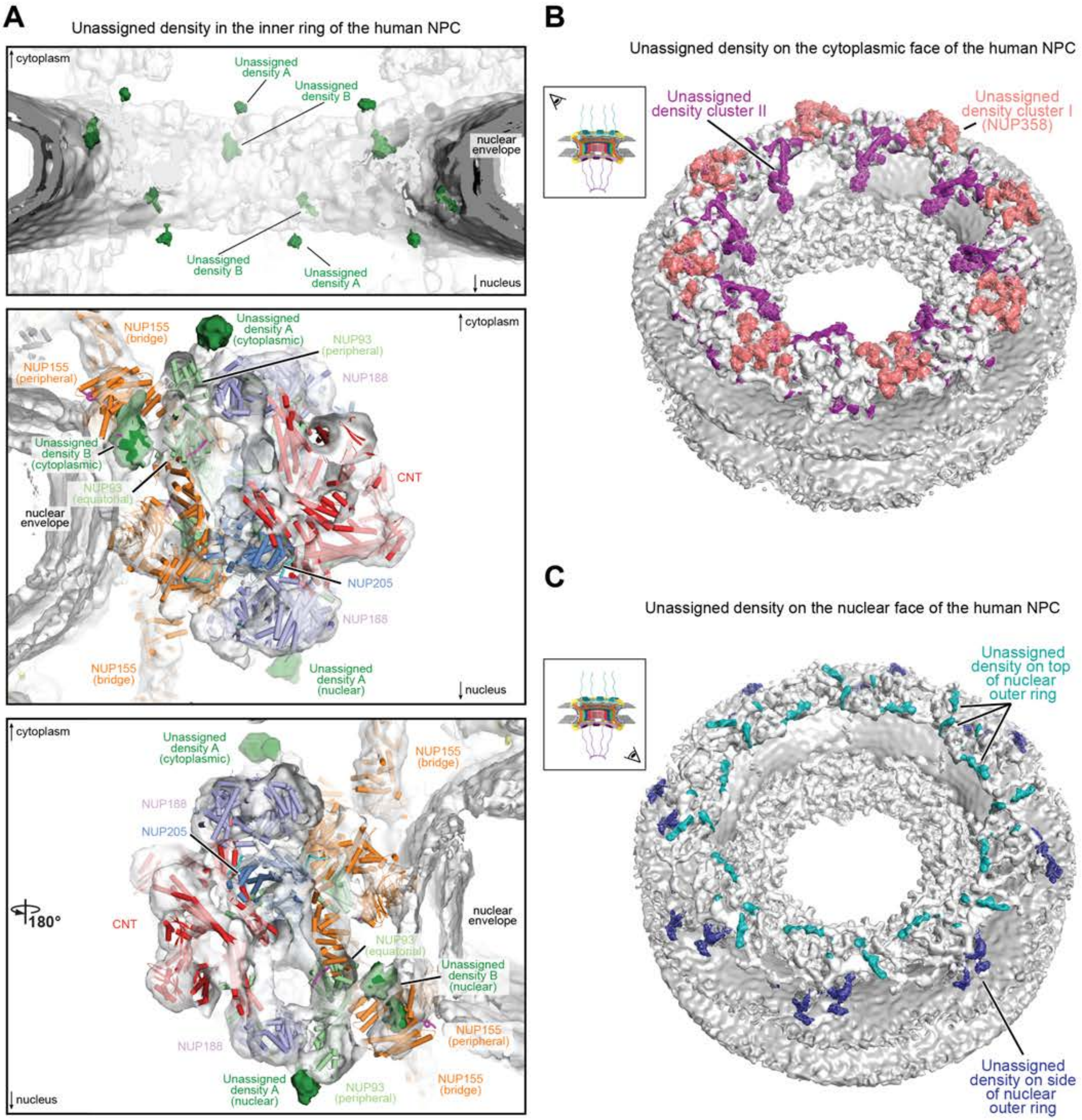

A
Unassigned density in the inner ring of the human NPC
cytoplasm
Unassigned density A
Unassigned density B
nuclear envelope
Unassigned density B
Unassigned density A
nucleus
Unassigned density A (cytoplasmic)
NUP155 (bridge)
NUP155 (peripheral)
NUP93 (peripheral)
NUP188
Unassigned density B (cytoplasmic)
NUP93 (equatorial)
CNT
nuclear envelope
NUP205
NUP188
NUP155 (bridge)
Unassigned density A (nuclear)
180°
NUP155 (peripheral)
Unassigned density B (nuclear)
NUP93 (peripheral)
B
Unassigned density on the cytoplasmic face of the human NPC
Unassigned density cluster II
Unassigned density cluster I (NUP358)
C
Unassigned density on the nuclear face of the human NPC
Unassigned density on top of nuclear outer ring
Unassigned density on side of nuclear outer ring

Supp Fig. 72

**Fig.S72. Unassigned density in an ~12 Å cryo-ET map of the intact human NPC.** Isosurface representation of the ~12 Å sub-tomogram averaged cryo-ET map of the intact human NPC (EMD-XXXX) (*41*) with assigned cryo-ET density (white) and nuclear envelope (gray) rendered as isosurfaces. (**A**) Cross sectional views from the center of the transport channel (*top*) or of a lateral transect of a single spoke (*middle* and *bottom*) of the inner ring with unassigned density isosurfaces indicated (dark green). Structures that explain the assigned density are shown in cartoon representation (*middle* and *bottom*). (**B**) View of the cytoplasmic face isosurface with two clusters of unassigned density indicated (salmon and magenta). (**C**) View of the nuclear face isosurface with two clusters of unassigned density present on top (cyan) and sides (dark blue) of the nuclear outer ring. Schematics indicate the orientation of the viewer with respect to the NPC.

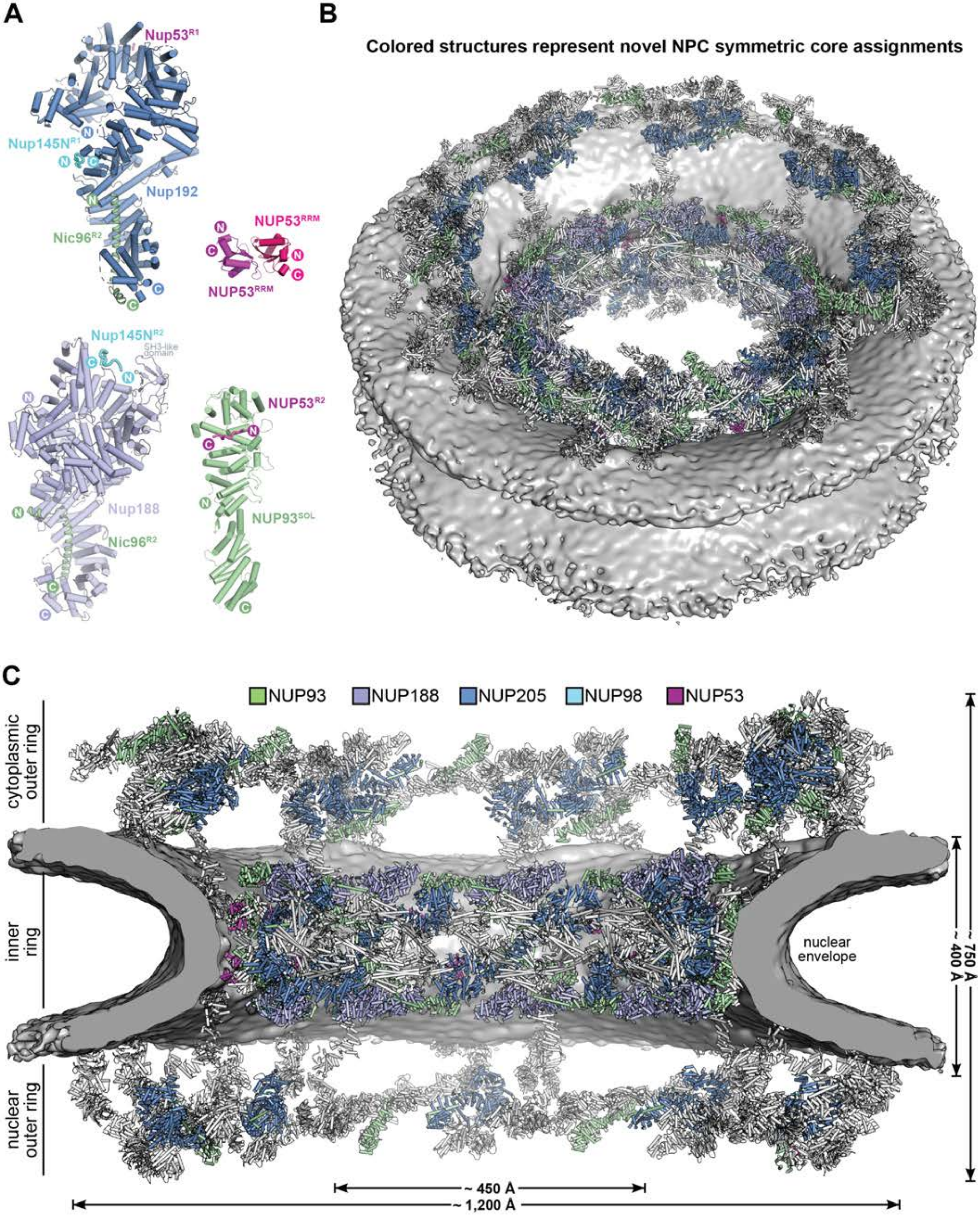

A
Nup53R1
Nup145NR1
Nup192
Nic96R2
NUP53RRM
NUP53RRM
Nup145NR2
SH3-like domain
Nup188
Nic96R2
NUP53R2
NUP93SOL
B
Colored structures represent novel NPC symmetric core assignments
C
NUP93
NUP188
NUP205
NUP98
NUP53
cytoplasmic outer ring
inner ring
nuclear outer ring
nuclear envelope
~ 750 Å
~ 400 Å
~ 450 Å
~ 1,200 Å

**Fig.S73. Summary of novel additions to the symmetric core composite structure.** (**A**) Novel high resolution structures of nups and nup complexes docked into intact human NPC symmetric core, shown as cartoons: $Nup192{\bullet}Nic96^{R2}{\bullet}Nup145N^{R1}{\bullet}Nup53^{R1}$ (blue, pale green, cyan, and magenta), $Nup188{\bullet}Nic96^{R2}{\bullet}Nup145N^{R2}$ (light purple, pale green, and cyan), $NUP93^{SOL}{\bullet}NUP53^{R2}$ (pale green and magenta), and the $NUP53^{RRM}$ homodimer (magenta). (**B**) View from above the cytoplasmic face and (**C**) a cross-sectional view from the central transport channel of the symmetric core of the intact human NPC with novel structures shown in cartoon representation and colored according to (A). The remainder of the symmetric core nups are colored white. Isosurface representations of the nuclear envelope are colored in gray.

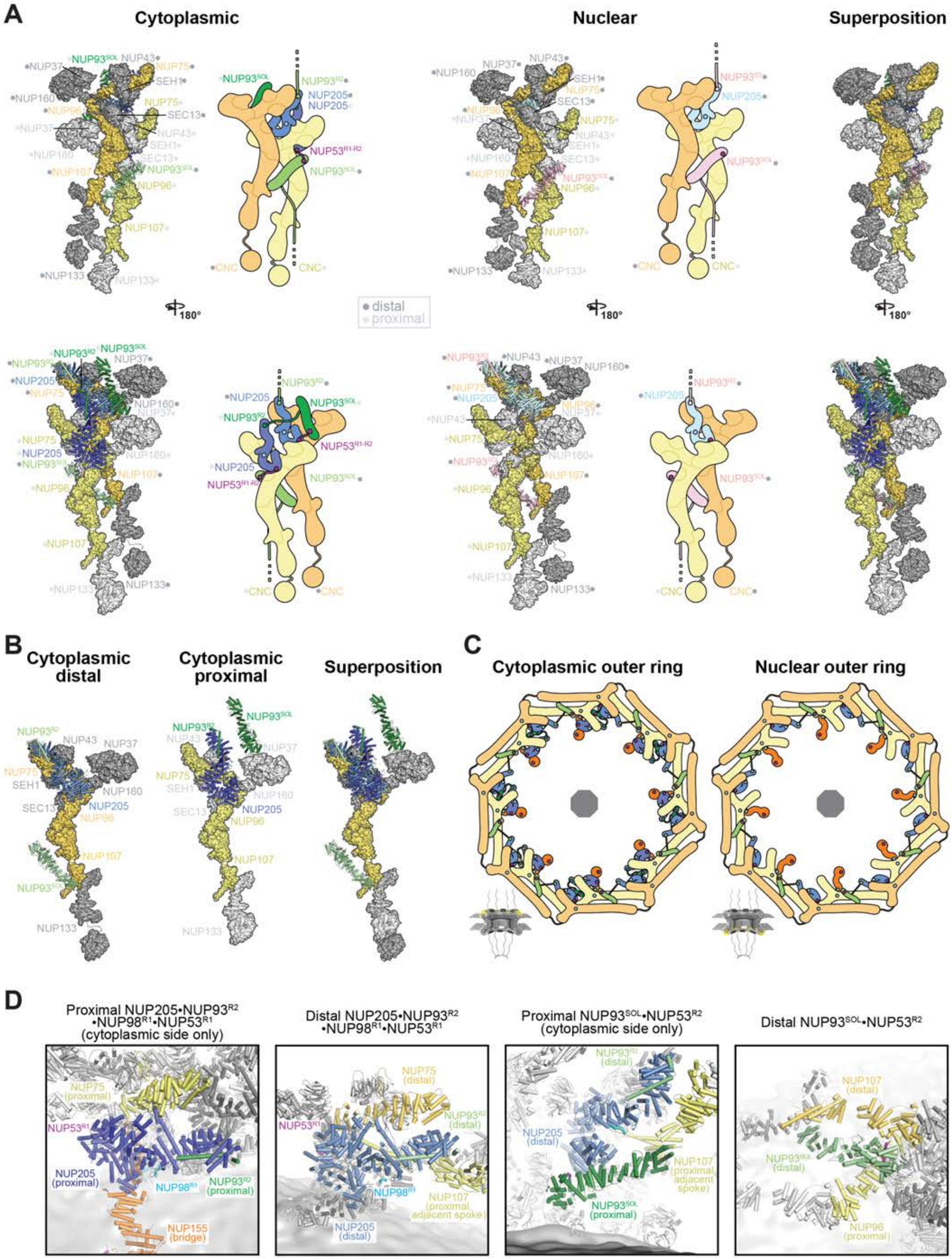

Supp Fig. S74

Hoelz et al., 2021

**Fig.S74. Architecture of the outer rings of the NPC symmetric core.** (**A**) Top and bottom views of the cytoplasmic and nuclear outer ring spoke protomers, and their structural superposition. NUP205 and NUP93 are shown in cartoon representation. CNCs are shown in surface representation and colored in gray, with the nups that interface with NUP205 and NUP93 highlighted in color. Circles by labels indicate if the nup is in a distal (dark gray) or proximal (light gray) position. The cytoplasmic spoke includes two copies of NUP205 (blue for distal and dark blue for proximal) and two copies of NUP93 (pale green for distal and dark green for proximal). The distal NUP205 (blue for cytoplasmic, light cyan for nuclear) and the distal NUP93 (pale green for cytoplasmic, pink for nuclear) are found at equivalent positions in the cytoplasmic and nuclear outer rings and form interfaces with both proximal (pale yellow) and distal (yellow) CNC. Schematics indicate linker connections: between distal $NUP93^{SOL}$ (pale green for cytoplasmic, pink for nuclear) and distal NUP205 of adjacent spokes (blue for cytoplasmic, light cyan for nuclear), between proximal NUP205 (dark blue) and proximal $NUP93^{SOL}$ (dark green) of the same cytoplasmic spoke, between NUP53 binding sites (magenta) on proximal NUP205 and distal $NUP93^{SOL}$ and distal NUP205 and proximal $NUP93^{SOL}$. (**B**) Structural superposition of the distal and proximal nups of the cytoplasmic outer ring. NUP205 (blue for distal and dark blue for proximal) and NUP93 (pale green for distal and dark green for proximal) are shown in cartoon representation. CNCs are shown in surface representation and colored in gray, with the nups that interface with NUP205 and NUP93 highlighted (pale yellow for proximal and yellow for distal). (**C**) Schematic representations of top views of the cytoplasmic (*left*) and nuclear (*right*) outer rings. Gray octagons indicate the eight-fold symmetry. The CNC double rings (yellow for proximal and ochre for distal) assume a head-to-tail circular arrangement with eight spokes. A staple between spokes is provided by the distal NUP205 (blue) interacting with a distal NUP93 (pale green) from an adjacent spoke. Bridge NUP155 (orange) provide an interface between the outer rings and the inner ring (not shown). The cytoplasmic outer ring encompasses a proximal NUP205 (dark blue) and NUP93 (dark green) that interact within the same spoke. Linker nup binding sites are indicated with circles: NUP98 (cyan), NUP53 (magenta), and NUP93 (green). (**D**) Closeup views of the interfaces accommodating outer ring NUP205 and NUP93. Nups are shown as cartoons and the nuclear envelope as isosurface (dark gray). The nups interfacing with indicated NUP205 or NUP93 molecule are shown in color and the remainder of nups are indicated in gray.

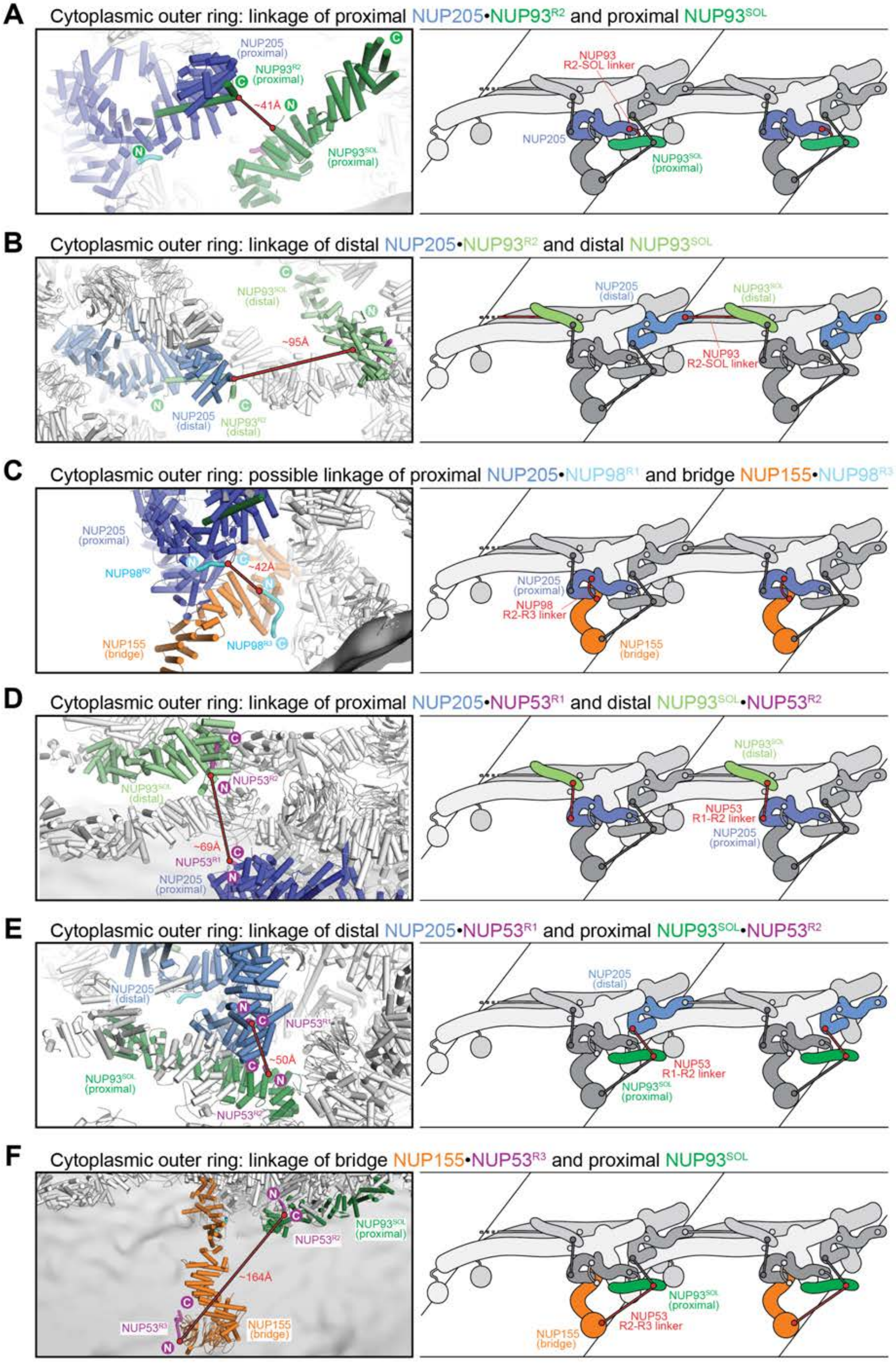
A
Cytoplasmic outer ring: linkage of proximal NUP205•NUP93$^{R2}$ and proximal NUP93$^{SOL}$
NUP205 (proximal)
NUP93$^{R2}$ (proximal)
~41Å
NUP93$^{SOL}$ (proximal)
NUP93 R2-SOL linker
NUP205
B
Cytoplasmic outer ring: linkage of distal NUP205•NUP93$^{R2}$ and distal NUP93$^{SOL}$
NUP93$^{SOL}$ (distal)
~95Å
NUP205 (distal)
NUP93$^{R2}$ (distal)
NUP93 R2-SOL linker
C
Cytoplasmic outer ring: possible linkage of proximal NUP205•NUP98$^{R1}$ and bridge NUP155•NUP98$^{R3}$
NUP205 (proximal)
NUP98$^{R2}$
~42Å
NUP155 (bridge)
NUP98$^{R3}$
NUP98 R2-R3 linker
D
Cytoplasmic outer ring: linkage of proximal NUP205•NUP53$^{R1}$ and distal NUP93$^{SOL}$•NUP53$^{R2}$
NUP53$^{R2}$
NUP93$^{SOL}$ (distal)
~69Å
NUP53$^{R1}$
NUP205 (proximal)
NUP53 R1-R2 linker
E
Cytoplasmic outer ring: linkage of distal NUP205•NUP53$^{R1}$ and proximal NUP93$^{SOL}$•NUP53$^{R2}$
NUP205 (distal)
NUP53$^{R1}$
~50Å
NUP93$^{SOL}$ (proximal)
NUP53$^{R2}$
NUP53 R1-R2 linker
F
Cytoplasmic outer ring: linkage of bridge NUP155•NUP53$^{R3}$ and proximal NUP93$^{SOL}$
NUP93$^{SOL}$ (proximal)
NUP53$^{R2}$
~164Å
NUP53$^{R3}$
NUP155 (bridge)
NUP53 R2-R3 linker

**G** Nuclear outer ring: linkage of distal NUP205•Nic96$^{R2}$ and distal NUP93$^{SOL}$

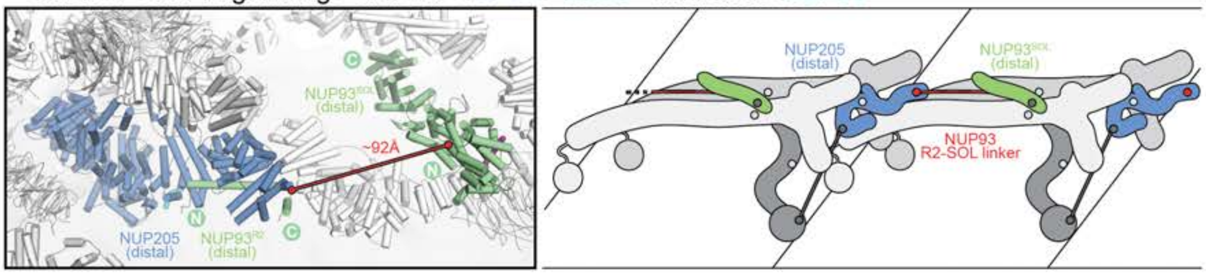

**H** Nuclear outer ring: linkage of distal NUP205•NUP53$^{R1}$ and bridge NUP155•NUP53$^{R3}$

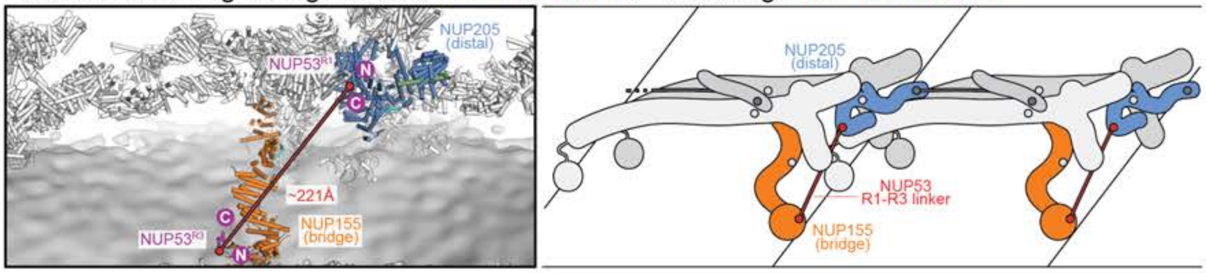

**I** Nuclear outer ring: linkage of distal NUP93$^{SOL}$•NUP53$^{R2}$ and bridge NUP155•NUP53$^{R3}$

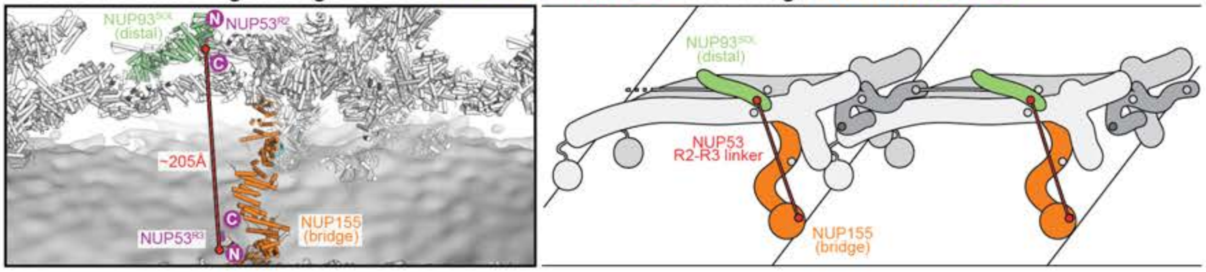

**Fig.S75. Shortest-distance analysis of linker network in the outer rings of the NPC.** Spatial distances in the composite structure of human NPC were measured between two copies of linker nup segments that are adjacent in their primary sequence. (**A-I**) In the box on the left, linker nup segments and the scaffold surface that they are bound to are shown in cartoon representation and highlighted in color. The remainder of nups are colored white. The nuclear envelope is rendered as dark gray isosurface. The straight-line linkage between the two closest copies of linker nup segments adjacent in their primary sequence is indicated by a solid red line. The measured distance is reported in red. On the right, a schematic representation of the scaffold nup binding sites (highlighted in color) and the connecting linker nup segment (red), illustrates the topology of the linkage within the context of two outer ring NPC spokes, with the remainder of the nups shown in gray.

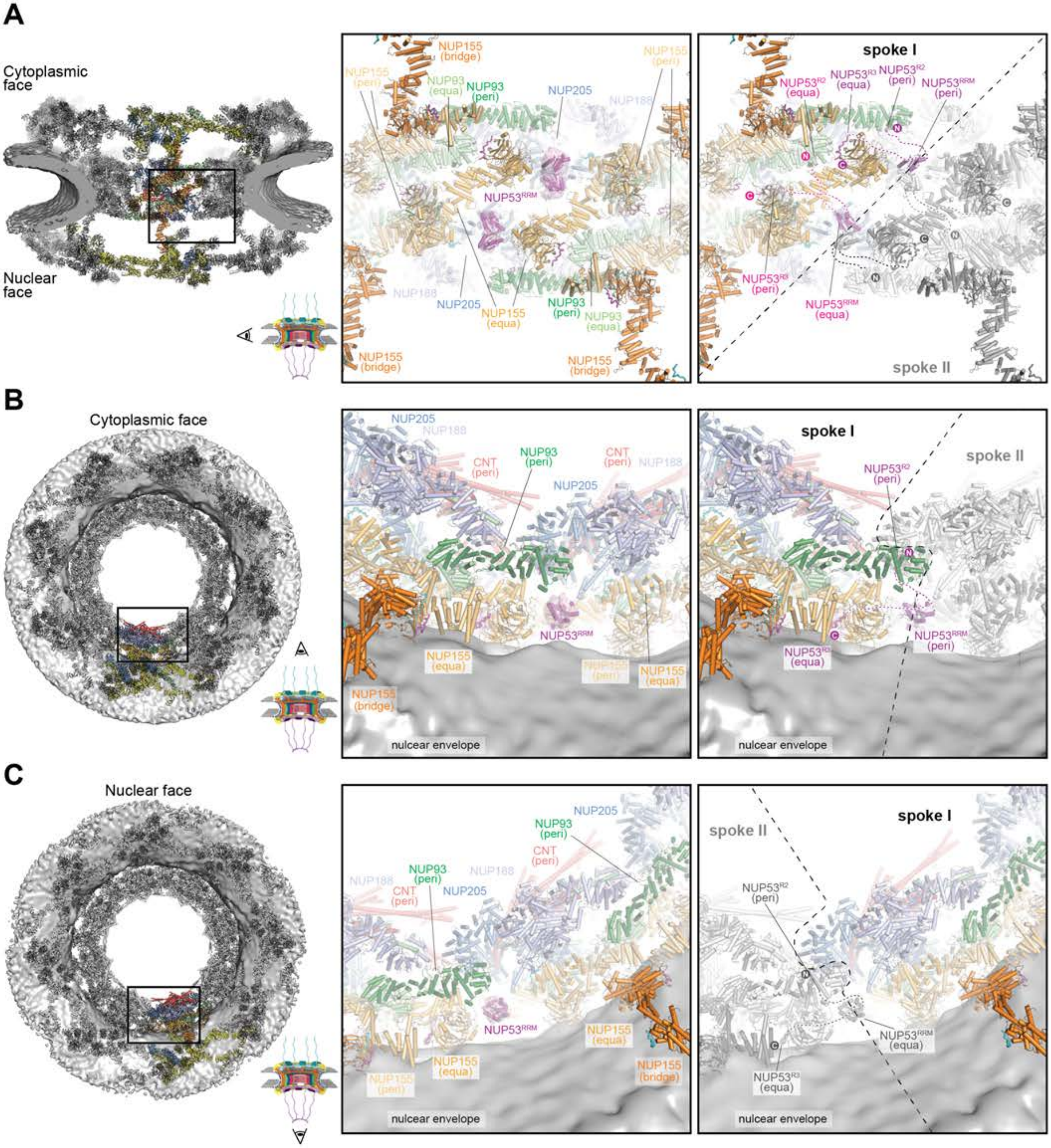

A
Cytoplasmic face
Nuclear face
NUP155 (bridge)
NUP155 (peri)
NUP93 (equa)
NUP93 (peri)
NUP205
NUP188
NUP53RRM
NUP155 (equa)
spoke I
spoke II
NUP53R2 (equa)
NUP53R3 (equa)
NUP53R2 (peri)
NUP53RRM (peri)
NUP53R3 (peri)
NUP53RRM (equa)
B
Cytoplasmic face
CNT (peri)
nulcear envelope
C
Nuclear face

**Fig.S76. Placement of the NUP53$^{RRM}$ homodimer between inner ring spokes.** (**A**-**C**) Different views of the interface between inner spokes of the human NPC are shown, as indicated by the inset box and schematic representation of the NPC (*left*). The composite NPC structures are shown as cartoons and the nuclear envelope as a dark gray isosurface. In the middle panel, globular cryo-ET densities, discontinuous with the rest of the ~12 Å human cryo-ET map, are shown as pink isosurfaces and fit with the docked NUP53$^{RRM}$ homodimer crystal structure (magenta). On the right, two spokes are delimited by a dashed line and one spoke is shown in gray. Shortest-distance connections with NUP53$^{R2}$ and NUP53$^{R3}$ peptides, bound to NUP93 and NUP155, respectively, are drawn as dashed lines. The NUP53$^{R2}$ and NUP53$^{R3}$ binding regions are N- and C-terminally adjacent to the NUP53$^{RRM}$ domain in the primary sequence. The NUP53$^{RRM}$ homodimer interfaces connect two inner ring spokes.

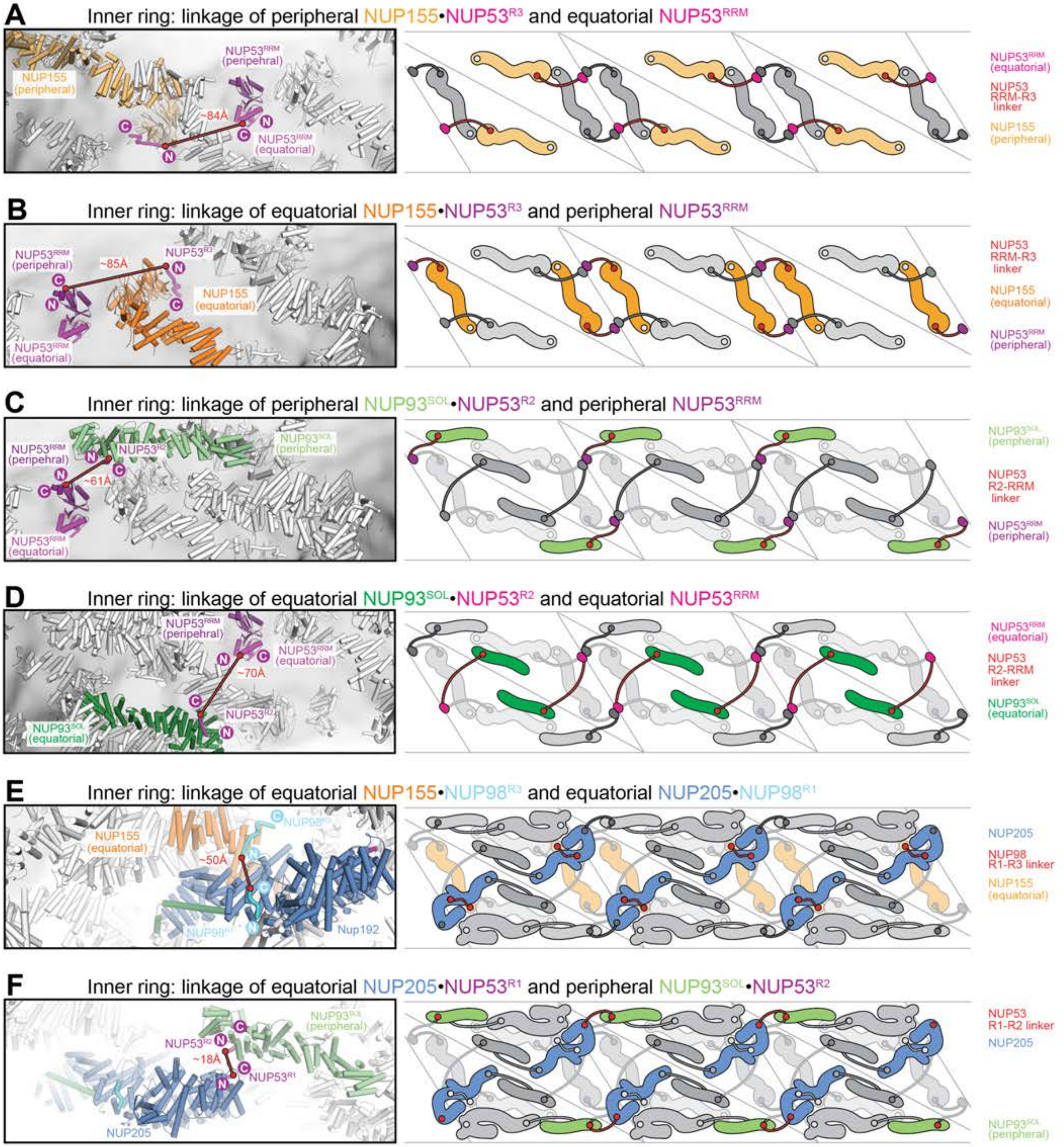

A
Inner ring: linkage of peripheral NUP155•NUP53$^{R3}$ and equatorial NUP53$^{RRM}$
NUP155 (peripheral)
NUP53$^{RRM}$ (peripehral)
~84Å
NUP53$^{RRM}$ (equatorial)
NUP53 RRM-R3 linker
NUP155 (peripheral)
B
Inner ring: linkage of equatorial NUP155•NUP53$^{R3}$ and peripheral NUP53$^{RRM}$
NUP53$^{RRM}$ (peripehral)
~85Å
NUP53$^{R3}$
NUP155 (equatorial)
NUP53$^{RRM}$ (equatorial)
NUP53 RRM-R3 linker
NUP53$^{RRM}$ (peripheral)
C
Inner ring: linkage of peripheral NUP93$^{SOL}$•NUP53$^{R2}$ and peripheral NUP53$^{RRM}$
NUP53$^{RRM}$ (peripehral)
NUP53$^{R2}$
NUP93$^{SOL}$ (peripheral)
~61Å
NUP53$^{RRM}$ (equatorial)
NUP53 R2-RRM linker
D
Inner ring: linkage of equatorial NUP93$^{SOL}$•NUP53$^{R2}$ and equatorial NUP53$^{RRM}$
NUP53$^{RRM}$ (peripehral)
NUP53$^{RRM}$ (equatorial)
~70Å
NUP53$^{R2}$
NUP93$^{SOL}$ (equatorial)
NUP53 R2-RRM linker
E
Inner ring: linkage of equatorial NUP155•NUP98$^{R3}$ and equatorial NUP205•NUP98$^{R1}$
NUP155 (equatorial)
NUP98$^{R3}$
~50Å
NUP98$^{R1}$
Nup192
NUP205
NUP98 R1-R3 linker
F
Inner ring: linkage of equatorial NUP205•NUP53$^{R1}$ and peripheral NUP93$^{SOL}$•NUP53$^{R2}$
NUP93$^{SOL}$ (peripheral)
NUP53$^{R2}$
~18Å
NUP53$^{R1}$
NUP205
NUP53 R1-R2 linker

**G** Inner ring: linkage of peripheral NUP188•NUP93$^{R2}$ and peripheral NUP93$^{SOL}$

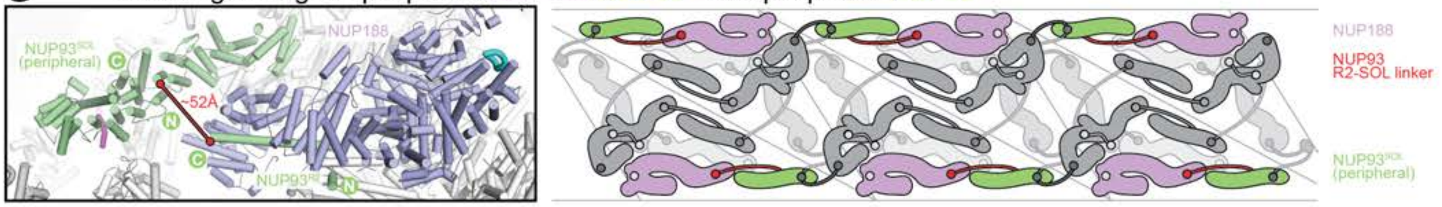

**H** Inner ring: linkage of equatorial NUP205•NUP93$^{R2}$ and equatorial NUP93$^{SOL}$

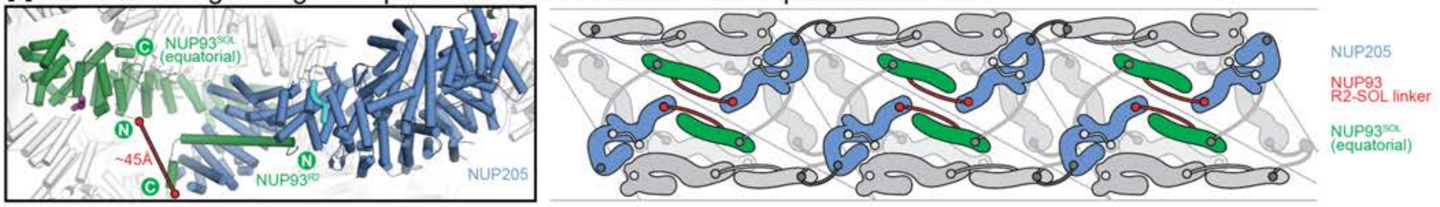

**I** Inner ring: linkage of peripheral NUP188•NUP93$^{R2}$ and peripheral CNT•NUP93$^{R1}$

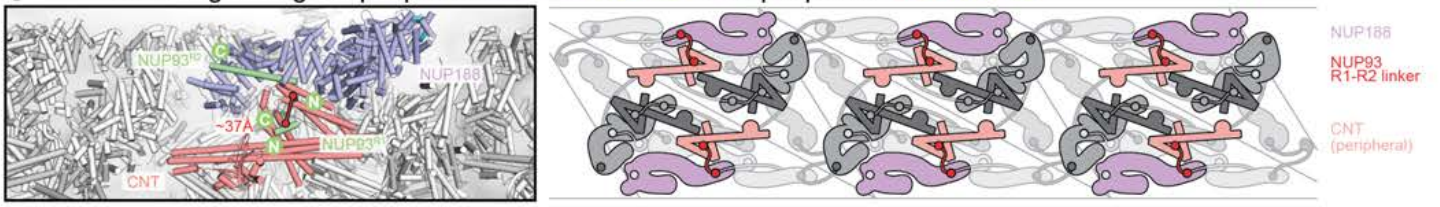

**J** Inner ring: linkage of equatorial NUP205•NUP93$^{R2}$ and equatorial CNT•NUP93$^{R1}$

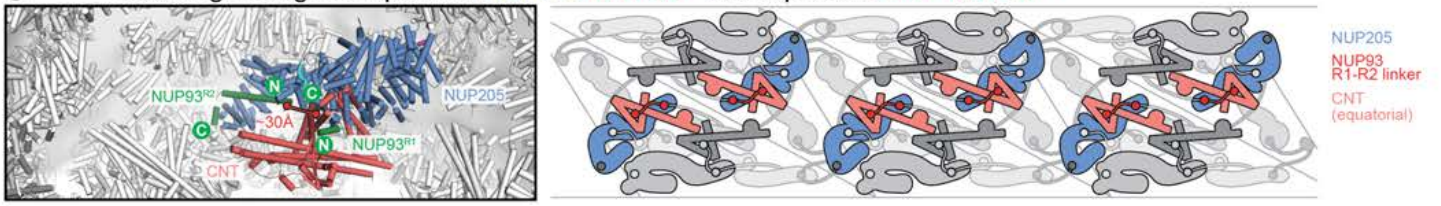

**K**

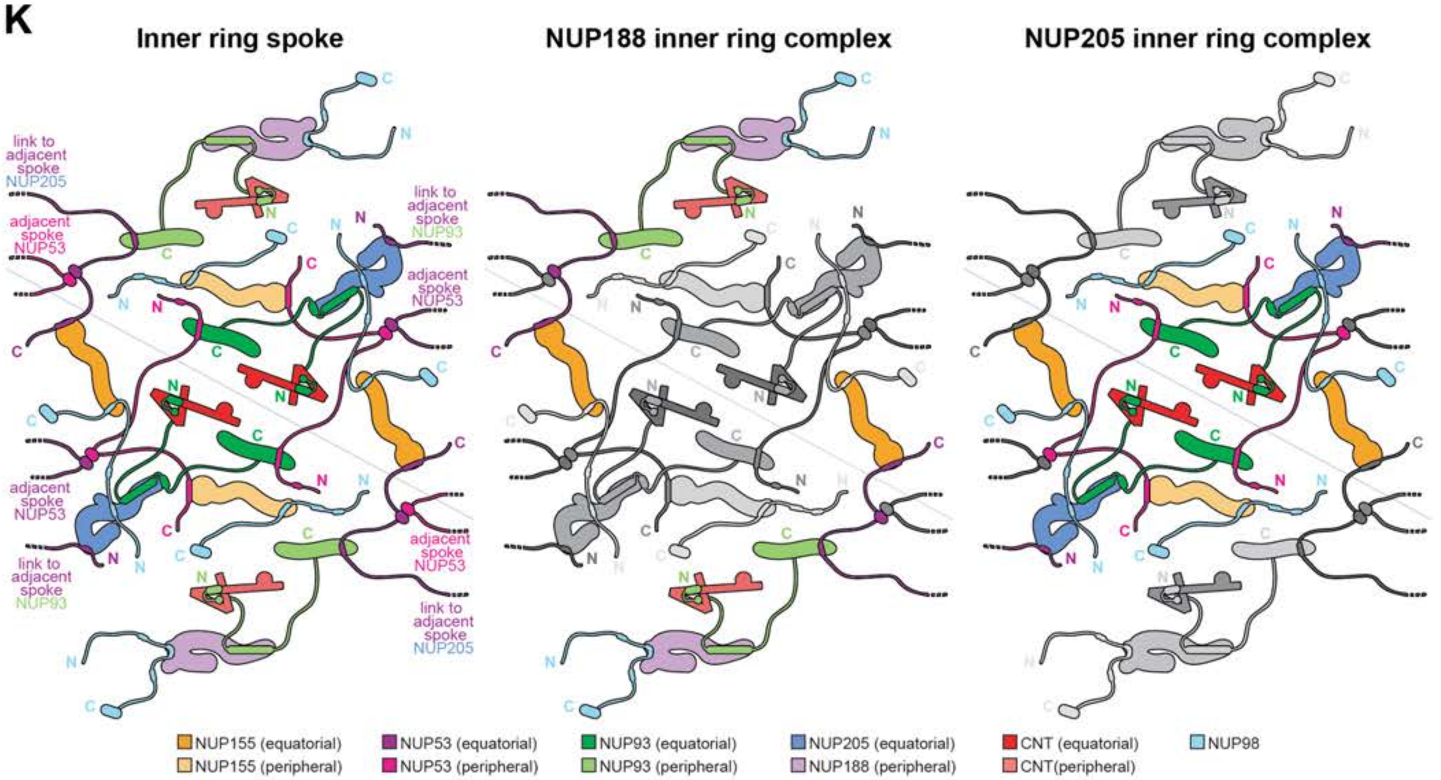

**Fig.S77. Shortest-distance analysis of linker network in the inner ring of the NPC.** Spatial distances in the composite structure of human NPC were measured between two copies of linker nup segments that are adjacent in their primary sequence. (**A-I**) In the box on the left, linker nup segments and the scaffold surface that they are bound to are shown in cartoon representation and highlighted in color. The remainder of nups are colored white. The nuclear envelope is rendered as a dark gray isosurface. The straight-line linkage between the two closest copies of linker nup segments adjacent in their primary sequence is indicated by a solid red line. The measured distance is reported in red. On the right, a schematic representation of the scaffold nup binding sites (highlighted in color) and the connecting linker nup segment (red), illustrates the topology of the linkage within the context of three inner ring NPC spokes (remainder of nups shown in gray).

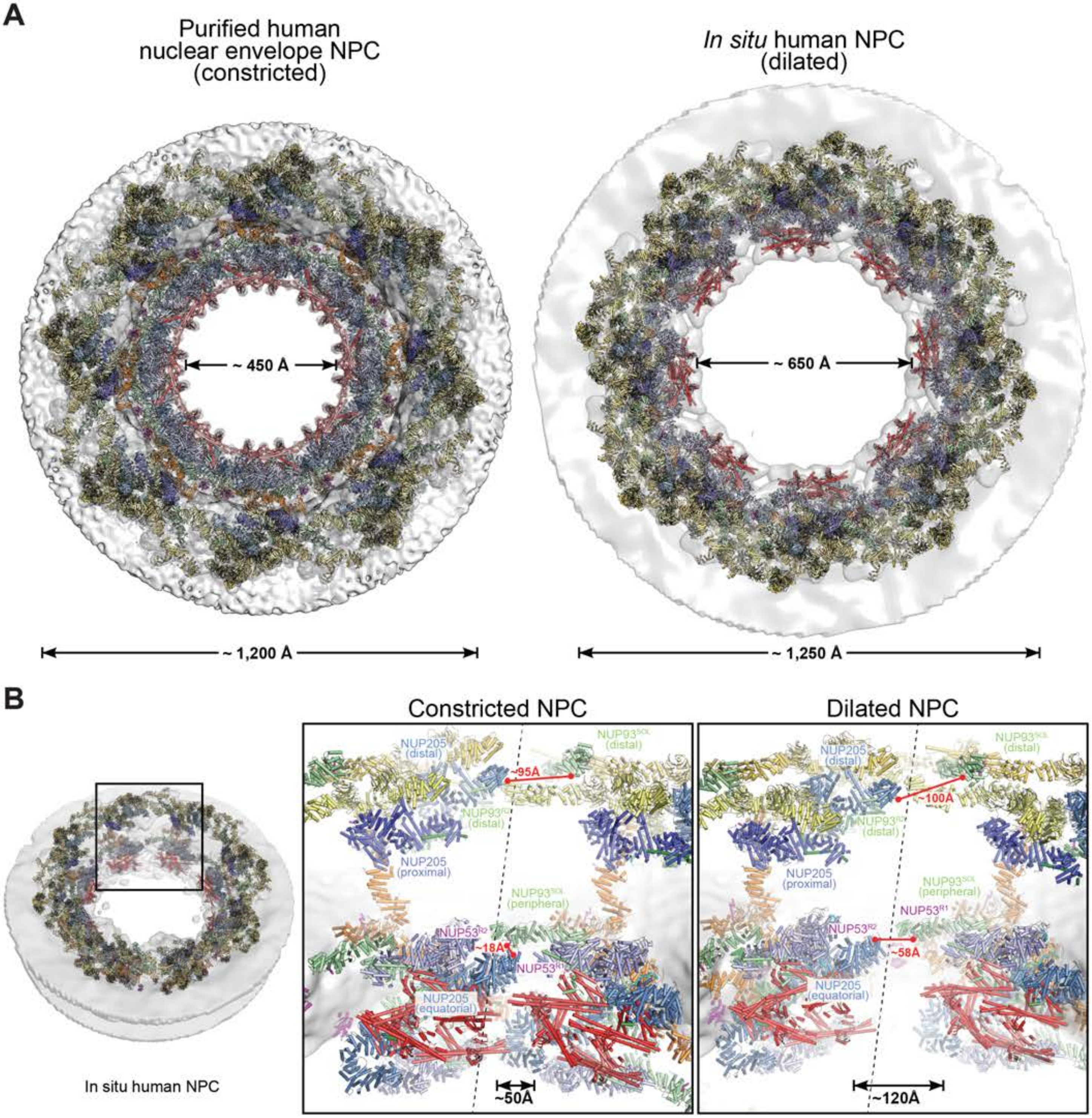

A
Purified human nuclear envelope NPC (constricted)
In situ human NPC (dilated)
~ 450 Å
~ 650 Å
~ 1,200 Å
~ 1,250 Å
B
Constricted NPC
Dilated NPC
In situ human NPC
NUP205 (distal)
NUP93 (distal)
~95Å
~100Å
NUP205 (proximal)
NUP93 (peripheral)
NUP53
~18Å
~58Å
NUP205 (equatorial)
~50Å
~120Å

**Fig.S78. Composite structure of the dilated *in situ* human NPC.** (**A**) Comparison of the constricted human NPC imaged in purified HeLa cell nuclear envelopes and the dilated human NPC imaged *in situ* in SupT1-R5 cells. Outer and inner ring spoke subcomplexes from the composite NPC structure built into the constricted ~12 Å cryo-ET sub-tomogram averaged human NPC map (EMD-XXXX) (*41*) were manually docked and locally fit as rigid units in the dilated *in situ* ~37 Å cryo-ET sub-tomogram averaged human NPC map (EMD-11967) (*40*). The composite NPC structures are shown as cartoons superposed to isosurfaces of the constricted and dilated NPCs (grey). In transitioning between constricted and dilated states, the outer ring diameter of the human NPC increases by ~4%, whereas the diameter of the central transport channel increases by ~31%. (**C**) Closeup views the cytoplasmic outer ring and inner ring at the interface (dashed line) between two spokes in the constricted and dilated human NPCs. Measurements of the distance spanned by the NUP93 linker between NUP205 (blue) and NUP93$^{SOL}$ (pale green) of adjacent spokes in the dilated and constricted NPCs show little dilation between spokes in the outer rings. On the contrary, measurements of the distance spanned by the NUP53 linker between NUP205 (blue) and the peripheral NUP93 (pale green) of adjacent spokes of the inner ring, as well as the distance between two equivalent points in the CNT (red) layer closest to the central transport channel, suggest that the dilation of the NPC results in a gap between inner ring spokes.

**Table S1.** Materials and reagents

| REAGENT or RESOURCE | SOURCE | IDENTIFIER |
|---|---|---|
| ***C. thermophilum* and *H. sapiens* nucleoporin cDNAs** | | |
| Nup192 | (*30*) | N/A |
| Nup188 | (*30*) | N/A |
| Nic96 | (*25*) | N/A |
| Nup145N | (*25*) | N/A |
| Nup53 | (*30*) | N/A |
| NUP98 | Gift, Beatriz Fontoura (*83*) | N/A |
| NUP155 | Gift, Susan Wente (*84*) | N/A |
| Nup42 | (*127*) | N/A |
| NUP53 | Open Biosystems | MHS6278-202758913 |
| NUP93 | Open Biosystems | MHS6278-202786560 |
| NUP58 | Open Biosystems | MHS1011-59697 |
| NUP54 | Open Biosystems | MHS1011-74585 |
| NUP205 | Kazusa Genome Technology | ORK06210 |
| NUP188 | DNASU | HsCD00295457 |
| Cre | Gift, David Baltimore (*122*) | N/A |
| **Molecular biology reagents** | | |
| PfuUltra II Fusion HotStart DNA Polymerase | Agilent Technologies | Cat# 600674 |
| Phusion High-Fidelity DNA Polymerase | New England Biolabs | Cat# M0530L |
| AseI | New England Biolabs | Cat# R0526L |
| BamHI | New England Biolabs | Cat# R0136L |
| EcoRI | New England Biolabs | Cat# R0101L |
| HindIII | New England Biolabs | Cat# R0104L |
| NcoI | New England Biolabs | Cat# R0193L |
| NdeI | New England Biolabs | Cat# R0111L |
| NotI | New England Biolabs | Cat# R0189L |
| SacII | New England Biolabs | Cat# R0157L |
| SalI | New England Biolabs | Cat# R0138L |
| XbaI | New England Biolabs | Cat# R3642L |
| XhoI | New England Biolabs | Cat# R0146L |
| T4 DNA Ligase | New England Biolabs | Cat# M0145L |
| Agarose | Invitrogen | Cat# 16500500 |
| ***S. cerevisiae* culture reagents** | | |
| Peptone | BD | Cat# 211677 |
| Yeast Extract | BD | Cat# 212750 |
| Raffinose | Gold Biotechnology | Cat# 030500 |
| Complete Supplement Mixture minus histidine (CSM-His) | Sunrise Science Products | Cat# 1006-100 |
| Complete Supplement Mixture minus leucine, tryptophan and uracil ( CSM-Leu-Trp-Ura) | Sunrise Science Products | Cat# 1017-100 |
| Complete Supplement Mixture Dropout (CSM-His-Leu-Met-Ura) | Sunrise Science Products | Cat# 1145-010 |
| L-Leucine | Sigma-Aldrich | Cat# L8000-100G |
| L-Tryptophan | Sigma-Aldrich | Cat# T0254-100G |
| Uracil | Sigma-Aldrich | Cat# U1128-25G |
| D-(+)-Galactose | Sigma-Aldrich | Cat# G0625-100G |
| Lithium acetate | Fluka | Cat# 62393 |
| G418 | Gold Biotechnology | Cat# G418-5 |
| Nourseothricin | Gold Biotechnology | Cat# N-500-100 |
| Agar | Fisher BioReagents | Cat# BP1423-500 |
| **Bacterial culture reagents** | | |
| Luria-Bertani (LB) media | Fisher BioReagents | Cat# BP9723-5 |
| Isopropyl-$\beta$-D-thiogalactopyranoside (IPTG) | Gold Biotechnology | Cat# 12481C-1KG |
| Ampicillin | Gold Biotechnology | Cat# A-301-100 |
| Kanamycin | Gold Biotechnology | Cat# K-120-100 |
| Chloramphenicol | Gold Biotechnology | Cat# C-105-100 |
| Seleno-L-methionine (SeMet) | TCL | Cat# S0442 |
| **Protein purification reagents / general chemicals** | | |

| REAGENT or RESOURCE | SOURCE | IDENTIFIER |
|---|---|---|
| TRIS (trometamol, 2-amino-2-(hydroxymethyl)-1,3-propanediol) | Sigma-Aldrich | Cat# 93362 |
| Sodium chloride | Sigma-Aldrich | Cat# 71376-5KG |
| Sodium acetate | Sigma-Aldrich | Cat# S3272-1KG |
| Potassium phosphate monobasic | Sigma-Aldrich | Cat# 60353-1KG |
| Potassium phosphate dibasic, trihydrate | Sigma-Aldrich | Cat# 7088-06 |
| Magnesium acetate | Fluka | Cat# 63049 |
| Ammonium sulfate | MP Bio | Cat# 808211 |
| Imidazole | Sigma-Aldrich | Cat# 56750 |
| 2-Mercaptoethanol (β-ME) | Sigma-Aldrich | Cat# M6250 |
| Dithiothreitol (DTT) | Gold Biotechnology | Cat# DTT100 |
| Glycerol | Sigma-Aldrich | Cat# 29767-100ML |
| Aprotinin from bovine lung | Sigma-Aldrich | Cat# A6279 |
| Phenylmethylsulfonyl fluoride (PMSF) | Gold Biotechnology | Cat# P-470-50 |
| EDTA-free protease inhibitor cocktail | Roche | Cat# A32965 |
| DNase I | Sigma-Aldrich | Cat# D4527 |
| Adenosine 5'-Triphosphate disodium salt (ATP) | Sigma-Aldrich | Cat# A7699 |
| Biotin | Pierce | Cat# 21335 |
| Bicine | Sigma-Aldrich | Cat# 14871 |
| Ethylenediaminetetraacetic acid (EDTA) | Sigma-Aldrich | Cat# 03620-1KG |
| Ubl-specific protease 1(ULP1) | Invitrogen | Cat# 12588018 |
| PreScission protease (Pres) | Sigma-Aldrich | Cat# GE27-0843-01 |
| **SDS-PAGE reagents** | | |
| 40% Acrylamide/Bis solution (37.5:1) | Bio-Rad | Cat# 1610148 |
| Tetramethylethylenediamine (TEMED) | Bio-Rad | Cat# 161-0801 |
| Glycine | Gold Biotechnology | Cat# G-630-5 |
| Sodium dodecyl sulfate | Sigma-Aldrich | Cat# L3771-1KG |
| Ammonium persulfate | Sigma-Aldrich | Cat# 09913-100g |
| Acetic acid | Sigma-Aldrich | Cat# 695092 |
| Brilliant Blue R | Sigma-Aldrich | Cat# B7920-50G |
| **FISH mRNA export assay reagents** | | |
| Poly-L-lysine | Sigma-Aldrich | Cat# P8920 |
| 10% (v/v) formaldehyde | Polysciences | Cat# 04018 |
| Sorbitol | Sigma-Aldrich | Cat# 85529 |
| T20 zymolyase | US Biological | Cat# Z1000 |
| Triethanolamine | Sigma-Aldrich | Cat# T9534-100G |
| Acetic anhydride | Fluka | Cat# 45830-250ML-F |
| Dextran sulfate | Sigma-Aldrich | Cat# 6001 |
| Denhardt's solution | Sigma-Aldrich | Cat# D2532 |
| Acetone | VWR Analytical | Cat# BDH1101 |
| Formamide | Sigma-Aldrich | Cat# 9037-100ML |
| Methanol | Avantor | Cat# 0000275255 |
| *E. coli* tRNA | Roche | Cat# 10109541001 |
| Salmon sperm DNA | Invitrogen | Cat# 15632-011 |
| Alexa Fluor 647-labeled 50-mer oligo (dT) probe | Integrated DNA Technologies | N/A |
| DAPI | Invitrogen | Cat# R37606 |
| **Crystallization reagents** | | |
| HEPES | Sigma-Aldrich | Cat# 54457-250G-F |
| PEG 20,000 MME | Sigma-Aldrich | Cat# 81275 |
| PEG 3,350 | Sigma-Aldrich | Cat# 88276 |
| 100% Ethylene glycol | Fluka | Cat# 03750 |
| 2-propanol | Sigma-Aldrich | Cat# 67630 |
| **Western Blot** | | |
| Superblock-PBS Blocking Buffer | Thermo Fisher Scientific | Cat# 37517 |
| Tween-20 | Anatrace | Cat# T1003 |
| Immobilon P transfer membrane | Millipore | Cat# IPVH00010 |
| **Commercial crystallization screens** | | |
| Crystal Screen | Hampton Research | Cat# HR2-110 |

| REAGENT or RESOURCE | SOURCE | IDENTIFIER |
|---|---|---|
| Crystal Screen 2 | Hampton Research | Cat# HR2-112 |
| Index Screen | Hampton Research | Cat# HR2-144 |
| PEG/Ion Screen | Hampton Research | Cat# HR2-126 |
| PEG/Ion 2 Screen | Hampton Research | Cat# HR2-098 |
| ProPlex Screen | Molecular Dimensions | Cat# BN043 |
| PEGs Suite | Qiagen | Cat# 130704 |
| PEGs II Suite | Qiagen | Cat# 130716 |
| **Antibodies** | | |
| Mouse monoclonal anti-HA | BioLegend | Covance Cat# MMS-101P; RRID: AB_231467 |
| Mouse monoclonal anti-FLAG | Sigma-Aldrich | Cat# F1804; RRID: AB_262044 |
| Rabbit polynoclonal anti-hexokinase | US Biological | Cat# H2035-01; RRID: AB_2629457 |
| IRDye 800CW Goat anti-Mouse IgG | LI-COR | P/N 925-32210; RRID: AB_2687825 |
| IRDye 800CW Goat anti-Rabbit IgG | LI-COR | P/N 926-32211; RRID: AB_2651127 |
| **Software** | | |
| XDS | (*90*) | https://xds.mr.mpg.de/ |
| COOT | (*91*) | http://www2.mrc-lmb.cam.ac.uk/personal/pemsley/coot/ |
| PHENIX | (*92*) | http://phenix-online.org/ |
| MolProbity | (*93*) | http://molprobity.biochem.duke.edu/ |
| DIALS | (*95*) | http://dials.github.io/ |
| RESOLVE | (*102*) | http://resolve.lanl.gov/ |
| MotionCor2 | (*105*) | https://msg.ucsf.edu/software |
| cryoSPARC | (*106*) | https://cryosparc.com/ |
| RELION | (*107*) | http://www2.mrc-lmb.cam.ac.uk/relion |
| 3DFSC | (*108*) | https://3dfsc.salk.edu/ |
| UCSF Chimera | (*109*) | http://www.cgl.ucsf.edu/chimera/ |
| PyEM | (*110*) | https://github.com/asarnow/pyem |
| EMRinger | (*112*) | https://github.com/cctbx/cctbx_project/blob/master/mmtbx/command_line/emringer.py |
| PROMALS3D | (*116*) | http://prodata.swmed.edu/promals3d |
| MAFFT | (*117*) | https://mafft.cbrc.jp/alignment/software/ |
| ALSCRIPT | (*118*) | http://www.cryst.bbk.ac.uk/CCSG/info/software/prot-sequ/alscript/alscript.html |
| Fiji | (*128*) | http://imagej.net/software/fiji/ |
| ClustalX | (*129*) | http://www.clustal.org/clustal2/ |
| ASTRA 6 | Wyatt Technology | http://www.wyatt.com/products/software/astra.html |
| Prism | GraphPad Software | http://www.graphpad.com/scientific-software/prism/ |
| PYMOL | Schrödinger | http://pymol.org/2/ |
| IGOR Pro | WaveMetrics | http://www.wavemetrics.com/ |
| Adobe illustrator | Adobe | http://www.adobe.com/products/illustrator.html |
| Adobe photoshop | Adobe | http://www.adobe.com/products/photoshop.html |
| **Expression vectors** | | |
| pET28a-SUMO | (*86*) | N/A |
| pET28a-Avi-SUMO | This study | N/A |
| pET28a-SUMO-Avi | This study | N/A |
| pET28a-His-PreS-Avi | (*26*) | N/A |
| pET28a-PreS | (*85*) | N/A |
| pETDuet1 | Novagen | Cat# 71146-3 |
| pETDuet1-SUMO2 | This study | N/A |
| pET-MCN | (*87*) | N/A |
| pET-MCN-SUMO | (*25*) | N/A |
| pRS411 | (*119*) | N/A |
| pRS413 | (*119*) | N/A |
| pRS415 | (*119*) | N/A |
| pRS416 | (*119*) | N/A |
| pFA6a-natNT2 | (*120*) | N/A |
| pFA6a-kanMX6 | (*121*) | N/A |
| pRS426 | (*130*) | N/A |
| pRS413-$P_{GAL1\text{-}10}$-Cre | This study | N/A |
| **Columns and resins** | | |

| REAGENT or RESOURCE | SOURCE | IDENTIFIER |
|---|---|---|
| HiLoad 16/60 Superdex 200 PG | GE Healthcare | N/A |
| HiLoad 16/60 Superdex 75 PG | GE Healthcare | N/A |
| Superdex 200 10/300 GL | GE Healthcare | N/A |
| Superdex 75 10/300 GL | GE Healthcare | N/A |
| Superdex 200 Increase 10/300 GL | GE Healthcare | N/A |
| HiTrap Q HP | GE Healthcare | N/A |
| HiTrap SP HP | GE Healthcare | N/A |
| MonoQ 10/100 GL | GE Healthcare | N/A |
| HiTrap Heparin HP | GE Healthcare | N/A |
| HiPrep 26/20 Desalting | GE Healthcare | N/A |
| HiTrap Desalting 5 mL | GE Healthcare | N/A |
| Ni-NTA agarose | Qiagen | Cat# 30230 |
| **Cell lines** | | |
| *E. coil* BL21-CodonPlus (DE3)-RIL | Agilent Technologies | Cat# 230245 |
| *E. coli* Max Efficiency DH5α | Invitrogen | Cat# 18258012 |
| *S. cerevisiae* BY4741 | ATCC | Cat# 201388 |

### **Table S2.** Bacterial expression constructs

| | Protein | Residues | Expression vector | Restriction sites (5', 3') | N-terminal overhang | C-terminal overhang | Expression conditions | Reference |
|---|---|---|---|---|---|---|---|---|
| 1 | Nup192 ❄ | 1-1756 | pET28a-PreS-Avi | NdeI, NotI | GPLMSGLNDIFEAQKIEWHEGSAGGS GH | None | 37 °C / 2 hours or 18 °C / 18 hours if co-expressed with Nic96$^{R2}$ | (*36*) |
| 2 | Nup192 (L1071A, I1075A, F1125A, H1126A) | 1-1756 | pET28a-PreS-Avi | NdeI, NotI | GPLMSGLNDIFEAQKIEWHEGSAGGS GH | None | 18 °C / 18 hours co-expressed with Nic96$^{R2}$ | this study |
| 3 | Nup192 (L1071A) | 1-1756 | pET28a-PreS-Avi | NdeI, NotI | GPLMSGLNDIFEAQKIEWHEGSAGGS GH | None | 18 °C / 18 hours co-expressed with Nic96$^{R2}$ | this study |
| 4 | Nup192 (I1075A) | 1-1756 | pET28a-PreS-Avi | NdeI, NotI | GPLMSGLNDIFEAQKIEWHEGSAGGS GH | None | 18 °C / 18 hours co-expressed with Nic96$^{R2}$ | this study |
| 5 | Nup192 (F1125A) | 1-1756 | pET28a-PreS-Avi | NdeI, NotI | GPLMSGLNDIFEAQKIEWHEGSAGGS GH | None | 18 °C / 18 hours co-expressed with Nic96$^{R2}$ | this study |
| 6 | Nup192 (H1126A) | 1-1756 | pET28a-PreS-Avi | NdeI, NotI | GPLMSGLNDIFEAQKIEWHEGSAGGS GH | None | 18 °C / 18 hours co-expressed with Nic96$^{R2}$ | this study |
| 7 | Nup192$^{\Delta Head}$ ♦ | 153-1756 (Δ167-184, replaced with GSGS) | pET28a-SUMO | NdeI, Not1 | GPHM | None | 18 °C / 18 hours co-expressed with Nic96$^{187\text{-}301}$ or Nic96$^{R2}$ | (*30*) |
| 8 | Nup192 (L1584A) | 1-1756 | pET28a-PreS-Avi | NdeI, NotI | GPLMSGLNDIFEAQKIEWHEGSAGGS GH | None | 37 °C / 2 hours | this study |
| 9 | Nup192 (F1735A) | 1-1756 | pET28a-PreS-Avi | NdeI, NotI | GPLMSGLNDIFEAQKIEWHEGSAGGS GH | None | 37 °C / 2 hours | this study |
| 10 | Nup192 (L1584A F1735A) | 1-1756 | pET28a-PreS-Avi | NdeI, NotI | GPLMSGLNDIFEAQKIEWHEGSAGGS GH | None | 37 °C / 2 hours | this study |
| 11 | Nup192 (L1584E A1648E) | 1-1756 | pET28a-PreS-Avi | NdeI, NotI | GPLMSGLNDIFEAQKIEWHEGSAGGS GH | None | 37 °C / 2 hours | this study |
| 12 | Nup192 (F1735E) | 1-1756 | pET28a-PreS-Avi | NdeI, NotI | GPLMSGLNDIFEAQKIEWHEGSAGGS GH | None | 37 °C / 2 hours | this study |
| 13 | Nup192 (L1584E A1648E F1735E) | 1-1756 | pET28a-PreS-Avi | NdeI, NotI | GPLMSGLNDIFEAQKIEWHEGSAGGS GH | None | 37 °C / 2 hours | this study |
| 14 | Nup188 ❄ ♦ | 1-1858 | pET28a-PreS | AseI, BamHI | GPHN | None | 18 °C / 18 hours co-expressed with Nic96$^{R2}$ | (*30*) |
| 15 | Nup188 (H409A H412A M485A I535A) | 1-1858 | pET28a-PreS | AseI, BamHI | GPHN | None | 18 °C / 18 hours co-expressed with Nic96$^{R2}$ | this study |
| 16 | Nup188 | 1-1858 | pET28a-PreS | AseI, BamHI | GPHN | AAALEHHHHHH | 30 °C / 3 hours | this study |
| 17 | Nup188 (F1487A) | 1-1858 | pET28a-PreS | AseI, BamHI | GPHN | AAALEHHHHHH | 30 °C / 3 hours | this study |
| 18 | Nup188 (L1689A) | 1-1858 | pET28a-PreS | AseI, BamHI | GPHN | AAALEHHHHHH | 30 °C / 3 hours | this study |
| 19 | Nup188 (V1760A) | 1-1858 | pET28a-PreS | AseI, BamHI | GPHN | AAALEHHHHHH | 30 °C / 3 hours | this study |
| 20 | Nup188 (F1487A L1689A V1760A) | 1-1858 | pET28a-PreS | AseI, BamHI | GPHN | AAALEHHHHHH | 30 °C / 3 hours | this study |
| 21 | Nup188 (F1487E) | 1-1858 | pET28a-PreS | AseI, BamHI | GPHN | AAALEHHHHHH | 30 °C / 3 hours | this study |
| 22 | Nup188 (L1689E V1760E) | 1-1858 | pET28a-PreS | AseI, BamHI | GPHN | AAALEHHHHHH | 30 °C / 3 hours | this study |
| 23 | Nup188 (F1487E L1689E V1760E) | 1-1858 | pET28a-PreS | AseI, BamHI | GPHN | AAALEHHHHHH | 30 °C / 3 hours | this study |
| 24 | Nup188$^{NTD}$ ♦ | 1-1134 | pET28a-PreS | AseI, BamHI | GPHN | None | 18 °C / 18 hours | (*30*) |
| 25 | Nup188$^{NTD}$ (H409A H412A) | 1-1134 | pET28a-PreS | AseI, BamHI | GPHN | None | 18 °C / 18 hours | this study |
| 26 | Nup188$^{NTD}$ (I477A) | 1-1134 | pET28a-PreS | AseI, BamHI | GPHN | None | 18 °C / 18 hours | this study |
| 27 | Nup188$^{NTD}$ (M485A) | 1-1134 | pET28a-PreS | AseI, BamHI | GPHN | None | 18 °C / 18 hours | this study |
| 28 | Nup188$^{NTD}$ (I535A) | 1-1134 | pET28a-PreS | AseI, BamHI | GPHN | None | 18 °C / 18 hours | this study |
| 29 | Nup188$^{NTD}$ (H409A H412A M485A I535A) | 1-1134 | pET28a-PreS | AseI, BamHI | GPHN | None | 18 °C / 18 hours | this study |
| 30 | Nic96$^{187\text{-}301}$ ♦ | 187-301 | pET-MCN-SUMO | BamHI, NotI | His$_6$-SUMO / S | None | 18 °C / 18 hours | this study |
| 31 | Nic96$^{R2}$ ❄ ♦ | 240-301 | pET-MCN-SUMO | BamHI, NotI | His$_6$-SUMO / S | None | 18 °C / 18 hours alone, or co-expressed with Nup188 or Nup192 | this study |
| 32 | Nic96$^{R2}$ (I94M) ♦ | 240-301 | pET-MCN-SUMO | BamHI, NotI | S | None | 18 °C / 18 hours co-expressed with Nup192 | this study |
| 33 | Nic96$^{R2}$ (F275A) | 240-301 | pET-MCN-SUMO | BamHI, NotI | His$_6$-SUMO | None | 18 °C / 18 hours | this study |
| 34 | Nic96 (F278A) | 240-301 | pET-MCN-SUMO | BamHI, NotI | His$_6$-SUMO | None | 18 °C / 18 hours | this study |
| 35 | Nic96$^{R2}$ (F298A) | 240-301 | pET-MCN-SUMO | BamHI, NotI | His$_6$-SUMO | None | 18 °C / 18 hours | this study |
| 36 | Nic96$^{R2}$ (F275E F278E) | 240-301 | pET-MCN-SUMO | BamHI, NotI | His$_6$-SUMO | None | 18 °C / 18 hours | this study |
| 37 | Nic96$^{R2}$ (F275A F278A F298A) | 240-301 | pET-MCN-SUMO | BamHI, NotI | His$_6$-SUMO | None | 18 °C / 18 hours | this study |
| 38 | Nic96$^{R2}$ (F298E) | 240-301 | pET-MCN-SUMO | BamHI, NotI | His$_6$-SUMO | None | 18 °C / 18 hours | this study |
| 39 | Nic96$^{R2}$ | 240-301 | pET-MCN-SUMO | BamHI, NotI | His$_6$-SUMO | None | 18 °C / 18 hours | this study |

| | Protein | Residues | Expression vector | Restriction sites (5’, 3’) | N-terminal overhang | C-terminal overhang | Expression conditions | Reference |
|---|---|---|---|---|---|---|---|---|
| | (F275E F278E F298E) | | | | | | | |
| 40 | Nup53 | 1-361 | pET28a - SUMO | BamHI, XhoI | S | LEHHHHHH | 23 °C / 18 hours | (*30*) |
| 41 | Nup53$^{R1}$ ❄ | 31-67 | pET28a - SUMO | BamHI, NotI | S | Y | 37 °C / 2 hours | this study |
| 42 | Nup145N | 606-993 | pET28a-SUMO | BamHI, NotI | S | None | 23 °C / 18 hours | this study |
| 43 | Nup145N | 606-683 | pET28a-SUMO | BamHI, XhoI | His$_6$-SUMO | AAALEHHHHHH | 37 °C / 1 hour | (*30*) |
| 44 | Nup145N | 606-673 | pET28a-SUMO | BamHI, NotI | His$_6$-SUMO | AAALEHHHHHH | 37 °C / 1 hour | this study |
| 45 | Nup145N | 606-663 | pET28a-SUMO | BamHI, NotI | His$_6$-SUMO | AAALEHHHHHH | 37 °C / 1 hour | this study |
| 46 | Nup145N | 606-657 | pET28a-SUMO | BamHI, NotI | His$_6$-SUMO | AAALEHHHHHH | 37 °C / 1 hour | this study |
| 47 | Nup145N | 606-650 | pET28a-SUMO | BamHI, NotI | His$_6$-SUMO | AAALEHHHHHH | 37 °C / 1 hour | this study |
| 48 | Nup145N | 606-639 | pET28a-SUMO | BamHI, NotI | His$_6$-SUMO | AAALEHHHHHH | 37 °C / 1 hour | this study |
| 49 | Nup145N$^{R1}$❄ | 616-683 | pET28a-SUMO | BamHI, NotI | His$_6$-SUMO / S | AAALEHHHHHH | 37 °C / 1 hour | this study |
| 50 | Nup145N | 626-683 | pET28a-SUMO | BamHI, NotI | His$_6$-SUMO | AAALEHHHHHH | 37 °C / 1 hour | this study |
| 51 | Nup145N | 632-683 | pET28a-SUMO | BamHI, NotI | His$_6$-SUMO | AAALEHHHHHH | 37 °C / 1 hour | this study |
| 52 | Nup145N | 616-673 | pET28a-SUMO | BamHI, NotI | His$_6$-SUMO | AAALEHHHHHH | 37 °C / 1 hour | this study |
| 53 | Nup145N | 616-663 | pET28a-SUMO | BamHI, NotI | His$_6$-SUMO | AAALEHHHHHH | 37 °C / 1 hour | this study |
| 54 | Nup145N | 606-750 | pET28a-SUMO | BamHI, NotI | His$_6$-SUMO | None | 37 °C / 1 hour | this study |
| 55 | Nup145N | 606-732 | pET28a-SUMO | BamHI, NotI | His$_6$-SUMO | None | 37 °C / 1 hour | this study |
| 56 | Nup145N | 606-720 | pET28a-SUMO | BamHI, NotI | His$_6$-SUMO | None | 37 °C / 1 hour | this study |
| 57 | Nup145N | 606-710 | pET28a-SUMO | BamHI, NotI | His$_6$-SUMO | None | 37 °C / 1 hour | this study |
| 58 | Nup145N | 606-700 | pET28a-SUMO | BamHI, NotI | His$_6$-SUMO | None | 37 °C / 1 hour | this study |
| 59 | Nup145N | 649-750 | pET28a-SUMO | BamHI, NotI | His$_6$-SUMO | None | 37 °C / 1 hour | this study |
| 60 | Nup145N | 660-750 | pET28a-SUMO | BamHI, NotI | His$_6$-SUMO | None | 37 °C / 1 hour | this study |
| 61 | Nup145N | 675-750 | pET28a-SUMO | BamHI, NotI | His$_6$-SUMO | None | 37 °C / 1 hour | this study |
| 62 | Nup145N | 695-750 | pET28a-SUMO | BamHI, NotI | His$_6$-SUMO | None | 37 °C / 1 hour | this study |
| 63 | Nup145N | 715-750 | pET28a-SUMO | BamHI, NotI | His$_6$-SUMO | None | 37 °C / 1 hour | this study |
| 64 | Nup145N$^{R2}$ ❄ | 640-732 | pET28a-SUMO | BamHI, NotI | His$_6$-SUMO / S | None | 37 °C / 1.5 hours | this study |
| 65 | Nup145N | 650-732 | pET28a-SUMO | BamHI, NotI | His$_6$-SUMO | None | 37 °C / 1 hour | this study |
| 66 | Nup145N | 660-732 | pET28a-SUMO | BamHI, NotI | His$_6$-SUMO | None | 37 °C / 1 hour | this study |
| 67 | Nup145N | 700-732 | pET28a-SUMO | BamHI, NotI | His$_6$-SUMO | None | 37 °C / 1 hour | this study |
| 68 | Nup145N (D606A, D607A, K608A, I609A, Q610A) | 606-993 | pET28a-SUMO | BamHI, NotI | S | None | 23 °C / 18 hours | this study |
| 69 | Nup145N (N611A, P612A, G613A, P614A, L615A) | 606-993 | pET28a-SUMO | BamHI, NotI | S | None | 23 °C / 18 hours | this study |
| 70 | Nup145N (N616A, P617A, G618A, P619A, L620A) | 606-993 | pET28a-SUMO | BamHI, NotI | S | None | 23 °C / 18 hours | this study |
| 71 | Nup145N (G621A, K622A, K624A, V625A) | 606-993 | pET28a-SUMO | BamHI, NotI | S | None | 23 °C / 18 hours | this study |
| 72 | Nup145N (K626A, S627A, R628A, S629A, I630A) | 606-993 | pET28a-SUMO | BamHI, NotI | S | None | 23 °C / 18 hours | this study |
| 73 | Nup145N (L631A, P632A, M633A, Y634A, K635A) | 606-993 | pET28a-SUMO | BamHI, NotI | S | None | 23 °C / 18 hours | this study |
| 74 | Nup145N (L636A, S637A, P638A, N640A) | 606-993 | pET28a-SUMO | BamHI, NotI | S | None | 23 °C / 18 hours | this study |
| 75 | Nup145N (S642A, R643A, L644A, V645A) | 606-993 | pET28a-SUMO | BamHI, NotI | S | None | 23 °C / 18 hours | this study |
| 76 | Nup145N (T646A, T647A, P648A, Q649, K650A) | 606-993 | pET28a-SUMO | BamHI, NotI | S | None | 23 °C / 18 hours | this study |
| 77 | Nup145N (R651A, Y653A, G654A, F655A) | 606-993 | pET28a-SUMO | BamHI, NotI | S | None | 23 °C / 18 hours | this study |
| 78 | Nup145N (S656A, F657A, S658A, Y660A) | 606-993 | pET28a-SUMO | BamHI, NotI | S | None | 23 °C / 18 hours | this study |

| | Protein | Residues | Expression vector | Restriction sites (5', 3') | N-terminal overhang | C-terminal overhang | Expression conditions | Reference |
|---|---|---|---|---|---|---|---|---|
| 79 | 606-993 (G661A, S662A, P663A, T664A, S665A) | 606-993 | pET28a-SUMO | BamHI, NotI | S | None | 23 °C / 18 hours | this study |
| 80 | Nup145N (P666A, S667A, S668A, S669A) | 606-993 | pET28a-SUMO | BamHI, NotI | S | None | 23 °C / 18 hours | this study |
| 81 | Nup145N (S671A, S672A, T673A, P674A, G675A) | 606-993 | pET28a-SUMO | BamHI, NotI | S | None | 23 °C / 18 hours | this study |
| 82 | Nup145N (F677A, G678A, Q679A, S680A) | 606-993 | pET28a-SUMO | BamHI, NotI | S | None | 23 °C / 18 hours | this study |
| 83 | Nup145N (I681A, L682A, S683A, S684A, S685A) | 606-993 | pET28a-SUMO | BamHI, NotI | S | None | 23 °C / 18 hours | this study |
| 84 | Nup145N (I686A, N687A, R688A, G689A, L690A) | 606-993 | pET28a-SUMO | BamHI, NotI | S | None | 23 °C / 18 hours | this study |
| 85 | Nup145N (N691A, K692A, S693A, I694A, S695A) | 606-993 | pET28a-SUMO | BamHI, NotI | S | None | 23 °C / 18 hours | this study |
| 86 | Nup145N (S697A, N698A, L699A, R700A) | 606-993 | pET28a-SUMO | BamHI, NotI | S | None | 23 °C / 18 hours | this study |
| 87 | Nup145N (R701A, S702A, L703A, N704A, V705A) | 606-993 | pET28a-SUMO | BamHI, NotI | S | None | 23 °C / 18 hours | this study |
| 88 | Nup145N (E706A, D707A, S708A, I709A, L710A) | 606-993 | pET28a-SUMO | BamHI, NotI | S | None | 23 °C / 18 hours | this study |
| 89 | Nup145N (Q711A, P712A, G713A, F715A) | 606-993 | pET28a-SUMO | BamHI, NotI | S | None | 23 °C / 18 hours | this study |
| 90 | Nup145N (S716A, N718A, S719A, S720A) | 606-993 | pET28a-SUMO | BamHI, NotI | S | None | 23 °C / 18 hours | this study |
| 91 | Nup145N (M721A, R722A, L723A, L724A, G725A) | 606-993 | pET28a-SUMO | BamHI, NotI | S | None | 23 °C / 18 hours | this study |
| 92 | Nup145N (G726A, P727A, G728A, S729A, H730A) | 606-993 | pET28a-SUMO | BamHI, NotI | S | None | 23 °C / 18 hours | this study |
| 93 | Nup145N (K731A, K732A, L733A, V734A, I735A) | 606-993 | pET28a-SUMO | BamHI, NotI | S | None | 23 °C / 18 hours | this study |
| 94 | Nup145N (N736A, K737A, D738A, M739A, R740A) | 606-993 | pET28a-SUMO | BamHI, NotI | S | None | 23 °C / 18 hours | this study |
| 95 | Nup145N (T741A, D742A, L743A, F744A, S745A) | 606-993 | pET28a-SUMO | BamHI, NotI | S | None | 23 °C / 18 hours | this study |
| 96 | Nup145N (L619A) | 606-993 | pET28a-SUMO | BamHI, NotI | S | None | 23 °C / 18 hours | this study |
| 97 | Nup145N (K622A) | 606-993 | pET28a-SUMO | BamHI, NotI | S | None | 23 °C / 18 hours | this study |
| 98 | Nup145N (K624A) | 606-993 | pET28a-SUMO | BamHI, NotI | S | None | 23 °C / 18 hours | this study |
| 99 | Nup145N (K626A) | 606-993 | pET28a-SUMO | BamHI, NotI | S | None | 23 °C / 18 hours | this study |
| 100 | Nup145N (R628A) | 606-993 | pET28a-SUMO | BamHI, NotI | S | None | 23 °C / 18 hours | this study |
| 101 | Nup145N (I630A, L631A) | 606-993 | pET28a-SUMO | BamHI, NotI | S | None | 23 °C / 18 hours | this study |
| 102 | Nup145N (M633A) | 606-993 | pET28a-SUMO | BamHI, NotI | S | None | 23 °C / 18 hours | this study |
| 103 | Nup145N (Y634A) | 606-993 | pET28a-SUMO | BamHI, NotI | S | None | 23 °C / 18 hours | this study |
| 104 | Nup145N (K635A) | 606-993 | pET28a-SUMO | BamHI, NotI | S | None | 23 °C / 18 hours | this study |
| 105 | Nup145N (L636A) | 606-993 | pET28a-SUMO | BamHI, NotI | S | None | 23 °C / 18 hours | this study |
| 106 | Nup145N (S637A) | 606-993 | pET28a-SUMO | BamHI, NotI | S | None | 23 °C / 18 hours | this study |
| 107 | Nup145N (P632A, P638A) | 606-993 | pET28a-SUMO | BamHI, NotI | S | None | 23 °C / 18 hours | this study |
| 108 | Nup145N (R643A) | 606-993 | pET28a-SUMO | BamHI, NotI | S | None | 23 °C / 18 hours | this study |
| 109 | Nup145N (K650A, R651A) | 606-993 | pET28a-SUMO | BamHI, NotI | S | None | 23 °C / 18 hours | this study |
| 110 | Nup145N (F655A) | 606-993 | pET28a-SUMO | BamHI, NotI | S | None | 23 °C / 18 hours | this study |
| 111 | Nup145N (R700A) | 606-993 | pET28a-SUMO | BamHI, NotI | S | None | 23 °C / 18 hours | this study |
| 112 | Nup145N (R701A) | 606-993 | pET28a-SUMO | BamHI, NotI | S | None | 23 °C / 18 hours | this study |
| 113 | Nup145N (L703A) | 606-993 | pET28a-SUMO | BamHI, NotI | S | None | 23 °C / 18 hours | this study |

| | Protein | Residues | Expression vector | Restriction sites (5', 3') | N-terminal overhang | C-terminal overhang | Expression conditions | Reference |
|---|---|---|---|---|---|---|---|---|
| 114 | Nup145N (N704A) | 606-993 | pET28a-SUMO | BamHI, NotI | S | None | 23 °C / 18 hours | this study |
| 115 | Nup145N (E706A) | 606-993 | pET28a-SUMO | BamHI, NotI | S | None | 23 °C / 18 hours | this study |
| 116 | Nup145N (D707A) | 606-993 | pET28a-SUMO | BamHI, NotI | S | None | 23 °C / 18 hours | this study |
| 117 | Nup145N (I709A) | 606-993 | pET28a-SUMO | BamHI, NotI | S | None | 23 °C / 18 hours | this study |
| 118 | Nup145N (L710A) | 606-993 | pET28a-SUMO | BamHI, NotI | S | None | 23 °C / 18 hours | this study |
| 119 | Nup145N (Q711A) | 606-993 | pET28a-SUMO | BamHI, NotI | S | None | 23 °C / 18 hours | this study |
| 120 | Nup145N (F715A) | 606-993 | pET28a-SUMO | BamHI, NotI | S | None | 23 °C / 18 hours | this study |
| 121 | Nup145N (N718A) | 606-993 | pET28a-SUMO | BamHI, NotI | S | None | 23 °C / 18 hours | this study |
| 122 | Nup145N (K624A, K626A, R628A) | 606-993 | pET28a-SUMO | BamHI, NotI | S | None | 23 °C / 18 hours | this study |
| 123 | Nup145N (M633A, Y634A, K635A, L636A) | 606-993 | pET28a-SUMO | BamHI, NotI | S | None | 23 °C / 18 hours | this study |
| 124 | Nup145N (R643A K650A, R651A) | 606-993 | pET28a-SUMO | BamHI, NotI | S | None | 23 °C / 18 hours | this study |
| 125 | Nup145N (K624A, K626A, R628A, M633A, Y634A, K635A, L636A) | 606-993 | pET28a-SUMO | BamHI, NotI | S | None | 23 °C / 18 hours | this study |
| 126 | Nup145N (M633A, Y634A, K635A L636A, R643A, K650A, R651A) | 606-993 | pET28a-SUMO | BamHI, NotI | S | None | 23 °C / 18 hours | this study |
| 127 | Nup145N (K624A, K626A, R628A, R643A, K650A, R651A) | 606-993 | pET28a-SUMO | BamHI, NotI | S | None | 23 °C / 18 hours | this study |
| 128 | Nup145N (K624A, K626A, R628A M633A, Y634A, K635A L636A, R643A, K650A, R651A) | 606-993 | pET28a-SUMO | BamHI, NotI | S | None | 23 °C / 18 hours | this study |
| 129 | Nup145N (E706A, D707A, S708A, I709A, L710A, F715A) | 640-732 | pET28a-SUMO | BamHI, NotI | S | None | 37 °C / 1.5 hours | this study |
| 130 | NUP54 | 118-507 | pET28a-SUMO | BamHI, XhoI | S | None | 18 °C / 18 hours co-expressed with NUP58 and NUP62 | this study |
| 131 | NUP58<br>NUP62 | 233-454<br>317-522 | pET-Duet1-SUMO2 | NcoI, NotI<br>NdeI, XhoII | S<br>S | None<br>None | 18 °C / 18 hours co-expressed with NUP54 | this study |
| 132 | NUP93$^{R1}$ | 2-51 | pET28a-SUMO | BamHI, NotI | $His_6$-SUMO(G-Y) | None | 37 °C / 1 hour | this study |
| 133 | NUP93$^{R1}$ (L33A, I36A, L43A) | 2-51 | pET28a-SUMO | BamHI, NotI | $His_6$-SUMO(G-Y) | None | 37 °C / 1 hour | this study |
| 134 | NUP93$^{\Delta R2\text{-}SOL}$ | 2-93 | pET28a-SUMO | BamHI, NotI | $His_6$-SUMO(G-Y) | None | 37 °C / 1 hour | this study |
| 135 | NUP93$^{\Delta R2\text{-}SOL}$ (L33A, I36A, L43A) | 2-93 | pET28a-SUMO | BamHI, NotI | $His_6$-SUMO(G-Y) | None | 37 °C / 1 hour | this study |
| 136 | NUP93$^{R2}$ | 95-150 | pET28a-Avi-SUMO | BamHI, NotI | $His_6$-Avi-SUMO(G-R) | None | 37 °C / 1 hour | this study |
| 137 | NUP93$^{R2}$ (I114E, Y128E, W137E, I144E, L148E) | 95-150 | pET28a-Avi-SUMO | BamHI, NotI | $His_6$-Avi-SUMO(G-R) | None | 37 °C / 1 hour | this study |
| 138 | NUP93$^{SOL}$ ♦ | 174-819 | pET28a-SUMO-Avi | BamHI, NotI | SSGLNDIFEAQKIEWHEGSAGGSGGS | None | 18 °C / 18 hours | this study |
| 139 | NUP93$^{SOL}$ (Y269A) | 174-819 | pET28a-SUMO-Avi | BamHI, NotI | SSGLNDIFEAQKIEWHEGSAGGSGGS | None | 18 °C / 18 hours | this study |
| 140 | NUP93$^{SOL}$ (Y413A) | 174-819 | pET28a-SUMO-Avi | BamHI, NotI | SSGLNDIFEAQKIEWHEGSAGGSGGS | None | 18 °C / 18 hours | this study |
| 141 | NUP93$^{SOL}$ (Y448A) | 174-819 | pET28a-SUMO-Avi | BamHI, NotI | SSGLNDIFEAQKIEWHEGSAGGSGGS | None | 18 °C / 18 hours | this study |
| 142 | NUP93$^{SOL}$ (Y269E) | 174-819 | pET28a-SUMO-Avi | BamHI, NotI | SSGLNDIFEAQKIEWHEGSAGGSGGS | None | 18 °C / 18 hours | this study |
| 143 | NUP93$^{SOL}$ (Y413E) | 174-819 | pET28a-SUMO-Avi | BamHI, NotI | SSGLNDIFEAQKIEWHEGSAGGSGGS | None | 18 °C / 18 hours | this study |
| 144 | NUP93$^{SOL}$ (Y448E) | 174-819 | pET28a-SUMO-Avi | BamHI, NotI | SSGLNDIFEAQKIEWHEGSAGGSGGS | None | 18 °C / 18 hours | this study |
| 145 | NUP53 | 84-250 | pET-MCN-SUMO | BamHI, NotI | $His_6$-SUMO / S | None | 18 °C / 18 hours | this study |
| 146 | NUP53 | 94-250 | pET-MCN-SUMO | BamHI, NotI | $His_6$-SUMO | None | 18 °C / 18 hours | this study |
| 147 | NUP53 | 84-105 | pET-MCN-SUMO | BamHI, NotI | $His_6$-SUMO | None | 18 °C / 18 hours | this study |
| 148 | NUP53 | 89-105 | pET-MCN-SUMO | BamHI, NotI | $His_6$-SUMO | None | 18 °C / 18 hours | this study |
| 149 | NUP53 ♦ | 84-150 | pET-MCN-SUMO | BamHI, NotI | $His_6$-SUMO / S | None | 18 °C / 18 hours | this study |
| 150 | NUP53 (I94M) ♦ | 84-150 | pET-MCN-SUMO | BamHI, NotI | S | None | 18 °C / 18 hours | this study |

| | Protein | Residues | Expression vector | Restriction sites (5', 3') | N-terminal overhang | C-terminal overhang | Expression conditions | Reference |
|---|---|---|---|---|---|---|---|---|
| 151 | NUP53 | 84-140 | pET-MCN-SUMO | BamHI, NotI | $His_6$-SUMO | None | 18 °C / 18 hours | this study |
| 152 | NUP53 | 84-130 | pET-MCN-SUMO | BamHI, NotI | $His_6$-SUMO | None | 18 °C / 18 hours | this study |
| 153 | NUP53 | 84-120 | pET-MCN-SUMO | BamHI, NotI | $His_6$-SUMO | None | 18 °C / 18 hours | this study |
| 154 | NUP53 | 84-110 | pET-MCN-SUMO | BamHI, NotI | $His_6$-SUMO | None | 18 °C / 18 hours | this study |
| 155 | NUP53 | 84-105 | pET-MCN-SUMO | BamHI, NotI | $His_6$-SUMO | None | 18 °C / 18 hours | this study |
| 156 | NUP53 | 84-100 | pET-MCN-SUMO | BamHI, NotI | $His_6$-SUMO | None | 18 °C / 18 hours | this study |
| 157 | NUP53 | 70-110 | pET-MCN-SUMO | BamHI, NotI | $His_6$-SUMO | None | 18 °C / 18 hours | this study |
| 158 | NUP53 | 90-110 | pET-MCN-SUMO | BamHI, NotI | $His_6$-SUMO | None | 18 °C / 18 hours | this study |
| 159 | $NUP53^N$ | 1-169 | pET28a-SUMO | BamHI, NotI | $His_6$-SUMO | None | 18 °C / 18 hours | this study |
| 160 | $NUP53^N$ (G71A, G72A, S73A, P74A, P75A) | 1-169 | pET28a-SUMO | BamHI, NotI | $His_6$-SUMO | None | 18 °C / 18 hours | this study |
| 161 | $NUP53^N$ (Q76A, P77A, V78A, V79A, P80A) | 1-169 | pET28a-SUMO | BamHI, NotI | $His_6$-SUMO | None | 18 °C / 18 hours | this study |
| 162 | $NUP53^N$ (A81A, H82A, K83A, D84A, K85A) | 1-169 | pET28a-SUMO | BamHI, NotI | $His_6$-SUMO | None | 18 °C / 18 hours | this study |
| 163 | $NUP53^N$ (S86A, G87A, A88A, P89A, P90A) | 1-169 | pET28a-SUMO | BamHI, NotI | $His_6$-SUMO | None | 18 °C / 18 hours | this study |
| 164 | $NUP53^N$ (V91A, R92A, S93A, I94A, Y95A) | 1-169 | pET28a-SUMO | BamHI, NotI | $His_6$-SUMO | None | 18 °C / 18 hours | this study |
| 165 | $NUP53^N$ (D96A, D97A, I98A, S99A, S100A) | 1-169 | pET28a-SUMO | BamHI, NotI | $His_6$-SUMO | None | 18 °C / 18 hours | this study |
| 166 | $NUP53^N$ (P101A, G102A, L103A, G104A, S105A) | 1-169 | pET28a-SUMO | BamHI, NotI | $His_6$-SUMO | None | 18 °C / 18 hours | this study |
| 167 | $NUP53^N$ P89A | 1-169 | pET28a-SUMO | BamHI, NotI | $His_6$-SUMO | None | 18 °C / 18 hours | this study |
| 168 | $NUP53^N$ P90A | 1-169 | pET28a-SUMO | BamHI, NotI | $His_6$-SUMO | None | 18 °C / 18 hours | this study |
| 169 | $NUP53^N$ V91A | 1-169 | pET28a-SUMO | BamHI, NotI | $His_6$-SUMO | None | 18 °C / 18 hours | this study |
| 170 | $NUP53^N$ R92A | 1-169 | pET28a-SUMO | BamHI, NotI | $His_6$-SUMO | None | 18 °C / 18 hours | this study |
| 171 | $NUP53^N$ S93A | 1-169 | pET28a-SUMO | BamHI, NotI | $His_6$-SUMO | None | 18 °C / 18 hours | this study |
| 172 | $NUP53^N$ I94A | 1-169 | pET28a-SUMO | BamHI, NotI | $His_6$-SUMO | None | 18 °C / 18 hours | this study |
| 173 | $NUP53^N$ Y95A | 1-169 | pET28a-SUMO | BamHI, NotI | $His_6$-SUMO | None | 18 °C / 18 hours | this study |
| 174 | $NUP53^N$ | 1-169 | pET28a-Avi-SUMO | BamHI, NotI | $His_6$-Avi-SUMO(G-R) | None | 18 °C / 18 hours | this study |
| 175 | $NUP53^{N\,\Delta R1}$ | 71-169 | pET28a-Avi-SUMO | BamHI, NotI | $His_6$-Avi-SUMO(G-R) | None | 18 °C / 18 hours | this study |
| 176 | NUP53 | 1-313 | pET28a-SUMO | BamHI, NotI | S | AAALEHHHHHH | 18 °C / 18 hours | this study |
| 177 | $NUP53^{R1-RRM}$ | 1-250 | pET28a-SUMO | BamHI, NotI | S | None | 18 °C / 18 hours | this study |
| 178 | $NUP53^{RRM-R3}$ | 170-313 | pET28a-SUMO | BamHI, NotI | S | AAALEHHHHHH | 18 °C / 18 hours | this study |
| 179 | $NUP53^{R3}$ | 251-313 | pET28a-SUMO | BamHI, NotI | $His_6$-SUMO | None | 18 °C / 18 hours | this study |
| 180 | $NUP155^{NTD}$ | 1-886 Δ260-269 | pET28a-SUMO | BamHI, NotI | S | None | 18 °C / 18 hours | this study |
| 181 | NUP98 | 464-880 | pET28a-Avi-SUMO | BamHI, NotI | $His_6$-Avi-SUMO(G-R) | None | 18 °C / overnight | this study |
| 182 | $NUP98^{R3-APD}$ | 596-880 | pET28a-Avi-SUMO | BamHI, NotI | $His_6$-Avi-SUMO(G-R) | None | 18 °C / 18 hours | this study |
| 183 | Avi-SUMO | -- | pET28a-Avi-SUMO | -- | $His_6$-Avi-SUMO | None | 18 °C / 18 hours | this study |

♦ Constructs that were used for crystallization

❄ Constructs that were used for cryo-EM analysis

$His_6$-SUMO: MGSSHHHHHHSSGLVPRGSHMASMSDSEVNQEAKPEVKPEVKPETHINLKVSDGSSEIFFKIKKTTPLRRLMEAFAKRQGKEMDSLRFLYDGIRIQADQTPEDLDMEDNDIIEAHREQIG**G**S

$His_6$-SUMO(G-Y): MGSSHHHHHHSSGLVPRGSHMASMSDSEVNQEAKPEVKPEVKPETHINLKVSDGSSEIFFKIKKTTPLRRLMEAFAKRQGKEMDSLRFLYDGIRIQADQTPEDLDMEDNDIIEAHREQIG**Y**S

$His_6$-SUMO(G-R): MGSSHHHHHHSSGLVPRGSHMASMSDSEVNQEAKPEVKPEVKPETHINLKVSDGSSEIFFKIKKTTPLRRLMEAFAKRQGKEMDSLRFLYDGIRIQADQTPEDLDMEDNDIIEAHREQIG**R**S

$His_6$-Avi-SUMO: MGSSHHHHHHSSGLVPRGSHMSGLNDIFEAQKIEWHEGSAGGSGHMASMSDSEXNQEAKPEVKPEVKPETHINLKVSDGSSEIFFKIKKTTPLRRLMEAFAKRQGKEMDSLRFLYDGIRIQAD QTPEDLDMEDNDIIEAHREQIG**G**S

$His_6$-Avi-SUMO(G-R): MGSSHHHHHHSSGLVPRGSHMSGLNDIFEAQKIEWHEGSAGGSGHMASMSDSEXNQEAKPEVKPEVKPETHINLKVSDGSSEIFFKIKKTTPLRRLMEAFAKRQGKEMDSLRFLYDGI RIQADQTPEDLDMEDNDIIEAHREQIG**R**S

### **Table S3.** Protein purification protocols

| Protein(s) | Expression constructs | Purification step | Buffer A | Buffer B |
|---|---|---|---|---|
| **Nup192** | Individual | 1. Ni-NTA | 1. Ni-A1 | 1. Ni-B1 |
| wildtype and mutants | (#1, #8-13) | 2. Dialysis/Cleavage | 2. Ni-A2 / PreS | 2. N/A |
| | | 3. Ni-NTA | 3. Ni-A2 | 3. Ni-B1 |
| | | 4. MonoQ 5/50 GL | 4. IEX-A1 | 4. IEX-B1 |
| | | 5. HiLoad Superdex 200 16/60 PG | 5. SEC-A | 5. N/A |
| **Nup192•SUMO-Nic96$^{R2}$** | Co-expression | 1. Ni-NTA | 1. Ni-A1 | 1. Ni-B1 |
| wildtype and mutants | (#1-7 and #30 or #31) | 2. Dialysis/Cleavage | 2. IEX-A1 / PreS | 2. N/A |
| | | 3. Ni-NTA | 3. Ni-A2 | 3. Ni-B1 |
| | | 4. MonoQ 5/50 GL | 4. IEX-A1 | 4. IEX-B1 |
| | | 5. HiLoad Superdex 200 16/60 PG | 5. SEC-A | 5. N/A |
| **Nup188** | Individual | 1. Ni-NTA | 1. Ni-A1 | 1. Ni-B1 |
| wildtype and mutants | (#16-23) | 2. Dialysis/Cleavage | 2. Ni-A2 / PreS | 2. N/A |
| | | 3. Ni-NTA | 3. Ni-A2 | 3. Ni-B1 |
| | | 4. Desalting | 4.IEX-A1 | 4. N/A |
| | | 5. MonoQ 5/50 GL | 5. IEX-A1 | 5. IEX-B1 |
| | | 6. HiLoad Superdex 200 16/60 PG | 6. SEC-A | 6. N/A |
| **Nup188•SUMO-Nic96$^{R2}$** | Co-expression | 1. Ni-NTA | 1. Ni-A1 | 1. Ni-B1 |
| wildtype and mutants | (#14-15 and #30) | 2. Dialysis/Cleavage | 2. IEX-A1 / PreS | 2. N/A |
| | | 3. Ni-NTA | 3. Ni-A2 | 3. Ni-B1 |
| | | 4. Desalting | 4. IEX-A1 | 4. N/A |
| | | 5. MonoQ 5/50 GL | 5. IEX-A1 | 5. IEX-B1 |
| | | 6. HiLoad Superdex 200 16/60 PG | 6. SEC-A | 6. N/A |
| **Nup188$^{NTD}$** | Individual | 1. Ni-NTA | 1. Ni-A1 | 1. Ni-B1 |
| wildtype and mutants | (#24-29) | 2. Dialysis/Cleavage | 2. Ni-A2 / PreS | 2. N/A |
| | | 3. Ni-NTA | 3. Ni-A2 | 3. Ni-B2 |
| | | 4. MonoQ 10/100 GL | 4. IEX-A1 | 4. IEX-B1 |
| | | 5. HiLoad Superdex 200 16/60 PG | 5. SEC-A | 5. N/A |
| **SUMO-Nic96$^{187-301}$** | Individual | 1. Ni-NTA | 1. Ni-A1 | 1. Ni-B1 |
| | (#30) | 2. Dialysis | 2. IEX-A1 | 2. N/A |
| | | 3. HiTrap Q HP | 3. IEX-A1 | 3. IEX-B1 |
| | | 4. HiLoad Superdex 75 16/60 PG | 4. SEC-A | 4. N/A |
| **SUMO-Nic96$^{R2}$** | Individual | 1. Ni-NTA | 1. Ni-A1 | 1. Ni-B1 |
| wildtype and mutants | (#31-39) | 2. Dialysis | 2. IEX-A1 | 2. N/A |
| | | 3. HiTrap Q HP | 3. IEX-A1 | 3. IEX-B1 |
| | | 4. HiLoad Superdex 75 16/60 PG | 4. SEC-A | 4. N/A |
| **Nup53** | Individual | 1. Ni-NTA | 1. Ni-A1 | 1. Ni-B1 |
| | (#40) | 2. HiPrep 26/20 Desalting/Cleavage | 2. Ni-A2 / ULP1 | 2. N/A |
| | | 3. Ni-NTA | 3. Ni-A2 | 3. Ni-B2 |
| | | 4. HiPrep 26/20 Desalting | 4. IEX-A2 | 4. N/A |
| | | 5. HiTrap S HP | 5. IEX-A2 | 5. IEX-B2 |
| | | 6. HiLoad Superdex 200 16/60 PG | 6. SEC-A | 6. N/A |
| **Nup53$^{R1}$** | Individual | 1. Ni-NTA | 1. Ni-A1 | 1. Ni-B1 |
| | (#41) | 2. HiPrep 26/20 Desalting/Cleavage | 2. Ni-A2 / ULP1 | 2. N/A |
| | | 3. Ni-NTA | 3. Ni-A2 | 3. Ni-B2 |
| | | 4. HiPrep 26/20 Desalting | 4. IEX-A2 | 4. N/A |
| | | 5. HiTrap S HP | 5. IEX-A2 | 5. IEX-B2 |
| | | 6. HiLoad Superdex S75 16/60 PG | 6. SEC-A | 6. N/A |
| **SUMO-Nup145N** | Individual | 1. Ni-NTA | 1. Ni-A1 | 1. Ni-B1 |
| N/C-terminal truncations | (#68-129) | 2. Dialysis | 2. SEC-A | 2. N/A |
| | | 3. HiLoad Superdex 75 16/60 PG | 2. SEC-A | 2. N/A |
| **Nup145N** | Individual | 1. Ni-NTA | 1. Ni-A1 | 1. Ni-B1 |
| wildtype, 5-Ala and point mutants | (#68-129) | 2. Desalting/Cleavage | 2. IEX-A1/ ULP1 | 2. N/A |
| | | 3. MonoS 10/100 GL | 3. IEX-A1 | 3. IEX-B1 |
| | | 4. HiLoad Superdex 200 16/60 PG | 4. SEC-A | 4. N/A |
| **NUP54•NUP58•NUP62 (CNT)** | Co-expression | 1. Ni-NTA | 1. Ni-A1 | 1. Ni-B1 |
| | (#130-131) | 2. HiPrep 26/20 Desalting/Cleavage | 2. Ni-A2 / ULP1 | 2. N/A |
| | | 3. Ni-NTA | 3. Ni-A2 | 3. Ni-B2 |
| | | 4. MonoQ 5/50 GL | 5. IEX-A1 | 5. IEX-B1 |
| | | 5. HiLoad Superdex 200 10/300 GL | 6. SEC-A | 6. N/A |
| **SUMO-NUP93$^{R1}$** | Individual | 1. Ni-NTA | 1. Ni-A1 | 1. Ni-B1 |
| wildtype and mutant | (#132-133) | 2. HiPrep 26/20 Desalting | 2. IEX-A1 | 2. N/A |
| | | 3. HiTrap S HP | 3. IEX-A1 | 3. IEX-B1 |
| | | 3. HiLoad Superdex 75 16/60 PG | 2. SEC-A | 2. N/A |
| **SUMO-NUP93$^{\Delta R2SOL}$** | Individual | 1. Ni-NTA | 1. Ni-A1 | 1. Ni-B1 |
| wildtype and mutant | (#134-135) | 2. HiPrep 26/20 Desalting | 2. IEX-A1 | 2. N/A |
| | | 3. HiTrap S HP | 3. IEX-A1 | 3. IEX-B1 |
| | | 3. HiLoad Superdex 200 16/60 PG | 2. SEC-A | 2. N/A |
| **SUMO-NUP93$^{R2}$** | Individual | 1. Ni-NTA | 1. Ni-A1 | 1. Ni-B1 |
| wildtype and mutants | (#136-137) | 2. Dialysis | 2. IEX-A1 | 2. N/A |
| | | 3. HiTrap Q HP | 3. IEX-A1 | 3. IEX-B1 |
| | | 4. HiLoad Superdex 75 16/60 PG | 4. SEC-A | 4. N/A |
| **NUP93$^{SOL}$** | Individual | 1. Ni-NTA | 1. Ni-A1 | 1. Ni-B1 |
| wildtype and mutants | (#138-144) | 2. Dialysis/Cleavage | 2. Ni-A4 / ULP1 | 2. N/A |
| | | 3. Ni-NTA | 3. Ni-A4 | 3. Ni-B1 |
| | | 4. HiTrap Q HP | 4. IEX-A4 | 4. IEX-B1 |
| | | 5. HiLoad Superdex 200 16/60 PG | 5. SEC-A | 5. N/A |
| **SUMO-NUP53** | Individual | 1. Ni-NTA | 1. Ni-A1 | 1. Ni-B1 |
| N/C-terminal truncations | (#145-158, #179) | 2. Dialysis | 2. SEC-A | 2. N/A |
| | | 3. HiLoad Superdex 75 16/60 PG | 3. SEC-A | 3. N/A |
| **NUP53$^{R2}$** | Individual | 1. Ni-NTA | 1. Ni-A1 | 1. Ni-B1 |
| wildtype and mutant | (#149-150) | 2. Dialysis/Cleavage | 2. Ni-A3 | 2. N/A |
| | | 3. Ni-NTA | 3. Ni-A3 | 1. Ni-B1 |
| | | 4. HiLoad Superdex 75 16/60 PG | 4. SEC-A | 4. N/A |
| **SUMO-NUP53$^{N}$** | Individual | 1. Ni-NTA | 1. Ni-A1 | 1. Ni-B1 |
| wildtype and mutants | (#159-173) | 2. Dialysis | 2. SEC-A | 2. N/A |
| | | 3. HiLoad Superdex 75 16/60 PG | 3. SEC-A | 3. N/A |

| Protein(s) | Expression constructs | Purification step | Buffer A | Buffer B |
|---|---|---|---|---|
| **His6-Avi-SUMO-NUP53$^{N}$**<br>wildtype and mutant | Individual<br>(#174-175) | 1. Ni-NTA<br>2. HiPrep 26/20 Desalting<br>3. HiTrap S HP<br>3. HiLoad Superdex 200 16/60 PG | 1. Ni-A1<br>2. IEX-A4<br>3. IEX-A4<br>2. SEC-A | 1. Ni-B1<br>2. N/A<br>3. IEX-B1<br>2. N/A |
| **NUP53**<br>wildtype and truncations | Individual<br>(#176-178) | 1. Ni-NTA<br>2. HiPrep 26/20 Desalting<br>3. HiTrap S HP<br>3. HiLoad Superdex 200 16/60 PG | 1. Ni-A1<br>2. IEX-A4<br>3. IEX-A4<br>2. SEC-A | 1. Ni-B1<br>2. N/A<br>3. IEX-B1<br>2. N/A |
| **NUP155$^{NTD}$** | Individual<br>(#180) | 1. Ni-NTA<br>2. Dialysis/Cleavage<br>3. Ni-NTA<br>4. MonoQ 5/50 GL<br>5. HiLoad Superdex 200 16/60 PG | 1. Ni-A1<br>2. Ni-A2 / ULP1<br>3. Ni-A2<br>4. IEX-A1<br>5. SEC-A | 1. Ni-B1<br>2. N/A<br>3. Ni-B1<br>4. IEX-B1<br>5. N/A |
| **His6-Avi-SUMO-NUP98**<br>wildtype and truncations | Individual<br>(#181-183) | 1. Ni-NTA<br>2. Dialysis<br>3. HiTrap Q HP<br>4. HiLoad Superdex 200 16/60 PG | 1. Ni-A1<br>2. IEX-A1<br>2. IEX-A1<br>4. SEC-A | 1. Ni-B1<br>2. N/A<br>3. IEX-B1<br>4. N/A |

Ni-A1: 20 mM TRIS (pH 8.0), 500 mM NaCl, 20 mM imidazole, 5 mM β-ME
Ni-A2: 20 mM TRIS (pH 8.0), 100 mM NaCl, 20 mM imidazole, 5 mM β-ME
Ni-A3: 20 mM TRIS (pH 8.0), 150 mM NaCl, 20 mM imidazole, 5 mM β-ME
Ni-A4: 20 mM TRIS (pH 8.0), 25 mM NaCl, 20 mM imidazole, 5 mM β-ME

Ni-B1: 20 mM TRIS (pH 8.0), 500 mM NaCl, 500 mM imidazole, 5 mM β-ME
Ni-B2: 20 mM TRIS (pH 8.0), 100 mM NaCl, 500 mM imidazole, 5 mM β-ME

IEX-A1: 20 mM TRIS (pH 8.0), 100 mM NaCl, 5 mM DTT
IEX-A2: 20 mM TRIS (pH 7.0), 100 mM NaCl, 5 mM DTT
IEX-A3: 20 mM TRIS (pH 8.0), 25 mM NaCl, 5 mM DTT
IEX-A4: 20 mM MES (pH 6.0), 100 mM NaCl, 5 mM DTT

IEX-B1: 20 mM TRIS (pH 8.0), 2.0 M NaCl, 5 mM DTT
IEX-B2: 20 mM TRIS (pH 7.0), 2.0 M NaCl, 5 mM DTT

SEC-A: 20 mM TRIS (pH 8.0), 100 mM NaCl, 5 mM DTT
SEC-B: 20 mM TRIS (pH 8.0), 150 mM NaCl, 5 mM DTT

**Table S4.** SEC-MALS analysis

| Figure | Nucleoporin or Nucleoporin Complex | Experimental Mass (kDa) | Theoretical Mass (kDa) | Stoichiometry |
|---|---|---|---|---|
| Fig. 2F<br>Fig. S6A | Nup192 | 183 | 197 | Stoichiometric |
| | SUMO-Nic96$^{R2}$ | 21 | 21 | |
| | Nup192•SUMO-Nic96$^{R2}$ | 196 | 218 | |
| Fig. 2F<br>Fig. S6B | Nup192 | 184 | 197 | No binding |
| | SUMO-Nic96$^{R2}$ FFF | 18 | 21 | |
| | Nup192 FFF•SUMO-Nic96$^{R2}$ | 182 | 218 | |
| Fig. 2F<br>Fig. S6C | Nup192 LAF | 185 | 197 | No binding |
| | SUMO-Nic96$^{R2}$ | 20 | 21 | |
| | Nup192 LAF•SUMO-Nic96$^{R2}$ | 185 | 218 | |
| Fig. 3F<br>Fig. S8A | Nup188 | 191 | 204 | Stoichiometric |
| | SUMO-Nic96$^{R2}$ | 20 | 21 | |
| | Nup188•SUMO-Nic96$^{R2}$ | 210 | 225 | |
| Fig. 3F<br>Fig. S8B | Nup188 | 191 | 204 | No binding |
| | SUMO-Nic96$^{R2}$ FFF | 20 | 21 | |
| | Nup188•SUMO-Nic96$^{R2}$ FFF | 194 | 225 | |
| Fig. 3F<br>Fig. S8C | Nup188 LFV | 191 | 204 | No binding |
| | SUMO-Nic96$^{R2}$ | 20 | 21 | |
| | Nup188 LFV•SUMO-Nic96$^{R2}$ | 191 | 225 | |
| Fig. 4H<br>Fig. S19A | Nup192•SUMO-Nic96$^{R2}$•Nup53 | 244 | 258 | Stoichiometric |
| | Nup145N | 42 | 42 | |
| | Nup192•SUMO-Nic96$^{R2}$•Nup53•Nup145N | 315 | 300 | |
| Fig. 4H<br>Fig. S19B | Nup192•SUMO-Nic96$^{R2}$•Nup53 | 238 | 258 | No binding |
| | Nup145N KKRMYKLRKR | 42 | 42 | |
| | Nup192•SUMO-Nic96$^{R2}$•Nup53•Nup145N KKRMYKLRKR | 230 | 300 | |
| Fig. 4H<br>Fig. S19C | Nup192 LIFH•SUMO-Nic96$^{R2}$•Nup53 | 242 | 258 | Stoichiometric |
| | Nup145N | 45 | 42 | |
| | Nup192 LIFH•SUMO-Nic96$^{R2}$•Nup53•Nup145N | 297 | 300 | |
| Fig. 4H<br>Fig. S19D | Nup192•SUMO-Nic96$^{R2}$•Nup53 | 244 | 258 | Stoichiometric |
| | SUMO-Nup145N$^{R1}$ | 22 | 21 | |
| | Nup192•SUMO-Nic96$^{R2}$•Nup53•SUMO-Nup145N$^{R1}$ | 264 | 279 | |
| Fig. 4H<br>Fig. S19E | Nup192•SUMO-Nic96$^{R2}$•Nup53 | 245 | 258 | No binding |
| | SUMO-Nup145N$^{R1}$ KKRMYKLRKR | 21 | 21 | |
| | Nup192•SUMO-Nic96$^{R2}$•Nup53•SUMO-Nup145N$^{R1}$ KKRMYKLRKR | 240 | 279 | |
| Fig. 4H<br>Fig. S19F | Nup192 LIFH•SUMO-Nic96$^{R2}$•Nup53 | 242 | 258 | No binding |
| | SUMO-Nup145N$^{R1}$ | 25 | 21 | |
| | Nup192 LIFH•SUMO-Nic96$^{R2}$•Nup53•SUMO-Nup145N$^{R1}$ | 239 | 279 | |
| Fig. S20A | Nup192•SUMO-Nic96$^{R2}$ | 203 | 218 | Stoichiometric |
| | Nup53 | 46 | 40 | |
| | Nup192•SUMO-Nic96$^{R2}$•Nup53 | 245 | 258 | |
| Fig. S24B | Nup188•SUMO-Nic96$^{R2}$•Nup145N | 250 | 267 | No binding |
| | Nup53 | 39 | 41 | |
| | Nup188•SUMO-Nic96$^{R2}$•Nup145N•Nup53 | 243 | 308 | |
| Fig. 5F<br>Fig. S30A | Nup188•SUMO-Nic96$^{R2}$ | 218 | 225 | Stoichiometric |
| | Nup145N | 48 | 42 | |
| | Nup188•SUMO-Nic96$^{R2}$•Nup145N | 280 | 267 | |
| Fig. 5F<br>Fig. S30B | Nup188•SUMO-Nic96$^{R2}$ | 216 | 225 | Stoichiometric |
| | Nup145N EDSILF | 50 | 42 | |
| | Nup188•SUMO-Nic96$^{R2}$•Nup145N EDSILF | 288 | 267 | |
| Fig. 5F<br>Fig. S30C | Nup188 HHMI•SUMO-Nic96$^{R2}$ | 214 | 225 | Stoichiometric |
| | Nup145N | 48 | 42 | |
| | Nup188 HHMI•SUMO-Nic96$^{R2}$•Nup145N | 251 | 267 | |
| Fig. 5F<br>Fig. S30D | Nup188•SUMO-Nic96$^{R2}$ | 213 | 225 | Stoichiometric |
| | SUMO-Nup145N$^{R2}$ | 26 | 23 | |
| | Nup188•SUMO-Nic96$^{R2}$•SUMO-Nup145N$^{R2}$ | 221 | 248 | |
| Fig. 5F<br>Fig. S30E | Nup188•SUMO-Nic96$^{R2}$ | 213 | 225 | No binding |
| | SUMO-Nup145N$^{R2}$ EDSILF | 26 | 23 | |
| | Nup188•SUMO-Nic96$^{R2}$•SUMO-Nup145N$^{R2}$ EDSILF | 213 | 248 | |
| Fig. 5F<br>Fig. S30F | Nup188 HHMI•SUMO-Nic96$^{R2}$ | 212 | 225 | No binding |
| | SUMO-Nup145N$^{R2}$ | 26 | 23 | |
| | Nup188 HHMI•SUMO-Nic96$^{R2}$•SUMO-Nup145N$^{R2}$ | 211 | 248 | |

### **Table S5.** Isothermal Titration Calorimetry

**Nup192 + Nic96 titrations**

| Cell (titrand) | Cell conc. (µM) | Syringe (titrant) | Syringe conc. (µM) | Buffer | *T* (°C) | *n* | $K_D$ (nM) | *Δ°H* (kJ mol$^{-1}$) | *Δ°S* (J mol$^{-1}$ K$^{-1}$) |
|---|---|---|---|---|---|---|---|---|---|
| Nup192 | 14.0 | SUMO-Nic96 (187-301) | 70.0 | ITC-A | 4 | 1.02 ± 0.06 | 75 ± 5 | -53 ± 5 | -31 ± 16 |
| Nup192 | 15.6 | SUMO-Nic96$^{R2}$ | 80.0 | ITC-A | 4 | 1.09 ± 0.07 | 73 ± 4 | -53 ± 4 | -31 ± 14 |
| Nup192 | 15.6 | SUMO-Nic96$^{R2}$ FFF | 80.0 | ITC-A | 4 | n.d. | n.d. | n.d. | n.d. |
| Nup192 LAF | 15.6 | SUMO-Nic96R2 | 80.0 | ITC-A | 4 | n.d. | n.d. | n.d. | n.d. |

**Nup188 + Nic96 titrations**

| Cell (titrand) | Cell conc. (µM) | Syringe (titrant) | Syringe conc. (µM) | Buffer | *T* (°C) | *n* | $K_D$ (nM) | *Δ°H* (kJ mol$^{-1}$) | *Δ°S* (J mol$^{-1}$ K$^{-1}$) |
|---|---|---|---|---|---|---|---|---|---|
| Nup188 | 17.0 | SUMO-Nic96 (187-301) | 100.0 | ITC-A | 4 | 1.12 ± 0.07 | 95 ± 8 | 39 ± 2 | 294 ± 7 |
| Nup188 | 16.8 | SUMO-Nic96$^{R2}$ | 100.0 | ITC-A | 4 | 1.08 ± 0.05 | 85 ± 18 | 33 ± 3 | 266 ± 11 |
| Nup188 | 16.8 | SUMO-Nic96$^{R2}$ FFF | 100.0 | ITC-A | 4 | n.d. | n.d. | n.d. | n.d. |
| Nup188 FLV | 16.8 | SUMO-Nic96$^{R2}$ | 100.0 | ITC-A | 4 | n.d. | n.d. | n.d. | n.d. |

**Nup192 + Nup145N titrations**

| Cell (titrand) | Cell conc. (µM) | Syringe (titrant) | Syringe conc. (µM) | Buffer | *T* (°C) | *n* | $K_D$ (nM) | *Δ°H* (kJ mol$^{-1}$) | *Δ°S* (J mol$^{-1}$ K$^{-1}$) |
|---|---|---|---|---|---|---|---|---|---|
| Nup192•Nic96$^{R2}$•Nup53 | 14.0 | Nup145N | 125.0 | ITC-B | 4 | 1.30 ± 0.10 | 825 ± 228 | 45 ± 2 | 291 ± 13 |
| Nup192•Nic96$^{R2}$•Nup53 | 14.0 | Nup145N KKRMYKLRKR | 125.0 | ITC-B | 4 | n.d. | n.d. | n.d. | n.d. |
| Nup192 LIFH •Nic96$^{R2}$•Nup53 | 15.0 | Nup145N | 118.3 | ITC-B | 4 | 1.07 ± 0.09 | 6620 ± 1500 | 110 ± 13 | 510 ± 50 |
| Nup192•Nic96$^{R2}$•Nup53 | 20.0 | SUMO-Nup145N$^{R1}$ | 180.0 | ITC-B | 4 | 1.28 ± 0.04 | 1600 ± 100 | 35 ± 9 | 260 ± 30 |
| Nup192•Nic96$^{R2}$•Nup53 | 20.0 | SUMO-Nup145N$^{R1}$ KKRMYKLRKR | 180.0 | ITC-B | 4 | n.d. | n.d. | n.d. | n.d. |
| Nup192 LIFH •Nic96$^{R2}$•Nup53 | 20.0 | SUMO-Nup145N$^{R1}$ | 180.0 | ITC-B | 4 | n.d. | n.d. | n.d. | n.d. |

**Nup188 + Nup145N titrations**

| Cell (titrand) | Cell conc. (µM) | Syringe (titrant) | Syringe conc. (µM) | Buffer | *T* (°C) | *n* | $K_D$ (nM) | *Δ°H* (kJ mol$^{-1}$) | *Δ°S* (J mol$^{-1}$ K$^{-1}$) |
|---|---|---|---|---|---|---|---|---|---|
| Nup188•Nic96$^{R2}$ | 28.0 | Nup145N | 250.0 | ITC-A | 4 | 1.01 ± 0.01 | 410 ± 240 | 14.2 ± 0.6 | 178 ± 5 |
| Nup188•Nic96$^{R2}$ | 20.0 | Nup145N EDSILF | 200.0 | ITC-A | 4 | 1.03 ± 0.09 | 2750 ± 1100 | 18.4 ± 0.1 | 181 ± 3 |
| Nup188 HHMI •Nic96$^{R2}$ | 21.2 | Nup145N | 148.3 | ITC-A | 4 | 1.23 ± 0.05 | 4710 ± 3200 | 22.4 ± 1.3 | 186 ± 7 |
| Nup188 HHMI •Nic96$^{R2}$ | 25.5 | Nup145N EDSILF | 172.3 | ITC-A | 4 | 1.22 ± 0.04 | 3610 ± 570 | 25.1 ± 0.8 | 210 ± 3 |
| Nup188•Nic96$^{R2}$ | 30.0 | SUMO-Nup145N$^{R2}$ | 250.0 | ITC-A | 21 | 1.04 ± 0.10 | 2600 ± 760 | -36 ± 5 | -4 ± 16 |
| Nup188•Nic96$^{R2}$ | 30.0 | SUMO-Nup145N$^{R2}$ EDSILF | 250.0 | ITC-A | 21 | n.d. | n.d. | n.d. | n.d. |
| Nup188 HHMI •Nic96$^{R2}$ | 30.0 | SUMO-Nup145N$^{R2}$ | 250.0 | ITC-A | 21 | n.d. | n.d. | n.d. | n.d. |

ITC-A = 20 mM TRIS (pH 8.0), 100 mM NaCl, 1 mM TCEP
ITC-B = 20 mM TRIS (pH 8.0), 150 mM NaCl, 1 mM TCEP
n.d., not detectable under the specified experimental conditions

**Table S6.** Crystallization and cryoprotection conditions

| Protein(s) | Concentration | Crystallization condition | Cryo-protection condition |
|---|---|---|---|
| $Nup188^{NTD}$<br>SeMet $Nup188^{NTD}$ | 25 mg/ml | 10 % (w/v) PEG 3,350<br>0.1 M potassium thiocyanate | Mother liquor gradually supplemented to 25% (v/v) ethylene glycol in 2% steps |
| SeMet Nup188•$Nic96^{R2}$ | 8 mg/ml | 0.05 M HEPES (pH 7.5)<br>6.5% (w/v) PEG 20,000<br>After appearance of crystals, PEG 20,000 concentration in well solution increased by in 1%/day steps over a week to gradually dehydrate crystals | Mother liquor gradually supplemented to 25% (v/v) ethylene glycol in 2% steps. |
| $Nup192^{\Delta Head}$ •$Nic96^{187-301}$<br>$Nup192^{\Delta Head}$ •$Nic96^{R2}$<br>SeMet $Nup192^{\Delta Head}$ •$Nic96^{R2}$ A289M | 12 mg/ml | 0.05 M HEPES (pH 7.5)<br>0.1 M ammonium sulfate<br>8% (w/v) PEG 3,350<br>1% (v/v) 2-propanol | Mother liquor gradually supplemented to 25% (v/v) ethylene glycol in 1% steps |
| $NUP93^{SOL}$ | 15 mg/ml | drop solution:<br>0.075 M TRIS (pH 8.5)<br>11% (w/v) PEG 20,000<br>well solution:<br>0.075 M TRIS (pH 8.5)<br>19% (w/v) PEG 20,000 | 20% (v/v) ethylene glycol |
| $NUP93^{SOL}$•$NUP53^{R2}$<br>$NUP93^{SOL}$• SeMet $NUP53^{R2}$ I94M | 35 mg/ml $NUP93^{SOL}$; 10-fold molar excess $NUP53^{R2}$ | drop solution:<br>0.075 M TRIS (pH 8.5)<br>11 % (w/v) PEG 20,000<br>well solution:<br>0.075 M TRIS (pH 8.5)<br>17% (w/v) PEG 20,000 | 20% (v/v) ethylene glycol |

**Table S7.** X-ray crystallography analysis of Nup192$^{\Delta Head}$•Nic96$^{187-301}$ and Nup192$^{\Delta Head}$•Nic96$^{R2}$

| | | | |
|---|---|---|---|
| **Data collection** | | | |
| Protein | Nup192$^{\Delta Head}$• Nic96$^{187-301}$ | Nup192$^{\Delta Head}$• Nic96$^{R2}$ | Nup192$^{\Delta Head}$• Nic96$^{R2}$ A289M |
| PDB ID | 7MVT | | |
| Synchrotron | SSRL[a] | SSRL[a] | SSRL[a] |
| Beamline | BL12-2 | BL12-2 | BL12-2 |
| Space group | $P4_12_12$ | $P4_12_12$ | $P4_12_12$ |
| Cell dimensions | | | |
| *a*, *b*, *c* (Å) | 76.8, 76.8, 712.5 | 76.9, 76.9, 708.9 | 76.4, 76.4, 702.5 |
| α, β, γ (°) | 90, 90, 90 | 90, 90, 90 | 90, 90, 90 |
| | | *Se-Peak* | *Se-Peak* |
| Wavelength | 1.1401 | 0.9793 | 0.9793 |
| Resolution (Å) | 40.0 – 3.6 | 40.0 – 4.2 | 40.0 – 4.0 |
| $R_{meas}$ (%)[b] | 14.6 (188.2) | 26.4 (164.3) | 21.9 (153.1) |
| $R_{pim}$ (%)[b] | 3.2 (41.6) | 5.4 (33.4) | 3.2 (35.3) |
| $CC_{1/2}$[b] | 1.00 (0.78) | 1.00 (0.62) | 1.00 (0.47) |
| $< I / \sigma I >$[b] | 12.35 (1.66) | 9.47 (1.93) | 15.47 (1.87) |
| Completeness (%)[b] | 99.6 (97.7) | 99.5 (97.0) | 97.8 (79.4) |
| No. of observations | 545,652 | 400,350 | 863,520 |
| No. of unique reflections[b,c] | 26,269 (2,539) | 16,714 (1,584) | 19,028 (1,480) |
| Redundancy[b] | 20.8 (18.9) | 24.0 (24.2) | 45.4 (17.7) |
| **Refinement** | | | |
| Resolution (Å) | 40.0 – 3.6 | | |
| No. of reflections | 26,238 | | |
| No. of reflections test set | 2,326 (8.9%) | | |
| $R_{work}$ (%) / $R_{free}$ (%) | 25.7 / 29.7 | | |
| No. atoms (non-hydrogen) | 11,077 | | |
| Protein | 11,077 | | |
| Water | 0 | | |
| Ligand/Ions | 0 | | |
| B-factors | 171 | | |
| Protein | 171 | | |
| Water | - | | |
| Ligand/Ions | - | | |
| RMSD | | | |
| Bond lengths (Å) | 0.002 | | |
| Bond angles (°) | 0.5 | | |
| **Ramachandran plot**[d] | | | |
| Favored (%) | 98.2 | | |
| Additionally allowed (%) | 1.8 | | |
| Outliers (%) | 0.0 | | |
| **MolProbity**[d] | | | |
| Clashscore | 0.9 (100$^{th}$ percentile) | | |
| MolPorbity score | 0.8 (100$^{th}$ percentile) | | |

[a]SSRL, Stanford Synchotron Radiation Lightsource
[b]Highest-resolution shell is shown in parentheses
[c]Friedel pairs were merged
[d]As determined by MolProbity (*93*)

**Table S8.** X-ray crystallography analysis of Nup188$^{NTD}$

| | | | | |
|---|---|---|---|---|
| **Data collection** | | | | |
| Protein | Nup188$^{NTD}$ | Nup188$^{NTD}$ | | |
| PDB ID | 7MVW | | | |
| Synchrotron | SSRL[a] | SSRL[a] | | |
| Beamline | BL12-2 | BL12-2 | | |
| Space group | $P6_4$ | $P6_4$ | | |
| Cell dimensions | | | | |
| *a*, *b*, *c* (Å) | 173.5, 173.5, 111.1 | 173.3, 173.3, 111.4 | | |
| α, β, γ (°) | 90, 90, 120 | 90, 90, 120 | | |
| | | *Se-Inflection* | *Se-Peak* | *Se-Remote* |
| Wavelength | 0.97941 | 0.97947 | 0.97963 | 0.96108 |
| Resolution (Å) | 45.0 – 2.75 | 35.0 – 3.5 | 35.0 – 3.9 | 35.0– 4.0 |
| $R_{meas}$ (%)[b] | 17.1 (222.5) | 42.2 (243.2) | 52.0 (250.7) | 54.2 (237.2) |
| $R_{pim}$ (%)[b] | 5.3 (71.5) | 9.4 (53.3) | 11.4 (55.8) | 11.9 (54.5) |
| $CC_{1/2}$[b] | 1.00 (0.57) | 0.99 (0.67) | 0.99 (0.46) | 0.99 (0.49) |
| $< I / \sigma I >$[b] | 13.67 (1.26) | 8.78 (1.53) | 6.5 (1.2) | 6.1 (1.2) |
| Completeness (%)[b] | 99.8 (99.6) | 99.7 (98.3) | 99.8 (99.9) | 99.8 (99.9) |
| No. of observations | 508,212 | 504,862 | 360,045 | 336,348 |
| No. of unique reflections[b,c] | 48,981 (4,880) | 24,294 (2,404) | 17,477 (1,743) | 16,250 (1,620) |
| Redundancy[b] | 10.4 (9.5) | 20.8 (20.6) | 20.6 (20.0) | 20.7 (19.7) |
| **Refinement** | | | | |
| Resolution (Å) | 45.0 – 2.75 | | | |
| No. of reflections | 48,929 | | | |
| No. of reflections test set | 2,449 (5.0%) | | | |
| $R_{work}$ (%) / $R_{free}$ (%) | 22.3 / 25.4 | | | |
| No. atoms (non-hydrogen) | 8,141 | | | |
| Protein | 8,089 | | | |
| Water | 34 | | | |
| Ligand/Ions | 37 | | | |
| B-factors | 81 | | | |
| Protein | 81 | | | |
| Water | 60 | | | |
| Ligand/Ions | 87 | | | |
| RMSD | | | | |
| Bond lengths (Å) | 0.002 | | | |
| Bond angles (°) | 0.4 | | | |
| **Ramachandran plot**[d] | | | | |
| Favored (%) | 98.5 | | | |
| Additionally allowed (%) | 1.5 | | | |
| Outliers (%) | 0.0 | | | |
| **MolProbity**[d] | | | | |
| Clashscore | 0.49 (100$^{th}$ percentile) | | | |
| MolPorbity score | 0.67 (100$^{th}$ percentile) | | | |

[a]SSRL, Stanford Synchotron Radiation Lightsource
[b]Highest-resolution shell is shown in parentheses
[c]Friedel pairs were merged
[d]As determined by MolProbity (*93*)

**Table S9.** X-ray crystallography analysis of Nup188•Nic96[R2]

| | |
|---|---|
| **Data collection** | |
| Protein | Nup188•Nic96[R2] |
| PDB ID | 7MVX |
| Synchrotron | SSRL[a] |
| Beamline | BL12-2 |
| Space group | $H3_2$ |
| | *Se-Peak* |
| Cell dimensions | |
| *a*, *b*, *c* (Å) | 302.7, 302.7, 152.7 |
| α, β, γ (°) | 90, 90, 120 |
| Wavelength | 0.9794 |
| Resolution (Å) | 25 – 4.35 |
| $R_{meas}$ (%)[b] | 25.3 (249) |
| $R_{pim}$ (%)[b] | 5.7 (55.9) |
| $CC_{1/2}$[b] | 0.96 (0.78) |
| $< I / \sigma I >$[b] | 11.3 (2.4) |
| Completeness (%)[b] | 99.5 (100.0) |
| No. of observations | 344,504 |
| No. of unique reflections[b,c] | 17,578 (4,981) |
| Redundancy[b] | 19.6 (19.7) |
| **Refinement** | |
| Resolution (Å) | 25 – 4.35 |
| No. of reflections | 17,565 |
| No. of reflections test set | 895 (5.1%) |
| $R_{work}$ (%) / $R_{free}$ (%) | 27.3 / 31.6 |
| No. atoms (non-hydrogen) | 13,265 |
| Protein | 13,265 |
| Water | 0 |
| Ligand/Ions | 0 |
| B-factors | 255 |
| Protein | 255 |
| Water | - |
| Ligand/Ions | - |
| RMSD | |
| Bond lengths (Å) | 0.003 |
| Bond angles (°) | 0.6 |
| **Ramachandran plot[d]** | |
| Favored (%) | 98.4 |
| Additionally allowed (%) | 1.6 |
| Outliers (%) | 0.0 |
| **MolProbity[d]** | |
| Clashscore | 7.1 (100th percentile) |
| MolPorbity score | 1.4 (100th percentile) |

[a]SSRL, Stanford Synchotron Radiation Lightsource
[b]Highest-resolution shell is shown in parentheses
[c]Friedel pairs were merged
[d]As determined by MolProbity (*93*)

**Table S10.** X-ray crystallography analysis of and *apo* $NUP93^{SOL}$ and $NUP93^{SOL}$•$NUP53^{R2}$

| **Data collection** | | | | |
|---|---|---|---|---|
| Protein | $NUP93^{SOL}$ | $NUP93^{SOL}$ | $NUP93^{SOL}$•$NUP53^{R2}$ | $NUP93^{SOL}$•$NUP53^{R2}$ I94M |
| PDB ID | 7MW0 | | 7MW1 | |
| Synchrotron | SSRL[a] | SSRL[a] | SSRL[a] | SSRL[a] |
| Beamline | BL12-2 | BL12-2 | BL12-2 | BL12-2 |
| Space group | $P2_12_12_1$ | $P2_12_12_1$ | $P2_1$ | $P2_1$ |
| Cell dimensions | | | | |
| *a*, *b*, *c* (Å) | 74.9, 66.1, 79.2 | 74.9, 66.0, 79.2 | 67.7, 181.4, 78.9 | 68.4, 183.4, 79.9 |
| $\alpha$, $\beta$, $\gamma$ (°) | 90, 116.3, 90 | 90, 116.3, 90 | 90, 94.0, 90 | 90, 94.7, 90 |
| | | *S-SAD* | | |
| Wavelength | 0.9795 | 1.7711 | 0.9795 | 0.9789 |
| Resolution (Å) | 40.0 – 2.0 | 40.0 – 2.2 | 38.0 – 3.4 | 38.0 – 3.8 |
| $R_{meas}$ (%)[b] | 9.0 (26.0) | 5.8 (11.5) | 13.0 (22.6) | 19.7 (117.9) |
| $R_{pim}$ (%)[b] | 2.0 (55.5) | 1.3 (30.7) | 4.91 (83.7) | 7.5 (44.1) |
| $CC_{1/2}$[b] | 1.00 (0.73) | 1.00 (0.88) | 1.00 (0.50) | 0.99 (0.73) |
| $< I / \sigma I >$[b] | 26.0 (1.54) | 35.5 (2.53) | 10.1 (0.90) | 7.9 (1.80) |
| Completeness (%)[b] | 99.1 (98.3) | 96.7 (94.5) | 98.5 (99.3) | 97.7 (93.6) |
| No. of observations | 973,955 | 646,488 | 179,880 | 129,312 |
| No. of unique reflections[b,c] | 46,644 (4,598) | 34,223 (3,297) | 25,795 (2,574) | 18,718 (1,715) |
| Redundancy[b] | 20.9 (21.5) | 18.9 (13.7) | 7.0 (7.2) | 6.9 (7.0) |
| **Refinement** | | | | |
| Resolution (Å) | 37.4 – 2.0 | | 33.5 – 3.4 | |
| No. of reflections | 46,580 | | 25,795 | |
| No. of reflections test set | 1,998 (4.3%) | | 2,005 (7.8%) | |
| $R_{work}$ (%) / $R_{free}$ (%) | 22.5 / 25.4 | | 23.3 / 27.6 | |
| No. atoms (non-hydrogen) | 5,258 | | 10,194 | |
| Protein | 5,062 | | 10,194 | |
| Water | 192 | | 0 | |
| Ligand/Ions | 4 | | 0 | |
| B-factors | 57 | | 144 | |
| Protein | 58 | | 144 | |
| Water | 53 | | - | |
| Ligand/Ions | 49 | | - | |
| RMSD | | | | |
| Bond lengths (Å) | 0.002 | | 0.003 | |
| Bond angles (°) | 0.4 | | 0.6 | |
| **Ramachandran plot**[d] | | | | |
| Favored (%) | 98.4 | | 98.0 | |
| Additionally allowed (%) | 1.6 | | 2.0 | |
| Outliers (%) | 0.0 | | 0.0 | |
| **MolProbity**[d] | | | | |
| Clashscore | 2.96 (99th percentile) | | 5.15 ($100^{th}$ percentile) | |
| MolPorbity score | 1.09 (100th percentile) | | 1.38 ($100^{th}$ percentile) | |

[a]SSRL, Stanford Synchrotron Radiation Lightsource
[b]Highest-resolution shell is shown in parentheses
[c]Friedel pairs were merged
[d]As determined by MolProbity (*93*)

**Table S11.**

Single particle cryo-EM analysis of Nup192•Nic96[R2] and Nup192•Nic96[R2]•Nup145N[R1]•Nup53[R1]

| | | |
|---|---|---|
| **Data collection** | | |
| Protein | Nup192•Nic96[R2] | Nup192•Nic96[R2]•Nup145N[R1]•Nup53[R1] |
| PDB ID | 7MVU | 7MVV |
| EMDB ID | 24056 | 24057 |
| Microscope | Titan Krios[a] | Titan Krios[a] |
| Voltage (kV) | 300 | 300 |
| Magnification | 165,000x | 105,000x |
| Focus range (μm) | -4.0 – -1.0 | -3.0 – -1.0 |
| Detector camera | Gatan K2 Summit | Gatan K3 |
| Detector camera mode | counting | super-resolution |
| Pixel size (Å) | 0.834 | 0.433 |
| Total dose ($e^-$/Å$^2$) | 54.0 | 100.0 |
| Exposure time (s) | 8.06 | 3.03 |
| Number of frames | 40 | 60 |
| **Reconstruction** | | |
| Particle number | 176, 609 | 484,910 |
| Symmetry imposed | C1 | C1 |
| Pixel size (Å) | 1.668 | 1.421 |
| Box size (Å) | 192, 192, 192 | 256, 256, 256 |
| Resolution (unmasked, Å)[b] | 6.5 | 4.1 |
| Resolution (masked, Å)[b] | 3.75 | 3.2 |
| Sphericity[c] | 0.778 | 0.807 |
| **Model Refinement** | | |
| Resolution for refinement (Å) | 3.75 | 3.2 |
| Map sharpening B-factor (Å$^2$) | -92.7 | locally sharpened |
| No. atoms (non-hydrogen) | 25,585 | 25,766 |
| Protein | 25,585 | 25,766 |
| Water | - | - |
| Ligand/Ions | - | - |
| B-factors | 149 | 161 |
| Protein | 149 | 161 |
| Water | - | - |
| Ligand/Ions | - | - |
| RMSD | | |
| Bond lengths (Å) | 0.001 | 0.001 |
| Bond angles (°) | 0.3 | 0.3 |
| $FSC_{overall}$ [d] | 3.80 | 3.25 |
| $CC_{MASK}$ | 0.70 | 0.74 |
| EMRinger score | 0.63 | 1.09 |
| Clashscore[e] | 3.24 | 3.88 (100th percentile) |
| MolProbity score[e] | 1.12 | 1.17 (100th percentile) |
| **Ramachandran plot[e]** | | |
| Favored (%) | 98.9 | 98.7 |
| Additionally allowed (%) | 1.1 | 1.3 |
| Outliers (%) | 0.0 | 0.0 |
| **Rotamers[e]** | | |
| Favored (%) | 99.3 | 98.1 |
| Additionally allowed (%) | 0.7 | 1.9 |
| Outliers (%) | 0.0 | 0.0 |

[a] California Institute of Technology cryoEM Facility
[b]As determined by the Gold Standard Fourier Shell Correlation (FSC) cutoff of 0.143
[c]As determined by 3DFSC.
[d]$FSC_{overall} = \Sigma(N_{shell}\ FSC_{shell})/\Sigma(N_{shell})$, where $FSC_{shell}$ is the FSC in a given shell, $N_{shell}$ is the number of structure factors in the shell. $FSC_{shell} = \Sigma(F_{model}\ F_{EM})/(\sqrt{(\Sigma(|F|^2_{model}))}\ \sqrt{(\Sigma(F^2_{EM}))})$. The $FSC_{overall}$ cutoff is 0.143.
[e]As determined by MolProbity (*93*)

**Table S12.** Single particle cryo-EM analysis of Nup188•Nic96$^{R2}$ and Nup188•Nic96$^{R2}$•Nup145N$^{R2}$

| | | |
|---|---|---|
| **Data collection** | | |
| Protein complex | Nup188•Nic96$^{R2}$ | Nup188•Nic96$^{R2}$•Nup145N$^{R2}$ |
| PDB ID | 7MVY | 7MVZ |
| EMDB ID | 24058 | 24059 |
| Microscope | Titan Krios[a] | Titan Krios[a] |
| Voltage (kV) | 300 | 300 |
| Magnification (x) | 130,000 | 130,000 |
| Focus range (µm) | -2.5 – -0.5 | -2.5 – -0.5 |
| Detector camera | Gatan K3 | Gatan K3 |
| Detector camera mode | super-resolution | super-resolution |
| Pixel size (Å) | 0.296 | 0.296 |
| Total dose ($e^-$/Å$^2$) | 103.0 | 103.0 |
| Exposure time (s) | 2.00 | 2.00 |
| Number of frames | 57 | 57 |
| **Reconstruction** | | |
| Particle number | 709,123 | 298,317 |
| Symmetry imposed | C1 | C1 |
| Pixel size (Å) | 0.972 | 1.296 |
| Box size (Å) | 340, 340, 340 | 256, 256, 256 |
| Resolution (unmasked, Å)[b] | 2.87 | 3.25 |
| Resolution (masked, Å)[b] | 2.4 | 2.8 |
| Sphericity[c] | 0.869 | 0.858 |
| **Model Refinement** | | |
| Resolution for refinement (Å) | 2.4 | 2.8 |
| Map sharpening B-factor (Å$^2$) | -10.0 | locally sharpened |
| No. atoms (non-hydrogen) | 26,577 | 26,735 |
| Protein | 26,577 | 26,735 |
| Water | - | - |
| Ligand/Ions | - | - |
| B-factors | 78.1 | 149 |
| Protein | 78.1 | 149 |
| Water | - | - |
| Ligand/Ions | - | - |
| RMSD | | |
| Bond lengths (Å) | 0.001 | 0.002 |
| Bond angles (°) | 0.4 | 0.5 |
| $FSC_{overall}$[d] | 2.40 | 3.23 |
| $CC_{MASK}$ | 0.69 | 0.67 |
| EMRinger score | 1.03 | 0.58 |
| Clashscore[e] | 2.94 (100th percentile) | 2.77 (100$^{th}$ percentile) |
| MolProbity score[e] | 1.08 (100th percentile) | 1.07 (100$^{th}$ percentile) |
| **Ramachandran plot[e]** | | |
| Favored (%) | 99.3 | 98.5 |
| Additionally allowed (%) | 0.7 | 1.5 |
| Outliers (%) | 0.0 | 0.0 |
| **Rotamers[e]** | | |
| Favored (%) | 99.9 | 98.2 |
| Additionally allowed (%) | 0.1 | 1.8 |
| Outliers (%) | 0.0 | 0.0 |

[a] Pacific Northwest Center for Cryo-EM (PNCC)
[b]As determined by the Gold Standard Fourier Shell Correlation (FSC) cutoff of 0.143
[c]As determined by 3DFSC.
[d]$FSC_{overall} = \Sigma(N_{shell}\ FSC_{shell})/\Sigma(N_{shell})$, where $FSC_{shell}$ is the FSC in a given shell, $N_{shell}$ is the number of structure factors in the shell. $FSC_{shell} = \Sigma(F_{model}\ F_{EM})/(\sqrt{(\Sigma(|F|^2{}_{model}))}\ \sqrt{(\Sigma(F^2{}_{EM}))}$. The $FSC_{overall}$ cutoff of 0.143.
[e]As determined by MolProbity (*93*)

**Table S13.** Yeast expression constructs

| Plasmid | Protein | Residues (Mutations) | Vector | Restriction sites (5', 3') | Select. Marker | Reference |
|---|---|---|---|---|---|---|
| pRS415-$P_{Nop1}$-*scNIC96* | *sc*Nic96 | 1-839 | pRS415 | BamHI, NotI | *LEU2* | (*25*) |
| pRS415-$P_{Nop1}$-*scnic96 FFF* | *sc*Nic96 | 1-839, F136E, F139E, F159E | pRS415 | BamHI, NotI | *LEU2* | this study |
| pRS415-$P_{Nop1}$-*scnic96 ΔR2* | *sc*Nic96 | 1-97, 164-839 | pRS415 | BamHI, NotI | *LEU2* | this study |
| pRS415-$P_{Nop1}$-*scnic96 R2/32×GS* | *sc*Nic96 | 1-97, 32-mer GS linker, 164-839 | pRS415 | BamHI, NotI | *LEU2* | this study |
| pRS415-$P_{Nop1}$-*scnic96 R2/66×GS* | *sc*Nic96 | 1-97, 66-mer GS linker, 164-839 | pRS415 | BamHI, NotI | *LEU2* | this study |
| pRS415-$P_{Nop1}$-3×FLAG-*scNIC96* | *sc*Nic96 | 1-839 | pRS415 | BamHI, NotI | *LEU2* | this study |
| pRS415-$P_{Nop1}$-3×FLAG-*scnic96 FFF* | *sc*Nic96 | 1-839, F136E, F139E, F159E | pRS415 | BamHI, NotI | *LEU2* | this study |
| pRS415-$P_{Nop1}$-3×FLAG-*scnic96 ΔR2* | *sc*Nic96 | 1-97, 164-839 | pRS415 | BamHI, NotI | *LEU2* | this study |
| pRS415-$P_{Nop1}$-3×FLAG-*scnic96 R2/32×GS* | *sc*Nic96 | 1-97, 32-mer GS linker, 164-839 | pRS415 | BamHI, NotI | *LEU2* | this study |
| pRS415-$P_{Nop1}$-3×FLAG-*scnic96 R2/66×GS* | *sc*Nic96 | 1-97, 66-mer GS linker, 164-839 | pRS415 | BamHI, NotI | *LEU2* | this study |
| pRS415-$P_{Nop1}$-eGFP-*scNIC96* | *sc*Nic96 | 1-839 | pRS415 | BamHI, NotI | *LEU2* | (*25*) |
| pRS415-$P_{Nop1}$-eGFP-*scnic96 FFF* | *sc*Nic96 | 1-839, F136E, F139E, F159E | pRS415 | BamHI, NotI | *LEU2* | this study |
| pRS415-$P_{Nop1}$-eGFP-*scnic96 ΔR2* | *sc*Nic96 | 1-97, 164-839 | pRS415 | BamHI, NotI | *LEU2* | this study |
| pRS415-$P_{Nop1}$-eGFP-*scnic96 R2/32×GS* | *sc*Nic96 | 1-97, 32-mer GS linker, 164-839 | pRS415 | BamHI, NotI | *LEU2* | this study |
| pRS415-$P_{Nop1}$-eGFP-*scnic96 R2/66×GS* | *sc*Nic96 | 1-97, 66-mer GS linker, 164-839 | pRS415 | BamHI, NotI | *LEU2* | this study |
| pRS415-$P_{Nop1}$-mCherry-*scNIC96* | *sc*Nic96 | 1-839 | pRS415 | BamHI, NotI | *LEU2* | (*25*) |
| pRS415-$P_{Nop1}$-mCherry-*scnic96 R2-SOL* | *sc*Nic96 | 98-839 | pRS415 | BamHI, NotI | *LEU2* | this study |
| pRS415-$P_{Nop1}$-mCherry-*scnic96 R2/32×GS* | *sc*Nic96 | 1-97, 32-mer GS linker, 164-839 | pRS415 | BamHI, NotI | *LEU2* | this study |
| pRS415-$P_{Nop1}$-*scNUP192* | *sc*Nup192 | 1-1683 | pRS415 | NotI, SacII | *LEU2* | this study |
| pRS415-P $_{Nop1}$-*scnup192 ΔTail* | *sc*Nup192 | 1-1316 | pRS415 | NotI, SacII | *LEU2* | this study |
| pRS415-P $_{Nop1}$-*scnup192 ΔTower-Tail* | *sc*Nup192 | 1-1114 | pRS415 | NotI, SacII | *LEU2* | this study |
| pRS415-$P_{Nop1}$-*scnup192 Tower-Tail* | *sc*Nup192 | 1115-1683 | pRS415 | NotI, SacII | *LEU2* | this study |
| pRS415-$P_{Nop1}$-*scnup192 LAF* | *sc*Nup192 | 1-1683, L1525E, A1610E, Y1679E | pRS415 | NotI, SacII | *LEU2* | this study |
| pRS415-$P_{Nop1}$-*scnup192 LIFH* | *sc*Nup192 | 1-1683, I1053E, E1057R, E1107R, F1106E | pRS415 | NotI, SacII | *LEU2* | this study |
| pRS415-$P_{Nop1}$-*scnup192 LAF+LIFH* | *sc*Nup192 | 1-1683, L1525E, A1610E, Y1679E, I1053E, E1057R, E1107R, F1106E | pRS415 | NotI, SacII | *LEU2* | this study |
| pRS415-$P_{Nop1}$-3×HA-*scNUP192* | *sc*Nup192 | 1-1683 | pRS415 | NotI, SacII | *LEU2* | this study |
| pRS415-P $_{Nop1}$-3×HA-*scnup192 ΔTail* | *sc*Nup192 | 1-1316 | pRS415 | NotI, SacII | *LEU2* | this study |
| pRS415-P $_{Nop1}$-3×HA-*scnup192 ΔTower-Tail* | *sc*Nup192 | 1-1114 | pRS415 | NotI, SacII | *LEU2* | this study |
| pRS415-$P_{Nop1}$-3×HA-*scnup192 Tower-Tail* | *sc*Nup192 | 1115-1683 | pRS415 | NotI, SacII | *LEU2* | this study |
| pRS415-$P_{Nop1}$-3×HA-*scnup192 LAF* | *sc*Nup192 | 1-1683, L1525E, A1610E, Y1679E | pRS415 | NotI, SacII | *LEU2* | this study |
| pRS415-$P_{Nop1}$-3×HA-*scnup192 LIFH* | *sc*Nup192 | 1-1683, I1053E, E1057R, E1107R, F1106E | pRS415 | NotI, SacII | *LEU2* | this study |
| pRS415-$P_{Nop1}$-3×HA-*scnup192 LAF+LIFH* | *sc*Nup192 | 1-1683, L1525E, A1610E, Y1679E, I1053E, E1057R, E1107R, F1106E | pRS415 | NotI, SacII | *LEU2* | this study |
| pRS415-$P_{Nop1}$-eGFP-*scNUP192* | *sc*Nup192 | 1-1683 | pRS415 | NotI, SacII | *LEU2* | (*36*) |
| pRS415-P $_{Nop1}$-eGFP-*scnup192 ΔTail* | *sc*Nup192 | 1-1316 | pRS415 | NotI, SacII | *LEU2* | (*36*) |
| pRS415-P $_{Nop1}$-eGFP-*scnup192 ΔTower-Tail* | *sc*Nup192 | 1-1114 | pRS415 | NotI, SacII | *LEU2* | this study |
| pRS415-$P_{Nop1}$-eGFP-*scnup192 Tower-Tail* | *sc*Nup192 | 1115-1683 | pRS415 | NotI, SacII | *LEU2* | this study |
| pRS415-$P_{Nop1}$-eGFP-*scnup192 LAF* | *sc*Nup192 | 1-1683, L1525E, A1610E, Y1679E | pRS415 | NotI, SacII | *LEU2* | this study |
| pRS415-$P_{Nop1}$-eGFP-*scnup192 LIFH* | *sc*Nup192 | 1-1683, I1053E, E1057R, E1107R, F1106E | pRS415 | NotI, SacII | *LEU2* | this study |
| pRS415-$P_{Nop1}$-eGFP-*scnup192 LAF+LIFH* | *sc*Nup192 | 1-1683, L1525E, A1610E, Y1679E, I1053E, E1057R, E1107R, F1106E | pRS415 | NotI, SacII | *LEU2* | this study |
| pRS416-$P_{Nop1}$-mCherry-*scNUP188* | *sc*Nup188 | 1-1655 | pRS416 | BamHI, NotI | *URA3* | this study |
| pRS415-$P_{Nop1}$-*scNUP188* | *sc*Nup188 | 1-1655 | pRS415 | BamHI, NotI | *LEU2* | this study |
| pRS415-$P_{Nop1}$-*scnup188 ΔTail* | *sc*Nup188 | 1-1030 | pRS415 | BamHI, NotI | *LEU2* | this study |
| pRS415-$P_{Nop1}$-*scnup188 Tail* | *sc*Nup188 | 1031-1655 | pRS415 | BamHI, NotI | *LEU2* | this study |
| pRS415-$P_{Nop1}$-*scnup188 FLV* | *sc*Nup188 | 1-1655, L1316E, L1504E, M1560E | pRS415 | BamHI, NotI | *LEU2* | this study |
| pRS415-$P_{Nop1}$-*scnup188 ΔHead* | *sc*Nup188 | 408-1655 | pRS415 | BamHI, NotI | *LEU2* | this study |
| pRS415-$P_{Nop1}$-*scnup188 Head* | *sc*Nup188 | 1-407 | pRS415 | BamHI, NotI | *LEU2* | this study |
| pRS415-$P_{Nop1}$-*scnup188 HHMI* | *sc*Nup188 | 1-1655, Y330E, R333E, L351E, I390E | pRS415 | BamHI, NotI | *LEU2* | this study |
| pRS415-$P_{Nop1}$-*scnup188 FLV+HHMI* | *sc*Nup188 | 1-1655, Y330E, R333E, L351E, I390E, L1316E, L1504E, M1560E | pRS415 | BamHI, NotI | *LEU2* | this study |
| pRS415-$P_{Nop1}$-3×FLAG-*scNUP188* | *sc*Nup188 | 1-1655 | pRS415 | BamHI, NotI | *LEU2* | this study |
| pRS415-$P_{Nop1}$-3×FLAG-*scnup188 ΔTail* | *sc*Nup188 | 1-1030 | pRS415 | BamHI, NotI | *LEU2* | this study |
| pRS415-$P_{Nop1}$-3×FLAG-*scnup188 Tail* | *sc*Nup188 | 1031-1655 | pRS415 | BamHI, NotI | *LEU2* | this study |
| pRS415-$P_{Nop1}$-3×FLAG-*scnup188 FLV* | *sc*Nup188 | 1-1655, L1316E, L1504E, M1560E | pRS415 | BamHI, NotI | *LEU2* | this study |
| pRS415-$P_{Nop1}$-3×FLAG-*scnup188 ΔHead* | *sc*Nup188 | 408-1655 | pRS415 | BamHI, NotI | *LEU2* | this study |
| pRS415-$P_{Nop1}$-3×FLAG-*scnup188 Head* | *sc*Nup188 | 1-407 | pRS415 | BamHI, NotI | *LEU2* | this study |
| pRS415-$P_{Nop1}$-3×FLAG-*scnup188 HHMI* | *sc*Nup188 | 1-1655, Y330E, R333E, L351E, I390E | pRS415 | BamHI, NotI | *LEU2* | this study |
| pRS415-$P_{Nop1}$-3×FLAG-*scnup188 FLV+HHMI* | *sc*Nup188 | 1-1655, Y330E, R333E, I351E, L390E, L1316E, L1504E, M1560E | pRS415 | BamHI, NotI | *LEU2* | this study |
| pRS415-$P_{Nop1}$-eGFP-*scNUP188* | *sc*Nup188 | 1-1655 | pRS415 | BamHI, NotI | *LEU2* | this study |
| pRS415-$P_{Nop1}$-eGFP-*scnup188 ΔTail* | *sc*Nup188 | 1-1030 | pRS415 | BamHI, NotI | *LEU2* | this study |

| Plasmid | Protein | Residues (Mutations) | Vector | Restriction sites (5', 3') | Select. Marker | Reference |
|---|---|---|---|---|---|---|
| pRS415-$P_{Nop1}$-eGFP-*scnup188 Tail* | *sc*Nup188 | 1031-1655 | pRS415 | BamHI, NotI | *LEU2* | this study |
| pRS415-$P_{Nop1}$-eGFP-*scnup188 FLV* | *sc*Nup188 | 1-1655, L1316E, L1504E, M1560E | pRS415 | BamHI, NotI | *LEU2* | this study |
| pRS415-$P_{Nop1}$-eGFP-*scnup188 ΔHead* | *sc*Nup188 | 408-1655 | pRS415 | BamHI, NotI | *LEU2* | this study |
| pRS415-$P_{Nop1}$-eGFP-*scnup188 Head* | *sc*Nup188 | 1-407 | pRS415 | BamHI, NotI | *LEU2* | this study |
| pRS415-$P_{Nop1}$-eGFP-*scnup188 HHMI* | *sc*Nup188 | 1-1655, Y330E, R333E, L351E, I390E | pRS415 | BamHI, NotI | *LEU2* | this study |
| pRS415-$P_{Nop1}$-eGFP-*scnup188 FLV+HHMI* | *sc*Nup188 | 1-1655, Y330E, R333E, L351E, I390E, L1316E, L1504E, M1560E | pRS415 | BamHI, NotI | *LEU2* | this study |
| pRS416-$P_{Nop1}$-*scnup116-nup145C* chimera | *sc*Nup116 / *sc*Nup145C | 1-967 (*scnup116*), 459-1157 (*scnup145*) | pRS416 | BamHI, NotI | *URA3* | this study |
| pRS415-$P_{Nop1}$-*scnup116-nup145C* chimera | *sc*Nup116 / *sc*Nup145C | 1-967 (*scnup116*), 459-1157 (*scnup145*) | pRS415 | BamHI, NotI | *URA3* | this study |
| pRS413-$P_{Nop1}$-*scNUP116* | *sc*Nup116 | 1-1109 | pRS413 | NdeI, NotI | *HIS3* | this study |
| pRS413-$P_{Nop1}$-*scNUP145* | *sc*Nup116 | 1-1317 | pRS413 | BamHI, NotI | *HIS3* | this study |
| pRS415-$P_{Nop1}$-*scNUP100* | *sc*Nup100 | 1-956 | pRS415 | NdeI, NotI | *LEU2* | this study |
| pRS415-$P_{Nop1}$-*scnup100 GLEBS* | *sc*Nup100 / *sc*Nup116 | 1-280 (*scnup100*), 101-166 (*scnup116*), 281-956 (*scnup100*) | pRS415 | NdeI, NotI | *LEU2* | this study[1] |
| pRS415-$P_{Nop1}$-*scNUP145* | *sc*Nup145 | 1-1317 | pRS415 | BamHI, NotI | *LEU2* | this study |
| pRS415-$P_{Nop1}$-*scnup145 GLEBS* | *sc*Nup145 | 1-44 (*scnup145*), 101-166 (*scnup116*), 45-1157 (*scnup145*) | pRS415 | BamHI, NotI | *LEU2* | this study |
| pRS415-$P_{Nop1}$-*scnup145 nup116FG* | *sc*Nup145 / *sc*Nup116 | 1-686 (*scnup116*), 210-1157 (*scnup145*) | pRS415 | BamHI, NotI | *LEU2* | this study |
| pRS416-$P_{Nop1}$-*scNUP116* | *sc*Nup116 | 1-1109 | pRS416 | NdeI, NotI | *URA3* | this study |
| pRS413-$P_{Nop1}$-*scnup145C* | *sc*Nup145C | 606-1317 | pRS413 | BamHI, NotI | *HIS3* | this study |
| pRS415-$P_{Nop1}$-*scNUP116* | *sc*Nup116 | 1-1109 | pRS415 | NdeI, NotI | *LEU2* | this study |
| pRS415-$P_{Nop1}$-*scnup116 ΔGLEBS* | *sc*Nup116 | 1-99, 167-1109 | pRS415 | NdeI, NotI | *LEU2* | this study |
| pRS415-$P_{Nop1}$-*scnup116 ΔFG* | *sc*Nup116 | 100-166, 716-1109 | pRS415 | NdeI, NotI | *LEU2* | this study |
| pRS415-$P_{Nop1}$-*scnup116 FG-GLEBS* | *sc*Nup116 | 1-99, 167-715, 100-166, 716-1109 | pRS415 | NdeI, NotI | *LEU2* | this study |
| pRS415-$P_{Nop1}$-*scnup116 ΔCTD* | *sc*Nup116 | 1-966 | pRS415 | NdeI, NotI | *LEU2* | this study |
| pRS415-$P_{Nop1}$-*scnup116 ΔR1* | *sc*Nup116 | 1-769, NheI, 811-1109 | pRS415 | NdeI, NotI | *LEU2* | this study |
| pRS415-$P_{Nop1}$-*scnup116 R1/40×GS* | *sc*Nup116 | 1-769, 40-mer GS linker, 811-1109 | pRS415 | NdeI, NotI | *LEU2* | this study |
| pRS415-$P_{Nop1}$-*scnup116 R1 mut.* | *sc*Nup116 | 1-1109, R776A, K777A, K778A, Y785A, K786A, M787A | pRS415 | NdeI, NotI | *LEU2* | this study |
| pRS415-$P_{Nop1}$-*scnup116 ΔR2* | *sc*Nup116 | 1-810, NheI, 851-1109 | pRS415 | NdeI, NotI | *LEU2* | this study |
| pRS415-$P_{Nop1}$-*scnup116 R2/40×GS* | *sc*Nup116 | 1-810, 40-mer GS linker, 851-1109 | pRS415 | NdeI, NotI | *LEU2* | this study |
| pRS415-$P_{Nop1}$-*scnup116 R2 mut.* | *sc*Nup116 | 1-1109, D837A, E838A, S839A, I840A, L841A | pRS415 | NdeI, NotI | *LEU2* | this study |
| pRS415-$P_{Nop1}$-*scnup116 ΔR3* | *sc*Nup116 | 1-855, NheI, 868-1109 | pRS415 | NdeI, NotI | *LEU2* | this study |
| pRS415-$P_{Nop1}$-*scnup116 R3/12×GS* | *sc*Nup116 | 1-855, 12-mer GS linker, 868-1109 | pRS415 | NdeI, NotI | *LEU2* | this study |
| pRS415-$P_{Nop1}$-*scnup116 R3 mut.* | *sc*Nup116 | 1-1109, L858A, I859A, M864A, L865A, I866A | pRS415 | NdeI, NotI | *LEU2* | this study |
| pRS415-$P_{Nop1}$-*scnup116 R1+R2 mut.* | *sc*Nup116 | 1-1109, R776A, K777A, K778A, Y785A, K786A, M787A, D837A, E838A, S839A, I840A, L841A | pRS415 | NdeI, NotI | *LEU2* | this study |
| pRS415-$P_{Nop1}$-*scnup116 R2+R3 mut.* | *sc*Nup116 | 1-1109, D837A, E838A, S839A, I840A, L841A, L858A, I859A, M864A, L865A, I866A | pRS415 | NdeI, NotI | *LEU2* | this study |
| pRS415-$P_{Nop1}$-*scnup116 R1+R3 mut* | *sc*Nup116 | 1-1109, R776A, K777A, K778A, Y785A, K786A, M787A, L858A, I859A, M864A, L865A, I866A | pRS415 | NdeI, NotI | *LEU2* | this study |
| pRS415-$P_{Nop1}$-*scnup116 R1+R2+R3 mut.* | *sc*Nup116 | 1-1109, R776A, K777A, K778A, Y785A, K786A, M787A, D837A, E838A, S839A, I840A, L841A, L858A, I859A, M864A, L865A, I866A | pRS415 | NdeI, NotI | *LEU2* | this study |
| pRS413-$P_{Nop1}$-*scnup145C*-3×HA | *sc*Nup145C | 606-1317 | pRS413 | BamHI, NotI | *HIS3* | this study |
| pRS415-$P_{Nop1}$-3×FLAG-*scNUP116* | *sc*Nup116 | 1-1109 | pRS415 | NdeI, NotI | *LEU2* | this study |
| pRS415-$P_{Nop1}$-3×FLAG-*scnup116 ΔGLEBS* | *sc*Nup116 | 1-99, 167-1109 | pRS415 | NdeI, NotI | *LEU2* | this study |
| pRS415-$P_{Nop1}$-3×FLAG-*scnup116 ΔFG* | *sc*Nup116 | 100-166, 716-1109 | pRS415 | NdeI, NotI | *LEU2* | this study |
| pRS415-$P_{Nop1}$-3×FLAG-*scnup116 FG-GLEBS* | *sc*Nup116 | 1-99, 167-715, 100-166, 716-1109 | pRS415 | NdeI, NotI | *LEU2* | this study |
| pRS415-$P_{Nop1}$-3×FLAG-*scnup116 ΔCTD* | *sc*Nup116 | 1-966 | pRS415 | NdeI, NotI | *LEU2* | this study |
| pRS415-$P_{Nop1}$-3×FLAG-*scnup116 ΔR1* | *sc*Nup116 | 1-769, NheI, 811-1109 | pRS415 | NdeI, NotI | *LEU2* | this study |
| pRS415-$P_{Nop1}$-3×FLAG-*scnup116 R1/40×GS* | *sc*Nup116 | 1-769, 40-mer GS linker, 811-1109 | pRS415 | NdeI, NotI | *LEU2* | this study |
| pRS415-$P_{Nop1}$-3×FLAG-*scnup116 R1 mut.* | *sc*Nup116 | 1-1109, R776A, K777A, K778A, Y785A, K786A, M787A | pRS415 | NdeI, NotI | *LEU2* | this study |
| pRS415-$P_{Nop1}$-3×FLAG-*scnup116 ΔR2* | *sc*Nup116 | 1-810, NheI, 851-1109 | pRS415 | NdeI, NotI | *LEU2* | this study |
| pRS415-$P_{Nop1}$-3×FLAG-*scnup116 R2/40×GS* | *sc*Nup116 | 1-810, 40-mer GS linker, 851-1109 | pRS415 | NdeI, NotI | *LEU2* | this study |
| pRS415-$P_{Nop1}$-3×FLAG-*scnup116 R2 mut.* | *sc*Nup116 | 1-1109, D837A, E838A, S839A, I840A, L841A | pRS415 | NdeI, NotI | *LEU2* | this study |
| pRS415-$P_{Nop1}$-3×FLAG-*scnup116 ΔR3* | *sc*Nup116 | 1-855, NheI, 868-1109 | pRS415 | NdeI, NotI | *LEU2* | this study |
| pRS415-$P_{Nop1}$-3×FLAG-*scnup116 R3/12×GS* | *sc*Nup116 | 1-855, 12-mer GS linker, 868-1109 | pRS415 | NdeI, NotI | *LEU2* | this study |
| pRS415-$P_{Nop1}$-3×FLAG-*scnup116 R3 mut.* | *sc*Nup116 | 1-1109, L858A, I859A, M864A, L865A, I866A | pRS415 | NdeI, NotI | *LEU2* | this study |
| pRS415-$P_{Nop1}$-3×FLAG-*scnup116 R1+R2 mut.* | *sc*Nup116 | 1-1109, R776A, K777A, K778A, Y785A, K786A, M787A, D837A, E838A, S839A, I840A, L841A | pRS415 | NdeI, NotI | *LEU2* | this study |

| Plasmid | Protein | Residues (Mutations) | Vector | Restriction sites (5', 3') | Select. Marker | Reference |
|---|---|---|---|---|---|---|
| pRS415-$P_{Nop1}$-3×FLAG-*scnup116 R2+R3* | *sc*Nup116 | 1-1109, D837A, E838A, S839A, I840A, L841A, L858A, I859A, M864A, L865A, I866A | pRS415 | NdeI, NotI | *LEU2* | this study |
| pRS415-$P_{Nop1}$-3×FLAG-*scnup116 R1+R3* | *sc*Nup116 | 1-1109, R776A, K777A, K778A, Y785A, K786A, M787A, L858A, I859A, M864A, L865A, I866A | pRS415 | NdeI, NotI | *LEU2* | this study |
| pRS415-$P_{Nop1}$-3×FLAG-*scnup116 R1+R2+R3 mut.* | *sc*Nup116 | 1-1109, R776A, K777A, K778A, Y785A, K786A, M787A, D837A, E838A, S839A, I840A, L841A, L858A, I859A, M864A, L865A, I866A | pRS415 | NdeI, NotI | *LEU2* | this study |
| pRS413-$P_{Nop1}$-*scnup145C*-mCherry | *sc*Nup145C | 606-1317 | pRS413 | BamHI, NotI | *HIS3* | this study |
| pRS415-$P_{Nop1}$-eGFP-*scNUP116* | *sc*Nup116 | 1-1109 | pRS415 | NdeI, NotI | *LEU2* | this study |
| pRS415-$P_{Nop1}$-eGFP-*scnup116 ΔGLEBS* | *sc*Nup116 | 1-99, 167-1109 | pRS415 | NdeI, NotI | *LEU2* | this study |
| pRS415-$P_{Nop1}$-eGFP-*scnup116 ΔFG* | *sc*Nup116 | 100-166, 716-1109 | pRS415 | NdeI, NotI | *LEU2* | this study |
| pRS415-$P_{Nop1}$-eGFP-*scnup116 FG-GLEBS* | *sc*Nup116 | 1-99, 167-715, 100-166, 716-1109 | pRS415 | NdeI, NotI | *LEU2* | this study |
| pRS415-$P_{Nop1}$-eGFP-*scnup116 ΔCTD* | *sc*Nup116 | 1-966 | pRS415 | NdeI, NotI | *LEU2* | this study |
| pRS415-$P_{Nop1}$-eGFP-*scnup116 ΔR1* | *sc*Nup116 | 1-769, NheI, 811-1109 | pRS415 | NdeI, NotI | *LEU2* | this study |
| pRS415-$P_{Nop1}$-eGFP-*scnup116 R1/40×GS* | *sc*Nup116 | 1-769, 40-mer GS linker, 811-1109 | pRS415 | NdeI, NotI | *LEU2* | this study |
| pRS415-$P_{Nop1}$-eGFP-*scnup116 R1 mut.* | *sc*Nup116 | 1-1109, R776A, K777A, K778A, Y785A, K786A, M787A | pRS415 | NdeI, NotI | *LEU2* | this study |
| pRS415-$P_{Nop1}$-eGFP-*scnup116 ΔR2* | *sc*Nup116 | 1-810, NheI, 851-1109 | pRS415 | NdeI, NotI | *LEU2* | this study |
| pRS415-$P_{Nop1}$-eGFP-*scnup116 R2/40×GS* | *sc*Nup116 | 1-810, 40-mer GS linker, 851-1109 | pRS415 | NdeI, NotI | *LEU2* | this study |
| pRS415-$P_{Nop1}$-eGFP-*scnup116 R2 mut.* | *sc*Nup116 | 1-1109, D837A, E838A, S839A, I840A, L841A | pRS415 | NdeI, NotI | *LEU2* | this study |
| pRS415-$P_{Nop1}$-eGFP-*scnup116 ΔR3* | *sc*Nup116 | 1-855, NheI, 868-1109 | pRS415 | NdeI, NotI | *LEU2* | this study |
| pRS415-$P_{Nop1}$-eGFP-*scnup116 R3/12×GS* | *sc*Nup116 | 1-855, 12-mer GS linker, 868-1109 | pRS415 | NdeI, NotI | *LEU2* | this study |
| pRS415-$P_{Nop1}$-eGFP-*scnup116 R3 mut.* | *sc*Nup116 | 1-1109, L858A, I859A, M864A, L865A, I866A | pRS415 | NdeI, NotI | *LEU2* | this study |
| pRS415-$P_{Nop1}$-eGFP-*scnup116 R1+R2 mut.* | *sc*Nup116 | 1-1109, R776A, K777A, K778A, Y785A, K786A, M787A, D837A, E838A, S839A, I840A, L841A | pRS415 | NdeI, NotI | *LEU2* | this study |
| pRS415-$P_{Nop1}$-eGFP-*scnup116 R2+R3 mut.* | *sc*Nup116 | 1-1109, D837A, E838A, S839A, I840A, L841A, L858A, I859A, M864A, L865A, I866A | pRS415 | NdeI, NotI | *LEU2* | this study |
| pRS415-$P_{Nop1}$-eGFP-*scnup116 R1+R3 mut.* | *sc*Nup116 | 1-1109, R776A, K777A, K778A, Y785A, K786A, M787A, L858A, I859A, M864A, L865A, I866A | pRS415 | NdeI, NotI | *LEU2* | this study |
| pRS415-$P_{Nop1}$-eGFP-*scnup116 R1+R2+R3 mut.* | *sc*Nup116 | 1-1109, R776A, K777A, K778A, Y785A, K786A, M787A, D837A, E838A, S839A, I840A, L841A, L858A, I859A, M864A, L865A, I866A | pRS415 | NdeI, NotI | *LEU2* | this study |
| pRS413-$P_{Nop1}$ | N/A | N/A | pRS413 | N/A | *HIS3* | (*119*) |
| pRS415-$P_{Nop1}$ | N/A | N/A | pRS415 | N/A | *LEU2* | (*119*) |
| pRS415-$P_{Nop1}$-3×FLAG | N/A | N/A | pRS415 | N/A | *LEU2* | this study |
| pRS415-$P_{Nop1}$-3×HA | N/A | N/A | pRS415 | N/A | *LEU2* | this study |
| pRS415-$P_{Nop1}$-eGFP | N/A | N/A | pRS415 | N/A | *LEU2* | (*119*) |
| pRS415-$P_{Nop1}$-mCherry | N/A | N/A | pRS415 | N/A | *LEU2* | (*119*) |
| pFA6a-kanMX6 | N/A | N/A | pFA6a | N/A | N/A | (*121*) |
| pFA6a-natNT2 | N/A | N/A | pFA6a | N/A | N/A | (*120*) |
| pFA6a-loxP-natNT2-loxP | N/A | N/A | pFA6a | N/A | N/A | this study |
| pRS413-$P_{GAL1\text{-}10}$-Cre | Cre | N/A | pRS413 | XbaI, NotI | *HIS3* | this study[2] |
| pRS411-$P_{Nop1}$-*scRPL25*-mCherry | *sc*Rpl25 | 1-142 | pRS411 | SacII, BamHI | *MET15* | this study |
| pRS426- $P_{GAL1\text{-}10}$-3×FLAG-6×His-NUP188 | NUP188 | 1-1749 | pRS426 | BamHI, NotI | *URA3* | this study |
| pRS426- $P_{GAL1\text{-}10}$-3×FLAG-6×His-NUP205 | NUP205 | 1-2012 | pRS426 | BamHI, NotI | *URA3* | this study |

[1]GLEBS insertion position designed as in (*44*).
[2]The *Cre* gene was kindly provided by David Baltimore.

### **Table S14.** *Saccharomyces cerevisiae* strains

| Strain | Parental strain | Genotype | Plasmid | Transformed with | Used for | Reference |
|---|---|---|---|---|---|---|
| *nic96Δ* | BY4741 | MATa, *his3Δ1, leu2Δ0, met15Δ0, ura3Δ0, scnic96::HIS3* | pRS416-P$_{Nop1}$-mCherry-*scNIC96* | pRS415-P$_{Nop1}$-*scNIC96* variants | growth analysis, poly(A)$^+$ RNA export | (*25*) |
| *nic96Δ* | BY4741 | MATa, *his3Δ1, leu2Δ0, met15Δ0, ura3Δ0, scnic96::HIS3* | pRS416-P$_{Nop1}$-mCherry-*scNIC96* | pRS415-P$_{Nop1}$-3×FLAG-*scNIC96* variants | growth analysis, western blotting | (*25*) |
| *nic96Δ* | BY4741 | MATa, *his3Δ1, leu2Δ0, met15Δ0, ura3Δ0, scnic96::HIS3* | pRS416-P$_{Nop1}$-mCherry-*scNIC96* | pRS415-P$_{Nop1}$-eGFP-*scNIC96* variants | growth analysis, localization of *sc*Nic96 variants | (*25*) |
| *nic96Δ* | BY4741 | MATa, *his3Δ1, leu2Δ0, met15Δ0, ura3Δ0, scnic96::HIS3* | pRS416-P$_{Nop1}$-mCherry-*scNIC96* | pRS415-P$_{Nop1}$-eGFP-*scNIC96* variants, pRS411-P$_{Nop1}$-*scRPL25*-mCherry | *sc*Rpl25 localization | (*25*) |
| *nic96Δ* *nup57-GFP* | BY4741 *sc*Nup57-GFP | MATa, *his3Δ1, leu2Δ0, met15Δ0, ura3Δ0, scnic96::*natNT2, *scNUP57*-GFP(S65T)::*HIS3*MX6 | pRS416-P$_{Nop1}$-mCherry-*scNIC96* | pRS415-P$_{Nop1}$-mCherry-*scNIC96* variants | growth analysis, colocalization of *sc*Nic96 variants and *sc*Nup57 | (*25*) |
| *nup192Δ* | BY4741 | MATa, *his3Δ1, leu2Δ0, met15Δ0, ura3Δ0, scnup192::*natNT2 | pRS416-P$_{Nop1}$-mCherry-*scNUP192* | pRS415-P$_{Nop1}$-*scNUP192* variants | growth analysis, poly(A)$^+$ RNA export | this study |
| *nup192Δ* | BY4741 | MATa, *his3Δ1, leu2Δ0, met15Δ0, ura3Δ0, scnup192::*natNT2 | pRS416-P$_{Nop1}$-mCherry-*scNUP192* | pRS415-P$_{Nop1}$-3×HA-*scNUP192* variants | growth analysis, western blotting | this study |
| *nup192Δ* | BY4741 | MATa, *his3Δ1, leu2Δ0, met15Δ0, ura3Δ0, scnup192::*natNT2 | pRS416-P$_{Nop1}$-mCherry-*scNUP192* | pRS415-P$_{Nop1}$-eGFP-*scNUP192* variants | growth analysis, localization of *sc*Nup192 variants | this study |
| *nup188Δ* | BY4741 | MATa, *his3Δ1, leu2Δ0, met15Δ0, ura3Δ0, scnup188::*natNT2 | none | pRS415-P$_{Nop1}$-eGFP-*scNUP188* variants | localization of *sc*Nup188 variants further knockout | this study |
| *nup188Δ* *pom152Δ* | BY4741 *nup188Δ* | MATa, *his3Δ1, leu2Δ0, met15Δ0, ura3Δ0, scnup188::*natNT2, *scpom152*::kanMX6 | pRS416-P$_{Nop1}$-mCherry-*scNUP188* | pRS415-P$_{Nop1}$-*scNUP188* variants | growth analysis | this study |
| *nup188Δ* *pom152Δ* | BY4741 *nup188Δ* | MATa, *his3Δ1, leu2Δ0, met15Δ0, ura3Δ0, scnup188::*natNT2, *scpom152*::kanMX6 | pRS416-P$_{Nop1}$-mCherry-*scNUP188* | pRS415-P$_{Nop1}$-3×FLAG-*scNUP188* variants | growth analysis, western blotting | this study |
| *nup188Δ* *pom34Δ* | BY4741 *nup188Δ* | MATa, *his3Δ1, leu2Δ0, met15Δ0, ura3Δ0, scnup188::*natNT2, *scpom34*::kanMX6 | pRS416-P$_{Nop1}$-mCherry-*scNUP188* | pRS415-P$_{Nop1}$-*scNUP188* variants | growth analysis, poly(A)$^+$ RNA export | this study |
| *nup188Δ* *pom34Δ* | BY4741 *nup188Δ* | MATa, *his3Δ1, leu2Δ0, met15Δ0, ura3Δ0, scnup188::*natNT2, *scpom34*::kanMX6 | pRS416-P$_{Nop1}$-mCherry-*scNUP188* | pRS415-P$_{Nop1}$-3×FLAG-*scNUP188* variants | growth analysis, western blotting | this study |
| *nup188Δ* *pom34Δ* | BY4741 *nup188Δ* | MATa, *his3Δ1, leu2Δ0, met15Δ0, ura3Δ0, scnup188::*natNT2, *scpom34*::kanMX6 | pRS416-P$_{Nop1}$-mCherry-*scNUP188* | pRS415-P$_{Nop1}$-eGFP-*scNUP188* variants | growth analysis, localization of *sc*Nup188 variants | this study |
| *nup188Δ* *pom34Δ* | BY4741 *nup188Δ* | MATa, *his3Δ1, leu2Δ0, met15Δ0, ura3Δ0, scnup188::*natNT2, *scpom34*::kanMX6 | pRS416-P$_{Nop1}$-mCherry-*scNUP188* | pRS415-P$_{Nop1}$-eGFP-*scNUP188* variants, pRS411-P$_{Nop1}$-*scRPL25*-mCherry | *sc*Rpl25 localization | this study |
| *nup100Δ* | BY4741 | MATa, *his3Δ1, leu2Δ0, met15Δ0, ura3Δ0, scnup100::*kanMX6, | none | none | further knockout | this study |
| *nup145Δ* | BY4741 | MATa, *his3Δ1, leu2Δ0, met15Δ0, ura3Δ0, scnup145::*kanMX6 | pRS416-P$_{Nop1}$-*scnup116-scnup145C chimera* | none | further knockout | this study |
| *nup100Δ* *nup116Δ* | BY4741 *nup100Δ* | MATa, *his3Δ1, leu2Δ0, met15Δ0, ura3Δ0, scnup100::*kanMX6, *scnup116::*loxP-natNT2-loxP | pRS416-P$_{Nop1}$-*scnup116-scnup145C chimera* | pRS413-P$_{Nop1}$-*scNUP116* variants, pRS415-P$_{Nop1}$-*scNUP100* variants | growth analysis | this study |
| *nup100Δ* *nup145Δ* | BY4741 *nup100Δ* | MATa, *his3Δ1, leu2Δ0, met15Δ0, ura3Δ0, scnup100::*kanMX6, *scnup145::*loxP-natNT2-loxP | pRS416-P$_{Nop1}$-*scnup116-scnup145C chimera* | pRS413-P$_{Nop1}$-*scNUP116* variants, pRS415-P$_{Nop1}$-*scNUP145* variants | growth analysis | this study |
| *nup116Δ* *nup145Δ* | BY4741 *nup145Δ* | MATa, *his3Δ1, leu2Δ0, met15Δ0, ura3Δ0, scnup116::*natNT2, *scnup145::*kanMX6 | pRS416-P$_{Nop1}$-*scnup116-scnup145C chimera* | pRS413-P$_{Nop1}$-*scNUP145* variants, pRS415-P$_{Nop1}$-*scNUP100* variants | growth analysis | this study |
| *nup100Δ* *nup116Δ* *nup145Δ* | BY4741 *nup100Δ* *nup116Δ* | MATa, *his3Δ1, leu2Δ0, met15Δ0, ura3Δ0, scnup100::* kanMX6, *scnup116::*loxP, *scnup145::*natNT2 | pRS416-P$_{Nop1}$-*scnup116-scnup145C chimera* | pRS413-P$_{Nop1}$-*scNUP145* variants, pRS415-P$_{Nop1}$-*scNUP116* variants | growth analysis | this study |
| *nup100Δ* *nup116Δ* *nup145Δ* | BY4741 *nup100Δ* *nup116Δ* | MATa, *his3Δ1, leu2Δ0, met15Δ0, ura3Δ0, scnup100::* kanMX6, *scnup116::*loxP, *scnup145::*natNT2 | pRS416-P$_{Nop1}$-*scNUP116,* pRS413-P$_{Nop1}$-*scnup145C* | pRS415-P$_{Nop1}$-*scnup145N* paralogs | growth analysis | this study |
| *nup100Δ* *nup116Δ* *nup145Δ* | BY4741 *nup100Δ* *nup116Δ* | MATa, *his3Δ1, leu2Δ0, met15Δ0, ura3Δ0, scnup100::* kanMX6, *scnup116::*loxP, *scnup145::*natNT2 | pRS416-P$_{Nop1}$-*scNUP116,* pRS413-P$_{Nop1}$-*scnup145C* | pRS415-P$_{Nop1}$-*scNUP116* variants | growth analysis, poly(A)$^+$ RNA export | this study |
| *nup100Δ* *nup116Δ* *nup145Δ* | BY4741 *nup100Δ* *nup116Δ* | MATa, *his3Δ1, leu2Δ0, met15Δ0, ura3Δ0, scnup100::* kanMX6, *scnup116::*loxP, *scnup145::*natNT2 | pRS416-P$_{Nop1}$-*scNUP116,* pRS413-P$_{Nop1}$-*scnup145C-3*×HA | pRS415-P$_{Nop1}$-3×FLAG-*scNUP116* variants | growth analysis, western blotting | this study |
| *nup100Δ* *nup116Δ* *nup145Δ* | BY4741 *nup100Δ* *nup116Δ* | MATa, *his3Δ1, leu2Δ0, met15Δ0, ura3Δ0, scnup100::* kanMX6, *scnup116::*loxP, *scnup145::*natNT2 | pRS416-P$_{Nop1}$-*scNUP116,* pRS413-P$_{Nop1}$-*scnup145C*-mCherry | pRS415-P$_{Nop1}$-eGFP-*scNUP116* variants | growth analysis, colocalization of *sc*Nup116 variants and *sc*Nup145C | this study |
| *nup100Δ* *nup116Δ* *nup145Δ* | BY4741 *nup100Δ* *nup116Δ* | MATa, *his3Δ1, leu2Δ0, met15Δ0, ura3Δ0, scnup100::* kanMX6, *scnup116::*loxP, *scnup145::*natNT2 | pRS416-P$_{Nop1}$-*scNUP116,* pRS413-P$_{Nop1}$-*scnup145C* | pRS415-P$_{Nop1}$-eGFP-*scNUP116* variants, pRS411-P$_{Nop1}$-*scRPL25*-mCherry | *sc*Rpl25 localization | this study |
